# Deep Learning-based Ligand Design using Shared Latent Implicit Fingerprints from Collaborative Filtering

**DOI:** 10.1101/2020.11.18.389213

**Authors:** Raghuram Srinivas, Niraj Verma, Elfi Kraka, Eric C. Larson

## Abstract

In their previous work, Srinivas et al.^1^ have shown that implicit fingerprints capture ligands and proteins in a shared latent space, typically for the purposes of virtual screening with collaborative filtering models applied on known bioactivity data. In this work, we extend these implicit fingerprints/descriptors using deep learning techniques to translate latent descriptors into discrete representations of molecules (SMILES), without explicitly optimizing for chemical properties. This allows the design of new compounds based upon the latent representation of nearby proteins, thereby encoding drug-like properties including binding affinities to known proteins. The implicit descriptor method does not require any fingerprint similarity search, which makes the method free of any bias arising from the empirical nature of the fingerprint models. ^1^ We evaluate the properties of the novel drugs generated by our approach using physical properties of drug-like molecules and chemical complexity. Additionally, we analyze the reliability of the biological activity of the new compounds generated using this method by employing models of protein ligand interaction, which assists in assessing the potential binding affinity of the designed compounds. We find that the generated compounds exhibit properties of chemically feasible compounds and are likely to be excellent binders to known proteins. Furthermore, we also analyze the diversity of compounds created using the Tanimoto distance and conclude that there is a wide diversity in the generated compounds.

**Graphical TOC Entry:** 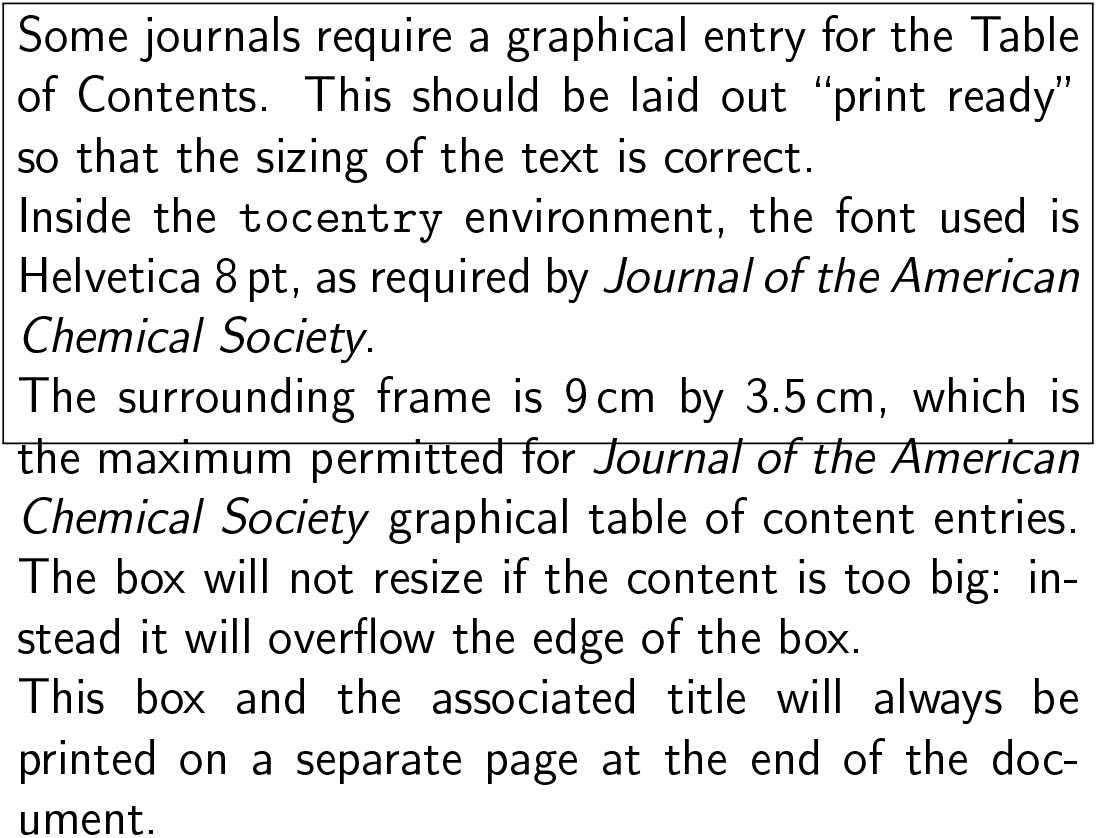

## Introduction

The field of virtual screening, a constituent part of the modern drug discovery process^2,3^ has been entrenched in the pharmaceutical industry for years and has developed into a sophisticated tool.^4–6^ A number of successful virtual screening strategies to identify novel hits have been reported, which serves as starting point for further investigation^7–9^. Many state-of-the-art protein-ligand interaction (PLI) models use machine learning that relies on abstract descriptors of compounds/proteins as input features.^10–15^ Virtual screening in drug discovery that use these models, while high performing, is not free of deficiencies—the limitations of representing drug compounds and targets abstractly also limits our ability to infer their binding properties. ^16,17^

We argue that a critical barrier is the lack of a universal fingerprinting model that can amass knowledge about drug compounds, protein targets, and assay characteristics in a shared latent space that can be used by a variety of machine learning models, visualization tools, and compound design tools. We further argue that if the representation is completely abstract, even if it performs well at PLI prediction, it is fundamentally limited because researchers cannot systematically create candidate compounds based on the featurization of the target.^18–20^

In their previous work Srinivas et al. ^1^, proposed the conception and development of an implicit mathematical representation that allows for a more accurate characterization of the drug compound and protein target in the same numeric latent space (as opposed to the current practice of separate descriptors for the compound and target), thus narrowing the model-associated bias down to that of the assay, i.e., real clinical (albeit in vitro) environment. ^21^ Additionally, to facilitate the ability to discover the physical structure of new compounds, in this work we propose decoding methods that map from the implicit representation of the candidate compounds to their physical structure. We believe this expanded capacity of the fingerprinting model will have significant impact on virtual screening and, consequently, drug discovery, as it will render drug discovery less dependent on costly clinical facilities and services. Moreover, the representation will provide new methods for creating and testing candidate drug compounds.

Several recent works have investigated the use of neural embedding on compound structure representations such as SMILES codes, showing this embedding is effective for exploring the chemical properties^22^ and generating novel compounds. Gómez-Bombarelli et al.^23^ refer to these embedded fingerprints as implicit representations. However, their methods work upon the raw SMILES textual representation and are therefore limited in their ability to discern more complicated relationships encoded by graphical fingerprints. The recent years have seen a plethora of deep learning based generative models for de-novo drug generation.^24–29^ The common theme in these techniques is to provide as input to the deep learning model, the molecules only to produce the same or similar molecules as output. The continuous vector representations of the input molecules in the intermediate layers produce a larger chemical property space, which is then sampled to produce novel molecules. In this work, we design and train deep learning methods that leverage the implicit compound fingerprints obtained from collaborative filtering based on the past bioactivity/assay data to map back to the physical structure of compounds. The implicit encoding of compounds are a continuous vector-valued representation and thus lend itself to the use of continuous optimization to generate novel compounds. We further assess the properties of the novel ligands generated in terms of the drug-like physical properties of molecules, chemical complexity and biological activity. We observed that our compounds exhibit properties similar to the known ligands even though our approach does not explicitly train the neural network for optimizing specific properties. Additionally, we compare our work to the prior work of Gómez-Bombarelli et al.^30^ on a set of chemical compounds with known binding affinities to cancer targets from the ChEMBL23 database.^31^ This comparative analysis investigates not only the potential binding affinity of the generated compounds to selected protein targets, but also the diversity of compounds generated. We provide evidence that our method is superior in both binding affinity and compound diversity. Finally, we conclude with a discussion of how our method could be integrated into a compound design tool and explore some of the advantages and limitations that such a tool would provide.

### Implicit Fingerprints from Collaborative Filtering

In their previous work, Srinivas et al.^1^ investigated implicit fingerprinting models that extend the existing virtual screening mechanisms by incorporating collaborative filtering. Collaborative filtering algorithms are used for designing recommendation systems such as movie recommendation engines. ^32,33^ In general, collaborative filtering is a method for making automatic predictions (filtering) about the interests of a user by collecting preferences or taste information from other users (collaborating). When applied to the field of virtual screening, this approach relies on modeling predictions based on assays measuring the interactions between compounds and targets.^34,35^ To intuit this,one can imagine building a recommendation system for matching movies to people. The direct approach might try to extract features specific to the person, like genre preferences and preferred actors, and features of the movie, like genre and runtime, to classify a match. This is the approach that is most similar to virtual screening where researchers directly featurize the compounds and targets based on their geometry and physicochemical properties. A more implicit approach groups users based on the movies they liked and groups movies based on the users that have seen them. Users and movies in similar groups could be implicitly found without attempting to featurize aspects of the users or movies directly.

In their previous work, Srinivas et al. ^1^ elucidated the performance of the collaborative filtering against the traditional approaches using evaluation criteria such as 1% enrichment factor (EF1%)^36^ for its ability to address specific properties of the early recognition problem specific to virtual screening, Boltzmann enhanced discrimination of the receiver operating characteristic (BEDROC20),^37^ and area under the curve (AUC) of the receiver operating characteristic.^38^ The collaborative filtering algorithm was found, at that time, to consistently and significantly outperform all the other methods using the evaluation criteria. Furthermore, the utility of the implicit continuous representation of the ligands obtained from collaborative filtering was illustrated in an example with cancer related targets as described in the next section.

#### Representation of Ligands and Proteins in Implicit Fingerprint Space

To help intuit the inherent properties of the implicit latent space, we randomly selected two cancer related targets from the ChEMBL23 database. We selected targets with ChEMBL23 IDs CHEMBL4899 and CHEMBL2150837 along with all the ligands with assays for these targets as shown in Figure 1. The 50-dimensional implicit fingerprints of the compounds are reduced into a 2-dimensional space using stochastic neighbor embedding (t-SNE).^39^ We visualize all compounds with available assays for the three selected cancer related protein targets in the ChEMBL23 database. The compounds are color-coded as either having demonstrated binding affinities to the target or not, on the basis of their standardized concentration levels in the assays, where a decreasing concentration level indicates stronger binding affinity. For the t-SNE plots, the ideal result would be perfect clustering for each concentration level, which would indicate the compounds cluster based on their binding affinity. Interestingly, the implicit ligand fingerprints in figure1 demonstrate a very clear separation between the compounds based on the concentration levels required to trigger binding affinities with the respective targets. This visual separation is striking for assays with excellent binding affinity (standard value below 100nM), indicating that the implicit representation is excellent in its ability to capture properties of similar compounds using an Euclidean distance.

**Figure 1:**
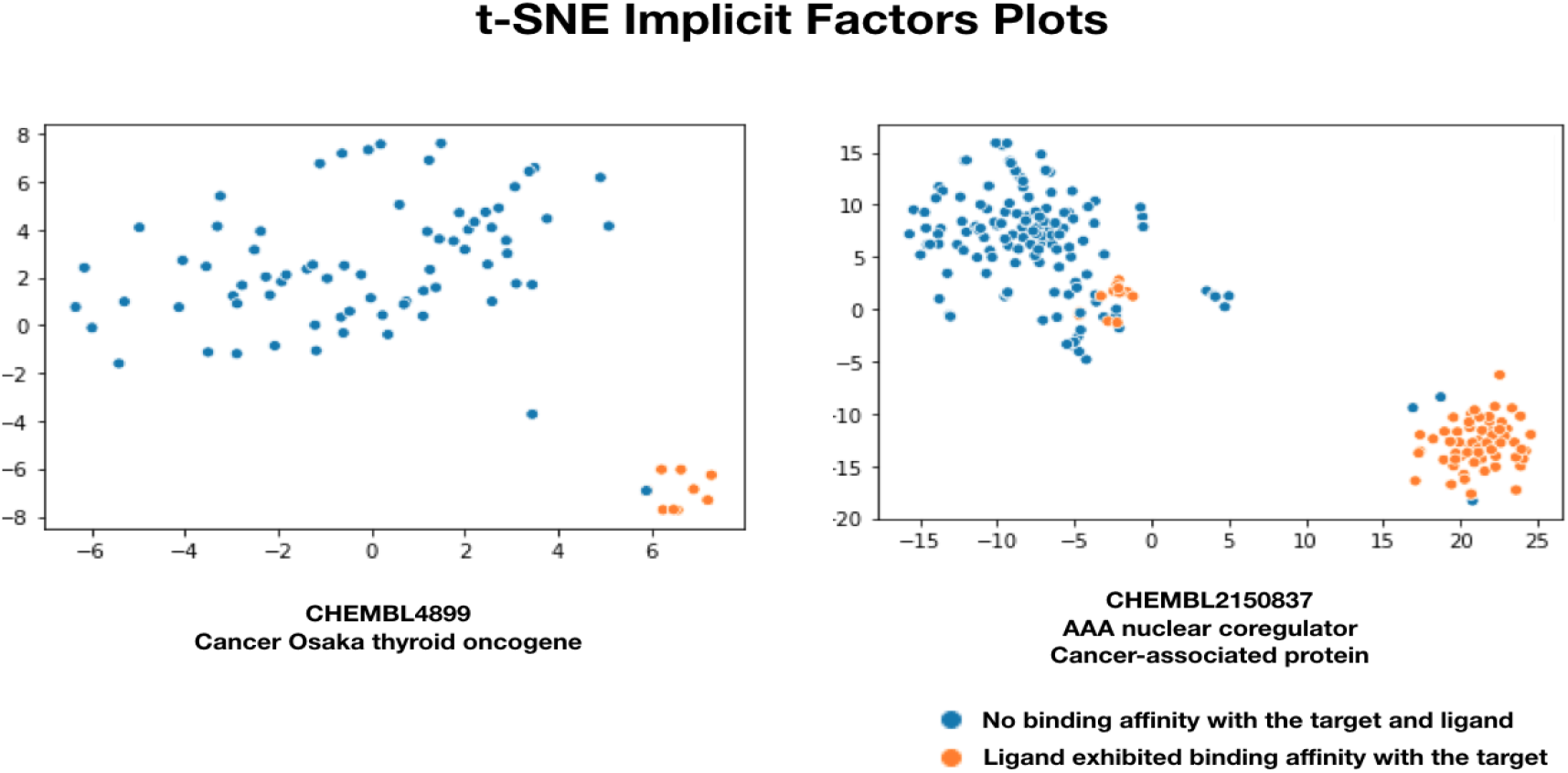
tSNE Plots of Implicit Ligand Fingerprints: Plots for two cancer targets are shown where each point represents a compound assayed from the ChEMBL database. The concentration results of the assays are color coded. tSNE plots of the 50 dimensional implicit representations, reduced to 2 dimensions preserving distance.

In order to further elucidate the power of implicit fingerprints for representing protein targets, we employed the t-SNE technique to embed 50 implicit *protein target fingerprints* into a 2-dimensional projection. That is, the fingerprints of protein targets are visualized, not compounds. Figure 2 illustrates the distribution of the protein targets when mapped into a 2-dimensional space, with the cancer related target proteins highlighted in blue. The graph demonstrates the presence of three potential clusters of known cancer related targets appearing close to each other. These cancer-related targets are visually separated from other biological targets in the latent space.

**Figure 2:**
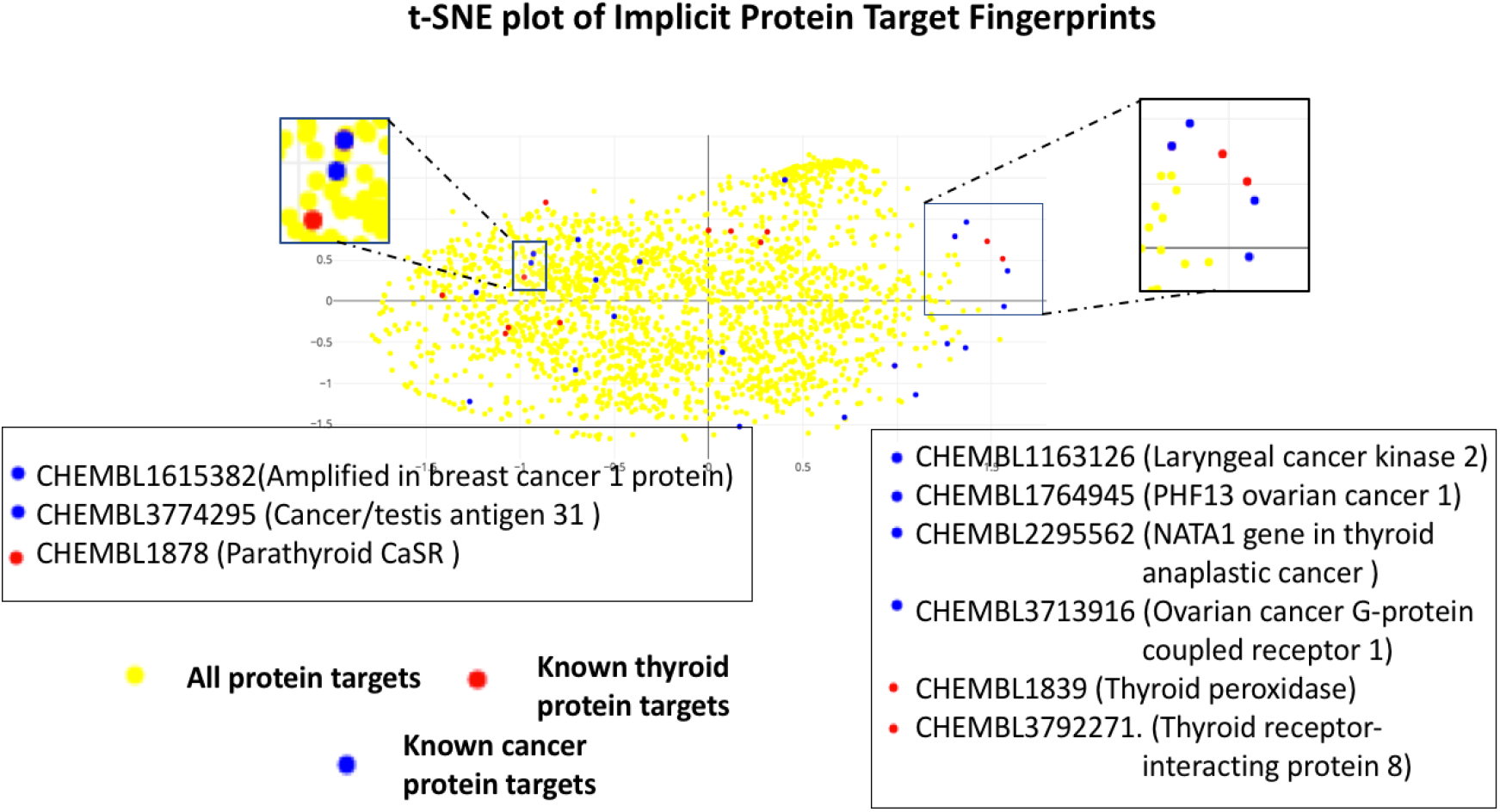
tSNE Plots of Implicit Protein Fingerprints^1^ (Left) t-SNE plot reducing the dimensions of the 50-dimensional implicit fingerprints into two dimensions for a subset of targets in the ChEMBL database. The method successfully clusters many known cancer related targets close to each other. (Right) A zoomed version of the largest cluster of cancer targets.

### Neural Network Architecture

Our method to generate novel ligands is comprised of two steps. (1) We generate implicit ligand and protein fingerprints using collaborative filtering and (2) train a neural network to generate the SMILES string from the implicit representation (i.e., a decoder that can map to a conventional representation from the implicit space). The first step involves generating the implicit fingerprints using known assays by applying the collaborative filtering algorithm. ^1^ This step yields the implicit fingerprint representations for both ligands and protein targets, as described above. The implicit fingerprints are continuous vectors that represent a point in 50 dimensional space.

The Implicit fingerprints of the ligands are then fed into a Gated Recurrent Unit (GRU)^40^ neural network to map the corresponding SMILES string encoding. The neural network is trained to minimize the error in reproducing the relevant SMILES string for each input implicit fingerprints of the ligands. The key aspect of the neural network is to learn the function to map the fixed-length continuous vector representation to the SMILES string. This architecture is illustrated in Figure 3. Additional details of the neural network design are discussed under the methods section.

**Figure 3:**
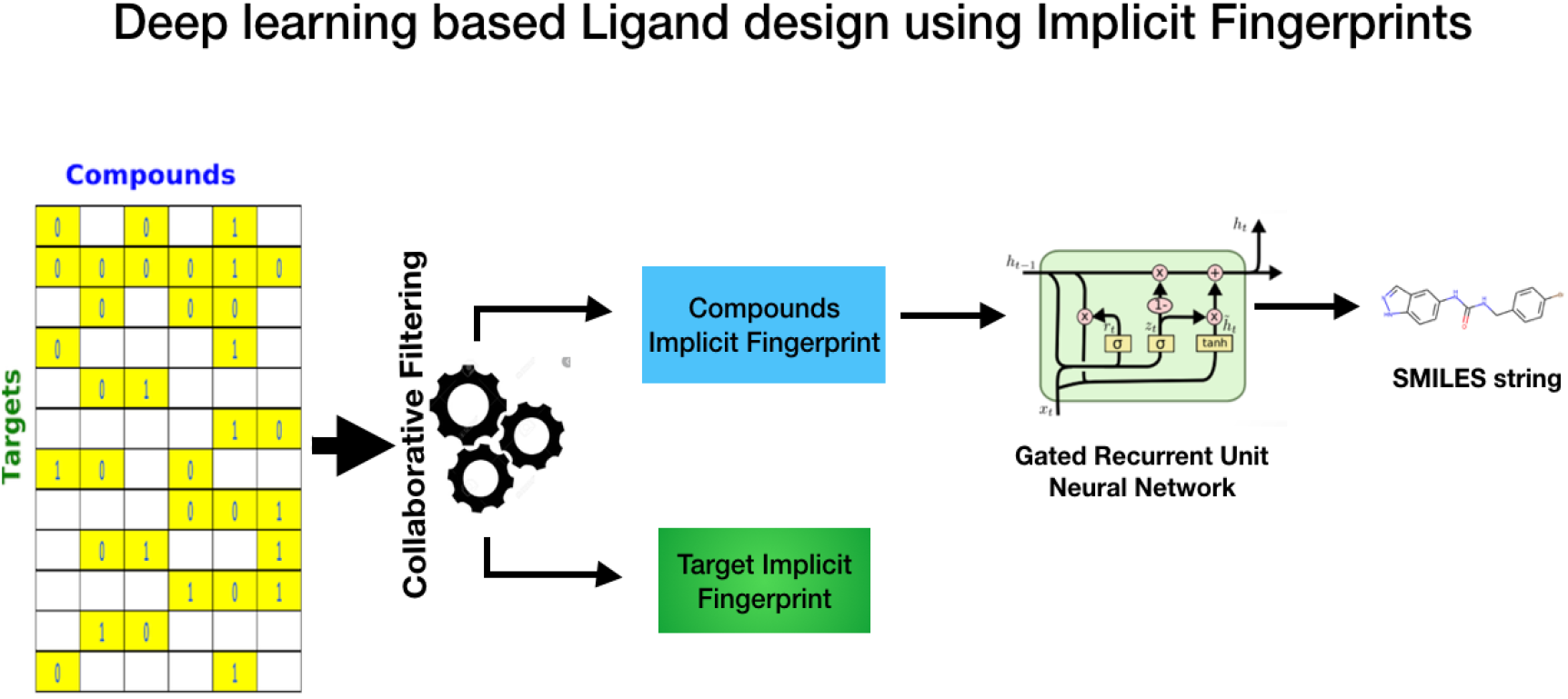
Deep learning based Ligand Design using Implicit Fingerprints from Collaborative Filtering - Architecture

As with other methodologies utilizing generative deep learning algorithms, ^30^ the neural network should ensure that the points in the latent space decode to valid SMILES strings. In order to avoid the latent space from being sparse and resolve to large “dead-areas” (areas in the space that are never trained to decode from and therefore behave unpredictably), we performed input data augmentation. The data augmentation involved adding randomness to the input layer of the neural network (i.e., adding random perturbations to the implicit vector). The data augmentation incentivizes the decoder to more fully represent the areas in the implicit latent space of the ligands, such that they can successfully resolve to the corresponding SMILES string. The intuition is that adding noise to the encoded molecules forces the decoder to learn how to decode a wider variety of latent points and find more robust representations. This approach follows the intuitions made popular by the variational auto-encoders(VAEs)^41^ by Bowman et al. The VAEs, instead of decoding from a single point in the latent space, sample from a location centered around the mean value and with spread corresponding to the standard deviation, before decoding. This ensures that a sample from anywhere in the area is treated similar to the original input. Even so, there are differences between the VAE approach and ours. In our approach, the latent space if fixed from the collaborative filtering; it is not trainable like in the VAE. Importantly, this means that the sampling incentivizes the decoder to reconstruct similar SMILES scores from given set of similar points. It does not incentivize the collaborative filtering algorithm to change its implicit representation.

The sequential nature of the output SMILES string required us to consider neural network architectures that are adept at handling such data. The application of neural network architectures such as recurrent neural networks and their enhanced variations such as gated recurrent neural networks (GRUs) for problems involving sequential data such as speech recognition, language translation have been very successful. ^40,42–44^ The GRU neural networks, with their innate abilities of learning long-term dependencies in sequences, are especially useful for handling SMILES strings.

### Tools for bioactivity prediction

Bioactivity of a drug is of critical importance to highlight the applicability of generated ligands. SSnet^15^ and smina^45^ were utilized to obtain a relative bioactivity of ligands towards various targets tested in this work.

#### SSnet

SSnet is a deep neural network based framework that requires a protein target in pdb format and a ligand as SMILEs string to predict their bioactivity (probability for binding). The protein structure is formatted to extract curvature and torsion patterns of the protein backbone that contains compact information about the fold information. The protein fold is a consequence of multitude of atomic interactions including the side chains and thus holds information about potential ligand interaction. SSnet had outperformed state-of-the-art machine learning models like Atomnet,^12^ 3D-CNN,^46^ and GNN-CNN^13^ and classical force field and knowledge based methods employed by Autodock Vina^47^ and smina^45^ in identifying positive protein-ligand pairs (protein ligand complex with high binding affinity). SSnet being pre-trained has fast execution time (18 minutes for 1M protein-ligand pairs) to curate high affinity ligands from a pool of large libraries such as ZINC (1+ billion ligands). SSnet is made available for public use on https://github.com/ekraka/SSnet.

#### Smina: Scoring and Minimization of Ligand Conformation

To perform virtual screening and docking, smina^45^ was used on a subset of ligands converted to 3D structures via openbabel.^48^ The docking was performed on the centre of a known ligand in a protein-ligand complex with a box size of 32Åx32Ax32A, and exhaustiveness of 36 on the default scoring function. The box size defines the space to consider in a protein for optimizing a ligand conformation resulting to a binding score. The exhaustiveness is an indication of the computation time for optimization. The exhaustiveness is required as Smina utilizes Monte Carlo method for optimization.

## Results and discussion

In this section, We present the details of the experiments conducted with their results. We begin with an exhaustive description of the data used for the experiments.

### Dataset Description

Our method involves translating the implicit ligand fingerprints into its corresponding SMILES string. The implicit fingerprints, however are derived from the ligand - target bioactivity data from the ChEMBL database (Version 23). The bioactivity data, keeping in line with previous studies^49,50^ was focused only on human targets. We restricted bioactivities to three types of binding affinities. The included half maximal inhibitory concentration *IC50,* maximal effective concentration (EC50), and inhibitory constant(ki). Following the precedence with previous works, ^1,49,50^ we converted the data into the binary active-inactive using the following conversation thresholds: lesser than 100 nM for “actives” and greater than 1000 nM as “inactives.” Furthermore, in order to be consistent with Srinivas et al.^1^, we retained only ligands which have at least two prior assays. This resulted in a bioactivity matrix of size 241,260 (ligands) by 2,739 (targets). The bioactivity matrix was subjected to the collaborative filtering method as described in Srinivas et al. ^1^. The resultant implicit fingerprints were then used as inputs to our deep learning model, with the goal to produce the respective canonical SMILES string as the output. Figure 4 illustrates the data distribution of the number of ligands against the known number of prior assays and known number of prior assays with known positive affinities. As evident from the plots, close to 50% of the ligands have only 2 prior assays. Additionally close to 62% of the ligands have only 1 prior assay with positive affinity. We also wish to note that the number of ligands (241k) used to model the deep learning model is comparable to previous works. ^24^

**Figure 4:**
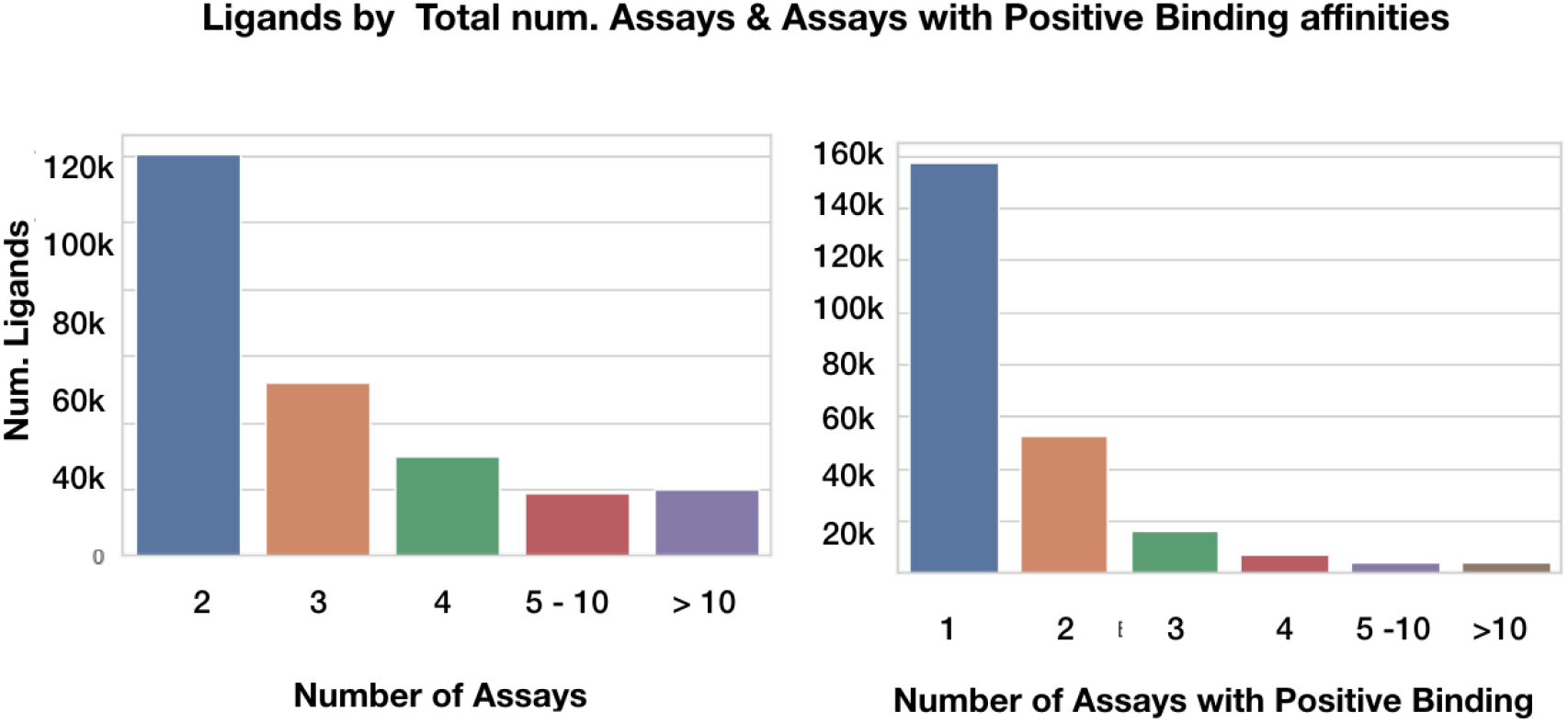
Data Distribution: The first figure illustrates the number of molecules against number of assays, binned at specified values or ranges. Close to 50% of the molecules have only 2 prior assays. The second figure illustrates the number of ligands against the number of known assays with positive affinities. It can be observed that 62% of the ligands with only one assay with positive binding affinity.

Considering that our approach relies on the prior assay history to determine the implicit ligand fingerprints, having more numerous examples of prior assays for each ligand may also result in better quality implicit fingerprints. This statement is further evidenced by results from the next section: (1) the ability of the decoder to accurately translate implicit fingerprints into the corresponding SMILES and (2) the abilities of the ligands to yield more novel ligands are both influenced by the number of available assays per ligand, as described next.

### De-novo generation of molecules from latent space

In this section, we discuss the outcomes of our method in the context of 5,000 randomly selected ligands from the validation set. Additionally we also present the outcomes of a scaffold analysis from novel ligands generated from ligands with known affinities to cancer targets from the ChEMBL23 database. To further analyze the practical applicability of our approach,the resulting novel ligands, specifically from approved cancer related drugs were further evaluated for their viability to be valid drugs with enhanced biological activities. The complete list of ligands is made available as a part of the supporting information.

Our method samples around the implicit latent space of the known ligands, or “anchor ligands” to generate (potentially novel) compounds. In our testing, we randomly sampled 100 points across the 50 dimensions in the implicit space around our anchor ligands. Each point was then processed through our neural network to obtain the corresponding SMILES string. The SMILES string was then validated using the RDKIT library. This process is discussed in more detail in the methods section.

We ran the aforementioned sampling and validation exercise on the 5,000 ligands (henceforth referred to as anchor ligands). A total of 4,632 out of the 5,000 (92.64%) anchor ligands resolved to at least one valid ligand, although not all resolved ligands were novel. As mentioned earlier, 100 points are randomly sampled for each anchor ligand. Depending on the information encoded in the continuous implicit vector space, multiple points around a given anchor ligand may resolve to the same ligand. Only those ligands that are generated at least twice, and can be resolved to a valid compound using the RDKIT library, are considered to be “valid” generated ligands. The frequency constraint of “at least twice “ is enforced to help ensure that the generated ligand is not generated spuriously. Additionally, 1,332 anchor ligands out of the 5000 yielded a total of 1,103 novel ligands (3,079 unique SMILES strings). We define a “novel ligand” as one which did not have sufficient similarity with the 1.2 million compounds from the ChEMBL23 database and the 1.3 billion compounds from the ZINC database. The similarity between compounds were measured by the Tanimoto Coefficient (TC) which measures a distance between fingerprints resulting in a score ranging from [0,1] (0 corresponds to least similar and 1 to exactly same).^51^ We obtained the TC based on 512 bit Morgan Fingerprints.^52^ 1,103 compounds were observed to have TC of less than 0.85, signifying 24 % of all anchor ligands to be novel. The entire list of TC scores for ChEMBL23 and ZINC database for the enumerated compounds are provided in the supporting information. These numbers are reproduced in table 1 for easier readability.

**Table 1.**

|  |  |
| --- | --- |
| Number of anchor ligands from validation set : | 5,000 |
| Number of anchor ligands yielding atleast 1 valid ligand : | 4,632 |
| Number of novel ligands generated from anchor ligands : | 1,103 |
| Number of anchor ligands yielding atleast 1 novel ligand : | 1,332 |

It was also observed that sampling around certain anchor ligands resulted in numerous novel ligands being generated, while sampling around other anchors did not yield *any* novel ligands. To further investigate this phenomenon, we analyzed the abilities of the associated anchor ligands to yield novel compounds by grouping according to known prior assays. Figure 5 (left) illustrates the relationship between the presence of assays of the anchor ligands and their ability to generated novel ligands. As can be seen, anchors that generated novel ligands tended to have a greater number of known assays. In Figure 5 (right), we can also observe that this relationship holds for the number of anchor ligands with positive affinities. Anchors with a more numerous positive assays also tended to generate novel ligands. This observation is perhaps not surprising considering that the implicit fingerprints are derived from the known assays. This provides evidence that the implicit fingerprints for anchor ligands encode more meaningful information when there are more numerous assays. One potential explanation for this is that the implicit representation can encode many desired properties that are difficult to measure as the number of assays increases, thereby giving a better blueprint for the decoder to generate novel ligands. However, because the implicit representation does not explicitly model chemical or experimental parameters, this hypothesis must be investigated through observation of known ligand properties, as discussed next.

**Figure 5:**
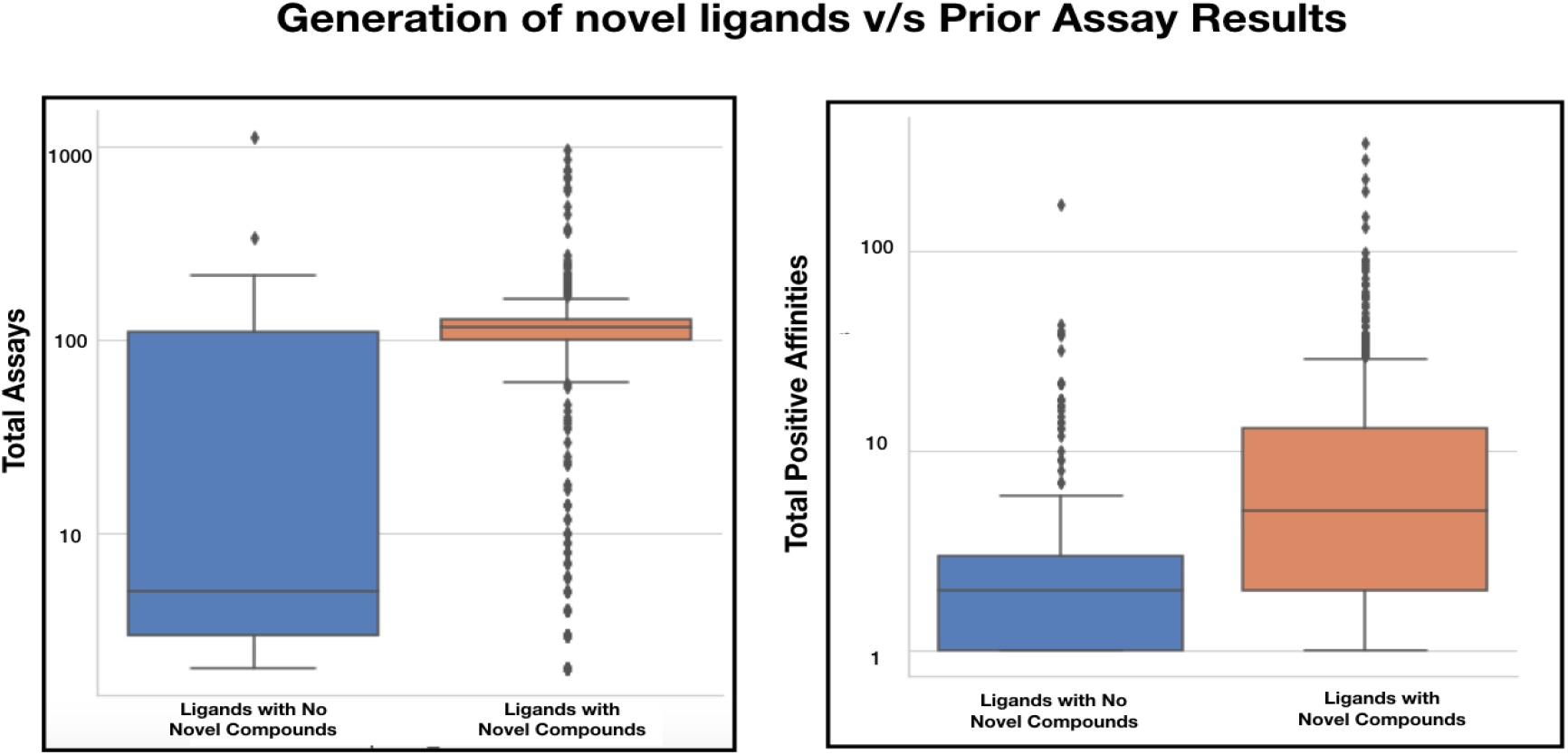
Correlation of the ability to generate novel ligands with prior assays: Box plots show the co-relation between the two sets of anchor ligands - one set from 1 or more novel ligands were generated and the second set which yielded no ligands when sampled in the implicit fingerprint latent space. The first figure visualizes the total number of known assays that exists for each set. The second box-plot visualizes the total number of positive binding affinities already recorded for each assay.

### Physical Properties of Novel and Anchor Ligands

In order to explore the similarity of novel and anchor ligands, we evaluated the properties of the compounds using a number of scoring measures. More specifically, we used the quantitative estimation of drug-likeness (QED), n-octanol water partition coefficient (LogP), synthetic accessibility score (SAS), and used the number of benzene rings as an indicator of the chemical complexity. Our approach to ascertain the similarities in the physical properties between the anchor ligands and its corresponding novel ligands considered the following approaches

- Compare the distribution of the populations of property values of the anchor ligands with the distribution of the generated ligands. This comparison is standard practice when evaluating the quality of generated ligands.^22,53^
- Additionally, to investigate similarity of generated ligands with their respective anchor ligand, we evaluated the magnitude of the difference in the values between the ligands for each of the 4 aforementioned properties. A residual value, which is the difference between the property values is calculated for each pair of anchor ligand and its corresponding generated ligand. A mean residual is then obtained for each anchor ligand as described in equation 1. The magnitude of the mean residual value was used as a method to determine the deviation of the properties between anchors and its generated ligands.

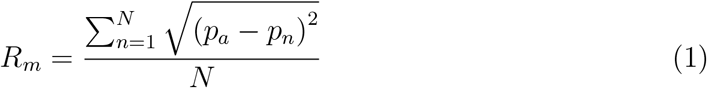 *R_m_*:mean residual property value for each anchor ligand *N*:number unique novel ligands generated for each anchor ligand *p_a_*: property value (QED,LogP,SAS and NumRings) for the anchor ligand *p_n_*: property value (QED,LogP,SAS and NumRings) for *n^th^* novel ligand for the corresponding anchor ligand.

The QED ranges between 0 and 1. The ligands with higher value indicate that the molecule is more drug-like. Additionally, the method also claims to capture the abstract notion of aesthetics in medicinal chemistry. ^54^ We leveraged the python based RDKit library to determine the QED scores of the generated novel compounds. As illustrated in Table 2, the average QED score of the novel ligands was found to be 0.57. Figure 6-A(i) illustrates the comparison of the distributions of the QED scores from the novel ligands with their anchors. It can be observed that the two distributions are very similar. The student t-test statistic of 0.99 with a p-value of 0.35 also confirms that there exists no statistical difference between the two distributions. Table 2 tabulates the mean, standard deviations and t-test scores of all the properties calculated as a part of our experiments. Additionally, figure 6-A(ii) illustrates the similarities of the QED scores between the anchor ligands and their respective generated ligands by measuring the mean residual value as described in equation 1. It is evident from the plot that a large number of mean residuals are less than .1 units. This indicates the QED scores of close to 80% of the anchor ligands are within .1 units of their generated novel ligands, and close to 96% of the anchor ligands have QED scores within .2 units of their generated ligands.

**Figure 6:**
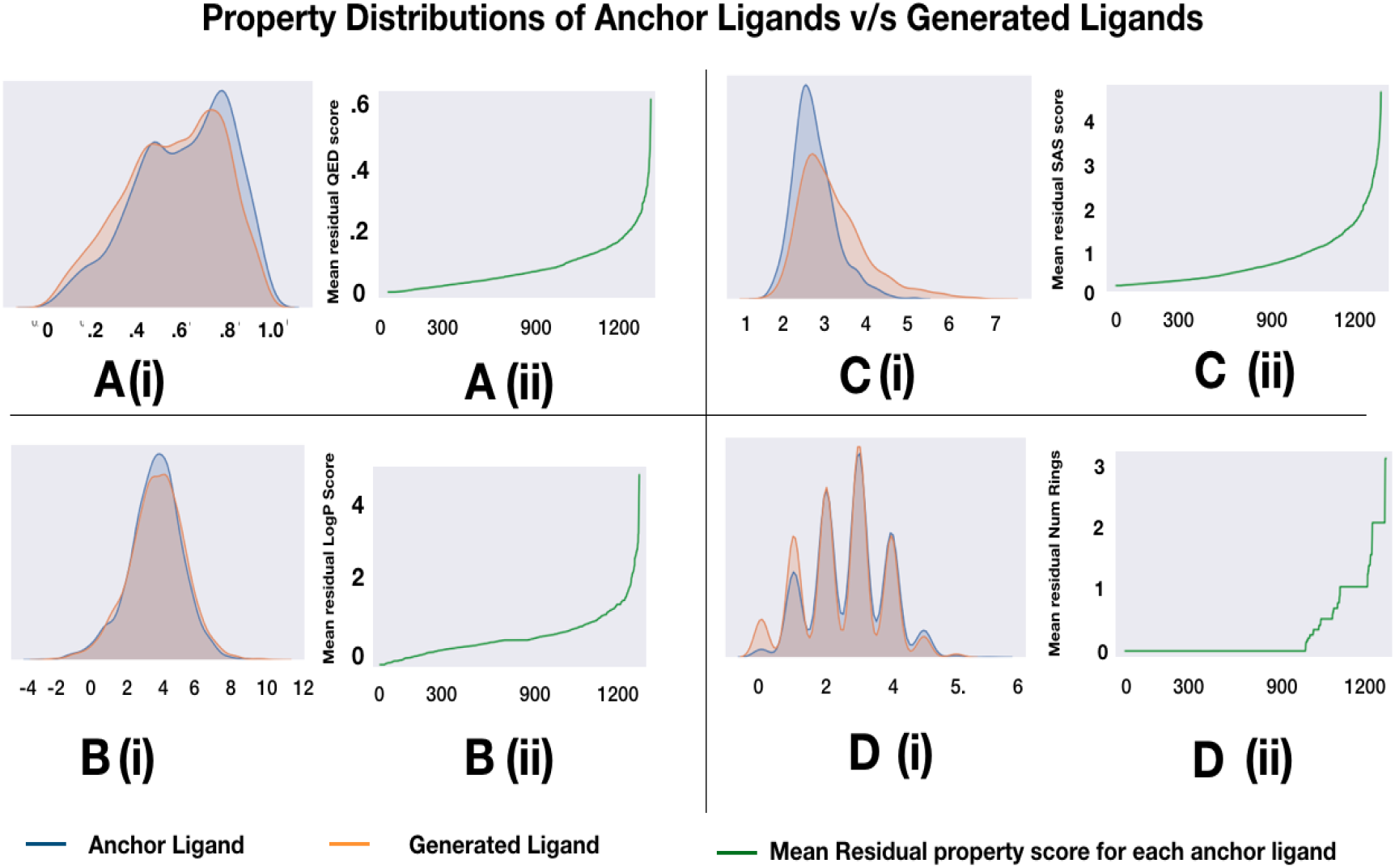
Property Distribution between anchor ligands and generated ligands: (A) Quantitative Estimate of DrugLikeness(QED) (B) Partition Coefficient (LogP) (C) Synthetic Accessability Score (SAS) (D) Number of Benzene rings. The figure demonstrates that the property distributions of the anchor ligands is similar to the novel ligands generated from the corresponding anchors across all 4 properties.

**Table 2:** Properties of anchor and novel ligands.

|  |  | Anchor Ligands | Novel Ligands | t-test |
| --- | --- | --- | --- | --- |
| QED | Mean | 0.69 | 0.57 | t-stat = .99<br>p-value = .35 |
|  | Stddev | 0.20 | .22 |  |
| LogP | Mean | 3.41 | 3.43 | t-stat = 0.33<br>p-value = .74 |
|  | Stddev | 1.69 | 1.95 |  |
| Benzene Rings | Mean | 3.45 | 3.12 | t-stat = -8.23<br>p-value = 2.34e-16 |
|  | Stddev | 1.24 | 1.33 |  |
| SAS Scores | Mean | 2.67 | 3.18 | t - stat = 21.62<br>p-value. = 6.7e-99 |
|  | Stddev | 0.55 | 0.85 |  |

The water-octanal partition coefficient (LogP) was an other property used to quantify the physical properties of the novel ligands. LogP describes the propensity of ligand to dissolve in an immiscible biphasic system of lipid (fats, oils, organic solvents) and water.^55^ A negative value for logP means the ligand has a higher affinity for the aqueous phase(hydrophilic);when logP = 0 the ligand is equally partitioned between the lipid and aqueous phases; a positive value for logP denotes a higher concentration in the lipid phase(lipophilic). The novel ligands tended to be more lipophilic with a mean LogP value of 3.43 with a standard deviation of 1.94. Figure 6-B(i) illustrates that distributions of LogP scores between the novel and anchor ligands. The two distributions appear to be visually similar and the student t-test score of .33 with p-value = .74 also confirms the same. Additionally figure 6-B(ii) illustrates the similarities of the LogP scores between the anchor ligands and their respective generated ligands by measuring the mean residual score as described in equation 1. It is observed that close to 87% of the anchor ligands have their LogP scores within 1 unit of their generated ligands.

The synthetic accessibility score (SAS), a method that is able to characterize molecule synthetic accessibility as a score between 1 (easy to make) and 10 (very difficult to make)^56^ was another property that was evaluated for the novel drugs generated by our method. The mean score was found to be at 3.17 with a standard deviation of .85. While the SAS scores between anchors and their novel ligands appear to be similar visually (6-C(i)), the t-statistic score of 21.62 with p-value=6.7e-99 indicate that the two distributions are statistically different. Nevertheless the mean score of 3.17 of the novel ligands indicate that the novel ligands are synthesizable to generate valid drugs. Figure 6-C(ii) further compares the individual SAS Scores between the generated ligands and their respective anchor ligands. It is observed that 87.3% anchor ligands have SAS Scores within 1 unit of the generated ligands. This indicates that an overwhelming majority of the anchor ligands share similar SAS Scores with their generated novel counterparts. Additionally the number of Benzene rings was evaluated as a measure of chemical complexity of the novel ligands. Figure 6-D(i) demonstrates that the complexities of the novel drugs are comparable to the complexities of their corresponding anchor ligands. Figure 6-D(ii) compares the similarities in the number of benzene rings between the anchor ligands with their respective generated novel ligands. It was observed that approximately 83% of the anchor ligands had the exact same number of benzene rings as their respective generated novel ligands.

We further evaluated the Lipinski’s rule of 5 (LR5) for all the generated ligands.^57^ The LR5 describes critical properties of a ligand in the human body such as absorption, distribution, metabolism, and excretion. The rule states that a ligand to be effective for therapeutics should have less than 5 hydrogen bond donors, less than 10 hydrogen bond acceptors, a molecular mass of less than 500 daltons and the LogP less than 5. The LR5 score was computed for all generated ligands based on Yao et al.^58^. We observed that 68% of generated ligands completely satisfies the LR5 rule and 22% of generated novel ligands satisfy atleast 3 out of the 4 rules. This is further illustrated in figure 7. The percentage of matches to Lipinski’s rule of 5 signifies that the generated ligands have properties to be an effective drug.

**Figure 7:**
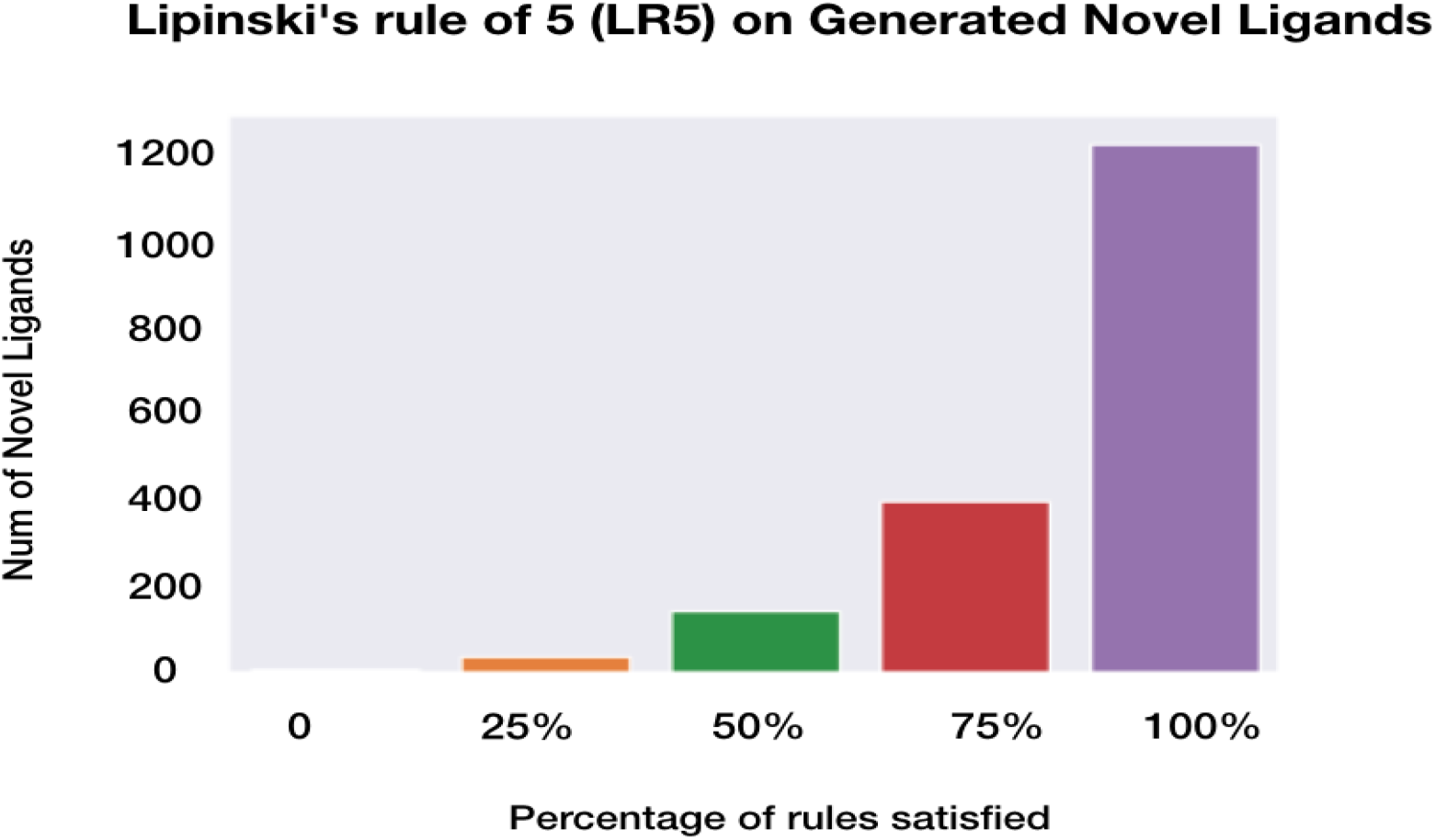
Lipinski’s Rule of 5 valuated on the novel ligands generated from implicit fingerprints. The figure demonstrates that 80% out of the 1,103 novel ligands satisfy 3 or more rules, signifying that the generated ligands have properties to be an effective drug.

Now that it is established that the novel and anchor ligands are likely to have similar and comparable physical properties, we turn our attention to answering whether the novel ligands are also likely to similarly bind to known targets.

### Binding Affinity predictions of novel ligands

The biological activities of the novel ligands were evaluated by inferring their binding affinities with 102 DUD-E protein targets. ^59^ The DUD-E targets consist of a variety of proteins exhibiting different mechanism of protein-ligand interactions. The relationship of bioactivities within the anchor ligand and generated ligands over the DUD-E targets will highlight the versatility of our model. Thus we used the anchor ligands to test their binding affinities with the DUD-E targets.

In order to validate the similarities of the binding affinity properties of the novel ligands with their respective anchors, the binding affinity scores were determined from SSnet for the anchor ligands with the 102 DUD-E proteins. Each ligand (anchor and novel ligands) yielded a distribution of binding affinity scores against each target from the set of 102 DUD-E protein targets. The similarities in the binding affinities of the novel and their respective anchor ligands were evaluated by comparing the aforementioned binding affinity distributions. Out of the total 1,332 unique combinations of novel and respective anchor ligands, approximately 84% demonstrated similar binding affinity behaviors. The similarity score or the measure of Intersection over Union, ^60^ in this exercise is calculated by evaluating the proportion of DUDE targets to which both the ligands demonstrate binding or lack of binding. An SSnet score of 0.5 or less is considered lack of binding, and a score greater than 0.5 as binding. Figure 8 illustrates this for 1,332 unique pairs of novel ligands and their anchor ligands. Each data point on the x - axis in figure 8 represents a unique anchor-novel ligand combination. The y-axis represents the Intersection over Union score calculated between the 2 distributions of binding affinity scores, the first distribution being binding affinity indicator of anchor ligand with 102 DUD-E proteins and the second distribution, the binding affinity indicator of the novel ligand with 102 DUD-E proteins. The figure further illustrates that a large majority of the anchor-ligand pairs exhibit similar binding affinities.

**Figure 8:**
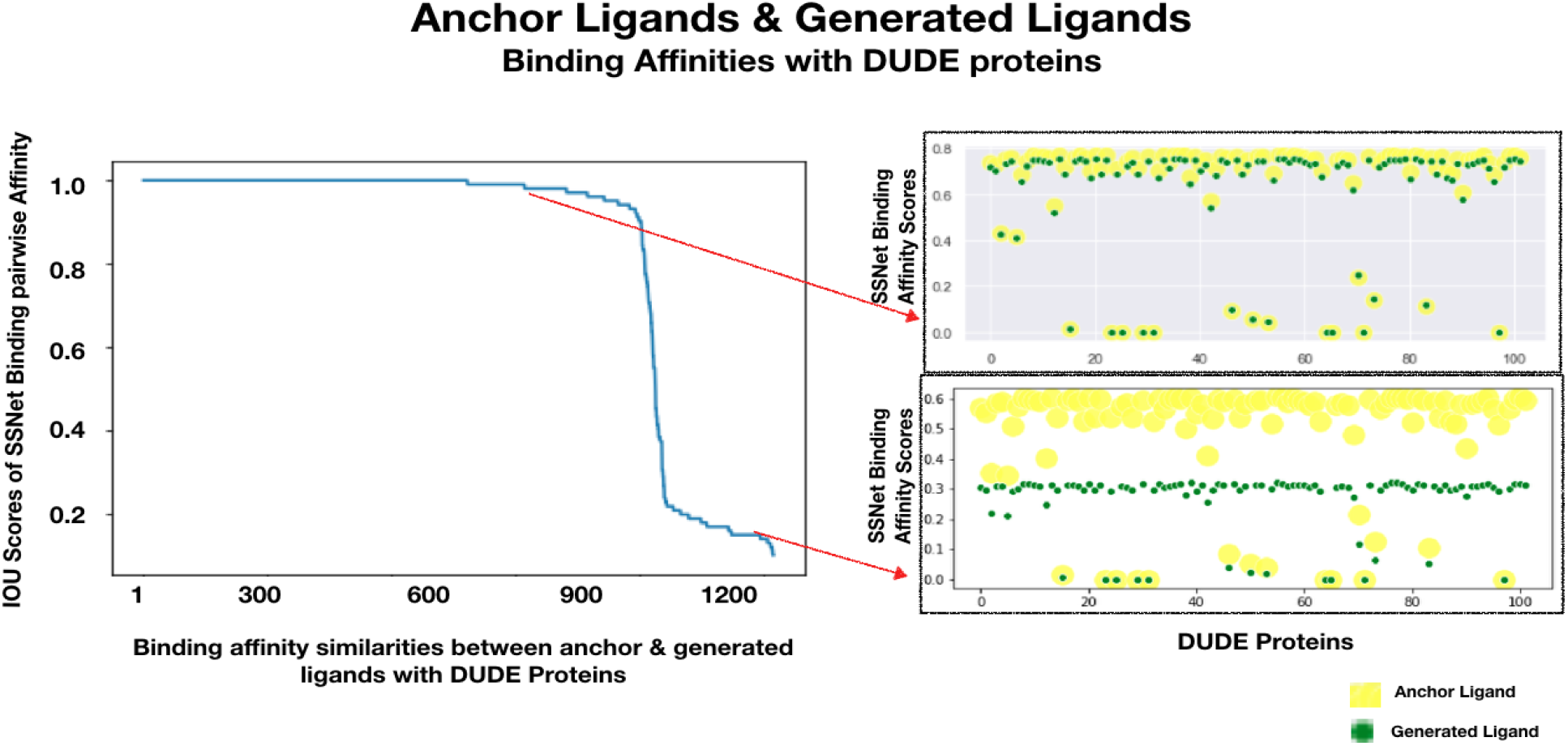
Pairwise binding affinity scores: The plot illustrates the similarities in the bioactivity between each pair of anchor ligands and their corresponding generated ligands to the 102 DUDE protein targets. The large regions of dark blue hue in the heat map demonstrate a strong co-relation between the binding affinities for most pairs with the DUDE targets. The scatter plot illustrates two sample pairs, with the top right plot representing a pair with very similar affinity scores, with the bottom right plot illustrating a pair where the affinities differ between the anchor and generated ligand.

The analysis on QED, LogP, and SAS provided an intuitive relationship of generated ligands and drug-likeliness. However, for a drug to be effective for specific target and show selectivity among other targets, should preserve the scaffold (core structure of a molecule^61–63^). To analyze if the generated ligands have similar scaffolds, we sorted all the anchor ligands by Tanimoto Coefficient (TC). The sorting was performed by recursively finding next most similar ligand from the anchor ligands starting from a random anchor ligand. The sorted list was then mapped to a pseudo-Hilbert space filling curve. The pseudo-Hilbert curve was used to observe molecular scaffolds directly from the map as pseudo-Hilbert curve preserves the spatial proximity of the sorted list. The pseudo-Hilbert map for the generated ligands were made similarly. Each anchor ligands were repeated to the same number of generated ligands in order to match one-to-one when comparing pseudo-Hilbert curve for generated ligands and anchor ligands. Figure 9a and 9b shows the pseudo-Hilbert map for anchor ligands and generated ligands respectively. The pseudo-Hilbert map is colored based on the SSnet scores obtained by docking the ligands with the DUD-E targets with PDB ID 1B9V and 3KBA. The pseudo-Hilbert map for all 102 DUD-E targets are provided in the supporting information. We observe that the clusters are majorly retained for the generated ligands when compared to the anchor ligands. This is further highlighted in Figure 9c, that shows the difference in SSnet scores for generated and anchor ligands. The map is mostly blue which represents a mere difference of SSnet scores in generated and anchor ligand of less than 0.1. The results highlight that the novel molecules generated preserves the scaffold that is essential in protein ligand binding.

**Figure 9:**
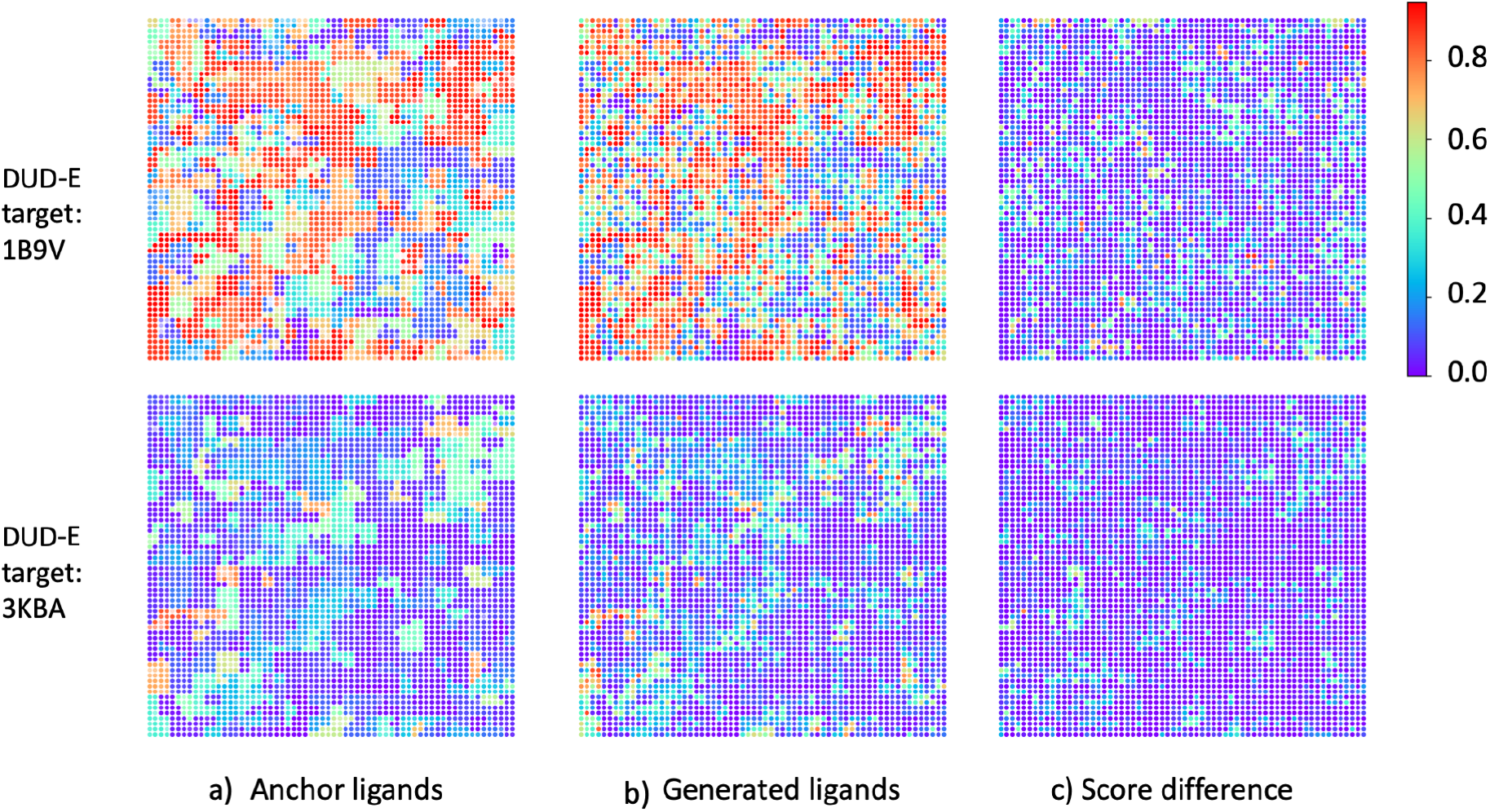
Scaffold analysis. A pseudo-hilbert curve is plotted for anchor ligands and generated ligands. The color denotes SSnet scores. Similarity between anchor and generated pseudo-hilbert curves and the low difference among them, signifies that our method retains scaffolds from the anchor ligands while also predicting similar bioactivities.

### Comparison with FDA Approved Drugs

In order to further hone in on the practical applicability of our methods and the novel drugs generated, we conducted analysis on novel drugs generated on known cancer related ligands. For this exercise we shortlisted 10 drugs approved for treating various forms of cancer also available in the ChEMBL23 database. We present detailed analysis of the novel ligands generated around a known cancer drug, DASATINIB. Sampling around the implicit fingerprint space of this anchor ligand yielded 10 novel ligands. Figure 10 illustrates the 10 novel ligands. The novelty of the compounds were tested from the ChEMBL23 dataset (1.4 million compounds) and the ZINC dataset (1.3 billion compounds). Across the 10 novel compounds, the maximum similarity score were 0.88 for ligands in the ChEMBL23 dataset and 0.92 for ligands in the ZINC dataset. Table S1 shows the largest TC obtained for each novel compounds. Interestingly, in this particular case, we observe that the scaffold for the anchor ligand is retained in most of the generated ligands. The results are in line with the scaffold analysis performed for the DUD-E protein targets provided in the previous section (Figure 9). Retention of scaffold is crucial for ligand binding as the protein pocket in general has confined space for docking. The scaffold provides both size and imperative interactions such as hydrogen bonding, π interactions etc. that contributes to the stability of the protein-ligand complex.

**Figure 10:**
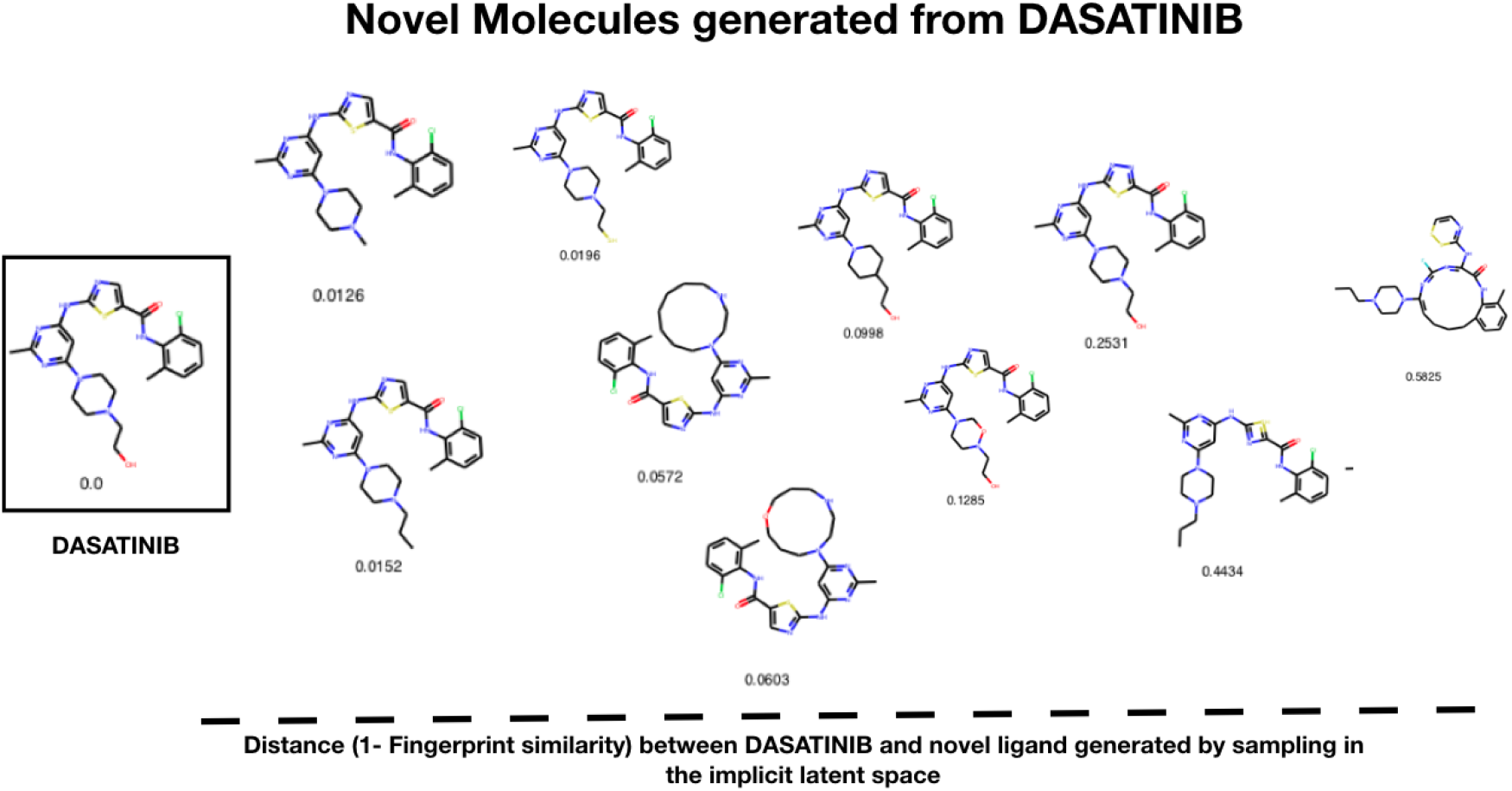
Novel ligands generated around known cancer drug, DATASINIB: Need to expand after analysis on Binding affinities.

To test the bioactivities for the novel ligands generated, we sorted 9 known targets for the anchor ligand, the details of which is provided in Table S2. We conducted a docking method Smina^45^ and a deep neural network based model SSnet^15^ for bioactivity score prediction. Figure 11 shows the results obtained by the two methods. The first five targets are labeled active and the remaining four as inactive for the anchor ligand used in the ChEMBL dataset. We observe that the generated ligands have similar Smina scores as the anchor ligands. A similar behaviour is observed when comparing the SSnet scores for anchor and generated ligands. It is important to note that both Smina and SSnet are sensitive to ligand and their complex interaction with a protein target. Many factors such as functional group, size of the molecule, molecular weight etc. govern the bioactivity. The fact that all the 10 novel generated ligands have similar bioactivities provides evidence that our ligand generation method produces ligands with similar binding characteristics to the anchor ligand.

**Figure 11:**
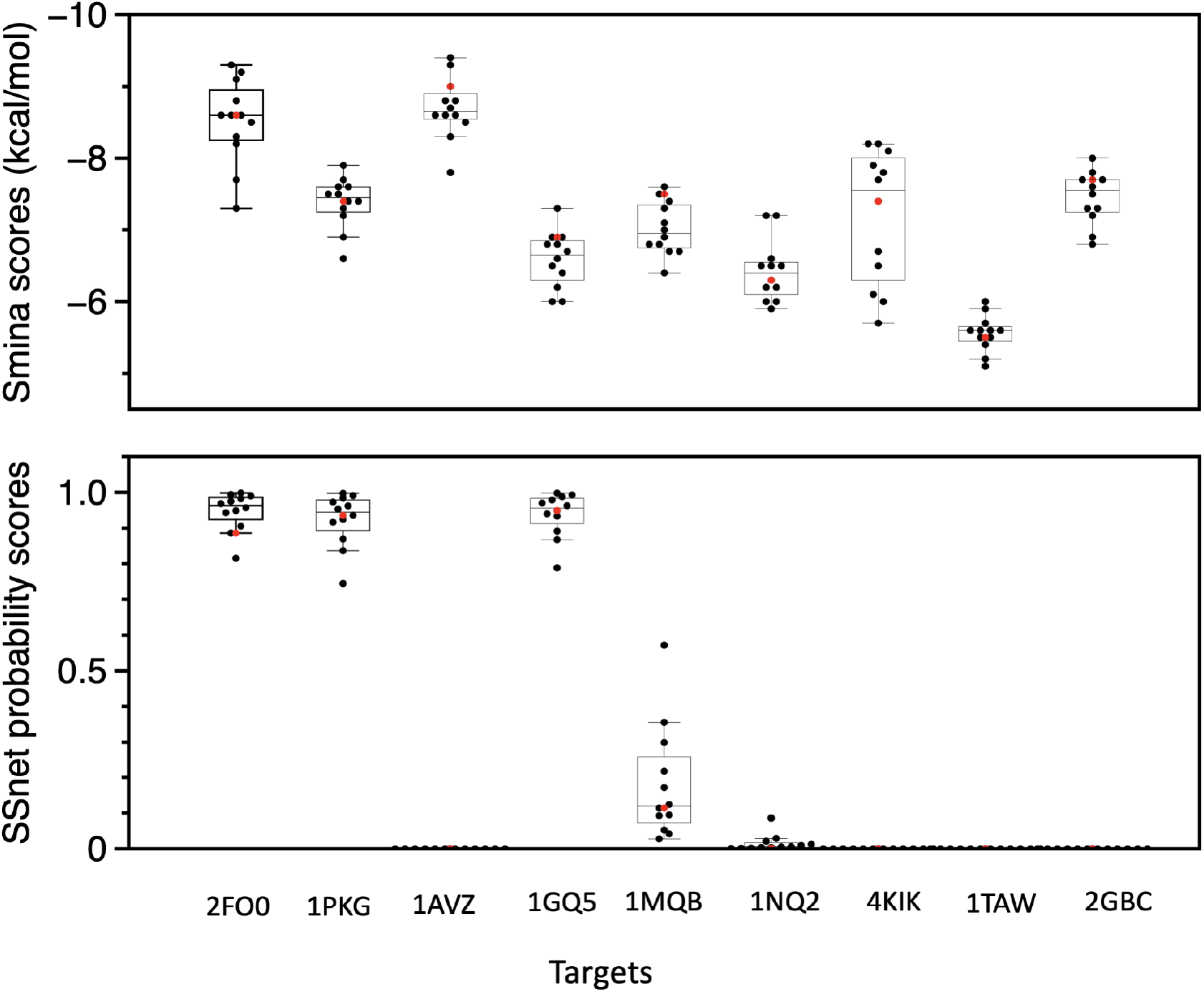
Sample test on various active/inactive targets for anchor ligands. The first 5 targets 2FO0, 1PKG, 1AVZ, 1GQ5 and 1MQB are active and the rest inactive. The red color denotes the anchor ligands and the black denotes generated ligands respectively.

We further compared the bioactivities of 6 FDA approved drugs and their corresponding generated ligands from the implicit fingerprint and latent space generated from the variational autoencoder (VAE) work from Gómez-Bombarelli et al.^30^ respectively. Each of the 6 ligands were docked towards their original intended target, the details of which is provided in Table S3. We observe high similarity in predicted bioactivities for implicit fingerprints compared to the latent space generated from VAE for both the Smina and SSnet scores shown in Figure 12 (Table S5-10). A visual inspection of the compounds generated from our method and the latent space from VAE shows that both retain the scaffold of the original anchor ligand (Figure S1-6). However, bioactivity is sensitive to small changes in the chemical structure such as a functional group. Our method is perceptive towards functional groups due to the way collaborative fingerprints was modeled, i.e. by considering the bioactivities.

**Figure 12:**
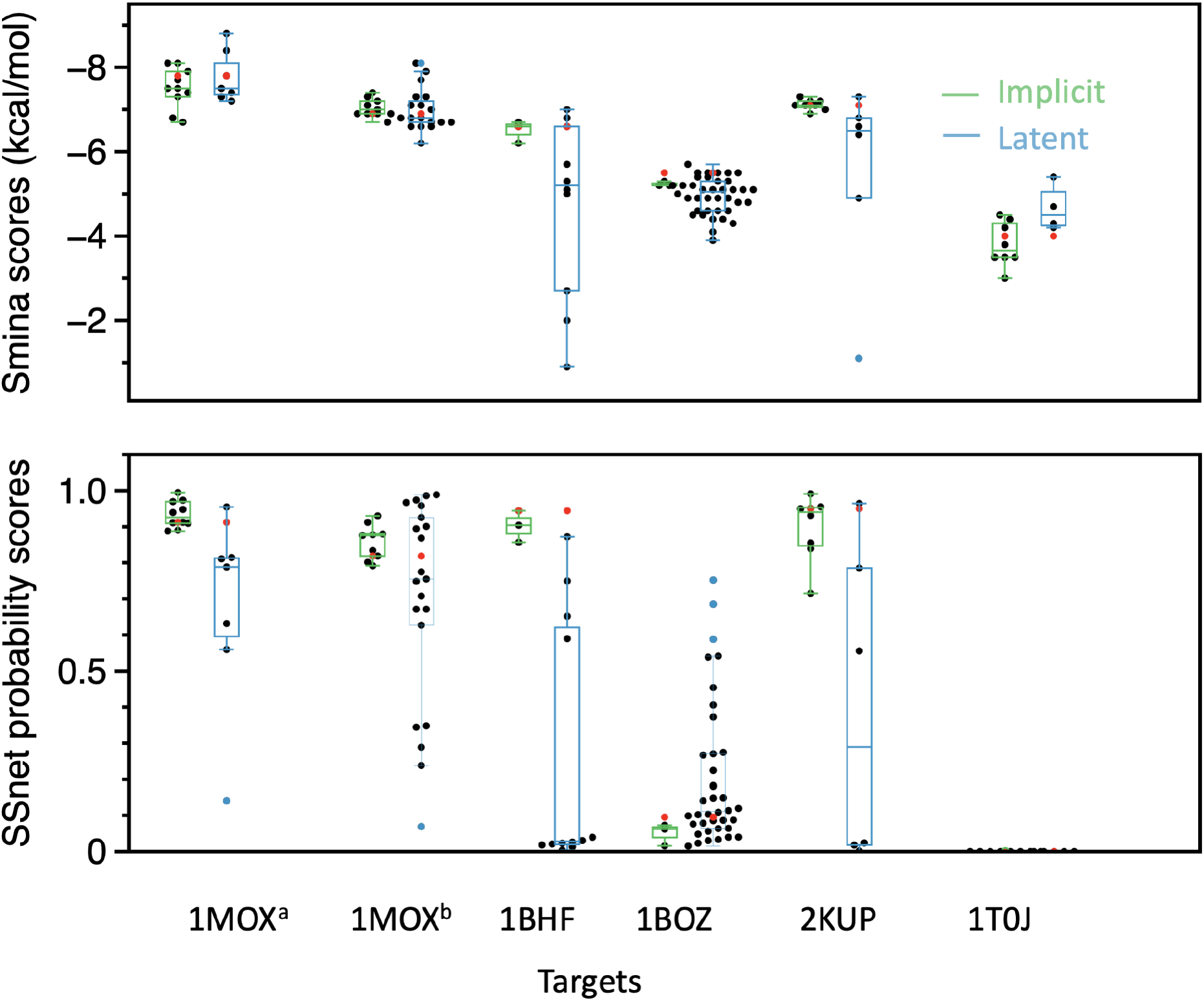
Comparison of Implicit and latent fingerprints on FDA approved drugs and their corresponding targets. The red color denotes the anchor ligand and the black color denotes generated ligands respectively. The latent label and implicit label shows the binding affinities for generated ligands from the method developed by Gómez-Bombarelli et al.^23^ (in blue) and our method (in green).

## Methods

The recent years have seen numerous deep learning based generative models for de-novo drug generation. The common theme in these techniques is to provide a deep learning model with anchor ligands to produce novel ligands with similar properties. Often the canonical SMILES notation of the ligand or a graphical-based fingerprint is used as input to these deep learning models. This representation is then translated into a continuous vector representation(s) of the input ligand, whereby the intermediate layers in the deep learning model are slightly perturbed (i.e., with additive noise) to produce novel molecules. Many previous works exist with the main distinguishing characteristic among the works being the architecture of the deep learning model and classification task employed for training. Some popular methods have been Recurrent Neural Networks,^24^ Variational AutoEncoders (VAEs),^25^ Generative Adversarial Networks (GANs),^26,27^ and graph-based neural networks.^28,29^ A survey of recent work is available from Chen et al.^64^.

Our approach stands apart in that we use the implicit ligand fingerprints obtained from the prior assay information (collaborative filtering) as inputs to a deep learning model, with the objective of producing the corresponding canonical SMILES representation as the output. This implicit representation can have a number of advantages because it is based solely on the observed behavior of the compound, rather than inherent measures of physical properties. Thus, formulating a decoding procedure from this implicit representation may have distinct advantages over previous methods. The implicit fingerprint, because it is a continuous vector of fixed length (50), also lends itself well to statistical sampling with simple procedures. We employ data augmentation of the input vector by employing a vector of means *μ* and another vector of standard deviations, σ. The input vector (implicit fingerprint vector) serves as the vector of means, which is then added to another vector,which is a random normal distribution centered at 0 with standard deviation σ, to yield a statistically sampled point around the implicit fingerprint. The stochastic sampling process ensures that the actual vector will vary on every single iteration due to sampling, while keeping the mean and standard deviations the same. Intuitively, the mean vector controls where the implicit fingerprint of a ligand is centered around, while the standard deviation controls the “area,” how much from the mean the encoding can vary. The decoder, hence learns that not only is a single point in latent space referring to ligand, but all nearby points refer to the same ligand. The decoder is exposed to a range of variations of the encoding of the same input during training. This process is illustrated in the decoder architecture, as shown in figure 13. The approach adopted here is similar to the data augmentation employed with Variational auto-encoders. ^65^

**Figure 13:**
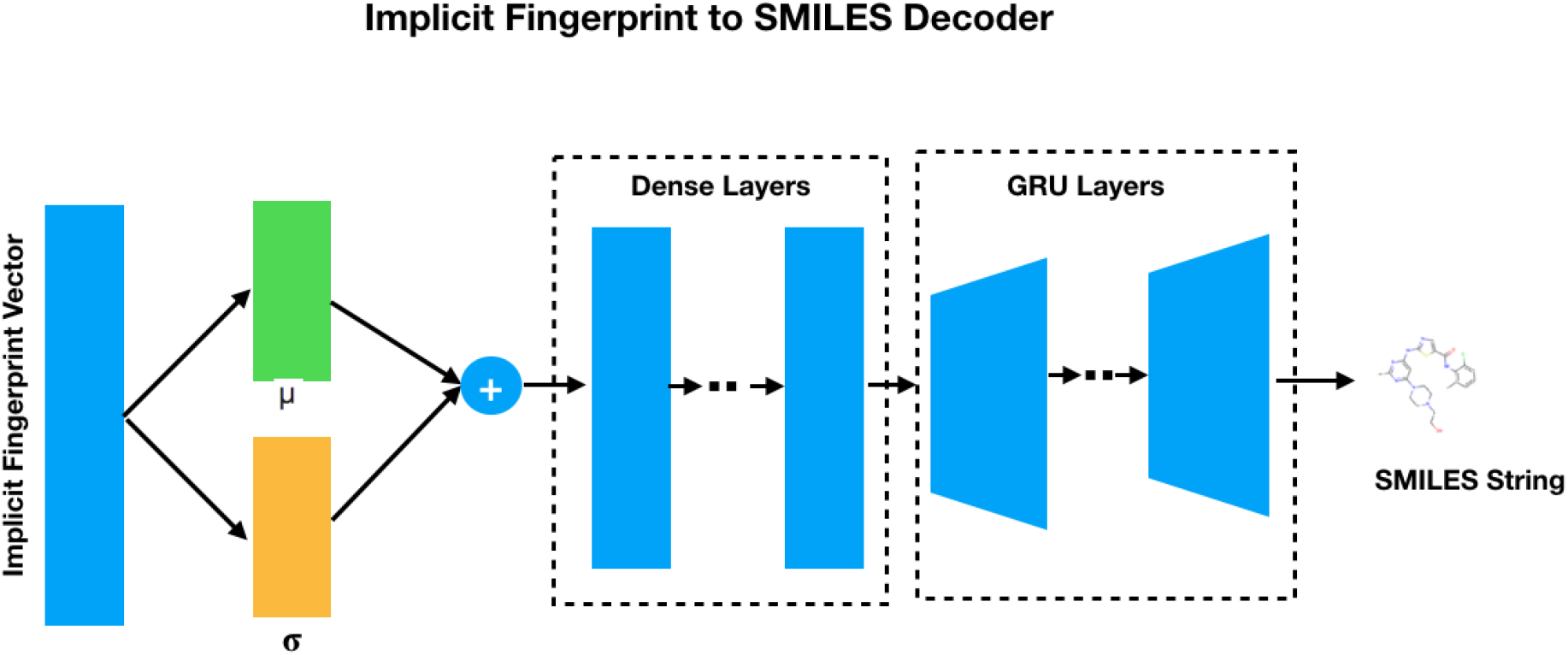
Implicit Fingerprints to SMILES Decoder: The deeplearning network learns ligand representations by employing data augmentation technique at the input layer.The continuous representation obtained is then fed into a series of dense layers followed by a Gated Recurrent Unit Neural Network to obtain the corresponding SMILES string.

In order to generate the SMILES string from the implicit fingerprint, we are motivated to use recurrent neural networks (RNNs) because of their success in modeling sequential data such as natural language. The SMILES strings lend themselves well to this model considering the sequential nature of the notation. Each unit in the RNN attempts to capture state information of the sequence by transforming all the elements that appeared before it. It does so by encapsulating this information in a hidden state vector, that is passed from one unit back into itself, recurrently. The hidden state *h^t^* of the RNNs can be represented as

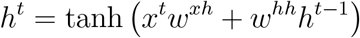

where *x^t^* represents the input at timestep *t, w^xh^* represents the weight from input node to the hidden node, *w^hh^* represents the weight on the feedback loop from the hidden node to itself, and *h*^*t*–1^ represents the previous hidden state. As evident from the equation,the hidden states from the earlier time steps get diluted over long sequences. This problem gets compounded with SMILES considering the long term dependencies (such as matching brackets etc.) that need to be maintained in order to resolve to a valid chemical compound. The Gated Recurrent Neural Network attempts to address this problem by introducing two gates called the “update” gate and a “reset” gate along with a memory which governs how much of the previous state is retained. Each of these units (update gates, reset gate, and memory) have their own trainable weights. The Update Gate at each unit decides the amount of new information to be added to the hidden states. The reset gate determines the past information to be forgotten or retained at each unit.

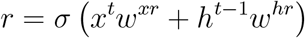

where *σ* represents a logistic or sigmoid function. These sigmoid values of the reset gate range from 0 to 1 and determine how much of the previous hidden node value is retained. A value of *r* = 0 implies the none of the previous node value is retained and a *r* =1 ensures the entirety of the previous node is retained. This memory *m,* can be signified by the following equation:

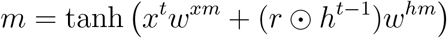

where Θ represents the Hadamard (or element-wise) multiplication of two vectors. Additionally the update gate is governed by the following equation:

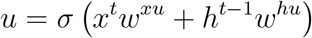

The update gate, with values ranging from 0 to 1, determines if the new hidden state should use the previous value or the new value. Tying all these together, hidden state is governed by the equation:

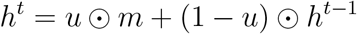

Intuitively, the GRUs are better suited than the RNNS to our problem considering the long term dependencies between symbols that must be maintained in the SMILES string. The Ring structures, for example, are represented by matching numeric symbols typically separated by two or more atoms within the SMILES string. The neural network should be able to remember these long term dependencies for effectively decoding to a valid SMILES string. Figure 13 illustrates multiple GRU layers that make up the decoder in order to effectively map to the SMILES code. These “layers” shown in the figure are visualized compactly—that is, they are actually two stacked sequences of GRU nodes.

### Neural Network Design

We performed extensive training and validation of the vanilla RNN and GRU models with the continuous implicit fingerprint vector as the input and the one hot encoded SMILES string as the output. We measured the outcome of the models by evaluating the categorical cross entropy loss and the accuracy. As a part of training, we explored a variety of architecture options with respect to the depth and the width of the deep learning models. We also trained with different composition of the training sets based on ligands with varying counts of past assay data. Across all training iterations, we noticed that the GRU based model performed better with,lower cross entropy and higher accuracy. We also noticed that the training loss converged faster compared to the validation loss. Our architecture comprised of a series of dense layers followed which consume the 50 vector wide implicit fingerprint representation of the ligands, followed by the GRU layers returning sequential information to map to the SMILEs representations. The exact makeup of the deeplearning architecture with the trainable parameters are provided in the supporting information.

Figure 14 illustrates the performance of the 3 different models trained with different datasets. The first dataset comprised of all the 241K ligands from the dataset. We additionally trained with training set comprised only of ligands with atleast 1 positive binding affinity (121k ligands) and another iteration comprising of ligands with atleast 5 positive binding affinities (8.5k ligands). As evident from the figure, the training and the corresponding validation loss was lower when trained with the filtered data sets as opposed to the entire population of 241k ligands. This can be attributed to the fact that the implicit fingerprints of the ligands that exhibited positive binding affinities in prior assays tend to encode more information on the ligands, and hence decodable into the explicit SMILES representations. While this implies that approximately half the ligands in our dataset do not resolve back to its corresponding SMILES representation, it does not however dent the utility of our approach. This is due to the fact our approach is able to resolve the implicit fingerprints of the ligands which have demonstrated bioactivities in the past, and hence such ligands are more desirable to be used as anchor ligands from which to generate novel ligands.

**Figure 14:**
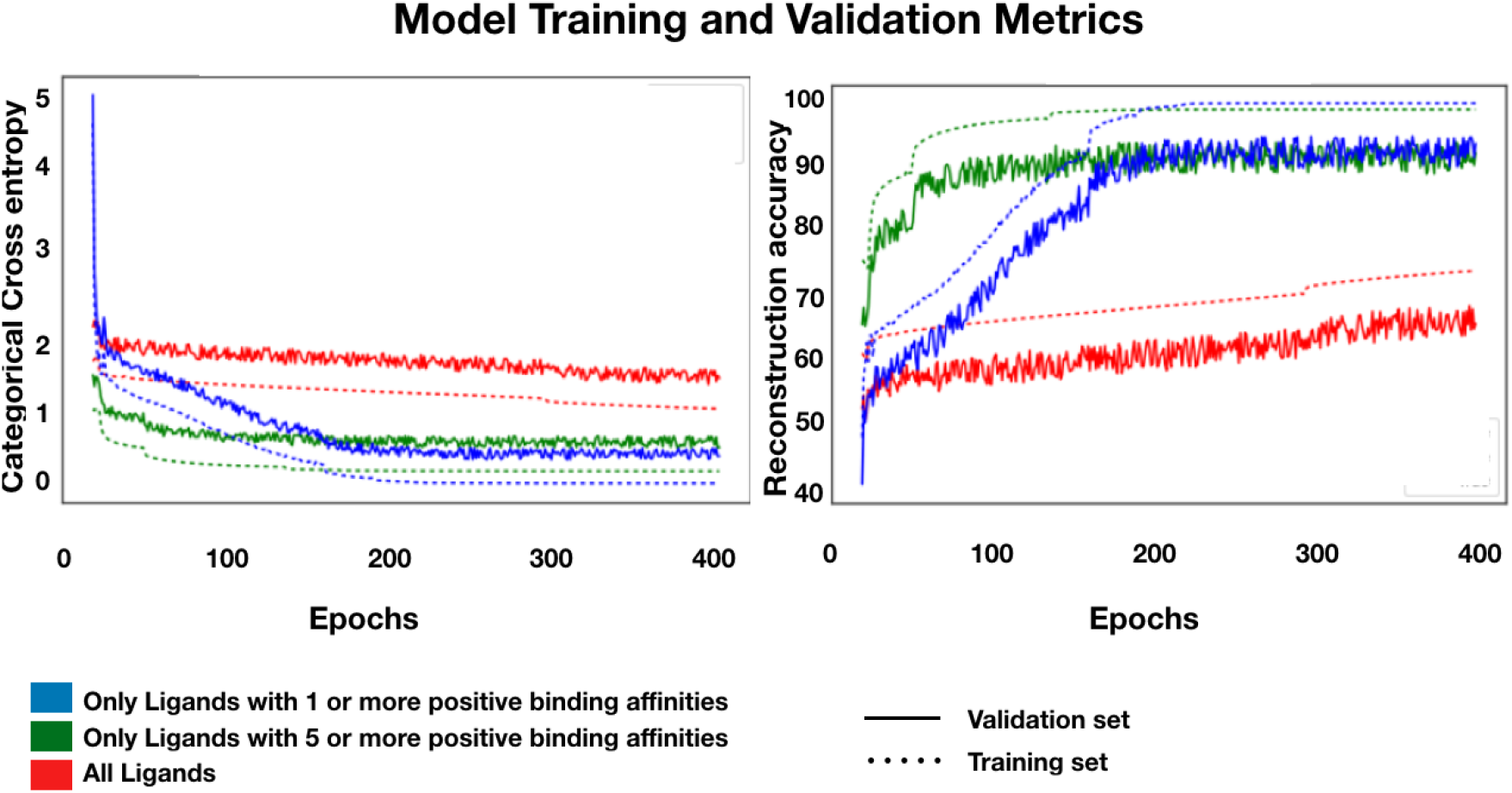
Train and Validation Losses of the Neural Network: Training and validation losses across multiple runs of the neural network.

## Conclusions

We conclude that our approach of marrying the proven collaborative filtering approach with generative deep learning models is a promising new method for de-novo drug generation. Our work shows that the implicit fingerprinting has a number of advantages in terms of encoding the desired properties of the ligands, including binding affinities to known proteins without explicitly optimizing for said chemical properties. The compounds from the implicit space also demonstrated a wide diversity when measured using the Tanimato distance. The collaborative filtering approach allows for the implicit fingerprints to be generated for any novel ligand with desired binding affinities to known target proteins. Leveraging these implicit fingerprints with encoded SMILES representations as the basis to generate useful novel drug-like compounds could further advance this exciting field of drug discovery using generative deeplearning models. We also note that our approach fundamentally relies on having training data for a particular anchor ligand and particular target. In order to create an implicit finger-print, the factorization employed in collaborative filtering requires assay examples. This requirement limits the scalability of the approach to ligands and targets for which assays are available or can be completed. We also point out that our analysis was completed on a large subset of the ChEMBL database, Version 23. Therefore, the consistency of the approach for ligands across different bio-activity databases needs to be further evaluated. Additionally,we note that we considered the implicit fingerprints based on binding affinities alone in this study,there are numerous desired properties (absorption, distribution, metabolism, excretion, toxicity, promiscuity, and pharmacovigilant properties) for which a ligand could be screened. These are typically referred to as secondary screens because they are most often (but not always) screened after affinity has been established. The cumulative generative capabilities by combining implicit fingerprints from these assays could also be evaluated in the future. We also note that further studies need to be conducted on the cumulative generative powers of the SMILES based generative algorithms and the implicit fingerprints generated from collaborative filtering. Future work in this space could leverage the implicit fingerprints with the other popular methods of sampling around the dense continuous representations of the SMILES vectors.

## Supporting Information Available

The supporting information is available for this project in SupportingInformation_ImplicitFPtoSMILESpdf. Additionally the source code and the trained deeplearning model is available at https://github.com/rsrinivas-repo/deepbind_molgen

