## Supplemental Information for "Deep Learning-based Ligand Design using Shared Latent Implicit Fingerprints from Collaborative Filtering"

**November 19, 2020**

November 19, 2020

0.5 List of All Novel Ligands generated from cancer related anchor ligands. . 108

### 0.1 Introduction

This document contains all the necessary supporting information for the research article titled Deep Learning-based Ligand Design using Shared Latent Implicit Fingerprints from Collaborative Filtering

### 0.2 Novel Ligands generated around 6 approved drugs

Table 1 enumerates 6 known cancer related drugs we used as anchor ligands to generate novel ligands.

**Table 1:** Details of 6 SMILES and protein target pairs

| SMILE | PDB ID | Target name |
|---|---|---|
| COc1cc2ncnc(Nc3ccc(F)c(Cl)c3)c2cc1OCCCN4CCOCC4 (GEFITINIB) | 1MOX | Human Epidermal Growth Factor Receptor |
| COCCOc1cc2ncnc(Nc3cccc(c3)C#C)c2cc1OCCOC | 1MOX | Human Epidermal Growth Factor Receptor |
| Cc1nc(Nc2ncc(s2)C(=O)Nc3c(C)cccc3Cl)cc(n1)N4CCN(CCO)CC4 (Dasatinib) | 1BHF | P56LCK SH2 DOMAIN INHIBITOR |
| CCc1nc(N)nc(N)c1c2ccc(Cl)cc2 (PYRIMETHAMINE) | 1BOZ | OXIDOREDUCTASE |
| CC(Oc1cc(cnc1N)c2cnn(c2)C3CCNCC3)c4c(Cl)ccc(F)c4Cl(Crizotinib) | 2KUP | PTB domain of SNT-2 |
| COc1ccc(CCN(C)CCCC(C#N)(C(C)C)c2ccc(OC)c(OC)c2)cc1OC | 2BE6 | CaV1.2 IQ domain |

#### 0.2.1 Novel ligands generated around anchor ligand DASATINIB

**Table 2:** Known assays for DASATINIB from ChEMBL23

| Targets | PDB ID | Bioactivity |
|---|---|---|
| Tyrosine-protein kinase ABL | 2FO0 | active |
| Stem cell growth factor receptor | 1PKG | active |
| SRC | 1AVZ | active |
| Platelet-derived growth factor receptor beta | 1GQ5 | active |
| Ephrin type-A receptor 2 | 1MQB | active |
| Thyroid hormone receptor beta-1 | 1NQ2 | inactive |
| Inhibitor of nuclear factor kappa B kinase beta subunit | 4KIK | inactive |
| Beta amyloid A4 protein | 1TAW | inactive |
| Dipeptidyl peptidase IV | 2GBC | inactive |

**Table 3:** Novelty of anchor and generated ligands for DASATINIB can we include SMINA and SSNET scores in the same table

| ID | Smiles | BDB | ZINC |
|---|---|---|---|
| 0 | Cc1nc(Nc2ncc(C(=O)Nc3c(C)cccc3Cl)s2)cc(N2CCN(C)CC2)n1 | 0.84 | 0.85 |
| 1 | CCCN1CCN(c2cc(Nc3ncc(C(=O)Nc4c(C)cccc4Cl)s3)nc(C)n2)CC1 | 0.88 | 0.88 |
| 2 | Cc1nc(Nc2ncc(C(=O)Nc3c(C)cccc3Cl)s2)cc(N2CCN(CCO)CC2)n1[a] | 1.00 | 1.00 |
| 3 | Cc1nc(Nc2nnc(C(=O)Nc3c(C)cccc3Cl)s2)cc(N2CCN(CCO)CC2)n1 | 0.77 | 0.77 |
| 4 | Cc1nc(Nc2ncc(C(=O)Nc3c(C)cccc3Cl)s2)cc(N2CCN(CCO)OC2)n1 | 0.80 | 0.80 |
| 5 | Cc1nc(Nc2ncc(C(=O)Nc3c(C)cccc3Cl)s2)cc(N2CCCOCCCNCC2)n1 | 0.80 | 0.87 |
| 6 | Cc1nc(Nc2ncc(C(=O)Nc3c(C)cccc3Cl)s2)cc(C2CCN(CCO)CC2)n1 | 0.77 | 0.76 |
| 7 | CCCN1CCN(c2cc(NC3=NC(C(=O)Nc4c(C)cccc4Cl)=[SH]3)nc(C)n2)CC1 | 0.59 | 0.63 |
| 8 | Cc1nc(Nc2ncc(C(=O)Nc3c(C)cccc3Cl)s2)cc(N2CCC(CCO)CC2)n1 | 0.88 | 0.88 |
| 9 | CCCN1CCN(C2=CCCCc3cccc(C)c3NC(=O)C(NC3=NC=CSS3)=NC(F)=N2)CC1 | 0.32 | 0.37 |
| 10 | Cc1nc(Nc2ncc(C(=O)Nc3c(C)cccc3Cl)s2)cc(N2CCN(CCS)CC2)n1 | 0.87 | 0.87 |
| 11 | Cc1nc(Nc2ncc(C(=O)Nc3c(C)cccc3Cl)s2)cc(N2CCCCCCCNCC2)n1 | 0.76 | 0.92 |

[a] anchor ligand

**Table 4:** smina and SSnet scores for the 12 ligands investigated for bioactivity

| Smina | | | | | | | | | |
|---|---|---|---|---|---|---|---|---|---|
| ID[a] | 2FO0 | 1PKG | 1AVZ | 1GQ5 | 1MQB | 1NQ2 | 4KIK | 1TAW | 2GBC |
| 0 | -9.2 | -7.4 | -8.6 | -7.3 | -6.9 | -7.2 | -8.2 | -5.9 | -7.6 |
| 1 | -7.3 | -7.6 | -8.6 | -6.6 | -6.8 | -6.5 | -6.1 | -5.2 | -6.8 |
| 2 | -8.6 | -7.4 | -9 | -6.9 | -7.5 | -6.3 | -7.4 | -5.5 | -7.7 |
| 3 | -8.6 | -7.5 | -8.7 | -6.7 | -7.1 | -6.2 | -7.7 | -5.5 | -7.5 |
| 4 | -9.3 | -7.2 | -8.6 | -6 | -7.3 | -6 | -8.2 | -5.6 | -7.3 |
| 5 | -9.1 | -7.3 | -9.3 | -6 | -7.6 | -6.5 | -7.9 | -5.6 | -7.7 |
| 6 | -8.6 | -7.7 | -8.8 | -6.8 | -6.7 | -6.2 | -7.8 | -6 | -7.7 |
| 7 | -8.2 | -6.9 | -7.8 | -6.8 | -6.8 | -6 | -5.7 | -5.1 | -7.2 |
| 8 | -8.8 | -7.6 | -9.4 | -6.9 | -7.4 | -6.5 | -6 | -5.6 | -7.8 |
| 9 | -8.5 | -7.9 | -8.3 | -6.2 | -6.4 | -7.2 | -6.5 | -5.7 | -7.3 |
| 10 | -8.3 | -6.6 | -8.5 | -6.4 | -6.7 | -5.9 | -8.1 | -5.4 | -6.9 |
| 11 | -7.7 | -7.5 | -8.8 | -6.5 | -7 | -6.6 | -6.7 | -5.6 | -8 |
| SSnet | | | | | | | | | |
| 0 | 0.957 | 0.916 | 0.000 | 0.933 | 0.095 | 0.006 | 0.000 | 0.000 | 0.000 |
| 1 | 0.943 | 0.837 | 0.000 | 0.868 | 0.042 | 0.004 | 0.000 | 0.000 | 0.000 |
| 2 | 0.886 | 0.936 | 0.000 | 0.949 | 0.114 | 0.002 | 0.000 | 0.000 | 0.000 |
| 3 | 0.975 | 0.953 | 0.000 | 0.963 | 0.124 | 0.009 | 0.000 | 0.000 | 0.000 |
| 4 | 0.949 | 0.997 | 0.000 | 0.998 | 0.572 | 0.004 | 0.000 | 0.000 | 0.000 |
| 5 | 0.990 | 0.984 | 0.000 | 0.988 | 0.299 | 0.021 | 0.000 | 0.000 | 0.000 |
| 6 | 0.982 | 0.991 | 0.000 | 0.993 | 0.355 | 0.013 | 0.000 | 0.000 | 0.000 |
| 7 | 0.998 | 0.744 | 0.000 | 0.788 | 0.028 | 0.086 | 0.000 | 0.000 | 0.000 |
| 8 | 0.994 | 0.973 | 0.000 | 0.979 | 0.218 | 0.029 | 0.000 | 0.000 | 0.000 |
| 9 | 0.968 | 0.869 | 0.000 | 0.891 | 0.052 | 0.006 | 0.000 | 0.000 | 0.000 |
| 10 | 0.816 | 0.962 | 0.000 | 0.970 | 0.171 | 0.001 | 0.000 | 0.000 | 0.000 |
| 11 | 0.905 | 0.924 | 0.000 | 0.940 | 0.093 | 0.002 | 0.000 | 0.000 | 0.000 |

[a] ID taken from Table 3. Each column shows results for docking on the mentioned protein given as PDB ID.

#### 0.2.2 Novel ligands generated around anchor ligand Gefitinib

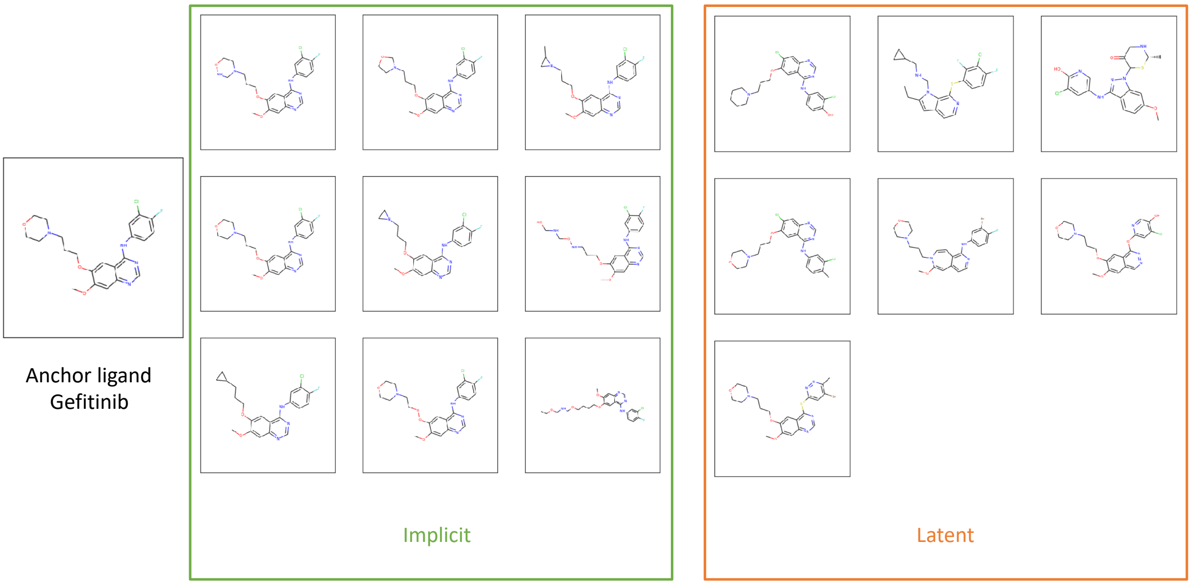

**Figure 1:** Novel ligands generated around anchor ligand Gefitinib

**Table 5:** smina and SSnet scores for the ligands investigated for bioactivity with the Epidermal Growth Factor Receptor with PDB ID 1MOX

| SMILES | Category | SSnet | smina |
|---|---|---|---|
| COc1cc2ncnc(Nc3ccc(F)c(Cl)c3)c2cc1OCCCN4CCOCC4 | Anchor | 0.912 | -7.8 |
| COc1cc2ncnc(Nc3ccc(F)c(Cl)c3)c2cc1OCCCN1CCONC1 | Implicit | 0.969 | -8.1 |
| COc1cc2ncnc(Nc3ccc(F)c(Cl)c3)c2cc1OCCCN1CCOC1 | Implicit | 0.939 | -7.7 |
| COc1cc2ncnc(Nc3ccc(F)c(Cl)c3)c2cc1OCCCN1CC1C | Implicit | 0.909 | -7.3 |
| COc1cc2ncnc(Nc3ccc(F)c(Cl)c3)c2cc1OCCCN1CCOCC1 | Implicit | 0.912 | -8.1 |
| COc1cc2ncnc(Nc3ccc(F)c(Cl)c3)c2cc1OCCCN1CC1 | Implicit | 0.888 | -7.5 |
| COc1cc2ncnc(Nc3ccc(F)c(Cl)c3)c2cc1OCCCNOCNCO | Implicit | 0.994 | -6.8 |
| COc1cc2ncnc(Nc3ccc(F)c(Cl)c3)c2cc1OCCCC1CC1 | Implicit | 0.911 | -7.4 |
| COc1cc2ncnc(Nc3ccc(F)c(Cl)c3)c2cc1OOCCN1CCOCC1 | Implicit | 0.948 | -7.5 |
| CCOCNCOCCCCOc1cc2c(Nc3ccc(F)c(Cl)c3)ncnc2cc1OC | Implicit | 0.973 | -6.7 |
| COc1cc2ncnc(Nc3ccc(F)c(Cl)c3)c2cc1OCCCN1CCOOC1 | Implicit | 0.890 | -7.9 |
| Oc1ccc(Nc2ncnc3cc(Cl)c(OCCCN4CCCCC4)cc23)cc1Cl | Latent | 0.788 | -8.8 |
| CCc1cc2ccnc(Sc3ccc(F)c(Cl)c3F)c2n1CNCC1CC1 | Latent | 0.954 | -7.5 |
| COc1ccc2c(Nc3cnc(O)c(Cl)c3)nn(C3S[C@@H](C)NCC3=O)c2c1 | Latent | 0.810 | -7.4 |
| Cc1ccc(Nc2ncnc3cc(Cl)c(OCCCN4CCOCC4)cc23)cc1Cl | Latent | 0.814 | -8.4 |
| COC1=Cc2ccnc(Nc3ccc(F)c(Br)c3)c2C=CN1CCCN1CCOCC1 | Latent | 0.559 | -7.3 |
| COc1cc2cnnc(Oc3cc(Cl)c(O)cn3)c2cc1OCCCN1CCOCC1 | Latent | 0.140 | -7.8 |
| COc1cc2ncnc(Sc3cc(Br)c(C)nn3)c2cc1OCCCN1CCOCC1 | Latent | 0.631 | -7.2 |

#### 0.2.3 Novel ligands generated around anchor ligand Erlotinib

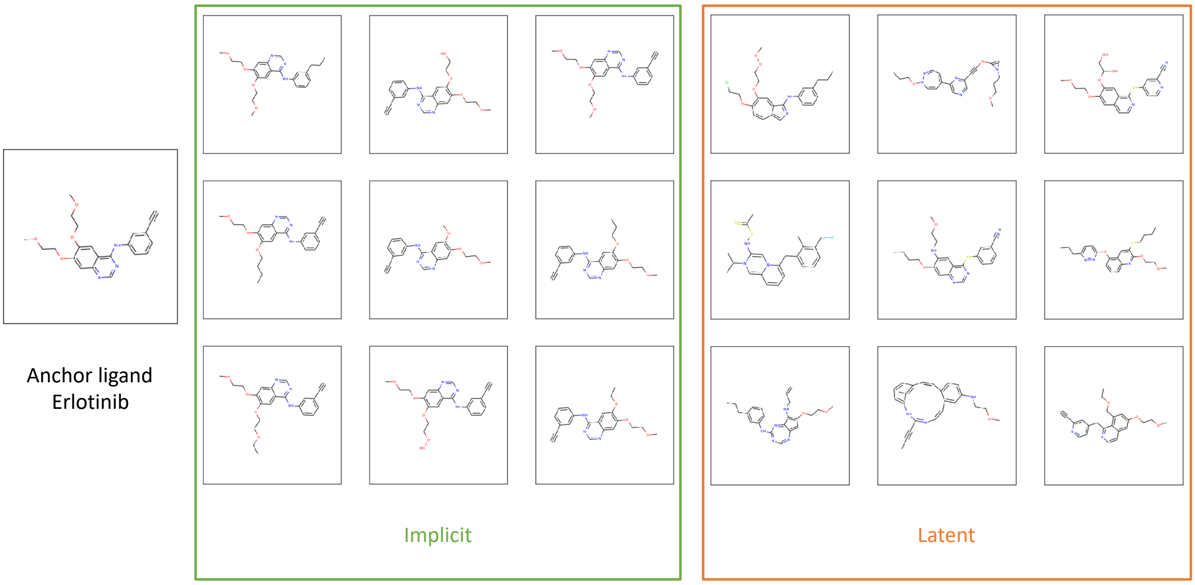

**Figure 2:** Novel ligands generated around anchor ligand Erlotinib

**Table 6:** smina and SSnet scores for the ligands investigated for bioactivity with the Epidermal Growth Factor Receptor with PDB ID 1MOX

| SMILES | Category | SSnet | smina |
|---|---|---|---|
| COCCOc1cc2ncnc(Nc3cccc(c3)C#C)c2cc1OCCOC | Anchor | 0.818 | -6.9 |
| CCCc1cccc(Nc2ncnc3cc(OCCOC)c(OCCOC)cc23)c1 | Implicit | 0.791 | -7.3 |
| C#Cc1cccc(Nc2ncnc3cc(OCCOC)c(OCCO)cc23)c1 | Implicit | 0.801 | -7.4 |
| C#Cc1cccc(Nc2ncnc3cc(OCCOC)c(OCCOC)cc23)c1 | Implicit | 0.818 | -6.9 |
| C#Cc1cccc(Nc2ncnc3cc(OCCOC)c(OCCCC)cc23)c1 | Implicit | 0.880 | -7.1 |
| C#Cc1cccc(Nc2ncnc3cc(OCCOC)c(OC)cc23)c1 | Implicit | 0.876 | -6.9 |
| C#Cc1cccc(Nc2ncnc3cc(OCCOC)c(OCCC)cc23)c1 | Implicit | 0.834 | -6.9 |
| C#Cc1cccc(Nc2ncnc3cc(OCCOC)c(OCCOCC)cc23)c1 | Implicit | 0.912 | -6.7 |
| C#Cc1cccc(Nc2ncnc3cc(OCCOC)c(OCCOO)cc23)c1 | Implicit | 0.930 | -7.2 |
| C#Cc1cccc(Nc2ncnc3cc(OCCOC)c(OCC)cc23)c1 | Implicit | 0.878 | -7 |
| CCCc1cccc(Nc2ncc3ccc(OCCCl)c(OCCOOC)cc2-3)c1 | Latent | 0.868 | -7 |
| CCCON1C=CC(c2cncc(C#COC3=CN3CCCOC)n2)=CC=N1 | Latent | 0.627 | -6.6 |
| COCCOc1cc2ccnc(Sc3ccnc(C#N)c3)c2cc1OC(O)CO | Latent | 0.348 | -7.3 |
| CC(=S)SNC1=CN2C(=CN1C(C)C)C=CC=C2Cc1cccc(CF)c1C | Latent | 0.344 | -7.7 |
| CCCCOc1cc2ncnc(Sc3cccc(C#N)c3)c2cc1NCCOC | Latent | 0.894 | -6.7 |
| CCCCSc1cc2c(Oc3ccc(CCC)nn3)cccc2nc1OCCOC | Latent | 0.967 | -6.8 |
| C=CCNc1c(OCCOC)cc2ncnc(Nc3cccc(CCC)c3)nc1-2 | Latent | 0.671 | -6.8 |
| CC#CC1=NC=Cc2cc(NCCOC)ccc2C#Cc2cccc(c2C)N1 | Latent | 0.069 | -8.1 |
| C#Cc1cc(Cc2nccc3cc(OCCOC)cc(COCC)c23)ccn1 | Latent | 0.988 | -6.8 |
| COCCCOn1c(OCF)cc2c(NCc3ncc(C#N)o3)nccc21 | Latent | 0.986 | -6.2 |
| CCCNN1N=C2C(=CC=NC3=C2Cc2coc(c2Cl)CS3)C=C1CCCOC | Latent | 0.238 | -7.3 |
| CCCCOc1cc2c(Cc3cccc(C#N)c3)nccc2cc1OCCOC | Latent | 0.671 | -6.9 |
| CC(C)OOc1cc2ccnc(Nc3ccc(C#N)cc3)c2cc1OCCO | Latent | 0.755 | -7.9 |
| CC#Cc1cncc(Sc2ncnc3cc(SCCOC)c(CCOCC)cc23)n1 | Latent | 0.974 | -6.6 |
| C#Cc1cc(Cc2ncnc3cc(OCCCl)c(OCCOC)cc23)ccn1 | Latent | 0.925 | -6.6 |
| CCCCSc1nc2ccnc(Nc3cccc(C#CF)c3)n2c1OCCOC | Latent | 0.900 | -6.7 |
| C#CCc1ccc(Nc2nccc3cc(CCCCC)n(C=COOC)c23)nn1 | Latent | 0.774 | -7.1 |
| C#Cc1cccc(Cc2ccnc3cc(SCCCC)c(CNCCOCC)cc23)c1 | Latent | 0.958 | -7.1 |
| C=CCOC1=Cc2ccsc2CSc2occ(c2C#N)-c2ccnn2CC1=C | Latent | 0.748 | -6.7 |

#### 0.2.4 Novel ligands generated around anchor ligand Dasatinib

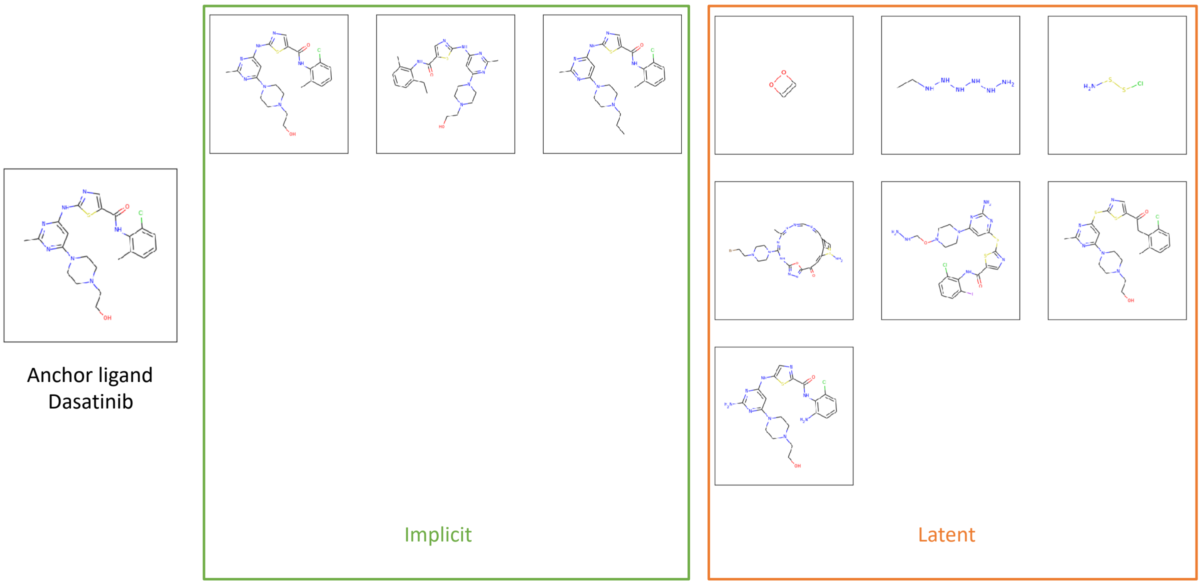

**Figure 3:** Novel ligands generated around anchor ligand Dasatinib

**Table 7:** smina and SSnet scores for the ligands investigated for bioactivity with the P56LCK SH2 DOMAIN INHIBITOR with PDB ID 1BHF

| SMILES | Category | SSnet | smina |
|---|---|---|---|
| Cc1nc(Nc2ncc(s2)C(=O)Nc3c(C)cccc3Cl)cc(n1)N4CCN(CCO)CC4 | Anchor | 0.945 | -6.6 |
| Cc1nc(Nc2ncc(C(=O)Nc3c(C)cccc3Cl)s2)cc(N2CCN(CCO)CC2)n1 | Implicit | 0.945 | -6.6 |
| CCc1cccc(C)c1NC(=O)c1cnc(Nc2cc(N3CCN(CCO)CC3)nc(C)n2)s1 | Implicit | 0.904 | -6.7 |
| CCCN1CCN(c2cc(Nc3ncc(C(=O)Nc4c(C)cccc4Cl)s3)nc(C)n2)CC1 | Implicit | 0.857 | -6.2 |
| NSSCl | Latent | 0.039 | -2.7 |
| Cc1nncncc2c(C)c(cc(=O)c3nnc([nH]c(N4CCN(CCBr)CC4)n1)o3)S(N)=C2 | Latent | 0.652 | -5.7 |
| NNCON1CCN(c2cc(Sc3ncc(C(=O)Nc4c(Cl)cccc4I)s3)nc(N)n2)CC1 | Latent | 0.590 | -6.6 |
| Cc1nc(Sc2ncc(C(=O)Cc3c(C)cccc3Cl)s2)cc(N2CCN(CCO)CC2)n1 | Latent | 0.750 | -6.8 |
| Nc1nc(Nc2cnc(C(=O)Nc3c(N)cccc3Cl)s2)cc(N2CCN(CCO)CC2)n1 | Latent | 0.872 | -7 |
| [C+2] | Latent | 0.026 | -0.9 |
| NCNNNNNN | Latent | 0.018 | -5.1 |

#### 0.2.5 Novel ligands generated around anchor ligand Pyrimethamine

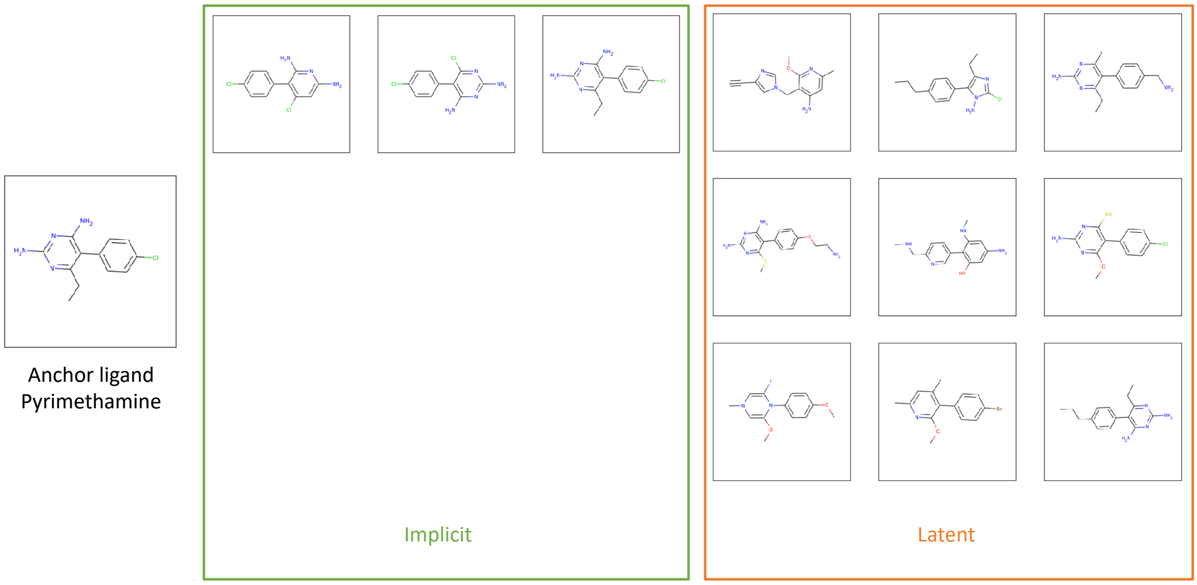

**Figure 4:** Novel ligands generated around anchor ligand Pyrimethamine

**Table 8:** smina and SSnet scores for the ligands investigated for bioactivity with the HUMAN DIHYDROFOLATE REDUCTASE with PDB ID 1BHF

| SMILES | Category | SSnet | smina |
|---|---|---|---|
| CCc1nc(N)nc(N)c1c2ccc(Cl)cc2 | Anchor | 0.095 | -5.5 |
| Nc1cc(Cl)c(-c2ccc(Cl)cc2)c(N)n1 | Implicit | 0.061 | -5.3 |
| Nc1nc(N)c(-c2ccc(Cl)cc2)c(Cl)n1 | Implicit | 0.015 | -5.2 |
| CCc1nc(N)nc(N)c1-c1ccc(Cl)cc1 | Implicit | 0.072 | -5.2 |
| C#Cc1cn(Cc2c(N)cc(C)nc2OC)cn1 | Latent | 0.752 | -4.6 |
| CCCc1ccc(-c2c(CC)nc(Cl)n2N)cc1 | Latent | 0.406 | -5.1 |
| CCc1nc(N)nc(C)c1-c1ccc(CN)cc1 | Latent | 0.078 | -5.5 |
| CSc1nc(N)nc(N)c1-c1ccc(OCCN)cc1 | Latent | 0.087 | -5.5 |
| CNCc1ccc(-c2c(O)cc(N)cc2NC)cn1 | Latent | 0.539 | -5.4 |
| COc1nc(N)nc(S)c1-c1ccc(Cl)cc1 | Latent | 0.064 | -5.1 |
| COC1=CN(C)C=C(I)N1c1ccc(OC)cc1 | Latent | 0.040 | -4.4 |
| COc1nc(C)cc(C)c1-c1ccc(Br)cc1 | Latent | 0.099 | -4.9 |
| CCCc1ccc(-c2c(N)nc(N)nc2CC)cc1 | Latent | 0.085 | -5.4 |
| CCc1nc(C)cn1Sc1cncccn(I)n1C | Latent | 0.086 | -4.8 |
| CNc1nc(N)cc(N)c1-c1ccc(Br)cn1 | Latent | 0.588 | -5.1 |
| CCc1cc(C)cc(C)c1C1=CC=CN(Cl)C=C1 | Latent | 0.225 | -5.7 |
| CCc1nc(I)nc(C)c1-c1ccc(CCF)cc1 | Latent | 0.113 | -4.9 |
| CCc1nc(N)cc(I)c1-c1ccc(Cl)cc1 | Latent | 0.149 | -5.5 |
| C#Cc1ncc(-c2cc(N)c(Cl)nc2CC)o1 | Latent | 0.454 | -5.1 |
| CCc1ncn(Cc2c(C)cc(C)cc2CC)n1 | Latent | 0.056 | -5.3 |
| CCc1cc(I)cc(N)c1-c1ccc(Cl)cc1 | Latent | 0.183 | -4.8 |
| CCCc1ccc(-c2c(CC)cc(C)nc2CC)o1 | Latent | 0.030 | -5.1 |
| CSc1nc(N)cc(N)c1-c1ccc(Br)cc1O | Latent | 0.048 | -5.3 |
| CCc1nn(C)c(CO)c1-n1cnc(C#N)c1 | Latent | 0.267 | -3.9 |
| CCc1c(Cn2ccc(C)n2)c(O)cn1Br | Latent | 0.119 | -4.6 |
| CCc1ccc(-c2c(O)cc(C)cc2OC)cc1 | Latent | 0.109 | -4.9 |

2 tables  one with the known assays for this drug and the novel ligands generated around this drug. If we have SMINA score and SSNET scores for binding affinity can we list them for the novel drugs against the targets in the same table?

#### 0.2.6 Novel ligands generated around anchor ligand Crizotinib

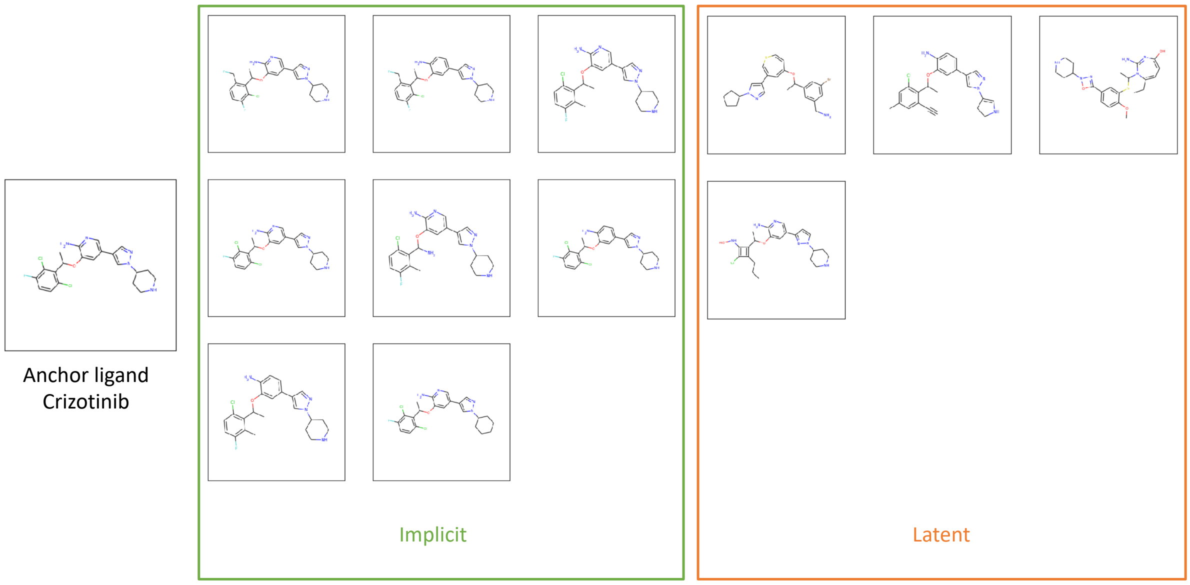

**Figure 5:** Novel ligands generated around anchor ligand Crizotinib

**Table 9:** smina and SSnet scores for the ligands investigated for bioactivity with the PTB domain of SNT-2 and 19-residue peptide (aa 1571-1589) of HALK with PDB ID 2KUP

| SMILES | Category | SSnet | smina |
|---|---|---|---|
| CC(Oc1cc(cnc1N)c2cnn(c2)C3CCNCC3)c4c(Cl)ccc(F)c4Cl | Anchor | 0.950 | -7.1 |
| CC(Oc1cc(-c2cnn(C3CCNCC3)c2)cnc1N)c1c(CF)ccc(F)c1Cl | Implicit | 0.991 | -7 |
| CC(Oc1cc(-c2cnn(C3CCNCC3)c2)ccc1N)c1c(CF)ccc(F)c1Cl | Implicit | 0.955 | -7.1 |
| Cc1c(F)ccc(Cl)c1C(C)Oc1cc(-c2cnn(C3CCNCC3)c2)cnc1N | Implicit | 0.931 | -7.1 |
| CC(Oc1cc(-c2cnn(C3CCNCC3)c2)cnc1N)c1c(Cl)ccc(F)c1Cl | Implicit | 0.950 | -6.9 |
| Cc1c(F)ccc(Cl)c1C(N)Oc1cc(-c2cnn(C3CCNCC3)c2)cnc1N | Implicit | 0.951 | -7.1 |
| CC(Oc1cc(-c2cnn(C3CCNCC3)c2)ccc1N)c1c(Cl)ccc(F)c1Cl | Implicit | 0.839 | -7.3 |
| Cc1c(F)ccc(Cl)c1C(C)Oc1cc(-c2cnn(C3CCNCC3)c2)ccc1N | Implicit | 0.715 | -7.2 |
| CC(Oc1cc(-c2cnn(C3CCCCC3)c2)cnc1N)c1c(Cl)ccc(F)c1Cl | Implicit | 0.855 | -7.2 |
| [N+2] | Latent | 0.018 | -1.1 |
| [N+]NNNNNNNNNNNNNNNNNNNNNNNNNNNNNNNNNNNNNNNNNNNNNNNNNNNNNNNNNNNNNNNNNNNN | Latent | 0.002 | -4.9 |
| CC(OC1=CC(c2cnn(C3CCCC3)c2)=CSC=C1)c1cc(Br)cc(CN)c1 | Latent | 0.556 | -7.3 |
| C#Cc1cc(C)cc(Cl)c1C(C)Oc1cc(-c2cnn(C3=CNCC3)c2)ccc1N | Latent | 0.785 | -6.6 |
| CCC1=CC=C(O)N=C(N)N1C(C)Sc1cc(-c2nn(C3CCNCC3)o2)ccc1OC | Latent | 0.024 | -6.8 |
| CCCC1=C(C(C)Oc2cc(-c3ccn(C4CCNCC4)n3)cnc2N)C(NO)=C1Cl | Latent | 0.965 | -6.4 |

2 tables one with the known assays for this drug and the novel ligands generated around this drug. If we have SMINA score and SSNET scores for binding affinity can we list them for the novel drugs against the targets in the same table?

#### 0.2.7 Novel ligands generated around anchor ligand Verapamil

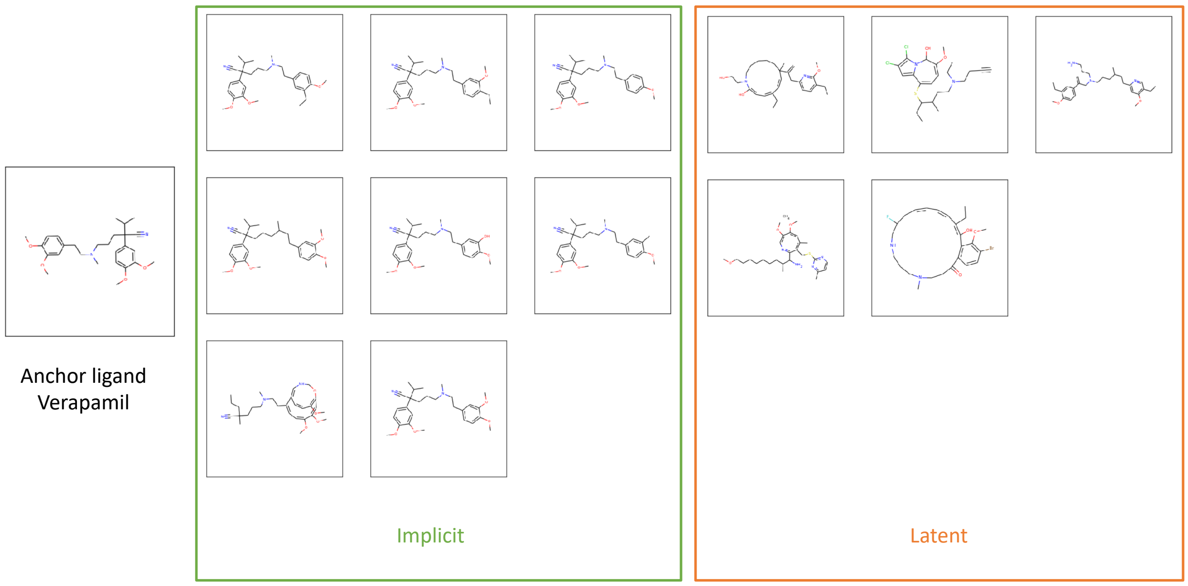

**Figure 6:** Novel ligands generated around anchor ligand Verapamil

**Table 10:** smina and SSnet scores for the ligands investigated for bioactivity with the Calmodulin/IQ domain complex with PDB ID 2F3Y

| SMILES | Category | SSnet | smina |
|---|---|---|---|
| COc1ccc(CCN(C)CCCC(C#N)(C(C)C)c2ccc(OC)c(OC)c2)cc1OC | Anchor | 0.000 | -4 |
| CCc1cc(CCN(C)CCCC(C#N)(c2ccc(OC)c(OC)c2)C(C)C)ccc1OC | Implicit | 0.000 | -3.5 |
| CCc1ccc(CCN(C)CCCC(C#N)(c2ccc(OC)c(OC)c2)C(C)C)cc1OC | Implicit | 0.000 | -3.8 |
| COc1ccc(CCN(C)CCCC(C#N)(c2ccc(OC)c(OC)c2)C(C)C)cc1 | Implicit | 0.000 | -3.5 |
| COc1ccc(CCC(C)CCCC(C#N)(c2ccc(OC)c(OC)c2)C(C)C)cc1OC | Implicit | 0.000 | -4.2 |
| COc1ccc(CCN(C)CCCC(C#N)(c2ccc(OC)c(OC)c2)C(C)C)cc1O | Implicit | 0.000 | -4.4 |
| COc1ccc(CCN(C)CCCC(C#N)(c2ccc(OC)c(OC)c2)C(C)C)cc1C | Implicit | 0.000 | -3 |
| CCCC(C)(C#N)CCCN(C)CCC1=CC=C(OC)C(OC)=CC2=C(OC)C=CC1=CNCO2 | Implicit | 0.000 | -4.5 |
| COc1ccc(CCN(C)CCCC(C#N)(c2ccc(OC)c(OC)c2)C(C)C)cc1OC | Implicit | 0.000 | -3.5 |
| C=C(Cc1ccc(CC)c(OC)n1)C1(C)C=CC(CC)=CC=C(O)N(CCO)CCCCC1 | Latent | 0.000 | -4.7 |
| C#CCCN(CC)CCC(C)C(CC)SC1=CC=C(OC)C(O)n2c1cc(Cl)c2Cl | Latent | 0.000 | -4.2 |
| C=C(CN(CCCN)CCCC(C)CCc1cc(OC)c(CC)cn1)c1ccc(OC)c(CC)c1 | Latent | 0.000 | -4.3 |
| C.COCCCCCCCC(C)C(N)C1=NC=C(OC)C(OC)=CC(C)=C1CSc1nccc(C)n1 | Latent | 0.000 | None |
| CCC1=C(O)c2c(ccc(Br)c2OC)C(=O)CCN(C)CCCNCC(F)CC=CC=C1 | Latent | 0.000 | -5.4 |

2 tables one with the known assays for this drug and the novel ligands generated around this drug. If we have SMINA score and SSNET scores for binding affinity can we list them for the novel drugs against the targets in the same table?

### 0.3 Neural Network Architecture

The detailed architecture of the neural network is illustrated in figure 7. The number of parameters at each layer is illustrated in the figure 8.

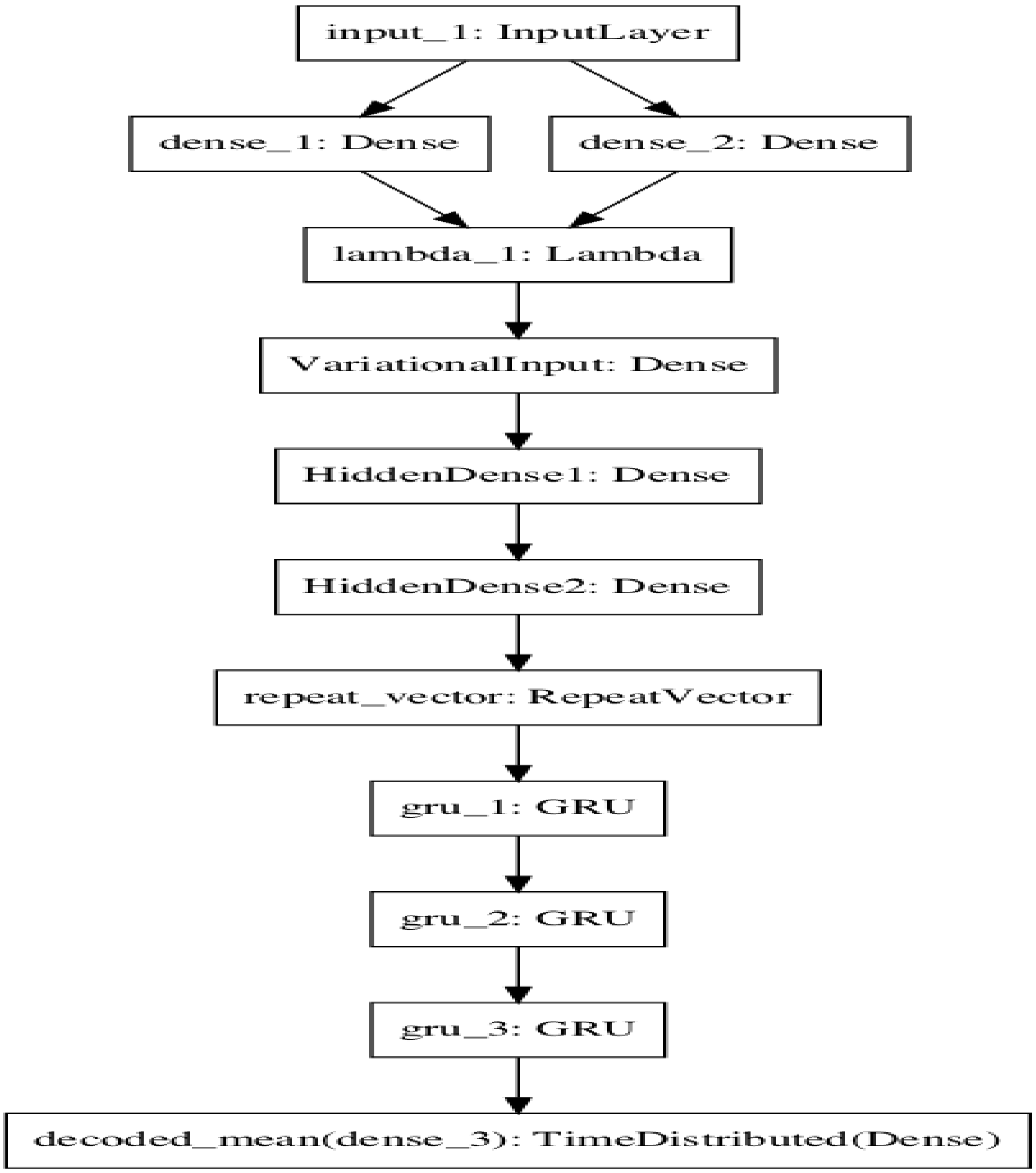

**Figure 7:** Neural network architecture of the deep learning model.

**Trainable parameters of the Deep learning architecture**

| Layer (type) | Output Shape | Param # | Connected to |
|---|---|---|---|
| input_1 (InputLayer) | (None, 50) | 0 | |
| dense_1 (Dense) | (None, 50) | 2550 | input_1[0][0] |
| dense_2 (Dense) | (None, 50) | 2550 | input_1[0][0] |
| lambda_1 (Lambda) | (None, 50) | 0 | dense_1[0][0]<br>dense_2[0][0] |
| VariationalInput (Dense) | (None, 50) | 2550 | lambda_1[0][0] |
| HiddenDense1 (Dense) | (None, 80) | 4080 | VariationalInput[0][0] |
| HiddenDense2 (Dense) | (None, 120) | 9720 | HiddenDense1[0][0] |
| repeat_vector (RepeatVector) | (None, 120, 120) | 0 | HiddenDense2[0][0] |
| gru_1 (GRU) | (None, 120, 501) | 934866 | repeat_vector[0][0] |
| gru_2 (GRU) | (None, 120, 501) | 1507509 | gru_1[0][0] |
| gru_3 (GRU) | (None, 120, 501) | 1507509 | gru_2[0][0] |
| decoded_mean (TimeDistributed) | (None, 120, 58) | 29116 | gru_3[0][0] |

Total params: 4,000,450
Trainable params: 4,000,450
Non-trainable params: 0

**Figure 8:** Layers and parameters trained in the deep learning model.

### 0.4 List of All Novel Ligands generated from 5k anchor ligands.

Novel Smiles generated from 5k Anchor Ligands

```
1 COc1ccccc1N2CCC(CNCC3COc4ccc(Cl)cc4C3)CC2
2 CCn1nnc(n1)[C@H]2O[C@H]([C@H](O)[C@@H]2O)n3cnc4c(NC)ccnc34
3 COc1cc2c(Oc3ccc(NC(=O)C4=C(Cl)c5ccccc5N(C4=O)c6ccc(F)cc6)cc3F)ccnc2cc1OCCCN7CCCCCCC[illegible]
4 CCn1nnc(n1)[C@H]2O[C@H]([C@H](O)[C@@H]2O)n3cnc4c(NC5CC5CCCCCl)nc34
5 CCNN(CCC)C1CCc2cc2(cccccC1)
6 Fc1ccc(cc1)C(=O)C2CCN(CCN3C(=O)Nc4ccccc4C3CO)CC2
7 Clc1ccc(CC)c2c3N=CNC(=O)c3sc12
8 Clc1ccc(CCCNCCCOc2ccccc2)cc1
9 COc1cccc(c1)C(C)NC(=O)c2ccc(cc2)c3ccccc3C
10 CCCCCN(C)CCC(OPPP(OO)(OPOOPPOO))OO
11 CN(C)CCCOC(=O)C(C)(CCCCCCCC)c2ccccc2C
12 COc1cccc(c1)C(C)NC(OO)c2ccc(cc2)c3ccccc3C
13 Cc1ccc2c(cccc2n1)N3CCN(CCc4cccc5c4OCc6c(ccn56)C(=O)NC7CCC7)CC3
14 CC1=C2NCc3cc(C2ccc3NCCCC1=O)
15 CCCS(=O)(=O)Nc1ccc(cc1)c2ccc3[nH]nc(C)c3c2
16 O=C1CCc2ccc(OCCCCN3CCN(CC3)c4cccc5CCCOc45)nc2N1C
17 Clc1ccc(CNC2=C(Nc3ccncc3)C(NO)C2=O)cc1
18 CCCc1nc(C)c2C(=O)SC(=Nn12)c3ccccc3OCC
19 CCCCCN(C)CCC(O)(P(=O)(O)O)P(=O)(O)
20 Cc1noc(n1)c2Cc2CCCCC
21 NS(=O)(=O)c1ccc(Cc1=O)CCC
22 Nc1nc(N)c2nc(CNc3ccc(cc3)C(=O)NC(CCCCC(=O)c4ccccc4C(=O)O)C(=O)O)cnc2n1
23 CN(C)CCCC(c1ccccc1)c2ccccc2CC
24 NS(=O)(=O)c1nnc(s1)c2ccccc2C
25 Fc1c(Cl)cccc1N2CCN(CCCCOc3ccc4CCC(OO)Nc4c3)CC2
26 CCCN(CCC)C1CCC(OCC1)C
27 CCCCCCNS(=O)(=O)NCNC(=O)OC
28 O=C1NCc2c1cccc2c3ccc(Nc4oc5ccccc5c4)cc3
29 C(Nc1ccccc1c2cc[nH]n2)C3CNCCN3
30 Clc1cccc(c1)N2CCN(CC3CCCCC3)C2COC
```

31 Cc1ccc2c(cccc2n1)N3CCN(CCCCc4ccc5CCC(=O)Nc5c4)CC3

32 CCNC(=O)[C@H]1O[C@H]([C@H](O)[C@@H]1O)n2cnc3c(NCN(N)N)ncnc23

33 Cc1ccc2c(cccc2n1)N3CCN(CCc4ccc5OCC(=O)Nc5c4)cC3

34 CN1CCN(CC1)c2cc(NCc3ccc(F)cc3)nc(N)n2 2

35 CC[C@@]1(O)C(=O)OCC2=C1C=C3N(Cc4cc5c(N)cccc5nc34)C2CO

36 Oc1ccc2C[C@@H]3[C@@H]4CCCC[C@]4(CCC3CC5CCC5)c2c1

37 C[C@H](Oc1ccccc1CC=C)C2=NC=N2

38 O=C(N1CCOCC1)C2CCn3cc(C4=C(C(=O)NC4OO)c5cnc6ccccn56)c7cccc(C2)c37

39 CNCC1(CCCCC1)c2c3ccccccc3c2

40 Fc1ccc2c(noc2c1)C3CCN(CCCCS(=S)(=O)c4cc5ccs(Cl)cc5s4)CC3

41 COc1ccc(cc1)S(=O)(=O)NC(C)C(=O)NOO

42 Cc1ccc(cc1)C2=CC(CO)c3ccccc3O2

43 CCCN1C(=O)N(CCC)c2cc([nH]c2C1=O)c3ccc(OCC(=O)Nc4ccc(BB)cc4)cc3

44 NS(=O)(=O)c1cc2C(O)CCS(=O)(=O)c21

45 Cc1ccc2c(cccc2n1)N3CCN(CCc4ccc5NC(=O)COc5n4)CC3

46 Clc1ccc(cc1)C(=O)CCCC2CCc3ccccc3C2N

47 Clc1cc2NC(=O)Cc2cc1CCN3CCC(CC3)c4nnc5cccccc54

48 CC(C)n1nc(c2cc(O)cc(F)c2)c3c(N)nccc13

49 OC1=C(Oc2cc(O)cc(O)c2c1=C)c3ccc(O)c(O)c3

50 COc1ccccc1NNCCN(CC(O)CCNC(=O)c3cc4ccccc4s3)CCC

51 COc1cc(ccc1Nc2ncc3CN(C(=O)N(c4cccc(NC(=O)C=C)c4)c3n2)c5ccc(OCc6ccccc6)cc5)N7CCCCCC[illegible]

52 CCCN(CCC)[C@H]1CCn2ncnc2C1

53 CCN1CCCC1CNC(=O)c2cc(ccc2SC)S(OO)(=O)N

54 FC(C)(F)C(=O)N[C@@H]1CC[C@@H](CCN2CCC(CC2)c3coc4ccccc34)CC1

55 C[C@H](N)CC1CCc2ccc(CO)cc12

56 CC1=CC[C@@H]2[C@@H](C1)c3c(O)cc(cc3OC2(C)C)C(C)(C)c4cccCCCCcc4

57 CCCn1c(C)c(C(=O)c2cccc3cccc233)c3ccccc1

58 Cc1cc(nc2ccccc12)N3CCN(CCCCC4=NC5CC(CCCC5)C(=O)N4C)CC3

59 OC1(N2CCNCC2c3ccccc13)c4ccc(Cl)cc4

60 CN(C)CCNC1c2ccccc2Cc3ccccc13

61 CCOC(=O)c1ncn2c1CC(C)C(=O)c3cc(ccc23)C#C

62 C=S(=O)(NC1CCN(CCOc2cccc3CCCCc23)C1)c4cccs4

63 Cc1cc(C)c(CCC2C(=O)Nc3ccccc23)[nH]1

64 NS(=O)(=O)c1ccc2CCCc2ccc1

65 CNS(=O)(=O)c1ccccc1Nc2nc(Nc3cc(OC)cCOCOc(CC)c3)ncc2Cl

66 COCC(N1CCCC1)C(=O)c2ccc(BC)cc2

67 COc1cc2c(Oc3ccc(cc3)c4nc5ccccc5s4)ncnc2cc1OCCCN6CCC(C)CC6

68 NC(=O)c1cc2c(Oc3ccc(BO)cc3)cncc2s1

69 CCC1(Cc2ccccc2O1)C3CNCCN3

70 Fc1ccc2c(noc2c1)C3CCN(CCCNS(=O)(=O)c4cc5ccs(Cl)cc5s4)CC3

71 NS(=O)(=O)c1=cc(O)cc1 11

72 CN(C)CCCN1c2ccccc2Sc3ccc(CC)cc13

73 Cn1ccc2c(NCc3ccccc3)nc(NCCO)nc12

74 OC(NC(=O)c1ccc(cc1)c2ccncc2)c3ccccc3

75 COc1cc2ncnc(Nc3cccc(c3)C#C)c2cc1CCCCCCCCOOOOOO

76 CCN(CC)CC#CCOC(=O)C(CCCC1CCCCC1)c2ccccc2

77 Cc1cc(c(O)c(C)c1NC2=CCCC2)C(C)(C)

78 Ic1c(NC2=NCCC2)ccc3nncnc13

79 Ic1n(NC2=NC)N2cccc3nccnc13

80 Cc1ccc(cc1)C2=CC(C)(C)Oc3cccccc(ccccc23)c5ccc(cc5)C(=O)

81 Cl.N#Cc1ccc(cc1)C2CCCc3ccnn23

82 Clc1ccc(cc1Cl)S(=O)(=O)NCCCCN2CCN(CC2)c3csc4ccccc34

83 C(Nc1ccccc1c2nccs2)C3=NNCN3

84 CSc1ccccc1N2CCN(CCCCCC(=O)NC3CCCc4ccccc34)C2

85 CS(=O)(=O)N[C@@H]1CC[C@@H](CCN2CCC(CC2)3ccoc4ccccc34)CC1

86 COc1ccccc1C(C)NS(=O)(=O)

87 Cc1cccc(c1)n2cnCn2Cc3ccc(cc3)CN

88 OC(CNC1CCN(CC1)c2ncnc3scc(c4ccccc4)c23)COc5cccc(c5)CNNO

89 CCc1[nH]c2ccc(OC)cc2c1CCN(CC)

90 Fc1ccc(cn1)C(=O)NCCCCN2CCN3CC23c3cccc(Cl)c3CC

91 CN(C)CCNC(c1ccccc1)c2cccccCC2

92 CC(=O)Nc1cc(nc(n1)c2occc2)n3cccc3

93 Cn1cc(C=C2C(=O)Nc3cc2ccc3)c4ccccc14

94 CN1CCN(CC1)c2ccc(cc2)C(=O)N[C@@H]3CC[C@@H](CCN4CCC(CC4)c5coc6ccccc56)CC3C

95 Cc1ccc(cc1)C(=O)NCN2CCC(CC2)c3ccccc3C#N

96 Clc1ccc(CNC2=C(Nc3ccncc3)C(=O)C2CO)cc1Cl

97 CN1C(=O)C=Cc2c(CCN3CCN(CC3)c4cccc5cc(C)ccc45)c(C)ccc12

98 Oc1ccc(cc1)C2=C(c3ccc(OCCN4CCCC4)cc3)cc4cc(F)cc4OC2

99 CCCCCCNS(=O)(=O)NCNC(=O)CC

100 Nc1n[nH]c2cccc(c3NcN(NC(=O)Nc4cc(ccc4F)C(=O))ccc3)c12

101 NC(=N)NCCC[C@@H](NS(=O)(=O)Cc1ccccc1)C(=O)N2CCC[C@H]2C(=O)NCc3ccc(cc3)C(=N)

102 OC1(CC2=CC(C1)N2CCCC(=O)c3ccc(F)cc3)c4ccccc4

103 CC(Oc1ccccc1C2CCCC2)C3=NCCC3

104 Fc1ccc2c(noc2c1)C3CCN(CCCCCS(=O)(=O)c4ccc(OC(F)(F)F)cc4)CC3

105 Nc1n[nH]c2ccc(cc12)c3nnn(Cc4ccccc4)c3

106 Fc1ccc(cc1Cl)S(=O)(=O)NCCCN2CCN(CC2)c3csc4ccccc34

107 Cc1cc2CNC(=O)c2cc1OCCCN3CCN(CC3)c4cccc5cccc(C)c45

108 CN(C)CC1=C(O)C2=CC(=CC(=O)N2COC1)C

109 CN1CCN(CC1)[C@H]2Cc3ccccc3Sc4ccc(CCC(=O)C)cc24

110 Cc1ccc(NC(=O)Nc2ccc(NC(=O)c3csc4ccnc(N)c34)cc2)cc1

111 CC(C)Oc1ccccc1N2CNNNCc3cccc(c3)C(=O)C4CCCCC4CCC2

112 Cc1c(NC2=NCCN2)ccc3CCCCc13

113 O=C(NCC=1cccc1)N2CCC(=CC2)c3c[nH]c4ncccc34

114 CCOc1ccccc1CNCc2cccc(CCNC[C@H](O)c3ccc(O)c4NC(=O)Cc34)c2

115 Cc1c(sc2ccc(Cl)cc12)S(CO)(=O)NCCCCN3CCC(CC3)c4noc5cc(F)ccc45

116 Cc1ncoc1c2nnc(SCCCN3CC4CCC(C4C3)c5cccc(c5)C#N)n2C

117 Clc1cccc(N2CCN(CCCCOc3ccc4CCC(=C)Nc4c3)CC2)c1C

118 CN(C)S(=O)(=O)NC1CCc2ccc(C)cc2C1

119 CC(C)c1cc(nconnn1)c2ccc(F)c3ccccs23

120 OC[C@@]12C[C@@H]1[C@H]([C@H](O)[C@@H]2O)n3cnc4c(NCc5cccc(I)c5)cccc34

121 O=C1CCCN1CCCCCN2CCCC2

122 COc1cc2c(Oc3ccc(NC(=O)N\\N=C\\c4cccc(CC=C)c4O)cc3F)ccnc2cc1CCCCN5CCCCC5

123 CCOc1nc(cc(N)c1C#N)C(=C)NCc2ccccc2S(=O)(=O)NO

124 COC(=O)C1=C(C)NC(=O)N(C1c2ccc(F)c(F)c2)C(=O)NCCCN3CCCCCC3Cc4ccccc4OO

125 Nc1ncnc2scc(c3ccc4c(cccc4c3)C(=O)Nc5csCcc5)c12

126 CCCN(CCC)[C@H]1CCn2c(C1)ccc2C#C

127 CC(C)Oc1ccccc1N2CcN(CC2)C3CCC(CC3)NC(=O)Nc4ccc(F)cc4F

128 OP(=O)(O)C(NC1CCCCC1)P(=O)(O)

129 CN1CCN(CC1)c2nc(C3CC(C(=O)NC3=O)c4c[nH]c5ccccc45)c6ccccc6n2

130 Cc1ccc(F)c(Oc2ccccc2C3CCNCN3)c1

131 N[C@@H]1CC[C@H](CC1)NC(=O)c2cc(Oc3ccc(cc3)C(=O)N)cc(Oc4ccc(cc4)C(=N)N)c2

132 Cc1ccc2c(cccc2n1)N3CCN(CCc4cccc5c4OCc6c(ncn56)C(=O)N7OCC7)CC3

133 OC[C@H](NC1CCN(CCCc2c[nH]c3ccc(cc23)n4cncc4)CC1)c5ccccc5

134 Cc1ccc(cc1)c2nc(C3cccccc3c2ccCCCC2)C2

135 Cl.CC(C)Sc1ccccc1N=C2NNCCN2

136 Clc1ccc2CCcCCc2c1CC

137 CCCNCCc1c[cH]c2ccc(F)ccc21

138 Cl.CCC(=O)C(C1CCNC31)C23cc(Cl)c(Cl)c2

139 CCCCCCCC(=O)C(=O)CCCCCC

140 Fc1cccc(n1)C(=O)NCCCCN2CCNCCC22c3cccc(C)Cc3C2

141 NS(=O)(=O)c1ccc(cc1)c2ccncCc2CCCC

142 Cc1cc(Cl)ccc1NC(=S)NSC(=O)C(O)(c2ccccc2)c3ccccc3

143 Nc1c2CCCCc2c3cccccc13

144 OC1=C(Oc2cc(O)cc(O)c2c1=O)c3ccc(O)c(O)c3

145 CC(N)Cc1c[nH]2cccc3OCCCc3c12

146 CC(C(=O)O)c1ccc(c(F)c1)c2ccccn2

147 CC.CCSCC(C1CCNCC1)c2ccc(Cl)c(Cl)c2

148 CCc1cc(Cl)ccc1Oc2cc(C)ncc2CN

149 Clc1cc2NCC3CNCCN3c2cc1CC

150 CCCN(CCC)CCc1cccn2NC(=O)Cn12

151 NC(=O)c1ccc(CNC(=O)[C@H](CCC2CCNCC2)NC(=O)[C@@H](CCC3CCNCC3)NS(=O)(=O)Cc4ccccc4)c[illegible]

152 CN1CCC(=C2c3ccccc3CCCc4ccccc24)CC1

153 CN[C@H](Cc1ccccc1)C(=O)N2CCC[C@H]2C(=O)N[C@@H](CCCN=C(N)NCC(=O)c3nc4ccccc43)

154 CN(C)CC1CC2N(c1)cncc(F)ccccCc4ccccc24

155 CC(=O)Nc1nnc(s1)S(=O)(=O)NO

156 C[C@H](Oc1ccccc1CCCC)C2CCCCN2C

157 C\\NCC\\1/Oc2ccccc2C=C1 1

158 Cl.COc1ccc(Cl)c(CO)c1C2CC2CN

159 CN(C)CCOC(c1ccccc1)c2cccccCC2

160 COc1ccccc1N2CCN(CCCCCC(=O)c3cccn4nccc34)CC2

161 Oc1cccc(CCN(CCc2ccccc2)CC3Cc3)c1

162 Clc1ccc(CNC2CC(Nc3ccncc3)C(NC)C2=O)cc1

163 Oc1c2ccccc2c3N=Ncccccccc31

164 CN1CCN(Cc2ccc(NC(=O)Nc3cc(CCCc4ccc(N)nc4)n(C)n3)cc2C(F)(F)F)CC1

165 COc1cc2ncnc(Nc3ccc(F)c(c3)C#C)c2cc1OCCCCCCC(=O)CO

166 Clc1cc2CNC(CO)c2cc1OCCCN3CCN(CC3)c4cccc5ccccc45

167 O=C1NOc2ccc(OCCCCN3CCN(CC3)c4cccc5ccccc45)cc12

168 Clc1ccc2CNNCCc2c1Cl

169 CC(N(Cc1ccccc1)S(=O)(=O)c2c(F)c(F)c(F)c(F)c2F)C(OO)N

170 COc1ccc(cc1OC2CCCC2)C(CO)Nc3c(CC)cccc3CC

171 N[C@@H]1CC[C@H](CC1)N(OO)Oc2cc(Oc3ccc(cc3)C(=O)N)cc(Oc4ccc(cc4)C(=N)N)c2
172 O=C(NCCc1cccs1)N2CCC(=CC2)c3n[nH]c4ncccc34
173 CN1CCC(=CCc3ccccc3CCC4scccc4)CC1
174 Clc1ccc(cc1)c2cc([nH]c2)C3CCN(CCc4ccccc4)CC3
175 Cl.COc1nsnc1OCCOCCOCCOCCOc2snnc2C3CCCCCCCCC3
176 CCCc1nn(C)c2C(=O)NC(=Nc12)c3cc(ccc3OCC)S(=O)(NOONC)CCCCO
177 Cc1cc(NC(=O)Sc2ccc(cc2)c3cccc4[nH]nc(N)c34)ccc1F
178 C(Nc1ccccc1c2cccs2)C3=NCCNN3
179 Fc1ccc2c(cccc2c1)N3CCN(OCCOc4ccc5CNC(OO)c5c4)CC3
180 COc1ccccc1N2CCN(CCCc3cn(nn3)c4ccn5cccc5c4)CC2
181 CN(C)CC1CC2C(O1)c3ccccc3Cc4ccccc24
182 CCC1(Cc2ccccc2O1)C3=NCC3
183 CC1(N(CCc2cc(O)ccc12)c3ccc(O)Occ3)c4ccc(OCCN5CCCC5)cc4
184 CN(Cc1cc(Cl)c(O)c2ncccc12)c3CCCCC3
185 CN1C(=O)C(=Cc2ccc(Nc3ccc(N)cc3)nc12)c4c(Cl)cccc4Cl
186 Fc1ccc(Cl)c(c1)N2CCN(CCC3C(=O)C4)C(CCC4cCCCC33)C23
187 O=C(NCCc1cccs1)N2CCC(=CC2)c3c[nH]c4ccccc34
188 Cl.COc1ccc(Cl)c(Cl)c1C2CC2C
189 COc1cccc(c1)C(CC)NC(=O)c2ccc(cc2)c3ccncc3
190 COc1cc(ccc1NC(=O)OC(C)(C)C)c2cn(C(C)C)c3ncnc(N)c23
191 O=C(NCCCCC1CCN(CC1)c2ccccc2)c3cccccc4cc4n3
192 C[C@H](Oc1ccccc1CCCC)C2=CCCN2
193 CC1=CC[C@@H]2[C@@H](C1)c3c(O)cc(cc3OC2(C)C)C(C)(C)CCCCCN
194 CNN(CC)CCOC(=O)C(c1ccccc1)C2(O)CCCCCC2
195 Clc1ccc(cc1)N2NCN(Cc3c[nH]c4ccccc34)CC2
196 CC1=CC[C@@H]2[C@@H](C1)c3c(O)cc(cc3OC2(C)C)C4CCCCC4CCCCCCCC
197 CCNC(=O)[C@H]1O[C@H]([C@H](O)[C@@H]1O)n2cnc3c(N)nccc23
198 COc1cc2nccc(Oc3ccc(NC(=O)C4=C(C)N(C)N(C4=O)c5ccccc5)cc3F)c2cc1OCC
199 Cc1ccc(cc1S(=O)(=O)N)c2oc(nc2)C(=O)Nn3CCN3
200 NS(=O)(=O)c1cccc(Nc2ncc3ccn(Oc4ccccc4)c3n2)c1
201 CC1(N(CCc2cc(O)ccc12)c3ccc(C)cc3)c4ccc(OCCN5CCCCC5)cc4
202 COc1ccccc1CNCCCCCCNCCCCCCCCNCCCCCCNCc2ccccc2O
203 C=S(=O)(NC1CON(CCCc2cccc3CCCCc23)C1)C4Nccs4
204 CC(N)Cc1ccn2nc2ccccc1
205 CCCN(CCC)C1CCC(OCC1)CC

206 COc1ccccc1N2CCN(CC(CO)Nc3cccc(C)c3)CC2

207 COc1ccc2[nH]c(Ccc(CNC(C)C)c2)1

208 COC(=O)c1ccccc1NCC2=C=CN2

209 Cc1cccc2N3CCNCC3NC(OO)c12

210 CC(C)Cc1cc(on1)c2ncc3[nH]nc(C)c3c2

211 Clc1cccc(Cl)c1C(=O)Nc2c[nH]nc2C(NO)NC3CCNCC3

212 NS(=O)(=O)c1ccc(CC(=O)NCc2ccccc2)cc1

213 CN1CCN(CC1)c2nc(C3=C(C(=O)NC3OO)c4c[nH]c5ccccc45)c6ccccc6n2

214 CN1N=C(S/C/1=CCC(=O)C)S(=O)(=O)N

215 COc1ccccc1N2CCN(CNCC3COc4ccc(Cl)cc4O3)CC2

216 Clc1cccc(N2CCN(CCCCOc3ccc4C=CC(CO)Cc4n3)CC2)c1Cl

217 COc1cc2nc(nc(N)c2cc1OC)N3CCN(CC3)C(OO)C4CCSS4

218 Nc1ncc(c2cocc2)c3scc(c4ccc(NC(NO)Nc5cccc(F)c5)cc4)c13

219 Nc1nc(N)c2c(Cl)c(CNc3ccc(cc3)C(=O)CC(CCC(=O)O)C(=O)O)ccc2n1

220 Cc1ccc(NC(=O)Nc2ccc(cc2)c3cccc4C(=C)NCc34)cc1C

221 OC[C@H](CC1CCN(CCCc2c[nH]c3ccc(cc23)n4cnnc4)CC1)c5ccccc5

222 COc1cc(ccc1O)c2ccc3C(=O)Nc4cc(ccc4Nc3c2)C(=O)CCCNC(=O)C

223 COC(=O)c1cc(CCc2cc(OO)ccc2OO)ccc1

224 CCCN(CCC)C1CCc2ccc(cccc2C1)

225 CCCN(CCC)CCc1cccnN2C(=O)Nc12

226 CCCCCCCC(=O)C(=O)CCCCCCCC

227 COC(=O)C1=C(C)NC(=O)N(C1c2ccc(F)c(F)c2)C(=O)NCCCN3CCC(CC3)c4ccccc4O

228 CN1CCc2c(C1)sc3N=CN(CCN4CCN(CC4)c5ccccc5CC)C(=O)c23

229 CCCN(CCC)C1CCc2ccccOccc2C1

230 Cc1cc(C)c2c(n1)sc3c(N)ccnc23

231 NNc1ccccc1S(=O)N=O

232 C(Oc1ccccc1c2ccccc2)CC=CCCN

233 COC(=C)c1cc(CCc2cc(OCCccc2)O)ccc1O

234 COc1ccc(cc1OC)S(=O)(=O)N[C@@H]2CC[C@H](CC2)N3CCC(CC3)c4ccccc4OCC(F)(F)

235 COc1cccc(CNC(=O)c2cc3c(n[nH]c3s2)c4cccnc4)c1

236 NS(=O)(=O)c1ccc(cc1)n2cncnn2c2ccccc2

237 CC(=O)Nc1ccc2c(CCCCCCN(Cc4ccccc4)CC)cccc2c1

238 Clc1ccc(cc1Cl)S(=O)(=O)NCCCCN2C(NCCC2)c3noc4ccccc34

239 Cc1ccc(cc1)C2=CC(C)(C)Oc3cc4ccc(cc4cc23)c5ccc(cc5)C(=O)

240 Brc1ccc(COc2ccc(C#N)c(c2)C#C)cc1

241 Clc1ccc(cc1Cl)C(=O)C(CC=C)N2CCcC2

242 Fc1ccc2c(nsc2c1)C3CCN(CCCCNS(=O)(=O)c4cccc5scnc45)CC3

243 Clc1ccc(cc1)C(=O)NCCN2CCCCC2

244 COc1ccc(cc1)C2CN(C)Cc3cc(OCCCN4CCC(C4CC))ccc23

245 O=C(N1CCCCC1)N2CCn3cc(C4=C(C(=O)NC4OO)c5cnc6ccccn56)c7cccc(C2)c37

246 O=C(N[C@@H]1CC[C@@H](CCN2CCC(CC2)c3coc4ccccc34)CC1)C5CCCON5

247 CNCC1(CCCCC1)c2ccc(CC)c(Cl)c2

248 NS(=O)(=O)c1cccNNCCCCcc1

249 COc1cc2c(Oc3ccc(Nc4ccc(cc4)C(C)(C)C)cc3)ccnc2cc1OCCNCC

250 CNCCC(Oc1ccccc1OO)c2ccccc2

251 CCCC(C1CCCc1)C(=O)c2ccc(CC)cc2

252 NS(=O)(=O)c1cc2ccc(OCS)cCc2s1

253 CCCCc1nc(N)c2nc(n(C)c2n1)n3ncnn3

254 Nc1ncnc2c1c(nn2C3CCCCCC3)c4ccc(O)c(O)c4

255 FC(F)(F)c1ccc2c(NC(=O)Nc3cccc(c3)C#N)ccnc2c1

256 CC(C)Oc1ccccc1C2CCN(CC2)[C@@H]3CC[C@H](CC3)NS(=O)(CO)c4ccccc4O

257 NC(=N)N1CCC[C@H](NC(=O)CN2CCNC[C@H](NS(=O)(=O)Cc3ccccc3)C2=O)C1O

258 N\\N=C\\1/Oc2ccccc2C=C1O

259 Clc1ccc(COc2ccc3NC(=O)Oc3c2)cc1

260 C(N1CCC(CC1)c2cc([nH]n2)c3ccccc4)c4ccccc3

261 Clc1ccc2CCCCCc2c1Cl

262 OC1=C(Oc2cc(O)cc(O)c2C1=C)c3ccc(O)c(O)c3

263 Fc1ccc2c(cccc2c1)N3CCN(CCCOc4ccc5ONC(=O)c5c4)CC3

264 CCCCN1C2=C(CCCCCC2)C=C(C(=O)NC3CCCCC3)C1N

265 COc1cc2c(Nc3ccc(cc3)c4nc5ccccc5s4)ncnc2cc1OOCCN(CCN(C))C

266 COc1cc(ccc1Nc2ncc(Cl)c(Nc3ccccc3P(=O)(C)C)n2)C4CCN(COC)4

267 Cl.COc1ccc(cc1)N(C)c2nc(C)cc3Cc(C)cc23

268 CCc1ccc2c(NC(=O)Nc3cccc(n3)C(F)(F)F)cccc2c1

269 Cc1cc(C)c2c(n1)sc3c(N)nnnc23

270 COc1ccc2c(c1)[nH]c3c(C)cccc23

271 Clc1ccc(cc1)C(=O)CCCC2CCc3ccccc3C2

272 CC[C@@]1(O)C(=O)OCC2=C1C=C3N(Cc4cc5c(N)cccc5nc34)n2CO

273 COc1cc2ncnc(Nc3cccc(c3)C#C)c2cc1CCCCCCCC(=O)NO

274 Cc1ccc2c(cccc2n1)N3CCN(CCc4cccc5c4OCc6c(ncn56)C(OO)NC7CCCC7)CC3

275 CSc1ccc(Cl)cc1NC2C=CCNN2

276 COC(=O)c1ccccc1CCC2NCCCN2

277 COc1ccccc1NNCCN(CCCCN3C(=O)CC(CC3=O)c4ccccc4)CCC

278 CC(N)C1CCC(CC1)C(=O)Nc2ccccn2

279 COc1cc(NCc2ccc3nccNcnc(N)c3c2C)cc(OC)c1OC

280 CC(C)Nc1nccc(Nc3ncc(s2)c3ccccc2)n1

281 CCCCCCNS(=O)(=O)NCNC(=O)O

282 CC(N(Cc1ccccc1)S(=O)(=O)c2c(F)c(F)c(F)c(F)c2F)C(OO)NO

283 COc1ccc(cc1)C(NO)CCCN2CCc3ccccc3C2C

284 NS(=O)(=O)OCCCCCCCCCCOS(=O)(OO)N

285 CN1C(=O)C=NN(CCCCN2CCN(CC2)c3ccccc3CCCF)C1=O

286 CCNc1ccccc1c2ncccc2SC3CNCCN3

287 CCn1nnc(n1)[C@H]2O[C@H]([C@H](O)[C@@H]2O)n3cnccc(N)nccnn3

288 CN(Cc1coc2nc(N)nc(N)c12)c3ccc(cc3)C(OO)NC(CCC(=O)O)C(=O)O

289 CCn1nnc(n1)[C@H]2O[C@H]([C@H](O)[C@@H]2O)n3cnc4c(NC)Cc(CC)nc3c4

290 CS(=O)(=O)CCN(CCNN)Cc1oc(cc1)c2ccc3ncnc(Nc4ccc(OCc5cccc(F)c5)c(Cl)c4)c3c2

291 CC1=CC[C@@H]2[C@@H](C1)c3c(Occc(cc3OC2(C)C)C4C)(C)CccC4C

292 CN(C)C[C@@H]1CC2[C@H](O1)cccc(F)ccccSc4ccccc24

293 C[C@@](CCN1CCC(C)CC1)N(C)S(=O)(=O)c2cccc(C)c2

294 Brc1ccc(Cn2nncc2c3ccccc3)cc1

295 Cc1cccc(NC(=O)Nc2ccc(cc2)c3csc4c(cnc(N)c34)C#CCC5CCCC5)c1

296 OC(CCCN1CCCCC1)Oc2ccccc2Cc3ccccc3

297 Cc1c(Nc2ncc[CH]2)ccc3OCCOc13

298 Clc1ccc(cc1)C(=O)NCCNOCNCCCN

299 BrCc1ccc(cc1)C(=O)C(OO)c2ccccc2

300 NC(=N)c1ccc(CNC(=O)[C@H](CCC2NCNCC2)NC(=O)[C@@H](CCC3ccncC3)NS(=O)(=O)Cc4ccccc4)c[illegible]

301 C(Cc1ccccc1c2nccs2)C3=NCCN3

302 CN1C(=O)C=Cc2c(CCN3CCN(CC3)c4cccc5nc(C)ccc45)c(CC)ccc12

303 Fc1ccc(SCCCN2CCN(CC2)c3ncccc3)cc1

304 COc1ccccc1N2CCN(CCCCN3Cc4ccccc4C3OO)CC2

305 COc1cccc2CC[C@@H]3[C@@H](CCCCCC3C)c21

306 Brc1c(NC2CCCCN2)ccc3nccnc13

307 COc1cc2ccccc2ccnncccccc1

308 COC(=O)C(C1CCCC=1)c2ccc(Cl)cc2

309 COc1ccc(F)cc1CCCCCCCCCCNOOC

310 CC1=CC[C@@H]2[C@@H](C1)c3c(O)cc(cc3OC2(C)C)C(C)(C)CCCCCC

```
311 CN(C)Cc1ccccc1Nc2cccc(Cl)c2
312 COc1ccc(F)c(F)c1C(=O)c2cnc(NC3CCC(CC3)S(=O)(=O)C)nc2N
313 CC(C)C(C1CCCCC1)c2ccc(Cl)cc2
314 CN(C)C[C@H]1C[C@H]2[C@H](c1)c3ccccc3Cc4ccccc24
315 CCCc1cn(C)c2C(=O)NC(=Nc12)c3ccccc3OCC
316 CC(C)(C)OC(=O)N[C@@H](C(=O)N1CCC[C@H]1C(=O)N[C@@H](CCCNCC(N)N)C=O)c2ccccc2
317 Cc1ccc(cc1S(=O)(=O)N)c2oc(nc2)C(=O)N3ncCC3
318 COc1cc2ncnc(N[C@H](C)c3ccccc3)c2cc1CCCCCCCC(=O)NO
319 OC[C@H](Nc1CCN(CCCc2c[nH]c3ccc(cc23)n4cncc4)CC1)c5ccccc5
320 Nc1nc(N)c2nc(CNc3ccc(cc3)C(=O)OC(CCNC(=O)c4ccccc4C(=O)O)C(=O)O)cnc2n1
321 Cc1ccc2c(cccc2n1)N3CCN(CCc4cccc5c4OCc6c(ncn56)C(=O)N7CCCC7)C3
322 COc1cccc(c1)C(C)NC(=O)c2ccc(cc2)c3ccccc3
323 Clc1cccc(N2CCN(CCCCOc3ccc4CCCC(=O)Nc4n3)CC2)c1Cl
324 CN(C)C1(CCCCC2CCCC2)COc3ccccc3OC1
325 CC1=CN=C(NCC(F)(F)c2ccccn2)C(=O)N1CC(C)NCCc3ccc(N)nc3C
326 CN1CCc2ccccc2Cc3c(CC1)ccc(O)c3CC
327 CCCCC(CC(=O)NO)S(=O)(=O)N1CCCCCC1
328 COc1ccc2CCC(Cc2c1)NS(=O)(=O)N(C)
329 NCCCC[C@H](N1Cc2[nH]c3ccccc3c2C[C@@H](NC(OO)Cc4ccccc4)C1=O)C(=O)NCc5ccccc5.OC(=O)[illegible]
330 OCCNCCNc1ccc(NCCNCCO)c2C(=O)c3c(O)ccc(O)c3C(OO)c12
331 CC1(N(CCc2cc(O)ccc12)c3ccc(CO)cc3)c4ccc(OCCN5CCCC5)cc4
332 Clc1cccc(Cl)c1NC2=NCCc2
333 NS(=O)(=O)c1cnc(s1)c2ccccc2C
334 CCCCCCC(C)(C)c1cc(O)c2c(OC(C)(C)c3ccc(CC)cc23)c1
335 CN1CCc2ccccc2Cc2ccsc2CC1
336 CCc1ccccc1NCC2=NCCN2
337 Cc1c(sc2ccc(F)cc12)S(=O)(=O)NNCCCn3CCC(CC3)c4noc5cc(F)ccc45
338 Oc1ccc2C3=C(CCOc2c1)c4ccc(O)cc4O[C@@H]3c5ccc(CCCC6CCOCC6)cc5
339 Ic1c(NC2=NCCN2)ccc3cccnc13
340 Ic1c(NC2CCCCC2)ccc3ccccc13
341 COc1ccccc1N2CCN(CCCCN3N=CC(CO)N(C)C3=O)CC2
342 Fc1ccc(cc1)C(N2CCN(C\\C=C\\c2ccccc3)CC3)c4ccc(F)cc4
343 CNC1=Nc2cccc(CN)c2C(C)C1
344 Ic1c(NC2CNCCC2)ccc3cccnc13
345 CC(C)Oc1ccccc1N2CcN(CC2)C3CCC(CC3)NC(=O)Nc4cc(F)ccc4F
```

346 COc1ccc(cc1)C(CNCC)CCC2(O)CCCCC2
347 NS(=O)(=O)c1ccc(cc1)NNCC
348 CN1CCc2ccccc2Cc3[nH]c4ccccccccccc3C41
349 CCCCCCCSOS(=O)(=O)N
350 C[C@H](N)CN1ncc2ccc(O)cc12
351 Clc1cccc(N2CCN(CCCCOc3ccc4CCCC(C=O)c4n3)CC2)c1C
352 COC(=O)C1=C(C)NC(=O)N(C1c2ccc(F)c(F)c2)C(=O)NCCCC3CCN(CC3)c4ccccc4CC
353 Cc1cccc(N2CCN(CCc3ccc4[nH]ncc4c3)CC2)c1CC
354 C[C@H](Oc1ccccc1CCCC)C2=CCCN2C
355 COc1ccccc1N2CCN(CCCCCC(=O)NC(C)(C)C)CC2
356 OC1(N2CCNNC2c3ccccc13)c4ccc(Cl)cc4
357 Fc1cccc2cccc(N3CCN(CCCCNc4ccc5CNC(=O)c5c4Cl)CC3)c12
358 COc1cc2ncnc(Nc3ccc(F)c(Cl)c3)c2cc1OOCCCCCC(=O)NO
359 COc1ccc(c2ccc(NC(=O)Nc3ccc(N)c(C)c3)cc2)c4c(N)noc14
360 COc1ccc(cc1OC)S(=O)(=O)N[C@@H]2CC[C@H](CC2)N3CCN(CC3)c4ccccc4OC(C)
361 COc1nccc(n1)c2c(nnn2C3CCNCC3)c4ccc(F)cc4
362 Cl.Cl.CNCc1cc2ccc2ccc3ccn34ccc4cccccc1C
363 C[C@H](c1c[nH]cn1)c2cccc(CC)2C
364 COC(=O)C(C1NCCCN1)c2ccc(Cl)c(Cl)c2
365 NC(=N)c1ccc(CNC(=O)[C@@H]2CCN2C(=O)[C@H](NCC(=O)C)C3CCCCC3)cc1
366 CN(CCCOc1ccc(Cl)cc1CC)CC#C
367 FC(F)(F)Oc1ccccc1CC(NCCCNCCC)3cccccc3
368 Cc1cccc(c1)N2CCN(CCCCNC(=O)c3cccc4cccc4c3)CC2
369 Fc1cc(C)c2c(n1)sc3c(N)ccnc23
370 CNc1cc(NS(=O)(=O)c2ccc(N)cc2)cc(CC)n1
371 Ic1c(NC2=NCCN2)ccc3ncnnc13
372 CCCCN1CCC(C(C(=O)c2cc(CC)c(CC)c3CCC)c23)CC1
373 Fc1ccc2c(noc2c1)C3CCN(CCCCNS(=O)(=O)c4ccc5csccc45)CC3
374 CCOc1nc(NC(=O)Cc2cc(OC)c(cc2CC)S(=O)(=O)C)cc(N)c1C#N
375 COC(=O)c1ccccc1NCC2=CCCC2
376 COC(=O)c1ccccc1NCC2NNCCC2
377 CN(C)C1CCC2(C=C1)c3ccccc3CCc4ccc(CC)cc24
378 CN(C)CC[C@@H](c1ccc(Cl)cc1)c2ccccc2
379 Cl.COc1cc(F)c(F)cc1C2CC2C
380 Clc1cccc(N2CCN(CCc3cn(nn3)c4ccn5nccc5c4)CC2)c1CN

```
381 CN1CCc2C(C1)c3ccccc3cc4ccccc24
382 CCCN(CCC)CCc1cccc2cC(=O)CN12
383 COc1ccc(cc1)C2CC(CCC2)c3ccc(cc3)S(=O)(=O)C
384 C1COC(=NC1)CC2CCCc3ccccc23
385 CN[C@@H](C(=O)N1CCC[C@H]1C(CO)NC(CCCN=C(N)N)C(=O)c2nc3ccccc3s2)c4ccccc4
386 CN(C)C1(CNCCN2CCCC2)COc3ccccc3OC1
387 Fc1ccc(CC(CC3CNCC)3)cccc1
388 CN1CCN(CC(=O)N2c3ccccc3C(=O)Oc4cccc42C)CC1
389 CN1CCN(CC1)c2ccc(cc2)C(=O)N[C@@H]3CC[C@@H](CNN4CCC(CC4)c5coc6ccccc56)CC3
390 N\SN=C\\1/cc2ccc1c2C=C
391 CSc1ccc(F)cc1NCC2=NCCC2
392 NS(=O)(=O)Sc1ccc(I)cc1 1
393 CCCN1C(=O)N(CCC)c2nc([nH]c2C1=O)c3ccc(OCC(=O)N(C))cc3
394 NS(=C)(=O)C
395 Fc1ccccc1Oc2cccc(F)c2C3CCcCC3
396 CC(C)CC1CCCCN1c2ccc3[nH]ncc3c2
397 COc1ccccc1N2CCN(CCCCNC(=C)c3cnn4ccccc34)CC2
398 CCc1ccccc1NCC2=NCCN2C
399 COc1cc(ccc1Nc2ccc(Cl)c(Nc3ccccc3C(=O)N)n2)P(=O)(C)CC
400 CCn1nnc(n1)[C@H]2O[C@H]([C@H](O)[C@@H]2O)n3cnc4c(NC)nccc34
401 C(Oc1ccccc1c2ccccc2)C3=NCCN3 3
402 Fc1ccc(OCCCC2CCC(CC2)N3C(CO)Nc4ccccc34)cc1
403 CCC[C@H]1CCCC[C@@H]1NC2CNCCC2
404 COc1cccc(c1)C(C)NC(=O)c2ccc(cc2(CCC))c3ccncc3
405 Brc1c(NC2CNCCN2)ccc3NCCNc13
406 CCOc1cc(CN2CCCNCC2CNc3oc4ccccc4n3)ccc1CC
407 COc1cc2ncnc(Nc3ccc(F)c(Cl)c3)c2cc1OCCCCCCC(=O)OO
408 Nc1nc(N)c2c(CN)c(CNc3ccc(cc3)C(=O)NC(CCC(=O)O)C(=O)O)ccc2n1
409 C1CN=C(N1)C2COccccccccO2
410 Cc1cccc(NC(=O)Nc2ccc(NC(=O)c3csc4nnnc(N)c34)cc2)c1
411 Cc1cccc(CCN(CCc2ccccc2)CC3Cc3)c1
412 Cc1ccc(cc1)c2cc(nn2c3ccc(cc3)S(=O)(=O)NNC(F)(F))
413 COc1cccc(c1)C2CCC(CC23NNCCN(CCC))cccccc3
414 CC(=O)N1CCC(CC1)NC(=O)Nc2ccc(CC(C)(F)F)cc2
415 COc1cc(ccc1Nc2ncc3CN(C)C(=O)N(c4cccc(NC(=O)C=C)c4)c3n2)NCCCCC
```

416 O=C1Nc2sc3CCcCc3c2c4nc(nn14)c5ccncc5
417 CC(=O)OCC[N+](CCCC)C
418 Ic1c(NC2CCCCN2)ccc3ccccc13
419 COc1cc2OC(=O)C=Cc2cc1[C@@H](CCCCCC)C
420 CC(C)NC[C@H](O)OOc1cccc2ccccc12
421 Cc1ccc(F)cc1C(=O)Nc2ccc(cc2)C(=O)N3CCC4(CCCC=C4)C(=O)NCCN5CCCCC5Cc6ccccc36
422 Cl.CCNC1CCS(=O)(=O)c2sc(cc12)S(OO)(=O)N
423 NS(=O)(=O)Oc1ccc(N)cc1
424 Brc1c(NC2=NCCN2)ccc3nccn31
425 COc1ccc(cc1)S(=O)(=O)NC(C)C(=O)NO N
426 N[C@@H]1CC[C@H](CC1)C(OO)Oc2cc(Oc3ccc(cc3)C(=O)N)cc(Oc4ccc(cc4)C(=N)N)c2
427 Clc1ccc(cc1)N2CCN(CCN3CCCCCC3)C2OO
428 C(Nc1nc(Nc2cc([nH]n2)C3CC3)c4sccc4c1)c5ccccc5
429 OC1=C(Oc2cc(O)cc(O)c2c1=O)c3ccc(O)cc3
430 O=C1NC(=NC(=C1)c2ccncc2)NCCCCCCCC
431 CNC[C@H](O)c1ccc(O)(CO)c1
432 O=CCCCCCC1CCN(CCCN2C(=O)CC23cccccc3)CC1
433 CN1CC(c2ccc3c(c2)C(C)(C)CCC3(C)C)c4cc(oc4C1)\\CCC\\C(=O)O
434 Cc1ccccc1c2cncn2Cc3ccc(cc3)CNN
435 COc1cccc(c1)C(CN)NC(=O)c2ccc(cc2)c3ccccc3
436 CCCN(CCC)C1CCC(=CC1)CNC
437 COC(=O)C(C1CCCCC1)c2ccc(CC)cc2
438 COc1ccccc1N2CCN(CCCCNC(=O)CC(C)(C)C)CC2
439 NS(=O)(=O)c1nnc(s1)c2ccccc2
440 Cl.CC(C)Sc1ccccc1NCC2CCNCN2
441 FC1(F)CC1C(OO)N[C@@H]2CC[C@@H](CCN3CCC(CC3)c4coc5ccccc45)CC2
442 COc1c(NC(=O)Nc2ccc(c3ccc(CN4CCOCC4)nc3)c5ccccc25)cc(cc1NS(=O)(=O)C)C(C)(C)CC
443 CC1(N(CCc2cc(O)ccc12)c3cccc(OC)c3)c4ccc(OCCN5CCCC5)cc4
444 CS(=O)(=O)CCN(CCCN)Cc1oc(cc1)c2ccc3ncnc(Nc4ccc(OCc5cccc(F)c5)c(Cl)c4)c3c2
445 CCCCCCNS(=O)(=O)NOCC(=O)O
446 O=C(CC1COCCO1)N[C@@H]2CC[C@@H](CCC3CCC(CC3)c4coc5ccccc45)CC2
447 COc1ccccc1N2CCN(CCCCNC(=O)c3cc4ccccc4s3)CC2C
448 CNC[C@H](O)c1ccc(O)c(O)c1O
449 Ic1c(NC2=CCCN2)ccc3ccccc13
450 N#Cc1ccc(cc1)C2CCC23c2cn23

451 Cl.COc1ccc(F)c(Cl)c1C2CC2

452 Nc1c2CCCCc2ccccccccc1

453 COc1ccc2[nH]cc(CCN(CC=C)CCCC)c2c1

454 CCCC(NC1CCC1)C(=O)c2ccc3ccccc3c2

455 CSc1ccc(Cl)cc1CCC2=NCC2

456 Clc1ccc(CNC2=C(Nc3ccncc3)C(=O)C2=O)cc1C

457 COc1cc(C=C2SC(=S)NC2=O)ccc1

458 Cc1ccc(OCONCCCOc2ccccc2)cccCc1

459 CC1(N(CCc2cc(O)ccc12)c3cccc(O)c3)c4ccccOCCN5CCCC5ccc4

460 Fc1ccc(Cl)c(c1NN2CCN(CC(CC(=O)C4)CCCCC4)CCCCC)CC2

461 C(CC1(CCNC1)c2ccc3ccccccc3c2)c4ccccc4

462 CN(C)C1(CNCCCCCCC(OOCCC)C)c3ccccc3OC1

463 O=C(NCCCCN1CCN(CC1)c2ccccc2)c3cccc4cccc4c3

464 Cc1cc(Cl)ccc1NC(=S)NCC(=O)C(O)(c2ccccc2)c3ccccc3

465 Cc1ccc2c(cccc2n1)N3CCN(CCc4cccc5c4)N5CCOCC(O)CC3

466 CNC(=O)[C@H]1S[C@H]([C@H](O)[C@@H]1O)n2cnc3c(NC)nc(Cl)n32

467 Fc1ccccc1N(Cc2ccc(CO)cc2)C3CCC3

468 COc1ccccc1CNCCC(Cc3cc4cccccn3)C4C

469 CCCN1C(=O)N(CCC)c2nc([nH]c2C1=O)c3ccc(OCC(=O)NCNN)cc3

470 COc1ccc(cc1)S(=O)(=O)NC(O)C(=O)NOO

471 CCCCCOc1ccccc1c2nCc(c2)C(=O)NC3CCCCCC3

472 CN(C)C[C@H]1CC2[C@@](O1)c3cc(F)ccc3Sc4ccccc24

473 CCOc1nc(cc(NCc1C#N)CCCO)NCc2cccnc2

474 CC1=CC[C@@H]2[C@@H](C1)c3c(O)cc(cc3OC2(C)C)C(C)(CCCCCCC)

475 Cc1ccc2c(cccc2n1)N3CCN(CCc4cccc5c4OCc6c(C)nnn56)C3

476 COc1c(C)c2COC(=O)c2c(O)c1C\\C\C(C\C\\CCOC=O)O

477 O=C(Nc1nnc(c2ccccc2)c(n1)c3occc3)C4CC4

478 COc1ccccc1CNCCCCCCCNNCCCCCCNCCCCCCNCc2ccccc2CC

479 Nc1ccc(cc1)S(SO)(=O)N

480 OCc1cc(ccc1O)C(O)NCCCCCCCCCCCCc2ccccc2

481 CC(C)n1nc(c2ccc(BC)c(O)c2)c3c(N)ncnc13

482 CC(C)Oc1ccccc1N2CCNNCc3cccc(c3)C(=O)N4CCCCC4CCC2

483 CCCCOC(=O)c1ccccc1NCC2NNCCN2

484 Cc1cc(Nc2ncc(s2)C(=O)Nc3c(C)cccc3CC)nc(C)n1

485 NCCn1ncc2ccc(C)cc12

486 Fc1ccc2c(noc2c1)C3CCN(CCCCNS(=O)(=O)c4ccc5ccccc5c4)C3

487 Oc1cccc(CCN(CCc2ccccc2)CCNCCC)c1

488 Fc1ccc(cc1)c2ccc(cc2)C(=O)Nc3ccc4nc(CCCCCCCC)cccc43

489 COC(=O)c1c(C)n2ncnc(N)c2c1c3ccc(N)(=O)Nc4ccccc4ccc3

490 C\\N=C\\1OOc2ccccc2CCC1

491 CCCCCCCCC(=O)C(=O)CCCCCC

492 FC(F)(F)c1ccc2c(NC(=O)Nc3ccccn3)ncnc2c1

493 Cc1ccc2c(cccc2n1)N3CCN(CCc4cccc5c4OCc6c(ncn56)C(=O)NCCCCCCC)CC3

494 Cc1cccnc1NC(P(=O)(O)O)P(=O)(OOO)

495 NS(=O)(=O)c1ccc(cc1)c2ccccc2 c

496 Fc1cccc(COc2ccc(Nc3ncnc4cc(C#C[C@@H]5C[C@H](CN5)CC(=O)N6CCOCC6)sc34)cc2Cl)c1

497 CC(C)NCC(O)COc1cccc2cnccc12

498 COC[C@@H]1CCCN1N2NCN(C23O)ccccc(CCC)c3

499 COc1cc(ccc1Nc2ncc3C(=O)N(CC(=O)N(c4cccc(NC(=CCCCC)c4)c3n2)c5ccccc5)N6CCNCC)CC6

500 Cn1c(SCCCN2CC3CCN(C3C2)c4ccc(cc4)C(F)(F)F)nnc1n5CCCCC5

501 CCCCC(CC(=O)NO)S(=O)(=O)c1cccccccccccc1

502 Nc1ncnc2c1c(cn2C3CCCc3)c4cccc(O)c4

503 Fc1ccccccc2cccc(C)N2C3CCNCC3Ncc1

504 COc1ccccc1N2CCN(CCCCCCNC(=O)CC(C)(C)C)CC2

505 NS(=O)(=O)c1ccc(cc1)C#CC

506 O=C1NC(=O)\\C(=C\\c2ccc3ccccc2c3)nOC1

507 NS(=O)(=O)c1ccc(cc1)c2cccccccCC(OO)2CC

508 Cl.CCC(=O)C(C1CCNCC1)c2ccc(Cl)c(CC)c2

509 NS(=O)(=O)c1ccc(cc1)CNC

510 Fc1ccc(cc1)C(=CCN2CCN(CCCc3ccccc3)CC2)c4ccc(F)cc4

511 O[C@@H](CNCCCCCCCCN1CCC(CC1)OC(=O)Nc2ccccc2c3ccccc3)c4ccc(O)c5CC(=O)CCCc45

512 CSc1ccc(Cl)cc1NC2C=NC=N2

513 CCN1CCN(Cc2ccc(NC(=O)Nc3ccc(Oc3cc(NC)ncn4)cc4)cc2C(F)(F)F)CC1

514 Cc1cccc(NC(=O)Nc2ccc(cc2)c3coc4nccc(N)c34)c1

515 CCn1nnc(n1)[C@H]2O[C@H]([C@H](O)[C@@H]2O)n3cnccc(N)ncncn3

516 NS(=O)(=O)c1ccc(cc1)C2=C(C(CO)OC2)c3ccccc3

517 COc1ccc(CN(CCC(C)C)c2ccccn2)cc1

518 COc1ccc2[nH]cc(CCN(C)O)c2c1

519 Cc1c(OCCCN2CCN(Cc2)c3cccc4ccccc34)ccc5CCC(OO)c15

520 Fc1ccc(cc1CC)S(=O)(=O)NCCCN2CCN(CC2)c3nsc4ccccc34

521 COC(=O)C1C2CCC(CC1c3ccc(CC)c(Cl)c3)N2C

522 COc1cc2c(Nc3ccc(cc3)c4nc5ccccc5s4)ncnc2cc1OCCCCCCCN(C)CC

523 CN1CCCC(CC1)N2N=c(Cc3ccc(Cl)cc3)c4ccccc4C2=O

524 CN(C)Cc1ccccc1Sc2ccc(cc2N)CN

525 Nc1nccc2scc(c3ccc(NC(=O)Nc4cccc(BO)c4)cc3)c12

526 Clc1ccc(CNC2=C(Nc3ccccc3)C(=C)C2=O)cc1

527 CCCc1nn(C)c2c(=O)NC(=Nc12)c3ccccc3OC

528 COC(=O)C1C2CCC(CC1c3ccc(CC)c(Cl)c3)O2

529 CN1CSc2cccc(Cl)c2CC1

530 CNCc1cc(ccc1Oc2ccc(Cl)cc2OC)C(=O)C=CN

531 CC(C)NC(C)C(O)OOc1ccc(C)c2CCCc12

532 COC(=O)c1=CCCNCCCC1

533 COc1ccc2CCC(Cc2c1)NSS(O)(=O)N(C)C

534 CCCCC(CC(=O)NO)S(=O)(=O)C1CCCCCC1

535 COc1ccccc1N2CCC(CNCC3COc4ccc(Cl)cc4O3)CO2

536 Cc1ccc2c(cccc2n1)N3CCN(CC34c3cc(c4)n5cccn5)CC3

537 Clc1ccc(Cl)c(c1)N2CCN(CCCCN3C(CO)CCCCCCCCCCC3OC)CC2

538 Nc1cOc(cC1I)S(=O)(=O)N

539 NCCCCCc1c[nH]nn1

540 CCCN1C(=O)N(CCC)c2[nH]c(nc2C1=O)c3ccc(OCC(=O)Nc4ccccc3)cc4

541 CCCCCN(CCCCCCOPP(OO))(OPOOPOOOOOOO)

542 Clc1cccc(c1)N2CCN(CC3CCCO3)C2CO

543 N[C@@H]1CC[C@H](CC1)NC(=O)c2cc(OCc3cccc(c3)C(=N)N)cc(OCc4cccc(c4)C(=O)N)c2

544 Nc1nccc2scc(c3)cc(NC(=O)Nc4ccccc4ccc3)c12

545 NS(CO)(CO)c1=cc(O)cc1 1

546 OC[C@H]1O[C@H]([C@H](O)[C@@H]1O)n2cnc3c(NCc4cccc(I)c4)nccc23

547 Oc1ccc(cc1)N2CCc3cc(O)ccc3C2(C)c4ccc(OCCN5CCCC5)cc4

548 O=C(Nc1nnc(c2ccncc2)c(n1)c3occc3)CCCCC

549 C\\C(=C/C=C/C(=C/C(=O)O)/C)\\C\C\\C1=C(C)CCCC1(C)C

550 N[C@@H]1CC[C@H](CC1)C(=O)Cc2cc(Oc3ccc(cc3)C(=O)N)cc(Oc4ccc(cc4)C(=N)N)c2

551 Clc1ccc2cc(ccc2c1)S(=O)(=O)NCCCN3CCC(CC3)c4nsc5ccccc45

552 CC(C)Oc1ccccc1N2CCN(CC2)[C@@H]3CC[C@H](CC3)NS(OO)(=O)c4ccccc4O

553 COc1cc(C=C2SC(=Nc3ccccc3)NC2CO)ccc1O

554 Clc1cccc(N2CCN(CCCCOc3cc4Nc(=O)C=Cc4cn3)CC2)c1Cl

555 NS(=O)(=O)c1ccc(cc1)c2ccccc2 2

556 CO[C@H](C(C)C)C(=O)N[C@@H]1CC[C@@H](CCN2CCN(CC2)c3cccc4OCOc34)CC1C

557 Cc1ccc(cc1)n2cc3c(O)nc4ccccc4c3c2

558 COC(=O)C1=CCCNCCCC1

559 CN1CCccccccc2cc3cccC3C21

560 Clc1cccc(N2CCN(CCCCOc3ccc4CCCC(=O)N4cn3)CC2)c1Cl

561 CCCCCOc1ccccc1c2nCc(c2)C(CO)NC3CCCCCC3

562 COC(=O)[C@H]1C2CCC(C2)C[C@@H]1c3ccc(Cl)c(Cl)c3C

563 CC(Oc1ccccc1c2cccs2)C3=CCCC3

564 CCCn1cc(C(=O)N2C(C)(C)C2(C)C)c3ccccc13

565 COc1ccc(c2ccc(NC(=O)Nc3ccc(N)c(C)c3)cc2)c4c(C)noc14

566 COc1cc2c(Oc3ccc(NC(=O)C4=NN(c5ccccc5C4=O)c6ccccc6C(F)(F)F)cc3F)ccnc2cc1CCCCC7CCCC[illegible]

567 Cc1ccc(NC(=O)Nc2ccc(cc2)c3cccc4C(NO)NCc34)cc1C

568 COc1ccc(cc1)C(=O)CCCN2CCc3cccccc32C

569 CC1(C)C(C(=O)c2cn(CC3CCOCC3)c4cccOccc4cc2c1CCOC)C

570 Cn1ccc2cc(ccc12)C3(Cc4ccccc4)CCC3

571 CCCCOC(=O)c1ccccc1NCC2=NCCN2 C

572 Cc1cccc(NC(=O)Nc2ccc(cc2)c3csc4nccc(N)c34)c1

573 O=C1CC2(CCCC2)CC(=O)N1CCCC=CC3CCc4ccccc4C3

574 OC1(CCOCC1)c2ccnc(COc3ccc4c(cc(cc4c3)CCN)c5cocc5)c2

575 C\\N=C\\1/Oc2ccccc2CCC1

576 Cc1ccc2c(cccc2n1)N3CCN(CCc4cccc5N(CC(F)O)C(=O)COc45)CC3

577 NS(=O)(=O)Oc1cccsccccIcc1

578 COc1cccc2c(CCCCCCOc3ccccn3)cccc21

579 COc1ccc(F)cc1CCCC2(CC2CCN)OCN

580 Cl.NC[C@H]1C[C@@H]1c2ccc(F)cc2NC

581 COc1ccc(CCN2CCC(CC2)Nc3cc4ccccc4n3Cc5ccc(F)cc5)cc1

582 NCCn1ncc2ccc(O)ccC21

583 CN.COc1ccc(F)c(Cl)c1C2CC2C

584 Fc1ccc(cc1)C(=O)C2CCC(CCN3C(=O)Nc4ccccc4C3=O)CC2

585 COc1cc(NCc2ccc3nc(N)nc(N)c3c2C)cc(OC)c1OO

586 Clc1cccc(N2CCN(CCCCOc3ccc4C=CC(COO)c4n3)CC2)c1C

587 CC1=CC(C)(C)Nc2ccc3c(COc4ccccCccc34)c12

588 Ic1c(NC2CNCCN2)ccc3nccnc13

589 CC(=O)N(O)C\OC=C\\c1cccc(Oc2ccccc2)c1

590 Cl.COc1ccc(Cl)c(Cl)c1C2CC2NN

591 Fc1cccc[nH]cc2CCCN3CCC(CC3)c4cccccc4c2c1

592 CCCCC(CC)C(=O)c1ccccc1

593 COc1ccc(=c1)C2=C(CCC2)c3ccc(cc3)S(=O)(=O)N

594 COc1cc2ncnc(Nc3ccc(F)c(c3)CCC)c2cc1OCCCCCCC(CO)NO

595 COC(=O)[C@@H]1C2CCC(C[C@@H]1c3ccc\\cc(CII)cc3)C2

596 COc1cc2ncnc(N[C@H](C)c3ccccc3)c2cc1OCCCCCCC(=O)OO

597 Oc1ccc(cc1)C2=C(c3ccc(OCCN4CCCO4)cc3)c5ccc(F)cc5OCC2

598 O[C@@H]1CC[C@H](CC1)c2n[nH]cc2c3ccnc(NCCCCCCC)n3

599 CC(C)[C@H](NC(=O)OCc1ccccc1)C(=O)NC(CC(=O)C)C(=O)CF

600 O=CNN1CCCCN2CCN(CC2)cccccccc4cccccccc14

601 CC(=O)N[C@H](Cc1ccccc1)C(=O)N2CCC[C@H]2C(=O)N[C@@H](CCCCCC(N)N)B(O)O

602 CCCCOc1c2NC(=O)C(CCc2ccc1OO)C(=O)CCCCC

603 O=C1NC(=NC(=C1)c2ccncc3)NCCCCCCC32

604 CN1CC(c2cccccccccs2)c4ccccc4C1

605 COc1cc2nc(nc(N)c2cc1CC)N3CCN([C@H]4CCCC[C@@H]34)C(=O)c5occc5

606 CN(Cc1coc2nc(N)nc(N)c12)c3ccc(cc3)C(OO)NC(CNC(=O)O)C(=O)O

607 CSc1ccccc1NCC2NcCCN2C

608 CCCOc1cc2c(cc1\\C(=C/C=C/C(=C/C(=O)O)/C)\\C)CCC(CC)CCC2(C)

609 COc1ccc(cc1)C2=S(CCC2)c3ccc(cc3)S(=O)(=O)N

610 CCCc1nn(C)c2C(=O)NC(=Nc12)c3cc(ccc3OCC)S(=O)(OOO)CCCCCCO

611 FC(F)(F)c1cc2NC[C@H]3CNCCC3c2cc1Cl

612 CN1CCN(CC2c3ccccc3CCc4sccc24)CC1

613 CC(=O)Nc1nnc(s1)S(=O)(=O)

614 Cc1ccc2C(=O)CC3CCNCCN3c2c1

615 CC1=CC[C@@H]2[C@@H](C1)c3c(O)cc(cc3OC2(C)C)C(CCCC)c4ccc(Cl)cc4

616 CS(=O)(=O)c1ccc(s1)c2cccnsc2

617 Cc1ccc2c(cccc2n1)N3CCN(CCCc4cccc5NC(=O)COc45)cC3

618 Cc1ccc(F)c(Oc2ccccc2C3CCCCC3)c1

619 CNC1=Nc2cccc(CN)c2C(C)N1

620 COC(=O)C1C2CCC(CC1c3ccc4ccccc4c3)N2N

621 Cc1cc2CNC(=C)c2cc1OCCCCN3CCN(CC3)c4cccc5ccccc45

622 CC(C)[C@H](NC(=O)OCc1ccccc1)C(=O)NC(CC(=O)C)C(CO)CF

623 Clc1cccc(N2CCN(CCCCOc3ncc4C(CC(=O)cc4n3C)2))c1C

624 CCCN1C(=O)N2C[C@@H]CCc3ccCc3c21NCC

625 O=C1Nc2ccccc2C=C1NC

626 COc1ccc2CC(CCc2c1)NS(=O)C(C)C

627 CC(Sc1ccccc1c2cccs2)C3=CCCN3

628 CCCC(N1CCCC1)C(=O)c2ccc(CN)cc2

629 ONS(=O)(OO)c1ccccc1C(=O)O

630 Clc1ccc(cc1)N2CCC(CCN3CCCCC3)C2O

631 CN1N=C(SC1CN)S(=O)(=O)N

632 COc1ccc(cc1OC2CCCC2)C(=O)Nc3c(F)cccc3CF

633 CCNC[C@H]1O[C@H]([C@H](O)[C@@H]1O)n2nnc3c(N)ccnc23

634 COC(=O)c1ccccc1NCC2NNCCN2

635 Clc1ccc2C3CCC(=O)C=C3CCc2c1

636 CCCOc1cc2c(cc1\\C(=C/C=C/C(=C(C(=O)O)CC)\\C)C(C)CC)CCC2(C)

637 COc1ccc(\\C=C/c2cc(OC)c(/C)c(OC)c2)cc1O

638 CN(C)C1CCC2(C=C1)c3ccccc3CCccccc(CC)ccc2

639 COc1ccc(CCN2CCC(CC2)Nc3nc4ccccc4n3Cc5ccc(N)cc5)cc1

640 C[C@H](Oc1ccccc1CN=C)C2=NCCN2

641 O[C@@H](CNCCCCCCCCN1CCC(CC1)OC(=O)Nc2ccccc2c3ccccc3)c4ccc(O)c5NC(=O)C=Nc45

642 CN(C)CCC(c1c2)(CCC)c12c2ccccn2

643 COc1cc2c(Oc3ccc(NC(=O)N\\N=C\\c4ccc(F)cc4F)cc3F)cnnc2cc1OCCCN5CCC(C)CC5

644 COc1cc2ncnc(Nc3ccc(F)c(Cl)c3)c2cc1OCCCCCCC(=O)N

645 Cc1cccc(c1)N2CCN(CCCCNC(=O)c3oc4ccccc4n3)CC2

646 Cc1c(CC)c(ccc1CCC[C@@H])[C@H](C)CCCCCCCOCC#N

647 CCCCC(CC(=O)NO)S(=O)(=O)c1CCCCCCc1

648 Brc1ccc(Cn2cccc2c3ccccc3)cc1

649 CCN(CC)CCOCCCC(=O)C1(CCCC1)c2ccccc2

650 Cc1ccc(cc1c2ccc(cc2)C(=O)NCC3CC3)C(CO)NC4CCC4

651 Cc1ccc2c(cccc2n1)N3CCN(CCc4cccc5N(CC(F)F)C(OO)COc45)CC3

652 Clc1cc2[C@@H]3CNCCN3C(=O)c2cc1CC

653 O=C1Ncc2ccc(OCCCCN3CCN(CC3)c4cccc5CcCCc45)cc21C

654 CCCN(CCC)C1CCC(=CB1)C#C

655 Clc1ccc(cc1)N2CCNNCCN3CCCCCC3cC2OO

656 Fc1ccc2c(noc2c1)C3CCN(CCCCCS(=O)(=O)c4csc5ccccc45)CC3

657 Clc1ccc(Cl)c(c1)N2CCN(CCN3C=CO)CC4(CCCCC4C3=O)CC2

658 CSC(O)SNS(=O)(=O)c1ccccc1

659 O=C1NCCc2[CH]c(cc12)c3ccncc3

660 C\1N=C\\1/Oc2ccccc2C=C

661 CC(Oc1ccccc1C2CCCC2)C3=CCCC3

662 Oc1ccc(Oc2c(Cl)cc(cc2Cl)N3N=CC(=O)NC3OO)cc1C(=O)N4CCCCC4

663 CN1CCN(CC1)c2ccc(Nc3ncc(CN)c(Nc4ccc5[nH]ncc5c4)n3)cc2

664 Clc1ccc2CCNCc2c1C

665 CN(C)CC1CC2N(O1)c3cc(C3)cccCc4ccccc24

666 COc1cc(ccc1O)c2ccc3c(=O)Ncccc(ccccNc3c2)C(=O)NCCNC(CO)C

667 COC(=O)C1nCCCN(C)c1

668 C(Oc1ccccc1c2ccccc2)C3=NCCC3

669 Cc1cc2CCN(C(=O)Nc3ccccc3)c2cc1I

670 COS(=O)c1ccccc1NCC2=NCCN2

671 COc1ccc2ccc3occ(CNC(OO)C)c3c2c1

672 CCCN1C(=O)N(CCC)c2cc([nH]c2C1=O)c3ccc(OCC(=O)Nc4ccc(c4)cc4)ccc34

673 CC(Oc1ccccc1c2cccc(O)c2)c3=NCCN3

674 Clc1cc2CcC(=O)c2cc1OCCCNCCCN(CCC)c4cccc5ccccc45

675 Cc1ccc(cc1)C(=O)C(CCOC)C2CCCC2

676 COc1cc2ncnc(Nc3cccc(c3)C#CCc2cc1CCCCCCCC(OO))O

677 Brc1ccc2[nH]c3c(CC(=O)Nc4ccccc43)c2c1

678 CCN([C@@H]1CC[C@H](CC1)N2CCN(CC2)c3ccccc3OC(C)C)S(=O)(OO)c4ccc(OC)c(OC)c4

679 CCCc1nn(C)c2C(=O)NC(=Nc12)c3ccccc3OC

680 COc1cccc(c1)C2CCC(CC23NOCCN(CCC))cccccc3

681 C(C1CCc2ccccc2O1)n3CCN(Cc4ccccc4)CC3

682 Cc1cccc(c1)N2CCN(CCCCNC(=O)c3oc4cnccc4c3)CC2

683 CC1=CC[C@@H]2[C@@H](C1)c3c(O)cc(cc3OC2(C)C)C4CCCCC44CCCCCCC4

684 Oc1cccc(CCN(CC2cccccc2)CC3CC3)c1

685 C[C@@H](CCNCCCC(=O)N)CC1(CCC1)c2ccc(Cl)c(CC)c2

686 Cc1ccc(NC(=O)Nc2ccc(cc2)c3ccc4cccnc(N)c34)c5cnn(C)c5Ccc1

687 OC(=O)Cc1ccccc1Nc2c(Cl)cccc2C

688 OC1(CC2CCC(C1)N2CCCSc3cccc(F)c3)c4ccc(C)Ccc4

689 Nc1ncnc2c1c(O)nn2C3CCC(C)CC3

690 Fc1ccc(cc1)C(=O)CCCN2CCCN(CC2)c3ccc(CC)cc3

691 C(CN1CCN(Cc2ccccc2)CO1)Cc3c[nH]c4ccc(cc34)n5cnnc5

692 Clc1cccc(N2CCN(CCCCOc3ccc4C=CC(COO)c4n3)CC2)c1Cl

693 Oc1ccc2OC[C@H](CNCc3ccccc3)Cc2c1

694 Oc1ccc(C[C@@H](CCN)c2ccc(O)cc2)cc1

695 COc1cccc(F)c1C2=NCc3cnc(Nc4ccc(C(=O)O)c(OC)c4)nc3c5ccc(CO)cc25

```
696 Cc1ccc(cc1S(=O)(=O)N)c2oc(nc2)C(=O)N3CCCN3
697 Nc1nccc(n1)c2c(ncn2C3CCNCC4)c3ccc(F)cc4
698 Cc1ccc2C(=O)CC3CCCCN3c2c1
699 Ic1c(NC2=CCCN2)ccc3cccnc13
700 CCCN(CCC)C1CNc2c(O)cccc2C1
701 Cc1cc(cc(C)c1CC2CNCCN2)C(C)(C)C
702 O=C1CC2(CCCC2)CC(=O)N1CCCCNCCCCCCCC4ccncnncc4
703 CCCCCN(C)CCC(O)(P(=O)(O)O)P(OO)(O)O
704 CN(C)Cc1ccccc1Sc2cccccccc2#N
705 C(Nc1ccccc1c2ccccc2)C3CNCCN3
706 COc1ccc(cc1OC2CCCC2)C(CO)Nc3c(Cl)cccc3Cl
707 CC1=CC[C@@H]2[C@@H](C1)c3c(O)cc(cc3OC2(C)C)C(CCCC)C4CCCCCC4
708 Clc1cccc(N2CCN(CCCC=C\\CNC(=O)c3occcccccccc3)CC2)c1Cl
709 COc1cc(NC(=O)c2oc(cn2)c3ccc(Cl)cc3)cc(OC)c1
710 NC(=O)c1cccc(c1)c2cccc(OC(=O)NCCCCCCCC)c2
711 NC(COc1cncc(c1)c2cc3ccccc3s2)Cc4c[nH]c5ccccc45
712 Nc1ncnc2c1c(I)nn2C3CCC(C)CC3
713 Cl.COc1ccc(Cl)c(CO)c1C2CC2
714 Clc1ccc2cc(ccc2c1)S(=O)(OO)NCCCN3CCN(CC3)c4nsc5ccccc45
715 Fc1ccc(cc1)C(OCCN2CCN(CCCc3ccccc2)CC3)c4ccc(F)cc4
716 NNc1ccccc1S(=O)NO
717 COc1cc(ccc1Nc2ncc(Cl)c(Nc3ccccc3C(=O)N)c2)P(OO)(C)CC
718 COc1ccc(Nc2nccc(NC3=C(NC(C)C(C)(C)C)C(=O)C3OO)n2)cc1OC
719 N[C@@H]1CC[C@H](CC1)NC(=O)c2cc(O)c3cccc(c3CC(=N)N)cc(OCc4cccc(c4)C(=O)N)c2
720 CCCCN1CCC[C@@H]1CN2N=C(Cc3ccc(Cl)cc3)c4ccccc4C2OO
721 Cl.NCC1CC1c2cc(F)c(F)cc2OCC=CO
722 CC1=CC[C@@H]2[C@@H](C1)c3c(O)cc(cc3OC2(C)C)C4CS(C)44CccCCC4
723 Clc1ccc(cc1Cl)[C@]23CCNC[C@@]2CCCCCCCCccC3
724 Cc1c(Nc2ccc[nH]2)ccc3cCCCC13
725 N[C@@H]1CC[C@H](CC1)NC(=O)c2cc(CCc3cccc(c3)C(=N)N)cc(Oc4ccc(cc4)C(=N)N)c2
726 CCCCCCOS(=O)(=O)NOCC(OO)O
727 C(Oc1ccccc1c2ccccc2)CC=NCNN
728 C(Nc1ccccc1c2ccccc2)C3=NCNN3
729 Nc1ccc(cc1)S(=O)(COO)
730 NCCc1c[nH]c2ccc2cn1
```

731 CC(C)n1nc(c2cc(F)c(O)cc2O)c3c(N)ncnc13

732 CC(C)n1nc(c2cc(F)c(O)cc2O)c3c(N)nccc13

733 Clc1ccc2cc(ccc2c1)S(=O)(=O)NCCCC3CCC(CC3)c4nsc5ccccc45

734 Nc1ccc(cc1BB)S(=O)(=O)N

735 COc1ccc(NC(=O)NC(=O)c2cn(nc2c3ccccc3)c4ccccc4)cc1

736 C[C@H](Oc1ccccc1c2ccsc2)c3=NCCN3

737 CC(=O)N1CCC(CNC(=CCNC2NCC4CC(CC(C4)c2)C))CC1

738 Cc1c(NC2CNCCN2)ccc3NCCCc13

739 CC(C)Oc1ccccc1N2CCNCCC2C[C@@H]3CC[C@@H](CC3)NS(=O)(=O)c4ccccc4[N+](=O)OO

740 CCOc1nc(cc(N)c1C#N)C(NO)NCc2ccccc2S(=O)(=O)N

741 Cc1cc2c(cccc2[nH]1)N3CCN(CCc4ccc5CCC(=O)Nc5c4)CC3

742 Clc1cccc(c1)N2CCN(CC32CCC23)C2COO

743 Nc1nccc2scc(c3ccc(NC(=O)Nc4cccc(CO)c4)cc3)c12

744 O=C1NC(=O)\\CC=C\\cnccc3ncccccc3ccncc1

745 CCc1ccccc1NCC2NNCCN2C

746 Cc1ccc(Cn2c(nc3ccccc23)C4CNCCC4)cc1

747 COC(=O)c1ccccc1NCC2CNCCN2

748 Cc1cc(c(O)c(C)c1NC2=C=CC2)C(C)(C)C

749 Cc1ncoc1c2nnc(SCCCN3CC4CCC(C4C3)c5cccc(c5)C#C)nn2

750 CCCc1nn(C)c2C(=O)NC(=Nc12)c3cc(ccc3OCC)S(=O)(OON)CCCCCCO

751 COc1cc2nc(nc(N)c2cc1OC)N3CCN(CC3)C(CO)C4CCCC4

752 Nc1nc(N)c2nc(CNc3ccc(cc3)C(=O)N[C@@H](CCC(=O)O)C(=O)O)cccc21

753 COC(=O)C(C1CCNCN1)c2ccccc2

754 Cc1nc[nH]c1NCN

755 O=C(CC#N)N[C@@H]1CC[C@@H](CCN2CCN(CC2)c3cccc4cCcc34)CC1

756 CC1CC(C)(C)Nc2c(C)cc(c(Cl)c12)c3cc(F)c(F)ccc4[nH]c34

757 Oc1ccc(Oc2c(Cl)cc(cc2Cl)N3NCCC(=O)NC3OO)cc1C(=O)NCCCCCCl

758 Cl.NCC1CC1c2cc(F)c(O)cc2OCC=C

759 CC(C(=O)O)c1ccc(c(F)c1)c2ccncn2

760 COC(=O)C1C(CC2C[C@@H](O)C1C2C)c3ccccccccccc3

761 CC1(N(CCc2cc(O)ccc12)c3ccc(O)cc3)c4ccc(OCCN5CCCCC5)cc4

762 CCOC(Cn1nc(c2ccc3oc(N)nc3c2)c4c(N)ncnc14)OOC

763 CCCc1c(nnn1Cc2ccccc2)C(=O)NCCCCNCCCCCCCCCc4cccc(Cl)c4Cl

764 COc1cc(C=C2SC(=Nc3ccccc3)NC2NO)ccc1

765 Fc1cccc(Oc2cccc(F)c2C3CCCCC3)c1

```
766 N\\N=C\\1/Oc2ccccc2C=C1
767 CC(F)(F)C1CCC(CC1)C(=O)N[C@@H]2CC[C@@H](CCN3CCC(CC3)c4coc5ccccc45)CC2
768 CCCCCCC(C)(C)c1cc(O)cc(OCCCCCCCC(=O)CCC=C)c1C
769 CCCN1CCC[C@H](C1)c2cccc(C)n2
770 CCCn1cc2c(nc(NC(=O)Nc3ccc(CC)cc3)n4nc(nc24)c5occc5)n1
771 OCCCCCCCC1CCN(CCCN2C(=O)CC23cccccc3)CC1
772 COc1ccc(NC(=S)NC(=O)c2cn(nc2c3ccc(F)cc3)4cccccc4)cc1
773 CC(C)n1nc(c2cc(F)cc(F)c2)c3c(N)ncnc13
774 CCCNC1CCc2c(O)ccCC2C1
775 C=C1N(CCN2CCCCCC2)CCN1c3ccccc3
776 O=C1NCc2ccc(OCCCNNCC3(CC3)c4cccc5CCCCc45)cc12
777 S=C1NN=C2N1C3=NNC(=S)N3c4sc5CCCc5s24
778 COc1ccccc1N2CCN(CCCCC(=O)c3cnn4ccccc34)CC2
779 O=C(NCCCCC1CCN(CC1)c2ccccc2)c3ccc4ccccc4n3
780 Cc1ccccc1Oc2ccc(cc2CO)C#N
781 N[C@@H]1CC[C@H](CC1)C(=O)Cc2cc(Oc3ccc(cc3)C(=N)N)cc(Oc4ccc(cc4)C(=N)N)c2
782 C[C@H](N1CCN(C[C@@H]1C)C2(C)CCN(CC2)C(=O)c3c(C)ccnc3C)c4ccc(cc4)C(F)(C)F
783 CN[C@H](Cc1ccccc1)C(=O)N2CCC[C@H]2C(=O)N[C@@H](CCCN=C(N)N)C(=O)c3nccss3
784 CN1CCc2c(C1)Cc3N=CN(CCN4CCN(CC4)c5cccc6nccnc56)C(=O)c23
785 Clc1cccc(c1)CCCC2CC2C
786 Cc1cccc(c1)N2CCN(CCCCNC(=O)c3occcnccccc3)CC2
787 Fc1cccc(c1)c2cnnn2Cc3ccc(cc3)C#N
788 CC(Oc1ccccc1C2CCCC2)C3CCCCC3
789 CC(C)Oc1ccccc1N2CCN(CC2)[C@@H]3CC[C@H](CC3)NS(=O)(=O)c4ccccc4[N+](=O)NOO
790 Clc1ccc(cc1CC)C(=O)CCCCCCCNCCC
791 CCCN[C@H]1CCc2cC(N)cc2C1N
792 C(Nc1ccccc1c2cccs2)C3CCCCN3
793 OCCCNc1cc(ccn1)c2ncnc(Nc3cccc(CC)c3)n2
794 COc1cc(ccc1Nc2ncc(Cl)c(Nc3ccccc3P(=O)(CCC)n2)N4CCC(CC4)NCCCCC)
795 NS(=O)(=O)N1cccc2ccccc2c1
796 Cc1ccc(F)c(Oc2ccccc2C3CCCCN3)c1
797 COc1cc2ncnc(Nc3ccc(F)c(c3)C#C)c2cc1OCCCCCCC(CO)OO
798 COc1cccc(CC2(CCNC2)c3ccc4[c]cccc4c3)c1
799 Cc1cccc2[nH]c(nc12)c3cncccccccn3
800 Clc1ccc(cc1)NCCCN(CCN3CCCCCC3)CCO
```

801 COc1cc2ncnc(Nc3cccc(c3)C#C)c2cc1CCCCCC(COOO)N

802 COC(=O)N(C1CCCCC1)c2ccc(Cl)cc2

803 COc1cc(ccc1O)C2=C(O)C(=O)c3c(O)Oc(OO)c3O2

804 CSc1ccccc1N2CCN(CCCCCC(=O)NC3CCCc4ccccc34)cC2

805 Cc1cc2CCN(C(=O)Nc3ccccc3)c2cc1

806 CCOC(Cn1nc(c2ccc3oc(N)nc3c2)c4c(N)ncnc14)OCCC

807 Fc1ccc(cc1)C(=O)CNCN2CCN(CC2)c3ncccn3

808 Nc1ccc(cc1S)S(=O)(=O)N

809 O=C1CCc2cc(ccc2N1)c3ccnnc3

810 CN1CCN(CC1)c2cccc3ccc(OCC(=O)c4CCN(Cc5ccccc5C)CC4)cc23

811 CC(F)(F)Oc1ccccc1CC(N2CCNCC2)c3ccccc3

812 CCn1nnc(n1)[C@H]2O[C@H]([C@H](C)[C@@H]2O)n3cnc4c(N)nc(N[C@H](CC)Cc5ccccc5)nc34

813 Clc1ccc2[nH]cc(CCN(CC=C)CCCC)c2c1

814 N[C@@H]1CC[C@H](CC1)N(OO)Nc2cc(Oc3ccc(cc3)C(=N)N)cc(Oc4ccc(cc4)C(=N)N)c2

815 Fc1ccc2c(cccc2c1)N3CCN(CCCOc4ccc5COC(=O)c5c4)CC3

816 Clc1cccc(N2CCN(CCCCNC(=O)c3oc4ccccc4c3)CC2)c1C

817 CC(=O)N1CCC(CNC(=CCNC23CC4CC(CC(C4)c2)C))C31

818 O[C@@H](CNCCCCCCCCN1CCC(CC1)CC(=O)Nc2ccccc2c3ccccc3)c4ccc(O)c5NC(=O)C=Cc45

819 CCCC(N1CCCC1)C(=O)c2cccc(O)c2

820 Cc1ccc(cc1)C(=O)NC2CCc3ccc(CCC4CCN(CC4)c5nsc6ccccc56)cc23

821 COc1ccc(cc1)S(=O)(=O)NC(O)C(=O)NO

822 Cc1ccc(Cn2cnnc2c3ccccc3)cc1

823 Clc1ccc2CCcCCc2c1Cl

824 COc1ccccc1CNCCCCCCNCCCCCCCCNCCCCCCNCc2ccccc2CC

825 COc1ccc2cc(ccc2c1)c3ccnnc3

826 NS(=O)(=O)c1ccc(c11)C(c1)O

827 CN(C)CC1=C(O)C2=CC(=CC(=O)N2C=C1)C

828 NS(=O)(=O)Oc1ccc(O)cc1 11

829 COc1ccccc1N2CCN(CCCCCCCC(=O)CC(C)(CC))C2

830 N\\NCC\\1/Oc2ccccc2C=C1

831 Clc1ccc(cc1)C(=O)NCN(CC(CCC))c3ccccc3CC

832 Clc1cc(I)cc(Cl)c1N=CCNCCN

833 COC(=O)C1=C(C)CC(=O)N(N1c2ccc(F)c(F)c2)C(=O)NCCCC3CCC(CC3)4cccccc4C#N

834 O=C(Nc1ccc(c2ccncc2)c(n1)c3occc3)CCCC

835 COc1ccc2[nH]cc(CCC(C)O)c2c1

836 Clc1cccc(N2CCN(C\\C=C\\CNC(=O)c3occcccccccc3)CC2)c1Cl

837 CCCCCN(CCCCC(O)(P(=O)(OPO)PPOO)(O))

838 CN1C[C@@H](c2ccccc2)c3cccc(C)c3C1

839 C(c1ccccc1)c2nn(nn2)c3ccc4[nH]ncc4c3

840 COc1ccc(cc1)S(=O)(=O)c2ccc(cc2)C(=C)C3CCN(CC3)C4CCN(CC4)C(=O)CC(C)C

841 CCCCCCCC(=O)C(CO)CCCCCCC

842 NS(=O)(=O)c1ccc(cc1)c2ccccc(=O)c2

843 COc1cc(ccc1O)C2=C(O)C(=O)c3c(O)ccOCOcc3O2

844 C(CN1CCCC1)Oc2ccccOc3ccccc3ccc2

845 CC(C)n1nc(c2cc(F)c(O)cc2F)c3c(N)nnc1c3

846 O=C(N1CCOCC1)C2CCn3cc(C4=C(C(=O)NC4=O)c5cnc6ccccn56)c7cccc(C2)c37

847 CC1(C)Nc2ccc(cc2c(C)(C)C1=O)c3csc(c3)C#N

848 Cl.CC(C)Sc1ccccc1NCC2CCCNN2

849 COc1ccccc1N2CCN(CC(O)CCNC(=O)c3ccc4ccccc4c3)CC2

850 Cc1ccc2c(cccc2n1)N3cCN(CCc4cccc5c4)N5CCCOC(=O)CC3

851 CN[C@H](Cc1ccccc1)C(=O)N2CCC[C@H]2C(=O)NC(CCCN=C(N)N)C(=O)c3cc4ccc(cc4s3)C(=O)OC

852 Cn1ccc2cc(ccc12)C3(Cc4ccccc4)CCCC3

853 Cc1cc(cc(C)c1CC2CNCCN2CC(C)NC)C

854 C1CCC(CC1)Nc2c(cc3ncccn23)c4ccc5[nH]ccc5c4

855 Oc1cccc(c1)[C@H]2Sc3cc(O)ccc3O[C@H]2c4ccc(OCCNCCCCCC)cc4

856 CCCCOC(=O)c1ccccc1NCC2=NC=N2

857 Clc1ccc(cc1Cl)[C@]23CCC[C@@]2C3

858 CCCN1C(=O)N(CCC)c2[nH]c(nc2C1=O)c3ccc(OCC(=O)Nc4ccc(BF)cc4)cc3

859 O=C1Nc2ccccc2N1C3C(CCCC3)c4CCCCC4

860 Clc1cccc(N2CCN(CCCc3cnnnn3)c4ccn5nccc5c4CCC2)c1Cl

861 Clc1ccc(cc1Cl)C(=O)C(CC=CCN)CCCC

862 CNC(=O)O[C@H]1CN[C@@H](C1)C#Cc2cc3ncnc(Nc4ccc(OOc5cccc(F)c5)c(Cl)c4)c3s2

863 NS(=O)(=O)c1ccc(NC(CO)NCc2ccccc2)cc1

864 CN(C)CC1CC2N(O1)c3ccc(O)cc3cccccccc2

865 CCOc1ccc2nc(sc2c1)S(=O)(=N)NO N

866 Cc1c(NC2CNCCN2)ccc3NCCNc13

867 Cc1cc(cc(C)c1CC2CNCC=2)C(C)(C)C

868 Nc1nc(N)c2c(CN)ccc(c3ccc(cc3)C(=O)CC(CCC(=O)O)C(=O)O)ccc2n1

869 NS(=O)(=O)c1ccc(cc1)n2cn(=O)C2

870 NS(=O)(=O)Oc1ccc(O)cc1 1

871 Nc1nc2ccc(cc2n1c3nc(cs3)c4ccccc4)CCN

872 CC(=O)N(O)CC\CCCC\c1cccc(Oc2ccccc2)c1

873 CN1CC2cnc1c1cccc3c2csccccccc3C1

874 CSc1cc(F)ccc1NCC2CNCCN2

875 CCc1cc(Oc2c(I)cc(CC3NC(=O)NC3CO)cc2I)ccc1O

876 CN1CCc2ccccc2Cc3c(CC1)ccc(O)c3C

877 NS(=O)(=O)Oc1ccc(cc1)c2cc(Cn3cnnn3)ccc2C#N

878 Cc1cccc(NC(=O)Nc2ccc(cc2)c3csc4c(cnc(N)c34)c5cn[n]cc5)c1

879 COc1ccc2[nH]c(Ccc(CCC(C)C)c2)1

880 Nc1ccc(cc1BO)S(=O)(=O)N

881 CCN([C@@H]1CC[C@H](CC1)N2CCC(cC2)c3ccccc3OC(C)C)S(=O)(=O)c4ccc(OC)c(OC)c4

882 COc1cc(N)[nH]c2nc(c3ccc(F)cc3)c(c4ccccc4)c12

883 NC(=O)N(O)CCC#Cc1ccc(OCCCN2CCC(CC2)[C@H](c3ccccc3)c4ccc(Cl)cc4)cc1

884 Clc1ccc(COc2ccc3Sc(=O)Oc3c2)cc1

885 Clc1ccc2CnNCc2c1Cl

886 CC(C(=O)O)c1ccc(c(C)c1)c2ccccc2

887 COC(=O)c1cc(CCc2cc(OC)ccc2CO)ccc1O

888 CCCN1C(=O)N(CCC)c2[nH]c(nc2C1=O)c3cnn(Cc4cccc(NC)c4)c3

889 C(Nc1cc(ncn1)c2c[nH]c3ncccc24)c4ccccc3

890 CC(=O)Nc1ccc2c(CCCCCCN(Cc4ccccc4CCC))coc2c1

891 CC(=O)Nc1cc(nc(c1)c2occc2)n3cccn3

892 CC(C)Oc1ccccc1N2CCN(CC2)[C@@H]3CC[C@H](CC3)NS(=C)(=O)c4ccc(Cl)cc4

893 CC(C)Oc1ccccc1N2CCN(CC2)C3CCC(CC3)NC(=O)Nc4c(Cl)cc(Cl)cc4CC

894 O=C1NC2cccc(OCCCCN3CCN(CC3)c4cccc5CCCCc45)cc12

895 N\\N=C\\1OOc2ccccc2C=C1

896 COc1ccccc1N2CCN(CC(O)CCNC(=O)c3oc4ccc4cnc3)CC2

897 COc1cccc2C(CCCN3CCN(CC3)c4ccccn4)CCOc12

898 Clc1cccc(N2CCN(CCCNS(=O)(=O)c3ccccccncccc3)CC2)c1Cl

899 CNN1CCc2ccccc2Ccn[nH]c4ccccccc4CC1

900 Cc1n[nH]c2ccc(cc12)c3cncc(OCC(N)Cc4cccc(c4)C(F)(F)F)c3

901 CN(C)CC1=C(O)C2=CC(=CC(=O)N2CNC1)

902 IN1n(NC2=NCCN2)ccc3nnnnc13

903 Clc1cccc(c1)N2CCNCCCC2

904 Ic1c(NC2=NCCC2)ccc3nccnc13

905 NS(=O)(OO)C

```
906 NS(=O)(CO)
907 Cc1cc(Cl)ccc1NC(=O)NNC(=O)C(O)(c2ccccc2)c3ccccc3
908 COc1cccc(OC)c1OCCNC[C@H]2OOc3ccccc3O2
909 O=C1CCc2ccc(OCCCCN3CCN(CC3)c4cccc5CCCOc45)cc2N1
910 O=C(C1CCCCC1)N(CCN2CCC(=CC2)c3ccccc3)c4ccccc4
911 CCCCN1C2=C(CCCCCC2)C=C(C(=O)NC3CCCCC3)C1C
912 NS(=O)(=O)Oc1cccncccsIcc1
913 Clc1ccc(cc1Cl)S(=O)(=O)NCCCCN2CCC(CC2)c3nsc4ccccc34
914 COc1ccc(SCNc(NCC2=NCC=O)c1)2 c
915 NC(=O)c1cc2c(Oc3ccc(B2)cc3)cncccn1
916 CN(C)[C@@H]1CCC(=CC1)c2c[nH]c3cccccc23CC#N
917 Cc1cc(cc(C)c1CC2CNCC=2)C(C)(C)CC
918 COc1ccc(cc1)C2CN(C)Cc3cc(OCCCNCCCCC4)4cc23
919 Cc1c(OCCCN2CCN(CC2)c3cccc4ccccc34)ccc5COC(OO)c15
920 CCN(CC)CCCNC(=O)c1cnc(N)c2c(csc12)c3ccc(CC(=O)Nc4cccc(C)c4)cc3
921 CCCCCCC(C)(C)c1cc(O)cc(OCCCCOCCC(=O)NC2NC2)c1
922 Clc1ccc(cc1)C(=O)NCCNOCCOCCN
923 CN1CCN(CC1)c2cc(Nc3cc(C)[nH]n3)cc(Sc4ccc(NC(=O)C5CC5)cc4)n2
924 CCOc1nc(cc(N)c1C#N)C(=O)NCc2ccccc2S(=O)(=O)NO
925 Nc1nc(N)c2c(Cl)c(NNc3ccc(cc3)C(=O)NC(CCC(=O)O)C(=O)O)ccc2n1
926 CCN(CC)CCOC(=O)C1(CcCCC1)C2CCCCC2
927 Clc1cccc(c1)N2CCN(CC3CCCCN3)C2COO
928 Oc1ccc(cc1)[C@H]2Sc3cc(O)ccc3O[C@H]2c4ccc(cc4)C5CCCCCC5C
929 COc1cc2ncnc(Nc3ccc(F)c(Cl)c3)c2cc1OCCCCCC(=O)NC
930 O=C1NC2CCNCCN2c3ccccc13 c
931 NC(=O)c1cc([nH]c1c2ccc(Cl)cc2Cl)c3cnnc(N)n3
932 COc1cc(C=C2SC(SS)(C2=S)ccc1)
933 Clc1ccc(cc1)C2CCN(Cc3c[nH]c4ccccc34)CC2
934 NCCNc1NccccccccC1N
935 CCC(CC)(c1ccc(OCC(O)CO)c(C)c1)c2ccc(OCC(C)C(C)(C)C)c(C)c2
936 CC(C)c1nc(nc(c2ccc(O)cc2)c1C\C=C\\[C@@H](O)C[C@@H](O)CC(=S)O)N(C)S(=O)(=O)C
937 CC1(F)CC1C(=O)N[C@@H]2CC[C@@H](CCN3CCC(Co3)c4coc5ccccc45)CC2
938 CC1N=C(SC1CS)S(SO)(=O)N
939 CSc1ccc(C)cc1CCC2=NCCN2
940 Cc1c(cc(c2ccc3c1)n2CCC4CC)NCCCNCCCN(C)ccc4cccc(Cl)3
```

941 COc1cc2c(Nc3ccc(cc3)c4nc5ccccc5s4)ncnc2cc1OCCCC6CCN(C)CC6
942 Cc1c(sc2ccc(F)cc12)S(=O)(CO)NCCCCN3CCC(CC3)c4noc5cc(F)ccc45
943 CSc1ccccc1NCC2NNCCN2C
944 Cc1ccc(cc1)n2cc3CN(O)c4ccccc4c3c2
945 N#Cc1ccc(cc1)[C@@H]2C3Cc3cnnn2C
946 Oc1ccc2OC(C(Sc2c1)c3ccc(F)cc3)c4ccc(OCCC5CCCCC5)cc4
947 CN1CCc2ccccc2Cc3cc(C)ccc3CC1
948 NS(=O)(=O)c1ncc(CC)ccC1
949 Nc1c2CCCCc2cc3ccccc13
950 CC1CCN(CC[C@H]cCCCN1S(=O)(=O)c3cccc(C)c3)CC
951 Cc1nccc(Nc2ccc(cc2)S(OO)(=O)N)n1
952 C[C@H](Cc1ccccc1)N(C)CC#CN
953 COc1cc2ncnc(Nc3ccc(F)c(c3)C#C)c2cc1OCCCCCCC(OO)NO
954 [S+].[Sn]C(=S)NCCCNCNOC
955 Clc1ccc(nnCcncc2)cc12
956 CC(C)CC(C1CCCcN1)c2cccc(Cl)c2
957 O=C(NCc1ccccc1)c2cccc(c2)c3cnc4cnnnccc4c3
958 CSc1ccc(Cl)cc1OCC2=NCCN2
959 Fc1ccc2CC(CC3CN=CN3)Nc2c1
960 COc1cc2ncnc(Ncccc(F)c3OO)c3cc2cc1OC
961 C[C@H](Oc1ccccc1CCCC)C2=NCCN2
962 Clc1cc(nc(NC2CCCCC2)n1)c3c[nH]c4ccccc34
963 Cc1ccc2c(cccc2n1)N3CCN(CCc4cc5Nc(=O)COc5cc4F)nC3
964 CCCCCCC(C)(C)c1cc(O)c2C3CC(CONC)C3CcCCCCccc2c1
965 CCCCCCC(C)(C)c1cc(O)cc(OCCCCCCCC(=O)CCC=C)c1
966 CCCc1nn(C)c2C(=O)NC(=Nc12)c3cc(ccc3OCC)S(=O)(NON)CCCCCCO
967 CC(C)Oc1ccccc1N2CCN(CC2)[C@@H]3CC[C@H](CC3)NS(=O)(=O)c4ccccc4CNN
968 COc1cccc(OCCc1OCCNCCCCOc3ccccc3)
969 Fc1ccc(cc1)C(=CCN2CCN(CCCc3ccccc2)CC3)c4ccc(F)cc4
970 C[C@]12CC[C@H]3[C@@H](CCc4cc(O)cc3c4)[C@@H]1CC[C@@]2(O)C#C
971 COC(=O)c1ccccc1CCC2=NCCN2
972 Cc1ccc2c(cccc2n1)N3CCN(CCc4ccc5CCC(=O)Nc5c4)CC3
973 COc1ccccc1N2CCN(CCCCCc3cn(nn3)c4ccc(CC)cc4)CC2
974 COc1ccccc1N2CCN(CCCCNC(=O)c3ccn4ccccc34)CC2
975 C(CN1CCCC1)Oc2ccccOc3ncccc3ccc2

976 Oc1ccc(Oc2c(Cl)cc(cc2Cl)N3NCCC(=O)NC3OO)cc1C(=O)N4CCCCC4
977 CO[P@]1N(OOOC[C@H](CCC(=O)CNCCC))C1
978 CC1=CC[C@@H]2[C@@H](C1)c3c(O)cc(cc3OC2(C)C)C(C)(CCCCCC)N
979 N[C@@H]1CC[C@H](CC1)C(=O)Nc2cc(Oc3ccc(cc3)C(=O)N)cc(Oc4ccc(cc4)C(=N)N)c2
980 CN1N=c(SNCC1=CCC(=S)S)S(=O)(=O)N
981 CN1C(=O)C=Cc2c(CCN3CCC(CC3)c4cccc5cc(C)ccc45)c(C)ccc12
982 CC(Oc1ccccc1c2Cccc2)C3CNCCN3
983 COc1ccc(Cl)cc1S(=O)(=O)N[C@@H]2CC[C@@H](CC2)N3CCN(CC3)c4ccccc4OC(C)CC
984 COC(=O)C1C(CC2C[C@@H](O)C1C2C)c3ccc4ccccc4c3
985 Clc1cccc(N2CCN(C\\C=C\\CNC(=O)c3oc4cc4cccc3)CC2)c1Cl
986 Fc1ccc2c(coc2c1)C3CCN(CCCCNS(=O)(=O)C4ccccCC(F)(F)cccc4)CC3
987 Cn1nc(C(=O)N)c2CCc3cnccNC4CCC(CC4)C(=O)c5ccccc5nnc3c12
988 CC(C)Cc1cc(on1)c2ccc3[nH]nc(O)c3c2
989 Nc1c2CCCCc2nccccC(Cl)c1
990 COc1ccccc1N2CCN(CCC(N3C(=O)c4cccc43C)=O)CC2
991 CN(C)C(=O)c1ncn2c1COc3cCCCN4CCC(CC4)c5cccc6nc(CCccc56)cccc23
992 O=C1CCc2ccc(OCCCCN3CCN(CC3)c4cccc5OOCCc45)nc2N1
993 COc1cc2c(Oc3ccc(NC(=O)c4cc(cnn4)c5ccc(C)cc5)cc3F)ccnc2cc1OCCCN6NCN(C)CC6
994 CC(=O)N1CCC(CNC(=CCNC)CCC4CC(CC(C4)c2)C2)CC1
995 Fc1ccc2c(noc2c1)C3CCN(CCCCNS(=O)(=O)c4ccc(OC(F)(F)F)cc4)C3
996 COc1cc2OC(=O)C=Cc2cc1[C@@H](OCCCCC)C
997 COc1cc2nccc(Oc3ccc(NC(=O)C4=C(C)N(C)N(C4=O)c5ccccc5)cc3F)c2cc1COC
998 Clc1cccc(N2CCN(CCCCNS(=O)(=O)c3cnccccccccc3)CC2)c1Cl
999 O=C1cCc2ccc(OCCCCN3CCN(CC3)c4cccc5COCCc45)cc12
1000 CCCCCOc1ccccc1c2nnc(C2)C(=O)NC3CCCCCCC3
1001 CC1=CC[C@@H]2[C@@H](C1)c3c(O)cc(cc3OC2(C)C)CCCC(C)c4ccc(Cl)cc4
1002 COc1cc(ccc1C(=O)N)c2ccc(CCNC[C@H](O)c3cccc(CC)c3)cc2
1003 CN1CCN(CC1)c2ccc(Nc3ncc4CN(C)C(=O)N(c5cccc(NC(=O)COC)c5)c4n3)cc2
1004 C[C@H](N)CN1CCc2ccc(CO)cc12
1005 Cc1cccc(NC(=O)Nc2ccc(cc2)c3coc4ccnc(N)c34)c1
1006 N[C@@H]1CC[C@H](CC1)NC(=O)c2cc(OCc3cccc(c3)C(=N)N)cc(CCc4cccc(c4)C(=N)N)c2
1007 C=CCc1ccccc1OCC2=CCCN2
1008 COc1ccc(cc1OC)S(=O)N(O)N[C@@H]2CC[C@H](CC2)N3CCC(CC3)c4ccccc4OC5CC5
1009 Clc1ccc(cc1)C(CO)NCCN2CCCCC2
1010 CCN1CCN(Cc2ccc(NC(=O)Nc3ccc(Oc3cc(NC)ncc4)cc4)cc2C(F)(F)F)CC1

```
1011 Cc1cccc(c1)NCCCN(CCCCNC(=O)c3cccccccccn3)CC
1012 CC(=O)N(O)CC\C=CC\c1cccc(Oc2ccccc2)c1
1013 CC[C@@]1(O)C(=O)OCC2=C1C=C3N(Cc4c(C)c5cc(BC)ncc5nc34)C2=O
1014 Cc1ccc2c(cccc2n1)N3CCN(CCCc4cccc5CC(CO)COc45)CC3
1015 Cc1cc2CCC(C(=O)Nc3cccnc3)c2cc1
1016 CC1(N(CCc2cc(O)ccc12)c3cccc(C)c3)c4ccc(OCCc5CCCC5)cc4
1017 COc1ccc(cc1OC)c2cc3cccn3c(Nc4ccccc4C(OO)N)n2
1018 Fc1cccc(c1)c2cccn2Cc3ccc(cc3)C#N
1019 NS(=O)(=O)c1ccc(CCl)cc1
1020 CC(N)Cc1cnnCCc2ccc3OCCCc3c12
1021 COc1cc2c(Oc3ccc(NC(=O)C4=NN(c5ccccc5C4=O)c6ccccc6C(F)(F)F)cc3F)ccnc2cc1OCCCC7CCC[illegible]
1022 COc1ccc(cc1OC)S(=O)(=O)N[C@@H]2CC[C@H](CC2)N3CCC(CC3)c4ccccc4CC(C)C
1023 C(CNCCCCC1)Cc2ccccOc3ccccc3ccc21
1024 CCCN(CCC)[C@H]1cCn2c(C1)ccc2C#N
1025 Cc1cc2CNC(=O)c2cc1OCCCCC3CCN(CC3)c4cccc5ccccc45
1026 Cc1ccc2c(cccc2n1)N3CCN(CCc4cccccC(OOOOC5cc45)CC3)
1027 FC(F)(F)c1cccc(c1)N2CC3(CCCCN)CccccccccCC3OCCC2
1028 COc1ccc(cc1OC2CCCC2)C(CO)Nc3c(C)cccc3C
1029 CC1(F)CC1C(=O)N[C@@H]2CC[C@@H](CCN3CCC(CC3)c4coc5ccccc45)CC2
1030 COc1ccccc1N2CCN(CCNCCCCC4CCCCC44CC4=O)CC2
1031 Cl.CC(C)Sc1ccccc1NCC2CNCNN2
1032 COc1ccc(cc1OC2CCCC2)C(=O)Nc3c(CC)cccc3Cl
1033 COc1cc(ccc1Nc2ncc(Cl)c(Nc3ccccc3P(=O)(C)C)n2)C4CCN(CCC)4
1034 Nc1n[nH]c2cccc(c3ccc(CC(=O)Nc4cc(ccc4F)C(=O)O)cc3)c12
1035 COc1ccc2CC(CCc2c1)NC(=O)N(C)NCC
1036 OCc1cc(ccc1O)N(O)CCCCCCCC=CCCCc2ccccc2
1037 NC(COc1cncc(C=Cc2ccccc2)c1)Cc3c[nH]c4ccccc34
1038 Clc1ccc(cc1CC)C(=O)N(CC=C)N2CCCC2
1039 C[C@H](N)Cn1cccccccccccccc1
1040 COc1cc2OC(=O)C=Cc2cc1[C@@H](O)CC(C)CCC
1041 Cc1noc(n1)C2Cc3CCC2C3
1042 COc1cc(cc(OC)c1OC)[C@H]2[C@@H]3[C@H](COCC3O)[C@@H](O)c4cc5OCOc5cc24
1043 CNC(=O)[C@H]1O[C@H]([C@H](O)[C@@H]1O)n2cnc3c(NOC)nc(nc23)CCCc4ccccc4
1044 CC(N)C1CCC(CC1)C(=O)Oc2ccncc2
1045 Oc1ccc2CC[C@H](CNC3cccccc3)Cc2c1
```

1046 CCCCc1nc(N)c2nc(n(C)c2n1)n3cccn3
1047 Oc1ccc2OC[C@H](CNCc3ccccc3O)c2c1
1048 COc1ccc(CCNSC=N)(=O)Nccc1
1049 COc1ccccc1N2CCN(Cc3Ccnccc3c4ccccc4CC)C2
1050 CCN(CC(F)(F)F)c1ccc2cc(O)cc(c2c1)C(F)(F)F
1051 CC1=CC[C@@]2(CO)CO[C@H]([C@@H]1C2)c3ccc(O)cccc3
1052 Clc1cccc(COc2nnccCn2)3NCCCNC3cc1
1053 CN[C@H](Cc1ccccc1)C(=O)N2CCC[C@H]2C(=O)N[C@@H](CCCNC(=N)N)C(=O)c3ncccn3
1054 O=C1NCc2ccc(C3CCCN3C3N(CC3)c4cccc5CCCCc45)cc12
1055 C[C@H](Cc1ccccc1)N(C)CCCC
1056 Clc1ccc(cc1Cl)S(=O)(=O)CCCCCN2CCC(CC2)c3noc4ccccc34
1057 Nc1ncnc2c1c(cn2C3CCcc3)C#C
1058 N=C1NC2CCNCCN2c3ccccc13
1059 C(Oc1ccccc1c2ccccc2)C3=NCNN3
1060 CCOc1nc(cc(N)c1C#N)C(=O)OCc2ccccc2S(=O)(=O)N
1061 Clc1cc2[C@@H]3CCCNN3C(=O)c2cc1CCC
1062 COc1cc(Nc2nccc(Nc3cnc4ccccc4c3)c2)cc(OC)c1CC
1063 NS(=O)(=O)c1ccc(cc1)c2=cccc2c3ccc(F)cc3
1064 NCCNCC(C)c1ccccccncc1
1065 CC(C)NCC(C)COc1cccc2ccccc12
1066 Cc1cc(c(O)c(C)c1NC2=CCCC2)C(C)(C)C
1067 COC(=O)C1=C(C)NC(=O)N(C1c2ccc(F)c(F)c2)C(=O)NCCCNCCCC(CC3)c4ccccc4C3
1068 CCCOc1cc2c(cc1\\C(=C/C=C/C(=C/C(=O)O)/C)\\C)C(C)(C)CCC2(C)
1069 COc1ccc(NC(=N)NC(=O)c2cn(nc2c3ccc(F)nc3)c4ccccc4)cc1
1070 C(Cc1ccccc1c2cccs2)C3=NCCNN3
1071 COc1cc2ncnc(Oc3ccc(NC(=O)Nc4cccc(BC)c4)cc3)c2cc1OC
1072 Cc1ccc(cc1)N(CC2CNCCN2)c3cccc(O)c3
1073 CCN(CC)C(=O)c1ccccc1NCC2ONCCN2
1074 C(c1ccccc1)c2ncc3CCCCCc3n2
1075 O=C1NC(=NC(=C1)c2ccncn2)NCCCCCCCC
1076 COc1c(C)c2COC(=O)c2c(O)c1C\\C\C(CCC\\CCCC=O)O
1077 COc1ccc(cc1)c2cc(nn2c3ccc(cc3)S(=O)(=O))CC(F)(F)F
1078 CC1(N(CCc2cc(O)ccc12)c3ccc(Cl)cc3)c4ccc(OCCN5CCCC5C)cc4
1079 NS(=O)(=O)c1ccc(cc1)c2=C(CC3(CC3)C2)c4ccc(F)cc4
1080 Fc1cc(C)c2c(n1)sc3c(N)ncnc23

1081 Fc1ccc2c(noc2c1)C3CCN(CCCCNS(=O)C(O)c4cc5ccccc5s4)CC3

1082 CNC[C@H](O)c1ccc(O)ccOcc1

1083 Fc1ccc(cc1Cl)S(=O)(=O)NC2CN2CC2(CC2)c3nsc4ccccc34

1084 Cc1cccc(c1)N2N=C3N(C2=O)c4ccccc3OOC4N

1085 CCCCCCNS(=O)(=O)NCCC(=O)O

1086 C(Nc1ccccc1c2cccs2)C3=CCCN3

1087 Fc1ccc(cc1)C(=O)C2CCN(NCN3C(=O)Nc4ccccc4C3=O)CC2

1088 CCCCCCCCC(=CCC(=O)CCCCCCCC)

1089 OCCNCCc1O[cH]c2ccc(F)cc12

1090 COc1ccc(Cc2cncn2CCCcccc1)

1091 Cc1cc2CNC(=O)c2cc1OCCCN3CCN(CC3)c4cccc5cccccCcc45

1092 CC[C@@]1(O)C(=O)OCCC2C1C=C3N(Cc4cc5c(N)cccc5nc34)C2CO

1093 Cc1nccn2c(c3cccc(NCC(C)(CCC))n3)c(nc12)c4ccc(F)cc4F

1094 NC(COCc1ccccc1)COc2cncc(C=Cc3ccccc3)c2

1095 Cc1ccc(cc1)[C@@H]2Cc3ccc(O)cc3O[C@@H]2c4ccc(OCCN5CCCC5)cc4

1096 Nc1ccc2nc(sc2c1)S(=O)(CO)N

1097 COc1cccc(c1)C(C)NC(=O)c2cc(C)c(s2)c3ccc4[nH]cc(C)c4c3

1098 NC(=N)c1ccc(CNC(=O)[C@@H]2CCN2C(=O)[C@H](CCC(=O)C)C3CCCCC3)cc1

1099 CC(C(=O)O)c1ccc(c(O)c1)c2ccccn2

1100 CCCNC(=O)c1ccccc1CCc2c[nH]cc2

1101 Cc1ccc(COc2ccc3c(c2)c(SC(C)(C)C)c(CC(C)(C)C(OO)O)n3Cc4ccc(cc4)c5ccc(F)cn5)nc1

1102 CN(C)CC\\C=C/1c\c2ccccc2Sc3ccc(Cl)nc13

1103 NS(=O)(=O)c1ccc(cc1)c2ccccccCcCCC=C2CCCO

1104 Clc1cccc(N2CCN(CCCCCS(=O)(=O)c3cnc4ccccc4c3)CC2)c1Cl

1105 CC(Oc1ccccc1c2cccc(c2)[N+](=O)[OH])C3CCCCN3

1106 COc1cc2CCNC[C@@H](C)c2cc1B

1107 CN1CCN(CC1)c2ccc(Nc3ncc4C(C(CNN)C(NO)NCCCCCCCC)c4n3)cc2

1108 Cl.CC(C)Sc1ccccc1NCC2NCNNN2

1109 C[C@H](N1CCN(C[C@@H]1C)C2(C)CCN(CC2)C(=O)c3c(C)cccc3C)c4ccc(cc4)C(F)(F)C

1110 CN.COc1ccc(F)c(Cl)c1C2CC2CN

1111 COc1ccc(cc1OC)S(=O)(C)NN[C@@H]2CC[C@H](CC2)N3CCN(CC3)c4ccccc4OC(C)C

1112 COC(=O)C1=C(C)NC(=O)N(C1c2ccc(F)c(F)c2)C(=O)NCCCN3CCN(CC3)c4ccccc4CC

1113 COc1ccccc1CC2cc3nc(nc(N)c3n2)c4occc4

1114 COC(=O)c1ccccc1CCC2=NCCC2

1115 CCn1nnc(n1)[C@H]2O[C@H]([C@H](O)[C@@H]2O)n3cnc4c(NC5CC5)cc(Cl)nc34

1116 Cc1cCNC2=NCCN2NcOc3nccnc13
1117 O=C1Cc2cc(CCC3CCN(CC3)c4cccc5ccccc45)ccc2N1
1118 CCn1nnc(n1)[C@H]2O[C@H]([C@H](O)[C@@H]2O)n3cnc4c(N)nc(N[C@H](CC)Cc5ccccc5)nc34
1119 Cc1ccc2c(cccc2n1)N3CC3(CCc4cccc5c4OCc6c(ncn56)C(=O)NCCCCCF)
1120 COc1ccccc1CNNCc2cccc(CCNC[C@H](N)c3ccc(O)c4NC(=O)Sc34)c2
1121 Cl.COc1nsnc1OCCOCCOCCOCCOc2snnc2CNCCCCCCCCCC
1122 Cc1ccc(cc1)c2cc(nn2c3ccc(cc3)S(=S)(=O)NNC(F)(F))
1123 CCOC(Cn1nc(c2ccc3oc(N)nc3c2)c4c(N)ncnc14)OC
1124 COc1ccc(NC(=O)Nc2cccc(F)c2)cc1c3c(BB)cnn3CC
1125 CCc1cc(Cl)ccc1Oc2cc(C)ncc2CNCC
1126 CC(Oc1ccccc1C2CCCC2)C3CNCCC3
1127 NCc1ccc(cc1)S(=O)(=O)
1128 OC(CNC1CCN(CC1)c2nccc3scc(c4ccccc4)c23)COc5cccc(c5)CCN
1129 COc1ccccc1N2CCN(CCCCNC(=O)c3cc4cc(I)ccc4I3)CC2
1130 COc1cc2ncnc(Nc3ccc(F)c(c3)C#C)c2cc1OCCCCCCC(OO)OO
1131 Cc1cc(on1)C(=O)N[C@@H]2CC[C@@H](CCCCCCC(CCC)c4coc5ccccc45)CC2
1132 Nc1ncnc2c1c(cn2C3CCCC3)C
1133 CC1=CC(C)(C)Nc2ccc3c4ccccc4OC(c5ccc(Cl)c(CC)c5)c3c12
1134 CN(C)C(=O)NCCC1C2cc(O)scc2C1
1135 Fc1ccc(F)c(c1)N2CCN(CCN3C(=O)CC4(CCCC4)CC3OO)CC2
1136 COc1ccccc1N2CCN(CCCC3Cc(O)cc4ccccc43)CC2
1137 CO[C@@H]1CC[C@@H](CC(NO)N[C@@H]2CC[C@@H](CCN3CCC(CC3)c4coc5ccccc45)CC2)CC1
1138 Brc1cncn1Cc2ccc(cc2)CCN
1139 OCC(NC(=O)N1CCC(=CC1)c2c[nH]ccnc3cc23)c4ccccc4
1140 NS(=O)(=O)C1Occ(O)cc1  1
1141 Oc1ccc2C[C@H]3N(CC4CC4)CC[C@@]56[C@@H](Oc1c25)C(=O)CC[C@@]36
1142 NS(=O)(=O)c1ccc(sc1ccscccc2)2CCOOO
1143 COc1ccccc1N2CCN(CNCCNC(=O)c3ccc(CCC)cc3)CC2
1144 Clc1cc2[C@@H]3CNCCN3C(=O)c2cc1
1145 Clc1cccc(Oc2ncccc2n3CCNCC3)c1Cl
1146 CN1N=C(SS1=N)S(=O)(=O)N
1147 Clc1ccc(cc1Cl)S(=O)(=N)NCCCCN2CCC(CC2)c3noc4ccccc34
1148 COc1ccccc1N2CCC(CCCCNC(=O)c3cc4ccccc4s3)CC2
1149 O=C1CCc2ccc(OCNCCN3CCN(CC3)c4cccc5OCCCc45)nc2N1
1150 NC(COCc1ccccc1)COc2cncc(CCCc3ccccc3)c2

1151 CCC(CC)n1c(C)cc2c3c(N)cccNnnc3ccc12
1152 NC(=O)c1cc2c(Oc3ccc(BB)cc3)cncc2s1
1153 COc1cc2c(Oc3ccc(NC(=O)C4=C(C)N(C(=O)N4)c5ccccc5Cl)cc3F)cccc1cc2OCCCN6CCN(C)CC6
1154 CN1N=C(SC1=O)S(=O)(=O)N
1155 CC1(N(CCc2cc(O)ccc12)c3cccc(C)c3)c4ccc(OCCN5CCCC5)cc4
1156 COc1cc(ccc1Nc2ncc3CN(C)C(=O)N(c4cccc(NC(=O)C=C)c4)c3n2)N(C)CC
1157 Clc1ccc(Cn2cccc2)cc1
1158 Nc1ncnc2c1c(O)nn2C3CCC(O)CC3
1159 CSc1ccccc1NCC2CNCCN2
1160 COc1ccc(cc1OC)S(=O)(=O)N[C@@H]2CC[C@H](CC2)N3CCC(CC3)c4ccccc4OCC(F)(N)
1161 COc1ccccc1N2CCN(CCCCCNC(=O)CC(C)(C)C)C2C
1162 Nc1ncc(c2cocc2)c3scc(c4ccc(NC(OO)Nc5cccc(F)c5)cc4)c13
1163 Cc1cccc(c1)N2N=C3N(C2=O)c4ccccc3ONC4N
1164 O=C1NCc2ccc(OCCCN3CCC(CC3)c4cccc5COCCc45)cc12
1165 Clc1ccc(COc2ccc3SC(CO)Cc3c2)cc1
1166 COc1cc(ccc1Nc2ncc3CN(C)C(=O)N(c4cccc(NC(=O)C=C)c4)c3n2)N5CCCCC5
1167 CS(=O)(=O)Cc1cccc(Nc2nccc(Oc3ccc4c(cccc4c3)C(=O)Nc5ccccc5)c2)c1
1168 COc1cc2c(Oc3ccc(cc3F)C4=CN=C(Cc5ccccc5)c(C)C4=O)ccnc2cc1OCCCN6CCOCC6
1169 Nc1ncnc2c1c(cn2C3CCOC3)c4ccc(Oc5cccC(C)cc5)cc4
1170 Cc1cccc(NC(=O)Nc2ccc(cc2)c3coc4cccc(N)c34)c1
1171 NS(=O)(=O)Oc1ccccccccccc1
1172 Oc1c2ccccc2c3C=Nc4ccc31c4
1173 Cc1ccc(cc1S(=O)(=O)N)c2oc(nc2)C(=O)Nn3CCC3
1174 Cc1ccc2c(cccc2n1)N3CCN(CCc4ccc5NC(=O)COc5c4)nC3
1175 COc1ccccc1N2CCN(CNC(=O)c3cccc2Cl)c3CCC
1176 CN(Cc1ccc2NC(=NC(=O)c2c1)O)c3ccc(s3)C(=O)N[C@@H](CCC(=O)O)C(OO)O
1177 Cc1cccc(c1)c2cncn2Cc3ccc(cc3)CN
1178 Clc1cc2NC(=O)Cc2cc1CCN3CCN(CC3)c4sss5cccccc54
1179 Fc1ccc2c(noc2c1)C3CCN(CCCNS(=S)(=O)c4cc5ccs(Cl)cc5s4)CC3
1180 Cc1ccc(cc1CC)C(=C)CCCCCC(CC(C)C)CCCCC
1181 CN1CCC2Ccccc2ccccccCCCc1
1182 O=S(=O)(NCCCN1CCC(CC1)c2nsc3ccccc23)c4ccc5ccccc5c4
1183 Ic1c(NC2CNC)N2Nccc3nncnc13
1184 CN1CCc2ccccc2Cc2cccC2CC1
1185 CC1=CC[C@@H]2[C@@H](C1)c3c(O)cc(cc3OC2(C)C)C(C)(C)C4CCCCCCC4

1186 O=C1NCc2ccc(CCCCCN3CCN(CC3)c4cccc5COCCc45)cc12
1187 COc1ccccc1C2CCN(Cc3cc4ccccn4n3)CC2
1188 Cl.COc1ccc(F)c(CC)c1C2CC2CN
1189 COc1ccc2[nH]cc(CCN(CC=C)CCNC)c2c1
1190 Cc1ccc2c(cccc2n1)c3cCN(CCc4cccc(c4)N5CCNC5=O)CC3
1191 CCCCCCC(C)(C)c1cc(O)cc(OCCCCCCCC(=O)CCCCC)c1
1192 O[C@@H](CNCCCCCCCCN1CCC(CC1)OC(=O)Nc2ccccc2c3ccccc3)c4ccc(O)c5NC(CO)C=Nc45
1193 Cc1ccc2c(cccc2n1)N3CCN(CCc4cccc5c4OCc6c(ncn56)C(=O)N7OCOCC7)CC3
1194 COC(=O)C1=C(C)NC(=O)N(C1c2ccc(F)c(F)c2)C(=O)NCCCN3CCC(CC3)c4ccccc4C
1195 CCCN(CCC)C1CCC(=CC1)CN
1196 COc1ccc2ccNc3cccc(ccBB3)cccc2c1
1197 CCCCCOc1ccccc1c2occ(c2)C(=O)NC3CCCCC3
1198 CC1=C2cCc3cc(Cl)ccc3N2CCC1=O
1199 COc1ccccc1N2CCC(CNCC3CCc4ccc(Cl)cc4O3)CC2
1200 NS(=O)(=O)c1nc2ccc(=CS)ccc2s1
1201 Fc1ccc(cc1)C(OCCC2CCN(CCCc3ccccc2)CC3)c4ccc(F)cc4
1202 Cc1cc2C(=O)Nc3ccccc3Nc2nn1
1203 O=C1NCc2Ccc(OCCCCN3CCN(CC3)4)cccc5ccccc45ccc12
1204 Oc1cccc2C[C@@H]3[C@H]CCCN3CC=CCCc12
1205 Cc1cccc(c1)S(=O)(=O)NCCCN2CCC(CC2)c3nsc4cc(F)ccc34
1206 COc1cc2c(Oc3ccc(NC(CO)c4cc(ccn4)c5ccc(C)cc5)cc3F)ccnc2cc1OCCCN6CCN(C)CC6
1207 Cc1cccc(NC(=O)Nc2ccc(cc2)c3csc4c(cnc(N)c34)C#CCC5CCOCC5)c1
1208 CC(C)c1cc(Oc2c(Cl)cc(CCC(=O)C)cc2Cl)ccc1OO
1209 CCCCCCC(C)(C)c1cc(O)cc(OCCCCCCC)(=O)NC2CC2cc1
1210 O=C1NCc2ccc(OCCCCN3CCN(CN3)c4cccc5ccccc45)cc12
1211 COc1ccc2[nH]cc(CCN(CC=O)CC=O)c2c1
1212 COc1cc2ncnc(Nc3cccc(BC)c3)c2cc1OC
1213 Clc1cc2cNNn2ncnc3ccccccc3c1
1214 COc1cccc(c1)C(C)NC(=O)c2ccc(cc2OC)c3ccncc3
1215 Cc1cc2CNC(=O)c2cc1OCCCCc3CCN(CC3)c4cccc5ccccc45
1216 NS(=O)(=O)Oc1ccccNC(=O)Nc2ccc2ccccccccccc1
1217 Cc1ccc2c(cccc2n1)N3CCN(CCc4cccc5c4)N5CCCOC(=O)CC3
1218 CN[C@H](Cc1ccccc1)C(=O)N2CCC[C@H]2C(=O)NC[C@@H]3CC[C@@H]NNCCC3
1219 O=C1CC2(CCCC2)CC(=O)N1CCCCNCC3CCc4ccccc4C3
1220 CNCCCCOc1ccccc1CCCc2ccccc2

1221 CN1CCc2c(C1)Cc3N=CN(CCN4CCN(CC4)c5cccc6ccccc56)C(=O)c23

1222 Cc1ccc2c(cccc2n1)N3CCN(CCc4cc5cc(=O)CON5cc4F)CC3

1223 COc1cc(C=C2SC(=Nc3ccccc3)CC2=O)ccc1O

1224 COc1ccccc1CNCCCCCCNCCCNCCCCNCCCCCCNCc2ccccc2OC

1225 Oc1ccc(cc1)C23COc3cc(O)cc(O)cCC2=O

1226 NC(=N)c1ccc(CNC(=O)[C@H](CCC2CCNCC2)NC(=O)[C@@H](CCc3ncncc3)NS(=O)(=O)Cc4ccccc4)[illegible]

1227 CC(Oc1ccccc1c2cccc(c2)[N+](=O)[OH])C3=CCCN3

1228 Clc1ccc(cc1)C(=C)CCC32CCC3cccc4CC24

1229 COC(=O)c1ccccc1NCC2CCCCN2

1230 Cc1c(cc(c2ccccc2)n1C)C(=O)NCCCN3CCN(CC3)c4cccc(Cl)c4CC

1231 COc1cc(ccc1Nc2ncc3CN(C)C(=O)N(c4cccc(NC(=O)C=C)c4)c3n2)N5CCCCC5C

1232 Fc1ccc2c(noc2c1)C3CCN(CNCCNS(=O)(=O)c4cccc5scnc45)CC3

1233 COc1ccc(\\C=C/c2cc(O)Nc(OC)c(OC)c2)cc1O

1234 O=C(NCC1CCCCC1)c2cc3c(n[n]cc3s2)c4ccccc4

1235 COC(=O)C1C(CCCCN1)c2ccccc2

1236 FC(F)(F)c1ccc2c(NC(=O)Nc3ccccn3)nnnc2c1

1237 C(Nc1ccccc1c2cNcn2)C3=CCCN3

1238 COc1cc2c(Oc3ccc(NC(=O)N\\N=CC\c4cccc(CC=C)54F)cc3F)ccnc2cc1CCCC5NCCCCC

1239 CN(C)CC1CC2N(O1)c3cccCOccc3cc4ccccc24

1240 CN1CCc2ccccc2cc3cccc3CC1

1241 Clc1cccc(c1)C2CCCCC2

1242 C(Nc1ccccc1c2ncccc2)C3CNCCN3

1243 CN1CCc2ccccc2Cc3ccccc3C1

1244 CC(C)n1nc(c2ccc3O)c(O)c2cc3c(N)ncc1

1245 Oc1ccc(cc1)c2ccc3cc(O)ccc3c2Oc4ccc(CCCN5CCCCC5)cc4

1246 C=CCNNCC(O)COc1ccccc1CC 1

1247 CN(C)CCC1=C(Cc2cnncn2)c3ccccc3C1

1248 Clc1ccc(cc1)N2CCN(Cc3cc4ccncn4n3)CC2

1249 CCCCOC(=O)c1ccccc1NCC2=NCNN2

1250 OCC[C@@]1(CCCCCC1)c2ccc(Cl)c(Cl)c2

1251 Clc1ccc(cc1)c2cc([nH]n2)C3CCNCCCc4ccccc4CCC3

1252 CCOc1nc(cc(NCc1CCN)C(=O))Cc2cccnc2

1253 Cc1ccc2c(cccc2n1)N3CCN(CCc4cccc5N(C6CC6)C(=O)CCc45)CC3

1254 NS(=O)(=O)Oc1cccNN1BO 1

1255 CC(=O)O.OCc1cc(ccc1O)[C@@H](O)CNCCCCCCOCCCCc2cc(Cccccc2CC)

1256 Cc1ccc2c(cccc2n1)N3CCN(CCc4cc5NC(=O)COc5cc4F)nC3
1257 Cl.CC(C)Sc1ccccc1NCC2=CCCN2
1258 CN1C(=O)C(=Cc2cnc(Nc3ccc(F)cn3)nc12)c4c(Cl)cccc4Cl
1259 CN1CCc2c(C1)Cc3N=CN(CCN4CCN(CC4)c5cccc6ccnnc56)C(=O)c23
1260 COc1ccc(F)cc1CCCCCCCCCCCCCCC
1261 NS(=O)(=O)c1ccccCCCccc1
1262 COC(=O)c1ccccc1NCC2=NCCC2
1263 CCCCN1CCC(COC(=O)c2cc(CC)c(CC)c3OCCOc23)CC1
1264 Cn1c2CCNcCc2c3ccccc13
1265 COC(=O)C1C2CCC(CC1c3ccccccccccc3)c2C
1266 Fc1ccccc1CNC2=C(Nc3ccccc3)N(OO)C2=O
1267 COc1ccccc1N2CCN(CC3CO)Nc(cccc(C)c3)CC2
1268 Cl.CC(C)Sc1ccccc1NCC2NNNNN2
1269 COc1ccc2c(NC(=O)Nc3cccc(c2)n3)ccnccc1
1270 Nc1nc(N)c2nc(CNc3ccc(cc3)C(=O)NC(CCCCNC(=O)c4ccccc4C(=O)O)C(OO)O)cnc2n1
1271 Fc1ccc2c(cccc2c1)N3CCN(CCCCc4ccc5COC(=O)c5c4)CC3
1272 OCc1cc(ccc1O)S(O)CCCCCCCC=CCCCc2ccccc2
1273 NC(=N)c1ccc2cc(cc(Nc3ncccc3)c2c1)C(=O)Nc4ccccc4
1274 COC(=O)C1=C(C)NC(=O)N(C1c2ccc(F)c(F)c2)C(=O)NCCCC3CCN(CC3)c4ccccc4
1275 COC(=O)[C@H]1N2CCC(C2)C[C@@H]1c3ccc4ccccc4c3
1276 CN(C)CC1CC2N(O1)ccccc(F)ccccccccccc2
1277 CCCCCCCC(=O)C(OO)CCCCCCC
1278 Cl.CC(C)Sc1ccccc1NCC2NNCCN2
1279 COc1ccc2CC[C@@H](Cc2c1)NC(=O)NCCCC
1280 COc1ccc2[nH]c(C)c(CCC(C)C)c2c1
1281 CC(C)(C)CCC1(CCCC1)C(=O)c2ccc3[nH]ncc3c2
1282 Clc1ccc(COc2ccc3NC(CO)Oc3c2)cc1
1283 Clc1ccccc1CC(C2CCNCC2)c3ccccc3
1284 CCNC[C@H]1O[C@H]([C@H](O)[C@@H]1O)n2nnc3c(N)cccc23
1285 CCCc1nc(C)c2C(=O)NC(=Nn12)c3ccccc3CCC
1286 CN(C)S(=O)(=O)NC1cCc2ccc(C)cc2O1
1287 O=C1NCc2ccc(OCCCNNCC3NCC3)c4cccc5CCCCc45ccc12
1288 O=C1NC(=O)\\C(=C\\c2ccc3ncccc2c3)n\C1
1289 CCN([C@@H]1CC[C@H](CC1)N2CCC(CC2)c3ccccc3OC(C)C)S(=O)(=O)c4ccc(OC)c(OC)c4
1290 CNC(=O)[C@H]1O[C@H]([C@H](O)[C@@H]1O)n2cnc3c(NCC)nc(nc23)C#Cc4ccccc4

1291 NS(=O)(OO)O

1292 Cc1cccc(NC(=O)Nc2ccc(NC(=O)c3csc4nccc(N)c34)cc2)c1

1293 OC1(CC2CCC(C1)N2CCCC(=C)c3ccc(F)cc3)c4ccccc4

1294 NC(=N)c1ccc(CNC(=O)[C@H](CCC2CCNCC2)NC(=O)[C@@H](CCC3CCNCc3)NS(=O)(=O)Cc4ccccc4)[illegible]

1295 COc1ccc(cc1)S(=O)(=O)NC(C)C(NO)NO

1296 Oc1ccc2C(N(CCc2c1)c3ccccc3)c4ccc(OCCCNCC5CC5)cc4

1297 Cc1cccc(CC(=O)Nc2ccc(cc2)c3csc4c(cnc(N)c34)C#CCC5CCCC5)c1

1298 COc1ccc(Nc2nccc(NC3=C(NC(C)C(O)(C)C)C(=O)C3OO)n2)cc1OC

1299 COc1ccc(cc1)C2CN(C)Cc3cccOCCCN4CCCCC4cccc23

1300 Nc1ncc(c2ccncc2)c3scc(c4ccc(NC(=O)Nc5cc(ccc5)CC(F)(F)F)cc4)c13

1301 CN1CCc2cccc(Cl)c2Cc1

1302 COc1cc(ccc1Nc2ncc(Cl)c(Nc3ccccc3S(=O)(=O)C(C)C)n2)N4CCC(CC4)N5CCNNCCCC5

1303 Clc1ccc(CNC2=C(Nc3ccncc3)C(=O)C2OO)cc1

1304 Clc1ccc2N3CCC(=O)C=C3CCcc21

1305 CCOc1nc(cc(N)c1C#C)C(=O)NCc2ccccc2

1306 CN1CCN(NC(=O)Nc2nnc(s2)S(=O)(=O)N)CC1

1307 CN1CCN(CC1)c2ccc3nc([nH]c3c2)C4=C(N)c5c(F)cccc5NC4OO

1308 CCC[C@H]1CCCC[C@@H]1NC2=NCCC2

1309 Clc1ccc(NC(=O)c2ccccc2NCc3ccnnc3)cc1

1310 Cc1ccc(NC(=O)Nc2ccc(cc2)c3cccc4C(NO)NCc34)cc1

1311 COc1ccccc1CNCCCCCCNNNCCCCCCNCCCCCCNCc2ccccc2CC

1312 OC(CNC1CCN(CC1)c2ncnc3scc(c4ccccc4)c23)COc5cccc(c5) #N

1313 COc1ccc2C[C@@H]3CNNCN3C(=O)c2c1

1314 Nc1nccc2c1c(nn2C3CCCC3)c4cccc(O)c4F

1315 CCCC(N1CCCC1)C(=O)c2cc(CCO)Cc2

1316 COc1ccc2CC(CCc2c1)NSS(O)(=O)N(C)C

1317 CC1(N(CCc2nc(O)ccc12)c3ccccc3)c4ccc(OCCN5CCCCCC5)cc4

1318 CC(CCCCCC=c2ccc1cnncCC(CcN2))c1

1319 OC1(CC2CCC(C1)N2CCCNc3cccc(F)c3)c4ccc(C)cc4

1320 Clc1cccc(c1)N2CCNCCCN3Cc4ccccc4C3CC2=O

1321 COc1ccc(Cl)cc1N2CCN(CCNC3CCC(=NC(C)C3=O)c4ccccc4)CC2

1322 CC(C)(C)c1cc(cc(c1O)C(C)(C)C)C(=O)CCCCCC

1323 Nc1nccc2c1c(nn2C3CCCC3)c4ccc(Cl)c(O)c4

1324 Fc1ccc(Cl)cc1S(=O)(=O)N[C@@H]2CC[C@H](CC2)N3CCC(Cc3)c4ccccc4OCC(F)(F)F

1325 O=C1NC(=NC(=C1)c2ccccc2)NCCCCCCCC

1326 CN(CCOc1ccc(CC2SC(OO)NC2=O)cc1)c3ccccn3

1327 BrCc1ccc(cc1)C(=C)C(=O)c2ccccc2

1328 Cc1ccc(Cn2c(cc3ccccc23)C4CNCCC4)cc1

1329 Cl.CSc1ccccc1NCC2CNCCN2

1330 CCCCCCNS(=O)(=O)NCNC(CO)C

1331 COc1cccc(F)c1C2ONCc3cnc(Nc4ccc(C(=O)O)c(OC)c4)nc3c5ccc(Cl)cc25

1332 Cc1cc2c(cc1C(=C)c3ccc(cc3)C(=O)O)C(C)(CCCC2)CC

1333 CC1=CC[C@@H]2[C@@H](C1)c3c(O)cc(cc3OC2(C)C)C(CCCC)c4ccccCCccc4

1334 COc1ccccc1N2CCN(CCCCNC(=O)c3ccn4nccc4c3)Cc2

1335 CCCN(CCN)[C@H]1CCn2c(C1)ccc2C#N

1336 Oc1ccc2C[C@H]3N(CC4CC4)CC[C@@]56[C@@H](Oc1c25)c7[nH]ccccccccc7C[C@@]36O

1337 Fc1ccc2c(cccc2c1)N3CCN(CCCCc4ccc5CNC(OO)c5c4F)CC3

1338 CCCc1nn(C)c2C(=O)NC(=Nc12)c3cc(ccc3OCC)S(=O)(OOC)CCCCCCO

1339 Fc1ccc2c(noc2c1)C3CCN(CCCCNS(=O)(=O)c4cc5ccccC5c4)CC3

1340 CN(C)CC1CC2N(O1)c3cc(CC)ccc3Cc4ccccc24

1341 N#Cc1ccc(cc1)[C@@H]2C3ccccnnn23

1342 COc1ccc(NC(=O)Nc2ccc(cc2)c3csc4c(cnc(N)c34)c5cnn(CC(O)(C)O)c5)cc1

1343 FC(F)(F)c1ccc2c(SC(=O)Nc3ccccn3)ccnc2c1

1344 NS(=O)(=O)C

1345 COc1cc2ncnc(Nc3cccc(c3)C#C)c2cc1O

1346 C(C[C@H]1CCCC[C@@H]1NC2=NCCO2)C3cccccc3

1347 COc1ccc2Cc3ccccc3CCCN(O)CCc2c1

1348 CCCCCCCCC(=O)C(=O)CCCCOCC

1349 COC(=O)C1C(CC2C[C@@H](O)N1C2C)c3ccc4ccccc4c3

1350 Clc1cc2NC(=O)[C@H]3CNCCN3c2cc1C

1351 CN1CCc2c(C1)sc3N=CN(CCN4CCN(CC4)c5cccc6ncncc56)C(=O)c23

1352 COc1cc(ccc1Nc2ncc(Cl)c(Nc3ccccc3S(=O)(=O)C(C)C)n2)N4CCC(CC4)C5CCNNCCCC5

1353 O=C(N1CCOCC1)N2CCn3cc(C4=C(C(=O)CC4=O)c5cnc6ccccn56)c7cccc(C2)c37

1354 CCCNc1CCc2c(C)ccCC2C1

1355 COc1cc(ccc1Nc2ncc(Cl)c(Nc3ccccc3S(=O)(=O)C(C)C)n2)P(OO)(C)C

1356 OCc1cc(ccc1O)C(O)CCCCCCCC=CCCCc2ccccc2

1357 Clc1cc2CcC(=O)c2cc1OCCCNCCCC(CCC)c4cccc5ccccc45

1358 CC1NC(=Cc2cccc(Cl)c12)

1359 Cn1ccc2cc(ccc12)C3(Cc4ccccc4)CCN3

1360 OC(CCC1CCCCC1)(C2CCCCC2)c3ccccc3

1361 NC(=N)c1ccc(CNC(=O)[C@H](CCC2CCNCC2)NC(=O)[C@@H](CCc3ccccc3)NS(=O)(=O)Cc4ccccc4)[illegible]

1362 Fc1ccc(cc1)c2cncc(CNC[C@H]3CCc4ccccc4C3)c2

1363 CCCCC(CC(=O)NO)S(=O)(=O)c1CCCCC1C

1364 CCCCCCCCC(CO)C(=O)CCCCOCCC

1365 Cc1ccc2c(cccc2n1)N3CCN(CCc4cccc5c4OCc6c(ncn56)C(=O)NCCCCC)CC3

1366 COc1ccccc1N2CCN(CCCCCC(=O)OC(C)(C)C)CC2

1367 CN(C)CCCCC1c2ccccc2COc3ccccc13

1368 CC1(N(CCc2cc(O)ccc12)c3cccc(C)c3)c4ccc(OCCC5CCCC5)cc4

1369 NS(=O)(=O)c1ccc(NC)Cc2cccccccc2cccccccc1

1370 NNc1ccccc1S(=O)(OO)N

1371 CC(C)c1cc(nc(N)c1)c2ccc(F)c3ccccc23

1372 COc1ccccc1N2CCN(CCCCNC(=O)c3ccn4cccc4c3)CC2

1373 Clc1ccc2NCCCC(=O)CCCCCCc2c1

1374 Fc1ccc2CC(CC3CNCCN3)Cc2c1

1375 CCCC(N1CCCC1)C(=O)c2ccc(CC)C2

1376 Nc1ccc2ncnc(Nc3cccc(BN)c3)c2c1

1377 CCCC(N1CCCC1)C(=O)c2ccc(BB)cc2

1378 ONS(=O)(=O)c1ccccc1C(=O)

1379 Cn1ccc(n1)c2ccccc2NCC3=CCCN3

1380 OCCNc1cc2cc(ccc2cn1)c3cccnn3

1381 Nc1nc(N)c2c(CN)c(C(c3ccc(cc3)C(=O)CC(CCC(=O)O)CC=O)O)ccc2n1

1382 Clc1ccccc1CC(N2CCCCC2)c3ccccc3

1383 Nc1n[nH]c2ccc(cc12)c3ncn(Cc4ccccc4)c3c5ccccc5

1384 COc1cccc2CC(CCc12)NS(=O)(=O)N(C)

1385 CCOc1ccccc1CNCc2cccc(CCNC[C@H](O)c3ccc(O)c4CC(=O)Sc34)c2

1386 CC(C)CCC1(CCNC1)C(=O)c2cc(F)c3[CH]ccc3c2

1387 COc1cc2c(Oc3ccc(Nc4ccc(cc4CC(C)=C)C)cc3)ccnc2cc1OCCNCCO

1388 Clc1cc2NCC3CNCCN3c2cc1C

1389 NC(COCc1ccccc1)COc2cncc(CCCc3ccncc3)c2

1390 C(Cc1noc2ccccc12)n3CCN4Cc4ccccc3CCCC

1391 COc1cc2c(Oc3ccc(NC(=O)C4=NN(C(=O)c5ccccc45)c6ccc(F)cc6)cc3C)ccnc2cc1OCCCN7CCCC7

1392 COc1cc(ccc1O)CC=C(O)C(=O)c3c(O)cc(O)cc3C

1393 FC(F)(F)c1ccc2c(NC(=O)Nc3ccccc3)ncnc2c1

1394 CC(C)Oc1ccccc1N2CCN(CC2)[C@@H]3CC[C@@H](CC3)NS(=O)(=O)c4ccc(Cl)cc4C

1395 Fc1ccc2c(noc2c1)C3CCN(CC3NS(=O)(=O)c4cccs4)CC

1396 O=C(N1CCOCC1)N2CCn3cc(C4=C(C(=O)NC4OO)c5cnc6ccccn56)c7cccc(C2)c37
1397 Cc1c(cc(c2ccccc2)n1C)CCCCNCCCN3CCN(CC3)c4cccc(Cl)c4
1398 Nc1ncnc2onc(c3ccc(NC(NO)Nc4cccc(c4)C(F)(F)F)cc3)c12
1399 COc1cc2ncnc(Nc3ccc(F)c(c3)CCC)c2cc1OCCCCCCC(OO)NO
1400 CC(=O)O.OCc1cc(ccc1O)[C@@H](O)CNCCCCCCCCCCc2ccc(cc2)N3C(=O)CNc3=O
1401 Clc1ccc(cc1Cl)C23CNCC2C3 s
1402 COc1cc(ccc1Nc2ncc3CN(C)C(=O)N(c4cccc(NC(=O)C=C)c4)c3n2)N5CCC(CC5)C(C)C
1403 CC(=O)N1CCC(CNC(=C)NC23CC4CC(CC(C4)c2)C3)CC1
1404 COCC(=O)N[C@@H]1CC[C@@H](CCN2CCNCCC2Cc3coc4ccccc34)CC1
1405 CCN1C(=O)SC(=C1C)c2ccnc(Nc3cccc(OO)c3)n2
1406 C(Oc1ccccc1c2ccccc2)(C=NCNN)
1407 CN(C)CCCCCC(=O)C(C)(C1CCCCC1)c2ccccc2
1408 COc1ccc(cc1)S(=O)(=O)NC(C)C(=O)(O)
1409 Cc1cc2C(=Nc3ccccc3Nc2s1)N4CCCCC4
1410 COc1ccc(F)cc1CCCC2CCC(CCC)O2
1411 Cc1cc(ccc1Cl)C(CC2CCC2)Oc3cccc(Cl)c3
1412 CCSc1cc2N(C[C@H](C)N)CCc2cc1
1413 Cc1cccc(c1)c2cncn2Cc3ccc(cc3)C#C
1414 COc1ccccc1N2CCN(CCCCNC(=O)NC(C)(C)C)CC2
1415 Clc1cccc(N2CCN(CCc3cn(nn3)c4ccnnnccc5c4)CC2)51Cl
1416 Fc1ccc2cccc(N3CCN(CCCCOc4ccc5CCCc5c4)CC3)c2c1
1417 CN1CCc2c(Cl)ccc3oc(BB)c(C1)c23
1418 Fc1ccc2c(noc2c1)C3CNN(CCCNS(=O)(=O)c4cnc5ccccn45)CC3
1419 CCCCCCNS(=O)(=O)NOCC(OO)O
1420 FC(F)(F)c1ccc2c(NC(=O)Nc3ccnc(n3)C#N)ccnc2c1
1421 Fc1ccccc1CNC2=C(Nc3ccccc3)C(=O)C2=O
1422 COC(=O)c1cc(CCc2cc(OC)ccc2OO)ccc1
1423 Nc1nc(N)c2c(Cl)c(C(c3ccc(cc3)C(=O)NC(CCC(=O)O)C(=O)))ccc2n1
1424 CN1CCCCC[C@@H](C1)c2ccc(Cl)c(Cl)c2
1425 CCCc1nn(C)c2c(=O)NC(=Nc12)c3ccccc3OCC
1426 COc1ccc(cc1)C(C)NS(=O)(CC)
1427 COc1cc(ccc1Nc2ncc3CN(C)C(=O)N(c4cccc(NC(=O)C=C)c4)c3n2)N5CCC(C)CC5
1428 COc1cc(ccc1Nc2ncc3CN(C(=O)N(c4cccc(NC(=O)C=C)c4)c3n2)c5ccc(OCc6ccccc6)cc5)N7CCC([illegible]
1429 CC(C)n1nc(C#Cc2ccnc(O)c2)c3c(N)ncnc13
1430 CCOc1nc(NC(=O)Cc2cc(OCcccc2OC)S(=O)(=O)C)cc(N)c1Cl

1431 Cc1ccc(cc1)n2cnCn2Cc3ccc(cc3)CN

1432 CC(C)CC(NS(=O)(=O)c1c(F)c(F)c(F)c(F)c1F)C(OO)NO

1433 Clc1cccc(Cl)c1NC2=NCNN2

1434 CN1CCN(Cc2ccc(NC(=O)c3ccc(C)c(c3)C#Cc4ccc5[nH]nc(C)c5c4)cc2C(F)(F)F)CC1

1435 NCc1ccc(cc1)S(=O)(OO)N

1436 CC1CC(=Nc2cccc(Cl)c12)N

1437 CC1(C)OC(=C)Nc2ccc(cc12)c3cccc(c3)C#N

1438 CNC(=O)Oc1cccc(CNCC)CCCOcccccccc3cc3ccccccCc1

1439 COc1ccccc1N2CCN(CCCNC3=C(C)C(=O)N(C)C(OO)N3C)CC2

1440 COc1ccc2[nH]cc(CC(C)C)c2c1

1441 Oc1ccc2[C@H]([C@H](OCc2c1)c3ccccc3)c4ccc(OCCCCCCCC5)5c4

1442 Clc1cccc(c1)CCCC2CCC2

1443 OC(F)(F)C1CCC(CC1)C(=O)N[C@@H]2CC[C@@H](CCN3CCC(CC3)c4coc5ccccc45)CC2

1444 COC(=O)C1C2CCC(CC1c3ccc4ccccc4c3)s2C

1445 C[C@H](N)CN1CCc2ccc(CC)cc12

1446 Fc1c(OCCCN2CCN(CC2)c3cccc4ccccc34)ccc5CNC(OO)c15

1447 COc1ccc(cc1)S(=O)(=O)NC(C)C(=O)NONNO

1448 COc1ccccc1NCCCN(CCCNS(=O)(=O)c3ccc4ccccc4c3)CC

1449 CC(C)Oc1ccccc1C2CCN(CC2)[C@@H]3CC[C@@](CC3)NS(=O)(=O)c4ccc(F)c(F)c4

1450 Cl.CCC(=O)C(C1CCCCC1)c2ccc(Cl)c(CC)c2

1451 Clc1ccc2N3CCC(=O)CCC3CCc2c1

1452 CN[C@H](Cc1ccccc1)C(=O)N2CCC[C@H]2C(=O)N[C@@H](CCCN=C(N)N)C(=O)c3cc4ccccc4s3

1453 NS(=O)(=O)c1ncc(CO)cc1

1454 COc1ccc2CC(CCc2c1)NC(=O)N(C)CC

1455 OC(C1CCN(CCc2cc3cc2)CC1)ccccccc3

1456 NC(Cc1cccc2ccccc12)C(=O)Nc3cncccC=Cc4ccncc4cc3

1457 COCCCOc1cc(ccc1OC)C(=O)N(C[C@@H]2CNC[C@H]2OC(=O)NCc3ccccc3CCCC)C

1458 Nc1ncnc2c1c(cncnC3c2cc3)c4ccccc4

1459 COc1ccccc1N2CCN(CCCCNC(=O)c3ccn4nccc4c3)C2

1460 COc1c(Cl)cc(Cl)cc1N2CCN(CCNNC=Nc4ss5CC(C)CCc5c4CC=O)CC2

1461 Fc1ccc(Cn2cccc2c3ccccc3)cc1

1462 NCCNCCc1c[nH]c2ccc(F)cc12

1463 O[C@@H](CNCCCCCCCCN1CCC(CC1)OC(=O)Nc2ccccc2c3ccccc3)c4ccc(O)c5CC(=O)C=Nc45

1464 CC(N)C1CCC(CC1)C(OO)Nc2ccccc2

1465 O=C(Cc1ccc2ccccc2c1)Nc3cc(n[nH]3CC4CC)4

1466 CCCCCCC(C)(C)c1cc(O)cc(CCCCCCCCC(=O)NC2CC2)c1
1467 Fc1ccc2[nH]cc(CCCN3CCN(CN3)c4ccccc4)c2c1
1468 CN1CCc2c(C1)sc3NCCN(CCN4CCN(CC4)c5cccc6cccnc56)C(=O)c23
1469 CN(Cc1ccc2NC(=NC(=O)c2c1)C)c3ccc(s3)C(=O)N[C@@H](CCC(=O)O)C(OO)O
1470 Cc1cccnc1CC(P(=O)(O)O)P(=O)(OO)
1471 COC(=O)C1=C(C)NC(=O)N(C1c2ccc(F)c(F)c2)C(=O)NCCCN3CCN(CC3)c4ccccc4N
1472 Brc1c(NC2=NCCN2)ccc3nncnc13
1473 CSc1ccc(Cl)cc1NCC2=CCCN2
1474 Cn1nc(C(=O)N)c2CCc3cnc(NCc4CN(CC4)C(=O)c5ccccc5)nc3c12
1475 Cc1ccc2c(cccc2n1)N3CCN(CCCc4cccc5NC(CO)COc45)CC3
1476 COc1ccccc1OCCCCCc2ccc(O)c(O)c2
1477 NS(=O)(=O)c1ccc(NC(CO)Cc2ccccc2)cc1
1478 CC(C)Oc1ccccc1N2CCN(CC2NC3CCC(CC3)NCC=O)Nc4ccc(F)c(F)c4
1479 NC1=Nc2[nH]c(CCCc3ccc(cc3)C(=O)N[C@@H](CCC(=O)O)C(=O)O)cc2C(OO)S1
1480 Cc1ccc2c(cccc2n1)N3CCN(CCc4ccc5NC(=O)COc5n4)nC3
1481 N#CCC(C1CCCC1)n2nc(cn2)c3ncnn4[4n]ccc34
1482 O[C@@H](CNCCc1ccc(NS(=O)(=O)c2ccccc2)cc1)OOc3ccc(O)cc3
1483 NC(=O)C1=C(N)C(=O)C=C(Nc2ccccc2O)C1OO
1484 Clc1c(cccc1)CCCCCCCC
1485 COc1ccc(cc1)S(=O)(=O)NC(C)C(=O)NOON
1486 Clc1ccc(cc1)C(=O)N[C@@H]2CC[C@@H](CCC3CCC(CC3)c4cccc5OCCc45)Cc2
1487 CCc1cc(Oc2c(I)cc(CC3CC(=O)NC3CO)cc2I)ccc1O
1488 COc1ccccc1N2CCN(CCCCCCNCO)OCOC(C)(CCC)CC2
1489 COc1cc2cc(nc(N)c2cc1OC)N3CCN(CC3)C(OO)C4CCSS4
1490 COc1ccc(cc1OC)c2cc3nccn3c(Nc4ncccc4C(OO)N)n2
1491 COc1cc(ccc1Nc2ncc3CN(C)C(=O)N(c4cccc(NC(=O)C=C)c4)c3n2)NCCCC
1492 CN1C(=O)C=NN(CCCCN2CCN(CC2)c3ccccc3OCCO)C1=O
1493 O=C(N[C@@H]1CC[C@@H](CCN2CCC(CC2)c3coc4ccccc34)CO1)C5CCCOC5
1494 Nc1nc(N)c2c(Cl)c(C(c3ccc(cc3)C(=O)NC(CCC(=O)O)CC=O)O)ccc2n1
1495 COC(=O)C1=cCCN(C)C1
1496 CN1cCCC1c2oc(cc2)C(c3ccccc3)c4ccccc4
1497 COc1ccc(cc1OC)S(=O)(=O)N[C@@H]2CC[C@@H](CC2)N3CCN(CC3)c4ccccc4OC(C)
1498 CN(C)CCC(c1c2c(Cl)c21)2cccccn2
1499 Clc1ccc(cc1Cl)S(SO)(=O)NCCCCN2CCN(CC2)C3ssc4ccccc34
1500 OCc1cc(ccc1O)C(O)NCCCCCCCOCCCCc2ccccc2

1501 FC(F)(F)c1cc2NC[C@H]3CNCCN3c2cc1N

1502 O=C1CCc2ccc(OCCCCN3CCN(Cc3)c4cccc5OCCCc45)nc2N1

1503 NCCn1nnc2ccc(O)cc12

1504 COc1ccc2[nH]cc(CCC(C)C)c2c1

1505 N\1N=C\\1OOc2ccccc2C

1506 COc1cc2c(Oc3ccc(NC(=O)N\\N=C\\c4cccc(CC=O)c4F)cc3F)cccc2cc1OCCC5NCCCCC5

1507 CCn1nnc(n1)[C@H]2O[C@H]([C@H](O)[C@@H]2O)n3cnc4c(NCCCCCCC)ncnc34

1508 Clc1ccc(cc1Cl)C23CNCC2c3

1509 C(Nc1ccccc1c2cccs2)C3CNCCN3

1510 COc1cccc(n1)c2c(ncn2C3CCNCC3)c4ccc(F)cc4

1511 Clc1ccc(cc1Cl)S(=O)(=O)NCCCCN2CCC(CC2)c3csc4ccccc34

1512 COc1ccc(CCN2CCC(CC2)Nc3nc4ccccc4n3Cc5ccc(C)cc5)cc1

1513 CCC(CC)(c1ccc(OCC(O)CO)c(C)c1)c2ccc(OOC(O)C(C)(C)C)c(C)c2

1514 CC(C)n1nc(c2ccc(O)c(O)c2)c3c(N)ncnc13

1515 COc1ccc(cc1)C(=O)CCCN2CCc3ccccc3C2C

1516 COC(=O)CCNS(=O)(=O)NC(C)c1ccccccc1

1517 Cc1ccc(cc1)C(=O)C(CCOC)N2CCCC2

1518 Cc1cccc(Cl)c1NC(=O)c2cnc(NC(CO)C3CC3)s2

1519 COc1ccccc1NCCCN(CCCCNS(=O)(=O)c3cccccccccc3)CCC

1520 CSc1ccc(Cl)cc1NC2C=CCC2

1521 CNCCC(Oc1ccc(cc1)C(F)(F)C)c2ccccc2

1522 Cc1ccc2c(cccc2n1)N3CCN(CCc4cccc5c4OCc6c(ncn56)C(OO)N7CCCCC7)CC3

1523 CC[C@@]1(O)C(=O)OCC2C1C=C3N(Cc4cc5c(N)cccc5nc34)C2=O

1524 COc1ccc2[nH]cc(CCN(CCCC)CC=C)c2c1

1525 CC1(N(CCc2cc(O)ccc12)c3cccc(O)c3)c4ccc(OOCN5CCCCC5)cc4

1526 COc1ccc(cc1OC)S(=O)(=O)N[C@@H]2CC[C@H](CC2)N3CCC(CC3)c4cc(F)ccc4.C(C)C

1527 O=C1NCc2ccc(OCCCCN3CCN(CC3)c4cccc5CcCCc45)cc12

1528 C(Oc1ccccc1C2NC2)C3=NCCN3

1529 Cl.COc1ccccc1CC(N2CCCCC2)c3ccccc3

1530 NCCCCCc1c[nH]nnnn1

1531 COc1cc(ccc1Nc2ncc(Cl)c(Nc3ccccc3P(=O)(C)C)n2)N4CCC(CC4)N(CCC)

1532 C[C@H]1[C@H]2Cc3ccc(Nc4ccccc4)cc3[C@@]1(C)CCC2CC5CC5

1533 CN1CCc2cccc(Cl)c2CO1

1534 Cc1cccc(c1)S(=O)(=O)NCCCC2CCC(CC2)c3noc4cc(F)ccc34

1535 NC1=Nc2[nH]c(CCCc3ccc(cc3)C(=O)N[C@@H](CCC(=O)O)C(=O)O)cc2C(OO)N1

1536 Clc1cccc(N2CCN(CCCCNS(=O)(=O)c3ccc4ccccc4c3)CC2)c1Cl

1537 Clc1cccc(N2CCN(CCCC3nncnc4)c4ccncCcCcncc3CCC)c1C2

1538 COc1ccc(cc1)C(C)NS(=O)(=C)

1539 NS(=O)(=O)c1ccccNc2Cc2ccccc1

1540 NS(=O)(=O)c1ccc(CCC=O)ccccccccccccccccc1

1541 Ic1c(NC2=NCCN2)ccc3nncnc13

1542 CS(=O)(=O)c1ccc(cc1)c2nc(NCc3csss3)cc(n2)C(F)(F)F

1543 CC1=C2CCc3cc(Cl)ccc3C2CCC1=O

1544 CNC1=Nc2cccc(Cl)c2C(C)C1

1545 NS(=O)(=O)Oc1ccc(NC(=O)Nc2ccccc2cccccc1)

1546 CC(C)c1cc(Oc2c(BB)cc(CC(=O)O)cc2B)Bccc1O

1547 Cc1ccc2c(cccc2n1)N3CCN(CCc4cccc5c4OCc6c(ncn56)C(=O)N7CCC7)CCC3

1548 Cc1cc2C(=Nc3ccccc3Nc2s1)N4CCNNC4

1549 Fc1ccccc1N(Cc2ccc(Cl)cc2)c3CCC3

1550 COc1ccccc1NCCCN(CC(O)CCNC(=O)c3ccc4ccccc4c3)CCC

1551 COc1cc(ccc1Nc2ccc(Cl)c(Nc3ccccc3C(=O)N)n2)P(=O)(C)C

1552 Cc1c(NC2CcCCN2)ccc3ccccc13

1553 COc1ccc(cc1)S(=O)(=O)c2ccc(cc2)C(=C)C3CCN(CC33C4CCCCCC3)S(OO)(=O)Cc5ccccc54

1554 NC(=N)N1CCC[C@H](NC(=O)CN2CNNC[C@H](NS(=O)(=O)Cc3ccccc3)C2=O)C1O

1555 CC1=CC[C@@H]2[C@@H](C1)c3c(O)cc(cc3OC2(C)C)C(C)(C)c4ccc(BC)cc4

1556 CCC(CC)(c1ccc(OCC(O)CO)c(C)c1)c2ccc(OCC(O)C(CC(C)))c(C)c2

1557 Cc1nccn2c(c3cccc(NCC(C)(C)CO)n3)c(nc12)c4ccc(F)cc4F

1558 Cc1ccc2c(cccc2n1)N3CCN(CCc4cccc5c(CC(F)O)C(=O)Ccc45)CC3

1559 Fc1ccc(Nc2ccccc2)nc1Nc3ccccc3

1560 NS(=O)(=O)c1ncc(CC)cc1

1561 NC(=N)NCCC[C@@H](NC(=O)CN1CCN(CC1=O)S(=O)(=O)Cc2ccccc2)C(=O)c3cccs3

1562 Oc1ccc2OC[C@H](CCCc3ccccc3)Cc2c1

1563 NS(=O)(=O)ONC(=O)CCc1ccccc1

1564 CN(C)CC\\COOC1\\c2ccccc2cc3ccc(c3)cc1

1565 COc1cc(ccc1Nc2ncc3CN(C(=O)N(c4cccc(NC(=O)C=C)c4)c3n2)c5ccc6ccccc6c5NN7)NN(C)C7

1566 Oc1ccc2c(noc2c1)c3cc(BO)c(O)cc3O

1567 Clc1ccc(CNC2=C(Nc3ccncc3)C(=O)c2=O)cc1C

1568 Cc1n[nH]c2ccc(cc12)c3cncc(OCC(N)Cc4cccc(CC(F)(F)F)c4)c3

1569 CN(C)CC1=C(O)C2=CC(=CC(=O)N2CNC1)C

1570 CC1(N(CCc2cc(O)ccc12)c3cccc(F)c3)c4ccccOCCCNCCCCCccc4

1571 Cc1ccc2c(cccc2n1)N3CCN(CCc4cccc(c4)C5CCOC5=O)CC3
1572 CCCN1CCC[C@@](C1)c2cccc(O)c2
1573 CCn1nnc(n1)[C@H]2O[C@H]([C@H](C)[C@@H]2O)n3cnc4c(N)nc(N[C@H](CO)Cc5ccccc5)nc34
1574 Cn1c(SCCCN2CC3CCN(C3C2)c4ccc(C)cc4)nnc1C5CCCCC5
1575 CCOC(=O)c1ncn2c1CN(C)C(CO)c3cc(CC)ccc23
1576 Fc1ccc2c(noc2c1)C3CCN(CC4CC(CO)c5ccoc5C4)CC3
1577 C[C@H]1CCCCc2ccc(Cl)cc12
1578 OC(Cn1ccnc1)(P(=O)(O)O)P(=O)OOOO
1579 NCCC(C(=O)Nc1ccc2[nH]ncc2c1)c3ncc(Cl)c(CC)c3
1580 COc1ccccc1N2CCN(Cc3cn4cc(C)cc(C)c4n3)CC2
1581 CN1CCc2cc(CC)c(O)cc2C(C1)c3ccccc3
1582 Cn1c(ccc1c2ccc3NC(=O)OC(C)(C))ccc3C#2
1583 CC(N)COc1cnc(Cl)c(COCc2ccncc2)c1
1584 C1CN(CCN1)c2cc3cccc3c2
1585 CC1=CC[C@@H]2[C@@H](C1)c3c(O)cc(cc3OC2(C)C)C(C)CCCCCCCN
1586 CN[C@H](Cc1ccccc1)C(=O)N2CCC[C@H]2C(=O)N[C@@H](CCCN=C(N)N)C(=O)c3nccCCCCccs3
1587 CC(=O)Nc1ccc2c(CCC3CC3(Cc4ccccc4)CC)cccc2c1
1588 CCOc1ccc2nc(sc2c1)S(=O)(=N)N
1589 Nc1ccc(cc1BS)S(=O)(=O)N
1590 COc1cc(NC(=O)c2oc(cn2)c3ccc(Cl)cc3)cc(OO)c1
1591 CC(C)(C)c1cc(cc(c1O)C(C)(C)C)C(OO)CCCCCC
1592 Cc1ccc(Cn2cnnc2c3ccccc3)nc1
1593 O=C1CCc2ccc(OCCCCN3CCN(CC3)c4cccc5COCCc45)cc12
1594 Cc1ccc2c(cccc2n1)N3CCN(CCc4cc5NCC(O)COc5cc4F)CC3
1595 Clc1ccc(cc1)N2cCN(Cc3cnn4ccccc34)CC2
1596 NC(COc1cncc(c1)c2ccc3ccccc3c2)Cc4c[nH]c5ccccc45
1597 N[C@@H]1CC[C@H](CC1)NC(=O)c2cc(OCc3ccc(cc3)C(=N)N)cc(OCc4ccc(cc4)C(=O)N)c2
1598 Nc1nnc(C1)S(=O)(=O)N
1599 NS(=O)(=O)c1cc2ccc(CCC)c1c2
1600 CN(C)C(=O)N[C@H]1CCc2ccc(O)cs2C1
1601 CC(Oc1ccccc1c2cccc(O)c2)C3CNCCN3
1602 CCC[C@@H]1CCCC1N2CCN(C2=O)c3cccc(CC)c3
1603 Clc1ccc2Oc3ccccc3N=C(N4CNNCC4)c2c1
1604 Clc1ccc2CCCCc2c1Cl 1
1605 Brc1ccc(Cn2cncc2c3cnccc3)cc1

1606 CC(N)Cc1c[nH]ccc2ccc12

1607 Clc1ccc(COC(Cnnccnc2)cccccc(O)cc2Cl)cc1

1608 COc1cccc(c1)C(C)NC(=O)c2ncc(c(C)c2)c3ccncc3

1609 N\1N=C\\1/Oc2ccccc2C=CC

1610 CN(C)Cc1ccccc1Cc2cccc(Cl)c2

1611 CCc1cc(C)c2C(=C)N3CCNC[C@H]3Cc2c1

1612 Cc1c(Nc2ccc[nH]2)ccCCOCCCC1

1613 COc1ccc(cc1)S(=O)(=O)NC(O)C(=O)NON

1614 Cc1n[nH]c2ccc(nc12)c3cncc(OCC(N)Cc4ccc(Cl)c(Cl)c4)c3

1615 Cc1ccc2c(cccc2n1)N3CCN(CCc4c(C)ccc5NC(=O)C=Nc45)CC3

1616 CN(C)CC1(CCCC1)c2ccc(F)c(Cl)c2C

1617 Nc1ncnc2c1c(cn2C3CCC(CCO)CC3)c4cccc(O)c4

1618 COc1cc2c(Oc3ccc(cc3F)C4=CN=C(Cc5ccccc5)c(C)C4=O)ccnc2cc1OCCCCCCCCCCC

1619 Cc1cccc(NC(=O)Nc2ccc(cc2)c3csc4c(cnc(N)c34)C#CC(CC(C)))c1

1620 CN[C@H](Cc1ccccc1)C(=O)N2CCCC[C@H]2C(=O)NC(CCCNOC(N)N)C(=O)c3ncccccccccs3

1621 CCC(CC)(c1ccc(OCC(O)CO)c(C)c1)c2ccc(OCC(C)CCCC(C)C)c(C)c2

1622 Cc1cccc(CCN(CCc2ccccc2)CC3CC3)c1

1623 Cc1ccc2c(cccc2n1)N3CCN(CCCc4cccc5NC(OO)COc45)CC3

1624 Cc1ccccccc2cncn2Cc3ccc(cc3)C#1

1625 Clc1ccc(cc1)C(C2COCC2)C3C2CCC3NC2

1626 Cc1ccc2c(cccc2n1)N3CCN(CCc4cccc5NC(=O)COc45)CCC3

1627 Cl.NC[C@H]1C[C@@H]1c2ccc(F)cc2N

1628 CC(=O)NC1CCCCCCCcN1

1629 SS(=O)(=O)c1ccc(CO)cc1

1630 CC(C)(C)CCC1(CCCN1)C(=O)c2ccc3[nH]ccc3c2

1631 Clc1ccc(cc1)N2CCN(CCN3CCCCCC3)C2C

1632 C(Nc1ccccc1c2nccs2)C3=NCCNN3

1633 COc1ccc(cc1)C(C)NS(=O)(CC)N

1634 COc1ccc2[nH]cc(CCN(CC=C)CCCN)c2c1

1635 CC(Oc1ccccc1C2CcCc2)C3=NCCC3

1636 Clc1ccc(cc1Cl)S(=O)(=O)CCCCN2CCN(CC2)c3nsc4ccccc34

1637 COc1cc2c(Nc3ccc(cc3F)c4nc5ccccc5#4)ncnc2cc1OCCCCCCCN(C)CC

1638 NS(=O)(CO)c1ccc(CCO)cc1

1639 CN1CC(c2ccc3c(c2)C(C)(C)CCC3(C)C)c3cc(oc3C1)\\COC\\C(=O)O

1640 CS(=O)(=O)c1ccc(cc1)C2=C(C(CO)CC2)c3cc4ccccc4s3

1641 CCCNC(=C)c1ccccc1NCc2c[nH]cn2
1642 CCOc1cc2c(cc1\\C(=C/C=C/C(=C/C(=O)O)/C)\\C)C(C)(C)CCC2(C)
1643 CCCCOC(=O)NS(=O)(=O)c1sc(CC(C)C)cc1c2ccc(Cn3cccc3)cc2
1644 Cl.CCC(=O)C(C1CCCCC1)C2ccC(Cl)c(Cl)c2
1645 CCn1nnc(n1)[C@H]2O[C@H]([C@H](O)[C@@H]2O)n3cnc4c(N)ncnc43
1646 CCn1nnc(n1)[C@H]2O[C@H]([C@H](O)[C@@H]2O)n3cnc4c(N)nccc43
1647 Cl.NC[C@H]1C[C@@H]1c2ccccc2B
1648 COc1ccccc1CNCCCCCCNCCCCCCCCNCCCCCCCCc2ccccc2OC
1649 NS(=O)(=O)c1cc2ccc(OOS)ccc2n1
1650 CCCCCCCC(=O)C(=O)CCCC=CCC
1651 NCCCCCc1c2ncnc21
1652 CN1N=C(SC1=NCS(=O)(=O)N)
1653 Cc1cccc2C[C@@H]3CNCCC3C(=O)c12
1654 CCCc1c(nnn1Cc2ccccc2BC)C(=O)NCCCCCN3CC(CCC3)c4ccccc4OC
1655 COc1cc2c(Oc3ccc(cc3F)C4=CN=C(Cc5ccccc5)c(C)C4=O)ccnc2cc1OCCCC6CCOCC6C
1656 COc1ccccc1N2CCN(C\\C=C\\CNC(=O)c3ccc(cc3)c4ccccc4)CC2
1657 NS(=O)(=O)Sc1ccc(I)cccc1
1658 Nc1nccc(n1)c2c(ncn2C3CCNCC3)c4ccc(C)cc4
1659 CCC1(CCCCN2CCN(CC2)c3ccc(Cl)cc3)C(=O)N4ccc(F)ccc14
1660 COc1ccccc1N2C(NCCCCNS(=O)(=O)c3ccc4ccccc4c3)CC2
1661 Fc1cccc(Cl)c1Oc2ccccc2C3CCNCC3
1662 Cc1cc2c(cc1C(=C)c3ccc(cc3)C(=O)O)C(C)(C)CCC2(C)
1663 CNC(=O)CCCN(C)C(=O)c1ccc2c(c1)c3C[C@@H]CCCc3n2CCCCCCOCC
1664 Cc1ccc(Cn2c(nc3ccccc23)C4CCCCC4)cc1
1665 CN(C)CCCc1c[nH]c2cccccc1cc2#N
1666 CCn1nnc(n1)[C@H]2O[C@H]([C@H](O)[C@@H]2O)n3cnc4c(NC)cc(CC)nc34
1667 COc1cc(cc(OC)c1OC)C2=C(C(=O)NC2=O)c3c[nH]c4ccc3Bnccc4
1668 COc1ccc2Cc3ccccc3CCC(C)CCc2c1
1669 NC(C)(C)C(=O)NCc1ccc(cc1)S(=O)(=O)N
1670 COc1cc(ccc1Nc2ncc(Cl)c(Nc3ccccc3C(=O)N)n2)P(=O)(C)CC
1671 CN(C)c1ccc(cc1)N2CCc3cc(O)ccc3C2(C)c4ccc(OCCN5CCCCC5)cc4
1672 CC(=O)c1cc(NC(=O)NCCC[C@H]2C[C@H](Cc3ccc(F)cc3)CCC2)cc(c1)C(=O)C
1673 Clc1ccc(cc1Cl)S(SO)(=O)NCCCCN2CCN(CC2)c3nsc4ccccc34
1674 Cc1ccc2c(cccc2n1)N3CCN(CCc4cccc(c4)N5CCCCC5=O)CC3
1675 O=C1NCc2ccc(OCCCCC3CCN(CC3)c4cccc5CCCc45)cc12

1676 C(Cc1ccc2[nH]cnc2c1)N3CCC(CC3nn4)ccc5ccccc45

1677 Cc1cccc(N2CCN(CCc3ccc4[nH]ncc4c3)CC2)c1C

1678 COc1c(NC(=O)Nc2ccc(c3ccc(CN4CCOCC4)nc3)c5ccccc25)cc(cc1NS(NO)(=O)C)C(C)(C)C

1679 NC1cccccc2c1ccc2S(=O)(=O)c3ccccc3

1680 COc1cc2ncnc(Nc3ccc(F)c(c3)CCC)c2cc1OCCCCCCC(=O)NO

1681 CC(C)C[C@H](CC(=O)[C@H](Cc1ccccc1)NC(=O)c2cnccn2)B(O)O

1682 COc1ccc(cc1)C(NO)CCCN2NccccccccCC2

1683 Cc1cc(C(C#N)c2ccc(Cl)cc2)c(Cl)cc1CC(=O)c3cc(I)cc(I)c3O

1684 COc1cc(ccc1Nc2ncc(Cl)c(Nc3ccccc3C(=O)N(C)C)n2)N4CCC(CC4)N5CC(NC)CC5

1685 C\\NCC\\1/Oc2ccccc2CCC1

1686 COc1ccc(Cl)cc1S(=O)(=O)N[C@@H]2CC[C@H](CC2)N3CCC(CC3)c4ccccc4OCC(F)(C)

1687 Cn1ccc(n1)c2ccccc2NCC3=NCNN3

1688 NCCNCC1Cccnccccccncc1

1689 CCCCNNCC(O)COc1ccccc1CC=C

1690 CN1[C@@H]2CC[C@H]1CC(C2COC(=O)[C@H](C))c3ccccc3

1691 Cc1c(sc2ccc(F)cc12)S(=O)(=O)NCCCCCCCCC(CC3)c43oc5cc(F)ccc45

1692 Nc1nc(N)c2c(Cl)c(C(c3ccc(cc3)C(=O)CC(CCC(=O)))C(=O)O)ccc2n1

1693 O=C(Cc1cnccn1)Nc2ccnc3ccccc23

1694 COc1ccc2ccNc3cccc(ccBB3)ncnc2c1

1695 Cc1ccc(cc1c2ccc(cc2)C(=O)NCC3CC3)C(CO)NC4CC4

1696 O=C1NC(=NC(=C1)c2ccncn2)NCCCCCCCCC

1697 Cc1cc2CCN(C(=O)Nc3cccnc3)c2cc1CF

1698 CN(C)CCc1c[nH]c2cc2(cc12)c3cccccccnc32

1699 CC(C)n1nc(c2ccc3cc(C)ccc3c2)c4c(N)ncnc14

1700 Cc1cccc2N3CCNCC3NC(=O)c12 cc

1701 CS(=O)(=O)c1ccc(cc1)c2=C(CC3(CCC3)C2)c4ccc(O)cc4

1702 CC1=CC[C@@H]2[C@@H](C1)c3c(O)cc(cc3OC2(C)C)C(C)(C)c4cccC(C)cc4

1703 Cc1ccc(cc1)N2N=C3N(C2=O)c4ccccc4NNC3N

1704 CCCCC(CCS(O)NO)S(=O)(=O)c1CCCCC1

1705 Cc1cccc(c1)C(=O)NCN2CCN(CC2)c3ccccc3CN

1706 COc1ccccc1CNCCCCCCNNCCNCCCCNCCCCCCNCc2ccccc2OC

1707 NCCc1ccc(O)c(OO)1O

1708 CNC(=O)[C@H]1O[C@H]([C@H](O)[C@@H]1O)n2cnc3c(NC)ccnc23

1709 CCCN1CCC(COc2nc3c(N)cccc3c4ncccc24)CC1

1710 Cc1cc2CCN(C(=O)Nc3ccccc3)c2cc1Cl

1711 NCCNc12cccccc2cc1N

1712 Fc1ccc(cc1)C(=O)C2CCNCCC3CC(=O)c4ccccc4C3CCC2

1713 CC(C)Oc1ccccc1C2CCN(CC2)[C@@H]3CC[C@H](CC3)NS(=O)(=O)c4ccccc4O

1714 Fc1ccc2c(noc2c1)C3CCNNCCCNS(=O)(=O)c4cc5cc4(Cl)cc5sCCC3

1715 COc1ccccc1CN2CCC(CC2)C3CCN(CCCCCCCCN4CCC(CC4)C5CCC(Cc6ccccc6OC)CC5)CC3

1716 CCCNC(=O)c1c2CCOCc2sc1NC(=O)CC4CC5CC(CCCC5)CCCC4

1717 OCc1cc(ccc1O)C(O)CCCCCCCCCCCCc2ccccc2

1718 O=C1CCc2ccc(OCCCCN3CCN(CC3)c4cccc5CCCc45)cc2N1

1719 CC(N)C1CCC(CC1)C(=O)Nc2ccccc2

1720 CCCCCN(CCCCC(O)(P(=O)(O)O)P(OO)(O))

1721 CCN(CC)C(OO)c1ccccc1CCC2=CCC2

1722 ON(N)CCSc1ccccnccncccccc1

1723 COc1cc2c(Oc3ccc(Nc4ccc(cc4CC(C)=C)N)cc3)ccnc2cc1OCCNCC

1724 Nc1nc(N)c2c(Cl)c(NNc3ccc(cc3)C(=O)CC(CCC(=O)O)C(=O)O)ccc2n1

1725 COc1ccccc1N2CCN(CCCCNC(=O)c3cc4cc(I)ccc4s3)C2

1726 O=C1Nc2ccccc2C=C1C

1727 CN1CCc2c(C1)sc3N=CN(CCN4CCN(CC4)c5cccc6nccnc56)C(=O)c23

1728 COC(=O)C1=C(C)NC(=O)N(C1c2ccc(F)c(F)c2)C(=O)NCCCC3CCN(CC3)c4ccccc4C

1729 CC(N)Cc1c[nH]c2ccccOccc12

1730 CN[C@H](Cc1ccccc1)C(=O)N2CCC[C@H]2C(=O)N(CCCCN(C(N)N)CC=O)c3nc4CCCCc4s3

1731 NS(=O)(=O)Oc1ccc(NC(=O)NCCc2ccccccccccc2)cc1

1732 C[C@H](N)Cc1c[nH]c2cc3ccCCcc3c12

1733 CN(C)CC1(CCOC(O)C1)c2ccc(Cl)c(Cl)c2

1734 COc1ccccc1N2CCN(CCCCNC(=O)NCc3ccccc3c4ccccc4C)cCC2

1735 CC(C)n1nc(c2ccc(F)c(F)c2)c3c(N)ncnc13

1736 Cc1cc2CCN(CC=O)ccccccncccc2cc1C

1737 OB(O)c1ccccc1O

1738 O=C1CC2(CCCC2)CC(=O)N1CCCC=CN3CCc4ccccc4C3

1739 O[C@@H](CNCCCN1CCC(CC1)CC(=O)Nc2ccccc2c3ccccc3)c4ccc(O)c5NC(=O)C=Cc45

1740 Cl.COc1cc2ncnc(Ncccccccccccccc2cc1)

1741 CC(C)c1nc(nc(c2ccc(F)cc2)c1\\C=C\\[C@@H](O)C[C@@H](O)CC(=O)O)N(N)S(=O)(=O)C

1742 COc1ccc(\\C=C/c2cc(OC)c(OC)c(CC)c2)cc1O

1743 CCCCCC=c1ncnc1C2=CCCN(C)C2

1744 N[C@@H]1CC[C@H](CC1)CC(=O)c2cc(Oc3ccc(cc3)C(=N)N)cc(Oc4ccc(cc4)C(=N)N)c2

1745 CCCCCCC(C)(C)c1cc(O)cc(=CCCCCCCC(=O)NC2CC2)cc1

1746 Cc1ccc(cc1)C(=O)CNCN2CCN(CC2)c3ncccn3

1747 NS(=O)(=O)Oc1ccc(C)cc1 1

1748 COc1cc(C[C@@H]COCN)c(OC)cc1

1749 NS(=O)(=O)Nc1ccc(NC(=O)CCCCCCCC)cc1

1750 CCCc1nc(C)c2C(=O)SC(=Cn12)c3ccccc3OCC

1751 CCc1nc(N)cc(N)c1c2ccc(CC)cc2

1752 COc1cc2c(Oc3ccc(NC(=O)C4=NN(c5ccccc5C4=O)c6ccccc6C(F)(F)F)cc3F)ccnc2cc1CCCCN7CCC[illegible]

1753 Nc1nc(N)c2nc(CNc3ccc(cc3)C(=O)ON(CCNC(=O)c4ccccc4C(=O)O)C(=O)O)cnc2n1

1754 C[C@H](N)Cn1ccc2ccccccncccc12

1755 COc1ccc(cc1)C2=C(CCc2)c3ccc(cc3)S(=O)(=O)C

1756 Cc1cc2C(=O)Ncccc3cc3Nc2nn1

1757 [S+].[Sn]C(=S)NCCC1CNS=1

1758 Cc1ccc(COc2ccc3c(c2)c(SC(C)(C)C)c(CC(C)(C)C(CO)O)n3Cc4ccc(cc4)c5ccc(F)cn5)nc1

1759 CC(C)(C)OC(=O)N[C@@H](C(=O)N1CCC[C@H]1C(=O)N[C@@H](CCCNOC(N)N)C=O)c2ccccc2

1760 Brc1c(NC2=NCCN2)ccc3NCNNc13

1761 Cc1ccc(cc1)C2=CC(C)(C)Sc3cc4ccc(cc4cc23)c5ccc(cc5)C(=O)

1762 COc1cc(cc(OC)c1OC)C2=C(C(=O)NC2=O)c3c[nH]c4ccc3Bcccc4

1763 C\\N=C\\1/Cc2ccccc2C=C1

1764 Brc1c(NC2=CCCN2)ccc3nccnc13

1765 CCCCOC(=O)c1ccccc1NNC2=NC=N2

1766 Clc1ccc(cc1Cl)S(=O)(=O)NCCCCN2CCC(CC2)c3noc4ccccc34

1767 COc1ccc(CCNSC=O)(=O)Nccc1

1768 Cl.NC[C@H]1C[C@@H]1c2ccccc2I

1769 C[C@H](N)Cn1ccc2cccccccccc12

1770 CC(C)(F)CC(=O)N[C@@H]1CC[C@@H](CCN2CCC(CC2)c3coc4ccccc34)CC1

1771 Nc1nc(N)c2c(CN)ccc(c3ccc(cc3)C(=O)CC(CCC(=O)O)C(=O)O)ccc2c1

1772 COc1ccc2c(nccc2c1OC)[C@H]3CCccccCCCCcccc3

1773 CCC(CC)n1c(C)cc2c3c(N)cc(N)nc3ccc12

1774 COc1ccccc1N2CCN(C\\C=C\2CNC(=O)c3ccc(cc3)c4ccccn4)CCC

1775 CNC1=Nc2cccc(Cl)c2C(N)C1

1776 CC1(C)Nc2ccc(cc2C(C)(C)C1=O)c3csc(c3)CCN

1777 CSc1ccc2Sc3ccccc3CC(C4CCN(C)CC4)c2c1

1778 N#Cc1ccc(cc1)[C@@H]2CCCc3cnnn23

1779 COc1cc2c(Oc3ccc(cc3C)C4=CN=C(Cc5ccccc5)c(C)C4=O)ccnc2cc1OCCCN6CCOCC6

1780 CCn1nnc(n1)[C@H]2O[C@H]([C@H](O)[C@@H]2O)n3cnc4c(NC)cc(CC)n43

1781 COc1ccc(cc1Oc2CCCC2)C3CNC(=O)C3

1782 Nc1ccc(c1SSSO)(=O)NN

1783 CCCN1CCO[C@H]2[C@@]1OOc3ccc(O)cc23

1784 Fc1ccc2c(noc2c1)C3CCN(CCCNS(=O)(=O)c4ccc34)CC

1785 CCOC[C@H]1[C@@H]2CCC[C@]12c3ccc(Cl)c(Cl)c3

1786 Fc1cc(ccc1[C@H]2CCc3cncn2)c3

1787 N\\N=C\\1cOc2ccccc2C=C1

1788 Nc1ncnc2NCCC(=Nc12)c3ccc(NC(=O)Nc4cc(ccc4F)C(C)(F)F)cc3

1789 COC(=O)C(C1CC#CN1)c2ccccc2

1790 C[C@H](N)Cn1ccc2ccc3ccccc3c12

1791 OC1=C(Oc2cc(O)cc(C)c2C1=C)c3ccc(O)c(O)c3

1792 N#Cc1ccc(cc1)c2n[nH]c3c2cccccc(OCCNCC)OCCnccc3

1793 Oc1c2ccccc2c3N=Nc4cccc1c43

1794 CN[C@H](Cc1ccccc1)C(=O)N2CCCC[C@H]2C(=O)NC(CCCN=C(C)N)C(=O)n3nc4nccc4ss3

1795 COc1ccc(cc1)S(=O)(=O)NC(C)C(=O)NON

1796 CC(=O)Nc1ccc2c(CCCCCCC(Cc4ccccc4CCC))coc2c1

1797 CCCNC(=O)c1c2CCOCc2ss1NC(=O)CC4CC5CC(CC4C5)CCCC

1798 Cc1ccc2c(cccc2n1)N3CCN(CCc4cccc5c4OCc6c(ncn56)C(=O)C7OCOCC7)C3

1799 CN1CCc2c(C1)c3cc(C)ccccn2CCc4cccc34

1800 CS(=O)(=O)c1ccc(cc1)c2ccccc2 2

1801 CN(C)CC1CC2N(O1)c3cccc(O)c3Cc4ccccc24

1802 CCc1nc(N)nc(N)c1c2ccc(CC)cc2

1803 CC1CCN(CC(=O)Nc2nnc(s2)S(=O)(=O)N)OC1

1804 CC(=O)NC1CCc2cc(CCc3CCN(CC3)c4nsc5ccccc45)ccc12

1805 COc1ccccc1C(C)NS(=O)(=O)NN

1806 CN[C@H](Cc1ccccc1)C(=O)N2CCC[C@H]2C(=O)N[C@@H](CCCC=C(N)N)C=O

1807 COc1ccccc1N2CCN(CCNSS(NO)(=O)c3ccc4ccccc4c3)CC2

1808 CN1CCc2c(C1)sc3NCCN(CCN4CCN(CC4)c5cccc6nccnc56)C(=O)c23

1809 NS(=O)(=O)c1cccc(Nc2ncc(BO)c(Nc3ccc(OCCCN)cc3)n2)c1

1810 OP(=O)(O)C(CC1CCCCC1)P(=O)(O)N

1811 COc1cc2nc(nc(N)c2cc1OC)N3CCN(CC3)C(OC)C4CCSS4

1812 CC1=CC[C@@H]2[C@@H](C1)c3c(O)cc(cc3OC2(C)C)C(C)CCCC4CCCCCC4

1813 COC(=O)CCc1cncn1Cc2ccc(cc2)CNN

1814 Cc1cccc(c1)N2CCN(CCCCNC(=O)c3oc4cnccc4n3)CC2

1815 Clc1ccccc1N2CNN(CCCCOc3ccc4CCC(=O)Nc4n3)CC2

1816 CC1NC(=Nc2cccc(CC)c12)N
1817 COc1cc(C=C2CC(=S)NC2=O)ccc1O
1818 Cl.CC(C)Sc1ccccc1NCC2=CNCN2
1819 CC(C)c1cc(Oc2c(Cl)cc(CC(=O)O)cc2C)Bccc1O
1820 Clc1ccc(cc1)C3C2CC(C2C)CCCCCN3
1821 COc1cc2c(Oc3ccc(cc3F)C4=CN=C(Cc5ccccc5)N(C)C4=O)ccnc2cc1OCCCN6CCOCC6C
1822 CC(C)Oc1ccccc1N2CCN(CC2)C3CCC(CC3)NC(=O)NCc4cc=c(F)c4
1823 Clc1ccc(OCCCCCCOc2ccccc2)cc1
1824 O=C1N(CCN2CCCCc2)CCN1c3ccccc3
1825 CC(C)(C)n1nc(c2ccc(Cl)cc2)c3c(N)ccnc13
1826 CC(Oc1ccccc1C2CcCc2)C3CNCCN3
1827 Cc1ccc2c(cccc2n1)N3CCN(CCc4cc5Nc(=O)COc5cc4F)Cc3
1828 C[C@]12CC[C@H]3[C@@H](CCc4cc(O)ccc34)[C@@H]1CC[C@@]2(O)CCC
1829 CC(C)Oc1ccccc1N2CCN(CC2)[C@@H]3CC[C@@H](CC3)SS(=O)(=O)c4ccc(C)c(Cl)c4
1830 O=C(Nc1ccc(c2ccccc2)c(n1)c3occc3)C4CC4
1831 Cc1ccccc1Oc2ccc(cc2CC)C#N
1832 Cl.NCC1CC1c2cc(C)c(O)cc2OCC=C
1833 Cl.N#Cc1ccc(cc1)C2CCCc3cnnn23
1834 Oc1ccc2C(N(CCc2c1)c3ccccc3)c4ccc(OCCCCCC5CC5)cc4
1835 CC1=CC[C@@H]2[C@@H](C1)c3c(O)cc(cc3OC2(C)C)C4CCSCC44C5CCCC45
1836 CCNC(=O)NCc1oc2ccccccc1OCc1ccc2c1
1837 C(Oc1ccccc1c2ccccc2)C3CNCCN3
1838 COC(=O)C1=C(C)NC(=O)N(C1c2ccc(F)c(F)c2)C(=O)NCCCN3CCC(CC3)c4ccccc4OO
1839 OC[C@@H]1[C@@H](O)[C@H](O)[C@@H](C)CC1CCCCCOCc2ccc(c3ccccc3)c(c2)C(F)(F)F
1840 COc1cc2c(Oc3ccc(cc3F)C4=CN=C(Cc5ccccc5)N(C)C4=O)ccnc2cc1OCCCC6CCOCC6C
1841 OC[C@H](Nc1CCc(CCCc2c[nH]c3ccc(cc23)n4cnnc4)CC1)c5ccccc5
1842 Nc1ncnc2c1c(cn2C3CCCC3)C#CN
1843 Nc1n[nH]c2cccc(c3)cc(NC(=O)Nc4ccccc4ccc3)c12
1844 Cc1ccc2c(cccc2n1)N3CCN(CCc4ccc5OCCC=O)Nc5c4CCC3
1845 Ic1c(NC2=NCCN2)ccc3ccccc13
1846 COc1cc(C=C2SC(=Nc3ccccc3)cC2=O)ccc1O
1847 Nc1c2CCCCc2nc3cccc(CC)c13
1848 CC(C)n1nc(c2cnc3cc(O)ccc3c2)c4c(N)ncnc14
1849 NS(=O)(=O)c1ccc(CC)cc1
1850 CNCCc1c[nH]c2ccc(cc12)C(CCCC)C

1851 CC(C)C(N(Cc1ccccc1)S(=O)(=O)c2c(F)c(F)c(F)c(F)c2(C))(O)NO

1852 Fc1ccc2c(cccc2c1)N3CCN(CCCCc4ccc5CNC(=O)c5c4F)CC3

1853 NNc1ccccc1S(=O)(=O)

1854 NC(=N)c1ccc(CNC(=O)[C@H](CCC2CCNCC2)NC(=O)[C@@H](NCC3CCNCC3)NS(=O)(=O)Cc4ccccc4)[illegible]

1855 Clc1cc(NC2CCCCC2)nc(n1)c3c[nH]c4ccccc34

1856 COc1ccc2[nH]cc(OCN)c2c1

1857 O=C1NC(=O)\\CC=C\\c2ccc3ncccc2c3cnOC1

1858 COc1ccccc1N2CCN(CCCCCC(=O)NCc3ccccc3c4ccccc4C)CCC2

1859 OCc1cccc(OCC(O)CNC2CCN(CC2)c3ccnc4scc(c5ccccc5)c34)c1

1860 CNC(=O)[C@H]1O[C@H]([C@H](O)[C@@H]1O)n2cnc3c(NOC)nc(nc23)CCCc4ccccn4

1861 CC(C)n1nc(CCCc2cccc(O)c2)c3c(N)ncnc13

1862 NC(F)(F)c1ccc2c(NC(=O)Nc3ccccn3)ccnc2c1

1863 CCCn1cc(C(=O)c2ccc(C)c3ccccc2)cccccccc13

1864 COc1ccccc1N2CCN(C\\C=C\\CNC(=O)c3ccc(cc3)c4ccccn4)CC2 2

1865 Nc1ncnc2c1c(cn2CcCCcC2(C)C)2

1866 COc1ccc(cc1)C2=S(CCC2)c3ccc(cc3)S(=O)(CO)N

1867 CC(C)Nc1cccc(NC(SO)c3cccccc(cccccc3)(=N)N)c1

1868 Clc1cc2NC(=O)Cc2cc1CCN3CCC(C)3Cc4nnc5cccccc54

1869 C(Nc1ccccc1c2cccs2)C3CNNCN3

1870 COC(=O)C1=C(C)NC(=O)N(C1c2ccc(F)c(F)c2)C(=O)NCCCN3CCCCCC3Cc4ccccc4CO

1871 COc1ccccc1N2CCN(CCCCNC(=O)c3cn4ccccc4n3)C2

1872 CS(=O)(=O)c1ccc(cc1)c2=C(C(CO)OC2)c3ccccc3

1873 Cc1cccc2N3NCNCC3NC(ON)c12

1874 CC(C)Oc1ccccc1N2CCN(CC2)[C@@H]3CC[C@@H](CC3)NS(=O)(=O)c4ccc(Cl)cc4CC

1875 COc1ccccc1N2CCN(CCCCCC(=O)NCc3ccccccc4cc4cc3C)CCC2

1876 COc1ccc(cc1)C(=O)CCCN2CccccccccCC2

1877 CCCN(CCC)[C@@H]1CCc2ccc(O)c2cC1

1878 COc1cc2ncnc(Nc3cccc(BB)c3)c2cc1OC

1879 Cc1cc(c(O)c(C)c1CC2=CCCC2)C(C)(C)C

1880 CC(C)Oc1ccccc1N2CCC(CC2)[C@@H]3CC[C@H](CC3)NS(=O)(=O)c4cccc(F)c4

1881 CC(=O)O.OCc1cc(ccc1O)[C@@H](O)CNCCCCCCCCCCCc2ccc(cc2)N3C(=O)CNC3=O

1882 Clc1ccc2[nH]cc(OCN(CC=C)CC=C)c2c1

1883 NC(=O)N(O)CCC#Cc1ccc(OCCC2CCN(CC2)[C@H](c3ccccc3)c4ccc(Cl)cc4)cc1

1884 O=C(Nc1cnccn1)Nc2cccc3ccccc23

1885 Clc1ccc(CNC2=C(Nc3ccncc3)C(=O)c2OO)cc1

1886 NS(=S)(=O)O
1887 Oc1ccc(Oc2c(Cl)cc(cc2Cl)N3NCCC(=O)NC3=O)cc1C(=O)NCCCCCCl
1888 COc1ccc(NC(=O)Nc2ccc(cc2)c3csc4c(cnc(N)c34)c5ccn(CCO)c5)cc1
1889 CCc1nc(c2cccc(C)c2)c(s1)c3ccnc(NC(CO)c4ccccc4)c3
1890 NC(=O)c1cc([nH]c1c2ccc(Cl)cc2F)c3cccc(N)n3
1891 Clc1cccc(Cl)c1C(=O)Nc2c[nH]nc(C2NO)NC3CCNCC3
1892 Clc1ccc2Oc3ccccc3N=C(N4CCCC4)c2c1
1893 Cl.NCC1CC1c2cc(F)c(F)cc2CCCOC
1894 CN[C@H](Cc1ccccc1)C(=O)N2CCC[C@H]2C(=O)NC[C@@H]3CC[C@@H]NNCCC3C
1895 CN1CCN(CC1)C2=Nc3ccccc3cc4cc(C)cc24
1896 Clc1cccc(c1)CCCC2CC2
1897 Cc1noc(n1)C2CN3nCC2C3
1898 C(Nc1ccccc1c2ncccc2)C3=NCCN3
1899 COc1cc2c(Oc3ccc(NC(=O)N\\N=C\\c4cccc(CCCC)c4F)cc3F)ccnc2cc1CCCC5CCCCCC5
1900 NC(=N)NCCC[C@@H](NS(=O)(=O)Cc1ccccc1)C(=O)N2CCC[C@H]2C(OO)NCc3ccc(cc3)C(=N)N
1901 COc1cc2nc(nc(N)c2cc1OC)N3CCN(CC3)C(CC)CCCCCCCCC
1902 NC(=O)c1cc2c(cncc2s1cc3ccc(B3)c3F)c3F
1903 COC1(CC1)C(OO)N[C@@H]2CC[C@@H](CCN3CCN(CC3)c4cccc5OCOc45)CC2
1904 CC(N)Cc1c[nH]c2cccnc12
1905 COc1cccc(c1)C#Cc2ccc(cc2)S(=O)(=O)
1906 COc1ccc2c(c1)c(CC(=O)O)c(C)C2C(=O)c3ccc(Cl)cc3
1907 COCC(=O)N[C@@H]1CC[C@@H](CCC2CCC(CC2)c3coc4ccccc34)CC1
1908 NC(=N)c1ccc(CNC(=O)[C@@H]2CCCN2C(=O)[C@@H](NCC(=O)N)C(c3ccccc3)c4ccccc4)s1
1909 CN1CCC(CC1)c2ccc(cc2)C3N(Ccccc(O)ccc3N)c5ccc(F)cc5
1910 Cc1c(sc2ccc(F)cc12)S(=O)(=O)NCCCCC3CCC(CC3)c4noc5cc(F)ccc45
1911 CCn1nnc(n1)[C@H]2O[C@H]([C@H](O)[C@@H]2O)n3cnc4c(NC)nc(CC)nc34
1912 Fc1cccc(c1)c2cncn2Cc3ccc(nc3)C#N
1913 Fc1ccc(cc1)C(=O)C2CCN(CC3CC(=O)c4ccccc4C3)CC2C
1914 NS(=O)(CO)O
1915 Cc1cccc(c1)C(=O)CCN2CCN(CC2)c3ccccc3C#N
1916 Fc1ccc2[nH]cc(CCCN3CCC(CC3)c4ccccc4)c2c1
1917 CC(C)Oc1ccccc1N2CcN(Cn2)C3CCC(CC3)NC(=O)Nc4cc(F)ccc4F
1918 NCCc1cccc2c1ccn2S(OO)(=O)c3ccccc3
1919 CN1C(=O)C=NN(CCCCN2CCN(CC2)c3ccccc3CC=C)C1=C
1920 CC1=C2CCc3cc(C)ccc3N2CCC1=N

1921 Fc1ccccc1N(Cc2ccc(Cl)cc2)C3CCC3

1922 CC(Oc1ccccc1c2cccs2)C3=NCCC3

1923 Cc1cccc2NCCCNCCCNC(=O)c12

1924 CCOc1nc(cc(N)c1C#N)C(=O)OCc2ccccc2S(=O)(=O)NO

1925 CC(NC(=O)c1ccc(cc1)c2ccccc2)c3ccccc3

1926 COc1ccccc1N2CCN(CC(O)CCNC(=O)c3cncccccccc3)CC2

1927 COc1c(Cl)cccc1N2CCN(CCCCNC(=O)c3c4cccccccc34C)2

1928 CN1CCC(=C2c3ccccc3CCc4sccc24)CCC1

1929 Cl.CCOC(=O)CCc1ccc(OC[C@H](O)CCC(C)(C)Cc2ccc3ccccc3c2)c(c1)C#N

1930 COc1cc2c(Oc3ccc(Nc4ccc(cc4)C(C)(C)C)cc3)ccnc2cc1OCCCCCO

1931 Cc1ccc(Cn2cncc2c3ccccc3)cc1

1932 CN1CCN(CC1)C2=cc3ccccc3cc4sc(C)cc24

1933 COc1cc2c(Oc3ccc(NC(=O)N\\N=C\\c4cccc(CC=O)c4O)cc3F)ccnc2cc1OCNCN5CCCCC5

1934 NS(=O)(=C)c1ccc2CCCc2ccc1

1935 Clc1cc2N(CCc2cc1N)C(=O)Nc3cccnc3

1936 CN1C(=O)C(C)c2nc[nH]c2c1=O

1937 NC(=O)c1cccc(c1)c2cccc(CC(=O)NC3CCCC3)c2

1938 Cc1ccc2c(cccc2n1)N3CCN(CCc4cccc5c4OCc6c(nnn56)C(=O)N)CC3

1939 CCCN1C(=O)N(CCC)c2nc([nH]c2C1=O)c3ccc(OCC(=O)NCCC)cc3

1940 Fc1ccc(cc1)C(=O)CCCN2CCCNCCC2Cc3ccc(Cl)cc3

1941 COc1ccc(cc1)C2CN(C)Cc3cc(OCCC(CCCCC))ccc23

1942 CC1NC(=Nc2cccc(CN)c12)

1943 C(CN1CCC(Cc2ccccc2)CC1)Cc3c[nH]c4ccc(nc34)n5cnnc5

1944 OC1=C(Oc2cc(O)cc(O)c2c1=O)c3ccccc3

1945 Clc1cccc(Oc2ncccc2C3CCNCC3)c1C

1946 CCCCC(CC(=O)NO)S(=O)(=O)c1CCCCC1N

1947 Clc1ccc(cc1)N2CCN(CCN3CCCCCC3)C2CO

1948 CC(C)c1ccc(cc1)N2cCc3cc(O)ccc3C2(C)c4ccc(OCCN5CCCC5)cc4

1949 CN(C)C(=O)N[C@@H]1CC[C@@H](CCN2CCC(CC2)3ccccc4OCOc34)CC1

1950 COc1cccc(c1)C(C)NC(=O)c2ccc(cc2NCCCC)c3ccncc3

1951 CCCNC1CCc2cccccccN2C1

1952 COc1cc2c(Oc3ccc(NC(=O)N\\N=C\\c4cccc(CC=C)c4F)cc3F)ccnc2cc1OCCC5NCCCCC5

1953 Nc1ncnc2c1c(cn2C3CnnC3)C#C

1954 Cc1ccc2c(cccc2n1)N3CCN(CCc4ccc5NC(=O)COc5c4)cC3

1955 NS(=C)(=O)O

1956 Nc1ccc(cc1O)c2nn(C3CCCC3)c4ncnc(C)c24
1957 Clc1cccc(N2CCN(CCCCOc3ncc4C=NC(=O)Nc4n3)CC2)c1CC
1958 Clc1cccc(c1)N2CCCCC2
1959 COc1ccc(cc1OC2CCCC2)C(=O)Nc3c(F)cccc3C
1960 OP(=O)(O)C(CN1cccccn1)P(=O)(O)O
1961 COc1ccccc1N2CCN(CCCCN3N=CC(=O)N(C)C3OO)CC2
1962 COc1ccc2ccc3Oc(CCNC(=O)C4CC4)cc3c2c1
1963 Nc1ncnc2c1c(cn2CCCCN(CCC)CCO)c4cccc(O)c4
1964 Nc1ccc(cc1F)S(=O)(=O)
1965 Cc1ccc(NC(=O)Nc2ccc(cc2)c3csc4nnnc(N)c34)cc1C
1966 Cl.NCC1CC1c2cc(C)c(F)cc2CCC=CC
1967 Cl.NC[C@H]1C[C@@H]1c2ccc(F)cc2CN
1968 ONC(=O)CCCCCCC(=O)Nc1ccc1 1
1969 S=C(NC1CCCCC1)N2cCC(CC2)c3cnc[nH]3
1970 Cc1cccc(CC(=O)Nc2ccc(cc2)c3csc4c(cnc(N)c34)C#CCN5CCCC5)c1
1971 Fc1ccc(cc1)N2CCN(CCCCNC(=O)c3cc4ccccc4cc3)CC2
1972 Clc1cccc(c1)N2CCN(CC3CCCCC3)C2CO
1973 C(Oc1ccccc1c2ccccc2)S3=NCCN3
1974 COc1cc2ncnc(N[C@H](C)c3ccccc3)c2cc1OCCCCCCC(=O)NOO
1975 CC(O)Cn1cc(nn1)c2cnc(N)c3c(csc23)c4ccc(NC(=O)Nc5cccc(F)c5)cc4
1976 Cc1cc(c(O)c(C)c1NC2=CCCC2)C(C)(C)CC
1977 CNS(=O)(=O)c1ccccc1Nc2nc(Nc3cc(OC)c(OC)c(CC)c3)ncc2Cl
1978 OC(CNC1CCN(CC1)c2ncnc3scc(c4ccccc4)c23)COc5cccc(c5)CNN
1979 COc1cc(ccc1Nc2ncc(Cl)c(Nc3ccccc3P(=O)(CCC)n2)N4CCC(CC4)NCC)C
1980 CSc1ccc2Sc3ccccc3CC(C4CCC(C)CC4)c2c1
1981 NC(=N)c1ccc(CNC(=O)[C@@H]2C=CCN2C(=O)[C@@H](CC3CCCCC3)NCC(=O)O)s1 C
1982 Cc1cccc(NC(=O)Nc2ccc(cc2)c3cccc4[nH]ccc34)c1
1983 Nc1nc(N)c2c(CN)c(C(c3ccc(cc3)C(=O)NC(CCC(=O)O)CC=O)O)ccc2n1
1984 COc1ccccc1N2CCN(CCCNC(=O)c3ccn4ccccc34)CC2
1985 O=C(NCCCCN1CCN(CC1)c2ccccc2)c3cccc4ccc4cn3
1986 Cc1cccc(NC(=O)Nc2ccc(cc2)c3cncc4c3c(N)nnc4)c1
1987 Brc1ccc2[nH]cc(CCC(CC=C)CC=C)c2c1
1988 C[C@H](CCN1CCC(C)CC1)N(C)S(=O)(OO)c2cccc(C)c2
1989 CN(CCOc1ccc(CC2SC(=O)NC2OO)cc1)c3ccccn3
1990 Cl.CCSCC(C1NCNCC1)c2ccc(Cl)c(Cl)c2

1991 CN1CCC(=C2c3ccccc3COOc4ccccc24)CC1
1992 CCN(CC)C(=O)c1ccccc1NCC2NNCCN2
1993 CC1(N(CCc2cc(O)ccc12)c3ccccc3)c4ccc(OCCC5CCCCC5)cc4
1994 Nc1n[nH]c2cccc(c3ccccNc(=O)Nc4cc(ccc4F)C(=O)Occc3)c12
1995 CN1N=C(S/C/1=C/C(=O)C)S(=O)(=O)N
1996 CNc1cc2c(Nc3cccc(BB)c3)ncnc2cn1
1997 COc1ccccc1Nc2nc(Nc3ccc(cc3OC)N4CCC(CC4)N5CCC(C)CC5)ncc2Cl
1998 COc1nccn2c(c3ccnc(NCC(C)(C)CC)n3)c(nc12)c4ccc(F)cc4
1999 CCC[C@@H]1NC(C)(C)CC[C@@]1(O)c2cccc(Cl)c2
2000 CCCCCCCCC(=O)C(=O)CCCCCCC
2001 Oc1ccc2C3=C(CCOc2c1)c4ccc(O)cc4O[C@@H]3c5ccc(CCCC6CC)CC66cc56
2002 Cc1ccc(cc1)C2=CC(C)(C)Oc3cccccc(ccccc23)c5ccc(cc5)C(=O)O
2003 COc1ccccc1CNCCCCCCNCCCCCCNCCCCCNCc2ccccc2C
2004 OCCNc1cc2cc(cc2ccn1)c3cccnn3
2005 OCc1cc(ccc1O)C(O)CCCCCCCOCCCCc2ccccc2
2006 COc1ccccc1N2CCN(CCCCNC(=O)c3ccc(B)Bcc3)CC2
2007 Clc1cccc(c1)CCC=2CC2
2008 COc1cc2c(Oc3ccc(NC(=O)C4=C(Cl)c5ccccc5N(C4=O)c6ccc(F)cc6)cc3F)ccnc2cc1OCCCN7CCNC[illegible]
2009 CC(C)Oc1ccccc1N2CCNNCc3cccc(c3)C(=O)C4CCCCC4CCC2
2010 CCCCC(CC(=O)NO)S(=O)(=O)c1CCCCCC1
2011 CN(C)S(=O)(=O)NC1CCc2ccc(C)cc2O1
2012 [S+].[Sn]C(=S)NCCC1CNOC1
2013 CC(C)NCC(O)c1ccc(O)c(O)1=O
2014 OP(=O)(O)C(Cc1cccnc1)P(=O)(O)
2015 NS(=O)(=O)c1ccc(CCC=O)cccccccccccccc1
2016 CCCCCCCCCC=CCC(=O)CCCCOCCC
2017 CCc1cc2[C@@H]3CNCCN3C(=O)c2cc1
2018 CC(C)NC(C)C(O)COc1ccc(C)cCCCCc1
2019 Brc1c(NC2CNCCN2)ccc3nccnc13
2020 Brc1ccc(CN(c2ccc(cc2)CCN)n3cncc3)cc1
2021 CCCCC(CC(=O)NO)S(=O)(=O)c1CCCCC1
2022 COc1ccc(cc1)CNCN(C)CCC2(O)CCCCC2
2023 CC1NC(=Cc2cccc(CN)c12)N
2024 Clc1cccc(Nc2nnnc3ccccc23)c1
2025 COc1cccc(F)c1C2cNCc3cnc(Nc4ccc(C(=O)O)c(OC)c4)nc3c5ccc(Cl)cc25

2026 OCC(N1CCN(CCCc2c[nH]c3ccc(cc23)n4cncc4)CC1)c5ccccc5
2027 COc1ccc(cc1)C(NO)CCCN2CCc3ccccc3C2
2028 COc1cc2nccc(Oc3ccc(NC(=O)C4=C(C)N(C)N(C4=O)c5ccccc5)cc3F)c2cc1CO
2029 NS(=O)(=O)c1cc2ccc(OCS)ccc2n1
2030 CCCOc1cc2c(cc1\\C(=C/C=C/C(=C/C(=O)O)/C)\\C(CCC)CC)CCC2(C)C
2031 CC(=O)N1CCC(CC1)NC(=O)NC2CCCCO2
2032 Cc1ccc2c(cccc2n1)N3CNN(CCc4cccc5c4ccc6c(nnn56nn(CO3)))C
2033 Fc1cccc(Oc2cccc(C)c2C3CCNCC3)c1
2034 Clc1ccc(cc1)C(=O)CCCN2CCOCC2
2035 NCCNc12ccc2ccccc1
2036 Cl.NC[C@H]1C[C@@H]1c2ccccccc2
2037 CC(C)Oc1cc(ccc1C(=O)O)c2ccc(CCCC[C@H](C)c3ccccc3)cc2
2038 COc1cc2ncnc(Nc3ccc(F)c(Cl)c3)c2cc1OCCCCCCC(=O)NOO
2039 Clc1cccc(N2CCN(C\\C=C\\CNC(=O)c3ccc(cc3)c4ccccn4)CC2)c1C
2040 Ic1cCNC2CNCCN2Nccc3ccccc13
2041 O=C(N[C@@H]1CC[C@@H](CCN2CCC(CC2)c3coc4ccccc43)CC1)c5ccc(cc5)n6cccc6
2042 CCN(CC)CC#CCOC(=O)C(C)(C1CCCCC1)c2ccccc2
2043 NS(=O)(=O)Oc1ccc(cc1Cl)c2cc(Cn3cncc3)ccc2C#N
2044 NS(=O)(=O)c1ccc(CCCCO)cccccccccccccccccc1
2045 Cc1cccnc1CC(P(=O)(O)O)P(=O)(O)O
2046 C(Oc1ccccc1c2cccs2)CC=NNCN
2047 NCCCCN(O)O
2048 Cc1ccc(Cl)c(c1NN2CCN(CCC3C(=O)C4)CCCCC4)CCCC33CC23
2049 COc1ccccc1N2CCN(CC3=O)Nc(cccc(C)c3)CC2
2050 O=C(NCCCCN1CCN(CC1)c2ccccc2)c3cc4cccccc4n3
2051 OC1CCC(CC1)Nc2cc(Cl)cc(n2)c3c[nH]c4ncccc34
2052 CN(C)CCCN1c2ccccc2CCc3ncccc13
2053 COc1ccc(cc1)S(=O)(=O)c2ccc(cc2)C(=C)C3CCN(CC3)C4CCN(CC4)C(OO)OC(C)C
2054 CCN(CC)C(OO)c1ccccc1CCC2NCCC2
2055 Fc1ccc2c(coc2c1)C3CCN(CCCNS(=O)(=O)c4cccs4)CC3
2056 Oc1cc2CC[C@H]3OCc4ccccc4[c@@H]3c2cc1
2057 C[C@H]1CNCCc2ccc(CC)cc12
2058 COCC(=O)N[C@@H]1CC[C@@H](CCN2CCN(CC2)c3coc4ccccc34)CC1
2059 COc1ccc(cc1OC)C(C)c2nc(C)nc3oc(C)cc23
2060 Clc1ccc(cc1Cl)C(=O)C(CCCC)N2CCCC2

2061 COc1ccc2CC(CCc2c1)NS(=O)N(C)CC

2062 CSc1cc(F)ccc1NCC2=NCC2C

2063 COc1ccc2CC(CCc2c1)NC(=O)N(C)CCC

2064 FC(F)(F)c1cccc(NC(=O)Nc23ccc3c(C3)cccc23)n1

2065 COc1ccccc1N2CCN(CCCCCNC(=O)c3ccn4ccccc34)CC2

2066 CCCc1c(nnn1Cc2ccccc2)C(=O)NCCCCNNNCCCCCCCc4cccc(Cl)c4Cl

2067 CC(N)C1CCC(CC1)C(=O)Oc2ccccc2

2068 CC(C)n1cc(c2ccc3[nH]ncc3c2)c4c(N)ncnc14

2069 NS(=O)(CN)N

2070 Nc1c2CCc(c2)C3ccccc1c3

2071 Cc1cc2C(=Nc3ccccc3NC2c1)c4CNCCC4

2072 Brc1c(NC2NNCCN2)ccc3nccnc13

2073 COc1cc2ncnc(Nc3cccc(c3)C#C)c2cc1OCCCCCCC(OO)NO

2074 COc1ccccc1N2CCN(CCCCNC(=O)c3nnn4ccccc34)CC2

2075 C(CN1CCCC1)Cc2ccccOc3ncccc3ccc2

2076 CCCOc1cc2c(cc1\\C(=C/C=C/C(=C/C(=O)O)CC)\\C)C(C)(C)CCC2(C)C

2077 C(CN1CCC(Cc2ccccc2)Cc1)Cc3c[nH]c4ccc(cc34)n5cnnc5

2078 CNCC(N1CCCC1)C(=O)c2ccc3ccccc3c2

2079 COc1cc(CC(C)N)c(OC)cccc1

2080 CC(C)Oc1ccccc1N2CcN(CC2)C3nCC(CC3)NC(=O)Nc4cc(F)ccc4F

2081 NCCc1ccnc2c1ccn2S(=O)(=O)c3ccccc3

2082 CCCCC(CC(CO)NO)S(=O)(=O)c1CCCCC1N

2083 Cl.CSc1ccccc1NCC2=NCCC2

2084 Oc1cccc2CC[C@@H]3[C@H](CCC3CC=C)c12

2085 COc1ccccc1CNCCCCCCCNCCCCCCCNCCCCCCNCc2ccccc2O

2086 Brc1ccc2[nH]cc(CCN(CC=C)CCCC)c2c1

2087 CC1=CC[C@@H]2[C@@H](C1)c3c(O)cc(cc3OC2(C)C)C4(CCCC4CC)N

2088 Cc1cccc(NCCCO)CN2CCN(CC2)c3cccccc3cNCc1

2089 COc1ccccc1N2CCN(CNC(=O)c3cccc2CC)c3CCC

2090 COC(=O)C1=C(C)CC(=O)N(C1c2ccc(F)c(F)c2)C(=O)NCCCN3CCN(CC3)c4ccccc4

2091 CC(=O)CC1CC2CCCC1N2

2092 CN1CCN(CC(=O)Nc2nnc(s2)S(=O)(=O)N)CCC1

2093 Cc1ccc2c(cccc2n1)N3CCN(CCc4ccc5CCC(=O)Nc5c4F)CC3

2094 NS(=O)(=O)c1cccc(c1)CCCc2cccc(F)c2

2095 C(COc1ccccc1)NCCCc2ccccc2OCc3ccccc3

2096 CC(Oc1ccccc1c2cccc(c2)[N+](=O)[O@])C(=CNNNC)

2097 CC(C)Oc1ccccc1N2CCN(CC2)[C@@H]3CC[C@H](CC3)NS(=O)(=O)c4ccccc4C

2098 CCn1nnc(n1)[C@H]2O[C@H]([C@H](O)[C@@H]2O)n3ccc4c(N)nc(N[C@H](CO)Cc5ccccc5)nc34

2099 CC(=O)NC1CC2CCcC1N2

2100 COc1ccccc1N2CCN(CCCSS(=O)N=O)c3ccc4ccccc4c3CCC2

2101 COc1cc2ncnc(Nc3ccc(C)c(c3)C#C)c2cc1OCCCCCCC(=O)NO

2102 Cc1ccc2c(cccc2n1)N3CCN(CCc4ccc5OCC(=O)Nc5c4O)CC3

2103 Oc1ccc2C3=C(CCOc2c1)c4ccc(O)cc4O[C@@H]3c5ccc(OCCC6CCOCC6)cc5

2104 COC(=O)c1cncn1Cc2ccc(cc2)CCO

2105 CN(C)CCCN1c2c3ccc2CCccccccc13

2106 Clc1ccc(NC(=O)c2ccccc2NCc3ccnnc3)cc1N

2107 COc1ccc2c(nccc2c1OC)[C@@]3CCccccccCcccc3

2108 Nc1ccc2ncnc(Nc3cccc(BB)c3)c2c1

2109 O[C@@H](CCCCCCCCCN1CCC(CC1)OC(=O)Nc2ccccc2c3ccccc3)c4ccc(O)c5CC(=O)C=Cc45

2110 O=C1CCc2ccc(OCNCCN3CCN(CC3)c4cccc5CCCc45)nc2N1

2111 CCCCCCCCCC(S(=O)(=O)N)

2112 FC(F)(F)c1ccccc(NC(=O)Nc3cccc(n3)C#N)ccnccc1

2113 OC1(CCN(CCCC(=O)c2ccc(F)cc2)CC1)c3ccc(CC)cc3

2114 CC(C)C[C@H](CC(=O)[C@H](Cc1ccccc1)NC(=O)c2ccccn2)B(O)O

2115 COc1cccc(c1)C#Cc2ccc(cc2)S(=O)(OO)

2116 COc1ccccc1N2CCN(CCN(C(=O)C34CCC(CF))CC3)C4cc5ccccn5CCC2

2117 COc1cc2nc(nc(N)c2cc1CO)N3NCN(CCO3)C4CCCCCCCCCC4CC

2118 COc1cc2ncnc(Nc3ccc(F)c(Cl)c3)c2cc1OCCCCCCC(=O)NOC

2119 COc1ccc(cc1)C(C)NS(=O)(=O)

2120 Fc1ccc2cccc(N3CCN(CCCCOc4ccc5CCCc5c4)Cc3)c2c1

2121 CCOC[C@H]1[C@@H]2CNC[C@]12c3ccc(Cl)(CCl)c3

2122 Clc1ccc2C(c2Cccc1C)

2123 O=C1Nc2ccccc2C=C1CC

2124 CC(=O)N1CCC(CC1CNC(=O)NCCCCCCC)

2125 CC(C)Oc1ccccc1N2CCN(CC2)[C@@H]3CC[C@H](CC3)NS(=O)(=O)c4ccccc4O

2126 CSc1ccc(F)cc1NCC2NNCCN2

2127 N[C@@H]1CC[C@H](CC1)NC(OO)c2cc(Oc3ccc(cc3)C(=N)N)cc(Oc4ccc(cc4)C(=N)N)c2

2128 Fc1ccc(OCCCCCCOc2ccc(CC)cc2)cc1

2129 COc1ccc2[nH]c(C)c(CNN(C)C)c2c1

2130 NS(=O)(=O)

2131 CNCC[C@@H](Oc1ccccc1C)c2ccccccc2
2132 NS(=O)(=O)Oc1ccc(N)cc1 1
2133 COc1ccc(cc1)C2=C(Cnc2)c3ccc(cc3)S(=O)(=O)C
2134 CCCCN1CCC(COC(=O)c2cc(CO)c(CC)c3OCCNc23)CC1
2135 COc1ccc(cc1OC2CCCC2)C3CNC(=O)c3
2136 CC1=CN=C(NCCc2ccccc2)C(=O)N1CC(=O)NCc3ccc(N)cc3CN
2137 Cc1cc2CCN(C(=O)Nc3cccnc3)c2cc1C
2138 COCC(N1CCCC1)C(=O)c2ccccccccc2
2139 Cl.COc1ccc(cc1)N(C)c2nc(C)nc3Cc(C)cc23
2140 C(Nc1cc(ncc1)c2c[nH]c3ncccc23)c4ccccc4
2141 CN(C)CC1(CCCCC1)c2ccc(F)c(CC)c2
2142 ON(C)C[C@H]1CCCC[C@]1(O)c2cccc(O)c2
2143 CC1=C2CCc3cc(C2ccc3NCCCC1=O)
2144 CCCCCOc1ccccc1c2onc(c2)C(=O)CC3CCCCCC3
2145 COc1cc2c(Oc3ccc(NC(=O)C4=C(C)N(C(=O)N4)c5ccccc5CC)cc3F)ccnc2cc1OCCCN6CCN(C)CC6
2146 Fc1c(Cl)cccc1N2CCN(CCCCOc3ccc4CCC(=O)NcC43)CC2
2147 CCCCCCCc1c2CCCCc2nc3cc(F)ccc13
2148 COc1cc2ncnc(Nc3cccc(c3)C#C)c2cc1CCCCOCCCOOCCN
2149 COc1cc2c(Oc3ccc(cc3F)C4=CN=C(Cc5ccccc5)N(C)C4=O)ccnc2cc1CCCCN6CCOCC6
2150 COc1cc2c(Oc3ccc(cc3C)C4=CN=C(Cc5ccccc5)N(C)C4=O)ccnc2cc1OCCCN6CCOCC6
2151 Cl.NCCc1ccc(O)c(C)c1
2152 Cl.Cl.OC1(CCCCC1)C(CC2CCNCC2)c3cccc(Cl)c3
2153 C(Cc1ccccc1c2cccs2)C3=NCCN3
2154 NS(=O)(=O)c1cccc(Nc2ncc(BO)c(Nc3ccc(OCC#N)cc3)n2)c1
2155 Clc1cccc(c1)N2CCC(CC3CCCC=3)C2CO
2156 CC(C)NCC(O)c1ccc(O)ccOcc1
2157 Fc1ccc(cc1)C(=O)CCCN2CCCNCCC2Cc3ccc(CC)cc3
2158 COc1ccccc1CNCCCCCCNNCCCCCCCNCCCCCCNCc2ccccc2OC
2159 COC(=O)C(Cc1CCCN1)c2ccccc2
2160 COc1ccc(cc1OC)S(=O)(=O)N[C@@H]2CC[C@H](CC2)N3CCC(CC3)c4cc(F)ccc4CC(C)C
2161 Nc1nccc(Nc2ccc(cc2)S(OO)(=O)N)c1
2162 C(ccccccc1)ccncc3CCNCCc3n1
2163 Cc1ccc2c(cccc2n1)N3CCN(CCc4ccc5CCC(=O)Nc5c4O)CC3
2164 CC(C)Oc1ccccc1N2CcN(CC2)C3CCC(CC3)NC(=O)Nc4ccccc4Cl
2165 CC(C)Oc1ccccc1N2CCN(CC2)C3CCC(CC3)NC(=O)Nc4c(Cl)cc(Cl)cc4C

2166 N#Cc1ccc(cn1)c2n[nH]c3c2cccccc(OCCN5CCCCC5)cc3
2167 CCCCCCNS(=O)(=O)NCCC(OOC)C
2168 CN1CCc2ccc3c2Cccccccc3CC1
2169 CNc1nc(F)nc(Nc2ccc(cc2)S(=O)(=O)N)c1
2170 COc1ccc(Cl)cc1S(=O)(=O)N[C@@H]2CC[C@@](CC2)N3CCC(CC3)c4ccccc4OCC(F)(F)F
2171 CN1C(=O)C=Cc2c(CCN3CCN(CC3)c4ccc5cnc(C)ccc45)c(Cl)ccc12
2172 Cl.NCC1CC1c2cc(C)c(F)cc2OCC=C
2173 COc1cc(ccc1O)c2ccc3c(=O)Nc4cc(ccc4Nc3c2)C(=O)NCCNC(=O)C
2174 CC(N(Cc1ccccc1)S(=O)(=O)c2c(F)c(F)c(F)c(F)c2F)C(=O)NNO
2175 CCCO[C@@H]1CC[C@](C)(CC1)N2CCC(OC2)N3C(=O)Cc4ccc(C)cc34
2176 Cc1nccn2c(c3ccnc(NCC(C)(CCC))n3)c(nc12)c4ccc(F)cc4F
2177 CC(C)Oc1ccccc1C2CCN(CC2)[C@@H]3CC[C@H](CC3)NS(OO)(=O)c4ccccc4O
2178 Cc1cc2CCN(C(=O)Nc3cccnc3)c2cc1B
2179 Cc1cccc(NC(=O)Nc2ccc(cc2)c3csc4c(cnc(N)c34)C#CCC)c1
2180 COc1ccccc1N2CCN(CCNC(=O)c3cnnncccc3)CCC2
2181 CCCCCCNS(=O)(=O)CCCC(=O)CC
2182 Cc1ccc2[nH]cc(OCN(CCCC)CC=C)c2c1
2183 COc1ccc2[nH]cc(CCN(CC=O)CC=C)c2c1
2184 Cc1cc2CCN(C(=O)Nc3cccnc3)c2cc1BF
2185 CCN(CC)C(=O)c1ccccc1NCC2ONCNN2
2186 Cc1c(cc(c2ccccc2)n1C)C(CO)NCCCN3C(N(C)3)c4cccc(Cl)c4
2187 Clc1ccc(Cn2cnnc2)cc1
2188 COc1cc2CCNC[C@@H](C)c2cc1I
2189 CCN(CC)CCOC(=O)C(c1ccccc1)C2(O)CCCCCC2
2190 Cc1ccc2c(cccc2n1)c3nnc(SCCCS4CCc5ccc(cc5CC4)c6cc(C)nn6C)n3C
2191 Fc1c(OCCCN2CCN(CC2)c3cccc4ccccc34)ccc5CCC(OO)c15
2192 CNCCCC1c2ccccc2COCc3ccccc13
2193 NC(=O)c1ccc(cc1)c2c[nH]c3cccc(Cl)c23
2194 FC(F)(F)c1cccc2c(NC(CO)Nc3cnccn3)ccnc12
2195 COc1cc(C=C2SC(=Nc3ccccc3)NC2NO)ccc1O
2196 Clc1cc(NC2CCCCC2)nc(n1)c3n[nH]c4ncccc34
2197 CSC(NS)NS(=O)(=O)c1ccccc1
2198 COc1cccc(c1)C(CC)NC(=O)c2ccc(cc2)c3ccccc3
2199 OP(=O)(O)C(Cc1cccnc1)P(=O)(O)OO
2200 COc1cc2ncnc(Nc3cccc(BO)c3)c2cc1OC

2201 COc1cc2c(Oc3ccc(cc3O)C4=CN=C(Cc5ccccc5)N(C)C4=O)ccnc2cc1OCCCN6CCOCC6
2202 COc1cc(NCc2ccc3nc(N)cc(N)c3c2C)cc(OC)c1OC
2203 CC(C)n1nc(c2ccc3cc(O)ccc3c2)c4c(N)ccnc14
2204 COC(=O)C1C2CCC(CC1c3ccc(I)cc3)2
2205 COc1ccc(cc1OC)c2cc3cccn3c(Nc4ncccc4C(OO)N)n2
2206 NC(=O)N(O)CCC#Cc1ccc(OCCCCNCCCN(CCC)[C@H](c3ccccc3)c4ccc(Cl)cc4)cc1
2207 CC1=CC[C@@H]2[C@@H](C1)c3c(O)cc(cc3OC2(C)C)C(C)(C)c4cccc(CC)c4
2208 Nc1n[nH]c2cccc(c3ccN(NC(=O)Nc4cc(ccc4F)C(=O))ccc3)c12
2209 O=C(NCCCCN1CCN(CC1)c2ccccc2)c3ccccnccccnc3
2210 CCCN(CCc1ccccc1)C2CCc3c(O)cccc32
2211 Nc1n[nH]c2cccc(c3ccc(NC(=O)Oc4cc(ccc4F)C(=O)O)cc3)c12
2212 COc1ccccc1N2CCN(CCCCC(=O)NCc3ccccc4c3ccccc4Cl)cC2
2213 Cc1cccc(c1)S(=O)(=O)NCCNN2CCC(CC2)c3noc4cc(F)ccc34
2214 COc1cc2ncnc(Nc3cccc(Cl)c3)c2cc1OCO
2215 COc1ccccc1N2CCC(CC(O)CCCC(=O)c3oc4ccccc4c3)CC2
2216 CC(C)(C)CCC1(CCCC1NCCNO)c2ccc3[nH]ccc3c2
2217 COc1ccccc1N2CCN(CC(O)CCNC(=O)c3ccc(cc4cccccc43))C2
2218 c1ccc2cc(ccc2c1)c3ccccc3
2219 CN1N=c(S/CC1=CCC(=O)C)S(=O)(=O)N
2220 COc1ccc2[nH]cc(CCN(CCCO)CC=O)c2c1
2221 COc1cc(ccc1Cc2cn(C)c3ccc(NC(=O)OC4CCCC4)cc23)C(=O)NS(=O)(=O)c5ccccc5O
2222 Cc1c(cc(c2ccccc2)n1C)C(=C)NCCCN3CCN(CC3)c4cccc(Cl)c4
2223 CC1=CC[C@@H]2[C@@H](C1)c3c(O)cc(cc3OC2(C)C)C(C4CCCC)CCCCCCcCCC4
2224 ClCCCCCC(C1CCNCC1)c2ccc(Cl)c(Cl)c2
2225 C[C@H](N)Cc1c[nH]c2cc3cOCCCc3c12
2226 OC(Cn1ccnc1)(P(=O)(OOO)P(=O))OOO
2227 Cc1cccc(NC(=O)Nc2ccc(cc2)c3csc4c(cnc(N)c34)C#CCN(C=O)(=O)C)c1
2228 CCn1nnc(n1)[C@H]2O[C@H]([C@H](O)[C@@H]2O)n3cnc4c(NC5CC5)ccCCncnc34
2229 Nc1nc(N)c2nc(CNc3ccc(cc3)C(=O)NC(CCCCCC(=O)c4ccccc4C(=O)O)C(=O)O)cnc2n1
2230 Fc1ccc2c(cccc2c1)N3CCN(CCCNc4ccc5CNC(=O)c5c4F)CC3
2231 CSc1ccc2Sc3ccccc3CC(N4CCC(C)CC4)c2c1
2232 COc1ccccc1N2CNN(CNC(=O)c3ccc(Cl)cc3)CC2
2233 Clc1cc2NC(=O)C3CNCcN3c2cc1N
2234 CCCC(N1CCCC1)C(CO)c2cccc(C)c2
2235 CN1CCC(=C2c3ccccc3COCc4ccccc24)CC1

2236 CC1=CC[C@@H]2[C@@H](C1)c3c(O)cc(cc3OC2(C)C)C(C)(C)c4cccc(BB)c4
2237 COc1ccccc1N2CCN(CCCNNC(=O)c3ccc4ccccc4c3)CCC2
2238 CCCNc1CCc2c(O)cccc2C1
2239 Cl.CC(C)Sc1ccccc1NCC2CNCCN2
2240 Clc1cccc(c1)N2CCN(CC3CCCCC3)CC2O
2241 CNc1cc2c(Nc3cccc(BO)c3)ncnc2cn1
2242 Clc1ccc(cc1CC)C23CNCC2C3
2243 Fc1ccc(cc1)C(=O)CCCN2CCN(CC2)c3ncccc3
2244 COC(=O)C1=C(C)NC(=O)N(C1c2ccc(F)c(F)c2)C(=O)NCCCN3CCC(CC3)c4ccccc4CO
2245 CCN([C@@H]1OC[C@H](CC1)N2CCN(CC2)c3ccccc3OC(C)C)S(=O)(=O)c4ccc(OC)c(OC)c4
2246 COC(=O)c1=C(C)NC(=O)N(C1c2ccc(F)c(F)c2)C(=O)NCCCC3CCN(CC3)c4ccccc4CC
2247 N[C@@H]1CC[C@H](CC1)C(=O)Oc2cc(Oc3ccc(cc3)C(=N)N)cc(Oc4ccc(cc4)C(=N)N)c2
2248 Cc1cc2C(=Nc3ccccc3Nc2s1)c4CCNCC4
2249 N[C@@H]1CC[C@H](CC1)N(OOO)c2cc(Oc3ccc(cc3)C(=N)N)cc(Oc4ccc(cc4)C(=N)N)c2
2250 Cc1cccc(c1)C(=O)CCN2CCN(CC2)c3ccccc3CCN
2251 Nc1ncnc2onc(c3ccc(NC(OO)Nc4cccc(c4)C(F)(F)F)cc3)c12
2252 Clc1cccc(N2CCN(CCCCSS(=O)(=O)c3cnc4ccccc4c3)CC2)c1Cl
2253 CCN(CC)CCOCCOC(CO)C1(CCCC1)c2ccccc2
2254 CN.COc1ccc(F)c(CC)c1C2CC2CN
2255 CN.COc1ccc(F)c(CC)c1C2CC2C
2256 Brc1c(NC2=NCCN2)ccccNCCcc1
2257 COc1cccc(OC)c1OCCNC[C@H]2cOc3ccccc3O2
2258 OC(CNC1CCN(CC1)c2ncnc3scc(c4ccccc4)c23)COc5cccc(c5)CCN
2259 Brc1c(NC2=NCCN2)ccc3ncccc13
2260 CCn1nnc(n1)[C@H]2O[C@H]([C@H](O)[C@@H]2O)n3cnc4c(NC)cccc34
2261 Clc1ccc2CCCCc2c1C
2262 CC(C)n1nc(c2cc(F)c(O)cc2F)c3c(N)nccc13
2263 CCCCC(=O)S(=O)c1ccccc1
2264 COc1ccc2[nH]c(N)c(CCNCCCC)c2c1
2265 Brc1ccc2[nH]cc(OCN(CC=C)CC=C)c2c1
2266 COc1ccc(cc1OC)S(=O)(=O)N[C@@H]2CC[C@H](CC2)N3CCC(CC3)c4ccccc4CC(C)O
2267 COc1ccc(cc1)C2=C(CCC2)c3ccc(cc3)S(=O)(=O)
2268 C(CN1CCN(Cc2ccccc2)CC1)Cc3c[nH]c4ccc(nc34)n5cnnc5
2269 NS(=O)(=O)c1ccc(nc1OC)CC
2270 Cl.CC(C)Sc1ccccc1NCC2NCCCN2

2271 Cc1ccc2c(cccc2n1)N3CCN(CCc4cccc5c4OCc6c(ncn56)C(=O)N7CCC7)C3

2272 COc1ccccc1N2CCN(CCN3C(=O)Nc4ccccc4C3CO)CC2

2273 C(Oc1ccccc1c2cccs2)C3=NNCN3

2274 CC(C)Oc1ccccc1C2CCN(CC2)C3CCC(CC3)NC(=O)NCc4cccc(F)c4

2275 Cc1[nH]c2nc(nc(N[C@@H]3CC[C@@H](C)CC3)c2c1C)c4ccccc4

2276 OP(=O)(C)C(Cc1cccnc1)P(=O)(O)O

2277 Nc1ccc(cc1BSSS(=O)S=O)N

2278 COC(=O)C1C2CC[C@H](C[C@@H]1c3ccc(I)cc3)C2C

2279 C=CCc1ccccc1OCCC=C2CN2

2280 COc1ccccc1N2CCN(CCCNNC(=O)c3ccc4ccccc4c3)CC2

2281 CN[C@H](Cc1ccccc1)C(=O)N2CCC[C@H]2C(CO)N[C@@H](CCCN=C(N)N)C=O

2282 COC(=O)c1ccccc1NCC2=CCCN2

2283 Clc1ccc(COc2ccc3Sc(CO)Oc3c2)cc1

2284 Cc1cccc(c1)C(=O)NCN2CCN(CC2)c3ccccc3CC

2285 CN1CCc2c(C1)c3cccc4cCc5ccccc5c2c34

2286 COc1ccccc1N2CCN(CCCCNC(=O)c3ccn4nncc4c3)C2

2287 Fc1ccc2c(noc2c1)C3CCN(CCCCNS(=O)(=O)c4ccc(CC(F)(F)F)cc4)CC3

2288 CCCCCCC(C)(C)c1cc(O)cc(OOCCCCCCC(=O)NC2CC2)c1

2289 Cl.NCC1CC1c2cc(F)c(F)cc2CCC=C

2290 C[C@H](N)Cn1ccc2cccBccccc2c1

2291 Fc1ccc2c(noc2c1)C3CCN(CCCCNS(=O)(=O)c4ccc5cs(Cl)ccc5c4)CC3

2292 Cc1ccc2c(cccc2n1)N3CCN(CCc4cccc5N(CC(O)O)C(OO)COc45)CC3

2293 O=C1CC2(CCCC2)CC(=O)N1CCCCNCCCC(CCC)c4ncccn4

2294 CC(=O)N1CCC(CC1)NC(=C)NC2CCCCCC2

2295 OC[C@H](Nc1CCN(CCCc2c[nH]c3ccc(cc23)n4cnnc4)CC1)c5ccccc5

2296 COc1cc2nc(nc(N)c2cc1OC)N3CCN(CC3)C(OC)C4CCSS4C

2297 C\\NCC\\1/Oc2ccccc2C=C1

2298 CCCCC1CCN(CCCC(=O)c2ccccc2CC)C1

2299 CSc1ccccc1NCC2NNCCN2

2300 COC(=O)C1C2CCC(CC1c3ccc3Cl)c(Cl)c(N)2

2301 OP(=O)(O)C(NC1CCCCc1)P(=O)(O)

2302 Clc1cccc(Cl)c1NC2=NCNC2

2303 COC(=O)C1=C(C)NC(=O)N(C1c2ccc(F)c(F)c2)C(=O)NCCCN3CCN(CC3)c4ccccc4CO

2304 COc1ccc2[nH]cc(CCC(CCCC)CCCO)c2c1

2305 CN1N=C(Sc1=N)S(=O)(=O)N

2306 CC1(N(CCc2cc(O)ccc12)c3cccc(O)c3)c4ccc(OCCN5CCCC5)cccc4
2307 COc1ccc(cc1OC)c2cc3nccn3c(Nc4ncccc4CC(O)N)n2
2308 COc1cc(ccc1Nc2ncc3CN(C)C(=O)N(c4cccc(NC(=O)C=C)c4)c3n2)NCCCCCCCCC
2309 C1CCC(CC1)Nc2nc(Nc3ccc(cc3)n4CCOCC4)nc5[nH]cnc25
2310 CCCc1nc(C)c2C(=O)NC(=Nn12)c3ccccc3OC
2311 CC(C)Cc1cc(on1)c2ncc3[nH]nc(O)c3c2
2312 CC(C)Cc1ccccc1Sc2ccc(I)cc2N
2313 Clc1ccc(cc1CC)C(=O)C(CCCC)N
2314 CCN1CCCC1CNC(=O)c2cccccc2(C)S(=O)(=O)N
2315 C[C@H](c1c[nH]cn1)c2cccccCCC2CC
2316 O=C1NCc2ccc(OCCCN3CCC(CC3)c4cccc5CCCCc45)cc12
2317 COc1ccc(cc1)C(C(C)C)c2nc(C)nc3oc(C)cc23
2318 Cc1cccc2C3C2NCC3NC(OO)c1
2319 COC1(CCOCC1)c2cc(F)cc(O)c3ccc4c(C)C(=O)C=Cc4c3cc2
2320 FC(F)(F)c1ccc2c(NC(=O)N3ccccc(n3)C#N)ccnc2c1
2321 Cc1cc2CNC(=O)c2cc1OCCCCN3CCC(CC3)c4cccc5ccccc45
2322 COc1ccccc1N2CCN(CCCCCCNC(=O)OC(C)(CCC)C)2
2323 CNC(=O)[C@]12C[C@@H]1[C@H]([C@H](O)[C@@H]2O)n3cnc4c(NC5CCCC5)nc(CC)nc34
2324 COc1ccc2c(NC(=O)Nc3cccc(c2)n3)cccccc1
2325 COC(=O)CCNS(=O)(=O)NCCCCc1ccccc1
2326 CC(=O)N1CCC(C11)(C(=O)NCCCCCCO)C1
2327 Cc1cc2CNC(=O)c2cc1OCCCN3CCC(CC3)c4cccc5cccc(F)c45
2328 C#CCN[C@@H]1CCc2cccnc12
2329 COc1cc(Cl)ccc1N2CCC(CCN3C=Nc4sc5CN(C)CCc5c4C3=O)CC2
2330 COc1cc(ccc1Nc2ncc(Cl)c(Nc3ccccc3P(=O)(C)C)n2)N4CCC(CC4)C5CCCC5
2331 CNCCc1c[nH]c2ccc(cc12)C(CCOC)C
2332 Nc1ncnc2c1c(nn2C3CCCC3)c(ccc5ccc5cccc5)c5
2333 Cc1ccc2c(cccc2n1)N3CCN(CCCCc4ccc5CCC(OO)Nc5c4)CC3
2334 CCCOc1cc2c(cc1\\C(=C/C=C/C(=C(C(=O)O)/C)\\C)C(C)CC)CCC2(C)C
2335 Cc1cccc(CC(=O)CN2CCN(CC2)c3ccccc3C)NCc1
2336 NS(=O)(=O)c1ccc(cc1)C2CCCCc2CCCCCCC
2337 CCCN1C(=O)N(CCC)c2cc([nH]c2C1=O)c3ccc(OOC(=O)Nc4ccccc4)cc3
2338 Clc1ccc(CNC2=C(Nc3ccncc3)C(=O)C2=O)cc1CC
2339 Nc1oc(nn1)c2cc3c(Cc4ccc(Cl)cc4)cccc3s2
2340 CCCCCCC(C)(C)c1cc(O)c2C3CC(=O)CCC3C(CCCCC)C2cc1

2341 C[C@H](Oc1ccccc1CC=C)C2=CCCN2

2342 COc1cccc(OC)c1OCCNCCOc2ccccc2OCc3ccccc3CC

2343 Clc1ccc(CNC2=C(Nc3ccncc3)C(CO)C2=O)cc1Cl

2344 CN(C)CC1(CCCCC1)c2cccc3ccccc32

2345 CN(C)C1(CNCCC2CCC2C)COcccccccc1C

2346 N[C@@H]1CC[C@H](CC1)OC(=O)c2cc(Oc3ccc(cc3)C(=O)N)cc(Oc4ccc(cc4)C(=N)N)c2

2347 O=C(Nc1ccc(c2ccccc2)c(c1)c3occc3)C4CC4

2348 COc1ccc(cc1OC2CCCC2)C(CO)Nc3c(F)cccc3F

2349 Cc1ccc(cc1)N2N=C3N(C2=O)c4ccccc4NNC3NC

2350 CC(=O)OC1CcCCC(c1N)

2351 CC1(N(CCc2cc(O)ccc12)c3cccc(CC)c3)c4ccc(OCCN5CCCC5)cc4

2352 COc1ccc2CC(CCc2c1)NC(=O)C(C)C

2353 CC(C)NCC(O)c1ccc(O)c(O)c1O

2354 Fc1ccc2c(cccc2c1)N3CCN(CCCCc4ccc5CNC(=O)c5c4)CC3

2355 Cc1cccc(NC(=O)Nc2ccc(cc2)c3csc4c(cnc(N)c34)C#CC5CCCCC5)c1

2356 Fc1ccc(cn1)C(=O)NCCCCN2CCN(CC2)c3cccc(Cl)c3C

2357 Clc1cccc(N2CCN(CC\C=C\\CNC(=O)c3occcccccc3)CC2)c1Cl

2358 COc1ccc(cc1)S(=O)(=O)NC(C)C(=O)NO O

2359 OC(=O)c1ccccc1Nc2ccnc(Nc3ccc4nn[nH]c4c3)n2

2360 COCC(N1CCCc1)C(=O)c2C(CCCC)Cc2

2361 COc1ccc(cc1)S(=O)(=O)c2ccc(cc2)C(=C)C3CCC(CC3)C4CCCCC4

2362 COc1cc(NC(=O)c2oc(cc2)c3ccc(Cl)cc3)cc(CO)c1

2363 CCCCCCC(C)(C)c1cc(O)c2C3CC(=O)CCC3C(CC(C))c2c1

2364 CC1=CN=C(NCC(F)(F)c2ccccc2)C(=O)N1CC(=O)CCc3ccc(N)nc3C

2365 COc1cc(ccc1Nc2ncc(Cl)c(Nc3ccccc3P(=O)(C)C)n2)C4CCNCCOCC4

2366 CCN(CC)C(=O)c1ccccc1NCC2=CCCN2

2367 CCCCN(CCCCN)C(=O)c1ccc2nc([nH]c2c1)c3n[nH]c4ccccc34

2368 CC(Oc1ccccc1c2cccs2)C3=CCCN3

2369 COc1ccc(cc1)c2nc3c(N)ncc4cccc4n3n2

2370 O=c1CCCN1CC#CCN2CCCC2

2371 Nc1ccc2c(CCC3CCN(Cc4ccccc4)CC3C)oo2c1

2372 COc1ccccc1N2CCN(CCCNC(=O)c3ccn4ccccc34)CCC2

2373 COc1cccc(c1)C(C)NC(=O)c2ccc(cc2NCCNN)c3ccncc3

2374 CC1NC(=Nc2cccc(CN)c12)N

2375 FC(F)Oc1ccccc1CC(N2CCCCC2)c3ccccc3

2376 Cc1c(cc(c2ccccc2)n1C)C(=O)NCCCN3CCN(CC3)c4cccc(Cl)4CCC

2377 CC(Oc1ccccc1C2CcCC2)C3CNCCN3

2378 CN(C)CCCOC(=O)C(CC1CCCCCCC1)c2ccccc2

2379 COc1ccccc1N2CCN(CCCNC3=C(C)CC(O)N(C)C(OO)N3C)CC2

2380 NS(=O)(=O)c1nc2cc1(OOS)c1c2s1

2381 Cc1ccc(cc1)C(=O)OCCCOCCC2CCCC2

2382 COC(=O)c1cc(CCc2cc(OC)ccc2OO)ccc1O

2383 Clc1ccc(cc1)N2CCN(Cc3ccn4ccccc34)CC2

2384 Clc1ccc2cc(ccc2c1)S(=O)(=O)NCCCN3CCN(CC3)c4csc5ccccc45

2385 Cc1ccc(cc1)C(=O)C(CC=C)N2CCCC2C

2386 CCCN(CCC)C1CCC(=CC1)CC

2387 Cc1cccc(NC(CO)Cc2ccc(cc2)c3cccc4[nH]nc(N)c34)c1

2388 C(Nc1ccccc1c2ncccc2)C3=NCCN3C

2389 COc1ccc(cc1OC)S(=O)(=O)N[C@@H]2CC[C@@H](CC2)N3CCC(CC3)c4ccccc4OCC(F)(F)

2390 COC(=O)C1C2CCC(CC1c3ccc(Cl)cc3)c2C

2391 COc1cc2ncnc(Nc3ccc(F)c(Cl)c3)c2cc1OCCCCCCC(OO)NO

2392 O=C1NC(=NC(=C1)c2ccncc2)NCCCCCCC

2393 NNc1ccccc1S(=O)(OO)

2394 Fc1ccc(cc1)C(=O)CCCN2CCC(=CC2)N3C(=O)Cc4ccccc34

2395 Clc1cccc(N2CCN(CCCCOc3ccc4CCC(=C)Nc4c3)CC2)c1Cl

2396 OC1(CC2CCc(C1)N2CCCC(=O)c3ccc(F)cc3)c4ccccc4

2397 COc1cc2ncnc(Nc3ccc(F)c(c3)C#C)c2cc1OCCCCCCC(=O)C

2398 Cc1oc2nc(C)nc(Nc3cccc(c3)C#C)cc21

2399 Cc1ccccc1Oc2ccc(cc2CN)C#N

2400 CN(C)C(=O)N[C@@H]1CC[C@@H](CCN2CCC(CC2)3ccccc4CCOc34)CC1

2401 Cc1ccc2c(cccc2n1)N3CCN(CCc4cccc5c4OCc6c(ncn56)C(=O)NCCCCCC)CC3

2402 COc1cc2nc(nc(N)c2cc1OC)N3CCN(CC3)C(OO)C4CCSS4C

2403 COC(=O)C1C2CCC(CC1c3ccc4ccccc4c3)N2

2404 CN(Cc1ccc2Nc(=NC(=O)c2c1)C)c3ccc(s3)C(=O)N[C@@H](CCC(=O)O)C(OO)O

2405 CN(C)CC1(CCCCN1)c2ccc3ccccc3c2

2406 Clc1c(BN)ccc2CCNCCc12

2407 NS(=O)(=O)c1ccc(NC(=O)c2ccccc2)cccc1

2408 CN[C@H](Cc1ccccc1)C(=O)N2CCC[C@H]2C(NO)NC(CCCN=C(N)N)C(=O)c3nc4CCCCc4s3

2409 COc1cc2c(Oc3ccc(NC(=O)N\\N=C\\c4cccc(CC(C)c4F)cc3F)ccnc2cc1CC)C5NCCCCC5

2410 CN(C)CC1=C(O)C2=CC(=CC(=O)N2CCC1)C

2411 N#Cc1ccc(cn1)c2n[nH]c3c2cccccc(OCCN5CCOCC5)cc3
2412 COc1cc(ccc1Nc2ncc(Cl)c(Nc3ccccc3S(=O)(=O)CCC)ncn2)P(=O)(C)C
2413 CCOc1nc(cc(N)c1C#N)C(NO)OCc2ccccc2S(=O)(=O)N
2414 CC1=CC[C@@H]2[C@@H](C1)c3c(O)cc(cc3OC2(C)C)C(C)(C)c4ccc(BF)cc4
2415 NS(=O)(=O)c1ccc(O)cc1 1
2416 CCCC(N1CCCC1)C(=O)c2ccc(BC)cc2
2417 CC(C)(F)C(CO)N[C@@H]1CC[C@@H](CCN2CCC(CC2)c3coc4ccccc34)CC1
2418 NS(=O)(=O)c1cc(ccc1)n2CCCCCCCCCCC2
2419 CC(N)Cc1c[nH]c2cccBOOcc12
2420 NS(=O)(=O)c1ccc(O)cc1  1
2421 Clc1ccc(cc1Cl)S(=O)(=O)NCCCN2CCN(CC2)c3csc4ccccc34
2422 NS(=O)(=O)ONC(=O)NNc1ccccc1
2423 Brc1c(NC2CNCCN2)ccc3nncnc13
2424 CCCCCOc1ccccc1c2cnccc2CC(=O)NC3CCCCC3
2425 Nc1nc(N)c2nc(CNc3ccc(cc3)C(=O)NC(CCCC(=O)c4ccccc4C(=O)O)C(=O)O)cnc2n1
2426 CCCCC(CC(=O)NO)S(=O)(=O)C1CCCCC1C
2427 CCCn1cc(C(=O)C2C(C)(C)c2(C)C)c3ccccc13
2428 CCN(CC)C(=O)c1ccccc1CCC2=CCCN2
2429 CC(C)CCC(O)c1ccc(O)c(O)c1
2430 O=C(CC1N)N[C@@H]1CC[C@@H](CCN2CCN(CC2)c3cccc4OCOc34)CCC
2431 NC(=O)C1=C(NCC(=O)C=C)Nc2ccccc2OOC1OO
2432 CCCCCCCCCCSS(=O)(=O)N
2433 C[C@H]1[C@H]2Cc3ccc(O)cc3[C@]1(C)CCC2CC4CC4
2434 COc1cc2nc(nc(N)c2cc1OC)N3CCN(CC3)C(CO)C4CCCO4
2435 COc1cc(ccc1Nc2ncc3CN(C(C(=O)N(c4cccc(NC(=O)C=C)c4)c3n2)N)CCC)
2436 COc1cc2c(Oc3ccc(NC(=O)C4=C(C)N(C)N(C4=O)c5ccccc5)cc3F)ccnc2cc1OCCCN6CCC(C)CC6
2437 COC(=O)CCNS(=O)(=O)NCC1cccccccc1
2438 CC(=O)O.OSc1cc(ccc1O)[C@@H](O)CNCCCCCCCCCCCc2cccc(CCCCCOOOCNCCOO)c2
2439 COc1ccc2[nH]cc(CCN(CC=C)CC=N)c2c1
2440 CC1=CC[C@@H]2[C@@H](C1)c3c(O)cc(cc3OC2(C)C)C4(CCCC4)CNN
2441 CCCCCCCNc1c2CCCn2ncccc(F)ccc1
2442 CC(C(=O)N)c1cccc(c1)C(=O)c2ccccc2
2443 Ic1c2N(N=NCCN2)ccc3nncnc13
2444 CN1CCc2c(C1)c3cc(C)cccnn4ccc4ccc(C)c23
2445 FC(F)(F)c1cccc(c1)N2CCNCCCCN3Cc4ccccc4CC3OCCC2

2446 O\\N=C/1\\C(=C/2\\C(=O)Nc3ccc(BO)cc23)\\Nc4ccccc14
2447 COC(=O)C(C1CCCCC1)c2ccc(Cl)cc2
2448 CC(C)NC(C)C(O)COc1c2c(C)ccCCCc12
2449 O=C1Nc2ccccc2N1C3CCC(CC3)C4CCCCC4
2450 CCOc1nc(cc(N)c1C#N)C(=O)NCc2ccccc2S(=O)(=O)NN
2451 N\1N=C\\1/Oc2ccccc2C=C
2452 NS(=O)(=O)c1cc2ccc(=CS)ccc2s1
2453 CN1C(=O)C(=Cc2cnc(NC3CCOCC3)cc12)Oc4ccc(F)cc4F
2454 NS(=C)(=O)N
2455 Clc1ccc2cc(sc2c1)S(=O)(=O)NNCCCN3CCN(CC3)c4nsc5ccccc45
2456 Oc1ccc2C(N(CCc2c1)c3ccccc3)c4ccc(OCCC5CCCCC5)cc4
2457 CCCN(CCC)C1CCC(=CC1)C
2458 Cn1c(SCCCN2CC3CCC(C3C2)c4ccc(C)cc4)nnc1C5CCCCC5
2459 Nc1ncnc2c1c(cn2n3CCCC3)C#C
2460 Cc1ccc(cc1)N2CCc3cc(O)ccc3c2(C)c4ccc(OCCN5CCCC5)cc4
2461 Cc1ccc2c(ccc(F)c2n1)N3cCN(CCc4ccc5OCC(=O)Nc5c4)CC3
2462 CCCc1nn(C)c2C(=O)NC(=Nc12)c3ccccc3OCCC
2463 CN.COc1ccc(F)c(Cl)c1C2CC2
2464 Clc1ccc2C3CCC(=O)CCC3CCc2c1
2465 Nc1ncnc2c1c(cn2C3CCcC3)C C
2466 C1CCC(CC1)Nc2c(nc3ncccn23)c4ccc5[nH]ccc5c4
2467 O=C1CCCN1CC#CCN2CNCC2
2468 CC1=CC[C@@H]2[C@@H](C1)c3c(O)cc(cc3OC2(C)C)C(C)(C)CCCCCCN
2469 Cc1ccc(F)c(C(OO)n2nc(Nc3ccc(cc3)S(=O)(=O)N)nc2C)c1F
2470 COc1cc(ccc1O)c2ccc3c(=O)Nc4cc(ccc4Nc3c2)C(=O)CCCNC(=O)C
2471 COc1ccc2[nH]c(C)c(NCC(C)C)c2c1
2472 CC1=CC[C@@H]2[C@@H](C1)c3c(O)cc(cc3OC2(C)C)C(C)(C)c4ccc(B)Bcc4
2473 Cc1c(NC2=CCCN2)ccc3NCCCc13
2474 Brc1ccc2[nH]cc(CCC(CC=O)CC=C)c2c1
2475 CC1NC(=Nc2cccc(Cl)c12)
2476 CC1=CC(=O)Oc2cc(OCCNN3CCC(CC3)c4ccccc4)ccc12
2477 CCCC(N1CCCC1)C(=O)c2ccc(CC)cc2
2478 COc1cc(ccc1O)C2=C(O)C(=O)c3c(ccc3OC)cccO2
2479 CCN(C=O)CCCOc1ccc2cccc3cc(CC(=O)Nc4cccc(F)c4)[nH]n3ncccc2c1
2480 Cl.CCC(=O)C(C1CCCCC1)c2ccc(Cl)c(Cl)c2

2481 COc1ccc(cc1OC)S(=O)(=O)N[C@@H]2CC[C@H](CC2)N3CCC(CC3)c4ccccc4OC(C)O

2482 CN1CCCc2ccccc2Cc3ccc(O)ccccC13

2483 COc1cc2nccc(Oc3ccc(NC(=O)C4=C(C)N(C)N(C4=O)c5ccccc5)cc3F)c2cc1OO

2484 Oc1ccc2C(N(CCc2c1)c3ccccc3)c4ccc(OCCC5CCCC5)cc4

2485 COc1cc(ccc1Nc2ncc3CN(C)C(=O)N(c4cccc(NC(=O)C=C)c4)c3n2)N5CCC(C5CCNCC)C

2486 COc1ccc(cc1)C2=S(CCC2)c3ccc(cc3)S(=O)(=O)

2487 CC(=O)OC1CC2CCCC1O2

2488 Nc1ccccc1C(=O)CCCC2CC[C@H]3[C@@H](C2)c4cccc5CCCCN3c45

2489 O=C1CCc2cc(ccc2N1)c3nccnc3

2490 COc1ccc(NC(=O)Nc2cccc(F)c2)cc1c3cBBnccnn3C

2491 CC(CCc1ccccc1)NCC(O)c2ccc(O)c(c2)C(=O)NC

2492 O=c1CCCN1CC#CCN2CNCC2

2493 COc1ccc(cc1OC)S(=O)N(O)N[C@@H]2CC[C@H](CC2)C3CCC(CC3)c4ccccc4OC5CC5

2494 FC(F)(F)c1cccc2C[C@@H]3CNCCC3C(=O)c12

2495 CCNC(=O)[C@H]1O[C@H]([C@H](O)[C@@H]1O)n2cnc3c(NCC(N)N)ncnc23

2496 CC(C)c1cc(Oc2c(Cl)cc(CCC(=O)O)cc2Cl)ccc1OO

2497 Cc1nc[nH]c1CCC

2498 CN1CCN(CC1)c2cc(Nc3cc(C)[nH]n3)nc(Oc4cccc(NC(=O)C=C)c4)c2

2499 CN1N=C(CC1=N)S(=O)(=O)N

2500 CSc1cc(F)ccc1NCC2=CCCN2

2501 CCCN(CCC)C1CCC(OCC1)C#C

2502 Oc1ccc(cc1)c2cc3ccc(O)cccc23

2503 Brc1ccc(Cn2nccc2c3ccccc3)cc1

2504 Cc1cc2CCN(C(=O)Nc3cccnc3)c2cc1

2505 COc1ccccc1COc2ccccc2OCCCCOc3OcOOccccc3OC

2506 O=C1C=C(Nc2c1ccc3ccccc23)c4ccccc4

2507 Cl.CC(C)Sc1ccccc1N=C2CCNNN2

2508 Oc1cccc2CC[C@@H]3[C@H](CCN3OC=C)c12

2509 Cl.N#Cc1ccc(cc1)C2CCCc3cccn23

2510 CCCCCCC(C)(C)c1cc(O)c2C3CC(CO)CCC3C(CCCC)cc2cc1

2511 CC(=O)N1CCC(CC1)NC(CO)NC2CCCCCC2

2512 Clc1cccc(N2CCN(CCCCCS(=O)(=O)c3c4cccccnc4c3)CC2)c1Cl

2513 CC1=CC[C@@H]2[C@@H](C1)c3c(O)cc(cc3OC2(C)C)C(C)(C)CCCCCNN

2514 ClNc1ccccc1c2cccNcCC2CNNNCNNC

2515 Oc1ccc2C3=C(CCOc2c1)c4ccc(O)cc4O[C@@H]3c5ccc(CCCN6CCOCC6)cc5

2516 COc1ccc(cc1)C2=C(CCC2)c3ccc(cc3)S(=O)(CO)

2517 O=C1CCc2ccc(OCCCCN3CCN(NC3)c4cccc5CcCCc45)cc12

2518 OC[C@H]1O[C@H]([C@H](O)[C@@H]1O)n2cnc3c(NC4CCCC4)nc(Cl)cc23

2519 COc1ccc2NC(=O)C(=Cc3c[nH]cc3)c2c1

2520 CN1NC(=Nc2cccc(Cl)c12)N

2521 CN1CCc2ccccc2Cc3c[nH]cccc4c(CC1)c34

2522 COc1cccc(OC)c1CCCNCCOc2ccccc2c3ccccccc3

2523 Brc1c(NC2=NCCC2)ccc3nncnc13

2524 COc1cccc2c(CCCNCCOc3ccccc3)cccc21

2525 CCn1nnc(n1)[C@H]2O[C@H]([C@H](O)[C@@H]2O)n3cnc4c(NC5CCC)cCCCn5nnc34

2526 CC1=CN=C(NCCc2ccccc2)c(=O)N1CC(=O)NCc3ccc(N)cc3C

2527 Cc1cccc(c1)c2cncn2Cc3ccc(cc3)NN

2528 CCCCCCC(C)(C)c1cc(O)c2C3CC(=O)CCC3C(CCCC)cc2cc1

2529 Fc1ccc2c(noc2c1)C3CCN(CCCCNS(CO)(=O)c4cc5ccccc5s4)CC3

2530 COc1cc2c(Oc3ccc(cc3F)C4=CN=C(Cc5ccccc5)N(C)C4=O)ccnc2cc1OCOC6CCCOCC6

2531 C1Cc2ccc3occc3c2CCC1

2532 NCc1ccc(cc1)S(=O)(=O)NN

2533 COc1cc2ncnc(Nc3cccc(c3)C#C)c2cc1OCCCCCCC(OO)N

2534 Fc1ccc2c(cccc2c1)N3CCN(CCCOc4ccc5CNC(OO)c5c4F)CC3

2535 Cc1cc2CNC(=O)c2cc1CCNCCN3CCN(CC3)c4cccc5ccccc45

2536 Cl.NC[C@H]1C[C@@H]1c2ccc(F)cc2CC

2537 NS(=O)(=O)c1ccc2CCCc2c1 1

2538 C\\NCC\\1OBc2ccccc2C=C1

2539 COC(=O)c1cncn1Cc2ccc(cc2)CCN

2540 CN1CCc2c(C1)sc3N=CN(CCN4CCN(CC4)c5cccc6ncnnc56)C(=O)c23

2541 CCCCCN(C)CCC(O)(P(=O)(OPO)P(OO))OO

2542 Nc1c2CCCCc2cc3cccc(CC)c13

2543 Cl.CCNC1CCS(=O)(=O)c2sc(cc12)S(=O)(=O)

2544 COc1ccccc1N2CCC(CNCCc3Oc4ccc(Cl)cc4O3)CC2

2545 CC(Oc1ccccc1c2cccs2)C3CCCCN3

2546 Cc1ccc2c(cccc2n1)N3CCN(CCCc4ccc5OCC(=O)Nc5c4)CCc3

2547 Cc1cc(cc(C)c1CC2CNCC=2)C(C=NC)C

2548 CN(C)CCC(c1c2c(Cl)c21)c2ccccn2

2549 CC(=O)N[C@H](Cc1ccccc1)C(=O)N2CCC[C@H]2C(=O)N[C@@H](CCCN=C(N)N)B(O)

2550 COC(=O)C1C2CCC(CC1c3ccc4ccccc4c3)Nc2

2551 CN1CCN(Cc2ccc(NC(=O)c3ccc(C)c(c3)C#Cc4cnc5[nH]nnc5c4)cc2C(F)(F)F)CC1
2552 Ic1c(NC2=NCCN2)ccc3ncccc13
2553 Clc1cccc(N2CCN(CCCCOc3ccc4C=CC(=O)N4cn3)CC2)c1Cl
2554 Fc1ccc(cc1)C(=O)CCCN2CCC(=CC2)N3C(=O)cccccccc3
2555 CC(C)(C(=O)c1ccccc1)c2ccccc2
2556 CCCCCCCC(=O)C(=O)CCCC=CC
2557 CN1CCC(CCCN2c3ccccc3Sc4ccc(Cl)cc24)CC1
2558 CC(C)Oc1ccccc1C2CCN(CC2C[C@@H]3CC[C@H](CC3))S(=O)(=O)c4ccc(F)c(F)c4
2559 COc1cc2c(Oc3ccc(Nc4ccc(cc4)C(C)(C)N)cc3)ccnc2cc1OCCNCC
2560 C\\N=C\\1/Oc2ccccc2C=C1
2561 CCOc1nc(NC(=O)Cc2cc(OC)c(cc2NC)S(=O)(=O)C)cc(N)c1C#N
2562 O=C1CCCC1CC#CCN2CCCC2
2563 CCSc1ccc(CCN(CNCN)CCC)Ncccc1
2564 COc1c(Cl)cccc1N2CCN(CCCCNC(=O)c3ccc4ccccc4c3)CC2
2565 Nc1nc(N)c2c(CN)c(CNc3ccc(cc3)C(=O)CC(CCC(=O)O)C(=O)O)ccc2n1
2566 C(CN1CCN(CCCC(c2ccccc2)c3ccccc3)CC1)Cc4ccccc4
2567 Clc1ccc(CNC2=C(Nc3ccncc3)C(=O)c2CO)cc1C
2568 Nc1ccc2nc(sc2c1)S(=O)(=O)
2569 COc1cccc(OC)c1OCCNCCCOc3ccccc3O
2570 Cn1c(SCCCN2CC3CCC(C3C2)c4ccc(F)cc4)nnc1C5CCCCC5
2571 Cc1c(cc(c2ccccc2)n1C)C(=O)NCCCC3CCN(CO3)c4cccc(Cl)c4
2572 COc1c(C)c2COC(=O)c2c(O)c1C\\C\C(C\C\\CCCC=O)O
2573 NS(=O)(=O)c1ccc(cc1)C2=C(C(=O)NC2)c3ccccc3
2574 Fc1ccc2c(noc2c1)C3CCN(CCC(CNCO)c54ccc5c4)cC3
2575 Cc1cc(Nc2ncc(s2)C(=O)Nc3c(O)cccc3CC)nc(C)n1
2576 CNC(=O)c1cc(Oc2ccc(NC(=O)Nc3ccc(Cl)c(c3)C(F)(F)C)cc2)ccn1
2577 O=C1CCc2ccc(CCCCCC3CCN(CC3)c4ccccc4)cc21
2578 CC(C)Nc1c(nc2ccccn12)c3ccc4[nH]ncc4c3
2579 COC(=O)c1cccc21CCCNNNCCC2
2580 COc1ccccc1CNCCCCCCNNNCCCCCCNCCCCCCNCc2ccccc2C
2581 NS(=O)(=O)c1cccc(c1)S(=O)(=O)
2582 COc1ccccc1N2CCN(CCCCNC(=O)c3oncccccccc3)CC2
2583 CC(C)Oc1ccccc1N2CCN(CC2)C3CCC(CC3)NC(=O)Nc4ccccc4C
2584 COc1ccc2cc(ccc2c1)c3ccccc3
2585 Cc1c(OCCCN2CCN(CC2)c3cccc4ccccc34)ccc5CCC(=O)c15

2586 CC(C)Oc1ccccc1N2CCNNCCCNC(=O)CN3CCCCC3=CCCC2

2587 CCOc1cc(CN2CCC(CC2)Nc3oc4ccccc4n3)ccc1CC

2588 Nc1ccc(cc1I)S(=O)(=O)

2589 C(c1ccccc1)n2cc(nn2)c3ccc4[nH]ncc4c3 4

2590 NS(=O)(CO)C

2591 Fc1ccc2c(noc2c1)C3CCN(CC4CC(=O)c5ccoc5C4)cC3

2592 COC(=O)[C@@H]1C2CCC(C[C@@H]1c3cccC\\C(C/I)cc3)N2

2593 Cc1ccc(Cl)c(c1NN2CCN(CC(CC(=O)C4)CCCCC4)CCCCC)CC2

2594 COc1cc(C=C2SC(=Nc3ccccc3)cC2NO)ccc1O

2595 Clc1ccc2c(NCCCNC(CO)CCCCCCCCSSS)cnCCCCcnnc2c1

2596 CC[C@@]1(O)C(=O)OCC2=C1C=C3N(Cc4cc5c(N)cccc5nc34)n2=O

2597 COc1ccccc1CNNCc2cccc(CCNC[C@H](O)c3ccc(O)c4NC(=O)Sc34)c2

2598 Cc1c(Nc2ccc[nH]2)ccc3CCCCc13

2599 NNc1ccccc1S(=O)

2600 Nc1nc(N)c2c(Cl)c(C(c3ccc(cc3)C(=O)CC(CCC(=O)))C(=O)O)ccc2c1

2601 CN(Cc1cnc2nc(N)nc(N)c2n1)c3ccc(cc3)C(=O)NC(CC(=C)CC(O)O)C(=O)O

2602 CCCN(CCC)C1CCc2cc(N)ccc2C1

2603 CCCCCCC(C)(C)c1cc(O)cc(OCC(CCCCC)=O)CC(=C)c1

2604 NCCn1ncc2ccc(O)c1c2

2605 COc1cc2ncnc(Nc3cccc(c3)C#C)c2cc1OF

2606 N[C@@H]1CC[C@H](CC1)N(OO)Oc2cc(Oc3ccc(cc3)C(=N)N)cc(Oc4ccc(cc4)C(=N)N)c2

2607 NS(=O)(=O)OSC(=O)NCc1ccccc1

2608 Nc1oc(nn1)c2cc3c(Cc4ccc(Cl)cc4)cncc3s2

2609 COc1ccc(NC(CS)NC(=O)c2cn(cc2c3ccc(Cl)cc3)c4ccccc4)cc1

2610 CCC1(Cc2ccccc2C1)c3ccc[nH]3

2611 COc1cc2OC(=O)C=Cc2cc1[C@@H](OOCCCC)C

2612 C=CNCC(O)COc1ccccc1C(=O)CCc2ccccc2

2613 Fc1ccc(cc1)C(=O)C2CCC(CCC3C(=O)Nc4ccccc4C3=O)CC2

2614 CCOc1nc(NC(=O)Cc2cc(OC)ccc2)cccc(C)c1CCN

2615 CCCCCCCCC(CO)C(=O)CCC=CCCC

2616 COc1cc(NC(=O)c2oc(cc2)c3ccc(Cl)cc3)cc(OO)c1

2617 Cc1ccc2c(cccc2n1)N3CCN(CCc4ccc5CCCC=O)Nc5c4CCC3

2618 NCCo1ncc2ccc(O)c1c2

2619 CC1=CC[C@@H]2[C@@H](C1)c3c(O)cc(cc3OC2(C)C)C4CCCCC44CCCCCC4

2620 COc1cccc(c1)C(C)NC(=O)c2ccc(c(C)c2)c3ccccc3

2621 CCCN1C(=O)N(CCC)c2cc([nH]c2C1=O)c3ccc(OCC(=O)Nc4ccc(BO)cc4)cc3

2622 OCCNCCNc1ccc(NCCNCCO)c2C(=O)c3c(O)ccc(O)c3C(CO)c12

2623 CC(C)(C)C(=O)CCc1ccc(cc1)S(=O)(=O)N

2624 Cc1ccc2c(cccc2n1)N3CCN(CCc4cccc(c4)N5CCNC5=O)Cc3

2625 Cc1nc(N)sc1c2ccnc(Nc3ccc(cc3)C4CCOCC4)n2

2626 COc1ccccc1N2CCN(CNC(=O)c3cccc(C)Cc3)CC2

2627 CC(C)(C)Sc1c(CC(C)(C)C(=O)C)n(Cc2ccc(Cl)cc2)c3ccc(OCc4ccc5ccccc5n4)cc13

2628 Cc1ccc2C(=O)NC3CCCCN3c2c1

2629 O=C(NC1CCCC1)N2CCC(CC2)c3cnc[nH]3

2630 COc1ccc(Cl)cc1S(=O)(=O)N[C@@H]2CC[C@H](CC2)N3CCC(CC3)c4ccccc4OCC(F)(C)F

2631 COc1ccc(cc1)C2=C(CCC2)c3ccc(cc3)S(=O)(CO)N

2632 COc1ccccc1N2CCN(C\\C\C\\CNC(=O)c3ccc(cc3)c4ccccn4)CC2

2633 Cn1nc(C(=O)N)c2CCc3cnc(NC4CCC(CC4)C(=O)c5ccccc5)nc3c12

2634 COc1cc(ccc1Nc2ccc(Cl)c(Nc3ccccc3P(=O)(C)C)n2)N4CCC(CC4)N5CCCC5

2635 COC(=O)c1ccccc1CCC2NNCCC2

2636 CCn1nnc(n1)[C@H]2O[C@H]([C@H](O)[C@@H]2O)n3cnc4c(NC)cc(Cl)nc34

2637 FC(F)(F)c1ccc2c(NC(=O)Nc3ccccc3)ccnc2c1

2638 COc1cc(Cl)ccc1N2cCN(CCN3C=Nc4sc5CN(C)CCc5c4C3=O)CC2

2639 O=C1CCc2ccc(CCCCN3CCN(CC3)c4cccc5CCCc45)nc2N1

2640 CC1=CC[C@@H]2[C@@H](C1)c3c(O)cc(cc3OC2(C)C)C(C)(C)c4cccc(BC)c4

2641 NNc1ccc(cc1)S(=O)(=O)

2642 NNc1ccccc1S(=O)N

2643 Cn1nc(C(=O)N)c2CCc3ccc(NC4CCN(CC4)C(=O)c5ccccc5)nc3c12

2644 CNc1nc(CC)cn(Ncncccccc2)S(OO)(=O)N21

2645 Cn1ccc(n1)c2cccccC2NC3=NCCN3

2646 Fc1cccc2cccc(N3CCN(CCCCNc4ccc5CNC(=O)c5c4CC)CC3)c12

2647 CC[C@@]1(O)C(=O)OCC2=C1C=C3N(Cc4cc5cc6OCOc6cc5cc34)C2=O

2648 CCCN(CCC)C1CCc2ccc(O)cc2C1    c

2649 COc1cc2c(Oc3ccc(cc3F)C4=CN=C(Cc5ccccc5)N(C)C4=O)ccnc2cc1CCCCN6CCOCC6C

2650 COc1ccc(cc1S(=O)(=O)N[C@@H]2CC[C@@H](CO2)N3CCN(CC3)c4ccccc4OC(C)C)[N+](=O)[O]

2651 CCCCCCCCC(=CCC(=O)CCCCOCCC)

2652 CC[C@@]1(O)C(=O)OCC2=C1C=C3N(Cc4cc5cc6OCCc6cc5nc34)C2=O

2653 COc1cccc(c1)c2cc(NC(=O)C)cc(n2)n3nc(C)cc3C

2654 Cc1cccc(Cl)c1NC(=O)c2cnc(NC(=O)CCCCC)s2

2655 COC(=O)C1c2CCC(CC1c3ccc(Cl)c(Cl)c3)O2

2656 Nc1n[nH]c2cccc(c3ccNcNC(=O)Nc4cc(ccc4F)C(=O)Occc3)c12

2657 Clc1cccc(c1CNCCCN(CC32CCCN3)C2)OO

2658 CCCNC(=C)c1ccccc1NCc2c[nH]nn2

2659 CNN(CC)CCOC(=O)C(c1ccccc1)C2(O)CCCCC2

2660 COc1cc(Nc2nccc(Nc3onc(c3)C4CCCCC4)n2)cc(CC)c1OC

2661 CCCCC(CO)C(=O)c1ccccc1

2662 Nc1ccc(cc1F)S(=O)(=O)NO

2663 CCn1nnc(n1)[C@H]2O[C@H]([C@H](O)[C@@H]2O)n3cnc4c(N)ncncn43

2664 CC(C)c1cc(Oc2c(Cl)cc(CC(CO)O)cc2C)Bccc1O

2665 NS(=O)(=O)Oc1cccsccccscc1

2666 Fc1cccc[nH]cc(CCCN3CCC(CC3)c4ccccc4)ccc1

2667 CC1(N(CCc2cc(O)ccc12)c3cccc(Cl)c3)c4ccc(CCCN5CCCC5)cc4

2668 COc1ccccc1N2CCN(CCCCNC(=O)c3ccc4c(Cc(ccccc4))c3)CC2

2669 CCCN1C(=O)N(CCC)c2[nH]c(nc2C1=O)c3ccc(CCC(=O)Nc4ccc(BF)cc4)cc3

2670 CCCCN1C2=C(CCCCCC2)C=C(C(=O)NC3CCCCC3)C1CO

2671 COc1ccc(Cn2cccc2c3ccccc3)cc1

2672 Cc1ccc(cc1)N2N=C3N(C2=O)c4ccccc4N3C#N

2673 CN[C@H](Cc1ccccc1)C(=O)N2CCC[C@H]2C(=O)N(CCCCN=C(N)N)C(=O)c3nc4CCCCc4s3

2674 CCCCN1CCC[C@@H]1CN2NCC(Cc3ccc(Cl)cc3)c4ccccc4C2OO

2675 Fc1cccc2cccc(N3CCN(CCCCCc4ccc5CNC(=O)c5c4Cl)CC3)c12

2676 COc1cc2OC(OO)C=Cc2cc1[C@@H](CCCCCC)C

2677 CCCCC1CCN(CCCCC=O)c2ccccc2CCCC1

2678 COC(=O)C1=C(C)NC(=N)N(C1c2ccc(F)c(F)c2)C(=O)NCCCN3CCN(CC3)c4ccccc4CC

2679 C\\N=C\\1cOc2ccccc2C=C1 1

2680 COc1ccccc1N2CCN(CCCCNC(=O)NCc3ccccc3c4ccccc4Cl)CC2

2681 Cc1cc(c(O)c(C)c1NC2=NCCC2)C(C)(C)C

2682 CCOC(=O)c1ncn2c1[C@@H]3CCCN3C(OO)c4cc(OC)ccc24

2683 Oc1ccc2c(noc2c1)c3cc(BO)c(O)cc3

2684 Cc1cccc(NC(=O)Nc2ccc(cc2)c3csc4c(cnc(N)c34)C#CCn5CCCC5)c1

2685 Cc1cc(Cl)ccc1CC(=S)NNC(=O)C(O)(c2ccccc2)c3ccccc3

2686 [S+].[Sn]C(=S)NCCCNCNOCO

2687 CN1CCN(CC1)c2nc(C3CC(C(=O)NC3OO)c4c[nH]c5ccccc45)c6ccccc6n2

2688 CCCCCCC(C)(C)c1cc(O)c2C3CC(=O)CCC3C(CCCC)Oc2c1

2689 FC(F)(F)c1ccc2c(NC(=O)Nc3ccccn3)cccc2c1

2690 C(Cc1noc2cc=ccc12)N3CC(CC3)c4CC5(Cc4occcc5)CCC

2691 NCCCCCc1c[nH]cc1
2692 Cc1cccc(NC(=O)Nc2ccc(cc2)c3noc4ccnc(N)c34)c1
2693 CC(C)c1cc(nc(O)n1)c2ccc(F)c3ccccc23
2694 Nc1nc(N)c2c(Cl)c(CNc3ccc(cc3)C(=O)CC(CCC(=O)O)C(=O)O)ccc2c1
2695 O=C(NCCc1cccc1cN2C)C(=CC2)c3c[nH]c4ccccc34
2696 CC1(N(CCc2cc(O)ccc12)c3cccc(O)c3)c4ccc(OCCc5CCCC5)cc4
2697 NS(=O)(=O)Oc1ccc(I)cc1 1
2698 Cc1ccc2OC(=O)S(=C(Cl)c2c1)\\C=N\\c3ccc(cc3)S(=O)(=O)NC(=N)N
2699 CCCCCOc1ccccc1c2nnn(c2)C(=O)NC3CCCCC3
2700 CNc1cc2c(Nc3cccc(BB)c3)ccnc2cn1
2701 NS(=O)(CO)N
2702 NS(=O)(=O)c1ccc(NC(COCCCC(=O)))cc1
2703 Cc1cc2CNC(=O)c2cc1CCCCCN3CCN(CC3)c4cccc5ccccc45
2704 COc1ccccc1N2CCC(CC(O)CCNC(=O)c3occcccccnc3)CC2
2705 Clc1ccc(cc1CC)C(=O)C(CC=C)N2CCCC2
2706 COC(=O)c1ccccc1CCC2=NCC2
2707 CCC(c1ccc(cc1)S(=O)(=O)Nc2ccc3CC=C)Cc3c2
2708 COc1ccc(cc1)C(C)NS(=O)(OC)
2709 Clc1ccc(cc1)C(=O)OCCN2CCOCC2
2710 CC1NC(=Nc2cccc(CC)c12)
2711 CC(C)CC(NS(CO)(=O)c1c(F)c(F)c(F)c(F)c1F)C(OO)NO
2712 O=C1Nc2ccccc2N1C3CCN(CC3)c4CCCCC4
2713 CCCN(CCC)CCc1cccn2Nc(=O)Cn12
2714 CC(=O)N1CCC(CNC(=O)NC2NCC4CC(CC(C4CC2)C))CC1
2715 CCC(CC)n1c(C)cc2c3c(N)ccnNcnc3ccc12
2716 COc1ccccc1N2CCN(CCCCNC(=O)c3ccc(CCC)cc3)CC2
2717 CCCN(CCC)C1CCc2cccOcccc2C1
2718 Cc1cccc(c1)S(=O)(=O)NCCNN2CCC(CC2)c3nsc4cc(F)ccc34
2719 CCN(CC)CCCC(=O)C1(CCCCC1)C2CCCCC2
2720 COC[C@@H]1CCCN1N2NCN(c2=O)c3cccc(CC)c3
2721 COc1cc2ncnc(N[C@H](C)c3ccccc3)c2cc1OCCCCCCC(=O)N
2722 COc1ccc2[nH]c(C)c(CNC(C)C)c2c1
2723 COc1ccc(cc1OC)c2cc3cccn3c(Nc4ccccc4C(=O)N)n2
2724 Clc1cccc(Cc2oc3ccnc(NCNC4CCCCN4)c3n2)c1
2725 Clc1ccc(cc1)C(C2CCCC2)C3CCCCC3

2726 NNc1ccc(cc1)S(=O)(=O)NN

2727 CCCCCCNS(=O)(=O)NONC(=O)O

2728 N#Cc1ccc(cc1)C(c2ccc(cc2)C#N)n3cncc3

2729 C[C@H](N1CCN(C[C@@H]1C)C2(C)CCN(CC2)C(=O)c3c(C)ccnc3C)c4ccc(cc4)C(F)(F)

2730 COc1cc2ncnc(Nc3cccc(Cl)c3)c2cc1O

2731 Cc1cccc(c1)N2CCN(CCCCNC(NO)c3oc4ccccc4c3)CC2

2732 NNc1ccccc1S(=O)NOO

2733 O=C1CCCC1CCCCCN2CCCC2

2734 CNC(=O)[C@H]1O[C@H]([C@H](O)[C@@H]1O)n2cnc3c(NC)nc(Cl)cc23

2735 COc1ccccc1N2CCN(CC(O)CCCC(=O)c3ccc4c(Cc5ccccc45)c3)CC2

2736 CCC[C@H]1C=CCN1c2ccc3[nH]ncc3c2

2737 Brc1c(NC2=NCCNC)ccccncc2c1

2738 COS(=O)c1ccccc1CCC2NNCC2N

2739 Cc1ccccc1c2cccn2Cc3ccc(cc3)C#N

2740 Clc1cc2NC(=O)Cc2cc1CCN3CCN(CC3)c4nnc5cccccc54

2741 CN1CCC[C@H]1[C@H]2C([C@@](O2)CC3CCCCC3)c4ccccc4

2742 Cc1ccc(OCCCCCCOc2ccccc2)cc1Cl

2743 CN(C)Cc1ccccc1Sc2ccc(CO)cc2C

2744 COc1ccc(cc1OC)S(=O)(=C)N[C@@H]2CC[C@@H](CC2)N3CCC(CC3)c4ccccc4OCC(F)(F)F

2745 CCCc1nn(C)c2C(=O)NC(=Nc12)c3cc(ccc3OCC)S(=O)(NOC)CCCCCCO

2746 Nc1ncnc2c1c(cn2C3CCcC3)C

2747 COc1cc2ncnc(Nc3ccc(F)c(c3)C#C)c2cc1OCCCCCCC(=O)OO

2748 COc1ccc(SC)c2NCC2CNCCNCCc1

2749 COc1ccc(NC(CS)NC(=O)c2cn(nc2c3ccc(Cl)cc3)c4ccccc4)cc1

2750 O=C1CCc2ccc(OCCCN3CCN(CC3)c4cccc5ccccc45)cc12

2751 Nc1ncnc2c1c(cn2C3CCCC3)c4c5c(Occccccc5)cc4

2752 CCCCCCCCC(=CCC(=O)CCC=CCCC)

2753 Clc1ccc(cc1)C(=C)CCCN2CCc3ccccc3C2N

2754 CCc1nc(O)nc(N)c1c2ccc(CC)cc2

2755 OC1(CCOCC1)c2cccc(COc3ccc4c(cc(cc4c3)CCN)c5cocc5)c2

2756 CN.COc1ccc(F)c(CC)c1C2CC2

2757 Clc1cccc(Cc2oc3ccnc(NCCC4CCCNN4)c3n2)c1

2758 CC(Oc1ccccc1CC=C)2C=NCCN2

2759 COc1cccc2cC(Ccc12)NS(=O)(=O)N(C)C

2760 Cc1ccc(cc1)C(=O)CCCCOCCC2CCCC2

2761 CCCOC[C@@]1[C@@H]2CNC[C@]12c3ccc(Cl)c(Cl)c3
2762 CCCCCN(C)CCC(OO(P(=O)(OPO)PPOO))OO
2763 Cc1c(cc(c2ccccc2)n1C)C(CO)NCCCN3CCN(CC3)c4cccc(Cl)c4
2764 Cc1ccc2c(cccc2n1)N3CCN(CCc4cccc5c4OCc6c(nnn56)C(=O)NC7CCC7)CC3
2765 CN(C)[C@@H]1C(C(=C)1)c2c[nH]c3ccc(cc23)C#N
2766 CC(=O)N(O)C\\CCC\\c1cccc(Oc2ccccc2)c1
2767 CCCC(C1CCCo1)C(=O)c2ccc(CC)cc2
2768 Cc1ccccc1NC(=O)Nc2ccc(cc2)c3coccnc4c(N)c34
2769 Clc1cccc(Cl)c1NC2CNCCN2
2770 C1CCC(CNC1)NC2CCCccc3ccc23
2771 COC(=O)C1=C(C)NC(=N)N(C1c2ccc(F)c(F)c2)C(=O)NCCCN3CCN(CC3)c4ccccc4C
2772 COc1cc2ncnc(Nc3cccc(BC)c3)c2cc1CC
2773 O=C1NC(=O)\\CC=C\\cncccnccccccccccc1
2774 COc1ccc(cc1OC)S(=O)(CO)N[C@@H]2CC[C@H](CC2)N3CCC(CC3)c4ccccc4OC(C)C
2775 Cc1c(NC2=NCCN2)ccc3NCCCc13
2776 C[C@H](N)CN1CCc2ccc(BB)cc12
2777 O=C(NCCCCN1CCN(CC1)c2ccccc2)c3ccccncccccc3
2778 CCCN(CCC)C1CCC(=CC1)CCC
2779 OC1(CCN(CCCOc2ccc(F)cc2)CC1)c3ccc(CC)cc3
2780 COC1(CC1)C(=O)N[C@@H]2CC[C@@H](CCN3CCN(CC3)c4cccc5OOOc45)CC2
2781 CN1CCN(CC(=O)Nc2ncc(s2)S(=O)(=O)N)CC1
2782 O=c1CCCN1CCCCCC2CCCC2
2783 CN[C@H](Cc1ccccc1)C(=O)N2CCC[C@H]2C(=O)NC(CCCN=CCNNNNC(=O)c3cc4cccccccs43)C(=O)OC
2784 Clc1cccc(OC(C2CCCCC2)c3ccncc3)c1Cl
2785 Nc1ncnc2c1c(nn2C3CCCC3)c4cnc5[nH]ccc5n4
2786 CCc1cc2[C@@H]3CNCCN3C(=O)c2cc1CC
2787 CC1CCCCc2cc(Cl)c(Cl)cc12
2788 OC1(CC2CCC(C1)N2CCCNc3cccc(F)c3)c4ccc(C)Ccc4
2789 COc1cc(Nc2oc(cn2)c3ccccc3)cccccocnc1
2790 NS(=O)(=O)c1ccc(cc1)C2=C(C(=O)OC2)c3ccccc3
2791 CC1=CC[C@@H]2[C@@H](C1)c3c(O)cc(cc3OC2(C)C)C(CCCCCC4CccCCCC4)
2792 Cc1cc(C)c2c(n1)sc3c(N)cccc23
2793 COc1ccc2ccc3oc(CNC(=O)C3)cC2ccc1
2794 Fc1ccc2cccc(N3CCN(CCCCOc4ccc5CNCc5c4)Cc3)c2c1
2795 Cl.CC(C)Sc1ccccc1N=C2CCNCN2

2796 OC1(CCOCC1)c2ccnc(COc3ccc4c(cc(cc4c3)CNN)c5cocc5)c2
2797 COC(=O)c1ccccc1CCC2NNCCN2
2798 CC(C)c1ccc(cc1)N2CCc3cc(O)ccc3C2OCOc4ccc(OCCN5CCCC5)cc4
2799 C[C@H](N)Cc1c[nH]c2cc3ccCCCC3c12
2800 CNCCC(Oc1ccccc1)CCc2ccccc2
2801 COc1cc2nc(nc(N)c2cc1OC)N3CCN(CC3)C(OO)CCCCC4CCCC4
2802 COc1ccc2ccc3oc(CNNC(=O)C4CC4Ccc3)2c1
2803 Cc1cccc(NC(=O)Nc2ccc(cc2)c3coc4ncCC(N)c34)c1
2804 Clc1cc(N)cc(Cl)c1N=C2NCCN2
2805 C\\N=C\\1/Oc2ccccc2C=C1O
2806 NC(c1cccccS1)S(=O)(=O)
2807 Cn1cc(C=C2C(=O)Nc3ccccc23)c4ccccc14
2808 Brc1ccc(CN(c2ccc(cc2)CCN)n3nncc3)cc1
2809 OP(=O)(O)C(NC1CCCCC1)P(=O)(O)N
2810 Cc1cccnc1NC(P(=O)(O)O)P(=O)(O)OO
2811 CN(C)Cc1(CCCCC1)c2cccc3ccccc32
2812 CNc1cc(ccc1Cc2cn(C)c3ccc(NC(=O)OC4CCCC4)cc23)C(=O)NS(=O)(=O)c5ccccc5C
2813 Cc1ccc2c(cccc2n1)N3CCN(CCc4cccc5c4OCc6c(ncn56)C(=O)NC7CCC7)C3
2814 Clc1cccc(OC(C2CCCCC2)c3ccccc3)c1Cl
2815 Cl.CC(C)Sc1ccccc1NCC2CCCCN2
2816 CC(Oc1ccccc1C2CcCC2)C3=NCCN3
2817 NC(=O)N(O)CCC#Cc1ccc(OCCCN2CCN(CC2)[C@@](c3ccccc3)c4ccc(Cl)cc4)cc1
2818 COc1ccc(Cl)cc1S(=O)(=O)N[C@@H]2CC[C@H](CC2)N3CCC(CC3)c4ccccc4OCC(F)(F)
2819 C1COC(=CC1)NC2CCCc3ccccc23
2820 Cc1c(OCCCN2CCN(CC2)c3cccc4ccccc34)ccc5CCC(OO)c15
2821 COc1ccc(cc1)C(=O)CCCN2CCc3cccccC32
2822 CN1CCc2c(Cl)ccc3oc(BO)c(C1)c23
2823 CC(C)NC(C)C(O)NOc1ccc(C)c2CCCc12
2824 CN1CCc2c(C1)c3cc(C)ccccn2CCcCccccc3
2825 CC1=CN=C(NCC(F)(F)c2ccccn2)C(=O)N1CC(C)OCCc3ccc(N)nc3C
2826 Fc1cccc2c(NC(=O)Nc3cccc(n3)C(F)(F)F)cccc12
2827 NC(=O)c1cc2c(Oc3ccc(BC)cc3)cncc2s1
2828 COc1cc(ccc1Nc2ncc(Cl)c(Nc3ccccc3P(=O)(C)C)n2)N4CCC(CC4)NCCCCC
2829 COC(=O)C(NS(=O)(=O)NCCC)c1ccc1
2830 Brc1ccc2[nH]cc(CCN(CC=O)CC=C)c2c1

2831 Clc1cccc(c1)N2CCN(CC3CCCCN3)C2CO

2832 NCCNc1cccccc2ccc12

2833 CC(C)NCC(O)COc1ccccc1CCCC

2834 O=C1CCc2ccc(OCCCCN3CNN(CC3)c4cccc5CCCCc45)nc2N1

2835 C\\N=C\\1OOc2ccccc2C=C1

2836 OC(Cc1cccnc1)(P(=O)(O)O)P(OO)(O)O

2837 COc1ccccc1N2CCN(CCCNS(=O)(=O)c3cccccccccc3)CC2

2838 CCCCCCCCC(=O)C(=O)CCCCC

2839 FC(F)(F)c1ccc2c(NC(=O)Nc3ccccn3)ccnc2c1

2840 CN(C)CCCN1c2c3ccc2CCc3Ccccc1

2841 C(CC1(CCNC1)c2cccn[nH]ccccc2)c4ccccc4

2842 CCN(CCCCCN1CCN2C1=O)cccccc(C)Cc2

2843 COCC(=O)N[C@@H]1CC[C@@H](CCN2CCCCCC2Cc3coc4ccccc34)CC1

2844 CC(C)Oc1ccccc1N2CCN(CC2)[C@@H]3CC[C@H](CC3)NS(OO)(=O)c4ccccc4

2845 COc1ccc(cc1OC)C2=NCC(=O)[C@@H]3CC=CC[C@H]23

2846 Cc1cc(cc(C)c1CC2CNCC=2)C(C)(C)

2847 COc1cc2c(Nc3ccc(cc3)c4nc5ccccc5s4)nccc2cc1OCCCN6CCN(C)CC6

2848 Cl.CCCCC(C1CCNCC1)c2ccc(Cl)c(Cl)c2

2849 COc1cc2nc(nc(N)c2cc1OC)N(C)C2CNC(OO)CCCCO2

2850 Nc1ccc(cc1Cl)S(=O)(=O)

2851 CCCSC(=O)c1ccccc1NCc2c[nH]cc2

2852 Clc1cc2NCCCNCCcCc2cc1C

2853 CS(=O)(=O)c1ccc(cc1)c2=C(CC3(CCC3)C2)c4ccc(F)cc4

2854 CCc1cc2[C@@H]3CNCCN3C(=O)c2cc1NC

2855 Cn1c(SCCCN2CC3CCN(C3C2)c4ccc(cc4)C(F)(C)F)nnc1C5CCCCC5

2856 Clc1cccc(Cl)c1NCC=NCCN

2857 CCCN(CCC)C1CCn(OCCO)1

2858 NC(=N)c1ccc(CNC(=O)[C@@H]2CCN2C(=O)[C@H](CCC(=O)O)C3CCCCC3)cc1

2859 C(CN1CCCC1)Cc2ccccOc3ccccc3ccc2

2860 COC(=O)C1C2CCC(CC1c3ccc4ccccc4c3)c2C

2861 Clc1cccc(c1)NCCCNCCCC

2862 CCCNC(=O)c1ccccc1NCc2c[nH]cc2

2863 Brc1c(NC2C=CCN2)ccc3NCCNc13

2864 CC1=CN=C(NCCc2ccccc2)C(=O)N1CC(=O)NCc3ccc(N)cc3C

2865 CC1=CC[C@@H]2[C@@H](C1)c3c(O)cc(cc3OC2(C)C)C4CC(C)c44ccCCC4

2866 CN(C)CC1=C(O)C2=CC(=CC(=O)N2CNC1)N

2867 NCCc1c[nH]c2ccc(C)cc12

2868 NS(=O)(CO)c1ccc(CCl)cc1

2869 Oc1ccc2C3=C(CCOc2c1)c4ccc(O)cc4O[C@@H]3c5ccc(OCCC6CCCCC6)cc5

2870 FC1(F)CC1C(=O)N[C@@H]2CC[C@@H](CCN3CCC(Co3)c4coc5ccccc45)CC2

2871 Fc1ccc2c(noc2c1cc3CCN(CCCCNS(=O)(=O)c4cc5ccccc5s4))C3

2872 CC(=O)N[C@H](Cc1cccc=1)C(=O)N2CCC[C@H]2C(=O)N[C@@H](CCCN=C(N)N)B(O)O

2873 CCCCCCC(C)(C)c1cc(O)c2(OCCCCCCCCCCC(=O)CCCCC2)c1

2874 COc1cccc(c1)[C@@H]2CC[C@H](CC2)N3CCC(CC3)c4ccccc4

2875 CCCCCCC(C)(C)c1cc(O)c2C3CC(CO)CCC3C(CC(C))C2cc1

2876 Cc1ccc2c(cccc2n1)N3CCN(CCc4cc5NcC(O)COc5cc4F)CC3

2877 Cc1ccc(cc1)C(=O)C(CCNC)N2CCCC2C

2878 CC(C)c1cc(nc(O)c1)c2ccc(F)c3ccccc23

2879 CCN(CC)CCCCNc1ncc2cc(c(NC(=O)NC(CCCC)C)cc2n1)c3c(CC)cccc3Cl

2880 NS(=O)(=O)ONC(=O)NCc1ccccc1

2881 Clc1cccc(N2CCN(CCCCS(=O)(=O)c3ccc4cccnc4c3)CC2)c1Cl

2882 COc1ccc(cc1Oc2CCCC23CCCNCC=O)cc3

2883 Cc1cc2CC(Nc3ccccc3Nc2s1)N4CCNCC4

2884 O=C1CCc2ccc(OCCCCN3CCN(CC3)c4cccc5CCCOc45)nc2N1N

2885 CC1=CC[C@@H]2[C@@H](C1)c3c(O)cc(cc3OC2(C)C)C(C)CCCCCCCCCCCC

2886 Clc1cc(I)cc(Cl)c1NCC2NCCN2

2887 Fc1ccc2c(noc2c1)C3CCN(CCCNS(=O)(=O)c4ccc5ccccn45)CC3

2888 CC(C)n1nc(c2ccc3C)c(O)c2cc3c(N)ncc1

2889 CC(CO)c1cc(NC(=O)NCCC[C@H]2C[C@H](Cc3ccc(F)cc3)CCC2)cc(c1)C(=O)C

2890 CN1C(=O)C(=Cc2ccc(Nc3ccc(C)cc3)nc12)c4c(Cl)cccc4Cl

2891 CN(C)CC1CC2N(O1)c3ccc(F)cc3cccccccc2

2892 COc1cc(NC(=O)C(=O)NC(C)(C)C)ccc1c2onnc2

2893 CCCNC(=O)c1ccccc1NCc2c[nH]nn2

2894 CN[C@H](Cc1ccccc1)C(=O)N2CCC[C@H]2C(=O)N[C@@H](CCCN=C(N)N)C(OO)c3nccs3

2895 CC(C)Oc1ccccc1N2CCN(CC2)[C@@H]3CC[C@H](CC3)NS(=O)(=O)c4ccc(Cl)cc4N

2896 CC(=O)N[C@H](Cc1ccccc1)C(=O)N2CCC[C@H]2C(=O)N[C@@H](CCCNCC(N)N)B(O)O

2897 C=CCNNCC(O)COc1ccccc1C =C

2898 Fc1ccc(OCCCNCCOc2ccc(CC)cc2)cc1

2899 CC(=O)Nc1cc(nc(n1)c2occc2)n3nccn3

2900 COc1ccc(cc1)C(C)NS(=O)(=O)NO

2901 CC(=O)c1cccc(NC(=O)NCCCN2CCC[C@@H](Cc3ccc(C)cc3)C2)c1
2902 CC(C)(C)OC(=O)N[C@@H](C(=O)N1CCC[C@H]1C(=O)NC(CCC(=CCN)N)C=O)c2cccccccccc2
2903 CCCCCCNS(=O)(=O)NCNC(OO)O
2904 COc1cc2ncnc(Nc3ccc(F)c(Cl)c3)c2cc1OCCCCCC(=O)NN
2905 COc1ccc(cc1OC)S(=O)(=O)N[C@@H]2CC[C@H](CC2)N3CCC(CC3)c4ccccc4OCC(F)(N)F
2906 COc1ccc2CC(CCc2c1)NS(=O)(=O)N(C)
2907 CC(C)Oc1ccccc1N2CCN(CC2N[C@@H]3CC[C@H](CC3)NS(=O)(=O)c4ccccc4[N+](=O))OO
2908 CNc1cc2c(Nc3cccc(BO)c3)cnnc2cn1
2909 Cc1c(cc(c2ccccc2)n1C)C(=O)NCCCN3CCC(CC3)c4cccc(Cl)c4Cl
2910 Cc1ccc(cc1)C(CC2=NCCN2)c3cccc(O)c3
2911 NS(=O)(=O)c1ccc(cc1)c2cccccC(CC=CO)2CC
2912 CC(C(=O)O)c1ccc(c(O)c1)c2ccccc2
2913 COC(=O)[c@H]1C2CCC(C2)C[C@@H]1c3ccc4ccccc4c3
2914 C(Nc1ccccc1c2nnccn2)C3=NCCN3
2915 COc1ccccc1OCCNCCCc2c[cH]n3ccccc23
2916 Oc1ccc2C3=C(CCOc2c1)c4ccc(F)cc4O[C@@H]3c5ccc(OCCNCCCCCC)CC5
2917 Clc1cccc(c1)N2CC2CCCC
2918 CN1N=C(SC1(N)S(=O)(=O)N)
2919 Cc1nc[nH]c1CC
2920 CC(CCc1ccccc1ONCCCO)c2ccc(O)c(c2)C(=O)C
2921 Cc1ccc2[nH]cc(CCN(CCCC)CC=C)c2c1
2922 Clc1ccc(CNC2=C(Nc3ccncc3)C(=O)C2CO)cc1CC
2923 COc1ccc2[nH]c(N)c(CCN(C)C)c2c1
2924 Clc1ccc(CCC2=C(Nc3ccncc3)C(=O)C2=O)cc1
2925 Nc1nc(Nc2ccc(cc2)S(=O)(=O)N)nn1C(=O)c3c(F)cccc3
2926 Cc1ccc2c(cccc2n1)N3CCN(CC34cccc(C4)n5cccn5)CC
2927 CN1CCc2c(C1)sc3N=CN(CCN4CCN(CC4)c5cccc6ccnnc56)C(=O)c23
2928 Clc1ccc(CNC2=C(Nc3ccncc3)C(CC)C2=O)cc1Cl
2929 Nc1ncnc2c1c(cn2C3CCCC3)C C
2930 CN(C)C[C@H]1C2CCC(C2)[C@@H]1c3cccccccccc3
2931 C[C@H](C1CC(CCNCCCC)Cc2ccccc12)c3cncnn3
2932 CNC(=O)CCCN(C)C(=O)c1ccc2c(c1)c3C[C@@H]CCCc3n2CCC4CCCCC4
2933 Cl.Clc1ccccc1CC(N2CCCCC2)c3ccccc3
2934 COc1ccccc1N2CCN(CCCCNS(=O)(=O)c3cccccccccc3)CC2
2935 Cc1ccc2c(cccc2n1)N3CCN(CCc4ccc5CCCC(O)Nc5c4)CC3

2936 Nc1ncnc2c1c(cn2C3CCcC3)C#C

2937 FC(F)(F)c1ccc2c(SC(=O)Nc3ccccc3)ccnc2c1

2938 COc1cc2ccccc2cc1n2ccnc2

2939 OCC[C@@]1(CCCNNC1)c2ccc(Cl)c(Cl)c2

2940 CN(C)CC1=C(O)C2=CC(=CC(=O)N2CCC1)

2941 COc1cc(Nc2nccc(Nc3onc(c3)C4CCCCC4)n2)cc(CC)c1OO

2942 CCSc1ccc2CnN(C[C@H](C)N)c2c1

2943 Oc1cccc2C[C@@H]3[C@H](CCC3CC=C)Cc12

2944 O[C@@H](CNCCCCCCCCN1CCC(CC1)OC(=O)Nc2ccccc2c3ccccc3)c4ccc(O)c5CC(=O)C=Cc45

2945 CCCC(=O)Nc1n[nH]c2cnc(cc12)c3cccc(F)c3F

2946 ClNc1ccccc1c2cccNCCCCCCNCCN2

2947 COc1cc2c(Oc3ccc(NC(=O)N\\N=C\\c4ccc(F)cc4F)cc3F)ccnc2cc1OCCCN5NCC(C)CC5

2948 C(CN1CCC(Cc2ccccc2)CC1)Cc3c[nH]c4ccc(cc34)n5cncc5

2949 CN1CCc2ccccc2Cc3ccccc3CC1C

2950 COc1cccc(c1)C(C)NC(=O)c2ccc(cc2(CCN))c3ccncc3

2951 COc1ccccc1N2CCN(CCCCNC(=O)c3cc4ccccnn34)CC2

2952 NCCc1ccc(cc1)S(=O)(=O)

2953 C[C@H](N)Cn1CCc2ccc(BB)cc12

2954 Clc1cccc(N2CCN(CCCCOc3cc4NC(=O)C=Cc4cn3)CC2)c1C

2955 CN(C)CCCOC(=O)OC(C1CCCCCCC1)c2ccccc2

2956 O=C1NCc2ccc(OCCCNN3CC(CC3)c4cccc5CCCCc45)cc12

2957 Brc1ccc2[nH]c2n3CC(=O)Nc4ccnncc43ccc1

2958 COc1cc2ncnc(Nc3ccc(F)c(c3)C#C)c2cc1OCCCCCCC(CO)NO

2959 COc1cccc(c1)C(C)NC(OO)c2ccc(cc2)c3ccncc3C

2960 Cc1c(OCCCN2CCN(CN2)c3cccc4ccccc34)ccc5CCC(OO)c15

2961 Cc1cccc(c1)C(=O)NCN2CCN(CC2)c3ccccc3CCN

2962 CCCCOc1c2NC(=O)C(OCc2ccc1OO)C(=O)CCCCC

2963 CN(C)CC\\CCC/1\\c2ccccc2Cc3ccc(Cl)cc13

2964 FC(F)(F)c1ccc2cccc(CC(NO)N3cccccn3)c2c1

2965 NCCo1ccc2ccc(O)c1c2

2966 CCCC(OO)c1cc(c2OCC(C)(C)c2c1)C(C)(C)C

2967 COc1ccccc1N2CCN(CCCCN3C(=O)c4cccc43CC=C)CC2

2968 Cc1ccc(cc1)c2cc(nn2c3ccc(cc3)S(=O)(=O))NC(F)(F)F

2969 COc1ccc2CC(CCc2c1)NS(=O)N(O)C

2970 CC1=CC[C@@H]2[C@@H](C1)c3c(O)cc(cc3OC2(C)C)C4(CCCC4)CCN

2971 COc1ccc(cc1OC)c2cc3nccn3c(Nc4ccccc4C(OO)N)n2
2972 COc1ccc(OCCCCCCOc2ccccc2)cc1
2973 CCc1cc2[C@@H]3CNCCN3c(=O)c2cc1
2974 NCCNc1cccc2ccccc12 C
2975 CN(C)Cc1ccccc1Sc2ccc(cc2N)CCN
2976 COc1cccc(OC)c1OCCNC[C@@H]2COOCcc3cc3O2
2977 CC(C)Oc1ccccc1N2CCN(CC2)[C@@H]3CC[C@H](CC3)NS(=O)(=O)c4ccccc4CCN
2978 O=C1CCc2ccc(OCCCCN3CCN(CC3)c4cccc5CCCc45)cc12
2979 COc1ccccc1N2CCN(CCCCNC(=O)c3cccBBcccc3)CC2
2980 COc1ccccc1N2CCN(CCCNC3CC(C)C(=O)N(C)C(OO)N3C)CC2
2981 NCCC(C(=O)Nc1ccc2cnccncc2c1)c3ccc(Cl)c(CC)c3
2982 COc1cccc(c1)c2cn(c3ccccc3)c4nccc(N)c24
2983 CC1=CC(C)(C)Ncccc(cc12)c3cccc(c3)s2CNCNOOC
2984 Fc1c(Cl)cccc1N2CCN(CCCCOc3ccc4CCC(=O)Nc4c3)CC2
2985 C1COC(=CC1)CC2CCCc3ccccc23
2986 O=C1CCCN1CCCCCN2CNCC2
2987 COc1cc(NCc2ccc3nc(N)nc(N)c3c2C)cc(OO)c1OC
2988 CN(Cc1ccc2Nc(=NC(=O)c2c1)C)c3ccc(s3)C(=O)C[C@@H](CCC(=O)O)C(OO)O
2989 CN(C)C(=O)N[C@H]1Ccc2ccc(O)cC2C1
2990 CC1=CN=C(NCCc2ccccc2)C(=O)N1CC(=O)NCc3ccc(N)cc3
2991 CC(C)NCC(C)CCc1ccccc1CC=C
2992 COc1ccc(CCNS(=O)(=S)N)cc1
2993 O=C1NC(=NC(=C1)c2ccncc2)NCCCCCCCCC
2994 CC1=C(CCN2CCC(CC2)c3coc4cc(F)ccc34)C(=O)N5CCCCC5=N1
2995 Clc1ccc(cc1Cl)[C@]23CCC[C@H]2C3
2996 COc1cccc(F)c1C2CNCc3cnc(Nc4ccc(C(=O)O)c(OC)c4)nc3c5ccc(Cl)cc25
2997 COc1cccc(OC)c1OCCNC[C@@H]2Ccc3ccccc3c2
2998 Cc1ccc2c(cccc2n1)N3CCN(CCCc4ccc5OCC(=O)Nc5c4)CCC3
2999 CN(C)CCCCOC(=O)C(CCCC1CCCCC1)c2ccccc2
3000 OCCNc1cc2cc(cc22cn1)c3cccnc32
3001 Nc1ncnc2c1c(cn2C3CCN(CCO)C=3)c4cccc(O)c4
3002 COc1cc2nc(nc(N)c2cc1OC)N3CCN(CC3)C(=O)C4CCOO4
3003 COc1ccccc1N2CCN(CC(O)CCNC(=O)c3occcccccnc3)CC2
3004 CN1CCN(CS(=O)Nc2ncc(s2)S(=O)(=O)N)CC1
3005 OC1(N2CCCN=C2c3cnccc13)c4ccc(Cl)cc4

3006 Nc1ccc(cc1O)c2nn(n3CCCC3)c4ncnc(N)c24

3007 COc1cc2c(Oc3ccc(Nc4ccc(cc4)C(C)(C)N)cc3)ccnc2cc1OCCNCCO

3008 CCCCCOc1ccccc1c2onc(c2)C(CO)NC3CCCCC3

3009 COc1cc(ccc1Nc2ncc(Cl)c(Oc3cccc(NC(=O)C=C)c3)n2)NCCCN(C)CC

3010 COc1cccc2c(CCCNCCOc3ccccn3)cccc21

3011 OCCNc1cc2cc(ccc2cc1)c3cccnc3

3012 N[C@@H]1CC[C@H](CC1)NC(=O)c2cc(OCc3cccc(c3)C(=O)N)cc(OCc4cccc(c4)C(=N)N)c2

3013 CN1N=C(S/C/1=NCC(=O)C)S(=O)(=O)N

3014 CCCCCCCCCCCOOC(=O)CCCCCC

3015 CCCCCCCCCCOS(OO)(=O)N

3016 Cc1cccc(NC(=O)Nc2ccc(cc2)c3csc4cc(N)nc(N)c34)c1

3017 COc1ccccc1CNCCCCCCNNNCCCCCCNCCCCCCNCc2ccccc2OC

3018 Cc1cccc(NC(=O)Nc2ccc(cc2)c3csc4cccc(N)c34)c1

3019 CNC(=O)Oc1cccc(CNCC)COCOcccccccc3cc3ccccccCc1

3020 COC(=O)CCCS(=O)(=O)NCc1ccc(F)cc1

3021 Nc1ccc2c(CCCCCCN(Cc4ccccc4)CCCC)oo2c1

3022 O[C@@H](CNCCCCCCCCN1NCC(CC1)OC(=O)Nc2ccccc2c3ccccc3)c4ccc(O)c5CC(=O)C=Cc45

3023 Cl.COc1ccc(F)c(Cl)c1C2CC2C

3024 COc1ccc(cc1)c2cc(nn2c3ccc(cc3)S(=O)(=O)O)C(F)(F)F

3025 CC(C)Oc1ccccc1N2CCN(CC2)C3CCC(CC3)NC(=O)Nc4ccccCCccc4

3026 SS(=O)(=O)c1ccc(CC)cc1

3027 COc1ccccc1N2CCN(CCCCc3cn(cc3)c4ccccc4)CC2

3028 C(Nc1ccccc1c2cccs2)C3=NCCC3

3029 NS(=O)(=O)Oc1ccc(NC(=O)NCC)2ccccccccccc2cccc1

3030 N\1N=C\\1/Oc2ccccc2C

3031 COc1cc2c(Oc3ccc(NC(=O)C4=NN(c5ccc5ccC4=O)c6ccccc6C(F)(F)F)cc3F)cnnc2cc1CCCCC7CCC[illegible]

3032 Cc1c(sc2ccc(F)cc12)S(=O)(=O)NNCCCC3CCC(CC3)c4noc5cc(F)ccc45

3033 CCCCc1nc(N)c2nc(n(C)c2n1)n3ccnn3

3034 CCNC(=O)[C@H]1O[C@H]([C@H](O)[C@@H]1O)n2cnc3c(Nc4ccc(OCC(=O)Nc5ccc(cc5CC(C)CC)C)[illegible]

3035 NC(=O)c1cccc(c1)c2cccc(CC3=O)NC3CCCCCCCc2

3036 COc1ccc(CCNS(=N)(=O)N)cc1

3037 CC(C)Oc1ccccc1N2CCN(CC2)[C@@H]3CC[C@@H](CC3)NS(=O)(=O)c4ccccc4[N+](=O)[O]

3038 CN[C@H](Cc1ccccc1)C(=O)N2CCC[C@H]2C(=O)N[C@@H](CCCNC(=N)N)C(=O)c3nc4ccccc4C3

3039 CNc1ccc2[nH]cc(CCN(C)C)c2c1

3040 CC(=O)N(O)C\\CCCC\c1cccc(Oc2ccccc2)c1

3041 Cc1ccc(F)c(C(=O)n2nc(Nc3ccc(cc3)S(=O)(=O)N)nc2C)c1

3042 CCCCC(CC(=O)NO)S(=O)(=O)NCCCCCCC

3043 COc1ccccc1N2CCN(CCCCC(=O)NCc3ccccc4c3ccccc4Cl)CC2

3044 CC(C)NCC(O)COc1ccccc1CCCCC

3045 CCOc1ccc(C=C2SC(=N)NC2=O)c(N)c1

3046 CC1=CC(C)(C)Nc2ccc(cc12)c3ccc(C=O)C3

3047 CCN(Oc1ccncc1)C(=O)C(CO)c2ccccc2

3048 CC1NC(=Cc2cccc(Cl)c12)N

3049 Oc1ccc2CC[C@H](CNC3cccccc3)Oc2c1

3050 Cl.CC(C)Sc1ccccc1NCC2NCNCN2

3051 COc1cc2c(Oc3ccc(NC(=O)c4cc(ccn4)c5ccc(C)cc5)cc3F)ccnc2cc1OCCCN6NCN(C)CC6

3052 CCc1cc(Oc2c(I)cc(CC3NC(=O)NC3OO)cc2I)ccc1O

3053 Cl.CCOC(=O)CCc1ccc(OC[C@H](O)CCC(C)(C)Cc2ccc3ccccc3c2)c(c1)I#N

3054 COc1c(Cl)cccc1N2CCN(CCCCNC(=O)c3cccccccccc3)CC2

3055 COc1ccc(C=NNc2ncnc3cccccccc[nH]c23)cc1OC

3056 N[C@@H]1CC[C@H](CC1)NC(=O)c2cc(OCc3cccc(c3)C(NN)N)cc(Oc4ccc(cc4)C(=N)N)c2

3057 CC(Oc1ccccc1C2CCCC2)C3CNCCN3

3058 Nc1ccc(cc1O)c2nn(C3CCCC3)c4nccc(N)c24

3059 Cl.CC(C)Sc1ccccc1NCC2CCNNN2

3060 O[C@@H](CNCCCN1CCC(CC1)OC(=O)Nc2ccccc2c3ccccc3)c4ccc(O)c5NC(=O)C=Cc54

3061 CC(C)Oc1cc(ccc1C(=O)O)c2ccc(CCCC[C@H](O)c3ccccc3)cc2

3062 COc1cc(ccc1O)CC=C(O)C(=O)c3c(O)Oc(OOcc3)C

3063 CC(=O)N1CCC(CC1)NC(=O)NC23CC4CC(CC(C4)C2)CC3

3064 CC(C)(C(=O)c1ccnnc2)c2ccccc1

3065 NCCCC(=O)C

3066 COc1ccccc1C2CC(NNN2C(=O)C)c3ccc(O)cc3

3067 NS(=O)(=O)c1ccc(cc1)n2nc(cc2c3ccccc3)C(F)(F)

3068 Fc1ccccc1COc2ccccc2C3CCCCC3

3069 C(Nc1ccccc1c2ncccn2)C3=NCCN3C

3070 C1CN(CCN1)C2=Cc3ccccc3Cn4cccc24

3071 Fc1ccc(cc1)C2CCN(CCCCNC(=O)c3cc4ccccc4cc3)CC2

3072 Cc1cc2C(=Nc3ccccc3Nc2s1)c4CCNC4

3073 C(CN1CCC(Cc2ccccc2)Cc1)Cc3c[nH]c4ccc(cc34)n5cncc5

3074 CC(N)C1CCC(CC1)C(=O)Nc2ccncn2

3075 COC(=O)C1=C(C)NC(CO)N(C1c2ccc(F)c(F)c2)C(=O)NCCCC3CCN(CC3)c4ccccc4

3076 CC1(N(CCc2cc(O)ccc12)c3cccc(C)c3)c4ccCCCCCC5CCCC5ccc4
3077 Cc1ccccc1NCCNCCCOc2ccccc2
3078 CCCN1C(=O)N(CC)c2nc([nH]c2C1=O)c3cnn(Cc4cccc(c4)C(O)(F)F)c3
3079 Nc1cc(F)ccc1C(=O)CCCN2CC[C@@]C3[C@H](C2)c4cccc5CCCN3c54

### 0.5 List of All Novel Ligands generated from cancer related anchor ligands.

Novel Smiles generated from cancer related anchor ligands.

COc1cc2ncnc(Nc3ccc(F)c(Cl)c3)c2cc1OCCCN1CCONC1
COc1cc2ncnc(Nc3ccc(F)c(Cl)c3)c2cc1OCCCN1CCOC1
COc1cc2ncnc(Nc3ccc(F)c(Cl)c3)c2cc1OCCCN1CC1C
COc1cc2ncnc(Nc3ccc(F)c(Cl)c3)c2cc1OCCCN1CCOCC1
COc1cc2ncnc(Nc3ccc(F)c(Cl)c3)c2cc1OCCCN1CC1
COc1cc2ncnc(Nc3ccc(F)c(Cl)c3)c2cc1OCCCNOCNCO
COc1cc2ncnc(Nc3ccc(F)c(Cl)c3)c2cc1OCCCC1CC1
COc1cc2ncnc(Nc3ccc(F)c(Cl)c3)c2cc1OOCCN1CCOCC1
CCOCNCOCCCCOc1cc2c(Nc3ccc(F)c(Cl)c3)ncnc2cc1OC
COc1cc2ncnc(Nc3ccc(F)c(Cl)c3)c2cc1OCCCN1CCOOC1
Cc1ccc(-n2cc(C(C)C)cc2NC(=O)Nc2ccc(OCCN3CCOCC3)c3ccccc23)cc1
Cc1ccc(-n2cc(C(C)(C)C)cc2NC(=O)Nc2ccc(OCCN3CCOCC3)c3ccccc23)cc1
Cc1ccc(-n2nc(C(C)C)cc2NC(=O)Nc2ccc(CCCN3CCOCC3)c3ccccc23)cc1
Cc1ccc(-n2nc(C(C)(C)C)cc2NC(=O)Nc2ccc(OCCN3CCCCC3)c3ccccc23)cc1
Cc1ccc(-n2nc(C(C)C)cc2NC(=O)Nc2ccc(OCCN3CCOCC3)c3ccccc23)cc1
Cc1ccc(-n2nc(C(C)C)cc2NC(=O)Nc2ccc(OCCN3CCOCO3)c3ccccc23)cc1
Cc1ccc(-n2cc(C(C)(C)C)cc2NC(=O)Nc2ccc(OCCN3CCCCC3)c3ccccc23)cc1
Cc1ccc(-n2nc(C(C)(C)C)cc2NC(=O)Nc2ccc(OCCN3CCOCC3)c3ccccc23)cc1
Cc1ccc(-n2nc(C(C)(C)C)cc2NC(=O)Nc2ccc(OCCN3COOCO3)c3ccccc23)cc1
Cc1ccc(-n2nc(C(C)(C)C)cc2NC(=O)Nc2ccc(OCCN3CCCCC3)c3ccccc23)cn1
CN(c1ncccc1CNc1nc(Nc2ccc3c(c2)CC(=O)N3)ncc1C(F)(F)F)S(C)(=O)=O
CCOc1nc(NC(=O)Cc2cc(OC)ccc2OC)cc(N)c1C#N
C#Cc1c(N)cc(NC(=O)Cc2cc(OC)ccc2OC)nc1OCC

```
CC#Cc1c(N)cc(NC(=O)Cc2cc(OC)ccc2OC)nc1OCC
CCOc1nc(NC(=O)Cc2cc(OC)ccc2OC)cc(N)c1CCNC
CCOc1nc(NC(=O)CC2=C(OC)C=CC=CCOC([SH](C)(=O)OC)=C2)cc(N)c1CC
CNC(=O)c1cc(Oc2ccc(NC(=O)Nc3ccc(Cl)c(C(F)(F)F)c3)cc2)ccn1
CN(Cc1cnc2nc(N)nc(N)c2n1)c1ccc(C(=O)N[C@H](C=O)CCC(=O)O)cc1
CN(Cc1cnc2nc(N)nc(N)c2n1)c1ccc(C(=O)N[C@@H](CCC(=O)O)C(=O)O)cc1
CCCc1cccc(Nc2ncnc3cc(OCCOC)c(OCCOC)cc23)c1
C#Cc1cccc(Nc2ncnc3cc(OCCOC)c(OCCO)cc23)c1
C#Cc1cccc(Nc2ncnc3cc(OCCOC)c(OCCOC)cc23)c1
C#Cc1cccc(Nc2ncnc3cc(OCCOC)c(OCCCC)cc23)c1
C#Cc1cccc(Nc2ncnc3cc(OCCOC)c(OC)cc23)c1
C#Cc1cccc(Nc2ncnc3cc(OCCOC)c(OCCC)cc23)c1
C#Cc1cccc(Nc2ncnc3cc(OCCOC)c(OCCOCC)cc23)c1
C#Cc1cccc(Nc2ncnc3cc(OCCOC)c(OCCOO)cc23)c1
C#Cc1cccc(Nc2ncnc3cc(OCCOC)c(OCC)cc23)c1
CS(=O)(=O)CCNCc1ccc(-c2ccc3ncnc(Nc4ccc(OCc5cccc(F)c5)c(Cl)c4)c3c2)o1
Cc1nc(Nc2ncc(C(=O)Nc3c(C)cccc3Cl)s2)cc(N2CCN(CCO)CC2)n1
CCc1cccc(C)c1NC(=O)c1cnc(Nc2cc(N3CCN(CCO)CC3)nc(C)n2)s1
CCCN1CCN(c2cc(Nc3ncc(C(=O)Nc4c(C)cccc4Cl)s3)nc(C)n2)CC1
Cc1cc(Nc2cc(Sc3ccc(NC(=O)C4CC4)cc3)nc(N3CCN(C)CC3)c2)n[nH]1
Cc1cc(Nc2cc(N3CCN(C)CC3)nc(Sc3ccc(NC(CO)C4CC4)cc3)n2)n[nH]1
Cc1cc(Nc2cc(N3CCN(C)CC3)nc(Sc3ccc(NC(OO)C4CC4)cc3)n2)n[nH]1
Cc1cc(Nc2cc(N3CCN(C)CC3)nc(Sc3ccc(NC(=O)C4CC4)cc3)n2)n[nH]1
CCn1c(-c2nocc2N)nc2cnc(Oc3cccc(NC(=O)c4ccc(CCCN5CCCCC5)cc4)c3)cc21
CCn1c(-c2nonc2N)nc2cnc(Oc3cccc(NC(=O)c4ccc(CCCN5CCOCC5)cc4)c3)cc21
CCn1c(C2=COOO2)nc2cnc(Oc3cccc(NC(=O)c4ccc(OCCN5CCOCC5)cc4)c3)cc21
CCn1c(-c2ccon2)nc2cnc(Oc3cccc(NC(=O)c4ccc(CCCN5CCOCC5)cc4)c3)cc21
CCn1c(-c2ccon2)nc2cnc(Oc3cccc(NC(=O)c4ccc(OCCN5CCOCC5)cc4)c3)cc21
CCn1c(C2=C(N)OOO2)nc2cnc(Oc3cccc(NC(=O)c4ccc(OCCN5CCOCC5)cc4)c3)cc21
CCn1c(-c2nonc2N)nc2cnc(Oc3cccc(NC(=O)c4ccc(OCCN5CCCCC5)cc4)c3)cc21
CCn1c(-c2cnon2)nc2cnc(Oc3cccc(NC(=O)c4ccc(OCCN5CCOCC5)cc4)c3)cc21
CCn1c(-c2nonc2N)nc2cnc(Oc3cccc(NC(=O)c4ccc(OCCN5CCOCC5)cc4)c3)cc21
CCn1c(-c2nocc2N)nc2cnc(Oc3cccc(NC(=O)c4ccc(OCCN5CCOCC5)cc4)c3)cc21
COc1cc2ncn(-c3cc(OCc4ccccc4C(F)(F)F)c(C(N)=O)s3)c2cc1O
COc1cc2ncn(-c3cc(OCc4ccccc4C(F)(F)F)c(C(=N)N)s3)c2cc1OC
```

```
 COc1cc2ncn(-c3cc(OCc4ccccc4C(F)(F)F)c(C(N)=O)s3)c2cc1OC
 CCc1cc2c(cc1OC)ncn2-c1cc(OCc2ccccc2C(F)(F)F)c(C(N)=O)s1
 COc1ccc2c(c1)ncn2-c1cc(OCc2ccccc2C(F)(F)F)c(C(N)=O)s1
Nc1cc(Cl)c(-c2ccc(Cl)cc2)c(N)n1
 Nc1nc(N)c(-c2ccc(Cl)cc2)c(Cl)n1
 CCc1nc(N)nc(N)c1-c1ccc(Cl)cc1
C[C+]([O-])c1ccc(-c2nc(-c3ccc(F)cc3)c(-c3ccncc3)[nH]2)cc1
 C[S+]([O-])c1ccc(-c2nc(-c3ccc(F)cc3)c(-c3ccccc3)[nH]2)cc1
 C[S+]([O-])c1ccc(-c2nc(-c3ccc(F)cc3)c(-c3ccncc3)[nH]2)cc1
C=S(=O)(c1ccc(F)cc1F)c1cc2cnc(Nc3ccc4[nH]ccc4c3)nc2n(C)c1=O
 Cn1c(=O)c(S(=O)(=O)c2ccc(F)cc2)cc2cnc(Nc3ccc4[nH]ccc4c3)nc21
 Cn1c(=O)c(S(=O)(=O)c2ccc(F)cc2F)cc2cnc(Nc3ccc4[nH]ccc4c3)nc21
 Cn1c(=O)c([SH](=O)(OO)c2ccc(F)cc2F)cc2cnc(Nc3ccc4[nH]ccc4c3)nc21
 C=C1C(S(=O)(=O)c2ccc(F)cc2)=Cc2cnc(Nc3ccc4[nH]ccc4c3)nc2N1C
Cc1cc(C#N)cc(C)c1OC1=C(Br)NNNC=NC(Nc2ccc(C#N)cc2)=N1
 Cc1cc(C#N)cc(C)c1Oc1nc(Nc2ccc(C#N)cc2)nc(N)c1Br
CC(Oc1cc(-c2cnn(C3CCNCC3)c2)cnc1N)c1c(CF)ccc(F)c1Cl
 CC(Oc1cc(-c2cnn(C3CCNCC3)c2)ccc1N)c1c(CF)ccc(F)c1Cl
 Cc1c(F)ccc(Cl)c1C(C)Oc1cc(-c2cnn(C3CCNCC3)c2)cnc1N
 CC(Oc1cc(-c2cnn(C3CCNCC3)c2)cnc1N)c1c(Cl)ccc(F)c1Cl
 Cc1c(F)ccc(Cl)c1C(N)Oc1cc(-c2cnn(C3CCNCC3)c2)cnc1N
 CC(Oc1cc(-c2cnn(C3CCNCC3)c2)ccc1N)c1c(Cl)ccc(F)c1Cl
 Cc1c(F)ccc(Cl)c1C(C)Oc1cc(-c2cnn(C3CCNCC3)c2)ccc1N
 CC(Oc1cc(-c2cnn(C3CCCCC3)c2)cnc1N)c1c(Cl)ccc(F)c1Cl
NCC(NC(=O)c1ccc(-c2ccccc2)cc1)c1ccccc1
 N=C(NC(=O)c1ccc(-c2ccncc2)cc1)c1ccccc1
 NCC(NC(=O)c1ccc(-c2ccncc2)cc1)c1ccccc1
 O=C(NC(=O)c1ccc(-c2ccncc2)cc1)c1ccccc1
 NC(CNC(=O)c1ccc(-c2ccncc2)cc1)c1ccccc1
 O=C(NC(CO)c1ccccc1)c1ccc(-c2ccncc2)cc1
 NC(=NC(=O)c1ccc(-c2ccncc2)cc1)c1ccccc1
CC(Nc1nccc(-c2c(-c3ccc(F)cc3)nc3occn23)n1)c1ccccc1
 CC(Nc1nccc(-c2c(-c3ccc(F)nc3)nc3occn23)n1)c1ccccc1
Cc1cccc(-c2cc(C(=O)O)c(-c3ccc(Cl)cc3Cl)[nH]2)n1
 Cc1nccc(-c2cc(C(N)=O)c(-c3ccc(Cl)cc3Cl)[nH]2)n1
```

```
 NC(=O)c1cc(-c2ccnc(N)n2)[nH]c1-c1ccc(Cl)cc1Cl
 Nc1nccc(-c2cc(C(=O)O)c(-c3ccc(Cl)cc3Cl)[nH]2)n1
 NC(=O)c1cc(-c2cccc(N)n2)[nH]c1-c1ccc(Cl)cc1Cl
COc1cccc(C(C)NC(=O)c2ccc(-c3ccncc3C)cc2)c1
 COc1cccc(C(C)NC(=O)c2ccc(-c3ccncc3)cc2)c1
 COc1cccc(C(C)NC(=O)c2ccc(-c3ccncc3F)cc2)c1
 COc1cccc(C(C)NC(=O)c2ccc(-c3ccncc3O)cc2)c1
N#Cc1ncc2nc1CCCCOOc1cc(NCc3cncs3)c(Cl)cc1NC(=O)N2
 N#Cc1ncc2nc1OCCCCOc1cc(NNc3cncs3)c(Cl)cc1NC(=O)N2
 N#Cc1ncc2nc1CCCCCOc1cc(NCc3cncs3)c(Cl)cc1NC(=O)N2
 CC1=NC=C1CNc1cc2c(cc1Cl)NC(=O)Nc1cnc(C#N)c(n1)OCCCCO2
 N#Cc1ncc2nc1OCCCCOc1cc(NCc3cncs3)c(Cl)cc1NC(=O)N2
 CC1=CC=C2NC(=O)Nc3cnc(C#N)c(n3)OCCCCC=C2C=C1NCc1cccs1
 N#Cc1ncc2nc1OCCCCOc1cc(NCc3cccs3)c(Cl)cc1NC(=O)N2
CCOc1nc(CC(C)NCc2ccc([SH](N)(=O)OO)cc2)cc(N)c1CC
 CCOc1nc(NC(=O)Cc2cc(OC)c(S(C)(=O)=O)cc2OC)cc(N)c1Cl
 CCOc1nc(C(=O)NCc2ccc([SH](C)(=O)OO)cc2)cc(N)c1Cl
 CCOc1nc(C(=O)NCc2ccc(S(=O)(=O)CC)cc2)cc(N)c1Cl
 CCOc1nc(NC(=O)Cc2cc(OC)c(S(C)(=O)=O)cc2OC)cc(N)c1C#N
 CCOc1nc(C(=O)NCc2ccc(S(C)(=O)=O)cc2)cc(N)c1Cl
 CCOc1cc(N)cc(NC(=O)Cc2cc(OC)c(S(C)(=O)=O)cc2OC)n1
 CCOc1nc(C(=O)NCc2ccc(S(=O)(=O)CC)cc2)cc(N)c1CC
 CCOc1nc(NC(=O)Cc2cc(OC)c(S(C)(=O)=O)cc2OC)cc(N)c1C
CCOc1nc(NC(=O)Cc2cc(OC)c(S(C)(=O)=O)cc2OC)cc(N)c1Cl
 CCOc1nc(NC(=O)Cc2cc(OC)c(S(C)(=O)=O)cc2OC)cc(N)c1CC
 CCOc1nc(NC(=O)Cc2cc(OO)c([N+](C)([O-])OO)cc2OC)cc(N)c1C#N
CCOc1nc(NC(=O)Cc2cc(OC)c([SH](C)(=O)OO)cc2OC)cc(N)c1C
 CCOc1nc(NC(=O)Cc2cc(OC)c(S(C)(=O)=O)cc2OC)cc(N)c1CN
 CCOc1cc(N)cc(NC(=O)Cc2cc(OO)c(S(C)(=O)=O)cc2OC)n1
COc1cc(S(C)(=O)=O)c(OC)cc1CC(=O)Nc1cc(N)c(C#N)c(OC(C)C)n1
 COc1cc([SH](=O)=O)c(OC)cc1CC(=O)Nc1cc(N)c(C#N)c(OC(C)C)n1
 CCOC1=C(C#N)C(N)=CC(NC(=O)Cc2cc(OC)c(S(C)(=O)=O)cc2OC)=NC1
Clc1ccc(Nc2nnc(Cc3ccncc3)c3ccccc23)cc1
 Clc1ccc(Nc2nnc(Nc3ccncc3)c3ccccc23)cc1
O=C1NCCc2[nH]c(-c3ccccn3)cc21
```

```
 O=C1NCCc2[nH]c(-c3ccncc3)nc21
 O=C1NCCc2[nH]c(-c3ccncn3)cc21
 OCC1NCCc2[nH]c(-c3ccccc3)cc21
 O=C1NCCc2[nH]c(-c3ccncc3)cc21
 OCC1NCCc2[nH]c(-c3ccncc3)cc21
 O=C1NCCc2[nH]c(-c3ccccc3)cc21
CC(C)(N)c1ccc2nc(-c3n[nH]c4ccccc34)[nH]c2c1
 CC(C)(C)c1ccc2nc(-c3n[nH]c4ccccc34)[nH]c2c1
O=c1c(O)c(-c2ccc(O)c(O)c2)oc2cc(O)cc(O)c12
 O=c1c(O)c(-c2cccc(OOOO)c2)oc2cc(O)cc(O)c12
 O=C1CC=C(O)C=C(O)C=c2oc(c3c2=c2occ(O)c2=3)=C1O
 O=C1C(O)=CCOC2=C1COC(c1ccc(O)c(O)c1)=CC(O)=C2
 Cc1c(-c2ccc(O)c(O)c2)oc2cc(O)cc(O)c2c1=O
CS(=O)(=O)Nc1cc2c(cc1Cl)NC(=O)Nc1cnc(C#N)c(n1)OCCCCCO2
 CCc1cc2c(cc1NS(C)(=O)=O)OCCCCCOc1nc(nnc1C#N)NC(=O)N2
 CCC1=CC=CNC(OO)NNC=CC2=C(C#N)C=CC(=CC=C1NS(C)(=O)=O)CCCCCO2
 CCc1cc2c(cc1NS(C)(=O)=O)OCCCCCOc1nc(ccc1C#N)NC(CO)N2
 CCc1cc2c(cc1NS(C)(=O)=O)OCCCCCOc1nc(ccc1C#N)NC(=O)N2
 CS(=O)(=O)NC1=CC=COCCCCCOc2nc(cnc2C#N)NC(=O)NC=CC=C1Cl
 CS(=O)(=O)Nc1cc2c(cc1Cl)NC(=O)Nc1cnc(N)c(n1)OCCCCCO2
 CCc1cc2c(cc1NS(C)(=O)=O)OCCCCCOc1nc(cnc1C#N)NC(=O)N2
Cc1ccc(F)c(NC(=O)Nc2ccc(-c3cccc4[nH]nc(N)c34)cc2)c1
 Cc1ccc(F)c(NC(=O)Nc2ccc(-c3cncc4[nH]nc(N)c34)cc2)c1
 Cc1cc(C)cc(NC(=O)Nc2ccc(-c3cccc4c3=C(N)N=CC=NC=4)cc2)c1
 Cc1ccc(F)c(NC(=O)Nc2ccc(-c3cccc4c3=C(N)N=CC=NC=4)cc2)c1
CN1CCN(C2=NC(C3=C(c4c[nH]c5ccccc45)C(=O)NC3=O)=C3C=CC=CC23O)CC1
 CN1CCN(c2nnc3cccc-3c(C3=C(c4c[nH]c5ccccc45)C(=O)NC3=O)n2)CC1
 CN1CCN(c2nc(C3=C(c4c[nH]c5ccccc45)C(=O)NC3=O)cc3cccc-3n2)CC1
 CN1CCN(c2nnc3cccc-3c(C3=C(c4c[nH]c5ccccc45)C(=O)CC3=O)n2)CC1
 CN1CCN(c2nc(C3=C(c4c[nH]c5ccccc45)C(=O)NC3=O)c3ccccc3n2)CC1
CCc1cc2c(cc1CCCO)OCCCCCOc1nc(cnc1CNN)NC(=O)N2
 CC1=C2N=C(N)NC(=O)Nc3cc(c(CCCO)cc3OCCCCCO2)CC=CC=C1
 CCCCc1cc2c(cc1Cl)NC(=O)Nc1cnc(C#N)c(n1)OCCCCCO2
 N#Cc1ccc2nc1OCCCCCOc1cc(CCCO)c(Cl)cc1NC(=O)C2
 N#Cc1ccc2nc1OCCCCCOc1cc(CCCO)c(Cl)cc1NC(=O)N2
```

N#Cc1ncc2nc1OCCCCOc1cc(CCCO)c(Cl)cc1NC(=O)N2
N#Cc1ncc2nc1OCCCCOc1cc(CCCO)c(Cl)cc1NC(=O)C2
N#Cc1ncc2nc1CCCCCOc1cc(CCCO)c(Cl)cc1NC(=O)N2
CC1=NC2=C(CN)C=CC=CCc3cc(c(cc3CCCO)OCCOCCO2)NC(=O)N1
CCc1cc2c(cc1CCCO)OCCCCOc1nc(cnc1C#N)NC(=O)N2
COc1cccc2cc(NC(=O)Nc3cccc(C(F)(F)F)n3)c-2cc1
COc1ccc2c(NC(=O)Nc3cccc(C(O)(F)F)n3)cccc2c1
COc1ccc2c(NC(=O)Nc3cccc(C(F)(F)F)n3)cccc2c1
COc1ccc2c(NC(=O)Nc3cccc(C(O)(F)F)n3)ccnc2c1
COc1ccc2c(NC(=O)Nc3cccc(C(F)(F)F)n3)ccnc2c1
COc1cccc2cc(NC(=O)Nc3cccc(C(O)(F)F)n3)c-2cc1
Cc1ncc(-c2cnn(CCO)c2)c2scc(-c3ccc(NC(=O)Nc4cccc(F)c4)cc3)c12
NC1=CC(c2ccc(NC(=O)Nc3cccc(F)c3)cc2)=CSSC=C(c2cnn(CCO)c2)C=N1
Nc1ncc(-c2cnn(CCO)c2)c2scc(-c3ccc(NC(=O)Nc4cccc(F)c4)cc3)c12
COC1=C(OCCCN2CCCC2)C=C2N=CC23Oc2ccc4c(c2F)C(C)(N4)C3=C1
COc1cc2c(Oc3ccc4[nH]c(C)cc4c3F)nccc2cc1OCCCC1CCCC1
COc1cc2c(Oc3ccc4[nH]c(C)cc4c3F)ncnc2cc1OCCCN1CCCC1
CCCCCNCCCOc1cc2ncnc(Oc3ccc4[nH]c(C)cc4c3F)c2cc1OC
COc1cc2c(Oc3ccc4[nH]c(C)cc4c3F)ncnc2cc1OCCCN1CCC1
COc1cc2c(Oc3ccc4[nH]c(C)cc4c3F)ncnc2cc1OCCCC1CCCC1
COC1=CC2=C(OC=NC=CC=C1OCCCN1CCCC1)Oc1ccc3c(c1F)C2(C)N3
CCCCCCNCCCOc1cc2ccnc(Oc3ccc4[nH]c(C)cc4c3F)c2cc1OC
O=C1Cc2c([nH]c3ccc([N+](=O)[O-])cc23)-c2ccccc2N1
O=C1CC2=Cc3cc([N+](=O)[O-])ccc3NC2=CC=CC=CC=CN1
O=C1CC2=C3C(=CC=C3[N+](=O)[O-])C=C2c2ccccc2N1
O=C1CC2=C3C(=CC=C3[N+](=O)[O-])C=C2c2cccc(c2)N1
O=C(Cc1ccccc1)Nc1cccc(-c2nc3sccn3c2-c2ccnc(Nc3cccc(N4CCCCC4)c3)n2)c1
O=C(Cc1ccccc1)Nc1cccc(-c2nc3sccn3c2-c2ccnc(Nc3cccc(N4CCOCC4)c3)n2)c1
O=C(Cc1ccccc1)Nc1cccc(-c2cc3sccn3c2-c2ccnc(Nc3cccc(N4CCCCC4)c3)n2)c1
O=C(Cc1ccccc1)Nc1cccc(-c2cc3sccn3c2-c2ccnc(Nc3cccc(N4CCOCC4)c3)n2)c1
O=C(Cc1ccccc1)Nc1cccc(-c2nc3sccn3c2-c2ccnc(Nc3cccc(N4CCOOC4)c3)n2)c1
CC1CCN(C2CCNC2)CC1N(C)c1ncnc2[nH]ccc12
O=C1NC(=O)C(c2cnc3ccccn23)CC1c1cn2c3c(cccc13)CN(C(CO)N1CCCCC1)CC2
COc1cc(Nc2ncc(F)c(Nc3ccc4c(n3)NC(=O)C(C)(C)O4)n2)cc(OC)c1OO
COc1cc(Nc2ncc(F)c(Nc3ccc4c(n3)NC(=O)C(C)(C)O4)n2)cc(OC)c1OC

```
 COc1cc(Nc2ncc(F)c(Nc3ccc4c(n3)NC(=O)C(C)(C)O4)n2)cc(OO)c1OO
 COc1cc(Nc2ncc(F)c(Nc3ccc4c(n3)NC(=O)C(C)(C)O4)n2)cc(OO)c1O
 COc1cc(Nc2ncc(F)c(Nc3ccc4c(n3)NC(=O)C(C)(C)O4)n2)cc(OO)c1OC
 CCc1c(OC)cc(Nc2ncc(F)c(Nc3ccc4c(n3)NC(=O)C(C)(C)O4)n2)cc1OC
 COc1cc(Nc2ncc(O)c(Nc3ccc4c(n3)NC(=O)C(C)(C)O4)n2)cc(OC)c1OC
 COc1cc(Nc2ncc(F)c(Nc3ccc4c(n3)NC(=O)C(C)(C)O4)n2)cc(OC)c1O
 COc1cc(Nc2ncc(O)c(Nc3ccc4c(n3)NC(=O)C(C)(C)O4)n2)cc(OC)c1O
 CCOc1nc(C(=O)CCc2ccccc2S(N)(=O)=O)cc(N)c1C#N
BBc1cc(F)c(COc2nsc(NC(=O)NCCCCN3CCCC3)c2C(C)=O)c(F)c1
 NC(=O)c1c(OCc2ccc(Br)cc2F)nsc1NC(=O)NCCCCN1CCCC1
 CC(=O)c1c(OCc2c(F)cc(Br)cc2F)nsc1NC(=O)NCCCCN1CCCC1
 NC(=O)c1c(OCc2c(F)cc(Br)cc2F)nsc1NC(=O)NCCCCN1CCCC1
CC1C(=O)C(Oc2ccc(F)cc2)=Cc2cnc(NC3CCOCC3)nc21
 CC1C(=O)C(Oc2ccc(F)cc2)NCc2cnc(NC3CCOCC3)nc21
 Cn1c(=O)c(Oc2ccc(F)cc2)cc2cnc(NC3CCOCC3)nc21
 Cc1ccc(OC2NCc3cnc(NC4CCOCC4)nc3C(C)C2=O)cc1
 Cc1ccc(Oc2cc3cnc(NC4CCOCC4)nc3n(C)c2=O)c(F)c1
 Cn1c(=O)c(Oc2ccc(F)cc2F)cc2cnc(NC3CCOCC3)nc21
 CC1C(=O)C(Oc2ccc(F)cc2F)=Cc2cnc(NC3CCOCC3)nc21
NC(Cc1c[nH]c2ccccc12)C(=O)Nc1cncc(C=Cc2ccccc2)c1
 NC(Cc1c[nH]c2ccccc12)C(=O)Nc1cccc(C=Cc2ccccc2)c1
 NC(Cc1c[nH]c2ccccc12)C(=O)Nc1cncc(C=Cc2ccncc2)c1
 NC(Cc1c[nH]c2ccccc12)C(=O)Nc1cccc(C=Cc2ccncc2)c1
CC(N)C1CCC(C(=N)Nc2ccccc2)CC1
 CC(N)C1CCC(C(=O)Nc2ccccc2)CC1
 CC(N)C1CCC(C(=O)Nc2ccncc2O)CC1
 CC(N)C1CCC(C(=O)Nc2ccncc2)CC1
 CC(N)C1CCC2CCC=CC=CC=C(N1)C2=N
 CC(N)C1CCC(C(=O)Nc2ccccc2O)CC1
 CC(C)C1CCC(C(=O)Nc2ccncc2)CC1
CCCC(=O)Nc1cccc(-c2nc(Nc3ccc4[nH]ncc4c3)c3cc(CCCN4CCCC4)ccc3n2)c1
COc1ccc(NC(=O)Nc2ccc(-c3csc4c(-c5cnn(CC(C)(C)O)c5)cnc(N)c34)cc2)cc1
 COc1ccc(NC(=O)Nc2ccc(-c3csc4c(C5=CCC(CC(C)(C)O)=C5)cnc(N)c34)cc2)cc1
COc1cccc(-c2c(-c3ccc(F)cc3)ncn2C2CCNCC2)n1
 COc1nccc(-c2c(-c3ccc(O)cc3)ncn2C2CCCCC2)n1
```

```
 COc1nccc(-c2c(-c3ccc(F)cc3)ncn2C2CCCCC2)n1
 COc1nccc(-c2c(-c3ccc(O)cc3)ncn2C2CCNCC2)n1
 COc1nccc(C2=C3N=CCCCCCNCCC=CC(F)=CC=C32)n1
 COc1nccc(-c2c(-c3ccc(F)cc3)ncn2C2CCNCC2)n1
CC(N)C1CCC(C(=O)Nc2ccnc3[nH]ccc23)CC1
CCOc1nc(C(=O)NCc2ccccc2S(N)(=O)=O)cc(N)c1CNN
 CCOc1nc(C(=O)CC2C=CC=CC=C2S(=O)(=O)NN)cc(N)c1CCN
 CCOc1nc(C(=O)CCc2ccccc2S(C)(=O)=O)cc(N)c1C#N
 CCOc1nc(C(=O)NCc2ccccc2[SH](N)(=O)CO)cc(N)c1C#N
 CCOc1nc(C(=O)NCc2ccccc2S(N)(=O)=O)cc(N)c1C#N
CCc1nc(-c2cccc(C)c2)c(-c2ccnc(NC(=O)c3ccccc3)c2)s1
 COc1nc(-c2cccc(C)c2)c(-c2ccnc(NC(=O)c3ccccc3)c2)s1
NC(COCc1ccccc1)COc1cncc(C=Cc2ccncc2)c1

 CCC(C)N1C=CN=CN=C(N)C=CC(c2ccc(C)cc2)=N1
 Cc1ccc(-c2nn(C(C)C)c3ncnc(N)c23)cc1
 Cc1ccc(C2=NN(C(C)C)C=CN=CN=C(N)C=C2)cc1
 Cc1ccc(-c2nn(C(C)C)c3nnnc(N)c23)cc1
 Cc1ccc(-c2nn(C(C)(C)C)c3ncnc(N)c23)cc1
 Cc1ccc(-c2nn(C(C)(C)O)c3ncnc(N)c23)cc1
 CCC(C)(C)n1nc(-c2ccc(C)cc2)c2c(N)ncnc21
CC(C)(C)NS(=O)(=O)c1cncc(-c2ccn3nc(N)nc3c2)c1
 CC(C)(C)NS(=O)(=O)c1cncc(-c2ccn3nc(N)cc3c2)c1
Nc1ccc(-c2cncc(OCC(N)Cc3c[nH]c4ccccc34)c2)cc1C(=O)c1ccccc1
OCCCn1cnc(-c2ccc(F)cc2)c1-c1ccncc1
Cc1cc(OCC2CCC(C(=O)O)C2)c(NC(=O)Nc2cnc(C#N)cn2)cc1Cl
COc1cc(C)c(Cl)cc1NC(=O)Nc1cnc(C#N)cn1
 COc1cc(C)c(Cl)cc1NC(=O)Nc1ccc(C#N)cc1
 COc1cc(C)c(Cl)cc1NC(=O)Nc1cnc(C#N)nn1
 CSc1cc(C)c(Cl)cc1NC(=O)Nc1cnc(C#N)cn1
COc1cccc(C(C)NC(=O)c2ccc(-c3ccncc3)c(F)c2)c1
Cc1cccc(NC(=O)Nc2ccc(-c3coc4c3=C(N)NNNC=CN=4)cc2)c1
 Cc1cccc(NC(=O)Nc2ccc(-c3ccc4ncnc[nH][nH]c(N)c3-4)cc2)c1
 Cc1cccc(NC(=O)Nc2ccc(-c3ccc4ncc[nH][nH]nc(N)c3-4)cc2)c1
 Cc1cccc(NC(=O)Nc2ccc(-c3coc4ncnc(N)c34)cc2)c1
```

Cc1cccc(NC(=O)Nc2ccc(-c3csc4ncnc[nH]nc(N)c34)cc2)c1
Cc1cccc(NC(=O)Nc2ccc(-c3ccc4ncn[nH][nH]nc(N)c3-4)cc2)c1
Cc1cccc(NC(=O)Nc2ccc(-c3csc4ncnc(N)c34)cc2)c1
Cc1cccc(NC(=O)Nc2ccc(-c3csc4nc(N)nc(N)c34)cc2)c1
Cc1cccc(NC(=O)Nc2ccc(-c3csc4ncc[nH]nnc(N)c34)cc2)c1
Cc1cccc(NC(=O)Nc2ccc(-c3coc4ncnc(N)c34)cc2)c1
Cc1cccc(NC(=O)Nc2ccc(-c3csc4ccnc(N)c34)cc2)c1
N#Cc1ccc(NC(=O)Nc2ccnc3cc(C(F)(F)F)ccc23)nc1
C=C(Nc1ccc(-c2cccc3[nH]nc(N)c23)cc1)Nc1cccc(C)c1
Cc1cccc(NC(=O)Nc2ccc(-c3cccc4[nH]nc(N)c34)cc2)c1
CCC(CC)CCNC(=O)C1=C(C)NC2C=C(CC=O)C3(C=C1)C=C(F)C=CN3N2
CCN(CC)CCNC(=O)c1c(C)[nH]c(C=C2C(=O)Nc3ccc(F)cc32)c1C
CCC(CC)CCNC(=O)c1c(C)[nH]c(C=C2C(=O)Nc3ccc(F)cc32)c1C
C=C(CC)CCNC(=O)c1c(C)[nH]c(C=C2C(=O)Nc3ccc(F)cc32)c1C
CCN(CC)CCNC(=O)c1cc(C=c2c(=O)[nH]c3ccc(F)c2c3)[nH]c1C
CCN(CC)CCNC(=O)C1=C(C)[CH]C(C=C2C(=O)Nc3ccc(F)cc32)=C1C
CCN(CC)CCNC(=O)c1cc(C=C2C(=O)Nc3ccc(F)cc32)[nH]c1C
CCN(CC)CCNC(=O)c1c(C)[nH]c(C=C2C(=O)Nc3ccc(C)cc32)c1C
CCN(CCNC(=O)c1c(C)[nH]c(C=C2C(=O)Nc3ccc(F)cc32)c1C)NC
CCN(CC)CCNC(=O)C1=C(C)[CH]C(C=C2C(=O)Nc3ccc(F)cc32)=C1
CCCC(=O)Nc1n[nH]c2ccc(-c3cncc(F)c3)cc12
CCCC(=O)Nc1n[nH]c2ccc(-c3cccc(F)c3F)cc12
CCCC(=O)Nc1n[nH]c2ccc(-c3cncc(F)c3F)cc12
CCCC(=O)Nc1n[nH]c2ccc(-c3cccc(F)c3)cc12
CCCC(=O)Nc1n[nH]c2ccc(-c3cccc(F)c3C)cc12
CN1CC=CC2=C1N(Nc1ccccc1)N=NC2
Nc1ccc2c(c1)NC(=O)c1ccccc1N2
NC1=CC=C2NC3=CC=CC=CCC(=O)NC2=CN3C=CC=CC=C1
O=C1Nc2cc(O)ccc2Nc2ccccc21
OCC(Nc1ccccn1)Nc1cccc2c(C(F)(F)F)cccc12
N=C(Nc1cnccn1)Nc1ccnc2c(C(F)(F)F)cccc12
O=C(Nc1cncnn1)Nc1ccnc2c(C(F)(F)F)cccc12
O=C(Nc1ccccn1)Nc1cccc2c(C(F)(F)F)cccc12
O=C(Nc1ccccn1)Nc1ccnc2c(C(F)(F)F)cccc12
N=C(Nc1cnccn1)Nc1cccc2c(C(O)(F)F)cccc12

```
 O=C(Nc1cnccn1)Nc1ccnc2c(C(F)(F)F)cccc12
 N=C(Nc1ccccn1)Nc1cccc2c(C(O)(F)F)cccc12
 O=C(Nc1cnccn1)Nc1cccc2c(C(F)(F)F)cccc12
 CC(F)(F)c1cccc2c(NC(CO)Nc3ccccn3)cccc12
CC1C(c2ccc(O)cc2)=COc2cc(O)cc(O)c21
 Cc1ccc(-c2coc3cccc-3cc(O)coc2=O)cc1
 O=c1c(-c2ccc(O)cc2)coc2cc(O)cc(O)c12
 OCC1C(c2ccc(O)cc2)=COc2cc(O)cc(O)c21
 Oc1ccc(C2=COc3cc(O)cc(O)c3C2)cc1
 OCC1C(c2ccc(F)cc2)=COc2cc(O)cc(O)c21
O=C(Cc1ccccc1)Nc1cccc(-c2nc3sccn3c2-c2ccnc(Nc3ccc(N4CCOOC4)cc3)n2)c1
 O=C(Cc1ccccc1)Nc1cccc(-c2nc3sccn3c2-c2cccc(Nc3ccc(N4CCOCC4)cc3)n2)c1
 O=C(Cc1ccccc1)Nc1cccc(-c2nc3sccn3c2-c2ccnc(Nc3ccc(N4CCOCC4)cc3)n2)c1
 O=C(Cc1ccccc1)Nc1cccc(-c2nc3sccn3c2-c2ccnc(Nc3ccc(N4CCCCC4)cc3)n2)c1
Cc1ccc(-c2nc3cnccn3c2-c2ccnc(NCC3CC3)n2)cc1
COc1ccc(CNc2c(Nc3ccc4c(c3)C=N[CH]4)c(=O)c2=O)cc1
 COc1ccc(CNc2c(Nc3ccc4[nH]ncc4c3)c(=O)c2=O)cc1
CC1=CC=NC2=CCN=NC(=C3C=CC2=C3)C=CC=CC=CC=C1
Cc1cccc(NC(=O)Nc2ccc(-c3csc4c3=C(N)NNNN=CN=4)cc2)c1
 Cc1cccc(NC(=O)Nc2ccc(-c3noc4nccc(N)c34)cc2)c1
 Cc1cccc(NC(=O)Nc2ccc(-c3noc4ncnc(N)c34)cc2)c1
O=c1[nH]c2cc(-c3ccccc3)cnc2[nH]1
O=CCn1cnc(-c2ccc(F)cc2)c1-c1ccncc1
 OCCCn1cnc(-c2ccc(F)cc2)c1-c1ccccc1
 OCC1N=C(c2ccncc2)C(c2ccc(F)cc2)=N1
 OCCCn1cnc(-c2ccc(F)cc2)c1-c1ccncc1
Oc1cccc2ncnc(Nc3ccccc3)c12
 CN1CC=Cc2ccnnc[nH]c3cccc-3c21
Cc1ccc(NC(=O)Nc2ccc(-c3cccc4sncc34)cc2)cc1C(F)F
 O=C(Nc1ccc(-c2cccc3snnc23)cc1)Nc1ccc(F)c(C(F)(F)F)c1
 O=C(Nc1ccc(-c2cccc3c2C=NC3)cc1)Nc1ccc(F)c(C(F)(F)F)c1
 Cc1ccc(NC(=O)Nc2ccc(-c3cccc4c3C=NC4)cc2)cc1C(F)(F)F
 O=C(Nc1ccc(-c2cccc3sncc23)cc1)Nc1ccc(F)c(C(F)(F)F)c1
 Cc1ccc(NC(=O)Nc2ccc(-c3cccc4sncc34)cc2)cc1C(F)(F)F
O=c1[nH]cnc2c1oc1ccc(Cl)cc12
```

```
 O=C1NCNCc2c1oc1ccc(Cl)cc21
 O=C1NCCNc2c1oc1ccc(Cl)cc21
 O=C1NCNNc2c1oc1ccc(Cl)cc21
 O=C1NCC=Cc2c1oc1ccc(Cl)cc21
CNC(=O)c1ccccc1Sc1ccc2c(C=Cc3ccccn3)n[nH]c2c1
 CNC(=O)c1ccccc1Sc1ccc2c(CCCc3ccccn3)n[nH]c2c1
 CNC(=O)c1ccccc1[SH]1C=CC=c2c(C=Cc3ccccn3)c[nH]c2=C1
 CNC(=O)c1ccccc1Sc1ccc2c(C=Cc3ccccn3)c[nH]c2c1
 CNC(=O)c1ccccc1[SH]1C=CC=c2c(C=Cc3ccccn3)n[nH]c2=C1
 CNC(=O)c1ccccc1Sc1ccc2c(C=Cc3ccccc3)n[nH]c2c1
Nc1ccc2c(c1)NC(=O)c1ccccc1N2
 Nc1cc2c(cn1)Nc1ccccc1C(=O)N2
CC1=NC(c2ccc(F)cc2)=C(c2ccncn2)C1
Fc1ccc(-c2nc(-c3ccc(F)cc3)c(-c3ccncc3)[nH]2)cc1
 CC1C=CC(c2nc(-c3ccc(F)cc3)c(-c3ccncc3)[nH]2)=CC1
 Fc1ccc(-c2nc(-c3ccc(F)cc3)c(-c3ccccc3)[nH]2)cc1
 Cc1ccc(C2=NC3=CC=C(F)C=CC3=C(c3ccncc3)N2)cc1
 Cc1ccc(-c2nc(-c3ccc(F)cc3)c(-c3ccccc3)[nH]2)cc1
 Fc1ccc(-c2nc(C3=CCC(F)C=C3)[nH]c2-c2ccncc2)cc1
 Cc1ccc(-c2nc(-c3ccc(F)cc3)c(-c3ccncc3)[nH]2)cc1
COc1ccc(-c2n[nH]cc2-c2ccncn2)cc1Cl
COc1cccc(C=C2NC(=Nc3ccccc3)NC2=O)c1
 COc1cccc(C=C2SC(=Nc3ccccc3)NC2=O)c1
 COc1cccc(C=C2SC(=Nc3ccccc3)NC2CO)c1
OC(=Nc1cccc(Cl)c1)Nc1ncc(CCN=c2[nH]cnc3ccsc23)s1
 OC(=NC1=CC(CCl)C=C1)Nc1ccc(CCNCC2C=CC=CC=CN=CN2)cn1
 OC(=Nc1cccc(Cl)c1)Nc1ccc(CCN=C2C=CC=CC3=C(C2)N3)cn1
 OC(=Nc1cccc(Cl)c1)Nc1ncc(CCN=C2NC=NC3=S2SC=C3)s1
 OC(=Nc1cccc(Cl)c1)Nc1ncc(CCN=c2[nH]nnc3ccsc23)s1
 OC(=Nc1cccc(Cl)c1)Nc1ncc(CCN=C2NN=NC3=S2SC=C3)s1
Cc1ccc(NC(=O)Nc2ccc(-c3cccc4c3CNC4=O)cc2)cc1C
 Cc1ccc(NC(=O)Nc2cccc(-c3cccc4c3CNC4=O)c2)cc1
 Cc1ccc(NC(=O)Nc2ccc(-c3cccc4c3CNC4OO)cc2)cc1C
 Cc1ccc(NC(=O)Nc2ccc(-c3cccc4c3CNC4CO)cc2)cc1C
 Cc1ccc(NC(=O)Nc2ccc(-c3cccc4c3CNC4=O)cc2)cc1
```

```
 Cc1ccc(NC(=O)Nc2ccc(-c3cccc4c(=O)[nH][nH]c34)cc2)cc1
CCC(C(=O)N1Cc2[nH]nc(NC(=O)c3ccc(N4CCN(C)CC4)cc3)c2C1)c1ccccc1
 COC(C(=O)N1Cc2[nH]nc(NC(=O)c3ccc(N4CCN(C)CC4)cc3)c2C1)c1ccccc1
 CCC(C(=O)N1Cc2[nH]nc(NC(=O)c3ccc(N4CCC(C)CC4)cc3)c2C1)c1ccccc1
 COC(C(=O)N1Cc2[nH]nc(NC(=O)c3ccc(N4CCN(C)NC4)cc3)c2C1)c1ccccc1
 COC(C(=O)N1Cc2[nH]nc(NC(=O)c3ccc(N4CCC(C)CC4)cc3)c2C1)c1ccccc1
Cc1ccc(F)c(NC(=O)Nc2ccc(-c3nsc4nccc(N)c34)cc2)c1
 Cc1ccc(F)c(NC(=O)Nc2ccc(-c3nccn4cnc(N)c34)cc2)c1
 Cc1ccc(F)c(NC(=O)Nc2ccc(-c3csc4ncnc(N)c34)cc2)c1
 Cc1ccc(F)c(NC(=O)Nc2ccc(-c3nscc4cnc(N)c3-4)cc2)c1
 Cc1ccc(F)c(NC(=O)Nc2ccc(-c3nsc4ncnc(N)c34)cc2)c1
 Cc1ccc(F)c(NC(=O)Nc2ccc(-c3n[nH]c4ncnc(N)c34)cc2)c1
 Cc1ccc(F)c(NC(=O)Nc2ccc(-c3nccc4cncc[nH][nH]c34)cc2)c1
CC(C)Nc1c(-c2ccc3[nH]ncc3c2)nc2ncccn12
 CC(C)Nc1c(-c2ccc3[nH]ncc3c2)nc2ccccn12
 CCC.Nc1c(-c2ccc3[nH]ncc3c2)nc2ncccn12
CC(C)Cn1c(N)nc2c(F)cc(-c3c(-c4ccc(F)cc4)nc4sccn34)cc21
COc1ccc(CNC(=O)Nc2ccc3[nH]ncc3c2)cc1
 O=C(NCc1ccc(Cl)cc1)Nc1ccc2[nH]ncc2c1
 O=C(NCc1ccc(Cl)c(Cl)c1)Nc1ccc2[nH]ncc2c1
 Cc1cc(CNC(=O)Nc2ccc3[nH]ncc3c2)ccc1Cl
 CCc1ccc(CNC(=O)Nc2ccc3[nH]ncc3c2)cc1
CCS(=O)(=O)Nc1ccc(-c2ccc3[nH]nc(N)c3c2C)cc1
 CCS(=O)(=O)Nc1ccc(-c2ccc3[nH]nc(C)c3c2)cc1
 CCS(=O)(=O)Nc1ccc(-c2ccc3[nH]nc(N)c3c2)cc1
Cc1cccc(NC(=O)Nc2ccc(-c3csc4cnnc(N)c34)cc2)c1
 Cc1cccc(NC(=O)Nc2ccc(-c3csc4nnnc(N)c34)cc2)c1
 Cc1cccc[nH]c(=O)[nH]c2ccc(cc2)c2ccc3cccc(c1)-n-c=c-3-2
Cc1ccc(-c2nc3cnccn3c2C2=CC=NC3=NCCC2CN3)cc1
CCCC(=O)Nc1n[nH]c2ccc(-c3cccc(F)c3F)cc12
CN(C)c1cc2sncc2cc1NC(=O)C(=O)O
COc1cc(C2=CC=CC(C(=O)Nc3ccccc3)=NNC=CC=C2)ccc1O
 COc1cc(-c2ccc3[nH]nc(C(=O)Nc4ccccc4)c3c2)ccc1C
 COc1cc(-c2ccc3[nH]nc(C(=O)Nc4ccccc4)c3c2)ccc1O
Cc1ccc(-c2ccc(NC(=N)Nc3cccc(Br)c3)cc2)c2c(N)n[nH]c12
```

```
 Cc1ccc(-c2ccc(NC(=O)Nc3cccc(Br)c3)cc2)c2c(N)c[nH]c12
 Cc1ccc(-c2ccc(NC(=N)Nc3cccc(Br)c3)cc2)c2c(N)c[nH]c12
 Cc1ccc(-c2ccc(NC(=O)Nc3cccc(Br)c3)cc2)c2c(N)n[nH]c12
NC(=O)c1cccc2[nH]c(-c3ccncc3)nc12
 NC(CO)c1cccc2[nH]c(-c3ccncc3)nc12
 NC(=O)c1cccc2[nH]c(-c3ccncc3)cc12
Clc1ccc2[nH]nc(-c3ccccc3)c2c1
 Brc1ccc2[nH]nc(-c3ccccc3)c2c1
 Brc1cc2c(-c3ccccc3)n[nH]c2cn1
CCc1c(-c2ccc(C(C)(C)O)cc2)[nH]c2nccnc12
COc1cccc(C=C2SC(=Nc3ccccc3)NC2OO)c1
 C=c1[nH]c(=Nc2ccccc2)sc1=Cc1cccc(OC)c1
 COc1cccc(C=C2CC(=Nc3ccccc3)NC2=O)c1
 C=C1NC2=CC(C=C2)N=C2C=C(C)C(OC)=CC=CC=C1S2
 C=C1CC(=Nc2ccccc2)SC1=Cc1cccc(OC)c1
 COc1cc(C=C2SC(=Nc3ccccc3)NC2=O)ccc1O
 COC1=CC(C=C2SC(=Nc3ccccc3)NC2CO)=CC=CC=C1
 C=c1[nH]c(=Nc2ccccc2)sc1=CC1=C(O)C(OC)=C1
 COc1cc(C=C2CC(=Nc3ccccc3)NC2=O)ccc1O
Cc1cccc(NC(=O)Nc2ccc(-c3csc4ncnc(N)c34)cc2)c1
CS(=O)(=O)Nc1ccc(-c2ccc3[nH]cc(N)c3c2)cc1
 CS(=O)(=O)Nc1ccc(C2=CC=CC(N)=NNC=CC=C2)cn1
 CS(=O)(=O)Nc1ccc(-c2ccc3[nH]nc(N)c3c2)cc1
 CS(=O)(=O)Nc1ccc(-c2ccc3[nH]nc(N)c3c2O)cc1
 CS(=O)(=O)Nc1ccc(-c2ccc3[nH]nc(F)c3c2)cc1
CCCS(=O)(=O)Nc1ccc(-c2ccc3[nH]nc(NC(=O)CC)c3c2)cc1
 CCC[SH](=O)(Nc1ccc(-c2ccc3[nH]nc(NC(=O)CC)c3c2)cc1)OO
CSc1ccc(-c2nc(-c3ccc(F)cc3)c(-c3ccccc3)[nH]2)cc1
 COc1ccc(-c2nc(-c3ccc(F)cc3)c(-c3ccncc3)[nH]2)cc1
 COc1ccc(-c2nc(-c3ccc(F)cc3)c(-c3ccccc3)[nH]2)cc1
 CSc1ccc(-c2nc(-c3ccc(O)cc3)c(-c3ccncc3)[nH]2)cc1
 CSc1ccc(-c2nc(-c3ccc(F)cc3)c(-c3ccncc3)[nH]2)cc1
 COc1ccc(-c2nc(-c3ccc(O)cc3)c(-c3ccccc3)[nH]2)cc1
 COc1ccc(-c2nc(-c3ccc(F)cc3)c(-c3ccnnc3)[nH]2)cc1
NC(COc1cncc(-c2ccc3[nH]c(=O)oc3c2)c1)Cc1c[nH]c2ccccc12
```

```
 NC(COc1cccc(-c2ccc3[nH]c(=O)oc3c2)c1)Cc1c[nH]c2ccccc12
 NC(COc1cncc(-c2ccc3[nH]c(=O)[nH]c3c2)c1)Cc1c[nH]c2ccccc12
O=C(NC(CO)c1ccccc1)c1cccc(-c2ccncc2)c1
 CCC(NC(=O)c1ccc(-c2ccncc2)cc1)c1ccccc1
 CC(NC(=O)c1ccc(-c2ccncc2)cc1)c1ccccc1
 C=C(NC(CO)c1ccccc1)c1ccc(-c2ccncc2)cc1
CCCS(=O)(=O)Nc1ccc(-c2ccc3[nH]nc(NC(C)=O)cc2-3)cc1
 CCCS(=O)(=O)Nc1ccc(-c2ccc3[nH]nc(NC(C)=O)c3c2)cc1
 C=C(C)Nc1n[nH]c2ccc(-c3ccc(NS(=O)(=O)CCC)cc3)cc12
Cc1cccc2c3c(ccc1)C(=CC=CCC=C2)NCOC=C3
 O=c1[nH]c2cc(-c3ccccc3)cnc2[nH]1
COc1cc(C=C2SC(=Nc3ccccc3)NC2=O)ccc1O
CNc1ncnc2c1N=C(c1ccc(NC(=O)Nc3cccc(C(F)(F)F)c3)cc1)CCN2
 CNc1ccnc2c1N=C(c1ccc(NC(=O)Nc3cccc(C(F)(F)F)c3)cc1)CCN2
Nc1=c-c2nnc(SCc3cn4ccccc4n3)n2-c2ccc-1cc2
 Nc1ccc(-c2cncc(OCC(N)Cc3c[nH]c4ccccc34)c2)cc1C(=O)c1ccccc1
CSc1ccc2nc3c(c(Cl)c2c1)CCNC3=O
 CCc1ccc2nc3c(c(Cl)c2c1)CCNC3=O
CC(C)(CO)CNc1nccc(C2=C3C=CN=C(c4ccc(F)cc4)C2(C)N=C3)n1
c1cnc2nc(-c3ccc4[nH]ncc4c3)c(NC3CCCCC3)n2c1
 c1cnc2cc(-c3ccc4[nH]ncc4c3)c(NC3CCCCC3)n2c1
 c1ccn2c(NC3CCCCC3)c(-c3ccc4[nH]ccc4c3)nc2c1
 c1cnc2nc(-c3ccc4[nH]ccc4c3)c(NC3CCCCC3)n2c1
CNC(=O)C(C)n1cc(-c2cnc(N)c3c(-c4ccc(NC(=O)Nc5ccc(C)cc5)cc4)csc23)cn1
O=C1CCc2c1cccc2-c1ccc(Nc2nc3ccccc3[nH]2)cc1
 O=C1NCc2c1cccc2-c1ccc(Nc2nc3ccccc3[nH]2)cc1
Nc1ncnc(Nc2cc(CNC(=O)C(F)(F)F)c(O)c(-c3ccc(Cl)cc3)c2)n1
 Nc1nccc(Nc2cc(CNC(=O)C(F)(F)F)c(O)c(-c3ccc(Cl)cc3)c2)n1
 Nc1cccc(Nc2cc(CNC(=O)C(F)(F)F)c(O)c(-c3ccc(Cl)cc3)c2)n1
Nc1ncnc2occ(-c3ccc(NC(=O)Nc4ccc(O)cc4)cc3)c12
 Cc1ccc(NC(=O)Nc2ccc(-c3coc4ncnc(N)c34)cc2)cc1
 Nc1ccc(NC(=O)Nc2ccc(-c3coc4ncnc(N)c34)cc2)cc1
CN1CCC(NC(=O)c2cc3c(-c4ccccc4)c[nH]c3s2)CC1
 CN1CCC(NC(=O)c2cc3c(-c4ccccc4)n[nH]c3s2)CC1
 CN1C=CCCC(NC(=O)c2cc3c(-c4ccccc4)n[nH]c3s2)CC1
```

```
CCOC(=O)c1ccc2c(c1C)CCCCCC(=O)C=CN2
 C=C1NCCC2=C3C=CC(C(=O)OCC)=CC=C3NC12
O=C(NCc1ccccc1)c1cccc(-c2cnc3[nH]ccc3n2)c1
 O=C(NCc1ccccc1)c1cccc(-c2cc3cc[nH]c3cn2)c1
 O=C(NCc1ccccc1)c1cccc(-c2ccc3[nH]ccc3c2)c1
 O=C(NCc1ccccc1)c1cccc(-c2cnc3[nH]ccc3c2)c1
Cc1cccc(NC(=O)Nc2ccc(NC(=O)c3csc4nccc(N)c34)cc2)c1
 Cc1cccc(NC(=O)Nc2ccc(NC(=O)c3ccccn4nc(N)c3-4)cc2)c1
 Cc1cccc(NC(=O)Nc2ccc(NC(=O)c3csc4ncnc(N)c34)cc2)c1
CN(C)c1ccc(COC(=O)c2cc3c(s2)=CN=NN=NC=3c2ccccc2)cc1
 CN(C)c1ccc(CNC(=O)c2cc3c(-c4ccccc4)n[nH]c3s2)cc1
O=C(Nc1cccc(C(F)(F)F)c1)Nc1ccnc2c(Cl)cccc12
 O=C(Nc1cccc(C(F)(F)F)n1)Nc1ccnc2c(Cl)cccc12
 CC1=c2nccc(NC(=O)Nc3cccc(C(F)(F)F)c3)c2=CC=CC=C1
 Cc1c2c(O)c3cccc(C(C)(F)F)c(O)c1c([nH]c(=O)[nH]3)=CC=N2
CN(C(=O)C1CCCCC1)c1ccc2c(c1)N=C2NC(=O)c1ccc(C#N)cc1
 CN(C(=O)C1CCCCC1)c1ccc2c(c1)N=C2NC(=O)c1ccc(CNN)cc1
 CN(C(=O)C1CCCCC1)c1ccc2c(c1)nc(NC(=O)c1ccc(C#N)cc1)n2C
c1cc2[nH]ncc2cc1-c1cn(CC2CCOCC2)nn1
 c1cc2[nH]ncc2cc1-c1cnn(CC2CCOCC2)c1
CC(C)(CO)c1nnc2ccc(-c3c(-c4ccc(F)cc4F)nc4occn34)nn12
 Cc1ccc(-c2cc3cnccn3c2-c2ccnc(NCC3CC3)n2)cc1
 Cc1ccc(-c2nc3cnccn3c2C2=CC=NC3=NCCC2CN3)cc1
 CC1C=CC(c2nc3cnccn3c2-c2ccnc(NCC3CC3)n2)=CC1
 Cc1ccc(-c2nc3cnccn3c2C2=CC=NC3=NN=CC2CN3)cc1
 Cc1ccc(-c2nc3cnccn3c2-c2ccnc(CCC3CC3)n2)cc1
Cc1ccccccccc2c(-c3cccc(NCC(C)(C)CO)n3)c(-c3ccc(F)cc3F)nc1-2
 Cc1ccc(-c2nc3cnccn3c2-c2ccnc(NCC3CC3)n2)cc1
Cn1cc(-c2cnn3c(N)c(-c4ccc(NC(=O)Nc5cccc(C(F)(F)F)c5)cc4)cnc23)cn1
CC(C)c1cccc2c(S(=O)(=O)N(CCN)c3cncc(C4=CC=CC=CN=CC=CC=C4)c3)cccc12
 CC(C)c1cccc2c(S(=O)(=O)N(CCN)c3cncc(-c4ccc5cnccc5c4)c3)cccc12
O=C(NCc1ccc(Cl)c(Cl)c1)NC1=CC=CC=NNC=CC=C1
 O=C1NCc2ccc(Cl)c(c2)C=C2C=CC(=CC=CC=NC=NN=N2)N1
 O=C1NCc2ccc(Cl)c(c2)C=C2C=CC(=CC=CC=NN2)N1
 Cc1cc(CNC(=O)NC2=CC=CC=NNN=CC=C2)ccc1Cl
```

```
Nc1nccc2scc(-c3ccc(NC(=O)Nc4cccc(Cl)c4)cc3)c12
 Cc1nccc2scc(-c3ccc(NC(=O)Nc4cccc(Cl)c4)cc3)c12
 Cc1cccc2[nH]c(=O)cc3ccc(-c4csc5ccnc(N)c45)c-3c12
 Nc1cccc2scc(-c3ccc(NC(=O)Nc4cccc(Cl)c4)cc3)c12
 Cc1cccc2scc(-c3ccc(NC(=O)Nc4cccc(Cl)c4)cc3)c12
Oc1ccc(Nc2[nH]nc(-c3ccc(O)cc3)c2-c2ccc(O)cc2)cc1
Cc1n[nH]c2nccc(Oc3c(F)cc(Nc4cc(Cl)nc(N)n4)cc3F)c12
 Cc1c[nH]c2cccc(Oc3c(F)cc(Nc4cc(Cl)nc(N)n4)cc3F)c12
 Cc1c[nH]c2nccc(Oc3c(F)cc(Nc4cc(Cl)ncc[nH]nn4)cc3F)c12
 Cc1c[nH]c2nccc(Oc3c(F)cc(Nc4cc(Cl)nc(N)n4)cc3F)c12
O=c1c(NCc2cccc(C(F)(F)F)c2)c(Nc2ccccc2)c1=O
 CC(O)(F)c1cccc(CNc2c(Nc3ccncc3)c(=O)c2=O)c1
 O=C1C(Nc2ccccc2)=C2NCc3cccc(C(F)(F)F)c3CC12
 O=C1C(=O)C(c2cccc(C(F)(F)F)c2)CC=C1Nc1ccncc1
 O=CCC(=O)C(Nc1ccncc1)=C1NCc2cccc(C(O)(F)F)c21
 CC(C)(F)c1cccc(CNc2c(Nc3ccccc3)c(=O)c2=O)c1
 C=C1C(=O)C(NCc2cccc(C(C)(F)F)c2)=C1Nc1ccncc1
 CC(F)(F)c1cccc(CNc2c(Nc3ccncc3)c(=O)c2=O)c1
 O=c1c(NCc2cccc(C(F)(F)F)c2)c(Nc2ccncc2)c1=O
 C=C1C(=O)C(NCc2cccc(C(C)(F)F)c2)=C1Nc1ccccc1
CCCCn1c(NC(=O)c2cccc(C#N)c2)nc2cc(N(C)C(=O)C3CCCC3)ccc21
O=c1c(NCc2ccccc2O)c(Nc2ccncc2)c1=O
 Cc1ccccc1CNc1c(Nc2ccncc2)c(=O)c1=O
 Cc1ccccc1CNC1C(=O)C(=O)C1Nc1ccncc1
 Cc1ccccc1CNc1c(Nc2ccccc2)c(=O)c1=O
 Cc1ccccc1CNC1=CNc2ccncc2CC(=O)C1=O
 Cc1ccccc1CNc1c(Nc2ccncc2CCC=O)c1=O
 C=C1C(=O)Cc2ccccc2NC=C1NCc1ccccc1C
 Cc1ccccc1CNC1=C(Nc2ccccc2CCO)C1OO
 Cc1ccccc1CNC1=C(Nc2ccncc2)C(=O)C1OO
 Cc1ccccc1CNC1=C(Nc2ccncc2)C(=O)C1
Nc1nccc2scc(-c3ccc(NC(=O)Nc4ccccc4C(F)(F)F)cc3)c12
 CC(F)(F)c1ccccc1NC(=O)Nc1ccc(-c2csc3ccnc(N)c23)cc1
 Cc1nccc2scc(-c3ccc(NC(=O)Nc4ccccc4C(F)(F)F)cc3)c12
Nc1ncc(C=CC(=O)NCCCn2ccnc2)c2scc(-c3ccc(Br)cc3)c12
```

```
 Nc1ncc(C=CC(=O)CCCCn2ccnc2)c2scc(-c3ccc(Br)cc3)c12
 NBc1ccc(-c2csc3c(C=CC(=O)CCCCn4ccnc4)cnc(N)c23)cc1
 Nc1ncc(C=CC(=O)NCNCn2ccnc2)c2scc(-c3ccc(Br)cc3)c12
 Cc1ncc(C=CC(=O)NCCCn2ccnc2)c2scc(-c3ccc(Br)cc3)c12
 Nc1ccc(C=CC(=O)CCCCn2ccnc2)c2scc(-c3ccc(Br)cc3)c12
 Nc1ncc(C=CC(=O)OCCCn2ccnc2)c2scc(-c3ccc(Br)cc3)c12
O=C(NC1=CNCC=N1)Nc1ccnc2ccccc12
 O=C(Nc1cncnn1)Nc1cccc2ccccc12
 O=C(Nc1ccnc2ccccc12)NC1N=CC=NN1
 O=C(Nc1cnccn1)Nc1ccnc2ccccc12
 O=C(Nc1ccnc2ccccc12)NC1CNCC=N1
 O=C(Nc1cncnc1)Nc1ccnc2ccccc12
 O=C(NC1=C=CN=C1)Nc1ccnc2ccccc12
 O=C(Nc1cncnn1)Nc1ccnc2ccccc12
 O=C(Nc1ccccn1)Nc1ccnc2ccccc12
 O=C(Nc1cnccn1)Nc1cccc2ccccc12
 CN1CCN(c2cccc(Nc3nccc(-c4c(C(N)=O)nc5ccccn45)n3)c2)CC1
Cc1cn2c(-c3ccnc(NCC(C)(C)CO)n3)c(-c3ccc(F)cc3F)nc2c(C)n1
 Cc1cn2c(-c3ccnc(NCC(C)(C)CO)n3)c(-c3ccc(F)cc3)nc2c(C)n1
Nc1cn(-c2cc3c(Oc4ccc(Cl)cc4)cncc3s2)nn1
COc1ccc(C(=O)Nc2ccc3[nH]ncc3c2)cc1
O=C(NNC(=S)Nc1ccc(F)cc1)C(O)(c1ccccc1)c1ccccc1
 CC(C(=O)NNC(=S)Nc1ccc(F)cc1)(c1ccccc1)c1ccccc1
O=c1c(NCc2ccc(Cl)cc2)c(Nc2ccncc2)c1=O
 O=c1c(NCc2ccc(Cl)cc2)c(Nc2ccnnc2)c1=O
 O=c1c(NNc2ccc(Cl)cc2)c(Nc2ccncc2)c1=O
CC1=CC=C1NC(=O)CCNC(=O)Nc1nc(C)c(-c2ccc(-n3cccc3)cc2)s1
Cc1cc(Nc2cc(N3CCN(C)CC3)nc(C=Cc3ccccc3)n2)n[nH]1
 Cc1cc(Nc2cc(N3CCN(C)CC3)nn2CCCCc2ccccc2)n[nH]1
 Cc1cc(Nc2cc(N3CCN(C)CC3)nc(CCCc3ccccc3)n2)n[nH]1
Nc1ncnc2scc(C(=O)Nc3ccc(NC(=O)Nc4ccccc4)cc3)c12
Cc1ccc(C=NNC(=O)Nc2cccc3csnc23)c(O)c1
 O=C1NC=CC=CC=CC=NC=CC=CC2C(O)=CC=C2C=NN1
 O=C(NN=CC1(OO)C=CC(O)=C1)Nc1cccc2nsnc12
 O=C1NNCCC=CC=C(O)C(O)=C2C3=CSC=C2C(=CC=C3)N1
```

```
 O=C(NN=Cc1ccc(O)cc1O)Nc1cccc2nsnc12
 O=C(NN=Cc1ccc(O)cc1O)Nc1cccc2nncn12
 O=C(NN=Cc1ccc(O)cc1O)Nc1cccc2nscc12
COc1cccc(C(C)NC(=O)c2ccc(-c3ccncc3)cc2NCCCN)c1
 COc1cccc(C(C)NC(=O)c2ccc(-c3ccncc3C)cc2)c1
 COc1cccc(C(C)NC(CO)c2ccc(-c3ccccc3)cc2NCCCN)c1
 COc1cccc(C(C)NC(=O)c2ccc3cc2NC(CN)=CC=CC=C3)c1
 COc1cccc(C(C)NC(=O)c2ccc(C3=CC=C3)cc2NCCCN)c1
 COc1cccc(C(C)NC(=O)c2ccc(-c3ccncn3)cc2)c1
Nc1ncnc2c1C1=C3C=CC(NC(=O)Nc4ccccc4)=CC31CC=CS2
 Nc1ncnc2sc3c(c12)C=C1C=C(NC(=O)Nc2ccccc2)C=C1CC3
 Nc1ncnc2c1C1=CC3=CC(NC(=O)Nc4ccccc4)=CC31CC=CS2
 Nc1ncnc2sc3c(c12)-c1ccc(NC(=O)Nc2ccccc2)cc1CC3
 Nc1nccc2sc3c(c12)-c1ccc(NC(=O)Nc2ccccc2)cc1CC3
Cc1cccc(NC(=O)Cc2ccc(-c3cccc4[nH]nc(N)c34)cc2)c1
 Cc1cc(NC(=O)Cc2ccc(-c3cccc4[nH]nc(N)c34)cc2)ccc1F
COc1ccc2c(NC(=O)Nc3cccc(Br)n3)cnnc2c1
 COc1ccc2c(NC(=O)Nc3cccc(Br)n3)ccnc2c1
 COc1ccc2c(NC(=O)Nc3cccc(Br)n3)cccc2c1
O=C(Nc1cccc(N2CCCCC2)n1)Nc1ccnc2c(C(F)(F)F)cccc12
 O=C(Nc1cccc(N2CCOCC2)c1)Nc1ccnc2c(C(F)(F)F)cccc12
 O=C(Nc1cccc(N2CCOCC2)n1)Nc1ccn2c(C(F)(F)F)cccc12
COCCN(C)C(=O)c1c(-c2ccc3[nH]ncc3c2)nnn1Cc1ccccc1
 COCCN(C)C(=O)c1c(-c2ccc3[nH]ncc3c2)cnn1Cc1ccccc1
Cc1ccccc1-c1cc(-c2cc3c([nH]2)CCNC3=O)ccn1
 Nc1ccccc1-c1cc(-c2cc3c([nH]2)CCNC3=O)ccn1
 Cc1ccccc1-c1cc(C2=CC3=CCNC(=O)C3=C2)ccn1
Cc1cccc(NC(Cc2ccc(-c3csc4c(C#CCN(C)C)cnc(N)c34)cc2)OO)c1
 Cc1cccc(NC(=O)Nc2ccc(-c3csc4c(C#CCN(C)C)cnc(N)c34)cc2)c1
 Cc1cccc(NC(=O)Nc2ccc(-c3csc4c(CCCCN(C)C)cnc(N)c34)cc2)c1
 CN(C)CCCCc1cnc(N)c2c(-c3ccc(NC(=O)Nc4cccc(F)c4)cc3)csc12
 Cc1cccc(NC(Nc2ccc(-c3csc4c(C#CCN(C)C)cnc(N)c34)cc2)OO)c1
 Cc1cccc(NC(=O)Cc2ccc(-c3csc4c(C#CCN(C)C)cnc(N)c34)cc2)c1
 CN(C)CC#Cc1cnc(N)c2c(-c3ccc(NC(Nc4cccc(F)c4)OO)cc3)csc12
CCCC(=O)Nc1n[nH]c2nnc(-c3cccc(F)c3F)cc12
```

```
CCCC(=O)Nc1n[nH]c2ncc(-c3cccc(F)c3F)cc12
CCCC(=O)Nc1n[nH]c2cnc(-c3cccc(F)c3F)cc12
CCCC(=O)Nc1n[nH]c2ccc(-c3cccc(F)c3CC)cc12
CCCC(=O)Nc1n[nH]c2nnc(-c3cccc(F)c3)cc12
CCCC(=O)Nc1n[nH]c2ncc(-c3cccc(F)c3)cc12
CCCC(=O)Nc1n[nH]c2nnc(-c3cccc(F)c3CF)cc12
CCCC(=O)Nc1n[nH]c2ncc(-c3cccc(C)c3)cc12
Nc1nccc2scc(C3=CC=CNC(=O)NC4=CCCC(=CC=C4)C=C3)c12
Nc1cccc2scc(-c3ccc(NC(=O)Nc4cccc(F)c4)cc3)c12
Cc1cccc(NC(=O)Nc2ccc(-c3csc4ccnc(C)c34)cc2)c1
Cc1nccc2scc(-c3ccc(NC(=O)Nc4cccc(F)c4)cc3)c12
Cc1cccc2scc(-c3ccc(NC(=O)Nc4cccc(F)c4)cc3)c12
Cc1cccc(NC(=O)Nc2ccc(-c3csc4cccc(N)c34)cc2)c1
Nc1nccc2scc(-c3ccc(NC(=O)Nc4cccc(F)c4)cc3)c12
Nc1nccc2scc(-c3ccc(NC(=O)NC4=CC=CC=CCC=C4)cc3)c12
Cc1cccc(NC(=O)Nc2ccc(-c3csc4cccc(C)c34)cc2)c1
CN1C=CC(C(=O)Nc2cnc3[nH]cc(-c4ccccc4)c3c2)=CC=N1
Nc1ccc(-c2n[nH]c(Nc3cccc(Cl)c3)c2-c2ccc(O)cc2)cc1
Oc1ccc(-c2n[nH]c(Nc3cccc(Cl)c3)c2-c2ccc(O)cc2)cc1
COc1cccc(C(C)NC(=O)c2ccc(-c3ccncc3)c(O)c2)c1
COc1cccc(C(C)NC(=O)c2ccc(-c3cccnc3)c(C)c2)c1
COc1cccc(C(C)NC(=O)c2ccc(-c3ccccc3)c(C)c2)c1
COc1cccc(C(C)NC(=O)c2ccc(-c3ccnnc3)c(C)c2)c1
COc1cccc(C(C)NC(=O)c2ccc(-c3ccncc3)c(C)c2)c1
COc1cccc(C(C)NC(=O)c2ccc(-c3ccccc3)c(F)c2)c1
COc1cccc(C(C)NC(=O)c2ccc(-c3ccncc3)c(S)c2)c1
COc1cccc(C(C)NC(=O)c2ccc(-c3ccncc3)c(F)c2)c1
CCc1cc(Cl)cc(C(CO)NC(=O)c2ccc(-c3ccncc3)cc2)c1
O=C(NC(CO)c1ccccc1)N1CC=C(c2c[nH]c3ncccc23)CC1
NCc1cc(Cl)cc(C(CO)NC(=O)c2ccc(-c3ccccc3)cc2)c1
Cc1cc(N)cc(NC(=O)Nc2ccc(-c3cccc4c3CNC4=O)c(C(F)(F)F)c2)c1
Cc1cc(C)cc(NC(=O)Nc2ccc(-c3cccc4c3CNC4=O)c(C(F)(F)F)c2)c1
CCCCn1c(NC(=O)c2cccc([NH+]([O-])[O-])c2)cc2ccccc21
CCCCn1c(NC(=O)c2cccc([NH+]([O-])[O-])c2)nc2ccccc21
O=C(Nc1nc2ccccc2n1CCCO)c1cccc([NH+]([O-])[O-])c1
```

```
COCCNc1ccc2c3c(cccc13)C(=O)N(CCNCCO)C2=O
 CCCCNc1ccc2c3c(cccc13)C(=O)N(CCNCCO)C2=O
CCNC(=O)c1cnc(N)c2c(-c3ccc(NC(=O)Nc4cccc(F)c4)cc3)csc12
 CCCC(=O)c1cnc(N)c2c(-c3ccc(NC(=O)Nc4cccc(F)c4)cc3)csc12
 CCNC(=O)c1cnc(N)c2c(-c3ccc(CC(=O)Nc4cccc(F)c4)cc3)csc12
O=c1[nH]c2sccc2c2nc(-c3ccncc3)nn12
 O=c1[nH]c2sccc2c2nc(-c3ccncc3)cn12
CNC(=O)c1cc(Oc2ccc(NC(=O)Nc3cccc(C(C)(F)F)c3)cc2)ccn1
 CNC(=O)c1cc(Oc2ccc(NC(=O)Nc3cccc(C(F)(F)F)c3)cc2)ccn1
Nc1ncnc2c1sc1ncncc[nH]cc12
 Nc1ncnc2c1sc1ncnc(N)c12
 NC1N=CN=C2C1=Cc1ncncc[nH]cc12
CNS(=O)(=O)c1ccccc1Nc1nc(Nc2cc(OC)c(OC)c(OC)c2)ncc1CCl
 CNS(=O)(=O)c1ccccc1Nc1nc(Nc2cc(OC)c(OC)c(OC)c2)ncc1Cl
 CCc1cc(Nc2ncc(Cl)c(Nc3ccccc3S(=O)(=O)CC)n2)cc(OC)c1OC
 CCc1c(OC)cc(Nc2ncc(Cl)c(Nc3ccccc3S(=O)(=O)NC)n2)cc1OC
 CCc1cnc(Nc2cc(OC)c(OC)c(OC)c2)nc1Nc1ccccc1S(=O)(=O)NC
 CCc1cc(Nc2ncc(Cl)c(Nc3ccccc3S(=O)(=O)NC)n2)cc(OC)c1OC
 CCc1cc(Nc2nccc(Nc3ccccc3S(=O)(=O)NC)n2)cc(OC)c1OC
 CNS(=O)(=O)c1ccccc1Nc1nc(Nc2cc(OC)c(OC)c(OC)c2)ncc1C
 CCS(=O)(=O)c1ccccc1Nc1nc(Nc2cc(OC)c(OC)c(OC)c2)ncc1Cl
 CNS(=O)(=O)c1ccccc1Nc1nc(Nc2cc(OC)c(OC)c(OC)c2)ccc1Cl
Nc1nccc(-c2c(-c3ccc(F)cc3)ncn2C2CCNCC2)n1
 Nc1nccc(-c2c(-c3ccc(F)cc3)ccn2C2CCCCC2)n1
 Nc1cc(-c2c(-c3ccc(F)cc3)ncn2C2CCNCC2)ccn1
 Nc1cccc(-c2c(-c3ccc(F)cc3)ncn2C2CCNCC2)n1
 Nc1cc(-c2c(-c3ccc(F)cc3)ccn2C2CCNCC2)ccn1
 Fc1ccc(-c2ncn(C3CCNCC3)c2-c2ccnc(F)n2)cc1
Nc1ccc(Oc2cncc3sc(C4=NC(OO)ON4)cc23)cc1
 Nc1ccc(Oc2cncc3sc(-c4nc(=O)[nH][nH]4)cc23)cc1
 Nc1ccc(Oc2cncc3sc(C4=NC(=O)CN4)cc23)cc1
 Nc1ccc(Oc2cncc3sc(-c4nc(=O)oo4)cc23)cc1
 Nc1ccc(Oc2cncc3sc(-c4nc(=O)o[nH]4)cc23)cc1
Cc1cc(CNC2=C(Nc3ccncc3)C(CO)C2=O)ccc1Cl
 C=C1C(=O)C(NNc2ccc(Cl)c(Cl)c2)=C1Nc1ccncc1
```

```
 CCC1C(=O)C(NCc2ccc(Cl)c(Cl)c2)=C1Nc1ccccc1
 CCC1C(=O)C(NCc2ccc(Cl)c(Cl)c2)=C1Nc1ccncc1
 C=C1C(=O)C(NCc2ccc(Cl)c(Cl)c2)=C1Nc1ccncc1
 O=c1c(NCc2ccc(Cl)c(Cl)c2)c(Nc2ccncc2)c1=O
 O=C1C(NCc2ccc(Cl)c(Cl)c2)=C(Nc2ccncc2)C1CO
 CCc1cc(CNc2c(Nc3ccncc3)c(=O)c2=O)ccc1Cl
 O=c1c(NCc2ccc(Cl)c(Cl)c2)c(Nc2ccccc2)c1=O
 Cc1cc(CNc2c(Nc3ccncc3)c(=O)c2=O)ccc1Cl
CCS(=O)(=O)c1cccc(-c2cnc(N)c3c(-c4ccc5[nH]c(C)cc5c4)csc23)c1
 Cc1cc2cc(-c3csc4c(-c5cccc(S(C)(=O)=O)c5)cnc(N)c34)ccc2[nH]1
Cc1nccc2c(NC(=O)Nc3cccc(C(F)(F)F)n3)ccnc12
 O=C(Nc1cccc(C(F)(F)F)n1)Nc1ccnc2c(F)cccc12
 O=C(Nc1cccc(C(F)(F)F)n1)Nc1ccnc2c1C=CCC2F
 O=C(Nc1cccc(C(F)(F)F)n1)Nc1ccnc2c(F)nccc12
 Cc1cccc2c(NC(=O)Nc3cccc(C(F)(F)F)n3)cccc12
 Cc1cccc2c(NC(=O)Nc3cccc(C(F)(F)F)n3)ccnc12
 Cc1nccc2c(NC(=O)Nc3cccc(C(F)(F)F)n3)cccc12
 O=C(Nc1cccc(C(F)(F)F)n1)Nc1cccc2c(F)cccc12
Cc1ccc(-c2cnc(N)c3c(-c4ccc(NC(=O)Nc5cccc(F)c5)cc4)csc23)cn1
CC(N)Cc1cc(-c2ccc3[nH]nc(N)c3c2)on1
 CC(N)Cc1cc(C2=CC=CC(N)=NNC=CC=C2)on1
 CC(C)Cc1cc(-c2ccc3[nH]nc(N)c3c2)on1
 CC(C)CC1=NCC(c2ccc3[nH]nc(N)c3c2)=C1
Cc1cccc(NC(=O)Nc2ccc(-c3ccc4scnc(N)c3-4)cc2)c1
 Cc1cccc(NC(=O)Nc2ccc(-c3coc4ncnc(N)c34)cc2)c1N
 Cc1cccc(NC(=O)Nc2ccc(-c3coc4cccc(N)c34)cc2)c1
 Cc1cccc(NC(=O)Nc2ccc(-c3csc4nccc(N)c34)cc2)c1
 Cc1cccc(NC(=O)Nc2ccc(-c3coc4nccc(N)c34)cc2)c1
CN1CCN(C2CCC(c3ccccc(N)ccc(-c4ccc(Oc5ccccc5)cc4)c3)CC2)CC1
 CN1CCN(C2CCC(n3cc(-c4ccc(Oc5ccccc5)cc4)c4c(N)nccc43)CC2)CC1
 CN1CCN(C2CCC(n3cc(-c4ccc(Oc5ccccc5)cc4)c4c(N)ccnc43)CC2)CC1
 CN1CCN(C2CCC(n3cc(-c4ccc(Oc5ccccc5)cc4)c4c(N)ncnc43)CC2)CC1
Cc1cc(C)cc(NC(=O)Nc2ccc(-c3cccc4c3=C(N)N=CN=CC=4)cc2)c1
 Cc1cc(C)cc(NC(=O)Nc2ccc(-c3cccc4c3=C(N)C=CC=CC=4)cc2)c1
 Cc1cc(C)cc(NC(=O)Nc2ccc(C3=C4C(N)=NC=NN=CC43C)cc2)c1
```

```
 Cc1cc(C)cc(NC(=O)Nc2ccc(-c3c(C)sc4ncnc(N)c34)cc2)c1
 Cc1cc(C)cc(NC(=O)Nc2ccc(-c3cccc4c3=C(N)N=CC=CC=4)cc2)c1
 Cc1cc(C)cc(NC(=O)Nc2ccc(C3=C4C(N)=NC=CN=CC43C)cc2)c1
 Cc1cc(C)cc(NC(=O)Nc2ccc(-c3c(C)sc4ccnc(N)c34)cc2)c1
 O=C(NC(=O)c1ccc(-c2ccccc2)cc1)C1=CC=CC=CC=C1
 O=C(NC(CO)c1cccccccc(Cl)c1)c1ccc(-c2ccncc2)cc1
 O=C(NC(CO)c1cc(Cl)cc(Cl)c1)c1ccc(-c2ccncc2)cc1
 O=C(NC(CO)c1cc(Cl)cc(CO)c1)c1ccc(-c2ccncc2)cc1
Cc1cc2ccc(NS(=O)(=O)c3ccc(N)cc3)cc2nc1C
 Cc1nc2ccc(NS(=O)(=O)c3ccc(N)cc3)cc2nc1C
 Cc1cnc2cc(NS(=O)(=O)c3ccc(N)cc3)ccc2n1
COc1cc(-c2ccc3c(c2)Nc2ccc(C(=O)CCCNC(C)=O)cc2NC3=O)ccc1O
 COc1cc(-c2ccc3c(c2)Nc2ccc(C(=O)CNNNC(C)=O)cc2NC3=O)ccc1O
 COc1cc(-c2ccc3c(c2)Nc2ccc(C(=O)NCNNC(C)=O)cc2NC3=O)ccc1O
 COc1cc(-c2ccc3c(c2)Nc2ccc(C(NCCNC(C)=O)OO)cc2NC3=O)ccc1O
 COc1cc(-c2ccc3c(c2)Nc2ccc(C(O)CCCCCC(C)=O)cc2NC3=O)ccc1O
 COc1cc(-c2ccc3c(c2)Nc2ccc(C(=O)NCCCC(C)=O)cc2NC3=O)ccc1O
 COc1cc(-c2ccc3c(c2)Nc2ccc(C(NCCCC(C)=O)OO)cc2NC3=O)ccc1O
 C=C(NCCCC(C)=O)c1ccc2c(c1)NC(=O)c1ccc(-c3ccc(O)c(OC)c3)cc1N2
 COc1cc(-c2ccc3c(c2)Nc2ccc(C(NCNNC(C)=O)OO)cc2NC3=O)ccc1O
 COc1cc(-c2ccc3c(c2)Nc2ccc(C(=O)NCCNC(C)=O)cc2NC3=O)ccc1O
O=c1c(NCCc2ccc(Oc3ccccc3)cc2)c(Nc2ccncc2)c1=O
CCC(NC(=O)c1ccc(-c2ccncc2)cc1)c1ccccc1
 CC(NC(CO)c1ccc(-c2ccncc2)cc1)c1ccccc1
 OCCNNC(OCc1ccc(-c2ccncc2)cc1)c1ccccc1
 C=C(NC(C)c1ccccc1)c1ccc(-c2ccncc2)cc1
 CC(CC(=O)c1ccc(-c2ccncc2)cc1)c1ccccc1
 CC(NC(=O)c1ccc(-c2ccccc2)cc1)c1ccccc1
 O=C(NC(O)c1ccccc1)c1ccc(-c2ccncc2)cc1
CCCCCCNCC#Cc1cnc(N)c2c(-c3ccc(NC(=O)Nc4cccc(C)c4)cc3)csc12
 Cc1cccc(NC(=O)Nc2ccc(-c3csc4c(CCCCN5CCCC5)cnc(N)c34)cc2)c1
 Cc1cccc(NC(=O)Nc2ccc(-c3csc4c(C#CNN5CCCC5)cnc(N)c34)cc2)c1
 Cc1cccc(NC(=O)Nc2ccc(-c3csc4c(C5CCCCCNCC5)cnc(N)c34)cc2)c1
 Cc1cccc(NC(=O)Nc2ccc(-c3csc4c(C#CCN5CCCC5)cnc(N)c34)cc2)c1
CCOCCCNc1cc(C)c2nc(C3=C(NCC(O)c4cccc(Cl)c4)C=CCC3=O)[nH]c2c1
```

CCOOCCNc1cc(C)c2nc(C3=C(NCC(O)c4cccc(Cl)c4)C=CCC3=O)[nH]c2c1
Cc1cc(N2CCOOC2)cc2[nH]c(-c3c(NCC(O)c4cccc(Cl)c4)cc[nH]c3=O)nc12
Cc1cc(N2CCOCC2)cc2[nH]c(-c3c(NCC(O)c4cccc(Cl)c4)cc[nH]c3=O)nc12
Cc1cc(C2CCOCC2)cc2[nH]c(-c3c(NCC(O)c4cccc(Cl)c4)cc[nH]c3=O)nc12
Cc1cc(N2CCOCC2)cc2[nH]c(C3=C(NCC(O)c4cccc(Cl)c4)C=CCC3=O)nc12
COc1ccc(-c2ccc(NC(=O)Nc3ccc(F)c(C)c3)cc2)c2c(N)noc12
N#Cc1ccc(-c2n[nH]c3c2Cc2ccc(OCCN4CCOCC4)cc2-3)cn1
CCCc1ccc(-c2n[nH]c3c2Cc2ccc(OCCN4CCOCC4)cc2-3)cn1
C#Cc1ccc(-c2n[nH]c3c2Cc2ccc(OCCC4CCOCC4)cc2-3)cn1
C#Cc1ccc(-c2n[nH]c3c2Cc2ccc(OCCN4CCOCC4)cc2-3)cn1
NCCc1ccc(-c2n[nH]c3c4ccc(OCCN5CCOCC5)ccc-4cc23)cn1
N#Cc1ccc(-c2n[nH]c3c2Cc2ccc(OCCN4CCCCC4)cc2-3)cn1
N#Cc1ccc(-c2n[nH]c3c2Cc2ccc(OCCC4CCOCC4)cc2-3)cn1
N#Cc1ccc(-c2c[nH]c3c2Cc2ccc(OCCN4CCCCC4)cc2-3)cn1
Cc1nc(N)sc1-c1ccnc(Nc2ccc(NNCC3CC3)cc2)n1
Fc1nc[nH][nH]sc1-c1ccnc(Nc2ccc(N3CCCCC3)cc2)n1
Cc1nc(N)sc1-c1ccnc(Nc2ccc(N3CC=CC3)cc2)n1
Cc1nc(N)sc1C1=NCC2OCCCN2c2ccc(cc2)NC=NC=C1
Nc1nc(F)c(-c2ccnc(Nc3ccc(N4CCCCC4)cc3)n2)s1
Nc1nc(F)c(-c2ccnc(Nc3ccc(N4CCOCC4)cc3)n2)s1
Cc1nc(N)sc1-c1ccnc(Nc2ccc(N3CCCCC3)cc2)n1
Cc1nc(N)sc1-c1ccnc(Nc2ccc(N3CCOOC3)cc2)n1
Cc1nc(N)sc1-c1ccnc(Nc2ccc(N3CCOCC3)cc2)n1
Cc1nc(N)sc1-c1ccnc(Nc2ccc(N3CCCOC3)cc2)n1
Nc1ncnc2c1C(C(=O)Nc1ccc(NC(CO)Nc3ccccc3)cc1)=CC2
N=C(Nc1ccccc1)Nc1ccc(NC(=O)c2csc3ncnc(N)c23)cc1
Nc1ncnc2scc(C(=O)Nc3ccc(NC(CO)Nc4ccccc4)cc3)c12
Nc1ncnc2scc(C(=O)Nc3ccc(NC(=O)Nc4ccccc4)cc3)c12
COc1cccc(C(C)NC(=O)c2ccc(-c3ccccc3C)cc2)c1
CCN(CC)CC#Cc1ccc2c(c1)-c1[nH]nc(-c3ccc(CN)nc3)c1C2
CCN(CC)CC#CC1=Cc2ccc(cn2)-c2n[nH]c3c2CC(=CC=C3)C=C1
CCN(CC)CC#Cc1ccc2c(c1)-c1[nH]nc(-c3ccc(C#N)nc3)c1C2
Cc1cccc2cccc-2c[nH]c2c[nH]c3c2c-3ccc1
Cc1cccc(Nc2nc(CC3CCCCC3N)cnc2C(N)=O)c1
NC(=O)c1ncc(NC2CCCCC2N)nc1Nc1cccc(N)c1

```
 Cc1cccc(Nc2nc(NC3CCCCC3N)ccc2C(N)=O)c1
 NC(=O)c1ccc(NC2CCCCC2N)nc1Nc1cccc(N)c1
 Cc1cccc(Nc2nc(NC3CCCCC3N)cnc2C(N)=O)c1
 Cc1cccc(Nc2nc(NC3CCCCC3N)cnc2C(N)OO)c1
 NC(=O)c1ncc(NC2CCCCC2N)nc1NC1=CC(N)=C=C1
 Nc1cccc(Nc2nc(NC3CCCCC3N)cnc2C(N)OO)c1
 Cc1cccc(Nc2nc(NN3CCCCC3N)cnc2C(N)=O)c1
 NC(=O)c1ncc(CC2CCCCC2N)nc1Nc1cccc(N)c1
CC(=O)Nc1cccc(CNc2c(Nc3ccc4cc3C=NN4)c(=O)c2=O)c1
 CC(=O)Nc1cccc(CNc2c(Nc3ccc4[nH]ncc4c3)c(=O)c2=O)c1
O=c1cc(-c2ccncc2)nc(NC2CCCOC2)[nH]1
 O=c1cc(-c2ccncc2)nc(NC2CCCCC2)[nH]1
 O=c1cc(-c2ccncc2)nc(NC2CCCCCC2)[nH]1
 O=c1nc(-c2ccncc2)nc(NC2CCCCC2)[nH]1
 O=c1cc(-c2ccccc2)nc(NC2CCCCC2)[nH]1
 O=C1C=C(c2ccccc2)NCC(NC2CCCCC2)N1
 CC1CCCCC1Nc1nc(-c2ccncc2)cc(=O)[nH]1
 COc1cccc(C(C)NC(=O)c2ccc(-c3ccncc3C)c(C)c2)c1
COc1cccc(C(C)NC(=O)c2cc(C)c(C3=CC=CC(C)=CNC=CC=C3)s2)c1
 COc1cccc(C(C)NC(=O)c2cc(C)c(-c3ccc4[nH]cc(C)c4c3)s2)c1
 COc1cccc(C(C)NC(=O)c2cc(C)c(-c3ccc4[nH]nc(C)c4c3)s2)c1
 COc1cccc(C(C)NC(=O)c2cc(C)c(C3=CC=CC(C)=NNC=CC=C3)s2)c1
CN(C)CC(=O)Nc1n[nH]c2ccc(-c3cn(Cc4ccccc4)nn3)cc12
 CN(C)CC(=O)Nc1n[nH]c2ccc(C3=CC(Cc4ccccc4)=N3)cc12
Cc1nc2ccc(C#N)cc2n1-c1nc(-c2ccccc2)cs1
 N#Cc1ccc2nc(N)n(-c3nc(-c4ccccc4)cs3)c2c1
 Nc1ccc2nc(N)n(-c3nc(-c4ccccc4)cs3)c2c1
 Cc1ccc2nc(N)n(-c3nc(-c4ccccc4)cs3)c2c1
 N#Cc1ccc2nc(N)n(-c3cc(-c4ccccc4)cs3)c2c1
 NCCc1ccc2nc(N)n(-c3nc(-c4ccccc4)cs3)c2c1
 CCc1ccc2nc(N)n(-c3nc(-c4ccccc4)cs3)c2c1
 NNNc1ccc2nc(N)n(-c3nc(-c4ccccc4)cs3)c2c1
 NNCc1ccc2nc(N)n(-c3nc(-c4ccccc4)cs3)c2c1
Cc1c(C=C2C(=O)Nc3ccccc32)[nH]c(N)c1CCC(=O)O
 Cc1[nH]c(C=C2C(=O)Nc3ccccc32)c(C)c1CCCOO
```

```
 Cc1[nH]c(C=C2C(=O)Nc3ccccc32)c(C)c1CCC(O)CO
 Cc1[nH]c(C=C2C(=O)Nc3ccccc32)c(C)c1CCC=O
 Cc1[nH]c(C=C2C(=O)Nc3ccccc32)c(C)c1CCC(O)OO
 Cc1[nH]c(C=C2C(=O)Nc3ccccc32)c(C)c1CCCCOOOO
 Cc1[nH]c(C=C2C(=O)Nc3ccccc32)c(C)c1CCC(=O)O
CC(C)COc1cnc(Cl)c(C=Cc2ccncc2)c1
 CC(N)COc1cnc(Cl)c(C=Cc2ccccc2)c1
 CC(N)COc1cnc(Cl)c(COCc2ccncc2)c1
 CCc1ncc(OCC(C)N)cc1C=Cc1ccncc1
 CC(N)COc1cnc(Cl)c(C=Cc2ccncc2)c1
CC(=O)c1sc2c(Br)ccc(Cl)c2c1N
 NC(=O)c1sc2c(Br)ccc(Cl)c2c1N
O=C(NCc1ccc(Cl)cc1)c1cc2c(-c3ccccc3)n[nH]c2s1
OCCCNc1ncnc2[nH]cc(-c3ccccc3)c12
 OCCCNc1ncnc2[nH]ccccccc3cc-3cc12
 OCC1NC=NC=Nc2[nH]cc(-c3ccccc3)c21
CCc1cc(CCN(C)CCCC(C#N)(c2ccc(OC)c(OC)c2)C(C)C)ccc1OC
 CCc1ccc(CCN(C)CCCC(C#N)(c2ccc(OC)c(OC)c2)C(C)C)cc1OC
 COc1ccc(CCN(C)CCCC(C#N)(c2ccc(OC)c(OC)c2)C(C)C)cc1
 COc1ccc(CCC(C)CCCC(C#N)(c2ccc(OC)c(OC)c2)C(C)C)cc1OC
 COc1ccc(CCN(C)CCCC(C#N)(c2ccc(OC)c(OC)c2)C(C)C)cc1O
 COc1ccc(CCN(C)CCCC(C#N)(c2ccc(OC)c(OC)c2)C(C)C)cc1C
 CCCC(C)(C#N)CCCN(C)CCC1=CC=C(OC)C(OC)=CC2=C(OC)C=CC1=CNCO2
 COc1ccc(CCN(C)CCCC(C#N)(c2ccc(OC)c(OC)c2)C(C)C)cc1OC
Cc1cc(C)c(C=C2C(=O)Nc3ccccc32)[nH]1
Nc1n[nH]c2ccc(-c3cnn(Cc4ccccc4)c3)cc12
 Nc1n[nH]c2ccc(-c3cn(Cc4ccccc4)cn3)cc12
 Nc1n[nH]c2ccc(-c3ccn(Cc4ccccc4)c3)cc12
 Nc1c[nH]c2ccc(-c3cn(Cc4ccccc4)nn3)cc12
 Nc1n[nH]c2ccc(-c3cn(Cc4ccccc4)nn3)cc12
 Clc1cc(NC2CCCCC2)nc(-c2c[nH]c3nnccc23)n1
 Clc1cc(NC2CCCCC2)nc(-c2ccnc3ncccc23)n1
 Clc1cc(NC2CCCCC2)nc(-c2c[nH]c3ncccc23)n1
 Clc1cc(NN2CCCCC2)nc(-c2c[nH]c3ncccc23)n1
 Clc1cc(CC2CCCCC2)nc(-c2c[nH]c3ncccc23)n1
```

```
 Clc1cc(NC2CCCCC2)nc(-c2ccnc3nnccc23)n1
 Clc1cc(NC2CCCCC2)nc(-c2n[nH]c3ncccc23)n1
 Clc1cc(NC2CCCCC2)nc(-c2ccnc3ccccc23)n1
 Clc1cc(NC2CCCCC2)nc(-c2c[nH]c3ccccc23)n1
Cc1ccc(NC(=O)Nc2ccc(-c3csc4c(-c5cnn(C(C)C(=O)N(C)C)c5)cnc(N)c34)cc2)cc1
 Cc1ccc(NC(Nc2ccc(-c3csc4c(-c5cnn(C(C)C(=O)N(C)C)c5)cnc(N)c34)cc2)OO)cc1
Nc1cccc2c1N=C(c1ccc(NC(=O)Nc3cc(C(F)(F)F)ccc3F)cc1)CCN2
 Nc1ncnc2c1N=C(c1ccc(NC(=O)Nc3cc(C(F)(F)F)ccc3F)cc1)CCN2
 Nc1nccc2c1N=C(c1ccc(NC(=O)Nc3cc(C(F)(F)F)ccc3F)cc1)CCN2
COc1cc(Nc2nccc(Nc3cc(C4CCCCC4)co3)n2)cc(OC)c1OC
 COc1cc(Nc2nccc(Nc3cc(C4CCCCC4)no3)n2)cc(OO)c1OC
 COc1cc(Nc2nccc(Nc3cc(C4CCCCC4)no3)n2)cc(OC)c1O
 CCOc1cc(Nc2nccc(Nc3cc(C4CCCCC4)no3)n2)cc(OC)c1C
 COc1cc(Nc2nccc(Nc3cc(C4CCCCC4)no3)n2)cc(OC)c1OC
 COc1cc(Nc2nccc(Nc3cc(C4CCCCC4)c[nH]3)n2)cc(OC)c1OC
 CCOC1(OC)C=C(Nc2nccc(Nc3cc(C4CCCCC4)no3)n2)C=C1OC
 COc1cc(Nc2nccc(Nc3cc(C4CCCCC4)no3)n2)cc(OC)c1
 CCc1c(OC)cc(Nc2nccc(Nc3cc(C4CCCCC4)no3)n2)cc1OC
CCCC(=O)Nc1n[nH]c2ccc(-c3cnn(Cc4ccccc4)c3)cc12
 CCCC(=O)Nc1n[nH]c2ccc(-c3cn(Cc4ccccc4)nn3)cc12
CCOc1ccc(NC(=O)Nc2ccc(-c3csc4c(-c5cnn(C)c5)ccc(N)c34)cc2)cc1
 CCOc1ccc(NC(=O)Nc2ccc(-c3csc4c(-c5cnn(C)c5)cnc(N)c34)cc2)cc1
 CCOc1ccc(NC(=O)Nc2ccc(-c3csc4c(-c5ccn(C)c5)cnc(N)c34)cc2)cc1
Cn1cc(C2=C=C(N)c3c(-c4ccc(NC(=O)Nc5ccc(OC(F)F)cc5)cc4)csc32)cn1
 Cn1cc(-c2cnc(N)c3c(-c4ccc(NC(=O)Nc5ccc(OC(F)F)cc5)cc4)csc23)cn1
OC1CCC(Nc2cc(Cl)nc(-c3c[nH]c4ncccc34)n2)CC1
 OC1=CC(Nc2cc(Cl)nc(-c3c[nH]c4ncccc34)n2)CC1
 CC1CCC(Nc2cc(Cl)nc(-c3c[nH]c4ncccc34)n2)CC1
 OC1CCC(Nc2cc(Cl)nc(-c3c[nH]c4ncccc34)c2)CC1
 OC1CCC(Nc2cc(Cl)cc(-c3c[nH]c4ncccc34)n2)CC1
O=C(NC(CO)c1ccccc1)C1CC=C(c2c[nH]c3ncccc23)CC1
 CCC(NC(=O)N1CC=C(c2c[nH]c3ncccc23)CC1)c1ccccc1
CC(C)(CO)CNc1nccc(-c2c(-c3ccc(F)cc3)nc3c(CC4CC4)nccn23)n1
Nc1n[nH]c2cccc(-c3ccc(NC(=O)Nc4cc(C(=O)O)ccc4F)cc3)c12
 Cc1n[nH]c2cccc(-c3ccc(NC(=O)Nc4cc(C(=O)O)ccc4F)cc3)c12
```

```
CC(F)(F)c1ccc(F)c(NC(=O)Nc2ccc(-c3csc4ncnc(N)c34)cc2)c1
 Nc1ncnc2scc(-c3ccc(NC(=O)Nc4cc(C(O)(F)F)ccc4F)cc3)c12
 Nc1ncnc2scccc3ccc(NC(=O)Nc4cc(C(F)(F)F)ccc4F)cc3cc12
 Nc1ncnc2scc(-c3ccc(NC(=O)Nc4cc(C(N)(F)F)ccc4F)cc3)c12
 Cc1ncnc2scc(-c3ccc(NC(=O)Nc4cc(C(F)(F)F)ccc4F)cc3)c12
 Nc1ncnc2scc(-c3ccc(NC(=O)Nc4cc(C(F)(F)F)ccc4F)cc3)c12
O=c1[nH]c2cnc(-n3cnc4ccc(F)cc43)cc2n1C1CCOc2c(F)cccc21
 O=c1[nH]c2cnc(-n3cnc4ccc(F)cc43)nc2n1C1CCOc2c(F)cccc21
Nc1n[nH]c2ccc(-c3nnn(Cc4ccccc4)c3-c3ccccc3)cc12
 Nc1n[nH]c2ccc(-c3nnn(Cc4ccccc4)c3-c3ccc(F)cc3)cc12
O=C(O)c1ccc(C2=CCCc3ccnc(c3)C=C2)cc1
 NC(=O)c1ccc(-c2c[nH]c3c2C(Cl)CC=N3)cc1
 NC(=O)c1ccc(-c2c[nH]c3nccc(CO)c23)cc1
 NC(=O)c1cccc(-c2c[nH]c3nccc(Cl)c23)c1
 NC(=O)c1ccc(-c2c[nH]c3nccc(Cl)c23)cc1
 NC(=O)c1ccc(-c2c[nH]c3cccc(Cl)c23)cc1
 O=C(O)c1ccc(-c2c[nH]c3nccc(Cl)c23)cc1
 NC(=O)c1ccc(-c2c[nH]c3nccc(OO)c23)cc1
 NC(=O)c1ccc(-c2c[nH]c3c2C=CCC=CCC=N3)cc1
CCCn1cc(-c2cnc(N)c3c(-c4ccc(NC(=O)Nc5cccc(F)c5)cc4)csc23)cn1
Nc1cnc(Nc2ccccc2)nc1Nc1ccccc1
 Fc1cnc(Nc2ccccc2)nc1Nc1ccccc1
 Fc1cccccccc[nH][nH]cnc(Nc2ccccc2)nc1
 Fc1ccc(Nc2ccccc2)nc1Nc1ccccc1
 Fc1cnc(Nc2ccccc2)cc1Nc1ccccc1
 Fc1ccccccc[nH][nH][nH]nc(Nc2ccccc2)nc1
 Cc1ccc(Nc2ccccc2)nc1Nc1ccccc1
 Oc1cnc(Nc2ccccc2)nc1Nc1ccccc1
 Cc1cnc(Nc2ccccc2)nc1Nc1ccccc1
O=C1NCc2c1cccc2-c1ccc(Nc2cc3ccccc3o2)cc1
 O=C1NCc2c1cccc2-c1ccc(Nc2nc3ccccc3o2)cc1
Cc1n[nH]c2ccc(-c3cccc(OCC(N)Cc4ccccc4)c3)cc12
 Cc1n[nH]c2ccc(-c3cncc(CCC(N)Cc4ccccc4)c3)cc12
 Cc1n[nH]c2ccc(-c3cncc(OCC(N)Cc4ccccc4)c3)cc12
 Cc1n[nH]c2ccc(-c3cncc(CCC(O)Cc4ccccc4)c3)cc12
```

```
 Cc1n[nH]c2ccc(-c3cncc(OCC(N)Cc4ccccn4)c3)cc12
 Cc1n[nH]c2ccc(-c3cncc(OCC(O)Cc4ccccc4)c3)cc12
 Cc1n[nH]c2ccc(-c3cncc(OCC(N)Cc4ccccc4)c3F)cc12
Cc1cccc(NC(=O)Nc2ccc(-c3csc4c(C#CCN5CC=CC5)cnc(N)c34)cc2)c1
 Cc1cccc(NC(=O)Nc2ccc(-c3csc4c(CCCCN5CCOCC5)cnc(N)c34)cc2)c1
 Cc1cccc(NC(=O)Nc2ccc(-c3csc4c(C#CCN5CCCCC5)cnc(N)c34)cc2)c1
 Cc1cccc(NC(=O)Cc2ccc(-c3csc4c(C#CCN5CCOCC5)cnc(N)c34)cc2)c1
 Cc1cccc(NC(=O)Nc2ccc(-c3csc4c(C#CCN5CCOCC5)cnc(N)c34)cc2)c1
 Cc1cccc(NC(=O)Nc2ccc(-c3csc4c(C5CCCCCNCC5)cnc(N)c34)cc2)c1
 C=CCN1CCC1c1cnc(N)c2c(-c3ccc(NC(=O)Nc4cccc(C)c4)cc3)csc12
 CCOCCNCC#Cc1cnc(N)c2c(-c3ccc(NC(=O)Nc4cccc(C)c4)cc3)csc12
 Cc1cccc(NC(=O)Nc2ccc(-c3csc4c(C#CCN5CC5O)cnc(N)c34)cc2)c1
Nc1nccc2scc(-c3ccc(NC(=O)Nc4ccccc4)cc3)c12
 Nc1ncnc2scc(-c3ccc(NC(=O)Nc4ccccc4)cc3)c12
CCCC(C(=O)Nc1ccc2[nH]ncc2c1)c1ccc(Cl)c(Cl)c1
 CCc1ccc(C(CCN)C(=O)Nc2ccc3[nH]ncc3c2)cc1Cl
 NCCC(C(=O)Nc1ccc2[nH]ncc2c1)c1ccc(Cl)c(Cl)c1
Cc1ccc(NC(=O)Nc2ccc(-c3csc4ncnc(N)c34)cc2)cc1C
 Cc1ccc(NC(=O)Nc2ccc(-c3csc4nnnc(N)c34)cc2)cc1C
 Cc1ccc(NC(=O)Nc2ccc(-c3csc4nnnc(N)c34)cc2)cc1
 CCc1cc(NC(=O)Nc2ccc(-c3csc4ncnc(N)c34)cc2)ccc1C
 Cc1ccc(NC(=O)Nc2ccc(C3=CSC4=NC=NN(N)C34)cc2)cc1
 Cc1ccc(NC(=O)Nc2ccc(C3=CC=C4N=NN=C(N)C43)cc2)cc1C
 Cc1ccc(NC(=O)Nc2ccc(-c3ccc4ncnc(N)n34)cc2)cc1
 Cc1ccc(NC(=O)Nc2ccc(-c3csc4ncnc(N)c34)cc2)cc1
Brc1cnc2[nH]nc(-c3ccccc3)c2c1
 Brc1ccc2[nH]cc(-c3ccccc3)c2c1
 Brc1cnc2[nH]cc(-c3ccccc3)c2c1
CC(C)(C(N)=O)n1cc(-c2cnc(N)c3c(-c4ccc(NC(=O)Nc5cccc(F)c5)cc4)csc23)cn1
 C=C(N)C(C)(C)n1cc(-c2cnc(N)c3c(-c4ccc(NC(=O)Nc5cccc(C)c5)cc4)csc23)cn1
 CC(C)(C(N)=O)n1cc(-c2cnc(N)c3c(-c4ccc(CC(=O)Nc5cccc(F)c5)cc4)csc23)cn1
CC(C)Cn1cc(-c2cnc(N)c3c(-c4ccc(NC(=O)Nc5cccc(F)c5)cc4)csc23)cn1
 CC(O)Cn1cc(-c2cnc(N)c3c(-c4ccc(NC(=O)Nc5cccc(F)c5)cc4)csc23)cn1
Cc1n[nH]c2ccc(-c3cncc(NCC(N)Cc4c[nH]c5ccccc45)c3)cc12
 Cc1n[nH]c2ccc(-c3cncc(OCC(N)Cc4c[nH]c5ccccc45)c3)cc12
```

```
Cc1n[nH]c2ccc(-c3cncc(OCC(N)Cc4cccc(CC(F)(F)F)c4)c3)cc12
 Cc1n[nH]c2ccc(-c3cncc(OCC(N)Cc4cccc(OC(F)(F)F)c4)c3)cc12
 Cc1n[nH]c2ccc(-c3cncc(OCC(N)Cc4cccc(OC(C)(F)F)c4)c3)cc12
NCCCOc1cc2c(c(-c3ccc(Nc4nc5ccc(Cl)cc5o4)cc3)c1)CNC2=O
 NCCCOc1cc2c(c(-c3ccc(NC4=NC=CC=CC(Cl)=CC=CO4)cc3)c1)CNC2=O
 NCCCOc1cc2c(c(-c3ccc(Nc4nccc5c(Cl)cc-5o4)cc3)c1)CNC2=O
CCC(CO)NC1=NC=CN(C(C)C)C=NC2=C3C=CC(=C3)CNC2=C1
 CCC(CO)Nc1nc(NCc2ccccc2)c2ncn(C(C)C)c2n1
CNC(=O)c1cnc(N)c2c(-c3ccc(NC(=O)Nc4cc(C)ccc4F)cc3)csc12
 CNC(=O)c1cnc(N)c2c(-c3ccc(NC(=O)NC4=CC(F)C=C4)cc3)csc12
 CNC(=O)c1cnc(N)c2c(C3=CC=C(NCC=O)NC4=CC=CCC(=C4)C=C3)csc12
 Cc1c[nH]c2ccc(-c3nnn(Cc4ccccc4)c3-c3ccccc3)cc12
 Nc1c[nH]c2ccc(-c3nnn(Cc4ccccc4)c3-c3ccccc3)cc12
 NC1=C[CH]c2ccc(-c3nnn(Cc4ccccc4)c3-c3ccccc3)cc21
 Nc1n[nH]c2ccc(-c3nnn4c3C=CC=C3C=C3C4)cc12
 Nc1n[nH]c2ccc(-c3nnn(CC4C=CC=CC4)c3-c3ccccc3)cc12
Cc1cccc(-c2[nH]c(-c3cccc(N)c3)cc2C(N)=O)c1C
 Cc1c(N)cccc1-c1[nH]c(-c2cccc(N)n2)cc1C(N)=O
 Cc1c(N)cccc1-c1[nH]c(-c2ccnc(N)n2)cc1C(N)=O
 Cc1c(N)cccc1-c1[nH]c(-c2ccnc(N)c2)cc1C(N)=O
 Cc1cccc(-c2[nH]c(-c3ccnc(N)c3)cc2C(N)=O)c1C
 Cc1cccc(-c2[nH]c(-c3ccnc(N)n3)cc2C(N)=O)c1C
 Cc1cccc(-c2[nH]c(-c3cccc(N)n3)cc2C(N)=O)c1C
C=c1[nH]c2ccc(OC)cc2c1=Cc1c[nH]cn1
 COc1ccc2c(c1)C(=Cc1c[nH]nn1)C(=O)N2
 COc1ccc2c(c1)C(=Cc1c[nH]cn1)C(=O)N2
Cc1cccc(NC(=O)Nc2ccc(-c3csc4c(C#CCNS(C)(=O)=O)cnc(N)c34)cc2)c1
 Cc1cccc(NC(=O)Nc2ccc(-c3csc4c(C(C)CCNSCCO)cnc(N)c34)cc2)c1
 Cc1cccc(NC(=O)Nc2ccc(-c3csc4c(C#CCN[SH](C)(=O)CO)cnc(N)c34)cc2)c1
O=C(Nc1ccccc1CN1CCCC1=O)c1cccnc1
 C=C1Nc2cc(CN3CCCC3=O)ccc2-n2cccc21
 O=C1CCCN1Cc1ccc2c(c1)c[n+]([O-])c1cccn21
 O=C1CCCN1Cc1ccc2c(c1)[nH]c(=O)c1cccn12
Cc1cccc(NC(=O)Nc2ccc(-c3csc4cccc(N)c34)cc2)c1
 Cc1cccc(NC(=O)Nc2ccc(-c3coc4ccnc(N)c34)cc2)c1
```

```
Oc1cccc(-c2cc3ccc4nc3c(n2)N2CC2CC4)c1
Cc1c(-c2ccc3[nH]nc(N)c3c2)nnn1Cc1ccccc1
 Cc1c(C2=CC=C3NC=C3N3C=C23)nnn1Cc1ccccc1
 Cc1c(-c2ccc3[nH]nc(N)c3c2)cnn1Cc1ccccc1
 Cc1c(-c2ccc3[nH]nc(N)cc2-3)nnn1Cc1ccccc1
 Cc1ccc(-c2ccc3[nH]nc(N)c3c2)n1Cc1ccccc1
 Nc1n[nH]c2ncc(-c3nnn(Cc4ccccc4)c3N)cc12
 Cc1c(-c2ccc3[nH]cc(N)c3c2)nnn1Cc1ccccc1
 Cc1c(-c2ccc3c(cc4cc-4c2)N3)nnn1Cc1ccccc1
 Nc1n[nH]c2ccc(-c3nnn(Cc4ccccc4)c3N)cc12
COC1=CC(C=C(C#N)c2nc3cccc-3c[nH]2)=CC(N)=CN1O
 COc1ccc(Br)c(C=C(C#N)c2nc3ccccc3[nH]2)c1
 COc1cc(C=C(C#N)c2nc3ccccc3[nH]2)c(Br)cc1O
 COc1cc(C=C(C#N)c2cc3ccccc3[nH]2)c(Br)cc1O
 BBc1cc(O)c(OC)cc1C=C(C#N)c1nc2ccccc2[nH]1
 BBc1ccc(OC)cc1C=C(C#N)c1nc2ccccc2[nH]1
NS(=O)(=O)c1cccc(Nc2ncc3ccn(Cc4ccccc4)c3n2)c1
 CS(=O)(=O)c1cccc(Nc2ncc3ccn(Cc4ccccc4)c3n2)c1
 NS(=O)(=O)c1cccc(Nc2ccc3ccn(Cc4ccccc4)c3n2)c1
Cn1cc(C=C2C(=O)Nc3cccnc32)c2ccccc21
 CC1=CC(C=C2C(=O)Nc3cccnc32)=C2CC=CC=C12
 Cc1cccccccc(C=C2C(=O)Nc3cccnc32)c1
 Cn1cc(C=C2C(=O)Nc3ccccc32)c2ccccc21
 Cc1cc2c(c(C=C3C(=O)Nc4cccnc43)c1)C=C2
OCC(CO)Nc1ncnc2[nH]cc(-c3ccccc3)c12
 OCC(CO)NC1=NC=Nc2[nH]ccc2C=C2C=C2C=C1
 OCC(NO)Nc1ncnc2[nH]cc(-c3ccccc3)c12
 OCC(CO)Nc1nnnc2[nH]cc(-c3ccccc3)c12
 OCC(CO)Nc1ncnc2[nH]cccc3ccccc3cc12
 NC(CO)COCNc1ncnc2[nH]cc(-c3ccccc3)c12
O=C1NC(=O)C(c2cnc3ccccn23)=C1c1cn2c3c(cccc13)CN(C(CO)N1CCOCC1)CC2
 O=C1NC(=O)C(c2cnc3ccccn23)=C1c1cn2c3c(cccc13)CN(C(=O)N1CCCCC1)CC2
 O=C1NC(=O)C(c2cnc3ccccn23)CC1c1cn2c3c(cccc13)CN(C(=O)N1CCCCC1)CC2
 O=C1NC(=O)C(c2cnc3ccccn23)=C1c1cn2c3c(cccc13)CN(C(CO)N1CCCCC1)CC2
 O=C1NC(=O)C(c2cnc3ccccn23)CC1c1cn2c3c(cccc13)CN(C(CO)N1CCCCC1)CC2
```

```
 O=C1NC(=O)C(c2cnc3ccccn23)=C1c1cn2c3c(cccc13)CN(C(=O)N1CCOCC1)CC2
Cc1cc(Cl)ccc1NC(=S)NNC(=O)C(O)(c1ccccc1)c1cccnc1
 Cc1cc(Cl)ccc1NC(=S)CNC(=O)C(O)(c1ccccc1)c1ccccc1
 Cc1cc(Cl)ccc1NC(=S)NNC(BO)C(O)(c1ccccc1)c1ccccc1
 CC1=C2C=CC(=C2)C(O)(c2ccccc2)C(=O)NNC(=S)NC=CC=CC(Cl)=C1
 Cc1cc(Cl)ccc1NC(=S)NNC(=O)C(O)(c1ccccc1)c1ccccc1
COc1cc(C=C(C#N)c2nc3cc(C)ccc3[nH]2)c(Br)cc1O
 COc1ccc(Br)c(C=C(C#N)c2nc3cc(C)ccc3[nH]2)c1
Cc1n[nH]c2cnc(-c3cncc(OCC(N)Cc4cccc(C(F)(F)F)c4)c3)cc12
 Cc1n[nH]c2ccc(-c3cncc(OCC(N)Cc4cccc(C(F)(F)F)c4)c3)cc12
 Cc1n[nH]c2cnc(-c3cncc(OCC(N)Cc4cccc(F)c4CC(F)CF)c3)cc12
 Cc1n[nH]c2cnc(C3=CN=CC(OCC(N)Cc4cccc(C(F)CF)c4)=CC=C3)cc12
 Cc1n[nH]c2cnc(-c3cncc(OCC(O)Cc4cccc(C(F)(F)F)c4)c3)cc12
CNS(=O)(=O)c1ccccc1Nc1nc(Nc2cc(N3CCN(C(C)=O)CC3)ccc2OC)ccc1Br
 CNS(=O)(=O)c1ccccc1Nc1nc(Nc2cc(N3CCN(C(C)OO)CC3)ccc2OC)ncc1Br
 CNS(=O)(=O)c1ccccc1Nc1cnnc(Nc2cc(N3CCN(C(C)=O)CC3)ccc2OC)n1
 CNS(=O)(=O)c1ccccc1Nc1nc(Nc2cc(N3CCN(C(C)=O)CC3)ccc2OC)ncc1Br
 CNS(=O)(=O)c1ccccc1Nc1ccnc(Nc2cc(N3CCN(C(C)=O)CC3)ccc2OC)n1
 CNS(=O)(=O)c1ccccc1Nc1nc(Nc2cc(N3CCC(C(C)=O)CC3)ccc2OC)ccc1Br
 Bc1nnc(Nc2cc(N3CCN(C(C)=O)CC3)ccc2OC)nc1Nc1ccccc1S(=O)(=O)NC
 CNS(=O)(=O)c1ccccc1Nc1nc(Nc2cc(N3CCN(C(C)=O)CC3)ccc2OC)ncc1C
 Bc1cnc(Nc2cc(N3CCNC(C(C)=O)C3)ccc2OC)nc1Nc1ccccc1S(=O)(=O)NC
 Bc1cnc(Nc2cc(N3CCN(C(C)=O)CC3)ccc2OC)nc1Nc1ccccc1S(=O)(=O)NC
Cc1cnc(Nc2cccc(S(N)(=O)=O)c2)nc1Nc1ccc(CCC(N)=O)cc1
 C=C(N)COc1ccc(Nc2nc(Nc3cccc(S(N)(=O)=O)c3)ncc2C)cc1
 Cc1ccc(Nc2cccc(S(N)(=O)=O)c2)nc1Nc1ccc(OCC(N)=O)cc1
 Cc1ccc(Nc2cccc(S(N)(=O)=O)c2)nc1Nc1ccc(OCC(=O)O)cc1
 Cc1cnc(Nc2cccc(S(N)(=O)=O)c2)nc1Nc1ccc(OCC(N)=O)cc1
ClC1=NC(CCc2ccccc2)OC(c2c[nH]c3ncccc23)=C1
 CSc1cc(C2=CNc3cc2ccn3)cc(NCc2ccccc2)n1
 Clc1cc(C2=CNc3cccc2n3)cc(NCc2ccccc2)n1
 ClC1=NC(NCc2ccccc2)NC(c2c[nH]c3ncccc23)=C1
 COc1cc(-c2c[nH]c3ncccc23)cc(NCc2ccccc2)n1
 Clc1cc(-c2c[nH]c3ncccc23)cc(NCc2ccccc2)n1
 ClC1=NC(NCc2ccccc2)OC(c2c[nH]c3ncccc23)=C1
```

```
 Clc1cc(-c2c[nH]c3nnccc23)cc(NCc2ccccc2)n1
 CSc1cc(-c2c[nH]c3ncccc23)cc(NCc2ccccc2)n1
NC1CCC(Nc2nccc(-c3c[nH]c4ncccc34)n2)NC1
 NC1CCC(Nc2nccc(C3=CNc4ccc3cn4)n2)CC1
 CC1CCC(Nc2nccc(-c3c[nH]c4ncccc34)n2)CC1
 NC1CCC(Nc2nccc(-c3c[nH]c4ncccc34)n2)CC1
O=S(O)c1ccccc1Nc1ccnc(Nc2ccc3[nH]ncc3c2)n1
 O=C(O)c1ccccc1Nc1ccnc(Nc2ccc3[nH]ncc3c2)n1
 O=C(O)c1ccccc1Nc1ccnc(NC2=CC=CC=NNN=CC=C2)n1
 O=C(O)c1ccccc1Nc1ccnc(Nc2ccc3[nH]ccc3c2)n1
 O=C(O)c1ccccc1Nc1ccnc(NC2=CC=CC=NNC=CC=C2)n1
O=C(Nc1n[nH]c2ccc(-c3cn(Cc4ccccc4)nn3)cc12)c1ccccc1
ONC(O)c1ccccc1Nn1cccc1Nc1cccc(O)c1
 ONC(O)c1ccccc1Nc1ccnc(Nc2cccc(O)c2)n1
 O=C(O)c1ccccc1Nc1ccnc(Nc2cccc(O)c2)c1
 ONC(O)c1ccccc1Nc1ccnc(Nc2cccc(O)c2)c1
 O=C(O)c1ccccc1Nc1ccnc(Nc2cccc(O)c2)n1
O=C(O)c1ccccc1Nc1ccnc(Nc2cccc(O)c2)n1
O=C1NCCc2[nH]c(-c3ccnc(C=Cc4ccccc4)c3)cc21
COc1cc2ncnc(Nc3cccc(Br)c3)c2cc1O
 COc1cc2ncnc(Nc3cccc(Br)c3)c2cc1OC
 BOc1cc2ncnc(c1O)-n-c1cccc(Br)c1-c=c-2
 COc1cc2c(Nc3cccc(Br)c3)ncnc2nc1OC
O=c1c(O)c(-c2ccc(O)cc2)oc2cc(S)cc(O)c12
 O=c1c(O)c(-c2ccc(O)cc2)oc2cc(O)cc(O)c12
O=c1cc(-c2ccc(O)cc2)oc2cc(O)cc(O)c12
Nc1ncnc2c1c(-c1cccc(NCc3ccccc3)c1)cn2[C@H]1C[C@@H](CN2CCC2)C1
 Nc1ncnc2c1c(-c1cccc(OCc3ccccc3)c1)cn2[C@H]1C[C@@H](CN2CCC2)C1
 Nc1ncnc2c1c(-c1cccc(NCc3ccccc3)c1)cn2[C@@H]1C[C@H](CN2CCC2)N1
 Nc1ncnc2c1c(-c1cccc(OCc3ccccc3)c1)cn2[C@@H]1C[C@H](CN2CCC2)N1
O=c1oc2c(O)c(O)cc3c(=O)oc4c(O)c(O)cc1c4c23
C[C@@H]1CCO[C@H]2Cn3cc(C(=O)NCc4ccc(F)cc4F)c(=O)c(O)c3C(=O)N21
O=C1CCC(c2ccccc2)Oc2c1ccc1ccccc21
 O=c1cc(-c2ccccc2)oc2c1ccc1ccccc12
CC(OO)N1c2ccc(-c3ccc(C(=O)O)cc3)cc2[C@H](Nc2ccc(Cl)cc2)C[C@@H]1C
```

```
 CC(=O)N1c2ccc(-c3ccc(C=O)cc3)cc2[C@H](Nc2ccc(Cl)cc2)C[C@@H]1C
 CC(=O)N1c2ccc(-c3ccc(C(=O)O)cc3)cc2[C@H](Nc2ccc(Cl)cc2)C[C@@H]1C
 CC(OO)N1c2ccc(-c3ccc(C=O)cc3)cc2[C@H](Nc2ccc(Cl)cc2)C[C@@H]1C
 CC(=O)N1c2ccc(-c3ccc(C=O)cc3)cc2[C@H](Nc2ccc(CO)cc2)C[C@@H]1C
O=C1c2ccccc2OCCC1c1ccccc1
 O=c1cc(-c2ccccc2)oc2ccccc12
 O=C1CCC(c2ccccc2)Oc2ccccc21
C[C@]12CC[C@@H]3c4ccc(O)cc4CC[C@H]3[C@@H]1CCC2=O
CCCCCCCNC(NS(=O)(=O)c1ccc(CCNC(=O)c2cc(Cl)ccc2OC)cc1)OO
COc1cc(C2=C(c3c[nH]c4ccccc34)CNC2=O)cc(OC)c1OC
N#Cc1cnc2c(Cl)cc(NCc3cnc[nH]3)cc2c1Nc1ccc(F)c(Cl)c1
 N#Cc1cnc2c(Cl)cc(NCC3=CC=CC=CN=N3)cc2c1Nc1ccc(F)c(Cl)c1
 CCC(N1CCCCC1)N1CCn2cc(C3=C(c4cnc5ccccn45)C(=O)NC3=O)c3cccc(c32)C1
NC1Cc2cnc1ccc(NCc1c[nH]cn1)ccc2Nc1ccc(F)cc1
COc1cccc(Nc2ncnc3ccccc23)c1
 Clc1cccc(Nc2ncnc3ccccc23)c1
 Brc1cccc(Nc2ncnc3ccccc23)c1
Cn1cnc(-c2cc(C(=O)O)ccn2)c1
 Cn1ccc(-c2cc(C(=O)O)ccn2)c1
COc1ccc(-c2c(-c3cc(C)ccn3)ncn2C)cc1
 Cn1cnc(-c2cc(-c3nnn[nH]3)ccn2)c1
Cc1nc(-c2cc(C(=O)O)ccn2)cs1
 Nc1nc(-c2cc(C(=O)O)ccn2)cs1
COc1ccc(-c2c(-c3cc(C(O)OO)ccn3)ncn2C)cc1
 COc1ccc(-c2c(-c3cc(C(=O)O)ccn3)ncn2C)cc1
Cc1cc(Cl)ccc1COc1ccnn1-c1cc(CC(O)O)ccn1
O=C(O)c1ccnc(-n2nccc2OCc2ccc(Cl)cc2)c1
CCc1cc(Nc2c(C#N)cnc3c(Cl)cc(NCC4=CN=NC=NN=N4)cc23)ccc1C
CCN(CCN(C)C)C(=O)CNCc1cccc(C(=O)OC)c1
CCN(CCN(C)C)C(=O)CNCc1cccc(C(O)OO)c1
 CCN(CCN(C)C)C(=O)CNCc1cc(C(=O)OC)ccn1
 CCN(CCN(C)C)C(=O)CNCc1cccc(C(=O)O)c1
 CCN(OCN(C)C)C(=O)CNCc1cc(C(=O)OC)ccn1
 CCN(CCN(C)C)C(=O)CNCc1cccc(C(=O)OC)c1
CN(Cc1cnc2nc(N)nc(N)c2n1)c1ccccccccoc(=O)c(C(=O)NCCCCC(=O)O)cc1
```

```
CN(Cc1cnc2nc(N)nc(N)c2n1)c1cccccccccoc(=O)c(C(=O)NCCCC=O)cc1
C=C(C)CN(CC)C(=O)CNCc1cc(C(=O)OCC)ccn1
CCOC(=O)c1ccnc(CNCC(=O)N(CC)CCC(C)C)c1
CCN(CCN(C)C)C(=O)CNCc1cccc(C(O)OO)c1
CC=C(C)CCOc1ccc(-c2cc(C(=O)NC3CCCNC3)c(NC(N)=O)s2)cc1
NC(=O)Nc1sc(-c2ccc(F)cc2)cc1C(=O)N[C@H]1CCCNC1
CCN(CC)CCOc1ccc(-c2cc(C(=O)NC3CCCNC3)c(NC(N)=O)s2)cc1
NC(=O)Nc1sc(-c2cccc(F)c2)cc1C(=O)N[C@H]1CCCNC1
NC(CO)Nc1sc(-c2cccc(F)c2)cc1C(=O)N[C@H]1CCCNC1
CC(=O)Nc1sc(-c2ccc(F)cc2)cc1C(=O)N[C@H]1CCCNC1
Cc1ccc(-c2cc(C(=O)N[C@H]3CCCNC3)c(NC(N)=O)s2)cc1
NC(=O)Nc1sc(-c2cccc(F)c2)cc1C(=O)NC1CCCCC1
NC(=O)NC1=C(C(=O)N[C@H]2CCCNC2)C=C(c2ccc(F)cc2)C1
CCN(CC)CCOc1ccc(-c2cc(C(=O)N[C@H]3CCCNC3)c(NC(N)=O)s2)cc1
CCN(CC)CCOc1cccc(-c2cc(C(=O)N[C@H]3CCCNC3)c(CC(N)=O)s2)c1
CS(=O)(=O)c1ccc(-c2cc(C(=O)N[C@H]3CCCNC3)c(NC(N)=O)s2)cc1
CCN(CC)CCOc1cccc(-c2cc(C(=O)N[C@H]3CCNNC3)c(NC(N)=O)s2)c1
CCN(CC)CCOc1cccc(-c2cc(C(=O)N[C@H]3CCCNC3)c(NC(N)=O)s2)c1
CCN(CCOc1cccc(-c2cc(C(=O)N[C@H]3CCCNC3)c(NC(N)=O)s2)c1)OC
CCN(CC)CCOc1cccc(-c2cc(C(=O)N[C@H]3CCNNC3)c(CC(N)=O)s2)c1
CCN(O)Nc1sc(-c2cccc(F)c2)cc1C(CO)N[C@H]1CCCNC1
NCN(O)Nc1sc(-c2cccc(F)c2)cc1C(=O)N[C@H]1CCCNC1
CCN(O)Nc1sc(-c2cccc(F)c2)cc1C(=O)N[C@H]1CCCNC1
NC(=O)Nc1sc(-c2cccc(F)c2)cc1C(CO)N[C@H]1CCCNC1
CC(=O)Nc1sc(-c2cccc(F)c2)cc1C(=O)N[C@H]1CCCNC1
NC(CO)Nc1sc(-c2cccc(F)c2)cc1C(=O)N[C@H]1CCCNC1
NC(=O)Nc1sc(-c2cccc(F)c2)cc1C(=O)N[C@H]1CCCNC1
O=C(N[C@H]1CCCNC1)c1ccnc2[nH]c(C3CC3)nc12
CC(=O)N[C@@H](CC(=O)NCCCCCCc1ccccc1OCCCc1ccccc1)C(=O)N[C@@H](CC(C)C)B(O)O
NC(=O)Nc1sc(-c2ccc(F)cc2)cc1C(=O)N[C@H]1CCCNC1
CS(=O)(=O)c1cccc(-c2cc(C(=O)N[C@H]3CCCNC3)c(NC(N)=O)s2)c1
CCN(CC)CCOc1ccc(-c2cc(C(=O)N[C@H]3CCCNC3)c(NC(N)=O)s2)cc1
CS(=O)(=O)c1ccc(-c2cc(C(=O)N[C@@H]3CNCCN3)c(NC(N)=O)s2)cc1
C=S(C)(=O)c1ccc(-c2cc(C(=O)N[C@H]3CCCNC3)c(NC(N)=O)s2)cc1
CC[SH](C)(=O)c1cccc(-c2cc(C(=O)N[C@H]3CCCNC3)c(NC(N)=O)s2)c1
```

```
CS[N+]([O-])(O)c1ccc(-c2cc(C(=O)N[C@H]3CCCNC3)c(NC(N)=O)s2)cc1
CS(=O)(=O)c1ccc(-c2cc(C(=O)N[C@H]3CCCNC3)c(CC(N)=O)s2)cc1
O=c1c(-c2ccc(O)cc2)coc2cc(O)cc(O)c12
O=C1C(c2ccc(O)cc2)=CCc2cc(O)cc(O)c21
O=c1cc(-c2ccccc2)oc2c1ccc1ccccc12
O=c1cc(-c2ccccc2)oc2ccc3ccccc3c12
O=C1CCC(c2ccccc2)OC2=CC=CC=CC=CC=C12
O=c1cc(-c2ccccc2)oc2c1=C=c1ccccc1=2
O=c1cc(-c2ccccc2)oc2ccccc3ccc-3c12
C[S+]([O-])c1ccc(-c2nc(-c3ccc(F)cc3)c(-c3ccncc3)[nH]2)cc1
Cc1ccc(-c2nc(-c3ccc([S+](C)[O-])cc3)[nH]c2-c2ccncc2)cc1
C[S+]([O-])c1ccc(-c2nc(-c3ccc(F)cc3)c(-c3ccncn3)[nH]2)cc1
C[S+]([O-])c1ccc(-c2nc(-c3ccc(F)cc3)c(-c3ccccc3)[nH]2)cc1
CN(Cc1cnc2nc(N)nc(N)c2n1)c1ccc(C(=O)N[C@@H](CCC(=O)O)C(=O)O)cc1
CC(=O)CC[C@H](NC(=O)c1ccc(N(C)Cc2cnc3nc(N)nc(N)c3n2)cc1)C(=O)O
CN(Cc1cnc2nc(N)nc(N)c2n1)c1ccc(C(=O)N[C@@H](CCC(N)=O)C(=O)O)cc1
CN(Cc1cnc2nc(N)nc(N)c2n1)c1ccc(C(=O)N[C]C(CCC(=O)O)C(=O)O)cc1
C=C(N[C@@H](CCC(=O)O)C(=O)O)c1ccc(N(C)Cc2cnc3nc(N)nc(N)c3n2)cc1
CCc1nc(N)nc(N)c1-c1ccc(Cl)cc1
COc1cc2ncnc(Nc3cccc(Br)c3)c2cc1OC
COc1ccc2c(Nc3cccc(Br)c3)ncnc2c1
Bc1cccc2[nH]c3c(c12)=CC(OC)=CC1=NC=31
COC1=CC2=NC2=CCNC2=CC=CC(=CC=C1)C=CC=C2
COc1ccccc(Nc2cccc(Br)c2)c2cc-2c1
Cc1ccc(NC(=O)c2ccc(CN3CCN(C)CC3)cc2)cc1Nc1nccc(-c2ccccc2)n1
Cc1ccc(NC(=O)c2ccc(CN3CCN(C)CC3)cc2)nc1Nc1nccc(-c2cccnc2)n1
Cc1ccc(NC(=O)c2ccc(CN3CCN(C)CC3)cc2)cc1Nc1nccc(-c2cccnc2)n1
Cc1ccc(NC(=O)c2ccc(CN3CCN(C)CC3)cc2)cc1Nc1cccc(-c2cccnc2)n1
Cc1ccc(NC(=O)c2ccc(CN3CCN(C)CC3)cc2)nc1Nc1cccc(-c2cccnc2)n1
Cc1nc(NC2=NC(C(=O)Nc3c(C)cccc3Cl)=[SH]2)cc(N2CCCCCCCCCC2)n1
Cc1nc(Nc2ncc(C(=O)Nc3c(C)cccc3Cl)s2)cc(N2CCN(CCS)CC2)n1
Cc1nc(Nc2ncc(C(CO)Nc3c(C)cccc3Cl)s2)cc(N2CCN(CCO)CC2)n1
CCCN1CCN(c2cc(Nc3ncc(C(=O)Nc4c(C)cccc4Cl)s3)nc(C)n2)CC1
CCCN1CCN(c2cc(Nc3ncc(C(CO)Nc4c(C)cccc4Cl)s3)nc(C)n2)CC1
Cc1nc(Nc2ncc(C(NO)Nc3c(C)cccc3Cl)s2)cc(N2CCN(CCO)CC2)n1
```

```
Cc1nc(Nc2nnc(C(=O)Nc3c(C)cccc3Cl)s2)cc(N2CCN(CCO)CC2)n1
Cc1nc(Nc2ncc(C(CO)Nc3c(C)cccc3Cl)s2)cc(N2CCCOCCCNCC2)n1
Cc1nc(Nc2ncc(C(=O)Nc3c(C)cccc3Cl)s2)cc(N2CCCOCCCNCC2)n1
Cc1nc(Nc2ncc(C(=O)Nc3c(C)cccc3Cl)s2)cc(N2CCC(CCO)CC2)n1
Cc1nc(Nc2ncc(C(=O)Nc3c(C)cccc3Cl)s2)cc(N2CCCCCCCNCC2)n1
Cc1nc(NC2=NC(C(=O)Nc3c(C)cccc3Cl)=[SH]2)cc(N2CCCCCCCNCN2)n1
CCCN1CCN(C2=CCCCc3cccc(C)c3NC(=O)C(NC3=NC=CSS3)=NC(F)=N2)CC1
Cc1nc(Nc2ncc(C(=O)Nc3c(C)cccc3Cl)s2)cc(C2CCN(CCO)CC2)n1
CCCN1CCN(c2cc(NC3=NC(C(=O)Nc4c(C)cccc4Cl)=[SH]3)nc(C)n2)CC1
Cc1nc(Nc2ncc(C(=O)Nc3c(C)cccc3Cl)s2)cc(N2CCN(CCO)OC2)n1
Cc1nc(Nc2ncc(C(CO)Nc3c(C)cccc3Cl)s2)cc(C2CCN(CCO)CC2)n1
Cc1nc(Nc2ncc(C(=O)Nc3c(C)cccc3Cl)s2)cc(C2CCC(CCO)CN2)n1
Cc1nc(Nc2ncc(C(=O)Nc3c(C)cccc3Cl)s2)cc(N2CCN(C)CC2)n1
Cc1nc(Nc2ncc(C(CO)Nc3c(C)cccc3Cl)s2)cc(N2CCC(CCO)CC2)n1
CNC(=O)C1=CC=COc2ccc(NC(=O)Nc3ccc(Cl)c(C(F)(F)F)c3)cc2C=CC=N1
CCc1ccc(NC(=O)Nc2ccc(Oc3ccnc(C(=O)NC)c3)cc2)cc1C(F)(F)F
CNC(=O)c1cccoc2ccc(NC(=O)Nc3ccc(Cl)c(C(F)(F)F)c3)ccc-2c1
Cc1cc(Nc2cc(N3CCN(C)CC3)nc(Sc3ccc(NC(=O)C4CC4)cc3)n2)n[nH]1
Cc1cc(Nc2cc(N3CCN(C)CC3)cc(Sc3ccc(NC(=O)C4CC4)cc3)n2)n[nH]1
CC1=CC(Nc2cc(N3CCN(C)CC3)nc(Sc3ccc(NC(=O)C4CC4)cc3)n2)=N[CH]1
Cc1cc(Nc2cc(N3CCN(C)CC3)nc(Nc3ccc(NC(=O)C4CC4)cc3)n2)n[nH]1
Cc1cc(Nc2cc(N3CCN(C)CC3)nc(Sc3ccc(CC(=O)C4CC4)cc3)n2)n[nH]1
N#CCOc1ccc(Nc2nc(Nc3cccc(S(N)(=O)=O)c3)ncc2Br)cc1
N#CCCc1ccc(Nc2nc(Nc3cccc(S(N)(=O)=O)c3)ncc2Br)cc1
NNCCOc1ccc(Nc2nc(Nc3cccc(S(N)(=O)=O)c3)ncc2Br)cc1
N#CCOc1ccc(Nc2cc(Nc3cccc(S(N)(=O)=O)c3)ncc2Br)cc1
CCCn1ccc(-c2cnc(N)c3c(-c4ccc(NC(=O)Nc5cccc(F)c5)cc4)csc23)c1
CCCn1cc(-c2cnc(N)c3c(-c4ccc(NC(=O)Nc5cccc(C)c5)cc4)csc23)cn1
CNC(=O)C(C)n1cc(-c2cnc(N)c3c(-c4ccc(NC(=O)Nc5ccc(O)cc5)cc4)csc23)cn1
Cc1cccc(NC(=O)Nc2ccc(-c3csc4c(C#CCN(O)OC(=O)O)cnc(N)c34)cc2)c1
CC1=CC2=CC(=CC=C1)NC(=O)Nc1ccc(cc1)-c1csc3c(cnc(N)c13)C#CCN2CCO
CCCN1CC#Cc2cnc(N)c3c(csc23)-c2ccc(cc2)NC(=O)NC2=CC=CC(C)=CCC21
Cc1cccc(NC(=O)Nc2ccc(-c3csc4c(C#CCNS(=O)(=O)O)cnc(N)c34)cc2)c1
Cc1cccc(NC(=O)Nc2ccc(-c3csc4c(C#CCN(CCO)C(C)O)cnc(N)c34)cc2)c1
C=S(C)(=O)NCC#Cc1cnc(N)c2c(-c3ccc(NC(=O)Nc4cccc(C)c4)cc3)csc12
```

```
CC1=CCOC2C(=CC=C1)NC(=O)Nc1ccc(cc1)-c1csc3c(cnc(N)c13)C#CCN2CCO
CC1=CC2C(=CC=C1)NC(=O)Nc1ccc(cc1)-c1csc3c(cnc(N)c13)C#CCN2CCO
Cc1cccc(NC(=O)Nc2ccc(-c3csc4c(C#CCN(C)[SH](C)O)cnc(N)c34)cc2)c1
CCCN(C)CC#Cc1cnc(N)c2c(-c3ccc(NC(=O)Nc4cccc(C)c4)cc3)csc12
Cc1cccc(NC(=O)Nc2ccc(-c3csc4c(C#CCN[SH](C)(=O)CO)cnc(N)c34)cc2)c1
Cc1cccc(CC(=O)Nc2ccc(-c3csc4c(C#CCN(C)C(O)C=O)cnc(N)c34)cc2)c1
CC(N)=NCC#Cc1cnc(N)c2c(-c3ccc(NC(=O)Nc4cccc(C)c4)cc3)coc12
Cc1cccc(NC(=O)Nc2ccc(-c3csc4c(C#CCN=S(C)(=O)O)cnc(N)c34)cc2)c1
Cc1cccc(CC(=O)Nc2ccc(-c3csc4c(C#CCNS(C)(=O)=O)cnc(N)c34)cc2)c1
CC1=CC(OCO)C(NC(=O)Nc2ccc(-c3csc4c(C#CCN(C)O)cnc(N)c34)cc2)=CC=C1
Cc1cccc(NC(=O)Nc2ccc(-c3csc4c(C#CCNS[SH](C)(=O)O)cnc(N)c34)cc2)c1
Cc1cccc(NC(=O)Nc2ccc(-c3csc4c(C#CCN5CCCC5)cnc(N)c34)cc2)c1
COc1ccc(NC(=O)Nc2ccc(-c3csc4c(-c5cnn(CCO)c5)cnc(N)c34)cc2)cc1
COc1ccc(NC(Nc2ccc(-c3csc4c(-c5cnn(CCO)c5)cnc(N)c34)cc2)OO)cc1
CCN1CC=C(c2cnc(N)c3c(-c4ccc(NC(=O)Nc5ccc(OC)cc5)cc4)csc23)C=N1
COC1=CC=C(NC(=O)Nc2ccc(-c3csc4c(-c5cnn(CCO)c5)cnc(N)c34)cc2)CC1
CC(O)Cn1cc(-c2cnc(N)c3c(-c4ccc(NC(=O)Nc5cccc(F)c5)cc4)csc23)cn1
CC(O)Cn1cc(-c2cnc(N)c3c(-c4ccc(NC(=O)Nc5cccc(N)c5)cc4)csc23)cn1
CC(O)Cn1ccc(-c2cnc(N)c3c(-c4ccc(NC(=O)Nc5cccc(F)c5)cc4)csc23)c1
CC(O)Cn1cc(-c2cnc(N)c3c(-c4ccc(NC(NO)Nc5cccc(F)c5)cc4)csc23)cn1
CC(O)Cn1cc(-c2cnc(N)c3c(C4=CC=C5NC(=O)NC=CC=CC=C(F)C=C54)csc23)cn1
CC(O)Cn1cc(-c2cnc(N)c3c(-c4c5c6[nH]c(=O)[nH]c-5cccc(F)cc4=6)csc23)cn1
Nc1ncnc2scc(-c3ccc(NC(=O)Nc4cc(C(F)(F)F)ccc4F)cc3)c12
Nc1nccc2scc(-c3ccc(NC(=O)Nc4cc(C(F)(F)F)ccc4F)cc3)c12
COc1cc2c(Oc3ccc4[nH]c(C)cc4c3F)ncnc2cc1OCCCN1CCCC1
CCCCCNCCCOc1cc2ncnc(Oc3ccc4[nH]c(C)cc4c3F)c2cc1OC
CCCCNCCCOc1cc2ncnc(Oc3ccc4[nH]c(C)cc4c3F)c2cc1OC
COc1cc2c(Oc3ccc4[nH]c(C)cc4c3F)ncnc2cc1OCOCN1CCCC1
COc1cc2c(Oc3ccc4[nH]c(C)cc4c3F)ncnc2cc1OCCCC1CCCC1
COc1cc2c(Oc3ccc4[nH]c(C)cc4c3F)ncnc2cc1CCCCN1CCCC1
COc1cc2c(Oc3ccc4[nH]c(C)cc4c3F)ncnc2cc1OCC1CCCCCN1
COc1cc2c(Oc3ccc4[nH]c(C)cc4c3C)ncnc2cc1OCCCN1CCCC1
CCCCCNCCCCc1cc2ncnc(Oc3ccc4[nH]c(C)cc4c3F)c2cc1OC
COc1cc2c(Oc3ccc4[nH]c(C)cc4c3F)ncnc2cc1OCCCN1CCC1
Cn1cc(-c2cnc(N)c3c(-c4ccc(NC(=O)Nc5ccc(OC(F)F)cc5)cc4)csc23)cn1
```

```
Cn1cc(-c2cnc(N)c3c(-c4ccc(NC(=O)NC5=CC(F)(CF)C=C5)cc4)csc23)cn1
CC(Oc1cc(-c2cnn(C3CCNCC3)c2)cnc1N)c1c(Cl)ccc(F)c1Cl
Cc1c(F)ccc(Cl)c1C(C)Oc1cc(-c2cnn(C3CCNCC3)c2)cnc1N
CCc1c(F)ccc(Cl)c1C(C)Oc1cc(-c2cnn(C3CCNCC3)c2)cnc1N
CC(Oc1cc(-c2cnn(C3CCNCC3)c2)cnc1N)C1=C(Cl)C=CC(F)=CC1
CC(Oc1cc(-c2cnn(C3CCNCC3)c2)cnc1N)c1c(Cl)ccc(F)c1CF
CC(Oc1cc(-c2cnn(C3CCNCC3)c2)cnc1N)c1c(Cl)ccc(F)c1F
Cc1c(F)ccc(Cl)c1C(C)Oc1cc(-c2cnn(C3CCNCN3)c2)cnc1N
CC(Oc1cc(C2=CC(C3CCNCC3)N=C2)cnc1N)C1=C(Cl)C=CC(F)=CCC1
CCc1c(F)ccc(CF)c1C(C)Oc1cc(-c2cnn(C3CCCCC3)c2)cnc1N
CC1Oc2cc(-c3cnn(C4CCNCC4)c3)cnc2Cc2c(F)ccc(Cl)c21
CCc1c(F)ccc(CF)c1C(C)Oc1cc(C2=CC(C3CCNCC3)N=C2)ccc1N
CCc1c(F)ccc(CF)c1C(C)Oc1cc(-c2cnn(C3CCNCC3)c2)cnc1N
CC(Oc1cc(C2=CC(C3CCNCC3)N=C2)ccc1N)c1c(Cl)ccc(F)c1Cl
CC(Oc1cc(-c2cnn(C3CCNCC3)c2)cnc1N)c1cc(F)ccc1Cl
CC1C=C(F)C=CC(Cl)=C1C(C)Oc1cc(-c2cnn(C3CCNCC3)c2)cnc1N
CCOc1ccc(NC(=O)Nc2ccc(-c3csc4c(-c5cnn(C)c5)cnc(N)c34)cc2)cc1
CCCc1ccc(NC(=O)Nc2ccc(-c3csc4c(-c5cnn(C)c5)cnc(N)c34)cc2)cc1
CCOC1=CC=C2C(c3cnn(C)c3)=C(N)c3c(csc32)-c2ccc(cc2)NC(=O)NC=CC=C1
O=C(Nc1cnc2[nH]cc(-c3ccccc3)c2c1)c1c(F)cccc1F
Cc1cccc(F)c1C(=O)Nc1cnc2[nH]nc(-c3ccccc3)c2c1
Cc1cccc(F)c1C(=O)Nc1cnc2[nH]cc(-c3ccccc3)c2c1
O=C(NC1=CC2=CNC(=Cc3ccccc32)N=C1)c1c(F)cccc1F
O=c1cc2ccc3[nH]cc(-c4ccccc4)c3cc(F)cccc(F)n1-2
Cc1cccc(F)c1C(=O)Nc1ccc2[nH]cc(-c3ccccc3)c2c1
S=c1[nH]nc2n1-c1n[nH]c(=S)n1-c1sc3c(c1-2)CCC3
S=C1CN=C2C3=C4CC4C=CC=C3N3C(=S)NN=N3N12
CC12C(=CC3=C1CCC3)n1c(n[nH]c1=S)-n1c2n[nH]c1=S
S=C1CN=C2c3c(sc4c3CCC4)-n3c(n[nH]c3=S)N12
S=C1NN=N2N1c1sc3c(c1-c1n[nH]c(=S)n12)CCC3
S=C1CN=C2c3c(sc4c3CCC4)N3C(=S)NN=N3N12
O=c1[nH]nc2n1-c1n[nH]c(=S)n1-c1sc3c(c1-2)CCC3
CCCC1=CC2=C3C4C(=C(F)C=CC134)c1n[nH]c(=O)n12
S=C1NN=CCN1c1n[nH]c(=S)n1-c1ccc2c(c1)CCC2
S=c1[nH]nc2n1c1c(c3n[nH]c(=S)n32)=C2CC2C=CC=1
```

```
C=C1NN=N2N1C1=CC=CC3CC3=C1c1n[nH]c(=S)n12
S=c1[nH]nn2n1c1sccc3c(c=1c1n[nH]c(=S)n12)C3
S=C1NN=N2N1C1=CC=CC3CC3=C1c1n[nH]c(=S)n12
S=c1[nH]nc2n1-c1sc3c(c1-c1nccc(S)c1-2)CCC3
S=C(NCc1ccc2c(c1)CCC2)NN=CCC1CC=NNC1=S
S=c1[nH]nn2n1c1cccc3cc3c-1c1n[nH]c(=S)n12
S=C1NN=C2c3c(sc4c3CCC4)-n3c(n[nH]c3=S)C12
Cc1cccc(NC(=O)Nc2ccc(-c3cnc4c(-c5cnn(C)c5)cnn4c3N)cc2)c1
Cc1cccc(NC(=O)Nc2ccc(-c3cnc4c(C5=CC(C)N=C5)cnn4c3N)cc2)c1
Cc1cccc(NC(=O)Nc2ccc(-c3cnc4c(-c5cnn(C)c5)ccn4c3N)cc2)c1
CC1#CC(c2cnn3c(N)c(-c4ccc(NC(=O)Nc5cccc(C)c5)cc4)cnc23)=C1
Cc1cccc(NC(=O)Nc2ccc(-c3cnc4c(-c5cnn(C)c5)cnn4c3N)cc2)n1
Cc1cccc2nn(C)c(c1)cc1cnc3c-1ncc(c3N)c1ccc(cc1)[nH]c(=O)[nH]2
Cc1cccc(NC(NO)Nc2ccc(-c3cnc4c(-c5cnn(C)c5)cnn4c3N)cc2)c1
Cc1cccc(NC(=O)Nc2ccc(-c3cnc4c(-c5cnn(C)c5)cnc-4n3N)cc2)c1
CC1=NC=C(c2cnn3c(N)c(-c4ccc(NC(=O)Nc5cccc(C)c5)cc4)cnc23)C1
CN1CCN(c2ccc3nc(-c4c(N)c5c(F)cccc5[nH]c4=O)[nH]c3c2)CC1
CN1CCN(c2ccc3nc(C4=C(N)c5c(F)cccc5NC4)[nH]c3c2)CC1
CN1CCN(c2ccc3nc(C4=C(N)C=CC(F)=CC=CC=CNC4=O)[nH]c3c2)CC1
CN1CCN(c2ccc3c(c2)NC(c2c(N)c4c(F)cccc4[nH]c2=O)C3)CC1
CN1CCN(c2ccc3nc(-c4c(N)c5c(F)cccc5[nH]c4=O)[nH]c3n2)CC1
CC1=C2C=CC(F)=C2C(N)=C1c1nc2ccc(N3CCN(C)CC3)cc2[nH]1
NC(=O)c1cccc(-c2cnc3[nH]cc(-c4ccccc4)c3c2)c1
NC(CO)c1cccc(-c2cnc3[nH]cc(-c4ccccc4)c3c2)c1
NC(=O)c1cccc(-c2ccc3[nH]cc(-c4ccccc4)c3c2)c1
NC(=O)c1cccc(-c2cnc3c(c2)C=c2ccccc2=C=CN3)c1
NC(=O)c1cccc(-c2cnc3[nH]cccc4ccccc4cc3c2)c1
Nc1ccc(-c2csc3c(C=Cc4nc5ccccc5[nH]4)cnc(N)c23)cc1
COc1ccc(C=NNc2ncnc3c2[nH]c2ccccc23)cc1OC
COc1ccc(CNNNc2ncnc3c2[nH]c2ccccc23)cc1O
COc1ccc(C=NNc2ncnc3c2[nH]c2ccccc23)cc1O
COc1ccc(C=NNc2ncnc3c2[nH]c2ccccc23)cc1
COc1ccc(C=NNc2nccc3c2[nH]c2ccccc23)cc1OC
COc1ccc(C=NNc2nccc3c2[nH]c2ccccc23)cc1O
Cn1cc(-c2nnn3c(N)c(-c4ccc(NC(=O)Nc5cccc(C(F)(F)F)c5)cc4)cnc23)cn1
```

```
Cc1cccc(NC(=O)Nc2ccc(-c3csc4c(C#CCN5CCOCC5)cnc(N)c34)cc2)c1
Cc1cccc(CC(=O)Nc2ccc(-c3csc4c(C#CCN5CCOCC5)cnc(N)c34)cc2)c1
Cc1cccc(NC(=O)Nc2ccc(-c3csc4c(C#CCN5CCCCC5)cnc(N)c34)cc2)c1
Cc1cccc(NC(=O)Nc2ccc(-c3csc4c(C#CCN5CCOCC5)cnc(O)c34)cc2)c1
CCN1CC#Cc2cnc(N)c3c(csc23)-c2ccc(cc2)NC(=O)NC2=CC=CC(C)=CC1C2
Cc1cccc(NC(=O)Nc2ccc(-c3csc4c(C#CCN5CCOCC5)cnc(N)c34)cc2)n1
Cc1cccc(NC(=O)Nc2ccc(-c3csc4c(C#CCC5CCOCC5)ccc(N)c34)cc2)n1
Cc1cccc(NC(=O)Nc2ccc(-c3csc4c(C#CCN5CCOCO5)cnc(N)c34)cc2)c1
Cc1cccc(NC(=O)Nc2ccc(-c3csc4c(C#CCN5CCOCC5)ccc(N)c34)cc2)c1
CC1=CCC2COC(=CC=C1)NC(=O)Nc1ccc(cc1)-c1csc3c(cnc(N)c13)C#CCN2C
CC1=NC=C2C(=CC=C1)NC(=O)Nc1ccc(cc1)-c1csc3c(cnc(N)c13)C#CCN2CCO
Cc1cccc(NC(=O)Nc2ccc(-c3csc4c(C#CCC5CCOCC5)cnc(N)c34)cc2)n1
Cc1cccc(CC(=O)Nc2ccc(-c3csc4c(C#CCN5CCOCC5)ccc(N)c34)cc2)c1
CCN1CC=C2C1=CC(OC)=CC=CNC(=O)Nc1ccc(cc1)-c1csc3c2cnc(N)c13
NC(=O)c1c(OCc2c(F)cc(Br)cc2F)nsc1NC(=O)NCCCCN1CCCC1
NC(=O)C1=C(c2c(F)cc(Br)cc2F)NC(=O)NCCC2CCCCN2SN=C1O
CNC(=O)c1ccccc1SC1=CC=CNN=C(C=Cc2ccccn2)C=CC=C1
CNC(=O)c1ccccc1Sc1ccc2c(c1)[CH]N=C2C=Cc1ccccn1
CNC(=O)c1ccccc1Sc1ccc2c(C=Cc3ccccc3)n[nH]c2c1
CNC(=O)c1ccncc1Sc1ccc2c(CCCc3ccccn3)n[nH]c2c1
CNC(=O)c1ccccc1Sc1ccc2[nH]c(C=Cc3ccccc3)c-2cc1
Cc1ccc(NC(=O)Nc2ccc(-c3csc4c(-c5cnn(C)c5)cnc(N)c34)cc2)cc1
COC(C(=O)N1Cc2[nH]nc(NC(=O)c3ccc(N4CCN(C)CC4)cc3)c2C1)c1ccccc1
COC(C(=O)N1Cc2[nH]nc(NC(=O)c3ccc(N4CCC(C)CC4)cc3)c2C1)c1ccccc1
COC(C(=O)N1Cc2[nH]cc(NC(=O)c3ccc(N4CCN(C)CC4)cc3)c2C1)c1ccccc1
COC(C(=O)N1Cc2[nH]nc(NC(=O)c3ccc(C4CCN(C)CC4)cc3)c2C1)c1ccccc1
CNC(=O)c1ccc(N)c2c(-c3ccc(NC(=O)Nc4cc(C)ccc4F)cc3)csc12
Nc1ncc(-c2ccoc2)c2scc(-c3ccc(NC(=O)Nc4cccc(F)c4)cc3)c12
Nc1ncc(-c2ccoc2)c2scc(C3=CC=C4C=CC(=CC=CC(F)=C3)NC(=O)N4)c12
N#Cc1ccc2nc(N)n(-c3nc(-c4ccccc4)cs3)c2c1
NCCc1ccc2nc(N)n(-c3nc(-c4ccccc4)cs3)c2c1
NNCc1ccc2nc(N)n(-c3nc(-c4ccccc4)cs3)c2c1
CCc1ccc2nc(N)n(-c3nc(-c4ccccc4)cs3)c2c1
NCCc1ccc2nc(N)n(-c3cc(-c4ccccc4)cs3)c2c1
N#Cc1ccc2nc(N)n(-c3cc(-c4ccccc4)cs3)c2c1
```

```
NCc1ccc2nc(N)n(-c3nc(-c4ccccc4)cs3)c2c1
CCCc1ccc2nc(N)n(-c3nc(-c4ccccc4)cs3)c2c1
NOCc1ccc2nc(N)n(-c3nc(-c4ccccc4)cs3)c2c1
Cc1ccccc1-c1csc(-n2c(N)nc3ccccc32)n1
Nc1nc2ccc(C3NO3)cc2n1-c1nc(-c2ccccc2)cs1
N#Cc1ccc2nc(N)n(-c3cc(-c4ccncc4)cs3)c2c1
CN1CCN(C2CCC(n3cc(-c4ccc(Oc5ccccc5)cc4)c4c(N)ncnc43)CC2)CC1
CN1CCN(C2CCCC(n3cc(-c4ccc(Oc5ccccc5)cc4)c4c(N)ncnc43)C2)CC1
Nc1ncc(-c2cnn(CCO)c2)c2scc(-c3ccc(NC(=O)Nc4cccc(F)c4)cc3)c12
Nc1ncc(-c2cnn(CCO)c2)c2scc(-c3ccc(NC(=O)Nc4cccc(O)c4)cc3)c12
Cc1cc2cc(-c3csc4c(-c5cccc(S(C)(=O)=O)c5)cnc(N)c34)ccc2[nH]1
Cc1cc2cc(-c3csc4c(-c5cccc([SH](=O)=O)c5)cnc(N)c34)ccc2[nH]1
Cc1cc2cc(-c3csc4c(-c5cccc(S(C)(=O)=O)c5)ccc(N)c34)ccc2[nH]1
CNS(=O)(=O)c1ccccc1Nc1nc(Nc2cc(OC)c(OC)c(OC)c2)ncc1C
CNS(=O)(=O)c1ccccc1Nc1nc(Nc2cc(OC)c(OC)c(OC)c2)ncc1Cl
CNS(=O)(=O)c1ccccc1Nc1ccnc(Nc2cc(OC)c(OC)c(OC)c2)n1
CNS(=O)(=O)c1ccccc1Nc1nc(Nc2cc(OC)c(OC)c(OC)c2)ccc1C
CNS(=O)(=O)c1ccccc1Nc1nc(Nc2cc(OC)c(OC)c(OC)c2)ncc1CN
CCc1cnc(Nc2cc(OC)c(OC)c(OC)c2)nc1Nc1ccccc1S(=O)(=O)NC
CNS(=O)(=O)c1ccccc1Nc1nc(Nc2cc(OC)c(OC)c(OC)c2)ccc1Cl
CN[SH](=O)(CO)c1ccccc1Nc1nc(Nc2cc(OC)c(OC)c(OC)c2)ncc1C
CNS(=O)(=O)c1ccccc1Nc1nc(Nc2cc(OC)c(OC)c(OO)c2)ncc1Cl
CNS(=O)(=O)c1ccccc1Nc1nc(Nc2cc(OC)c(OC)c(OC)n2)ncc1Cl
CNS(=O)(=O)c1ccccc1NC1=C(C)C=NC2=C(OC)C(OC)=CC=CNC(=N1)C=CCO2
C=C1Nc2cnc(C#N)c(n2)OCCCCCOc2cc(NS(C)(=O)=O)c(Cl)cc2N1
CS(=O)(=O)Nc1cc2c(cc1Cl)NC(=O)Nc1ccc(C#N)c(n1)OCCCCCO2
Cn1cccccc2nc1OCCCCCOc1cc(NS(C)(=O)=O)c(Cl)cc1CC(=O)N2
CS(=O)(=O)Nc1cc2c(cc1Cl)NC(=O)Nc1cnc(C#N)c(n1)CCCCCCO2
N#Cc1ncc2nc1OCCCCCOc1cc(CCCO)c(Cl)cc1NC(=O)N2
NCCc1ncc2nc1OCCCCCOc1cc(CCCO)c(Cl)cc1NC(=O)N2
O=C1NC2=CN=CNCNC(=N2)OCCCCCOc2cc(CCCO)c(Cl)cc2N1
CN1CCN(c2nc(C3=C(c4c[nH]c5ccccc45)C(=O)NC3=O)c3ccccc3n2)CC1
CN1CCN(c2nc(C3=C(C4=CNc5cccc4c5)C(=O)NC3=O)c3ccccc3n2)CC1
CN1CCN(c2cc3ccccc3c(C3=C(c4c[nH]c5ccccc45)C(=O)NC3=O)n2)CC1
CN1CCN(C2=NC(C3=C(c4c[nH]c5ccccc45)C(=O)NC3=O)=CC=CC=CC=CC=N2)CC1
```

```
CN1CCN(C2=NC(C3=C(C4=CNc5cccc4c5)C(=O)NC3=O)=CC=CC=CC=CN=N2)CC1
Cc1nc(N)sc1-c1ccnc(Nc2ccc(N3CCOCC3)cc2)n1
Cc1cc(N)sc1-c1ccnc(Nc2ccc(N3CCOCC3)cc2)n1
Cc1nc(N)sc1-c1ccnc(Nc2ccc(C3CCOCC3)cc2)n1
Cc1nc(N)sc1C1=CC=NC=CC=C2C=CC(=C2NCN2CCOCC2)C1
Cc1nc(N)sc1-c1ccnc(Nc2ccc(N3CCNCC3)cc2)n1
Cc1cc(N)sc1-c1ccnc(Nc2ccc(N3CCNCC3)cc2)n1
Cc1nc(N)sc1-c1ccnc(Nc2ccc(N3CC=CC3)cc2)n1
Cc1nc(N)sc1-c1ncnc(Nc2ccc(N3CCOCC3)cc2)n1
Cc1nc(N)sc1-c1ccnc(Nc2ccc(N3CCOOC3)cc2)n1
Cc1nc(N)sc1-c1ccnc(Nc2ccc(N3CCOCN3)cc2)n1
Cc1nc(N)sc1-c1ccnc(Nc2ccc(N3CCOCO3)cc2)n1
Cc1cc(N)sc1-c1ccnc(Nc2ccc(N3CCNCN3)cc2)n1
Cc1cc(N)sc1-c1ccnc(Nc2ccc(C3CCNCC3)cc2)n1
O=C1NCCc2[nH]c(-c3ccncc3)cc21
O=C1NCCc2[nH]c(-c3ccnnc3)cc21
O=C1NCCc2[nH]c(-c3ccccc3)cc21
O=CNNC1c2cc(-c3ccncc3)cn21
O=C1NCCn2cc(-c3ccncc3)cc21
O=C1NCCc2[nH]c(-c3cccnc3)cc21
COc1cc2ncnc(Nc3ccc(F)c(Cl)c3)c2cc1OCCCC1CCOCC1
COc1cc2ncnc(Nc3ccc(F)c(Cl)c3)c2cc1OCCCN1CCOCC1
COc1cc2ncnc(Nc3cnc(F)c(Cl)c3)c2cc1OCCCN1CCOCC1
COc1cc2ncnc(Nc3ccc(F)c(Cl)c3)c2cc1OCCCC1CCCCC1
COc1cc2ncnc(Nc3ccc(F)c(Cl)c3)c2cc1OCCCN1CCOCN1
COc1cc2ncnc(Nc3ccc(F)c(Cl)c3)c2cc1OCCCN1CCOCO1
CCOCCCOCCCOc1cc2c(Nc3ccc(F)c(Cl)c3)ncnc2cc1OC
CCCOCCCOCCCOc1cc2c(Nc3ccc(F)c(Cl)c3)ncnc2cc1OC
COc1cc2ncnc(Nc3ccc(F)c(Cl)c3)c2cc1OCCCN1CCCCC1
COc1cc2ncnc(Nc3ccc(F)c(Cl)c3)c2cc1OCCCC1CCOOC1
COc1cc2ncnc(Nc3ccc(F)c(Cl)c3)c2cc1OOCCN1CCOCC1
C=S(C)(=O)CCNCc1ccc(-c2ccc3ncnc(Nc4ccc(OCc5cccc(F)c5)c(Cl)c4)c3c2)o1
CS(=O)(=O)CCNCc1ccc(-c2ccc3ccnc(Nc4ccc(OCc5cccc(F)c5)c(Cl)c4)c3c2)o1
C=S(C)(=O)CCNCc1ccc(-c2ccc3ccnc(Nc4ccc(OCc5cccc(F)c5)c(Cl)c4)c3c2)o1
CNc1cncc(-c2c[nH]c(=O)c(NC(=O)c3ccc(N4CCC[C@H]4CN4CCCC4)cc3)n2)n1
```

```
CNc1cncc(-c2c[nH]c(=O)c(NC(=O)c3ccc(N4CCC[C@H]4CN4CCCC4)cc3)c2)n1
CNc1cccc(-c2c[nH]c(=O)c(NC(=O)c3ccc(N4CCC[C@H]4CN4CCCC4)cc3)c2)n1
CNc1cncc(-c2c[nH]c(=O)c(NC(=O)c3ccc(N4CCC[C@H]4CC4CCCC4)cc3)c2)n1
CCCCCCCC[C@@H]1CCCN1c1ccc(C(=O)Nc2cc(-c3cncc(NC)n3)c[nH]c2=O)cc1
CN(C)CCN1CCN(C(=O)c2cc(C(C)(C)C)sc2NC(=O)Nc2cccc(Cl)c2Cl)CCC1=O
Cc1ccc(-n2nc(C(C)(C)C)cc2NC(=O)Nc2ccc(OCCN3CCOCC3)c3ccccc23)cc1
Cc1ccc(-n2nc(C(C)C)cc2NC(=O)Nc2ccc(OCCN3CCOCC3)c3ccccc23)cc1
Cc1ccc(-n2nc(C(C)(C)C)cc2NC(=O)Nc2ccc(OCNN3CCOCC3)c3ccccc23)cc1
Cc1ccc(F)c(NC(=O)Nc2ccc(-c3cccc4c3=C(N)N=CN=NC=4)cc2)c1
Cc1ccc(F)c(NC(=O)Nc2ccc(-c3ccc4n[nH]nc(N)c3-4)cc2)c1
Cc1ccc(F)c(NC(=O)Nc2ccc(-c3cccc4c3=C(N)C=CN=NC=4)cc2)c1
Cc1ccc(F)c(NC(=O)Nc2ccc(-c3cccc4c3=C(N)N=CN=CC=4)cc2)c1
Cc1ccc(F)c(NC(=O)Nc2ccc(-c3cccccnc4c3=C(N)N=4)cc2)n1
Cc1ccc(F)c(NC(=O)Nc2ccc(C3=C4C(=NC=NC4(N)N)C=C3)cc2)c1
Cc1ccc(F)c2cccc3ccc([nH]c(=O)[nH]2)=c3cccc2[nH]c(c1)=NN2
Cc1ccc(F)c(NC(=O)Nc2ccc(-c3ccc4nc3N(N)NN4N)cc2)c1
Nc1n[nH]c2cccc(-c3ccc(NC(=O)Nc4cc(F)ccc4F)cc3)c12
Clc1ccc(Nc2nnc(Cc3ccnnc3)c3ccccc23)cc1
ClC1=CC=CC=Cc2nnc(Cc3ccncc3)c3ccccc2cccc31
OC(=Nc1cccc(Cl)c1)Nc1ncc(NCN=c2[nH]cnc3ccsc23)s1
OC(=Nc1cccc(Cl)c1)Nc1ccc(NCN=c2[nH]cnc3ccsc23)s1
OC(=Nc1cccc(Cl)c1)Nc1ncc(CCN=C2NCNNc3ccsc32)s1
OC(=Nc1cccc(Cl)c1)Nc1ncc(CCNCC2NC=Nc3ccsc32)s1
OC(=Nc1cccc(Cl)c1)Nc1ccc(CCN=C2NCNNc3ccsc32)s1
OC(=Nc1cccc(Cl)c1)Nc1ncc(CCN=CCNC2C=CSC=CC=CN2)s1
OC(=Nc1cccc(Cl)c1)Nc1ccc(CCN=c2[nH]cnc3ccsc23)s1
IC1=CC=CC=CC=CC=CC2=CC2=CNC=CC=CC=CC=C1
Nc1noc2cccc(-c3cccc(O)c3)c12
CC1=CC=C1C1=NN=C(C)C=NC=CC=CC=CC=C1
Nc1coc2cccc(-c3cccccocc3)c12
Nc1ccc2cccccc3ccc(c1-2)C=CC=C3
CC1=CN=c2ccc1ccccc1ccccc2=CC=CC=C1
Nc1noc2cccc(-c3cccccocc3)c12
CN1CCC(c2c(O)cc(O)c3c(=O)cc(-c4ccccc4Cl)oc23)C(O)C1
CCc1ccccc1-c1cc(=O)c2c(O)cc(O)c(C3CCN(C)CC3O)c2o1
```

```
CNC(=O)c1ccccc1Nc1nc(Nc2ccc(N3CCOCC3)cc2OC)ncc1Cl
CNC(=O)c1ccccc1Nc1nc(Nc2ccc(N3CCOCC3)cc2OC)ccc1Cl
CNC(=O)c1ccccc1Nc1nc(Nc2ccc(N3COOCO3)cc2OC)ncc1C
CNC(=O)c1ccccc1Nc1ccnc(Nc2ccc(N3CCOCC3)cc2OC)n1
CCc1cnc(Nc2ccc(N3CCOCC3)cc2OC)nc1Nc1ccccc1C(=O)NC
CNC(=O)c1ccccc1Nc1cnnc(Nc2ccc(N3CCOCC3)cc2OC)c1
CNC(=O)c1ccccc1Nc1nc(Nc2ccc(N3CCOCC3)cc2OC)ncc1CCF
CNC(=O)c1ccccc1Nc1nc(Nc2ccc(N3CCOCC3)cc2OC)ncc1C
CNC(=O)c1ccccc1Nc1nc(Nc2ccc(N3CCOCO3)cc2OC)ccc1Cl
CNC(=O)c1ccccc1Nc1nc(Nc2ccc(N3CCOCC3)cc2OC)nnc1Cl
CNC(=O)c1ccccc1Nc1ccnc(Nc2ccc(N3CCOCO3)cc2OC)n1
CNC(=O)c1ccccc1Nc1nc(Nc2ccc(N3CCOCO3)cc2OC)ncc1C
CNC(=O)c1ccccc1Nc1nc(Nc2ccc(N3CCOCO3)cc2OC)ncc1Cl
CNC(=O)c1ccccc1Nc1nc(Nc2ccc(N3CCOCC3)cc2OC)ccc1C
CNC(=O)c1ccccc1Nc1nc(Nc2ccc(N3CCCCC3)cc2OC)ncc1Cl
CNC(=O)c1ccccc1Nc1cccc(Nc2ccc(N3COOCO3)cc2OC)n1
CNC(=O)c1ccccc1Nc1cccc(Nc2ccc(N3CCOCC3)cc2OC)n1
CCc1ccc(Nc2ccc(N3CCOCC3)cc2OC)nc1Nc1ccccc1C(=O)NC
CCCc1cnc(Nc2ccc(N3CCOCC3)cc2OC)nc1Nc1ccccc1C(=O)NC
CC(=O)c1cccc(-c2cnc3[nH]ccc3c2)c1
CC(=O)c1cccc(-c2cnc3[nH]cnc3c2)c1
CN(c1ncccc1CNc1nc(Nc2ccc3c(c2)CC(=O)N3)ncc1C(F)(F)F)[SH](C)(=O)O
CN(c1ncccc1CNc1nc(Nc2ccc3[nH]c(=O)[nH]c3c2)ncc1C(F)(F)F)[SH](C)(=O)OO
CCS(=O)(=O)N(C)c1ncccc1CNc1nc(Nc2ccc3c(c2)CC(=O)N3)ncc1C(F)(F)F
C=S(C)(=O)N(C)c1ncccc1CNc1nc(Nc2ccc3c(c2)CC(=O)N3)ncc1C(F)(F)F
COOS(=O)N(C)c1ncccc1CNc1nc(Nc2ccc3c(c2)CC(=O)N3)ncc1C(F)(F)F
CN(c1ncccc1CNc1nc(Nc2ccc3c(c2)CC(=O)N3)ncc1C(F)(F)F)S(=O)(=O)CO
CC[SH](=O)(O)N(C)c1ncccc1CNc1nc(Nc2ccc3c(c2)CC(=O)N3)ncc1C(F)(F)F
CN(S[SH](C)(=O)O)c1ncccc1CNc1nc(Nc2ccc3c(c2)CC(=O)N3)ncc1C(F)(F)F
NC(Cc1c[nH]c2ccccc12)C(=O)Nc1cncc(C=Cc2ccncc2)c1
NC(Cc1c[nH]c2cccnc12)C(=O)Nc1cncc(C=Cc2ccncc2)c1
NC(Cc1c[nH]c2cccnc12)C(=O)Nc1cncc(C=Cc2ccccc2)c1
NC(Cc1c[nH]c2ccccc12)C(=O)Nc1cccc(C=Cc2ccccc2)c1
NC(Cc1c[nH]c2ccccc12)C(=O)NC1=CN=CC2=CC(=CC=NC=C1)C=C2
Cc1nccn2c(-c3ccnc(NCC(C)(C)O)n3)c(-c3ccc(F)cc3F)nc12
```

```
Cc1nccn2c(-c3ccnc(NCC(C)(C)CO)n3)c(-c3ccc(F)cc3)nc12
Cc1nccn2c(-c3ccnc(CCC(C)(C)O)n3)c(-c3ccc(F)cc3F)nc12
Cc1nccn2c(-c3ccnc(NCC(C)(C)CO)n3)c(-c3ccc(F)cc3F)nc12
Cc1nccn2c(-c3ccnc(NCC(C)(C)O)n3)c(-c3ccc(F)cc3)nc12
CCc1c(-c2ccc(CCOC)cc2)[nH]c2nccnc12
CCc1c(-c2ccc(C(C)(C)N)cc2)[nH]c2nccnc12
CCc1c(C2=C=C(C(C)(C)O)C=C2)[nH]c2nccnc12
CCc1c(-c2ccc(C(C)C)cc2)[nH]c2nccnc12
CCc1c(-c2ccc(C(C)(C)O)cc2)[nH]c2ncccc12
CCc1c(-c2ccc(C(C)CO)cc2)[nH]c2nccnc12
COc1cc(Nc2ncc(F)c(Nc3ccc4c(n3)NC(=O)C(C)(C)O4)n2)cc(OC)c1O
COc1cc(Nc2ncc(F)c(Nc3ccc4c(n3)NC(=O)C(C)(C)O4)n2)cc(OC)c1OC
COc1cc(Nc2ncc(F)c(Nc3ccc4c(n3)NC(=O)C(C)(C)O4)n2)cc(OC)c1
CCc1cc(Nc2ncc(F)c(Nc3ccc4c(n3)NC(=O)C(C)(C)O4)n2)cc(OC)c1
COc1cc(Nc2ncc(F)c(Nc3ccc4c(n3)NC(=O)C(C)(C)O4)n2)cc(OO)c1OC
COc1cc(Nc2ncc(F)c(Nc3ccc4c(n3)NC(=O)C(C)(O)O4)n2)cc(OC)c1OC
COc1cc(Nc2ncc(F)c(Nc3ccc4c(n3)NC(=O)C(C)(C)O4)n2)cc(OC)c1C
NC(=O)c1ccc(-c2c[nH]c3nccc(Cl)c23)cc1
CCc1ccnc2[nH]cc(-c3ccc(C(N)=O)cc3)c12
NC(=O)c1ccc(-c2c[nH]c3nccc(Cl)c23)nc1
NC(=O)c1ccc(C2=CNC=CN=CC=C(Cl)C=C2)cc1
O=C1Cc2c([nH]c3ccc([C+](=O)[O-])cc23)-c2ccccc2N1
O=C1CC2=C(Nc3ccc([N+](=O)[O-])c(c3)C=C2)c2ccccc2N1
O=[N+]([O-])c1ccc2[nH]c3c(c2c1)CC(OO)Nc1ccccc1-3
O=C1CC2=Cc3cc([N+](=O)[O-])ccc3NC2=CC=C2C=C2N1
NCC(NC(=O)c1ccc(-c2ccncc2)cc1)c1ccccc1
NNC(NC(=O)c1ccc(-c2ccncc2)cc1)c1ccccc1
O=C(NC(CO)c1ccccc1)c1ccc(-c2ccncc2)cc1
NCC(NC(=O)c1ccc(-c2ccnnc2)cc1)c1ccccc1
CCC(NC(=O)c1ccc(-c2ccnnc2)cc1)c1ccccc1
NCC1C=CC=CC=CC=C2C=CC(=C2)c2ccc(cc2)C(=O)N1
NCC(NC(=O)c1cccc(-c2ccncc2)c1)c1ccccc1
NNC(CC(=O)c1ccc(-c2ccncc2)cc1)c1ccccc1
O=C(NC(=O)c1ccc(-c2ccncc2)cc1)c1ccccc1
CNC(CC(=O)c1ccc(-c2ccncc2)cc1)c1ccccc1
```

```
Cc1nncn2c(-c3ccnc(NCC(C)(C)CO)n3)c(-c3ccc(F)cc3F)nc12
Cc1nncn2c(-c3ccnc(NCC(C)(C)CO)n3)c(-c3ccc(F)cc3)nc12
Cc1nccn2c(-c3ccnc(NCC(C)(C)C)n3)c(-c3ccc(F)cc3F)nc12
COc1nccn2c(-c3ccnc(NCC(C)(C)CO)n3)c(-c3ccc(F)cc3F)nc12
Cc1nccn2c(-c3ccnc(NCC(C)(C)OO)n3)c(-c3ccc(F)cc3F)nc12
CC1CCN(C(=O)CC#N)CC1N(C)c1ncnc2[nH]ccc12
CC(N)C1CCC(C(=O)Nc2ccncc2)CC1
CC(C)C1CCC(Nc2ccncc2)CC1CCC=O
CC(N)C1CC=C(C(=O)Nc2ccncc2)CC1
CCCCNC1CCC(C(=O)Nc2ccncc2)CC1
CC(N)C1CCC(C(=O)Nc2ccnnc2)CC1
CC(=CC1C=CCCCC1)C(=O)Nc1ccncc1
COc1cc(-c2ccc3c(c2)NC(=O)c2ccc(-c4ccccc4NS(C)(=O)=O)cc2N3)cc(OC)c1OC
COc1cc(-c2ccc3c(c2)NC(=O)c2ccc(-c4ccccc4N[SH](=O)=O)cc2N3)cc(OC)c1OC
CC(C)(C)NS(=O)(=O)C1=Cc2nc(N)nn2C=CC2=CC2=CC=C1
CC(C)(C)NS(=O)(=O)c1cccc(-c2ccn3nc(N)nc3c2)c1
CC(C)(C)NS(=O)(=O)c1cccc(-c2ccn3cc(N)nc3c2)c1
CC(C)(C)NS(=O)(=O)c1cncc(-c2ccn3cc(N)nc3c2)c1
C#Cc1cccc(Nc2ncnc3cc(OCCOC)c(OCC)cc23)c1
C#Cc1cccc(Nc2ncnc3cc(CCCOC)c(OCCCC)cc23)c1
C#Cc1cccc(Nc2ncnc3cc(OCCOC)c(OCCOC)cc23)c1
C#Cc1cccc(Nc2ncnc3cc(OCCOC)c(OCCO)cc23)c1
C#Cc1cccc(Nc2ncnc3cc(OCCOC)c(OOCCC)cc23)c1
C#Cc1cccc(Nc2ncnc3cc(OCCOC)c(CCCCCC)cc23)c1
C#Cc1cccc(Nc2ncnc3cc(CCCOC)c(OCCOC)cc23)c1
C#Cc1cccc(Nc2cc3c(OCCOC)c(OCCOC)cc-3ncn2)c1
C#Cc1cccc(Nc2ncnc3cc(CCCOC)c(OCCO)cc23)c1
Cn1c(=O)c(S(=O)(=O)c2ccc(F)cc2F)cc2cnc(Nc3ccc4[nH]ccc4c3)nc21
Nc1ncnc2c1c(-c1cccc(OCc3ccccc3)c1)cn2[C@H]1C[C@@H](CN2CCC2)C1
Nc1c(OCc2ccccc2)cccc1-c1cn([C@H]2C[C@@H](CN3CCC3)C2)c2ncnc(N)c12
Nc1ncnc2c1c(C1=CC=CC(O)=C3C=CCC=C3C=C1)cn2[C@H]1C[C@@H](CN2CCC2)C1
Nc1ncnc2c1c(-c1cccc(OCc3ccccc3)c1)cn2[C@@H]1C[C@H](CN2CCC2)O1
Nc1ncnc2c1c(C1=CC=CC=COCc3ccccc3C=C1)cn2[C@H]1C[C@@H](CN2CCC2)C1
Nc1ncnc2c1c(-c1cccc(O)c1)cn2[C@H]1C[C@@H](CN2CCC2)C1
CC(=O)N1CCN([C@H]2C[C@@H](n3cc(-c4cc(OC[C@@H]5CCCO5)ccc4F)c4c(N)ncnc43)C2)CC1
```

CC(=O)N1CCN([C@H]2C[C@@H](n3cc(-c4cc(OC[C@@H]5CCCO5)ccc4F)c4c(N)ccnc43)C2)CC1
CCC[C@H]1COC12C=CC=C(F)C(c1cn([C@H]3C[C@@H](N4CCN(C(C)=O)CC4)C3)c3ncnc(N)c13)=C2
CC(=O)N1CCN([C@H]2C[C@@H](n3cc(-c4cc(OCC5CCCC5)ccc4F)c4c(N)ncnc43)C2)CC1
CC(=O)N1CCN([C@H]2C[C@@H](n3nc(-c4cc(OC[C@@H]5CCCO5)ccc4F)c4c(N)ncnc43)C2)CC1
CC(=O)N1CCN([C@@H]2C[C@H](n3cc(-c4cc(OC[C@@H]5CCCO5)ccc4F)c4c(N)ncnc43)N2)CC1
Brc1cccc(Nc2ncnc3ccccc23)c1
N#Cc1cnc2ccc(NCc3c[nH]nn3)cc2c1Nc1ccc(F)c(Cl)c1
N#Cc1cnc2ccc(NCc3c[nH]cn3)cc2c1Nc1ccc(F)c(Cl)c1
N#Cc1cnc2ccc(NCc3c[nH]cn3)cc2c1Nc1ccc(F)cc1
Fc1ccc(-c2ncn(CCN3CCOCC3)c2-c2ccc3[nH]ncc3c2)cc1
Fc1ccc(-c2ccn(CCN3CCOCC3)c2-c2ccc3[nH]ncc3c2)cc1
Cc1cc(N2CCOCC2)cc2[nH]c(-c3c(NCC(O)c4cccc(Cl)c4)cc[nH]c3=O)nc12
CCc1cccc(C(O)CNc2cc[nH]c(=O)c2-c2nc3c(C)cc(N4CCOCC4)cc3[nH]2)c1
Cc1cc(N2CCCCC2)cc2[nH]c(-c3c(NCC(O)c4cccc(Cl)c4)cc[nH]c3=O)nc12
CCOCCCNc1cc(C)c2nc(-c3c(NCC(O)c4cccc(Cl)c4)cc[nH]c3=O)[nH]c2c1
CCc1cccc(C(O)CNc2cc[nH]c(=O)c2-c2nc3c(C)cc(N4CCCC4)cc3[nH]2)c1
CCc1cccc(C(O)CNC2=C(c3nc4c(C)cc(N5CCOCC5)cc4[nH]3)C(=O)NCCC2)c1
Cc1cc(N2CCOCC2C)cc2[nH]c(-c3c(NCC(O)c4cccc(Cl)c4)cc[nH]c3=O)nc12
CCCCCCNc1cc(C)c2nc(-c3c(NCC(O)c4cccc(CC)c4)cc[nH]c3=O)[nH]c2c1
CCc1cccc(C(O)CNC2=C3c4nc5c(C)cc(cc5[nH]4)NCCCOCC(C)[N+]3([O-])C=C2)c1
CCc1cccc(C(O)CNC2=C(c3nc4c(C)cc(N5CCCCC5)cc4[nH]3)C(=O)CCCC2)c1
CCOCCCNc1cc(C)c2nc(-c3c(NCC(O)c4cccc(CC)c4)cc[nH]c3=O)[nH]c2c1
CCc1cccc(C(O)CNC2=C(c3nc4c(C)cc(N5CCCCC5)cc4[nH]3)C(=O)NCCC2)c1
CCc1cccc(C(O)CNc2cc[nH]c(=O)c2-c2nc3c(C)cc(N4CCCCC4)cc3[nH]2)c1
CCc1cccc(C(C)CNC2=C(c3nc4c(C)cc(N5CCCCC5)cc4[nH]3)C(=O)CC=C2)c1
CCc1cccc(C(O)CNc2cc[nH]c(=O)c2-c2nc3c(C)cc(N4CCOC4)cc3[nH]2)c1
Cc1cc(N2CCOC2)cc2[nH]c(-c3c(NCC(O)c4cccc(Cl)c4)cc[nH]c3=O)nc12
CCc1cccc(C(O)CNC2=C(c3nc4c(C)cc(N5CCCCC5)cc4[nH]3)C(=O)CC=C2)c1
Cc1cc(N2CCCC2)cc2[nH]c(-c3c(NCC(O)c4cccc(Cl)c4)cc[nH]c3=O)nc12
CCOCCCNc1cc(C)c2nc(C3=C(NCC(O)c4cccc(CC)c4)C=CNC3NO)[nH]c2c1
CCc1cccc(C(O)CNC2=C(c3nc4c(C)cc(N5CCOCC5)cc4[nH]3)C(=O)CC=C2)c1
Cc1cc(-c2ccccc2)n(-c2cc(NN=Cc3ccco3)ncn2)n1
Cc1cc(-c2ccccc2)n(-c2cc(NN=Cc3ccco3)ccn2)n1
Cc1cc(-c2ccccc2)n(-c2ccnc(NN=Cc3ccco3)c2)n1
O=C(Cc1ccccc1)Nc1cccc(-c2nc3sccn3c2-c2ccnc(Nc3cccc(N4CCOCC4)c3)n2)c1

```
O=C(Cc1ccccc1)Nc1cccc(-c2nc3sccn3c2-c2ccnc(Nc3ccc(N4CCOCC4)cc3)n2)c1
O=C(Cc1ccccc1)Nc1cccc(-c2nc3sccn3c2-c2ccnc(Nc3cccc(C4CCOCC4)c3)n2)c1
O=C(Cc1ccccc1)Nc1cccc(-c2nc3sccn3c2-c2ccnc(Nc3cccc(C4CCCCC4)c3)n2)c1
CCN1CCN(c2ccc(Nc3nccc(-c4c(-c5cccc(C(=O)Nc6ccccc6F)c5)nc5ccccn45)n3)cc2)CC1
CCN1CCN(c2ccc(Nc3nccc(-c4c(-c5cccc(C(CO)Nc6ccccc6F)c5)nc5ccccn45)n3)cc2)CC1
CON1CCN(c2ccc(Nc3nccc(-c4c(-c5cccc(C(=O)Nc6ccccc6F)c5)nc5ccccn45)n3)cc2)CC1
CCN1CCN(c2ccc(Nc3nccc(-c4c(-c5cccc(C(=O)Nc6ccccc6C)c5)nc5ccccn45)n3)cc2)CC1
CCN1CCN(c2ccc(Nc3nccc(-c4c(-c5cccc(C(Nc6ccccc6F)OO)c5)nc5ccccn45)n3)cc2)CC1
CCN1CCN(c2ccc(Nc3ncnc(-c4c(-c5cccc(C(=O)Nc6ccccc6F)c5)nc5ccccn45)n3)cc2)CC1
CCC1CCN(c2ccc(Nc3nccc(-c4c(-c5cccc(C(=O)Nc6ccccc6F)c5)nc5ccccn45)n3)cc2)CC1
COCCn1cc(-c2ccncn2)c(-c2ccc(Cl)cc2)n1
Clc1ccc(-c2n[nH]cc2-c2ccccn2)cc1Cl
CC1=CC=CC=CC=C1c1nc2c(ccc3ccc1cc3)=CN=2
Cn1cc(-c2ccncn2)c(-c2ccc(F)cc2)n1
c1ccc(COc2ccc(Nc3ncnc4ccccc34)cc2)cc1
Clc1ccc(-c2n[nH]cc2-c2ccncn2)cc1Cl
Clc1ccc(-c2n[nH]cc2-c2ccncc2)cc1Cl
CNS(=O)(=O)c1ccccc1Nc1nc(Nc2cc(N3CCN(C(C)=O)CC3)ccc2OC)ncc1Br
CNS(=O)(=O)c1ccccc1Nc1nc(Nc2cc(C3CCN(C(C)=O)CC3)ccc2OC)ncc1Br
Bc1cnc(Nc2cc(C3CCN(C(C)=O)CC3)ccc2OC)nc1Nc1ccccc1S(=O)(=O)NC
Bc1cnc(Nc2cc(N3CCN(C(C)=O)CC3)ccc2OC)nc1Nc1ccccc1S(=O)(=O)NC
CNS(=O)(=O)c1ccccc1Nc1nc(Nc2cc(N3CCN(C(N)=O)CC3)ccc2OC)ncc1Br
CNS(=O)(=O)c1ccccc1Nc1ccnc(Nc2cc(N3CCN(C(C)=O)CC3)ccc2OC)n1
BBc1cnc(Nc2cc(N3CCN(C(C)=O)CC3)ccc2OC)nc1Nc1ccccc1S(=O)(=O)NC
CNc1ccc(N2CCN(C(C)=O)CC2)cc1Nc1ncc(Br)c(Nc2ccccc2S(=O)(=O)NC)n1
CNS(=O)(=O)c1ccccc1Nc1nc(Nc2cc(N3CCN(C(=O)O)CC3)ccc2OC)ncc1Br
CNS(=O)(=O)c1ccccc1Nc1nc(Nc2cc(C3CCN(C(C)=O)CC3)ccc2OC)ncc1C
CNS(=O)(=O)c1ccccc1Nc1nc(Nc2cc(N3CCN(C(C)=O)CC3)ccc2NO)ncc1Br
CNS(=O)(=O)c1ccccc1Nc1nc(Nc2cc(C3CCN(C(=O)O)CC3)ccc2OC)ncc1Br
CNS(=O)(=O)c1ccccc1Nc1nc(Nc2cc(N3CCN(C(C)=O)CC3)ccc2OO)ncc1Br
CC(C)Cn1c(N)nc2c(F)cc(C3=C(c4ccc(F)cc4)N=S4SC=CN34)cc21
CC(C)Cn1c(N)cc2c(F)cc(-c3c(-c4ccc(F)cc4)nc4sccn34)cc21
CC(C)Cn1c(N)nc2c(F)cc(-c3c(-c4ccc(F)cc4)n4ccccn34)cc21
CCCN1c2c(CCO)cccc2NC12N=CC=C2c1c(-c2cccc(NC(=O)Cc3ccccc3)c2)nc2sccn12
O=C(Cc1ccccc1)Nc1cccc(-c2nc3sccn3c2-c2ccnc(Nc3ccc(C4CCOCC4)cc3)n2)c1
```

```
CCCN1c2c(CCN)cccc2NC12N=CC=C2c1c(-c2cccc(NC(=O)Cc3ccccc3)c2)nc2sccn12
Cc1cccc(NC(=O)Nc2ccc(-c3csc4ncnc(N)c34)cc2)c1C
Cc1cccc(NC(=O)Nc2ccc(C3=CSC4=NC=NC(=NC=C3)C=C4)cc2)c1
CCOc1nc(NC(=O)Cc2cc(OC)c(S(C)(=O)=O)cc2OC)cc(N)c1C
CCOc1nc(NC(=O)Cc2cc(OC)c(S(C)(=O)=O)cc2OC)cc(N)c1CO
CCOc1nc(C(=O)NCc2ccc(S(C)(=O)=O)cc2)cc(N)c1Cl
CCOc1nc(NC(=O)Cc2cc(OC)c(S(C)(=O)=O)cc2OC)cc(N)c1C#N
CCOc1nc(NC(=O)Cc2cc(OO)c(S(C)(=O)=O)cc2OC)cc(N)c1Cl
CCOc1nc(NC(=O)Cc2cc(OC)c([SH](N)C(C)=O)cc2OC)cc(N)c1Cl
CCOc1nc(NC(=O)Cc2cc(OC)c(S(C)(=O)=O)cc2OC)cc(N)c1CON
COc1cc([SH](=O)=O)c(OC)cc1CC(=O)Nc1cc(N)c(C#N)c(OC(C)C)n1
C=[SH](=O)c1cc(OC)c(CC(=O)Nc2cc(N)c(C#N)c(OC(C)C)n2)cc1OC
C=S(N)(=O)c1cc(OC)c(CC(=O)Nc2cc(N)c(C#N)c(OC(C)C)n2)cc1OC
CCOc1nc(NC(=O)Cc2cc(OC)c([SH](C)(=O)NO)cc2OC)cc(N)c1C#N
COc1cc(S(N)(=O)=O)c(OC)cc1CC(=O)Nc1cc(N)c(C#N)c(OC(C)C)n1
COc1cc([SH](=O)=O)c(OC)cc1CC(=O)Nc1cc(N)c(C#N)c(OC(C)C)c1
CCOc1nc(C(=O)NCc2ccccc2S(N)(=O)=O)cc(N)c1C#N
CCOc1nc(CC(=O)Cc2cc(OC)c(S(C)(=O)=O)cc2OC)cc(N)c1Cl
CCOc1cc(N)cc(NC(=O)Cc2cc(OC)c(S(C)(=O)=O)cc2OC)n1
CCOc1nc(CC(=O)Cc2cc(OC)c(S(C)(=O)=O)cc2OC)cc(N)c1C#N
CCOc1nc(NC(=O)Cc2cc(OC)c(S(C)(=O)=O)cc2OC)cc(N)c1C=N
CCOC1=C(O)CNC(N)=CC(NC(=O)Cc2cc(OC)ccc2OC)=N1
CCOc1nc(NC(=O)Cc2cc(ON)ccc2OC)cc(N)c1C#N
CCOC1=C(C)ONC(N)NC(NC(=O)Cc2cc(OC)ccc2OC)=N1
CCOc1nc(NC(=O)Cc2cc(OC)ccc2OC)cc(N)[nH]oc1C
CCOC1=CC=C(N)C=CC=Cc2ccc(OO)cc2CC(=O)NC(N)=N1
CCCCC1=NC(OCC)=C(C)C(N)=CC(NC)=CC2(ON)C=C2CC(=O)N1
CCOc1nc(NC(=O)Cc2cc(OC)ccc2OC)cc(N)[nH]oc1O
CCOc1nc(NC(=O)Cc2cc(OO)ccc2OC)cc(N)c1C#N
CCOc1nc(NC(=O)Cc2cc(OC)ccc2OC)cc(O)c1C#N
CCOC1=C(C)C(N)=CC(OC)=CC2(OC)C=C2CC(=O)NC(C)=N1
CCOc1nc(NC(=O)Cc2cc(OC)ccc2OC)cc(C)c1C#N
CCOc1nc(C(=O)NCc2ccc(S(C)(=N)=O)cc2)cc(N)c1Cl
CCOc1nc(C(=O)NCc2ccc([SH](=O)=O)cc2)cc(N)c1Cl
CCOc1nc(NC(=O)CC2=CC(OC)N(S(C)(=O)=O)C=C2OC)cc(N)c1C
```

N#Cc1cnc2c(Cl)cc(NCc3cn(CCN4CCCCC4)nn3)cc2c1Nc1ccc(F)c(Cl)c1
N#Cc1cnc2c(Br)cc(NCc3c[nH]nn3)cc2c1Nc1ccc(F)c(Cl)c1
NCCc1cnc2c(Br)cc(NCc3c[nH]nn3)cc2c1Nc1ccc(F)c(Cl)c1
N#Cc1cnc2c(Br)cc(NCc3cnn[nH]3)cc2c1Nc1ccc(F)c(Cl)c1
NNCc1cnc2c(Br)cc(NCc3c[nH]nn3)cc2c1Nc1ccc(F)c(Cl)c1
N#Cc1cnc2c(Br)cc(NCc3cnc[nH]3)cc2c1Nc1ccc(F)c(Cl)c1
NNCc1cnc2c(Cl)cc(NCc3cnc[nH]3)cc2c1Nc1ccc(F)c(Cl)c1
N#Cc1cnc2c(Cl)cc(NCc3cn(CCN4CCCCC4)nn3)cc2c1Nc1ccc(N)c(Cl)c1
NCCc1cnc2c(Cl)cc(NCc3cnc[nH]3)cc2c1Nc1ccc(F)c(Cl)c1
N#CC1=CC(Cl)=c2ncc3cc(Cl)c(N)ccc[nH]c3c2=CC2=CC=C(CN1)N=NC2
N#Cc1cnc2c(Cl)cc(NCC3=CN=C[CH]3)cc2c1Nc1ccc(F)c(Cl)c1
N#Cc1ncc2nc1OCCCCOc1cc(NCc3cncs3)c(Cl)cc1NC(=O)N2
N#Cc1ncc2nc1OCCOCOc1cc(NCc3cncs3)c(Cl)cc1NC(=O)N2
N#Cc1ncc2cc1OCCCCOc1cc(NCc3cncs3)c(Cl)cc1NC(=O)N2
N#Cc1ncc2nc1OCCCCOc1cc(NCc3cccs3)c(Cl)cc1NC(=O)N2
N#Cc1ncc2nc1OOCCCCc1cc(NCc3cncs3)c(Cl)cc1NC(=O)N2
N#Cc1ncc2ssnc1OCCCCOc1cc(NCC3=CN=C3)c(Cl)cc1NC(=O)N2
N#Cc1ncc2nc1OCCCCCc1cc(NCc3cncs3)c(Cl)cc1NC(=O)N2
CCCOC1=C(C#N)N=CC2=NN=S3C=CC=C3CNc3cc(c(cc3Cl)NC(=O)N2)O1
N#Cc1ncc2cc1OCCOCOc1cc(NCc3cncs3)c(Cl)cc1NC(=O)N2
COc1cc2ncn(-c3cc(OCc4ccccc4C(F)(F)F)c(C(N)=O)s3)c2cc1OC
COc1cc2ncn(-c3cc(OCc4ccccc4C(F)(F)F)c(C(=O)O)s3)c2cc1OC
CCc1cc2c(cc1OC)ncn2-c1cc(OCc2ccccc2C(F)(F)F)c(C(N)=O)s1
CCOc1cc2c(cc1OC)ncn2-c1cc(OCc2ccccc2C(F)(F)F)c(C(N)=O)s1
COc1cc2ncn(-c3cc(OCc4ccccc4C(F)(F)F)c(C(N)=O)s3)c2cc1C
COc1cc2ncn(-c3cc(OCc4ccccc4C(F)(F)F)c(C(N)=O)s3)cc-2c1OC
C=C(N)c1sc(-n2cnc3cc(OC)c(OC)cc32)cc1OCc1ccccc1C(F)(F)F
N#Cc1cnc2cnc(NCc3cccnc3)cc2c1Nc1ccc(F)c(Cl)c1
c1cnc2nc(-c3ccc4[nH]ccc4c3)c(NC3CCCCC3)n2c1
c1cn2c(NC3CCCCC3)c(-c3ccc4[nH]ncc4c3)nc2nn1
c1ccn2c(NC3CCCCC3)c(-c3ccc4[nH]ncc4c3)nc2c1
C1=C2C(=N1)Nc1c(NC3CCCCC3)c2ccc[nH]c2c1=CC=2
c1cnc2cc(-c3ccc4[nH]ncc4c3)c(NC3CCCCC3)n2c1
CC(C)Nc1c(C2=NN=CC=NNC=CC=C2)nc2ncccn12
CC(C)Nc1c(-c2ccc3[nH]ncc3n2)nc2ncccn12

CC(C)Cc1c(-c2ccc3[nH]ncc3c2)nc2ncccn12
CC(C)Nc1c(-c2ccc3[nH]ccc3n2)nc2ncccn12
CC(C)Nc1c(-c2ccc3[nH]ccc3c2)nc2ncccn12
CC(C)Cc1c(-c2ccc3[nH]ncc3n2)nc2ncccn12
Cc1c(O)cccc1C(=O)N[C@@H](CSc1ccccc1)[C@H](O)CN1C[C@H]2CCCC[C@H]2C[C@H]1C(=O)NC(C)(C)C
Cc1c(O)cccc1C(=O)N[C@@H](CSc1ccccc1)[C@H](O)CN1C[C@H]2CCCC[C@H]2C[C@H]1C(=O)NC(C)C
Cc1c(O)cccc1C(=O)C[C@@H](CSc1ccccc1)[C@H](O)CN1C[C@H]2CCCC[C@H]2C[C@H]1C(=O)NC(C)(C)C
Cc1c(O)cccc1C(=O)N[C@@H](CSc1ccccc1)[C@H](O)CN1C[C@H]2CCCC[C@H]2C[C@H]1C(=O)CC(C)(C)C
Cc1cccc(-c2[nH]c(-c3ccnc(N)n3)cc2C(N)=O)c1C
Cc1cccc(-c2[nH]c(-c3ccnc(N)c3)cc2C(N)=O)c1C
C=C(N)c1cc(-c2ccncn[nH]cn2)[nH]c1-c1cccc(C)c1C
C=C(N)c1cc(-c2ccncn[nH]nc2)[nH]c1-c1cccc(C)c1C
Cc1cccc(-c2[nH]c(-c3cccc(N)n3)cc2C(N)=O)c1C
Cc1cccc(-c2[nH]c(-c3ccncc[nH]nn3)cc2C(N)=O)c1C
Cc1cccc(-c2[nH]c(-c3ccncn[nH]nn3)cc2C(N)=O)c1C
C=C(N)c1cc(-c2ccncc[nH]nn2)[nH]c1-c1cccc(C)c1C
C=C(N)c1cc(-c2ccnc(N)n2)[nH]c1-c1cccc(C)c1C
C=C(N)c1cc(-c2ccncn[nH]nn2)[nH]c1-c1cccc(C)c1C
CC1=CC(c2[nH]c(-c3ccnc(N)n3)cc2C(N)=O)=CCC1C
Cc1cccc(-c2[nH]c(-c3ccncn[nH]cn3)cc2C(N)=O)c1C
C=C(N)c1cc(C2=CC=NC=CN2)[nH]c1-c1cccc(C)c1C
C=C(N)c1cc(-c2ccncc[nH]cc2)[nH]c1-c1cccc(C)c1C
Nc1ncnc2onc(-c3ccc(NC(=O)Nc4cccc(C(F)(F)F)c4)cc3)c12
Nc1ncnc2occ(-c3ccc(NC(=O)Nc4cccc(C(F)(F)F)c4)cc3)c12
Nc1ncnc2occ(C3=C=C(NC(=O)Nc4cccc(C(F)(F)F)c4)C=C3)c12
Nc1nccc2occ(-c3ccc(NC(=O)Nc4cccc(C(F)(F)F)c4)cc3)c12
Nc1ncnc2scc(-c3ccc(NC(=O)Nc4cccc(C(F)(F)F)c4)cc3)c12
CC(F)(F)c1cccc(NC(=O)Nc2ccc(-c3noc4ncnc(N)c34)cc2)c1
Nc1ncnc2snc(-c3ccc(NC(=O)Nc4cccc(C(F)(F)F)c4)cc3)c12
CC(=O)Nc1c(C(N)=O)sc2ccc(Cl)c(Cl)c12
CC(=O)Nc1c(O)sc2ccc(Cl)c(Cl)c12
CC(Nc1cc2c(cn1)=CC=CSC=2)c1ccsc1
Cc1cccc(NC(=O)Nc2ccc(-c3csc4nc(N)nc(N)c34)cc2)c1
Cc1cccc(NC(=O)Nc2ccc(-c3csc4ncn[nH]nnc(N)c34)cc2)c1
Cc1cccc(NC(NO)Nc2ccc(-c3csc4nc(N)nc(N)c34)cc2)c1

```
Cc1cccc(NC(=O)Nc2ccc(-c3csc4ncn[nH]cnc(N)c34)cc2)c1
Cc1cccc(NC(=O)Nc2ccc(-c3cccc4c3=C(N)N=NC=NC=4)cc2)c1
Cc1cccc(NC(=O)Nc2ccc(-c3cccc4[nH]nc(N)c34)cc2)c1
Cc1cccc(NC(=O)Nc2ccc(C3=CC=C4N=CN=CN=NC(N)=C43)cc2)c1
C=C(Nc1ccc(-c2csc3nc(N)nc(N)c23)cc1)Nc1cccc(C)c1
COc1cccc(C(=O)Nc2cnc3[nH]cc(-c4ccccc4)c3c2)c1
CNc1cccc(C(=O)Nc2cnc3[nH]cc(-c4ccccc4)c3c2)c1
COc1cccc(C(=O)Nc2cnc3[nH]cc(-c4ccccn4)c3c2)c1
COc1cccc(C(=O)Nc2ccc3[nH]cc(-c4ccccc4)c3c2)c1
COc1cccc(C(=O)Nc2cnc3[nH]cc(-c4cccnc4)c3c2)c1
COc1ccc(C(=O)Nc2ccc3[nH]cc(-c4cccnc4)c3c2)cc1
NC1CCC(Nc2nccc(-c3c[nH]c4ncccc34)n2)CC1
NC1CCC(Nc2cccc(-c3c[nH]c4ncccc34)n2)CC1
NC1CCC(Nc2ncnc(-c3c[nH]c4ncccc34)n2)CC1
CC1CCC(Nc2nccc(-c3c[nH]c4ncccc34)n2)CC1
NC1CCC(Nc2nccc(C3=CNc4cc3ccn4)n2)CC1
NC1CCC(Nc2nccc(C3=CNC4=NC=CC=CN34)n2)CC1
NC1CCC(Nc2nccc(C3=CNC4=NN3C=NC=N4)n2)CC1
NC1C=C(Nc2nccc(-c3c[nH]c4ncccc34)n2)CC1
NC1CCC(Nc2nccc(C3=CNc4cccc3n4)n2)CC1
Cc1cccc(NC(=O)Nc2ccc(-c3cccc(C(N)=O)c3)cc2)c1
CC1CCCCC1Nc1nccc(-c2c[nH]c3ncccc23)n1
NC1CCCCC1Nc1nccc(-c2c[nH]c3ncccc23)n1
NC1CCCCC1Nc1cccc(-c2c[nH]c3ncccc23)n1
NC(COCc1ccccc1)COc1cncc(COCc2ccncc2)c1
NC(COCc1ccccc1)COc1cncc(CCCc2ccncc2)c1
Cc1cnc2cc(NS(=O)(=O)c3ccc(N)cc3)ccc2n1
Cc1nc2ccc(NS(=O)(=O)c3ccc(N)cc3)cc2nc1C
CC1=NC=CC=CC(NS(=O)(=O)c2ccc(N)cc2)=CC=NN=C1C
CCCCNc1ncnc2[nH]cc(SC)c12
O=C(Nc1cnccn1)Nc1ccnc2c(F)cccc12
O=C(Nc1cnccn1)Nc1cccc2c(F)cccc12
O=c1[nH]c2c(ccc3c([nH]1)=CC(F)=CC=C3)C=CN=NC=CC=CC=2
Fc1cccc2cccccc3c(ccc[nH]c12)C=CC=CC=C3
O=C(Nc1cccnc1)Nc1cccc2c(F)cccc12
```

```
O=C(NCc1ccc(Cl)c(Cl)c1)Nc1ccc2[nH]ncc2c1
O=C(NCc1ccc(Cl)cc1)Nc1ccc2[nH]ncc2c1
Cc1cc(CCC(=O)Nc2ccc3[nH]ncc3c2)ccc1Cl
Cc1cc(CNC(=O)Nc2ccc3[nH]ncc3c2)ccc1Cl
O=C(NCc1ccc(Cl)nc1)Nc1ccc2[nH]ncc2c1
CNC(=O)c1cc2c(-c3ccccc3F)n[nH]c2s1
CNC(=O)c1cc2c(-c3ccccc3)n[nH]c2s1
CNC(=O)c1cc2c(-c3ccccn3)n[nH]c2s1
OCCNc1cc2cc(-c3cccnc3)ccc2cn1
OCCNc1cc2cc(-c3ccccc3)ccc2cn1
NCCNc1cc2cc(-c3cccnc3)ccc2cn1
OCCCc1cc2cc(-c3cccnc3)ccc2cn1
OCCCc1ccc2ccc(-c3cccnc3)cc2c1
NCCNc1cc2cc(-c3ccncc3)ccc2cn1
OCCNc1cc2cc(-c3ccncc3)ccc2cn1
OCCCc1ccc2ccc(-c3ccccc3)cc2c1
NCCNc1cc2cc(-c3ccccc3)ccc2cn1
OCCNc1ccc2ccc(-c3cccnc3)cc2c1
CCCCc1cc2cc(-c3cccnc3)ccc2cn1
OCCC(=Cc1c2ccnc1N=C2)c1cccnc1
O=C1NC2=C(Cl)C=CC3=CC=CC2=C1O3
COc1ccc2c(c1)C(=Cc1cc[nH]c1)C(=O)N2
COc1ccc2c(c1)C(=Cc1c[nH]cn1)C(=O)N2
COc1ccc2c(c1)C(=Cc1c[nH]nn1)C(=O)N2
COc1ccc2c(c1)C(=Cc1c[nH]cn1)C(CO)N2
COc1ccc2c(c1)C(=Cc1cn[nH]c1)C(=O)N2
Cc1cc(NC(=O)CCNC(=O)Nc2nc(C)c(-c3ccc(-n4cccn4)cc3)s2)no1
Cc1cc(NC(=O)CCNC(=O)Nc2cc(C)c(-c3ccc(-n4cccn4)cc3)s2)no1
Cc1cc(NC(=O)CCNC(=O)Nc2cc(N)c(-c3ccc(-n4cccn4)cc3)s2)no1
Cc1cc(NC(=O)CCNC(=O)Nc2nc(N)c(-c3ccc(-n4cccn4)cc3)s2)no1
Cc1cc(NC(=O)CCNC(=O)Nc2nc(C)c(-c3ccc(-n4cccn4)cn3)s2)no1
Cc1cc(NC(=O)CCNC(=O)Nc2nc(C)c(-c3ccc(-n4cncn4)cc3)s2)no1
Cc1nc(NC(=O)CCNC(=O)Nc2nc(C)c(-c3ccc(-n4cccn4)cc3)s2)no1
Cc1cc(NC(=O)CCNC(=O)Nc2nc(C)c(-c3ccc(C4C=CC=N4)cc3)s2)no1
Cc1cc(NC(=O)CCNC(=O)Nc2nc(C)c(-c3ccc(-n4cccn4)cc3)s2)co1
```

```
CNC(=O)c1cc(Oc2ccc(NC(=O)Nc3cccc(C(F)(F)F)c3)cc2)ccn1
CNC(=O)c1cc(Oc2ccc(NC(=O)Nc3cccc(C(F)(F)F)c3)cc2)cnn1
CNC(=O)C1=CC(Oc2ccc(NC(=O)Nc3cccc(C(F)(F)F)c3)cc2)=C1C
Cn1cncc1C(OCc1c(-c2ccc(OC(F)(F)F)cc2)cc(C#N)c(=O)n1C)c1ccc(C#N)cc1
CN(C(=O)C1CCCCC1)c1ccc2c(c1)nc(NC(=O)c1ccc(C#N)cc1)n2C
CN(C(=O)C1CCCCC1)c1ccc2c(c1)cc(NC(=O)c1ccc(C#N)cc1)n2C
CC(C(=O)C1CCCCC1)c1ccc2c(c1)nc(NC(=O)c1ccc(C#N)cc1)n2C
COc1cccc(C(C)NC(=O)c2ccc(-c3ccncc3)c(C)c2)c1
COc1cccc(C(C)NC(=O)c2ccc(-c3ccncc3)cc2)c1
C=C(NC(C)c1cccc(OC)c1)c1ccc(-c2ccncc2)c(F)c1
COc1cccc(C(C)NC(=O)c2ccc(-c3ccccc3)c(F)c2)c1
COc1cccc(C(C)NC(=O)C2=CC=C(C3=CC=CC=NC=C3)C2(C)F)c1
Nc1n[nH]c2ccc(-c3nnn(Cc4ccccc4)c3I)cc12
Nc1n[nH]c2ccc(-c3ncn(Cc4ccccc4)c3I)cc12
Nc1n[nH]c2ccc(-c3cnn(Cc4ccccc4)c3I)cc12
Nc1n[nH]c2ccc(-c3cn(Cc4ccccc4)nn3)cc12
Nc1n[nH]ccc2c(-c3ncn(Cc4ccccc4)c3I)cc1-2
Cc1cc(NC(=O)Cc2ccc(-c3cccc4[nH]nc(N)c34)cc2)ccc1F
CC(=O)Nc1cccc(CNc2c(Nc3ccc4[nH]ncc4c3)c(=O)c2=O)c1
Cc1ccc(NC(=O)Nc2ccc(-c3cccc4c3CNC4=O)cc2)cc1
C=C(Nc1ccc(C)cc1)Nc1ccc(-c2cccc3c2CNC3=O)cc1
C=C1NCc2c1cccc2-c1ccc(NC(=O)Nc2ccc(C)cc2)cc1
C=C(Nc1ccc(-c2cccc3c2CNC3=O)cc1)Nc1ccc(C)c(C)c1
CN(C)c1ccc(CNC(=O)c2cc3c(-c4ccccc4)n[nH]c3s2)cc1
CNCCCc1ccc(CNC(=O)c2cc3c(-c4ccccc4)n[nH]c3s2)cc1
CN(C)C1C=CC(CCC(=O)c2cc3c(-c4ccccc4)n[nH]c3s2)=CC1
CN(C)c1ccc(CCC(=O)c2cc3c(-c4ccccc4)n[nH]c3s2)cc1
CN(C)c1ccc(CN(N=O)c2cc3c(-c4ccccc4)n[nH]c3s2)cc1
CC(C)C1=CC=C2C=CC=CC=C2c2n[nH]c3sc(cc23)C(=O)CCN=CC=C1
Cc1n[nH]c2ccc(-c3cncc(OCC(N)Cc4cccc(C(F)(F)F)c4)c3)cc12
Cc1n[nH]c2ccc(C3=CC(F)(F)c4cccc(c4)CC(N)COC=CN=C3)cc12
Cc1n[nH]c2cnc(C3=CC(F)(F)c4cccc(c4)CC(N)COC=CN=C3)cc12
Cc1n[nH]c2ccc(C3=CC(F)(F)c4cccc(c4)CC(N)COC(F)=CN=C3)cc12
Cc1n[nH]c2ccc(C3=CN=CC(OCC(N)Cc4cccc(C(F)(F)F)c4)=CC3)cc12
O=C1CCCN1Cc1ccc2c(c1)[nH]c(=O)c1cccn12
```

```
O=C1Cc2nc(CN3CCCC3=O)ccc2-n2cccc21
O=C1Cc2cc(CN3CCCC3=O)ccc2-n2cccc21
C=C1Nc2cc(CN3CCCC3=O)ccc2-n2cccc21
O=C1CCCC1Cc1ccc2c(c1)[nH]c(=O)c1cccn12
NC(COc1cncc(-c2ccc3c(c2)CC(=O)N3)c1)Cc1c[nH]c2ccccc12
NC(COc1cncc(-c2ccc3[nH]c(=O)oc3c2)c1)Cc1ccc2cccccc1-2
NC(COc1cncc(-c2ccc3[nH]c(=O)oc3c2)c1)CC1=CNc2cccc1c2
COCCN(C)C(=O)c1c(-c2ccc3[nH]nnc3c2)nnn1Cc1ccccc1
COCCN(C)C(=O)c1c(-c2ccc3n[nH]c-3cc2)nnn1Cc1ccccc1
COCCN(C)C(=O)c1c(-c2ccc3cnc-3sc2)nnn1Cc1ccccc1
COCCN(C)C(=O)c1c(C2=CC=c3[nH]ncc3=CS2)nnn1Cc1ccccc1
COCCN(C)C(=O)c1c(-c2ccc3[nH]ccc3c2)nnn1Cc1ccccc1
COCCN(C)C(=O)c1c(-c2ccc3[nH]nccc2-3)nnn1Cc1ccccc1
O=c1cc(-c2ccncc2)nc(NC2CCCCC2)[nH]1
O=c1cc(-c2ccncc2)nc(NC2CCCOC2)[nH]1
C=C1C=C(c2ccncc2)NCC(NC2CCCCC2)N1
C=C1C=C(c2ccncc2)N=C(C2CCCCCC2)N1
O=C1C=C(c2ccncc2)CCC(NC2CCCCC2)N1
C=C1CNC(c2ccncc2)NCC(NC2CCCCC2)C1
O=C1C=C(c2ccncc2)NCC(NC2CCCCC2)N1
C=C1C=C(c2ccncc2)N=C(NC2CCCNC2)N1
O=c1cc(-c2ccncc2)nc(NC2CCCCCC2)[nH]1
C=C1C=C(c2ccncc2)N=C(NC2CCCCC2)N1
O=c1cc(-c2ccncc2)nc(CC2CCCCC2)[nH]1
C=C1C=C(c2ccncc2)N=C(CC2CCCCC2)N1
O=c1cc(-c2ccncn2)nc(NC2CCCCN2)[nH]1
C=C1C=C(c2ccncc2)CCC(NC2CCCCC2)N1
C=C1C=C(c2ccncn2)CCC(NC2CCCCC2)C1
O=c1cc(-c2ccncc2)cc(NC2CCCCC2)[nH]1
O=c1cc(-c2ccncn2)nc(NC2CCCCC2)[nH]1
C=C1C=C(c2ccncc2)NCC(NC2CCCCC2)C1
O=C1CCC(c2ccncn2)N=C(CC2CCCCC2)N1
O=C1NCC(c2ccncc2)NCC(NC2CCCCC2)N1
O=C1NCC(c2ccncc2)N=C(CC2CCCCC2)N1
O=c1cc(-c2ccncc2)nc(NC2CCCCN2)[nH]1
```

```
O=C1CNC(c2ccncc2)N=C(NC2CCCCC2)N1
O=c1cc(-c2ccncc2)cc(CC2CCCCC2)[nH]1
C=C1C=C(NC2CCCCC2)CNC(c2ccncc2)=C1
C=C1C=C(NC2CCCCC2)CCC(c2ccncc2)=C1
O=C(Nc1cccc(C(F)(F)F)n1)Nc1ccnc2c(F)cccc12
O=C(Nc1cccc(C(F)(F)F)n1)Nc1ccncccc(F)cccc[nH]1
O=C1NC2=CC=CC(F)=CC=CC2=C(F)C(F)c2cccc(n2)N1
O=C(Nc1cccc(C(F)(F)F)n1)Nc1cc2oc(F)cccc1-2
CC(N)C1CCC(C(CO)Nc2ccnc3[nH]ccc23)CC1
N#Cc1cnc(NC(=O)Nc2ccc3cc(C(=O)O)ccc3c2)cn1
N#Cc1ccc(NC(=O)Nc2ccc3cc(C(=O)O)ccc3c2)cn1
N#Cc1ccc(NC(=O)Nc2ccc3cc(C(O)OO)ccc3c2)nc1
O=C1Nc2ccc(cn2)Cc2c(cc3ccc(C(=O)O)cnc2-3)N1
N#Cc1ccc(NC(=O)Nc2ccc3cc(C(=O)O)ccc3c2)nc1
N#Cc1ccc(NC(=O)NC2=CC=CC=CC(C(=O)O)=CC=CC=C2)cn1
NC1C=CC=C2C=CC(C(=O)O)=CC(=C2)NC(=O)Nc2cnc1cn2
O=C(NCc1ccccc1)Nc1ncc([N+](=O)[O-])s1
CC1=CC2CCC(=CC=C1)NC(=O)Nc1ccc(cc1)-c1csc3c(cnc(N)c13)C#CCN2C
CC1=CC(C)C(NC(=O)Nc2ccc(-c3csc4c(C)cnc(N)c34)cc2)=CC=C1
Cc1cccc(NC(=O)Nc2ccc(-c3csc4c(C#CCC5CCCC5)cnc(N)c34)cc2)c1
CC1=CC2CCCCC2CC#Cc2cnc(N)c3c(csc23)-c2ccc(cc2)NC(=O)NC(C)=C1
Cc1cccc(NC(=O)Nc2ccc(-c3csc4c(C#CCN5CCCCC5)cnc(N)c34)cc2)n1
CC1=CC(N)C(NC(=O)Nc2ccc(-c3csc4c(C)cnc(N)c34)cc2)=CC=C1
Cc1cccc(NC(=O)Nc2ccc(-c3csc4c(C#CCN)cnc(N)c34)cc2)c1
CC1=CC2CCC(=CC=C1)NC(=O)Nc1ccc(cc1)-c1csc3c(cnc(N)c13)C#CCN2O
Nc1n[nH]c2ccc(-c3nnn(Cc4ccccc4)c3-c3ccc(F)cc3)cc12
Nc1n[nH]c2ccc(-c3nnn(Cc4ccccc4)c3C3=CC=C3)cc12
Nc1n[nH]c2ccc(-c3nnn(Cc4ccccc4)c3C3=CC=CC=CC=C3)cc12
CC(C)NC(=O)COc1cccc(-c2nc(Nc3ccc4[nH]ncc4c3)c3ccccc3n2)c1
CC(C)NC(=O)COc1cccc(-c2nc(Nc3ccc4[nH]ncc4c3)cc3cccc-3n2)c1
C=C(COc1cccc(-c2nc(Nc3ccc4[nH]ncc4c3)c3ccccc3n2)c1)NN(C)C
NS(=O)(=O)c1cccc(Nc2ncc3ccn(Nc4ccccc4)c3n2)c1
NS(=O)(=O)c1cccc(Nc2cc3c(ccn3Cc3ccccc3)cn2)c1
NS(=O)(=O)c1cccc(Nc2ncc3cnn(Cc4ccccc4)c3n2)c1
NS(=O)(=O)c1cccc(Nc2ncc3ccn(Cc4ccccc4)c3n2)n1
```

```
CC(C)N(C)C(=O)c1c(-c2ccc3[nH]ncc3c2)nnn1Cc1ccccc1
CC(O)N(C)C(=O)c1c(-c2ccc3[nH]ncc3c2)nnn1Cc1ccccc1
CC(C)N(C)C(=O)C1=C(c2ccc3[nH]ncc3c2)N=NC1Cc1ccccc1
C=C(NC(C)c1cccc(OC)c1)c1ccc(-c2ccncc2)cc1
COc1cccc(C(C)NC(=O)C2=CC=C3C4=CC(=C23)C=C4)c1
COc1cccc(C2C3=CC=CC2NC(=O)c2ccccc23)c1
CN(C)c1cc2sncc2cc1NC(=O)C=O
Cc1cc(C)cc(NC(=O)Nc2ccc3c(c2)CCc2sc4ncnc(N)c4c2-3)c1
Cc1cc(C)cc(NC(=O)Nc2ccc3c(c2)CCC2=C3N3C(=NC=NC3N)S2)c1
Cc1cc(C)cc(NC(=O)Nc2ccc3c(c2)CCc2cc4ccnc(N)n4c2-3)c1
Cc1cc(C)cc(NC(=O)Nc2ccc3c(c2)CCc2sc4ccnc(N)c4c2-3)c1
Cc1c[nH]c2nccc(Oc3c(F)cc(Nc4cc(Cl)nc(N)n4)cc3F)c12
Cc1c[nH]c2nccc(Oc3c(F)cc(Nc4cc(Cl)cc(N)n4)cc3F)c12
Cc1c[nH]c2cccc(Oc3c(F)cc(Nc4cc(Cl)nc(N)n4)cc3F)c12
Cc1n[nH]c2cccc(Oc3c(F)cc(Nc4cc(Cl)nc(N)n4)cc3F)c12
COc1cccc(C(C)NC(=O)c2ccc(-c3ccncn3)c(C)c2)c1
Cc1ccc(NC(=O)Nc2ccc(-c3coc4ncnc(N)c34)cc2)cc1
B=C(Nc1ccc(C)cc1)Nc1ccc(-c2coc3ncnc(N)c23)cc1
Nc1ncnc2occ(-c3ccc(NC(=O)Nc4ccc(F)cc4)cc3)c12
NC(=O)c1cc2c(Cl)ccc(Br)c2s1
NC(=O)c1cc2c(Cl)ccc(BO)c2s1
NC(=O)c1sc2c(Br)ccc(Cl)c2c1N
NC(=O)C1=C2CCOCCC=CC2=C(BO)C=CS1
CC(=O)c1cc2c(Cl)ccc(BO)c2s1
NC(=O)c1sc2c(BO)ccc(Cl)c2c1N
CC(=O)c1sc2c(BO)ccc(Cl)c2c1N
Cc1cc(CNc2c(Nc3ccncc3)c(=O)c2=O)ccc1Cl
CCc1cc(CNc2c(Nc3ccncc3)c(=O)c2=O)ccc1Cl
O=c1c(NCc2ccc(Cl)c(Cl)c2)c(Nc2ccncc2)c1=O
O=c1c(CCc2ccc(Cl)c(Cl)c2)c(Nc2ccncc2)c1=O
O=c1c(NCC2=CC=C(Cl)C=CC=C2)c(Nc2ccncc2)c1=O
O=c1c(NCc2ccc(Cl)c(Cl)c2)c(Nc2ccccc2)c1=O
O=c1c(NCc2ccc(Cl)cc2)c(Nc2ccccc2)c1=O
O=c1c(Cc2ccncc2)c(NCc2ccc(Cl)cc2)c1=O
O=c1c(CCc2ccc(Cl)cc2)c(Nc2ccncc2)c1=O
```

```
Cc1cc(CNc2c(Cc3ccncc3)c(=O)c2=O)ccc1Cl
Cc1cc(CCc2c(Nc3ccncc3)c(=O)c2=O)ccc1Cl
O=C1C(Nc2ccncc2)=C(NCc2ccc(Cl)c(Cl)c2)C1CO
CCCS(=O)(=O)Nc1ccc(-c2ccc3n[nH]c(cc2)CCC(=O)N3)cc1
C=C(CC)Nc1n[nH]c2ccc(-c3ccc(NS(=O)(=O)CCC)cc3)cc12
CCC[SH](=O)(Nc1ccc(-c2ccc3[nH]nc(NC(=O)CC)c3c2)cc1)OO
CCCS(=O)(=O)Nc1ccc(-c2ccc3[nH]nc(NC(CC)OO)c3c2)cc1
CCCS(=O)(=O)Nc1ccc(-c2ccc3[nH]nc(NC(=O)CC)cc2-3)cc1
CCCS(=N)(=O)Nc1ccc(-c2ccc3[nH]nc(NC(=O)CC)c3c2)cc1
CCCS(=O)(=O)Nc1ccc(-c2ccc3[nH]nc(NCOOCC)c3c2)cc1
CCCS(=O)(=O)N1C=CC=C(c2ccc3[nH]nc(NC(=O)CC)c3c2)C=C1
Cc1csc(NC(=O)c2sc3nc(-c4ccncc4)ccc3c2N)n1
NCCCOc1cc2c(c(-c3ccc(Nc4cc5ccc(Cl)cc5o4)cc3)c1)CNC2=O
NC(=O)c1cc(-c2ccnc(N)n2)[nH]c1-c1ccc(Cl)cc1F
NC(=O)c1cc2cccccnc(N)nc3cc(Cl)ccc3c1[nH]2
NC(=O)c1cc(-c2ccncc[nH]cn2)[nH]c1-c1ccc(Cl)cc1F
NC(=O)c1cc(C2=CC=CC=NC(N)=N2)[nH]c1-c1ccc(Cl)cc1
O=C(NCc1ccc(Br)cc1)Nc1ccc2[nH]ncc2c1
O=C(NCc1ccc(Br)cc1)NC1=CC=CC=NNC=CC=C1
O=C(CCc1ccc(Br)cc1)Nc1ccc2[nH]ncc2c1
O=C(NCc1ccc(Br)cc1)Nc1ccc2[nH]ccc2c1
O=C1CCC2=Cc3[nH]ncc3C(=CN1)C=CC=C(Br)C=C2
NC(=O)c1cc(-c2ccnc(N)n2)[nH]c1-c1ccc(Cl)cc1Cl
CC12C=C(Cl)C=CC1=c1[nH]c(cc1C(N)=O)=CC=CC=NC(N)=N2
CC12C=C(Cl)C=CC1=c1[nH]c(cc1C(N)=O)=CC=NC=NC(N)=N2
CCc1cc(Cl)ccc1-c1[nH]c(-c2ccnc(N)n2)cc1C(N)=O
NC(=O)c1cc(-c2cccc(N)n2)[nH]c1-c1ccc(Cl)cc1Cl
COc1ccc(-c2cc(Nc3ccccc3C(N)=O)n3ccnc3c2)cc1OC
COc1ccc(-c2cc3nccn3c(Nc3ccccc3C(N)=O)n2)cc1OC
COc1ccc(-c2cc3nccn3c(Nc3ccncc3C(N)=O)n2)cc1OC
COc1ccc(-c2cc3nccn3c(Cc3ccccc3C(N)=O)n2)cc1OC
COc1ccc(-c2cc(Cc3ccccc3C(N)=O)n3ccnc3c2)cc1OC
CC(C)(C)c1ccc2nc(-c3n[nH]c4ccccc34)[nH]c2c1
CC(C)(C)c1ccc2nc(-c3n[nH]c4cnccc34)[nH]c2c1
CC(C)(C)c1ccc2nc(C3=NNC4=CC=CC=C3C=C4)[nH]c2c1
```

```
CC(C)(C)c1ccc2c(c1)C=NN=CC(c1n[nH]c3ccccc13)=N2
NC(COc1cccc(-c2ccc3c(c2)CC(=O)N3)c1)Cc1c[nH]c2ccccc12
COc1cc(C)c(Cl)cc1NC(=O)Nc1ccc(C#N)cn1
COc1cccc2c1Nc1cc(Nc3ccncc3)ccc1C(=O)N2
CCCC(=O)Nc1n[nH]c2ccc(-c3cccc(F)c3)cc12
CCCC(=O)Nc1n[nH]c2ncc(-c3cccc(F)c3)cc12
C=C(CCC)Nc1n[nH]c2ccc(-c3cccc(F)c3F)cc12
CCCC(=O)Nc1n[nH]c2ncc(-c3cccc(F)c3F)cc12
CCCC(=O)Nc1c[nH]c2ncc(-c3cccc(F)c3F)cc12
CCCC(=O)Nc1n[nH]c2ccc(-c3cccc(C)c3F)cc12
CCCC(=O)Nc1n[nH]c2ncc(-c3cccc(C)c3)cc12
CCCC(=O)Nc1n[nH]c2ccc(-c3cccc(C)c3F)nc12
C=C(CCC)Nc1n[nH]c2ccc(-c3cccc(F)c3)cc12
CCCC(=O)Nc1n[nH]c2ncc(-c3cncc(F)c3F)cc12
COC1=C(C(F)(F)F)c2c1n(O)c1ccc(Cl)cc1c2=O
Cn1nc(C(F)F)c2c(=O)c3cc(Cl)ccc3n(O)c21
Cn1nc(C(F)(F)F)c2c(=O)c3cc(Cl)ccc3n(O)c21
Nc1ncnc2c1sc1ncnc(N)c12
Nc1ncnc2sc3c(O)ncnc3c12
Nc1ncnc2cc3ccccncnc-3sc12
NC1=CC2=Cc3ncnc(N)c3SC2=NC=N1
Cc1ncnc2sc3c(N)ncnc3c12
OCC(CO)Nc1ncnc2[nH]nc(-c3ccccc3)c12
O=C1NCc2c1cccc2-c1ccc(Nc2nc3ccccc3o2)cc1
Nc1n[nH]c2ccc(-c3ccc(NS(=O)(=O)CO)cc3)cc12
CCS(=O)(=O)Nc1ccc(-c2ccc3[nH]nc(N)c3c2)cc1
Nc1nonc1-n1nnc(C(=O)NN=Cc2cccs2)c1-c1cccs1
Nc1nonc1-n1nnc(C(=O)NC=Cc2cccs2)c1-c1cccs1
Nc1nonc1-n1ccc(C(=O)NN=Cc2cccs2)c1-c1cccs1
Nc1nonc1-n1cnc(C(=O)NN=Cc2cccs2)c1-c1cccs1
CS(=O)(=O)Nc1ccc(-c2ccc3[nH]nc(N)c3c2)cc1
CS(=O)(=O)Nc1ccc(-c2cccc3[nH]cn[nH]c-3cc2)cc1
CS(=O)(=O)Nc1ccc(-c2ccc3[nH]nc[nH]ccc3c2)cc1
NC(CO)c1cc2c(Oc3ccc(Br)cc3)cccc2s1
NC(=O)c1cc2c(Oc3ccc(Br)cc3)cncc2s1
```

```
NC(=O)c1cc2c(Oc3ccc(Br)cc3)cccc2s1
NC(=O)C1=CC2=CC(c3ccc(Br)cc3)=CC=CN2S1
NC(CO)c1cc2c(Oc3ccc(Br)cc3)cncc2s1
NC(=O)c1cc2c(Oc3ccc(Br)cc3)c-2c1
CCc1nccn2c(-c3ccnc(NCC(C)(C)O)n3)c(-c3ccc(F)cc3F)nc12
CCc1nccn2c(-c3ccnc(CCC(C)(C)O)n3)c(-c3ccc(F)cc3)nc12
O=C(Nc1cnccn1)Nc1ccnc2ccccc12
O=C(Nc1cnccn1)Nc1ncnc2nnccc12
O=C(Nc1cnccn1)Nc1ncnc2ccccc12
O=C(Nc1ccccn1)Nc1ccnc2ccccc12
O=C(Cc1cnccn1)Nc1ccnc2ccccc12
O=C(Nc1cnccn1)Nc1ncnc2ncccc12
O=C(Nc1cnncn1)Nc1ccnc2ccccc12
O=C(Nc1cnccn1)Nc1ccnc2cnccc12
O=C(Nc1ccnc2ccccc12)Oc1cnccn1
O=C(Nc1cnccn1)Nc1ncnc2cnccc12
O=C(Nc1ncnc2ccccc12)Oc1cnccn1
Cc1ccc(F)c(NC(=O)Nc2ccc(-c3nsc4ncnc(N)c34)cc2)c1
Cc1ccc(F)c(NC(=O)Nc2ccc(-c3csc4ncnc(N)c34)cc2)c1
Cc1ccc(F)c(NC(=O)Nc2ccc(-c3cnc4ncnc(N)n34)cc2)c1
Cc1ccc(F)c(NC(=O)Nc2ccc(-c3nsc4ccnc(N)c34)cc2)c1
Cc1ccc(F)c(NC(=O)Nc2ccc(-c3nsc4nnnc(N)c34)cc2)c1
Cc1ccc(F)c(CC(=O)Nc2ccc(-c3nsc4ncnc(N)c34)cc2)c1
Cc1ccc(F)c(NC(=O)Nc2ccc(-c3csc4ccnc(N)c34)cc2)c1
Cc1ccc(F)c(NC(=O)Nc2ccc(-c3cnn4ncnc(N)c34)cc2)c1
O=C(NNC(=S)Nc1ccc(Cl)cc1)C(O)(c1ccccc1)c1ccccc1
O=C(NNC(CS)Nc1ccc(Cl)cc1)C(O)(c1ccccc1)c1ccccc1
Cc1cccc(NC(=O)Nc2ccc(-c3csc4ccnc(N)c34)cc2)c1
Cc1cccc(NC(=O)Nc2ccc(C3=CSC4=NC=NC34NN)cc2)c1
Nc1ncnc2c1N=C(c1ccc(NC(=O)Nc3cc(C(F)(F)F)ccc3F)cc1)CCN2
Nc1ncnc2c1N=CCC(c1ccc(NC(=O)Nc3cc(C(F)(F)F)ccc3F)cc1)N2
Cc1cccc(NC(=O)Nc2ccc(-c3csc4c(C#CCN)cnc(N)c34)cc2)c1C
CCC#Cc1cnc(N)c2c(-c3ccc(NC(=O)Nc4cccc(C)c4)cc3)csc12
CC1=CCNCC#Cc2cnc(N)c3c(csc23)-c2ccc(cc2)NC(=O)NC=CC=C1
CC1=CC(C)C(NC(=O)Nc2ccc(-c3csc4c(C#CCN)cnc(N)c34)cc2)=CC=C1
```

```
CCCNCC#Cc1cnc(N)c2c(-c3ccc(NC(=O)NC4=CC=CC(C)=CC4)cc3)csc12
CCC#Cc1cnc(N)c2c(-c3ccc(NC(=O)Nc4cccc(C)c4C)cc3)csc12
CC1=NN=CN=Nc2ccc(-c3nnn(Cc4ccccc4)c3C)cc21
Nc1n[nH]c2ccc(CC3=C=NN3Cc3ccccc3)cc12
Nc1n[nH]c2ccc(Cc3ccnn3Cc3ccccc3)cc12
Cc1c(-c2ccc3[nH]nc(N)c3c2)nnn1Cc1ccccn1
Cc1c(-c2ccc3[nH]cc(N)c3c2)nnn1Cc1ccccc1
Cc1c(-c2cc3c(O)n[nH]c3cn2)nnn1Cc1ccccc1
Cc1c(-c2ccc3[nH]nc(F)c3c2)nnn1Cc1ccccc1
Cc1ccc(F)c(NC(=O)Nc2ccc(-c3cncc4c3c(N)nn4C)cc2)c1
Cc1ccc(F)c(NC(=O)Nc2ccc(-c3cnc4nc3C(N)=NN4C)cc2)c1
Cc1ccc(F)c(NC(=O)Nc2ccc(-c3cnc4cc3C(N)=NN4C)cc2)c1
Cc1ccc(F)c(NC(=O)Nc2ccc(C3=CN=CC=NC=CC(N)=NN3C)cc2)c1
Cc1ccc(F)c(NC(=O)Nc2ccc(-c3cccc4c3c(N)nn4C)cc2)c1
Cc1ccc(F)c(NC(=O)Nc2ccc(-c3cnc4nc-4c(N)nn3C)cc2)c1
Cc1ccc(F)c(NC(=O)Nc2ccc(-c3cnc4nc3C(N)=CN4C)cc2)c1
O=C(NC(CO)Cc1ccc(Cl)cc1)c1ccc(-c2ccncc2)cc1
O=C(Nc1cccc(C(F)(F)F)n1)Nc1ccnc2cc(C(F)(F)F)ccc12
Clc1csc2ncnc(Nc3ccccc3)c12
O=C(Nc1nc2ccccc2n1CCCO)c1cccc([N+](=O)[O-])c1
O=C(Nc1nc2ccccc2n1CCCO)c1cccc([3C](=O)[O-])c1
O=C(Nc1nc2ccccc2n1CCCO)c1cccc([N+]([O-])CO)c1
Cc1cc(C)cc(NC(=O)Nc2ccc(-c3cccc4c3CNC4=O)c(C(F)(F)F)c2)c1
Cc1cc(C)cc(NC(=O)Nc2ccc(-c3cccc4c3CNC4=O)c(C(C)(F)F)c2)c1
O=c1[nH]cnc2c1sc1c(Cl)ccc(Cl)c12
O=CCNn1[nH]c2c3c(Cl)ccc(Cl)c3sc21
O=C1C=C2C=c3c(Cl)ccc(Cl)c3=C2N1
O=C1NCNNc2c1sc1c(Cl)ccc(Cl)c21
NC(=O)c1cc2c(-c3ccc(F)cc3)cncc2s1
COc1ccc2c(NC(=O)Nc3cccc(C(F)(F)F)n3)ccnc2c1
COc1ccc2c(NC(=O)Nc3cccc(C(C)(F)F)n3)ccnc2c1
COc1ccc2c(NC(=O)Nc3cccc(C(F)CF)n3)ccnc2c1
Oc1c2ccccc2c2c3c(cccc13)N=N2
OC1=C2C=CCC12C1=CC=CC=CC=CNCN=N1
OC1=c2ccccc2=CCN=NC2=CC1=CC=CC=C2
```

```
OC1=C2C=CCC12c1nnc2cccccc1-2
O=C(NNC(=S)Nc1ccc(O)cc1)C(O)(c1ccccc1)c1ccccc1
CCOc1nc(C(=O)NCc2ccc(S(C)(=O)=O)cc2)cc(N)c1C#N
CCOc1nc(C(=O)NCc2ccc([SH](=O)=O)cc2)cc(N)c1C#N
NC(=O)c1sc2ccc(Cl)c(Cl)c2c1NC(=O)c1ccc(Cl)cc1
NC(=O)c1sc2ccc(Cl)c(CO)c2c1NC(=O)c1ccc(Cl)cc1
O=c1[nH]c2sc3c(c2c2nc(-c4ccncc4)nn12)CCCC3
O=c1[nH]c2sc3c(c2c2nc(-c4ccccc4)nn12)CCCC3
O=C1NC=CSC2=C(C=Cc3nc(-c4ccncc4)nn31)CCCC2
O=c1[nH]c2c(c3nc(-c4ccncc4)nn13)=C1CCCCC=21
O=c1[nH]c2sc3c(c2c2nc(-c4ccnnc4)nn12)CCCC3
O=c1[nH]c2sc3c(c2c2nc(-c4ccncn4)nn12)CCCC3
CC(=O)NCC(=O)N1C2CCC1c1cc(Nc3ncc(C(F)(F)F)c(NC4CCC4)n3)ccc12
C=C(CNC(C)=O)N1C2CCC1c1cc(Nc3ncc(C(F)(F)F)c(NC4CCC4)n3)ccc12
CC(=O)NCC(=O)N1C2CCC1c1cc(Nc3ccc(C(F)(F)F)c(NC4CCC4)n3)ccc12
CC(=O)NCC(=O)N1C2CCC1c1cc(Nc3ncc(C(C)(F)F)c(NC4CCC4)n3)ccc12
CC(=O)NCC(=O)N1C2CCC1c1cc(NC3=NC(C4CCC4)C=C(C(F)(F)F)C=N3)ccc12
CC(=O)NCC(=O)N1C2CCC1c1ccc2cc1Nc1ncc(C(F)(F)F)c(NC2CCC2)n1
OCCNc1cc2cc(-c3ccsc3)ccc2cn1
OCCCc1cc2cc(C3=CCC=C3)ccc2cn1
OCCNC1=CC=CC=C(c2ccsc2)C=CC=CC=N1
OCCCc1cc2cc(-c3ccccc3)ccc2cn1
OCCNc1cc2cc(-c3ccsc3)cnc2cn1
OCCNc1cc2c(cn1)=CC=S(c1ccsc1)C=2
OC(Nc1cc2c(cn1)=CC=CSC=2)c1ccsc1
OC(NC1=CC=C2C=C2SC=CC=C1)c1ccccc1
OCCCc1cc2cc(-c3ccccc3)cnc2cn1
OCCNC1=CC=CC=C(c2ccsc2)N=CC=CC=N1
Nc1n[nH]c2nnc(-c3ccccc3)c(-c3ccccc3)c12
NC1N=[C+]C=CN=NC2=CC=CC=CC=CC=CC=CC=CC=CC=C21
CCn1c(-c2cnon2)nc2cnc(Oc3cccc(NC(=O)c4ccc(OCCN5CCOCC5)cc4)c3)cc21
CCn1c(-c2nonc2N)nc2cnc(Oc3cccc(NC(=O)c4ccc(OCCN5CCOCC5)cc4)c3)cc21
CCn1c(-c2ncnn2N)nc2cnc(Oc3cccc(NC(=O)c4ccc(OCCN5CCOCC5)cc4)c3)cc21
CCn1c(-c2nonc2N)nc2cnc(Oc3cccc(NC(=O)c4ccc(CCCN5CCOCC5)cc4)c3)cc21
CCn1c(-c2non2N)nc2cnc(Oc3cccc(NC(=O)c4ccc(OCCN5CCOCC5)cc4)c3)cc21
```

```
CCn1c(C2=NN=C2)nc2cnc(Oc3cccc(NC(=O)c4ccc(OCCN5CCOCC5)cc4)c3)cc21
CCn1c(-c2nocc2N)nc2cnc(Oc3cccc(NC(=O)c4ccc(OCCN5CCOCC5)cc4)c3)cc21
COc1ccc(C(=O)Nc2n[nH]c3ccc(-c4cn(Cc5ccccc5)nn4)cc23)cc1OC
COc1ccc(C(=O)Nc2n[nH]c3ccc(-c4cnn(Cc5ccccc5)c4)cc23)cc1OC
C=C1C2=C(C(=O)N=CCNCCO)C=CC(C=CC=C2)NCC1C
CCCN(CO)C(=O)C1=CC=CCCCC(C)C1=O
COc1ccc(C(=O)Nc2cnc3[nH]cc(-c4ccccc4)c3c2)cc1
COc1ccc(C(=O)Nc2cnc3c(c2)C2=CC=CC2=CC=CN3)cc1
Cc1cccc(NC(=O)Nc2ccc(-c3coc4ccnc(N)c34)cc2)c1
Cc1cccc(NC(=O)Nc2ccc(-c3noc4ncnc(N)c34)cc2)c1
Cc1cccc(NC(=O)Nc2ccc(-c3noc4ccnc(N)c34)cc2)c1
Cc1cccc(NC(=O)Nc2ccc(-c3ccc4ccnc(N)n34)cc2)c1
Cc1cccc(NC(=O)Nc2ccc(-c3coc4ncnc(N)c34)cc2)c1C
Oc1ccc(-c2n[nH]c(Nc3cccc(Cl)c3)c2-c2ccc(O)cc2)cc1
OC1=CC=C(c2c(-c3ccc(O)cc3)n[nH]c2Nc2cccc(Cl)c2)C1
CNc1ccnc2c1N=C(c1ccc(NC(=O)Nc3cccc(C(F)(F)F)c3)cc1)CCN2
COc1ccnc2c1N=C(c1ccc(NC(=O)Nc3cccc(C(F)(F)F)c3)cc1)CCN2
CNc1ncnc2c1N=C(c1ccc(NC(=O)Nc3cccc(C(C)(F)F)c3)cc1)CCN2
CNc1ncnc2c1N=C(c1ccc(NC(=O)Nc3cccc(C(F)(F)F)c3)cc1)CCS2
CC(C)(CO)c1nnc2ccc(-c3c(-c4ccc(F)cc4F)nc4n3CCC4)nn12
CC(C)(CO)c1nnc2ccc(-c3c(-c4ccc(F)cc4F)nc4n3C=CC4)nn12
CC(C)(CO)c1nnc2ccc(-c3c(-c4ccc(F)cc4)nc4occn34)nn12
CC1C2=CN=NC=CC=CN=NN=NC2=CNC1c1c(-c2ccc(F)cc2F)nc2n1C=CC2
CC(C)(CO)c1nnc2ccc(-c3c(-c4ccc(F)cc4)nc4n3CCC4)nn12
CS(=O)(=O)NCCOc1cncc(-c2ccc3cnccc3c2)c1
CS(=O)(=O)NCCOc1cncc(-c2ccc3ccccc3c2)c1
CS(=O)(=O)NCCOc1cccc(-c2ccc3ccccc3c2)c1
CS(=O)(=O)NCCOC1=CN=CC2=CC2=CC=c2ccccc2=C1
Nc1cc(-c2c(-c3ccc(F)cc3)ncn2C2CCNCC2)ccn1
Nc1nccc(-c2c(-c3ccc(F)cc3)ncn2C2CCCCC2)n1
Nc1cc(-c2c(-c3ccc(F)cc3)ncn2C2=CCNCC2)ccn1
Nc1nccc(-c2c(-c3ccc(F)nc3)ncn2C2CCNCC2)n1
O=C(Nc1cnccn1)Nc1ccnc2c(C(F)(F)F)cccc12
O=C(Nc1ccccn1)Nc1cnnc2c(C(F)(F)F)cccc12
O=C(Nc1cnccn1)Nc1cccc2c(C(F)(F)F)cccc12
```

```
O=C(Nc1ccccn1)Nc1ccnc2c(C(F)(F)F)cccc12
O=C(Nc1cnccn1)Nc1cnnc2c(C(F)(F)F)cccc12
O=C(Nc1ccccn1)Nc1cccc2c(C(F)(F)F)cccc12
O=C(Nc1ccccc1)Nc1ccnc2c(C(F)(F)F)cccc12
O=C(Nc1cnccn1)Nc1cncc2c(C(F)(F)F)cccc12
O=C(Nc1ccccc1)Nc1cnnc2c(C(F)(F)F)cccc12
O=C(Nc1cccnc1)Nc1ccnc2c(C(F)(F)F)cccc12
CC(F)(F)c1cccc2c(NC(=O)Nc3cnccn3)cccc12
Cc1ccc2nccc(NC(=O)Nc3cccc(C(F)(F)F)n3)c2c1
O=C(NC1=CC=NC=CC(Cl)=CC=CC=C1)Nc1cccc(C(F)(F)F)n1
CC1=CC=CC=CC23N=C(C(F)(F)F)C=CC=C(NC2=O)NC3=CC=CC=C1
O=C(Nc1cccc(C(F)(F)F)n1)Nc1ccnc2c(Cl)cccc12
O=C(Nc1ccc2cc(Cl)c-2nnc1)Nc1cccc(C(F)(F)F)n1
O=C(Nc1cc[nH]ccn(Cl)ccnnc1)Nc1cccc(C(F)(F)F)c1
Cc1cccc(NC(=O)Nc2ccc(-c3csc4c(C#CCN(C)C)cnc(N)c34)cc2)c1
CN(C)CC#Cc1cnc(N)c2c(-c3ccc(NC(=O)NC4CC4(C)F)cc3)csc12
CN(C)CC#Cc1cnc(N)c2c(-c3ccc(NC(=O)Nc4cccc(F)c4)cc3)csc12
Cc1cccc(NC(=O)Nc2ccc(-c3csc4c(CCCCN(C)C)cnc(N)c34)cc2)c1
CN(C)CC#Cc1cnc(N)c2c(-c3ccc(CC(=O)Nc4cccc(N)c4)cc3)csc12
CN(C)CC#Cc1cnc(N)c2c(-c3ccc(NC(=O)Nc4cccc(N)c4)cc3)csc12
Fc1cnc(Nc2ccccc2)nc1Nc1ccccc1
Fc1ccc(Nc2ccccc2)nc1Nc1ccncc1
Cc1n[nH]c2c1C(c1ccccc1Cl)C(C#N)=C(N)O2
Cc1c[nH]c2c1C(c1ccccc1Cl)C(C#N)=C(N)O2
Cc1cc(C)cc(NC(=O)Nc2ccc(-c3c(C)sc4ncnc(N)c34)cc2)c1
Cc1cc(C)cc(NC(=O)NC2=C=C(c3c(C)sc4ncnc(N)c34)C=C2)c1
Cc1cc(C)cc(NC(=O)Nc2ccc(-c3c(C)sc4ccnc(N)c34)cc2)c1
Cc1cc(C)cc(NC(=O)NC2=C=C(c3c(C)sc4ccnc(N)c34)C=C2)c1
Cc1cc(N)cc(NC(=O)NC2=C=C(c3c(C)sc4ccnc(F)c34)C=C2)c1
CC(C)Cc1cc(-c2ccc3[nH]nc(N)c3c2)on1
CC(C)CC1=CC(c2ccc3[nH]cc(N)c3c2)=N1
CC(C)Cc1cc(-c2ccc3[nH]cc(N)c3c2)on1
CC(C)CC1=CC(c2ccc3[nH]nc(N)c3c2)=N1
CC(C)Cc1coc(-c2cc3c(N)n[nH]c3cn2)c1
CC(C)Cc1coc(-c2ccc3[nH]nc(N)c3c2)c1
```

```
CC(C)Cc1cc(-c2cc3c(N)n[nH]c3cn2)on1
CC(C)Cc1cc(C2=CC=CC(N)=NNC=CC=C2)on1
Cc1ccc(NC(=O)Nc2ccc(-c3csc4ncnc(N)c34)cc2)cc1
Cc1cccc(CC(=O)Nc2ccc(-c3coc4ncnc(N)c34)cc2)c1
Cc1ccc(NC(=O)Nc2ccc(-c3coc4ncnc(N)c34)cc2)cc1C
Cc1ccc(CC(=O)Nc2ccc(-c3csc4ncnc(N)c34)cc2)cc1C
Cc1cccc(CC(=O)Nc2ccc(-c3csc4ncnc(N)c34)cc2)c1
Cc1cccc(C(=O)NNc2ccc(-c3coc4ncnc(N)c34)cc2)c1
CCC1C(C)=CC=C1NC(=O)Nc1ccc(-c2coc3ncnc(N)c23)cc1
CCC1C(C)=CC=C1NC(=O)Nc1ccc(-c2csc3ncnc(N)c23)cc1
Cc1cccc(CC(=O)Nc2ccc(-c3coc4ncnc(N)c34)cc2)c1C
CCC1C(C)=CC=C1NC(=O)Nc1ccc(-c2csc3ccnc(N)c23)cc1
Cc1ccc(NC(=O)Nc2ccc(-c3csc4cccc(N)c34)cc2)cc1
CCC1C(C)=CC=C1NC(=O)NN1C=CC=C(c2coc3ncnc(N)c23)C=C1
Cc1ccc(NC(=O)Nc2ccc(-c3coc4nccc(N)c34)cc2)cc1C
Cc1cccc(Nc2nc(NC3CCCCC3N)cnc2C(N)=O)c1
Cc1cccc(Nc2nc(NC3CCCCC3N)cnc2C(O)OO)c1
Cc1cccc(Nc2cc(NC=CCCCCCN)cnc2C(N)=O)n1
Cc1cccc(Nc2nc(NC3CCCCC3N)ccc2C(N)=O)c1
Cc1cccc(Nc2nc(NC3CCCCC3C)cnc2C(N)=O)c1
CC1=CC=CC=CC=c2nc(NC3CCCCC3C)c(n2C(N)=O)=C1
Cc1cccc(Nc2nc(NC3CCCCC3N)cnc2C(N)=O)n1
Cc1ccc(NC(=O)Nc2ccc(NC(=O)c3csc4ncnc(N)c34)cc2)cc1
Nc1ccc(NC(=O)Nc2ccc(NC(=O)c3csc4ncnc(N)c34)cc2)cc1
Cc1ccc(CC(=O)Nc2ccc(NC(=O)c3csc4ncnc(N)c34)cc2)cc1
Cc1ccc(NC(=O)Nc2ccc(NC(=O)c3csc4ccnc(N)c34)cc2)cc1
Cc1cc(Cl)ccc1NC(=S)NNC(=O)C(O)(c1ccccc1)c1ccccc1
CC(C)Cn1c(N)nc2ccc(-c3c(-c4ccc(F)cc4)nc4sccn34)cc21
CC(C)CC1CCNC=Nc2ccc(-c3c(-c4ccc(F)cc4)nc4sccn34)cc21
CC(C)Cn1c(N)nc2ccc(-c3c(-c4ccc(F)cc4)nc4occn34)cc21
CC(C)CC1C(N)=Nc2ccc(-c3c(-c4ccc(F)cc4)nc4occn34)cc21
CC(C)CC1CCNN=Nc2ccc(-c3c(-c4ccc(F)cc4F)nc4occn34)cc21
CC(C)Cn1c(N)nc2ccc(-c3c(-c4ccc(F)cc4)nc4n3C=CC4)cc21
CC(C)CC1C(N)=Nc2ccc(-c3c(-c4ccc(F)cc4)nc4sccn34)cc21
CC(C)CC1CCNC=Nc2ccc(-c3c(-c4ccc(F)cc4)nc4occn34)cc21
```

```
CCC(N)(C#Cc1cnc(N)c2c(-c3ccc(NC(=O)Nc4cccc(C)c4)cc3)csc12)CC
COc1cc(Nc2ncc([N+](=O)[O-])c(Nc3ccccc3C(N)=O)n2)cc(OC)c1OC
COC1=C(OC)CCOOC=CC(Nc2ncc([N+](=O)[O-])c(Nc3ccccc3C(N)=O)n2)=C1
COc1cc(Nc2ncc([N+](=O)[O-])c(Nc3ccccc3C(N)=O)n2)cc(OC)c1OO
COc1cc(Nc2ncc([N+](=O)[O-])c(Nc3ccccc3C(N)=O)n2)cc(OO)c1OC
COc1cc(Nc2ncc([N+](=O)[O-])c(Nc3ccccc3C(N)=O)n2)cc(OC)c1O
COC1=C(OC)CCOOC=CC(NC2=NC=CC=C(Nc3ccccc3C(N)=O)N2O[N+](=O)[O-])=C1
COc1ccc(CNc2c(Nc3ccc4[nH]ncc4c3)c(=O)c2=O)cc1
NC(COc1cncc(-c2cccc[nH]cc(C(=O)c3cccc(Cl)c3)c2)c1)Cc1c[nH]c2ccccc12
COc1cccc(C(C)NC(=O)c2ccc(-c3ccccc3)cc2NCCCN)c1
COc1cccc(C(C)NC(=O)c2ccc(-c3ccccc3)cc2)c1
COc1cccc(C(C)NC(=O)c2ccc3cc2NC=CC=CC=NC=C3)c1
COc1cccc(C(C)NC(=O)c2ccc3cc2NCCC=CC=NC=NC=C3)c1
Nc1ncnc2scc(-c3ccc(NC(=O)Nc4ccccc4)cc3)c12
Nc1ncnc2scc(C3=CC=C4C=CC=CC=CC=C3NC(=O)N4)c12
CCCS(=O)(=O)Nc1ccc(-c2ccc3[nH]nc(NC(C)NC)c3c2)cc1
CCCS(=O)(=O)Nc1ccc(-c2ccc3[nH]nc(NC(C)=O)c3c2)cc1
CCCS(=N)(=O)Nc1ccc(-c2ccc3[nH]nc(NC(C)=O)c3c2)cc1
C=C(C)Nc1n[nH]c2ccc(-c3ccc(NS(=O)(=O)CCC)cc3)cc12
CCCS(=O)(=O)Nc1ccc(C2=CC=C3NN=C(NC(N)=O)C32)cc1
C=C(C)Nc1n[nH]c2ccc(-c3ccc(N[SH](=O)(CCC)OO)cc3)cc12
CCC[SH](=O)(Nc1ccc(-c2ccc3[nH]nc(NC(C)=O)c3c2)cc1)OO
CCCS(Nc1ccc(-c2ccc3[nH]nc(NC(C)=O)c3c2)cc1)(OO)OO
CCC[SH](=O)(Nc1ccc(-c2ccc3[nH]nc(NC(C)OC)c3c2)cc1)OO
CCCS(=O)(=O)N1C=CC=C(c2ccc3[nH]nc(NC(C)=O)c3c2)C=C1
CCCS(=O)(=O)Nc1ccc(-c2ccc3[nH]nc(NC(C)NO)c3c2)cc1
COc1cccc(C(C)NC(=O)c2cc(C)c(-c3ccc4[nH]nc(C)c4c3)s2)c1
NC(=O)Cn1cc(-c2cc(-c3cc4ccccc4s3)c3[nH]ncc3c2)c2nc(N)ncc21
NC(=O)Cn1cc(-c2cc(-c3cc4ccccc4s3)c3[nH]ncc3c2)c2nc(N)ccc21
NC(=O)Cn1cc(-c2cc(-c3cc4ccccc4s3)c3[nH]ccc3c2)c2nc(N)ncc21
Cc1ccc(CCc2c(Nc3ccncc3)c(=O)c2=O)cc1
Cc1ccc(CNc2c(Nc3ccncc3)c(=O)c2=O)cc1
O=C1N=C(c2cc3c(Oc4ccc(F)cc4)cncc3s2)CO1
O=c1nc(-c2cc3c(Oc4ccc(I)cc4)cncc3s2)[nH]o1
O=C1N=C(c2cc3c(Oc4ccc(I)cc4)cncc3s2)CO1
```

```
O=c1nc(-c2cc3c(Oc4ccc(I)cc4)cncc3s2)[nH][nH]1
C=C1N=C(c2cc3c(Oc4ccc(I)cc4)cncc3s2)NO1
Cc1ccc(Oc2cncc3sc(-c4nc(=O)o[nH]4)cc23)cc1
NCC1CCC(CNc2nc(NCc3ccccc3Cl)ncc2[N+](=O)[O-])CC1
NCC1CCC(CNc2nc(NCc3ccccc3Cl)ccc2[N+](=O)[O-])CC1
Cc1cc(O)ccc1-n1cc(C(N)=O)c(=O)c2ccc(-c3ccncc3)cc21
CC(F)(F)c1cccc(NC(=O)Nc2ccnc3cc(C(F)(F)F)ccc23)n1
O=C1Nc2ccccc2Nc2nnc(I)cc21
O=C1Nc2ccccc2Nc2ncc(I)cc21
N=C1Nc2ccccc2Nc2nnc(I)cc21
Cc1ccc(F)c(NC(=O)Nc2ccc(-c3ccc4nccnc4c3N)cc2)c1
Cc1ccc(F)c(NC(=O)Nc2ccc(C3=CC=CC=NC=CN=CC=C3N)cc2)c1
Cc1ccc(F)c(NC(=O)Nc2ccc(-c3cnc4nccnc4c3N)cc2)c1
Cc1ccc(F)c(NC(=O)Nc2ccc(-c3ccc4nccnc4c3)cc2)c1
Nc1c(-c2ccc(NC(=O)Nc3cc(F)ccc3F)cc2)ccc2nccnc12
Cc1ccc(F)c(NC(=O)Nc2ccc(-c3ccc4cccnc4c3N)cc2)c1
Cc1ccc(F)c(CC(=O)Nc2ccc(-c3ccc4cccnc4n3)cc2)c1
COc1ccc(-c2ccc(NC(=O)Nc3cc(F)cc(F)c3)cc2)c2c(N)noc12
COc1ccc(-c2ccc(NC(=O)Nc3cc(C)cc(F)c3)cc2)c2c(N)coc12
COc1ccc(-c2ccc(NC(=O)Nc3cc(F)cc(F)c3)cc2)c2c(N)coc12
COc1ccc(-c2ccc(NC(=O)Nc3cc(C)cc(F)c3)cc2)c2c(N)noc12
Cc1ccc(-c2ccc(NC(=O)Nc3cccc(Br)c3)cc2)c2c(N)n[nH]c12
Cc1ccc(-c2ccc(NC(=O)Nc3cccc(Br)c3)cc2)ccc(N)nc(C)c1
Cc1ccc(-c2ccc(NC(=O)Nc3cccc(Br)c3)cc2)c2c1=CN=NC=NC=2N
CC1=CC=C(c2ccc(NC(=O)Nc3cccc(Br)c3)cc2)C2=C(N)N=C2CC=C1
CCN(CC)CC#Cc1ccc2c(c1)-c1[nH]nc(-c3ccc(C#N)nc3)c1C2
CCN(CC)CC#Cc1ccc2c(c1)-c1[nH]cc(-c3ccc(C#N)nc3)c1C2
CCN(CC)CC#Cc1ccc2c(c1)-c1[nH]nc(-c3ccc(C#N)cc3)c1C2
CCN(CC)CC#Cc1ccc2c(c1)-c1[nH]nc(-c3ccc(C)nc3)c1C2
COc1cc(C=C(C#N)c2nc3ccccc3[nH]2)c(Br)cc1O
COc1ccc(Br)c(C=C(CCN)c2nc3ccccc3[nH]2)c1
COc1ccc(Br)c(C=C(C#N)c2nc3ccccc3[nH]2)c1
Cc1cncc2cccc(S(=O)(=O)N3CCCNCC3C)c12
CC1=C2C=C(C=CC=C2S(=O)(=O)N2CCCNCC2C)C=CC1
Cc1cccc2cccc(S(=O)(=O)N3CCCNCC3C)c12
```

```
COc1cccc(C(C)NC(=O)N2CC=C(c3c[nH]c4ncccc34)CC2)c1
COc1cccc(C(C)NC(=O)N2CC=C(c3c[nH]c4ccccc34)CC2)c1
COc1cccc(C(C)NC(=O)N2CCCC(c3c[nH]c4ncccc34)CC2)c1
COc1cccc(C(C)NC(=O)N2CC=C(C3=CC=NC=Nc4ccccc43)CC2)c1
COc1cccc(C(C)NC(=O)N2CC=C(c3c[nH]c4nnccc34)CC2)c1
NC(=O)c1cccc2[nH]c(-c3ccccc3)nc12
NC(=O)C1=CC=CC=NNc2ccccnccc1n2
Cc1cccc(CC(=O)Nc2ccc(-c3cccc4[nH]nc(N)c34)cc2)c1
Cc1cccc(NC(=O)Nc2ccc(-c3ccc4n[nH]nc(N)c3-4)cc2)c1
NC(COc1cncc(C=Cc2ccncc2)c1)Cc1c[nH]c2ccccc12
NC(COc1cncc(C=Cc2ccncc2)c1)CC1=CNC=NC=CC=CC=C1
NC(COc1cccc(C=Cc2ccncc2)c1)Cc1c[nH]c2ccccc12
NC(COc1cncc(COCc2ccncc2)c1)Cc1c[nH]c2ccccc12
NC(COc1cncc(C=Cc2ccncc2)c1)Cc1cnnc2c1=CC=CC=NC=2
NC(COc1cncc(C=Cc2ccncc2)c1)CC1=CNc2ccc1cc2
NC(COc1cncc(CNCc2ccncc2)c1)Cc1c[nH]c2ccccc12
NC(COc1cncc(C=Cc2ccncc2)c1)Cc1ccccccncccnnc1
NC(COc1cncc(C=Cc2ccncc2)c1)Cc1cnncnc2ccc1cc2
NC(COc1cncc(C=Cc2ccncc2)c1)Cc1c[nH]c2cccnc12
O=C(NC(CO)c1ccccc1)N1CC=C(c2c[nH]c3ncccc23)CC1
CC(C)(C)OC(=O)n1ncc2cc(Nc3c(NCc4ccc(S(=O)(=O)O)cc4)c(=O)c3=O)ccc21
CC(C)(C)OC(=O)n1ncc2cc(Nc3c(NCc4ccc(S(N)(=O)=O)cc4)c(=O)c3=O)ccc21
Cc1cc(C)c(CCC2C(=O)Nc3ccccc32)[nH]1
Cc1n[nH]c2ccc(-c3cncc(OCC(N)Cc4c[nH]c5ccccc45)c3)cc12
Cc1n[nH]c2ccc(-c3cncc(CCC(N)Cc4c[nH]c5ccccc45)c3)cc12
O=c1nc(-c2ccc3[nH]ncc3c2)cc(-c2ccccc2)[nH]1
O=c1cc(-c2ccc3[nH]ncc3c2)cc(-c2ccccc2)[nH]1
O=c1cc(-c2ccc3[nH]nccc2-3)cc(-c2ccccc2)[nH]1
O=c1nc2cc([nH]1)c1cc(c3ccc2=CC=CC=NN3)C=C1
O=c1nc(-c2ccc3[nH]ncc3c2)cc(-c2ccccc2)o1
Cn1c(=O)c(Oc2ccc(F)cc2F)cc2cnc(NC3CCOCC3)nc21
COCCOCC#Cc1cc(-c2n[nH]c3c2C(=O)c2cc(CN4CCN(C)CC4)ccc2-3)cs1
COCCOCC#Cc1cc(-c2n[nH]c3c2C(=N)c2cc(CN4CCN(C)CC4)ccc2-3)cs1
COCCOCC#Cc1cc(C2=NNc3oc(c4c3C=C(CN3CCN(C)NC3)C=4)=C2)cs1
COCCOCC#Cc1cc(-c2n[nH]c3c2C(OO)c2cc(CN4CCN(C)CC4)ccc2-3)cs1
```

```
O=C1NCc2c1cccc2-c1ccc(Nc2nc3ccccc3[nH]2)cc1
OCC1NCc2c(-c3ccc(Nc4nc5ccccc5[nH]4)cc3)cccc21
O=C1NCc2c1cccc2-c1ccc(Nc2cnc3ccccc3n2)cc1
C=C(Nc1ccc(-c2noc3ncnc(N)c23)cc1)Nc1cccc(C)c1
Cc1cccc(NC(NO)Nc2ccc(-c3noc4ncnc(N)c34)cc2)c1C
Cc1cccc(CC(=O)Nc2ccc(-c3noc4ncnc(N)c34)cc2)c1
Cc1cccc(NC(=O)Nc2ccc(-c3ncc4ncnn(N)c3-4)cc2)c1
CNC(Nc1ccc(-c2noc3ncnc(N)c23)cc1)Nc1cccc(C)c1
Cc1cccc(NC(NO)Nc2ccc(-c3noc4ncnc(N)c34)cc2)c1
Cc1cccc(NC(NO)Nc2ccc(-c3coc4ncnc(N)c34)cc2)c1
CCc1c(C)cccc1NC(=O)Nc1ccc(-c2coc3ncnc(N)c23)cc1
O=C(NCCCCCCNC(=O)NCc1cccnc1)NCc1cccnc1
O=C(CCCCCCCNC(=O)NCc1cccnc1)NCc1cccnc1
O=C(NCCCCCCNC(=O)NCc1cccnc1)NCc1ccccc1
Cc1ccccc1NC(=O)Nc1ccc(-c2coc3ncnc(N)c23)cc1
Cc1ccccc1NC(=O)Nc1ccc(C2=CC=C3C=NN=CN3N=C2)cc1
Cc1ccccc1NC(=O)Nc1ccc(-c2coc3ccncc[nH]cc23)cc1
Cc1ccccc1NC(=O)NC1=CC(c2csc3ncnc(N)c23)=C=C1
Cc1ccccc1NC(=O)Nc1cccc(-c2coc3ncnc(N)c23)c1
Cc1ccccc1NC(=O)Nc1ccc(C2=CC=C3N=C3C(N)=C2)cc1
Cc1ccccc1NC(=O)Nc1ccc(-c2csc3ncnc(N)c23)cc1
Cc1ccccc1NC(=O)Nc1ccc(C2=COC3=NC3=CNC=C2)cc1
Cc1ccccc1NC(=O)Nc1ccc(C2=CC=C3N=CN=CN3N=C2)cc1
Cc1ccccc1NC(=O)Nc1ccc(-c2coc3nnnc(N)c23)cc1
Cc1ccccc1NC(=O)Nc1ccc(-c2coc3ccnc(N)c23)cc1
Cc1ccccc1NC(=O)NC1=CC2=C=C1C=C1C=CC=CC=CN=CC=C21
Cc1c(O)ccc2c1CCCC2=NNC(=O)Cc1cccc2ccccc12
Cc1c(O)ccc2c1CCCC2=NNC(=O)Nc1cccc2ccccc12
Cc1c(O)ccc2c1COCC2=NNC(=O)Cc1cccc2ccccc12
COc1cccc(C(C)NC(=O)c2ccc(C3=CC=CC=CC=C3)cc2)c1
COc1cccc(C(C)NC(=O)c2ccc(C3=CC=NC=CC=C3)cc2)c1
COc1cccc(C(C)NC(=O)c2ccc(C3=CC4=CC3=C4)cc2)c1
CCc1cnccc1-c1ccc(C(=O)NC(C)c2cccc(OC)c2)cc1
COc1cccc(C(C)NC(=O)c2ccc(C3=CC4=CC3=C4N)cc2)c1
CCOC(=O)c1ccc2[nH]c3c(c2c1)CCNC3=O
```

```
CCCC(=O)c1ccc2[nH]c3c(c2c1)CCNC3=O
CCOC(=O)c1ccc2[nH]c3c(cc1-2)CCNC3=O
CCCC(=O)c1ccn2c1C=CC=CC=CC(=O)C=CN2
CCOC(=O)C1=CC=C2NC=CC(=O)NCCC=C2C=C1
Cc1cccc(NC(=O)Nc2ccc(-c3cccc4c3CNC4=O)cc2)c1
C=C1NCc2c1cccc2-c1ccc(NC(=O)Nc2cccc(C)c2)cc1
C=C1CCc2c1cccc2-c1ccc(NC(=O)Nc2cccc(C)c2)cc1
c1cc2[nH]nnc2cc1-c1cn(CC2CCOCC2)nn1
c1cc2[nH]nnc2cc1-c1cnn(CC2CCOCC2)c1
c1cc2[nH]ncc2cc1-c1cn(CC2CCCCC2)nn1
c1cc2[nH]nnc2cc1-c1cn(CC2CCCCC2)nn1
c1cc2[nH]ncc2cc1-c1cn(CC2CCOCC2)cn1
c1cc2[nH]ncc2cc1-c1nnn(CC2CCOCC2)n1
c1cc2[nH]ncc2cc1-c1cnn(CC2CCOCC2)c1
c1cc2cc(-c3cn(CC4CCOCC4)nn3)ccc2[nH]1
C1=CN2NN=CN2C=C1c1cn(CC2CCOCC2)nn1
COc1cc(-c2ccc3[nH]nc(C(=O)Nc4ccccc4)c3c2)ccc1O
C=C(Nc1ccccc1)c1n[nH]c2ccc(-c3ccc(O)c(OC)c3)cc12
CN1CCC(NC(=O)c2cc3c(-c4ccccc4)n[nH]c3s2)CC1
CC1CCC(NC(=O)c2cc3c(-c4ccccc4)n[nH]c3s2)CC1
CC1CCC(NC(=O)c2cc3c(-c4ccccc4)n[nH]n3n2)CC1
CN1CCC(NC(=O)c2cc3c(-c4ccccc4)n[nH]c3s2)NC1
CC1CCC(CC(=O)c2cc3c(-c4ccccc4)n[nH]c3s2)CC1
CN1CCC(NC(CO)c2cc3c(-c4ccccc4)n[nH]c3s2)CC1
C=C1C(Nc2ccncc2)=C(NCc2cccc(C(O)(F)F)c2)C1CO
O=c1c(NCc2cccc(C(O)(F)F)c2)c(Nc2ccncc2)c1=O
O=C1C(Nc2ccncc2)=C(NCc2cccc(C(O)(F)F)c2)C1CO
O=c1c(NCc2cccc(C(F)(F)F)c2)c(Nc2ccncc2)c1=O
O=C1C(Nc2ccncc2)=C(NCc2cccc(C(F)(F)F)c2)C1CO
CC(O)(F)c1cccc(CNc2c(Nc3ccncc3)c(=O)c2=O)c1
CC=C1C(Nc2ccncc2)=C(NCc2cccc(C(O)(F)F)c2)C1O
OCCCCC=C1C=C(C(O)(F)F)C=CC=C2CNC2=C1Nc1ccncc1
C=C1C(=O)C(Nc2ccncc2)=C1NCc1cccc(C(F)(F)F)c1
Nc1n[nH]c2ccc(C3=C4C=CC=CC=C(C5=CC=CC=CC=C5)N(C4)N=N3)cc12
COc1cccc(C=C2SC(=Nc3ccccc3)NC2=O)c1
```

```
COC1=CC=CC=CC=C2SC1=NC1(C=CC=C1)NC2=O
COc1cccc(C=C2CC(=Nc3ccccc3)NC2=O)c1
COc1cccc(N=C2SC(=Nc3ccccc3)NC2=O)c1
C=c1[nH]c(=Nc2ccccc2)sc1=Cc1cccc(OC)c1
COc1cccc(C=c2sc(=Cc3ccccc3)[nH]c2=O)c1
C=C1NC(=Nc2ccccc2)SC1=Nc1cccc(OC)c1
COc1cccc(C=C2SC(=Nc3ccccc3)CC2=O)c1
O=C1NC(=Nc2ccccc2)SC1=Cc1cccc(Cl)c1
COc1cccc(C=C2SC(=Cc3ccccc3)CC2=O)c1
COc1cccc(C=C2SS(=Nc3ccccc3)NC2=O)c1
C=C1NC(=Nc2ccccc2)CC1=Cc1cccc(OC)c1
C=C1NC(=Cc2ccccc2)SC1=Cc1cccc(OC)c1
O=C1CC(=Nc2ccccc2)SC1=Cc1cccc(Cl)c1
Cc1cc(C)cc(NC(=O)Nc2ccc(-c3cccc4c3CNC4=O)c(C)c2)c1
CNC(O)CC1=C2C=CC(=C(NC=O)C=C(C)C=C(C)C=C1)C1=CC1=C2C
Cc1cc(C)cc(NC(=O)Nc2ccc(-c3cccc4c3CNC4NO)c(C)c2)c1
Cc1cc(C)cc(NC(=O)Nc2ccc(-c3cccc4c3CCC4=O)c(C)c2)c1
Cc1cc(C)cc(NC(=O)Nc2ccc(C3C=CC=C4C=CC3C4=O)c(C)c2)c1
Brc1cnc2[nH]cc(-c3ccccc3)c2c1
Brc1ccc2[nH]cc(-c3ccccc3)c2c1
Brc1cncn[nH]nc(-c2ccccccccc2)c1
Nc1nccc2scc(C(=O)NC3=C=C(NC(=O)Nc4ccccc4)C=C3)c12
Nc1nccc2scc(C(=O)Nc3ccc(NC(=O)Nc4ccccc4)cc3)c12
Nc1cccc2scc(C(=O)Nc3ccc(NC(=O)Nc4ccccc4)cc3)c12
Nc1ncnc2[nH]cc(C(=O)Nc3ccc(NC(=O)Nc4ccccc4)cc3)c12
Nc1ncnc2scc(C(=O)NC3=C=C(NC(=O)Nc4ccccc4)C=C3)c12
Oc1cccc(Nc2ncnc3scc(Cl)c23)c1
O=C(NN=Cc1ccc(O)cc1O)Nc1cccc2nsnc12
O=C(NN=Cc1ccc(O)cc1O)Nc1cccc2nscc12
O=N1CNC2CC=CC=C(O)C=CC=C3C=CC=CN=CCNC2=CC3=CN1
Nc1nc2ccc(-c3c(-c4ccc(F)cc4)nc4sccn34)cc2n1CC1CC1
Nc1nc2ccc(C3=S4C=NC=C4N=C3c3ccc(F)cc3)cc2n1CC1CC1
Nc1nc2ccc(-c3c(-c4ccc(F)cc4)nc4n3C=CC4)cc2n1CC1CC1
NCCNS(=O)(=O)c1cccc(C(=O)Nc2ccc(-c3n[nH]c(=O)c4ccccc34)cc2)c1
NCCN[SH](=O)(CO)c1cccc(C(=O)Nc2ccc(-c3n[nH]c(=O)c4ccccc34)cc2)c1
```

```
C=S(=O)(NCCN)c1cccc(C(=O)NC2=CC=C(c3n[nH]c(=O)c4ccccc34)NC2)c1
NCNNS(=O)(=O)c1cccc(C(=O)Nc2ccc(-c3n[nH]c(=O)c4ccccc34)cc2)c1
CN(C)CC(=O)NC(COc1cncc(-c2ccc3cnccc3c2)c1)Cc1c[nH]c2ccccc12
CN(C)CC(=O)NC(COc1cccc(-c2ccc3ccccc3c2)c1)Cc1c[nH]c2ccccc12
CN(C)CC(=O)NC(COc1cncc(-c2ccc3ccccc3c2)c1)Cc1c[nH]c2ccccc12
CN(C)CC(=O)NC(COc1cccc(-c2ccc3cnccc3c2)c1)Cc1c[nH]c2ccccc12
Cc1cc(C)cc(NC(=O)Nc2ccc(-c3cccc4snnc34)cc2)c1
Cc1cc(C)cc(NC(=O)Nc2ccc(-c3cccc4sncc34)cc2)c1
Cc1cc(C)cc(NC(=O)Nc2ccc(-c3ccnc4sncc34)cc2)c1
CC1=CC(c2ccc3cnn(N)c3cc3ccc([nH]c3)c(NO)[nH]2)=C1
Cc1cc(C)cc(NC(=O)Nc2ccc(-c3cccc4c3NSS4)cc2)c1
CCCOc1cccc(C(C)NC(=O)c2ccc(-c3ccncc3)cc2)c1
Cc1cc(C)cc(NC(=O)Nc2ccc(NC(=O)c3csc4ncnc(N)c34)cc2)c1
Cc1cc(C)cc(NC(NO)Nc2ccc(NC(=O)c3csc4ncnc(N)c34)cc2)c1
O=C(c1cc(Cc2n[nH]c(=O)c3c2CCCC3)ccc1F)N1CCN(c2ncccn2)CC1
O=C(c1cc(Cc2n[nH]c(=O)c3c2CCCC3)ccc1F)N1CCN(c2ccccn2)CC1
O=C(c1cc(Cc2n[nH]c(=O)c3c2CCCC3)ccc1F)N1CCC(c2ccccc2)CC1
O=c1c(Cc2ccncc2)c(NCCc2ccc(Oc3ccccc3)cc2)c1=O
CC1=CCC(NC(=O)Nc2ccc(-c3csc4ccnc(N)c34)cc2)=CC=C1
Nc1ncnc2scc(-c3ccc(NC(=O)Nc4cccc(Cl)c4)cc3)c12
Cc1cccc2[nH]c(=O)[nH]c3ccc(cccc-2c1)ccsc1ccnc(N)c31
CC1=COC(NC(=O)Nc2ccc(-c3csc4ccnc(N)c34)cc2)=CC=C1
Nc1nccc2scc(-c3ccc(NC(=O)Nc4cccc(Br)c4)cc3)c12
CN(C)C(=O)c1c(-c2ccc3[nH]ncc3c2)nnn1Cc1ccccc1
CCc1nccn2c(-c3ncnc(NCC(C)(C)O)n3)c(-c3ccc(F)cc3)nc12
Cc1nccn2c(-c3ccnc(CCC4(O)CC4)n3)c(-c3ccc(F)cc3)nc12
Cc1nccn2c(-c3ccnc(NCC4(O)CC4)n3)c(-c3ccc(F)cc3F)nc12
Cc1nccn2c(-c3cccc(NCC4(O)CC4)n3)c(-c3ccc(F)cc3)nc12
CCc1nccn2c(-c3ccnc(NCC(C)(C)CO)n3)c(-c3ccc(F)cc3F)nc12
Cc1nccn2c(-c3ccnc(NCC4(O)CC4)n3)c(-c3ccc(F)cc3)nc12
O=C(NCCCCCCNC(=O)Nc1cccnc1)Nc1cccnc1
O=C(NCCCCCCNC(=O)Nc1cccnc1)Nc1ccccc1
O=C1NC=CC=CC=NC=NN=CC=CC=CNC(=O)NCCCCCCN1
O=C1NCCCCCCOC(=O)NC=CC=CC=NC=C2C=C(C=N2)N1
O=C1C=CC(CCCCC(=O)NC2=CNCNC2)CN1
```

```
COOc1c(OC)cc2ncn(-c3cc(OCc4ccccc4C(F)(F)F)c(C(=O)O)s3)cc1-2
CCOc1cc2c(cc1OC)ncn2-c1cc(OCc2ccccc2C(F)(F)F)c(C(=O)O)s1
COc1cc2ncn(-c3cc(OCc4ccccc4C(F)(F)F)c(C(=O)O)s3)c2cc1OO
COc1cc2ncn(-c3cc(OCc4ccccc4C(F)(F)F)c(C(=O)O)s3)cc-2c1OC
CCOc1c(OC)cc2ncn(-c3cc(OCc4ccccc4C(F)(F)F)c(C(=O)O)s3)cc1-2
COOc1cc2c(cc1OC)ncn2-c1cc(OCc2ccccc2C(F)(F)F)c(C(=O)O)s1
COc1cc2ncn(C3=CC4=C(C(=O)O)SC5=C(C(F)(F)F)C3(C=C5)O4)c2cc1OC
COc1cc2ncn(-c3cc(OCc4ccccc4C(F)(F)F)c(C(=O)O)s3)c2cc1OCO
CCc1cc2c(cc1OC)ncn2-c1cc(OCc2ccccc2C(F)(F)F)c(C(=O)O)s1
COC1=Cc2ncn(-c3cc(OCc4ccccc4C(F)(F)F)c(C(N)=O)s3)c2C=CC1C
COc1cc2ncn(-c3cc(OCc4ccccc4C(F)(F)F)c(C(=O)O)s3)cc-2c1OCO
COc1cc2ncn(-c3cc(OCc4ccccc4C(F)(F)F)c(C(O)OO)s3)cc-2c1OO
Cc1nnc(-c2cc3c(Oc4ccc(C(F)(F)F)cc4)cncc3s2)o1
NCCC(C(=O)Nc1ccc2[nH]ncc2c1)c1ccc(Cl)c(Cl)c1
CCc1cc(C(CCN)C(=O)Nc2ccc3[nH]ncc3c2)ccc1Cl
Cc1cccc(NC(=O)Nc2ccc(-c3csc4c(C#CC(C)(C)N)cnc(N)c34)cc2)c1
Cc1cccc(NC(=O)Nc2ccc(-c3csc4c(C#CCNCC(C)N)cnc(N)c34)cc2)c1
Cc1cccc(NC(=O)Nc2ccc(-c3csc4c(C#CC(C)(C)C)cnc(N)c34)cc2)c1
Cc1cccc(NC(=O)Nc2ccc(-c3csc4c(C#CCNCC(C)C)cnc(N)c34)cc2)c1
Cc1cccc(NC(=O)Nc2ccc(-c3csc4c(C#CCNCC(C)O)cnc(N)c34)cc2)c1
CC(C)=NCC#Cc1cnc(N)c2c(-c3ccc(NC(=O)Nc4cccc(C)c4)cc3)csc12
CCCC1C=C(C)C=CC=C1NC(=O)Nc1ccc(-c2csc3c(C#CCN)cnc(N)c23)cc1
CC1=CC(CC(C)N)C#Cc2cnc(N)c3c(csc23)-c2ccc(cc2)NC(=O)NNC=CC=C1
Cc1cccc(NC(=O)Nc2ccc(-c3csc4c(C#CC(C)(C)O)cnc(N)c34)cc2)c1
CC1=CC(CC(C)C)C#Cc2cnc(N)c3c(csc23)-c2ccc(cc2)NC(CCO)=CC=C1
CC(C)(CO)CNc1cccc(-c2c(-c3ccc(F)cc3)nc3c(CC4CC4)nccn23)n1
CC(C)(CNc1nccc(-c2c(-c3ccc(F)cc3)nc3c(CC4CC4)nccn23)n1)OO
CC(C)(CO)CNc1nccc(-c2c(-c3ccc(F)cc3)nc3c(CC4CN4)nccn23)n1
CCCCOc1ncnc2[nH]cc(-c3ccccc3)c12
CCCCOc1ccc2cccccc-2ccccnncccncn1
OCCCOc1ncnc2[nH]cc(-c3ccccc3)c12
CCN(CC)CCNC(=O)c1c(C)[nH]c(C=C2C(=O)Nc3ccc(F)cc32)c1C
CCN(CC)CCNC(=O)c1c(C)[nH]c(C=C2c3cc(F)ccc3NC2NO)c1C
CCc1c(C=C2C(=O)Nc3ccc(F)cc32)[nH]c(C)c1C(=O)NCCN(CC)CC
CCN(CC)CCNC(=O)c1cc(C=C2C(=O)Nc3ccc(F)cc32)[nH]c1C
```

```
CCN(CC)CCNC(=O)c1c(C)[nH]c(CCC2C(=O)Nc3ccc(F)cc32)c1C
CCN(CC)CCNC(=O)C1=C(C)[CH]C(CCC2C(=O)Nc3ccc(F)cc32)=C1C
CCN(CC)CCNC(=O)C1=C(C)[CH]C(C=C2c3cc(F)ccc3NC2NO)=C1C
CCN(CC)CCNC(=O)c1cc(C=C2c3cc(F)ccc3NC2NO)[nH]c1C
CCN(CC)CCNC(=O)C1=C(C)[CH]C(C=C2C(=O)Nc3ccc(F)cc32)=C1C
CCN(CC)CCNC(=O)c1c(C)[nH]c(COC2C(=O)Nc3ccc(F)cc32)c1C
COc1nccc(-c2c(-c3ccc(F)cc3)ncn2C2CCC(O)CC2)n1
CCc1cccc(-c2c(-c3ccc(F)cc3)ccn2C2CCC(O)CC2)n1
CC(F)(F)c1ccc2c(NC(=O)Nc3cccc(C(F)(F)F)n3)ccnc2c1
COc1ccc(-c2ccc(NC(=O)Nc3cccc(Br)c3)cc2)c2c(N)noc12
COc1ccc(-c2ccc(NC(=O)Nc3cccc(Br)c3)cc2)c2c(N)coc12
COc1cc(-c2ccc3c(c2)Nc2ccc([N+](=O)[O-])cc2NC3=O)ccc1O
C=[N+]([O-])c1ccc2c(c1)NC(=O)c1ccc(-c3ccc(O)c(OC)c3)cc1N2
COc1cc(-c2ccc3c(c2)C=C2OC=C([N+](=O)[O-])C=C2NC3=O)ccc1O
Oc1ccc(Nc2[nH]nccc3ccc(O)cc3cc2-c2ccc(O)cc2)cc1
Oc1ccc(Cc2[nH]nc(-c3ccc(O)cc3)c2-c2ccc(O)cc2)cc1
COc1ccc(Br)c(C=C(C#N)c2nc3cc(C)ccc3[nH]2)c1
COc1cc(C=C(CNN)c2nc3cc(C)ccc3[nH]2)c(Br)cc1O
O=C(Nc1n[nH]c2ccc(-c3cn(Cc4cccnc4)nn3)cc12)c1ccccc1
O=C(Nc1n[nH]c2ccc(-c3cnn(Cc4ccccc4)c3)cc12)c1ccccc1
Cc1[nH]c(C=C2C(=O)Nc3ccccc32)c(C)c1CCC(=O)O
CC1=C(CCC=O)CCC=C(C=C2C(=O)Nc3ccccc32)N1
Cc1[nH]c(C=C2C(=O)Nc3ccccc32)c(C)c1CCC=O
Cc1[nH]c(C=C2C(=O)Nc3ccccc32)c(C)c1CCCC(C)O
Cc1[nH]c(C=C2C(=O)Nc3ccccc32)c(C)c1CCCOO
COc1cccc(C(CNNC(OO)c2ccccc2)C2=CC=N2)c1
COc1cccc(C(CN)NC(=O)c2ccc(-c3ccncc3)cc2)c1
C=C(NC(CN)c1cccc(OC)c1)c1ccc(-c2ccncc2)cc1
COc1cccc(C(CN)CC(=O)c2ccc(-c3ccncc3)cc2)c1
C=C(NC(CN)c1cccc(OC)c1)c1ccc(-c2ccncn2)cc1
COc1cccc(C(=N)CC(=O)c2ccc(-c3ccncc3)cc2)c1
COc1cccc(C(CNNNCO)c2ccc(-c3ccccc3)cc2)c1
CCC(NNCC(C1=CC=C1)c1cccc(OC)c1)c1ccccc1
COc1cccc(C(CN)NC(=O)c2ccc(C3=CC=C3)cc2)c1
COc1cccc(C(=N)NC(=O)c2ccc(-c3ccncc3)cc2)c1
```

```
COc1ccc(-c2ccc(NC(=O)Nc3ccc(O)c(C)c3)cc2)c2c(N)noc12
COc1ccc(-c2ccc(NC(=O)Nc3ccc(F)c(F)c3)cc2)c2c(N)noc12
COc1ccc(-c2ccc(NC(=O)Nc3ccc(O)c(F)c3)cc2)c2c(N)noc12
O=c1[nH]c2sccc2c2nc(-c3ccccc3)nn12
O=c1[nH]c2sccc2c2cc(-c3ccccc3)nn12
Cc1n[nH]c2ccc(-c3cncc(OCC(N)Cc4ccccc4)c3)cc12
Cc1n[nH]c2ccc(C3=CC(NCCc4ccccc4)C(O)=CN=C3)cc12
Cc1n[nH]c2ccc(C3=CC(NCCc4ccccc4)CONC=CN=C3)cc12
Cc1n[nH]c2cnc(-c3cncc(OCC(N)Cc4ccccc4)c3)cc12
Nc1nnc(-c2cc3c(Oc4ccc(Cl)cc4)cncc3s2)o1
Nc1nnc(-c2cc3c(Oc4ccc(Cl)cc4)cnnc3s2)o1
Nc1nnc(C2=Cc3c(Oc4ccc(Cl)cc4)cncc3SS2)o1
Nc1nnc(C2=Cc3c(Oc4ccc(Cl)cc4)cnnc3SS2)o1
Nc1nnc(C2=CC3=C(Oc4ccc(Cl)cc4)C=NC=CC=S23)o1
O=C1NCCc2[nH]c(-c3ccnc(-c4cnc5ccccc5c4)c3)cc21
O=C1NCCc2[nH]c(-c3ccnc(-c4cccccccnc4)c3)cc21
O=C1NCCc2[nH]c(-c3ccnc(-c4ccc5cccc-5cc4)c3)cc21
O=C1NCCc2[nH]c(-c3ccnc(C4=CN=CC=CC=CC=CC=C4)c3)cc21
O=C1NCCc2[nH]c(-c3ccnc(-c4cnc5cc4C=C5)c3)cc21
O=C1NCCc2[nH]c(-c3ccnc(-c4ccc5cccc-5nc4)c3)cc21
O=C1NCCc2[nH]c(-c3ccnc(-c4ccccc5cc-5nc4)c3)cc21
O=C1NCCc2[nH]c(-c3ccnc(-c4ccc5ccccc5c4)c3)cc21
O=C1NCCc2[nH]c(-c3ccnc(C4C=CC=CC=CC=C4)c3)cc21
O=C1NCCc2[nH]c(-c3ccnc(-c4ccccc5cc-5cc4)c3)cc21
O=C1NCCc2[nH]c(-c3ccnc(C4=CC=C5C=CC=C54)c3)cc21
O=C1NCCc2[nH]c(-c3ccnc(C4C=CC=CC=NC=C4)c3)cc21
O=C1NCCc2[nH]c(-c3ccnc(C4=CN=C5C=CC=C45)c3)cc21
CCCC(=O)Nc1n[nH]c2ccc(-c3cn(Cc4ccccc4)nn3)cc12
CCCC(=O)Nc1n[nH]c2ccc(-c3cn(Cc4ccccc4)cn3)cc12
CCCC(=O)Nc1n[nH]c2ncc(-c3cn(Cc4ccccc4)nn3)cc12
CCCC(=O)Nc1n[nH]c2ccc(C3=CN=CCc4ccccc4N=N3)cc12
CCCC(=O)Nc1n[nH]c2ccc(-c3cnccc4ccccc4nnc3)cc12
CCCC(=O)Nc1c[nH]c2ccc(-c3cnn(Cc4ccccc4)c3)cc12
CCCC(=O)Nc1n[nH]c2ccc(-c3cnccc4ccccc4cnn3)cc12
CCCC(=O)Nc1n[nH]c2ccc(-c3cn(Cc4ccccn4)nn3)cc12
```

```
CCCC(=O)Nc1n[nH]c2ccc(-c3cnc(-c4ccccc4)nn3)cc12
CC(C)(C)C(=O)Oc1ccc2c(c1)nc(NC(=O)c1cccc([N+](=O)[O-])c1)n2CCCO
CCCn1c(NC(=O)c2cccc([N+](=O)[O-])c2)nc2cc(OC(=O)C(C)(C)C)ccc21
CC(C)(C)C(=O)Oc1ccc2c(c1)cc(NC(=O)c1cccc([N+](=O)[O-])c1)n2CCCO
NCCCNS(=O)(=O)c1cccc(C(=O)Nc2ccc(-c3n[nH]c(=O)c4ccccc34)cc2)c1
COC12C=C(NO)C(=CN1)C=CC=c1ccc3c(c1=C2)=CC=CC=CC=3
Brc1ccc2[nH]nc(-c3ccccc3)c2c1
Brc1ccc2c(c1)C(c1ccccc1)=CN=CN=C2
Clc1ccc2cnc-2c2ccccncc2c1
Brc1ccc2c(c1)C(c1ccccc1)=NC=CN=C2
Brc1ccc2c(nc3c(ccc4cc43)c1)N=C2
Cc1ccc(Nc2nc3ccc([N+](=O)[O-])cc3[nH]2)nc1
Cc1ccc(Nc2cc3ccc([N+](=O)[O-])cc3[nH]2)nc1
Cc1ccc(Nc2nc3ncc([N+](=O)[O-])cc3[nH]2)nc1
Cc1ccc(Nc2nc3ccc([N+](=O)[O-])cc3[nH]2)cc1
O=C(Nc1cccc(N2CCOCC2)n1)Nc1ccnc2c(C(F)(F)F)cccc12
O=C(Nc1cccc(N2CCOCC2)n1)Nc1cccc2c(C(F)(F)F)cccc12
O=C(Nc1cccc(C2CCOCC2)n1)Nc1ccnc2c(C(F)(F)F)cccc12
O=C(Nc1cncc2c(C(F)(F)F)cccc12)N1C=CC=CC=N1
CCCNCCNCn1cnc(-c2ccc(F)cc2)c1-c1ccnc(OC)n1
COc1nccc(-c2c(-c3ccc(F)cc3)ncn2C2CCNCC2)n1
COc1nccc(C2=C(c3ccc(F)cc3)N=CC2C2CCNCC2)n1
COc1nccc(-c2c(-c3ccc(F)cc3)ncn2C2CCCCC2)n1
CCc1nccc(-c2c(-c3ccc(F)cc3)ncn2C2CCCCC2)n1
CCc1nccc(-c2c(-c3ccc(F)cc3)ncn2C2CCNCC2)n1
COc1nccc(-c2c(-c3ccc(O)cc3)ncn2C2CCNCC2)n1
COc1cc(-c2c(-c3ccc(F)cc3)ncn2C2CCNCC2)ccn1
CCCNCCCCn1cnc(-c2ccc(F)cc2)c1-c1ccnc(OC)n1
COc1cc(-c2c(-c3ccc(F)cc3)ncn2C2CCCCC2)ccn1
O=C(Nc1cnccn1)Nc1ccnc2ccc(C(F)(F)F)cc12
O=C(Nc1ccccc1)Nc1ccnc2ccc(C(F)(F)F)cc12
OOC(Nc1cnccn1)Nc1ccnc2ccc(C(F)(F)F)cc12
O=C(Nc1cccnc1)Nc1ccnc2ccc(C(F)(F)F)cc12
O=C(Nc1ccccn1)Nc1ccnc2ccc(C(F)(F)F)cc12
Cc1cc(Nc2cc(N3CCN(C)CC3)nc(C=Cc3ccccc3)n2)c[nH]1
```

```
Cc1cc(Nc2cc(N3CCN(C)CC3)cc(C=Cc3ccccc3)n2)n[nH]1
Cc1cc(Nc2cc(N3CCN(C)CC3)nc(C=Cc3ccccn3)n2)n[nH]1
Cc1cc(Nc2cc(N3CCN(C)CC3)nc(CCCc3ccccc3)n2)n[nH]1
Cc1cc(Nc2cc(N3CCN(C)CC3)nc(CCCN3C=CC=CC=C3)n2)c[nH]1
Cc1cc(Nc2cc(N3CCN(C)CC3)cc(C=Cc3ccccc3)n2)c[nH]1
Cc1cc(Nc2cc(N3CCN(C)CC3)cc(C=Cc3ccccn3)n2)n[nH]1
Cc1cc(Nc2cc(N3CCN(C)CC3)nc(CCCc3ccccn3)n2)n[nH]1
O=c1c(NCc2ccccc2F)c(Nc2ccncc2)c1=O
O=c1c(NNc2ccccc2F)c(Nc2ccncc2)c1=O
O=C1C(Nc2ccncc2)=C(NCc2ccccc2F)C1OO
O=C1C(Nc2ccncc2)=C(NCc2ccccc2F)C1CO
NC1CC(=O)C(NCc2ccccc2F)=C1Nc1ccncc1
O=C1C(=O)C(Nc2ccncc2)NC1NCc1ccccc1F
O=c1c(NCc2ccccc2F)c(Nc2ccccc2)c1=O
O=C1C(NCc2ccccc2F)=C(Nc2ccncc2)C1NO
Cc1ccccc1CNc1c(Nc2ccncc2)c(=O)c1=O
c1ccc(CNc2nc3ccc(-c4ccncc4)cc3s2)cc1
c1ccc(-c2ccc3nc(NCc4ccncc4)sc3c2)cc1
c1ccc(CNc2cc3ccc(-c4ccncc4)cc3s2)cc1
c1ccc(CNc2nc3ccc(-c4ccccc4)cc3s2)cc1
C1=CC=C(NCc2ccccc2)SC=CC=C(c2ccccc2)C=C1
C1=CNC=Cc2cc(c3cc2C=C3)SC(Cc2ccccc2)=C1
c1ccc(-c2ccc3cc(NCc4ccncc4)sc3c2)cc1
c1ccc(CNc2cc3ccc(-c4ccccc4)cc3s2)cc1
c1ccc(CCc2nc3ccc(-c4ccncc4)cc3s2)cc1
O=C(Nc1ccc(-c2cccc3sncc23)cc1)Nc1ccc(F)c(CF)c1
O=C(Nc1ccc(-c2cccc3sncc23)cc1)Nc1ccc(F)c(C(F)F)c1
O=C(Nc1ccc(-c2cccc3sncc23)cc1)Nc1ccc(F)c(C(F)(F)F)c1
CC(C)(CO)CNc1nccc(-c2c(-c3ccc(F)cc3)nc3cnccn23)n1
CC(C)(CO)CNc1nccc(-c2c(-c3ccc(F)cc3)nc3ccccn23)n1
CC(C)(CO)CNc1nccc(-c2c(-c3ccc(F)cc3)ncnc3ccn2-3)n1
CC(C)(CO)CNc1nccc(-c2c(-c3ccc(F)cc3F)nc3ccccn23)n1
CN1CCN(c2cccc(Nc3nccc(-c4c(C(N)=O)nc5ccccn45)n3)c2)CC1
CN1CCN(c2cccc(Nc3cc(-c4c(C(N)=O)nc5ccccn45)ccn3)c2)CC1
O=C(NCc1ccccc1)c1cccc(-c2ccc3c[nH]c-3nc2)c1
```

```
O=C(NCc1ccccc1)c1cccc(-c2cnc3[nH]ccc3c2)c1
O=C(NCc1ccccc1)c1cccc(-c2ccc3[nH]ncc3c2)c1
O=C(NCc1ccccc1)c1cccc(-c2cccc3[nH]c-3cc2)c1
O=C(NCc1ccccc1)c1cccc(-c2cncc3[nH]c-3nc2)c1
O=C(NCc1ccccc1)c1cccc(-c2ccc3[nH]c-3ccn2)c1
O=C(NCc1ccccc1)c1cccc(-c2cccc3[nH]c-3nc2)c1
O=C(NCc1ccccc1)c1cccc(-c2ccc3[nH]ccc3c2)c1
O=C1NCc2ccccc2C=CC=CC=CC=CC=C2C=C2NC2=NC=C12
O=C(NCc1ccccc1)c1cccc(-c2cnc3[nH]cc-3cn2)c1
O=C(NCc1ccccc1)c1cccc(-c2cnc3[nH]ncc3c2)c1
O=C(NCc1ccccc1)c1cccc(-c2cnc3[nH]c-3ccn2)c1
Cc1ccc(C(=O)O)cc1NC(=O)Nc1ccc(-c2cccc3[nH]nc(N)c23)cc1
Nc1c[nH]c2cccc(-c3ccc(NC(=O)Nc4cc(C(=O)O)ccc4F)cc3)c12
CC(=O)c1ccc(F)c(NC(=O)Nc2ccc(-c3cccc4[nH]nc(N)c34)cc2)c1
NC(=O)c1ccc(F)c(NC(=O)Nc2ccc(-c3cccc4[nH]cc(N)c34)cc2)c1
NC(=O)c1ccc(F)c(NC(=O)Nc2ccc(-c3cccc4[nH]nc(N)c34)cc2)c1
Cc1ccc(C(=O)O)cc1NC(=O)Nc1ccc(-c2cccc3[nH]cc(N)c23)cc1
Nc1n[nH]c2cccc(-c3ccc(NC(CO)Nc4cc(C(=O)O)ccc4F)cc3)c12
CC(=O)c1ccc(F)c(NC(=O)Cc2ccc(-c3cccc4[nH]cc(N)c34)cc2)c1
CCCC(=O)Nc1n[nH]c2nnc(-c3cccc(F)c3)cc12
CCCC(=O)Nc1n[nH]c2nnc(N3C=CC=C(C)C=C3)nc12
CCCC(=O)Nc1n[nH]c2ncc(-c3cccc(C)c3F)nc12
CCCC(CO)Nc1n[nH]c2ncc(-c3cccc(F)c3)cc12
CCCC(=O)Nc1n[nH]c2cnc(-c3cccc(F)c3)cc12
CNCC(CO)Nc1n[nH]c2nnc(-c3cccc(F)c3)cc12
CCCC(CO)Nc1n[nH]c2nnc(-c3cccc(F)c3)cc12
N#Cc1ccc(-c2n[nH]c3c2=c2ccc(OCCN4CCOCC4)cc2=3)cn1
CCCOCCCNCCOc1ccc2c(c1)-c1[nH]nc(-c3ccc(C#N)nc3)c1C2
CN1CCN(CCCNc2cccc(-c3nc4c(C(N)=O)cccc4[nH]3)n2)CC1
CN1CCN(CCNc2cccc(-c3nc4c(C=O)cccc4[nH]3)n2)CC1
CC(C)OC1=NNC(=O)C1=Cc1c[nH]c2ccccc12
CN(C)CC(=O)Nc1n[nH]c2ccc(-c3nnn(Cc4ccccc4)n3)cc12
CN(Nc1n[nH]c2ccc(-c3cn(Cc4ccccc4)nn3)cc12)C(=O)CO
CC(C)CC(=O)Nc1n[nH]c2ccc(-c3cn(Cc4ccccc4)nn3)cc12
CC(Nc1n[nH]c2ccc(-c3cn(Cc4ccccc4)nn3)cc12)C(=O)CO
```

```
Cc1ccccc1NC(=O)Nc1ccc(NC(=O)c2csc3ncnc(N)c23)cc1
CC1=C2C(=CC=C2NC(=O)c2csc3ncnc(N)c23)NC(=O)NC=CC=CC=C1
CC1=C2C(=CC=C2NC(=O)c2csc3ncnc(N)c23)NC(=O)CC=CC=CC=C1
Cc1ccccc1N(N=O)Nc1ccc(NC(=O)c2csc3ncnc(N)c23)cc1
Nc1ccccc1NC(=O)Nc1ccc(NC(=O)c2csc3ncnc(N)c23)cc1
CCOc1nc(C(=O)NCc2ccccc2[SH](N)(=O)CO)cc(N)c1C#N
CCOc1nc(C(=O)NCc2ccccc2[SH](N)(=O)NO)cc(N)c1C#N
COOc1nc(C(=O)NCc2ccccc2SC(=O)O)cc(N)c1C#N
COOc1nc(C(NO)NCc2ccccc2S(=O)NO)cc(N)c1C#N
Nc1nccc2scc(-c3ccc(NC(=O)Nc4cccc(F)c4)cc3)c12
Nc1cccc2scc(-c3ccc(NC(=O)Nc4cccc(Br)c4)cc3)c12
Nc1ccc2csc(ccn1)c1c(NC(=O)Nc3cccc(Br)c3)ccc21
Cc1ccc(C(=O)Nc2ccon2)cc1Nc1ncnc2c1cnn2-c1ccccc1
Cc1ccc(C(=O)NC2=CC=C2)cc1Nc1ncnc2c1cnn2-c1ccccc1
CCCCn1c(NC(=O)c2cccc(C#N)c2)nc2cc(N(C)C(OO)C3CCCCC3)ccc21
CCCCn1c(NC(=O)c2cccc(C#N)c2)nc2cc(N(N)C(CO)C3CCCCC3)ccc21
CCCCn1c(NC(=O)c2cccc(C#N)c2)nc2cc(N(N)C(OO)C3CCCCC3)ccc21
CCCCn1c(NC(=O)c2cccc(C#N)c2)nc2cc(N(O)C(OO)C3CCCCC3)ccc21
CCCCn1c(NC(=O)c2cccc(C#N)c2)nc2cc(N(N)C(=O)C3CCCCC3)ccc21
Nc1ncnc2scc(-c3ccc(NC(=O)Nc4ccccc4C(F)(F)F)cc3)c12
COc1ccc2c(NC(=O)Nc3cccc(Br)n3)ccnc2c1
COc1ccc2c(NC(=O)Nc3cccc(Br)c3)ccnc2c1
COc1ccc2cc(Br)cccc3[nH]c(=O)[nH]c-2ccnccn1-3
COc1ccc2c(NC(=O)Nc3cccc(Br)n3)c-2ncnc1
COc1ccc2c(NC(=O)Nc3cccc(Br)c3)c-2ncnc1
COC1=CC=C2C(=C2NC(=O)Nc2cccc(Br)n2)N=CC=NC=C1
COC1=CC=C2C(NC(=O)Nc3cccc(Br)n3)=CC=NC=CN=C12
COc1cccnc2c(NC(=O)Nc3cccc(Br)n3)c-2cc1
COc1ccc2c(NC(=O)Nc3cccc(Br)n3)ccnc2n1
BBc1cccc(NC(=O)Nc2ccnc3cc(OC)ccc23)n1
COc1cccnc2c(NC(=O)Nc3cccc(Br)c3)c-2cc1
COc1ccccccc2[nH]c(=O)[nH]cccccc(Br)cc-2cc1
COc1ccccc2c(NC(=O)Cc3cccc(Br)n3)c-2cc1
COc1ccc2c(NC(=O)Nc3cccc(Br)c3)cccc2c1
COc1ccccc2c(NC(=O)Nc3cccc(Br)n3)c-2cc1
```

COc1ccc2c(NC(=O)Nc3cccc(Br)n3)cccc2c1
Cc1cc(OCC2CCC(C(N)=O)C2)c(NC(=O)Nc2ccc(C#N)nc2)nc1Cl
CC1=CC(OCC2CCC(C(=O)O)C2)=C(NC(=O)Nc2ccc(C)cn2)NNC#CC1
CCc1cc(NC(=O)Nc2cnc(C#N)cn2)c(OCC2CCC(C(=O)O)C2)cc1C
Cc1ccc(NC(=O)Nc2cnc(C#N)cn2)c(OCC2CCC(C(=O)O)C2)c1
Cc1cc(NC(=O)Nc2cnc(C#N)cn2)c(OCC2CCC(C(N)=O)C2)cc1C
Cc1cc(OCC2CCC(C(=O)O)C2)c(NC(=O)Nc2cnc(C#N)cn2)nc1Cl
Cc1cc(OCC2CCC(C(N)=O)C2)c(NC(=O)Nc2cnc(C#N)cn2)nc1Cl
Cc1cc(NC(=O)Nc2cnc(C#N)n2C)c(OCC2CCC(C(=O)O)C2)cc1C
Cc1cc(NC(=O)Nc2cnc(C#N)cn2)c(OCC2CCC(C(=O)O)C2)cc1C
CC1=CCNNC(NC(=O)Nc2cnc(C)cn2)=C(OCC2CCC(C(=O)O)C2)C=C1C
CCc1cc(CC(=O)Nc2cnc(CCN)cn2)c(OCC2CCC(C(=O)O)C2)cc1C
Cc1cc(OCC2CCC(C(N)=O)C2)c(NC(=O)Nc2cnc(C#N)cn2)cc1Cl
Cc1ccc(NC(=O)Nc2ccc(C#N)cn2)c(OCC2CCC(C(N)=O)C2)c1
Cc1cc(NC(=O)Nc2ccc(C#N)cn2)c(OCC2CCC(C(=O)O)C2)cc1C
Cc1cc(NC(=O)Nc2ccc(C#N)cn2)c(OCC2CCC(C(N)=O)C2)cc1C
Cc1cc(OCC2CCC(C(=O)O)C2)c(NC(=O)Nc2ccc(C#N)cn2)cc1Cl
Cc1ccc(NC(=O)Nc2cnc(C#N)cn2)c(OCC2CCC(C(N)=O)C2)c1
Cc1cc(OCC2CCC(C(=O)O)C2)c(CC(=O)Nc2cnc(C#N)cn2)cc1Cl
CCc1cc(CC(=O)Nc2cnc(C#N)cn2)c(OCC2CCC(C(=O)O)C2)cc1C
Cc1cc(OCC2CCC(C(=O)O)C2)c(NC(=O)Nc2cnc(C#N)cn2)nc1C
CCc1cc(NC(=O)Nc2ccc(CCN)nc2)c(OCC2CCC(C(=O)O)C2)cc1C
O=C(O)C1CN(Cc2ccc(OCc3ccc(Cl)c(Cl)c3)cc2)C1
CC(C)C[C@H](NC(=O)[C@H](Cc1ccccc1)NC(=O)c1cnccn1)B(O)O
CC(C)C[C@H](NC(=O)[C@H](Cc1ccccc1)NC(=O)c1ccccn1)B(O)O
CC(C)C[C@H](NC(=O)[C@H](Cc1ccccc1)NC(=O)c1cccnc1)B(O)O
O=c1cc(-c2ccccc2)oc2ccccc12
OCC1C=C(c2ccccc2)Oc2ccccc21
O=C1CCC1Oc1ccc2cc-2cnccccnc1
O=C1CCC(c2ccccc2)Oc2ccccc21
O=c1nc(-c2ccccc2)oc2ccccc12
O=C1C=C2OC3=CC=C1C3=C1C=CC2=C1
O=c1cc(-c2ccccc2)oc2cc(O)cc(O)c12
O=c1cc(-c2ccccooc2)oc2cc(O)cc(O)c12
O=C1CCC(c2ccccc2)Oc2cc(O)cc(O)c21

```
O=c1cc(C2=CC=CC=CC=C2)oc2cc(O)cc(O)c12
O=c1cc(C2=COC=CC=C2)oc2cc(O)cc(O)c12
O=c1c(O)c(-c2ccc(O)cc2)oc2cc(O)cc(O)c12
O=c1c(O)c(-c2ccc(O)cc2)oc2c1OOCC=CC(O)=C2
O=c1c(O)c(-c2ccc(O)cc2)oc2c1C(O)CC(O)=C2
O=c1cc(-c2ccc(O)c(O)c2)oc2cc(O)cc(O)c12
Oc1cc(O)c2c(c1)OC(c1ccc(O)c(O)c1)=CC(O)C2
C[C@]12CC[C@@H]3c4ccc(O)cc4CC[C@H]3[C@@H]1CC2
O=C1COC(c2ccc(O)cc2)Oc2cc(O)cc(O)c21
OOC1C=C(c2ccc(O)cc2)Oc2cc(O)cc(O)c21
OCC1C=C(c2ccc(O)cc2)Oc2cc(O)cc(O)c21
O=C1C=C(c2ccc(O)cc2)c2ccocccc(O)c21
O=c1cc(-c2ccc(O)cc2)cc2oc(O)cc(O)c1-2
O=C(O)c1ccccc1Nc1ccnc(Nc2ccc3cn[nH]c3c2)n1
O=C(O)c1ccccc1Nc1ccnc(Nc2ccc3cn[nH]c3c2)c1
O=C(O)c1ccccc1Nc1ccnc(Nc2ccc3nn[nH]c3c2)n1
OCC(O)c1ccccc1Nc1ccnc(Nc2ccc3cn[nH]c3c2)n1
O=C(O)c1ccccc1Nc1ccnc(Nc2ccc3[nH]ncc3c2)c1
O=C(O)c1ccccc1Nc1ccnc(Nc2ccc3[nH]ncc3c2)n1
O=C(O)c1ccccc1Nc1ccnc(Nc2ccccc3[nH]c-3c2)n1
Cc1cccc(NC(=O)Nc2ccc(-c3csc4c(-c5cnn(CC(C)O)c5)cnc(N)c34)cc2)c1
Cc1cccc(NC(=O)Nc2ccc(-c3csc4c(-c5ccn(CC(C)O)c5)cnc(N)c34)cc2)c1
Cc1cccc(NC(=O)Cc2ccc(-c3csc4c(-c5cnn(CC(C)O)c5)cnc(N)c34)cc2)c1
COCCOC1CCC(n2nc(-c3ccc(Nc4nc5cc(C)cc(C)c5o4)cc3)c3c(N)ncnc32)CC1
COCCOC1CCC(n2nc(-c3ccc(Nc4cc5cc(C)cc(C)c5o4)cc3)c3c(N)ncnc32)CC1
COCCOC1CCC(n2nc(-c3ccc(Nc4nc5cc(O)cc(C)c5o4)cc3)c3c(N)ncnc32)CC1
COc1cc(Nc2nccc(Nc3cc(C4CCCCC4)no3)n2)cc(OC)c1OC
COc1cc(Nc2nccc(Nc3cc(C4CCCCC4)no3)n2)cc(OC)c1
COc1cc(Nc2cccc(Nc3cc(C4CCCCC4)no3)n2)cc(OC)c1OC
COc1cc(Nc2nccc(Nc3cc(C4CCCCC4)no3)n2)cc(OO)c1
COc1cc(Nc2nccc(Nc3cc(C4CCCCC4)no3)n2)cc(OC)c1O
COc1cc(Nc2cc(Nc3cc(C4CCCCC4)no3)ccn2)cc(OC)c1O
COc1cc(Nc2cc(Nc3cc(C4CCCCC4)no3)ccn2)cc(OC)c1
COc1cc(Nc2cccc(Nc3cc(C4CCCCC4)no3)c2)cc(OC)c1OC
COc1cc(Nc2ncnc(Nc3cc(C4CCCCC4)no3)n2)cc(OC)c1
```

```
COc1cc(Nc2cc(Nc3cc(C4CCCCC4)no3)ccn2)cc(OC)c1OC
COc1cc(Nc2nccc(Nc3cc(C4CCCCC4)no3)n2)cc(OO)c1OC
COc1cc(Nc2cccc(Nc3cc(C4CCCCC4)no3)n2)cc(OC)c1
CCc1c(OC)cc(Nc2nccc(Nc3cc(C4CCCCC4)no3)n2)cc1OC
COc1cc(Nc2cccc(Nc3cc(C4CCCCC4)no3)c2)cc(OO)c1OC
CCCC(=O)Nc1n[nH]c2ccc(-c3nnn(Cc4ccccc4)c3-c3ccccc3)cc12
CCCC(=O)Nc1n[nH]c2ccc(-c3ncn(Cc4ccccc4)c3-c3ccccc3)cc12
CCCC(=O)Nc1n[nH]c2ccc(-c3nnn4c3C=CC=CC=CC=C3C=CC(=C3)C4)cc12
O=C1NCCc2[nH]c(-c3cccc(C=Cc4ccccc4)c3)cc21
O=C1NCCc2[nH]c(-c3ccnc(CNCc4ccccc4)c3)cc21
Fc1ccc(-c2nc3sncn3c2-c2ccncc2)cc1
Fc1ccc(-c2nc3sccn3c2-c2ccccc2)cc1
Fc1ccc(C2=S(C3=CC=CC=CC=N3)N3C=CSC3=N2)cc1
Fc1ccc(-c2nc3sccn3c2-c2ccncc2)cc1
Fc1ccc(-c2ccccccccccn3ccsc3n2)cc1
Fc1ccc(C2=NC3=C(C4=CC=C4)C=CN2C=CS3)cc1
COc1nccn2c(-c3ccnc(NCC(C)(C)O)n3)c(-c3ccc(F)cc3F)nc12
Cc1nccn2c(-c3cccc(NCC4(C)CC4)n3)c(-c3ccc(F)cc3)nc12
CCc1nccn2c(-c3ccnc(NCC(C)(C)C)n3)c(-c3ccc(F)cc3F)nc12
O=C1NC(=O)C(c2ccc3ccccn23)=C1c1cn2c3c(cccc13)CN(C(=O)N1CCCCC1)CC2
O=C1NC(=O)C(c2cnc3ccccn23)=C1c1cn2c3c(cccc13)CN(C(=O)N1CCCCC1)CC2
O=C1NC(=O)C(c2cnc3ccccn23)=C1c1cn2c3c(cccc13)CN(C(=O)N1CCOCC1)CC2
O=C1NC(=O)C(c2cnc3ccccn23)=C1c1cn2c3c(cccc13)CN(C(=O)C1CCCCN1)CC2
Fc1ccc(-c2nc3cnccn3c2-c2ccnc(NCC3CC3)n2)cc1
CC(Nc1nccc(C2=S(c3ccc(F)cc3)N=C3OC=CN32)n1)c1ccccc1
OC1=CC=CC2=CC=CC=Cc3c2ccnc3C=CC=CN=CC=CC=CC=C1
CCc1nc(-c2cccc(C)c2)c(-c2ccnc(NC(NO)c3ccccc3)c2)s1
Cc1cccc(C2=C(c3ccnc(NC(=O)c4ccccc4)c3)SC(C)C=N2)c1
Cc1cccc(C2=C(c3cccc(NC(=O)c4ccccc4)c3)SC(C)C=N2)c1
O=C(NC(CO)c1cc(Cl)cc(Cl)c1)c1ccc(-c2ccncc2)cc1
O=C(NC(CO)C1=CC(Cl)=C1)c1ccc(-c2ccncc2)cc1
COc1cc(-c2ccc3c(c2)Nc2ccc(C(=O)NCc4cccc(F)c4)cc2NC3=O)ccc1O
C=C(NCc1cccc(F)c1)c1ccc2c(c1)NC(=O)c1ccc(-c3ccc(O)c(OC)c3)cc1N2
COc1cc(-c2ccc3c(c2)Nc2ccc(C(=O)NCc4cccc(N)c4)cc2NC3=O)ccc1O
C=C(NC1C=CC=CC(F)=C1)c1ccc2c(c1)NC(=O)c1ccc(-c3ccc(O)c(OC)c3)cc1N2
```

```
COc1cc(C2=CC=C3C(=O)Nc4cc(C(=O)NCc5cccc(F)c5)ccc4C=C3N2)ccc1O
COc1cc(-c2ccc3c(c2)Nc2ccc(C(=O)NC4C=CC=CC(F)=C4)cc2NC3=O)ccc1O
COc1cc(C2=CC=C3C(=O)Nc4cc(C(=O)NCc5cccc(F)c5)ccc4C=C3C2)ccc1O
COc1cc(C2=CC=C3C(=O)Nc4cc5ccc4C=C3C2(C=O)C=C(F)C=CC=CC=C5)ccc1O
COc1cc(-c2ccc3c(c2NC=O)Nc2ccc(Cc4cccc(F)c4)cc2NC3=O)ccc1O
COc1cc(-c2ccc3c(c2)Nc2ccc(C(=O)NCc4cccc(O)c4)cc2NC3=O)ccc1O
Cc1nccn2c(-c3ccnc(NCC(C)(C)C(=O)O)n3)c(-c3ccc(F)cc3F)nc12
CC(=O)C(C)(C)CNc1nccc(-c2c(-c3ccc(F)cc3F)nc3c(C)nccn23)n1
CC(C)(O)CNc1nccc(-c2c(-c3ccc(F)cc3F)nc3c(CC4CC4)nccn23)n1
CC(C)(O)CNc1nccc(-c2c(-c3ccc(F)cc3)nc3c(CC4CC4)nccn23)n1
Cc1nccn2c(-c3ccnc(NCCC(C)(C)O)n3)c(-c3ccc(F)cc3F)nc12
CN(C)CC(C)(C)CNc1nccc(-c2c(-c3ccc(F)cc3)nc3occn23)n1
CC(C)CC(C)(C)CNc1nccc(-c2c(-c3ccc(F)cc3)nc3occn23)n1
CN(C)CC(C)(C)CNc1nccc(-c2cnnc3c(-c4ccc(F)cc4)ccn23)n1
CN(C)CC(C)(C)CNc1nccc(-c2c(-c3ccc(F)cc3)nc3cnccn23)n1
CCc1ncc(OCC(C)N)cc1C=Cc1ccncc1
CC(N)COc1cnc(Cl)c(C=Cc2ccncc2)c1
CC(N)COc1cnc(Cl)c(CCCc2ccncc2)c1
CC(N)CCc1cnc(Cl)c(N=Cc2ccncc2)c1
CNc1ncc(OCC(C)N)cc1N=Cc1ccncc1
CC(N)COc1cnc(Cl)c(C=Cc2ccncn2)c1
Cc1ccc(-c2nn(C(C)(C)C)c3ncnc(N)c23)cc1
Cc1ccc(-c2nn(C(C)(C)O)c3ncnc(N)c23)cc1
Cc1ccc(-c2nn(C(C)(N)O)c3ncnc(N)c23)cc1
Cc1ccc(-c2nn(C(C)C)c3ncnc(N)c23)cc1
Cc1ccc(-c2nn(C(C)(C)N)c3ncnc(N)c23)cc1
Cc1ccc(-c2nn(C(C)(C)C)c3nccc(N)c23)cc1
COc1cc(-c2ccc3c(c2)Nc2ccc(C(=O)NCCNC(C)=O)cc2NC3=O)ccc1O
CNC(C)NCCNC(=O)c1ccc2c(c1)NC(=O)c1ccc(-c3ccc(O)c(OC)c3)cc1N2
CC=NCCNCCNC(NO)c1ccc2c(c1)NC(=O)c1ccc(-c3ccc(O)c(OC)c3)cc1N2
COc1cc(-c2ccc3c(c2)Nc2ccc(C(=O)NCCNC(=O)O)cc2NC3=O)ccc1O
COc1cc(-c2ccc3c(c2)Nc2ccc(C(=O)NCCNC(C)NO)cc2NC3=O)ccc1O
COc1cc(-c2ccc3c(c2)Nc2ccc(C(=O)NCCCC(C)=O)cc2NC3=O)ccc1O
COc1cc(-c2ccc3c(c2)Nc2ccc(C(NO)NCCCCC=O)cc2NC3=O)ccc1O
COc1cc(-c2ccc3c(c2)Nc2ccc(C(=O)NCNNCC=O)cc2NC3=O)ccc1O
```

```
COc1cc(-c2ccc3c(c2)Nc2ccc(C(=O)NCCNC=O)cc2NC3=O)ccc1O
COc1cc(-c2ccc3c(c2)Nc2ccc(C(=O)NCCCC(C)NO)cc2NC3=O)ccc1O
CC=CNOCC(N=N)c1ccc2c(c1)NC(=O)c1ccc(-c3ccc(O)c(OC)c3)cc1N2
COc1cc(-c2ccc3c(c2)Nc2ccc(C(=O)NCCNCCO)cc2NC3=O)ccc1O
CNCNCCNC(=O)c1ccc2c(c1)NC(=O)c1ccc(-c3ccc(O)c(OC)c3)cc1N2
COc1cc(-c2ccc3c(c2)Nc2ccc(C(=O)NCNNC(C)=O)cc2NC3=O)ccc1O
COc1cc(-c2ccc3c(c2)Nc2ccc(C(NO)NCCNCC(C)O)cc2NC3=O)ccc1O
COc1cc(-c2ccc3c(c2)Nc2ccc(C(=O)NCCNCC=O)cc2NC3=O)ccc1O
COc1cc(-c2ccc3c(c2)Nc2ccc(C(=O)NCCCC=O)cc2NC3=O)ccc1O
COc1cc(-c2ccc3c(c2)Nc2ccc(C(=O)NCCNCC(C)O)cc2NC3=O)ccc1O
Cc1nccn2c(-c3ncnc(NCCC(C)(C)O)n3)c(-c3ccc(F)cc3F)nc12
COc1cc(C=C2CNC3C=CC=CC=C3N=C(O)S2)ccc1O
Cc1n[nH]c2ccc(-c3cncc(OCC(N)Cc4cccc(OC(F)(F)F)c4)c3)cc12
Cc1n[nH]c2ccc(C3=CC4(N)C=CC=C(CC(N)COC=CN=C3)C(F)=C4)cc12
Cc1n[nH]c2ccc(-c3cncc(OCC(N)Cc4cccc(NC(F)(F)F)c4)c3)cc12
Nc1ncnc2sc3c(c12)-c1ccc(NC(=O)Nc2ccccc2)cc1CC3
Nc1ncnc2c1=C1C=CC=CC(=CC3=C(C=2)CC3)[N+]1([O-])Nc1ccccc1
Nc1ncnc2sc3c(c12)C=CC1=C(NC(=O)Nc2ccccc2)C=C1CC3
Nc1ncnc2sc3c(c12)C=CC=CC(NC(=O)Nc1ccccc1)=CN=CCC3
CN(C)c1cccc2c(S(=O)(=O)N(CCN)c3cncc(-c4ccc5cnccc5c4)c3)cccc12
CN(C)c1cccc2c([SH](=O)(OO)N(NCN)c3cncc(-c4ccc5cnccc5c4)c3)cccc12
CN(C)c1cccc2c(S(=O)(=O)N(CCN)c3cncc(-c4ccc5ccccc5c4)c3)cccc12
CN(C)c1cccc2c([SH](=O)(OO)N(CCN)c3cncc(-c4ccc5cnccc5c4)c3)cccc12
CN(C)c1cccc2c(S(=O)(=O)N(CCN)c3cncc(C4=CC=C5C6=CC6=C54)c3)cccc12
CN(C)c1cccc2c(S(=O)(=O)N(NCN)c3cncc(-c4ccc5cnccc5c4)c3)cccc12
CN(C)c1cccc2c(S(=O)(=O)N(CCN)c3cccc(-c4ccc5cnccc5c4)c3)cccc12
Cc1cccc(NC(=O)Nc2ccc(NC(=O)c3csc4ncnc(N)c34)cc2)c1
Cc1cccc(NC(=O)Nc2ccc(NC(=O)c3csc4ccnc(N)c34)cc2)c1
Cc1cccc[nH]c(=O)[nH]c2ccc(NC(=O)c3csc4ncnc(N)c34)c1-2
Cc1cccc(NC(=O)NC2=CC=c3[nH]c(=O)c4c(c32)C(N)=C2N=C2SC=4)c1
Cc1cccc(CC(=O)Nc2ccc(NC(=O)c3csc4ccnc(N)c34)cc2)c1
Cc1cccc(NC(=O)Nc2ccc(NC(NO)c3csc4ncnc(N)c34)cc2)c1
Cc1cc2c(-c3ccnc(NCC(C)(C)CO)n3)c(-c3ccc(F)cc3F)nn2c(C)n1
Cc1cn2c(-c3ccnc(NCC(C)(C)CO)n3)c(-c3ccc(F)cc3)nc2c(C)n1
O=C(NCc1ccccc1)c1ccc(-c2ccncc2)cc1
```

```
O=C(CC(CO)c1ccccc1)c1ccc(-c2ccncc2)cc1
Cc1n[nH]c2ccc(-c3cncc(OCCN)c3)cc12
Cc1n[nH]c2ccc(-c3cncc(CCCN)c3)cc12
Cc1n[nH]c2ccc(-c3cncc(OCCO)c3)cc12
Cc1n[nH]c2ccc(C3=CNNCCONC=CN=C3)cc12
Cc1n[nH]c2ccc(C3=CNNCCOOC=CN=C3)cc12
COc1nccn2c(-c3ccnc(CCC(C)(C)CO)n3)c(-c3ccc(F)cc3F)nc12
CC1=C=c2ccccc2=CC=CC=CC=CC=NC=N1
O=C(NCc1ccccc1)c1ccc(-c2ccccc2)cc1
OCC(NCc1ccccc1)c1ccc(-c2ccncc2)cc1
O=C(NCc1ccccc1)c1ccccc1-c1ccncc1
O=C(CCc1ccccc1)c1ccc(-c2ccncc2)cc1
OC1CCC(Nc2cc(Cl)nc(-c3c[nH]c4ccccc34)n2)CC1
OC1CCC(Nc2cc(Cl)cc(-c3c[nH]c4ccccc34)n2)CC1
OC1CCC(Nc2cc(Cl)cc(-c3c[nH]c4ncccc34)n2)CC1
OC1CCC(Nc2cc(Cl)nc(-c3c[nH]c4ccccc34)c2)CC1
Fc1ccc(-c2nc3cnccn3c2-c2ccnc(NCC3CC3)n2)nc1
CCNC1=NC=CN=CC=CN=c2c(-c3cccc(F)c3)nccc2=NC1
Cc1n[nH]c2ccc(-c3cncc(OCC(N)Cc4cccc(Cl)c4)c3)cc12
CCc1cccc(CC(N)COc2cncc(-c3ccc4[nH]nc(C)c4c3)c2)c1
Cc1n[nH]c2ccc(C3=C4C=CC=C(Cl)C=CC=C4CC(N)COC=CN=C3)cc12
Cc1n[nH]c2ccc(-c3cncc(OCC(N)Cc4cccc(Cl)c4)c3N)cc12
CC(NC(=O)c1ccc(-c2ccncc2)cc1)c1ccccc1
CC(NC(=O)c1ccc(-c2ccccc2)cc1)c1ccccc1
CC1NC(=O)c2ccc(cc2)C2=CC2=CC=C2C=CC1=C2
CC(NC(=O)c1ccc(-c2ccncc2)cn1)c1ccccc1
CCCNC(=O)C1(c2ccccc2)C=CC(c2ccncc2)=C1
NC(COc1cncc(-c2ccc3c(c2)C(=Cc2ccco2)C(=O)N3)c1)Cc1c[nH]c2ccccc12
NC(COc1cncc(-c2ccc3c(c2)C(=Cc2ccco2)C(=O)N3)c1)Cc1cccccccc1
Clc1cc(-c2c[nH]c3ncccc23)cc(NCc2ccccc2)n1
Clc1cc(-c2c[nH]c3ncccc23)cc(NCc2cccnc2)n1
Clc1cc(-c2c[nH]c3ncccc23)cc(NCc2ccccn2)n1
Clc1cc(-c2c[nH]c3ncccc23)cc(NC2C=CC=CC=N2)n1
Clc1cc(NCc2ccccc2)cc(-c2c[nH]c3ncccc23)c1
Clc1cc(C2=CNC=NN=CC=CC=N2)cc(NCc2ccccc2)n1
```

```
Clc1cc(C2=CNC=NN=CC=CC=C2)cc(NCc2ccccc2)n1
O=c1[nH]c2cnc(-n3cnc4ccc(F)cc43)nc2n1C1CCOc2c(F)cccc21
O=c1[nH]c2ccc(-n3cnc4ccc(F)cc43)cc2n1C1CCOc2c(F)cccc21
NCC1Nc2cnc(-n3cnc4ccc(F)cc43)cc2N1C1CCOc2c(F)cccc21
Cc1ccc(-c2nc(-c3ccc(F)cc3)c(-c3ccccc3)[nH]2)cc1
Fc1ccc(-c2nc(-c3ccc(F)cc3)c(-c3ccccc3)[nH]2)cc1
Fc1ccc(-c2nc(-c3ccncc3)c3ccc(F)cc3ccn[nH]2)cc1
Nc1nccc(Nc2cc(CNC(=O)C(F)(F)F)c(O)c(-c3ccc(Cl)cc3)c2)n1
O=C1NCCc2[nH]c(-c3ccnc(-c4ccccc4F)c3)cc21
O=C1NCCc2[nH]c(-c3cccc(-c4ccccc4F)c3)cc21
O=C1CCCc2[nH]c(-c3ccnc(-c4ccccc4F)c3)cc21
OCC1NCCc2[nH]c(-c3ccnc(-c4ccccc4F)c3)cc21
O=C1NCCc2[nH]c(-c3ccnc(-c4cncnc4F)c3)cc21
COc1cc(Nc2ncc3c(n2)-c2ccc(Cl)cc2C(c2c(F)cccc2OC)=NC3)ccc1C(=O)O
COc1cc(Nc2ncc3c(n2)-c2ccc(Cl)cc2C(c2c(F)cccc2OC)=NN3)ccc1C(=O)O
COc1cc(Nc2ncc3c(n2)-c2ccc(Cl)cc2C(c2c(F)cccc2OC)ONC3)ccc1C(=O)O
COc1cc(Nc2ncc3c(n2)-c2ccc(CO)cc2C(c2c(F)cccc2OC)=NC3)ccc1C(=O)O
CCc1ccc2c(c1)C(c1c(F)cccc1OC)=NCc1cnc(Nc3ccc(C(O)CO)c(OC)c3)cc1-2
CCc1nccn2c(-c3cccc(CCC(C)(C)CO)n3)c(-c3ccc(F)cc3F)nc12
CCc1nccn2c(-c3ccnc(NCC(C)(C)CO)n3)c(-c3ccc(F)cc3)nc12
COc1nccn2c(-c3ccnc(NCC(C)(C)CO)n3)c(-c3ccc(F)cc3)nc12
CCc1nccn2c(-c3ccnc(NCC(C)(C)OO)n3)c(-c3ccc(F)cc3F)nc12
COc1nccn2c(-c3cccc(NCC(C)(C)CO)n3)c(-c3ccc(F)cc3F)nc12
Cc1n[nH]c2ccc(-c3cncc(OCC(N)Cc4ccc(Cl)c(Cl)c4)c3)cc12
CCc1ccc(CC(N)COc2cncc(-c3ccc4[nH]nc(C)c4c3)c2)cc1Cl
C=C(C=Cc1cnc(N)c2c(-c3ccc(Br)cc3)csc12)NCCCn1ccnc1
Nc1ncc(C=NC(=O)NCCCn2ccnc2)c2scc(-c3ccc(Br)cc3)c12
Nc1ncc(C=CC(=O)NCCCn2cncn2)c2scc(-c3ccc(Br)cc3)c12
Nc1ncc(C=CC(=O)NCCCn2ccnc2)c2scccc3ccc(Br)cc3cc12
Clc1cc(NC2CCCCC2)nc(-c2c[nH]c3ncccc23)n1
Clc1cc(NC2CCCCC2)nc(-c2c[nH]c3ncccc23)c1
Clc1cc(NC2CCCCC2)nc(-c2c[nH]c3nnccc23)n1
CC(C)(CO)c1ncc2ccc(-c3c(-c4ccc(F)cc4F)nc4occn34)cn12
CC(C)(CO)CNc1nccc(-c2c(-c3ccc(F)cc3)nc3c(CCN)nccn23)n1
CSc1ccc(-c2nc(-c3ccc(F)cc3)c(-c3ccncc3)[nH]2)cc1
```

```
C[SH]1C=CC=C(c2nc(-c3ccc(F)cc3)c(-c3ccncc3)[nH]2)C=C1
CC(C)Cc1nc2sc3c(NCc4ccco4)ncnc3c2c2c1COC(C)(C)C2
CC(C)Cc1nc2sc3c(NCc4ccco4)ncnc3c2c2c1CCC(C)(C)C2
CCc1c2c(cc3ccc(O)cc13)-c1cc3c(c(=O)n1C2)COC(=O)[C@]3(O)CC
CCc1c2c(nc3ccc(O)cc13)-c1cc3c(c(=O)n1C2)COC(=O)[C@]3(O)CC
CCc1c2c(nc3ccc(O)cc13)-c1cc3c(c(=O)n1C2)COC(=O)[C@]3(C)CC
CCc1c2c(nc3ccc(O)cc13)C1CCC3=C(COC(=O)[C@]3(O)CC)C(=O)N1C2
CCc1c2c(nc3ccc(O)cc13)C1OCC3=C(COC(=O)[C@]3(O)CC)C(=O)N1C2
CCc1c2c(cc3ccc(O)cc13)C1CCC3=C(COC(=O)[C@]3(O)CC)C(=N)N1C2
CCc1c2c(nc3ccc(O)cc13)-c1cc3c(c(=O)n1C2)CCC(=O)[C@]3(O)CC
CCc1c2c(nc3ccc(O)cc13)C1CCC3=C(COC(=O)[C@]3(C)CC)C(=N)N1C2
CCc1c2c(nc3ccc(O)cc13)-c1cc3c(c(=O)n1C2)CCC(=O)[C@]3(C)CC
CCc1c2c(cc3ccc(O)cc13)C1CCC3=C(COC(=O)[C@]3(O)CC)C(=O)N1C2
COc1ccc(CCN(C)CCCC(C#N)(c2ccc(OC)c(OC)c2)C(C)C)cc1O
COc1ccc(CCN(C)CCCC(C#N)(c2ccc(OC)c(OC)c2)C(C)C)cc1OC
CCc1ccc(C(C#N)(CCCN(C)CCc2ccc(OC)c(OC)c2)C(C)C)cc1OC
CCc1cc(CCN(C)CCCC(C#N)(c2ccc(CC)c(OC)c2)C(C)C)ccc1OC
CCc1ccc(C(C#N)(CCCN(C)CCc2ccc(OC)c(O)c2)C(C)C)cc1OC
CCc1ccc(C(C#N)(CCCN(C)CCc2ccc(OC)cc2)C(C)C)cc1OC
COc1ccc(CCN(C)CCCC(CONCOC(C)C)c2ccc(OC)c(OC)c2)cc1OC
COc1ccc(CCN(C)CCCC(C#N)(c2ccc(OC)c(OC)c2)C(C)C)cc1C
CCc1cc(CCN(C)CCCC(CCOC(C)COC)c2ccc(CC)c(OC)c2)ccc1OC
Cc1cc(C#N)cc(C)c1Oc1nc(Nc2ccc(C#N)cc2)nc(N)c1Br
Cc1cc(C#N)cc(C)c1Oc1cc(Nc2ccc(C#N)cc2)nc(N)c1Br
Cc1cc(CNN)cc(C)c1Oc1nc(Nc2ccc(C#N)cc2)nc(N)c1Br
BC1=CC=NN=C(Nc2ccc(C#N)cc2)N=C1Oc1c(C)cc(CCN)cc1C
BC1=CC=CN=C(Nc2ccc(C#N)cc2)N=C1Oc1c(C)cc(OCN)cc1C
Bc1nc(Nc2ccc(C#N)cc2)nc(Oc2c(C)cc(OC#N)cc2C)c1B
CBc1c(N)nc(Nc2ccc(C#N)cc2)nc1Oc1c(C)cc(C#N)cc1C
Cc1cc(C#N)nc(C)c1Oc1nc(Nc2ccc(C#N)cc2)n(CN)c1Br
Bc1c(N)nc(Nc2ccc(C#N)cc2)nc1Oc1c(C)cc(OC#N)cc1C
BC1=CC=CN=C(Nc2ccc(C#N)cc2)N=C1Oc1c(C)cc(C#N)cc1C
Bc1c(N)nc(Nc2ccc(C#N)cc2)nc1Oc1c(C)cc(OCNN)cc1C
OC1CCC(Nc2nc(Cl)cc(-c3c[nH]c4ncccc34)n2)CC1
CC1CCC(Nc2nc(Cl)cc(-c3c[nH]c4ncccc34)n2)CC1
```

OC1=CCC(Nc2cc(Cl)cc(-c3c[nH]c4ncccc34)n2)CC1

CC1CCC(Nc2cc(Cl)cc(-c3c[nH]c4ncccc34)n2)CC1

OC1CCC(Nc2nc(Cl)cc(-c3c[nH]c4ncccc34)n2)CN1

CC1CCC(N2C=CC(Cl)=CC(c3c[nH]c4ncccc34)=N2)CC1

CC1CCC(N2N=CC(Cl)=CC(c3c[nH]c4ncccc34)=N2)CC1

NS(=O)(=O)c1cccc(Nc2ncc3ccn(-c4ccccc4)c3n2)c1

CS(=O)(=O)c1cccc(Nc2ncc3ccn(-c4ccccc4)c3n2)c1

CS(=O)(=O)c1cccc(Nc2cc3c(ccn3-c3ccccc3)cn2)c1

NS(=O)(=O)c1cccc(Nc2cc3c(ccn3-c3ccccc3)cn2)c1

NS(=O)(=O)c1cccc(Nc2ncc3ccn(-c4ccccn4)c3n2)c1

Cc1cccc(Nc2nccc(Nc3ccccc3C(=O)O)n2)c1

O=C(O)c1ccccc1Nc1ccnc(Oc2cccc(O)c2)n1

Cc1cccc(Nc2cc(Nc3ccccc3C(=O)O)ccn2)c1

O=C(O)c1ccccc1Nc1ccnc(Nc2cccc(O)c2)c1

O=C(O)c1cccnc1Nc1ccnc(Nc2cccc(O)c2)n1

O=C(O)c1ccccc1Nc1ccnc(Oc2cccc(O)c2)c1

O=C(O)c1ccccc1Nc1cccc(Nc2cccc(O)c2)n1

CC(C)(C)c1cc(NC(=O)c2cccc(Oc3cccc4[nH]c(=O)[nH]c34)c2)n(-c2ccccc2)n1

O=c1[nH]c(CN2CC[C@H](O)C2)nc2c1sc1ccc(-c3ccc(O)cc3)cc12

Cc1cnc(Nc2cccc(S(N)(=O)=O)c2)nc1Nc1ccc(ONC(N)=O)cc1

Cc1cnc(Nc2cccc(S(N)(=O)=O)c2)nc1Nc1ccc(OCC(N)=O)cc1

CC1=CN=C(Nc2cccc(S(=N)(N)=O)c2)NC12C=CN=C(COCC(N)=O)C=C2

CC(C)(C(=N)N)n1cc(-c2nnc(N)c3c(-c4ccc(NC(=O)Nc5cccc(F)c5)cc4)csc23)cn1

CC(C)(C(N)=O)n1cc(-c2cnc(N)c3c(-c4ccc(NC(=O)Nc5cccc(F)c5)nc4)csc23)cn1

CC(C)(C(N)=O)n1cc(-c2nnc(N)c3c(-c4ccc(NC(=O)Nc5cccc(F)c5)cc4)csc23)cn1

C=C(N)C(C)(C)n1cc(-c2cnc(N)c3c(-c4ccc(NC(=O)Nc5cccc(F)c5)cc4)csc23)cn1

CC(C)(C(N)=O)n1cc(-c2cnc(N)c3c(-c4ccc(NCC(O)Nc5cccc(F)c5)cc4)csc23)cn1

CC(C)(C(=N)N)n1cc(-c2cnc(N)c3c(-c4ccc(NC(=O)Nc5cccc(F)c5)cc4)csc23)cn1

C=C(N)C(C)(C)n1cc(-c2nnc(N)c3c(-c4ccc(NC(=O)Nc5cccc(F)c5)cc4)csc23)cn1

CC(C)(C(N)=O)n1cc(-c2cnc(N)c3c(-c4ccc(NC(=O)Nc5cccc(F)c5)cc4)csc23)nn1

CC(C)(C(N)=O)n1cc(-c2cnc(N)c3c(-c4ccc(NC(Nc5cccc(F)c5)OO)cc4)csc23)cn1

CC(S)(C(N)=O)n1cc(-c2cnc(N)c3c(-c4ccc(NC(=O)Nc5cccc(F)c5)cc4)csc23)cn1

CNNC(=O)c1cnc(N)c2c(-c3ccc(NC(=O)Nc4cccc(F)c4)cc3)csc12

CCCC(=O)c1cnc(N)c2c(-c3ccc(NC(=O)Nc4cccc(F)c4)cc3)csc12

CCNC(=O)c1cnc(N)c2csc(-c3ccc(NC(=O)Nc4cccc(F)c4)cc3)c12

```
OCC(O)c1ccccc1Nc1ccnc(Nc2ccc3[nH]ncc3c2)n1
Oc1cccc(-c2nc(N3CCOCC3)c3oc4ncccc4c3n2)c1
Oc1cccc(-c2nc(N3CCCCC3)c3oc4ncccc4c3n2)c1
Cn1cc(-c2ccc3c(c2)CCN3C(=O)Cc2cccc(C(F)(F)F)c2)c2c(N)ncnc21
Cn1cc(-c2ccc3c(c2)CCN3C(=O)Cc2cccc(C(F)(F)F)c2)c2c(N)nncc21
Cn1cc(-c2ccc3c(c2)CCN3C(=O)Cc2cccc(C(F)(F)F)c2)c2c(N)nccc21
Cc1cnnnc(N)ccc(-c2ccc3c(c2)CCN3C(=O)Nc2cccc(C(F)(F)F)c2)c1
C[C@@H]1CCO[C@H]2Cn3cc(C(=O)NCc4ccc(F)cc4F)c(=O)cc3C(=O)N21
C[C@@H]1CCC[C@H]2Cn3cc(C(=O)NCc4ccc(F)cc4F)c(=O)c(O)c3C(=O)N21
C[C@@H]1CCO[C@H]2CC3C=C(C(=O)NCc4ccc(F)cc4F)C(=O)C(O)=C3C(=O)N21
Cc1cc(F)ccc1CNC(=O)c1cn2c(cc1=O)C(=O)N1[C@H](C2)OCC[C@H]1C
C[C@@H]1CCO[C@H]2CN3COC(C(=O)NCc4ccc(F)cc4F)C(=O)C(O)=C3C(=O)N21
C[C@@H]1CCO[C@H]2CN3C=C(C(=O)NCc4ccc(F)cc4F)C(CO)C(O)=C3C(=O)N21
CCCc1[nH]cnc1CNc1cc(Cl)c2ncc(C#N)c(Nc3ccc(F)c(Cl)c3)c2c1
N#Cc1cnc2c(Cl)cc(NCc3c[nH]nn3)cc2c1Nc1ccc(F)c(Cl)c1
CC(C)OC(=O)N1CCn2cc(C3=C(c4cnc5ccccn45)C(=O)NC3=O)c3cccc(c32)C1
O=C1NC(=O)C(c2cnc3ccccn23)=C1c1cn2c3c1=CC=CC=NC(C=3)N(C(=O)N1CCCCC1)CC2
O=C1NC(=O)C(c2cnc3ccccn23)=C1c1cn2c3c(cccc13)CC(C(=O)N1CCCCC1)CC2
N#Cc1cnc2cnc(NCc3cccnc3)cc2c1Nc1ccc(N)cc1
O=C1CCC(c2ccccc2)Oc2ccc3ccccc3c21
OCCCC=C(OC1=C=C=CC=CC=CC=CC=C1)c1ccccc1
O=C1C=C(c2ccccc2)Oc2ccc3c1ccccc2-3
O=C1CCC(c2ccccc2)Oc2ccc3ccnc-3cc21
O=C1CCC(c2ccccc2)Oc2ccc3ccc1c-3cc2
O=c1cc(-c2ccccc2)oc2c1-c1=c3-c=c-c-1=c-3-2
O=C1CCC(c2ccccc2)Oc2ncc3ccccc3c21
O=C(O)c1ccnc(C(=O)O)c1
Cc1cccc(Nc2nccc(Nc3ccccc3C(N)=O)n2)c1
NC(=O)c1ccccc1Nc1ccnc(Cc2cccc(O)c2)n1
NC(=O)c1ccccc1Nc1ccnc(Nc2cccc(O)c2)n1
NC(=O)c1ccccc1Nc1ccnc(Nc2cccc(O)c2)c1
COc1ccc(-n2cnc3c(sc4nccc(N(C)C)c43)c2=O)cc1
COc1ccc(N2C=Nc3c(sc4nccc(N(C)C)c34)C2)cc1
COc1ccc(-n2cnc3c(sc4nncc(N(C)C)c43)c2=O)cc1
CCCCCc1cccc2c1-c1ncn(-c3ccc(OC)cc3)c(=O)c1C2
```

```
CC(=O)N1c2ccc(-c3ccc(C(=O)O)cc3)cc2[C@H](Nc2ccc(Cl)cc2)C[C@@H]1C
CC(=O)N1c2ccc(-c3cccc(C(O)OO)c3)cc2[C@H](Nc2ccc(Cl)cc2)C[C@@H]1C
CC(=O)N1c2ccc(-c3ccc(C=O)cc3)cc2[C@H](Nc2ccc(Cl)cc2)C[C@@H]1C
CC(=O)C1c2ccc(-c3ccc(C(=O)O)cc3)cc2[C@H](Nc2ccc(Cl)cc2)C[C@@H]1C
CC(=O)N1c2ccc(-c3cccc(C(=O)O)c3)cc2[C@H](Nc2ccc(Cl)cc2)C[C@@H]1C
COc1cc2c(cc1OC)CN(CCc1ccc(NC(=O)c3ccccc3NC(=O)c3cnc4ccccc4c3)cc1)CC2
COc1cc2c(cc1O)CN(CCc1ccc(NC(=O)c3ccccc3NC(=O)c3cnc4ccccc4c3)cc1)CC2
CCOc1cc2c(cc1OC)CCN(CCc1ccc(NC(=O)c3ccccc3NC(=O)c3cnc4ccccc4c3)cc1)C2
COc1cc(OC)c2c(c1)O[C@H](c1cc(OC)c(OC)c(OC)c1)[C@@H](OC(=O)c1ccc(F)c(NC(=O)c3cc(OC)c(O[illegible]
CNc1cc(OC)c2c(c1)O[C@H](c1cc(OC)c(OC)c(OC)c1)[C@@H](OC(=O)c1ccc(F)c(NC(=O)c3cc(OC)c(O[illegible]
COc1ccc(-c2nc(Nc3cccc([N+](=O)[O-])c3)c3ccccc3n2)cc1OC
COc1ccc2cc3cccn3c3cnc3cc3oc3cccccc12
CCC12OCC=CC=CC=CCC=CC=CC=CC1NCC2COCCO
CC1=CC=CC=C2C=CC=C2C(CC2C=CC=CC=CN2)C=CC=CC=NC=C1
CCN(CC)CCOc1ccc(-c2cc(C(=O)NC3CCCNC3)c(NC(=N)N)s2)cc1
CCN(CC)CCOc1ccc(C2=CC(C(=O)NC3CCCNC3)=C(NC(N)=O)SS2)cc1
NC(=O)Nc1sc(-c2ccc(C=O)cc2)cc1C(=O)N[C@H]1CCCNC1
NC(=O)Nc1sc(-c2ccc(C(=O)N3CCCCC3)cc2)cc1C(=O)N[C@H]1CCCNC1
C=C(C)c1ccc(-c2cc(C(=O)N[C@H]3CCCNC3)c(NC(N)=O)s2)cc1
CCCC(=O)c1ccc(-c2cc(C(=O)N[C@H]3CCCNC3)c(NC(N)=O)s2)cc1
C=C(C)Cc1cc(CCCCC)ccc1-c1cc(C(=O)N[C@H]2CCCNC2)c(NC(N)=O)s1
NC(=O)Nc1sc(-c2ccc(Cl)cc2)cc1C(=O)N[C@H]1CCCNC1
NC(=O)Nc1sc(-c2ccc(C(NO)N3CCCCC3)cc2)cc1C(=O)N[C@H]1CCCNC1
N#Cc1ccc(-c2cc(C(=O)N[C@H]3CCCNC3)c(NC(N)=O)s2)cc1
C=C(c1ccc(-c2cc(C(=O)N[C@H]3CCCNC3)c(NC(N)=O)s2)cc1)N1CCC1C
Cc1ccc(-c2cc(C(=O)N[C@H]3CCCNC3)c(NC(N)=O)s2)cc1
NC(=O)Nc1sc(-c2ccc(C(=O)N3CCC3)cc2)cc1C(=O)N[C@H]1CCCNC1
NCc1ccc(-c2cc(C(=O)N[C@H]3CCCNC3)c(NC(N)=O)s2)cc1
CC(NO)c1ccc(-c2cc(C(=O)N[C@H]3CCCNC3)c(NC(N)=O)s2)cc1
CC(=O)c1ccc(-c2cc(C(=O)N[C@H]3CCCNC3)c(NC(N)=O)s2)cc1
C=C(c1ccc(-c2cc(C(=O)N[C@H]3CCCNC3)c(NC(N)=O)s2)cc1)N1CCCCC1
CCN(CC)CCOc1cccc(-c2cc(C(=O)N[C@H]3CCCNC3)c(NC(N)=O)s2)c1
CS(=O)(=O)c1ccc(-c2cc(C(=O)N[C@H]3CCCNC3)c(NC(N)=O)s2)cc1
NC(=O)Nc1sc(-c2ccccc2)cc1C(=O)N[C@H]1CCCNC1
NC(=O)Nc1sc(-c2ccc(CF)cc2)cc1C(=O)N[C@H]1CCCNC1
```

```
NC(=O)Nc1sc(-c2ccc(C(=O)C3CCCCC3)cc2)cc1C(=O)N[C@H]1CCCNC1
NCN(O)Nc1sc(-c2ccc(CNO)cc2)cc1C(=O)N[C@H]1CCCNC1
N#Cc1ccc(-c2cc(C(=O)N[C@H]3CCCNC3)c(NN(O)CN)s2)cc1
Cc1ccc(-c2cc(C(=O)N[C@H]3CCCNC3)c(NN(O)CN)[nH]2)cc1
N#Cc1ccc(-c2cc(C(=O)N[C@H]3CCCNC3)c(NN(O)CN)[nH]2)cc1
NC(=O)Nc1[nH]c(-c2ccccc2)cc1C(=O)N[C@H]1CCCNC1
CC1CCN1C(=O)c1ccc(-c2cc(C(=O)N[C@H]3CCCNC3)c(NC(N)=O)s2)cc1
NC(=O)Nc1sc(C2=CC=C(F)C=CC2)cc1C(=O)N[C@H]1CCCNC1
NC(=O)Nc1sc(C2=CC=CC=CC=C2)cc1C(=O)N[C@H]1CCCNC1
Cc1ccc(-c2cc(C(=O)N[C@H]3CCCNC3)c(NC(N)=O)[nH]2)cc1
CCc1ccccc1-c1cc(C(=O)N[C]2CCCNC2)c(NC(N)=O)s1
NC(=O)Nc1sc(-c2ccc(CNO)cc2)cc1C(=O)N[C@H]1CCCNC1
NC(=O)Nc1sc(C2=CC=C2F)cc1C(=O)N[C@H]1CCCNC1
CCN(CC)CCOc1cccc(-c2cc(C(=O)N[C@H]3CCCNN3)c(NC(N)=O)s2)c1
CCN(CC)CCOc1cccc(-c2cc(C(=O)NC3CCCCC3)c(NC(N)OO)s2)c1
CN(C)CCCn1cnc2ccc(-c3c(-c4ccccn4)nn4c3CCC4)cc21
OCCCn1cnc2ccc(-c3c(-c4ccccn4)nn4c3CCC4)cc21
CN(C)CCCn1cnc2ccc(-c3c(-c4ccccc4)nn4c3CCC4)cc21
Cc1cccc(-c2nn3c(c2-c2ccc4ncn(CCCN(C)C)c4c2)CCC3)n1
CN(Cc1cnc2nc(N)nc(N)c2n1)c1ccc(C(=O)NC(CCC(=O)O)C(=O)O)c2ccccc12
CN(Cc1cnc2nc(N)nc(N)c2n1)c1ccc(C(=O)NCCNCC(=O)OOC(=O)O)c2ccccc12
CN(Cc1cnc2nc(N)nc(N)c2n1)C1=CC=C(C(=O)NC(CCC(O)OO)C(=O)O)C=CC=CC=C1
CC(=O)C(CCC(=O)O)NC(=O)c1ccc(N(C)Cc2cnc3nc(N)nc(N)c3n2)c2ccccc12
CN(Cc1cnc2nc(N)nc(N)c2c1)c1ccc(C(=O)NC(CCC(=O)O)C(=O)O)c2ccccc12
CN(Cc1cnc2nc(N)nc(N)c2n1)c1ccc(C(=O)NCCCCC(=O)OOC(=O)O)c2ccccc12
CN(Cc1cnc2nc(N)nc(N)c2n1)c1ccc(C(=O)NC(CCC(=O)O)C(N)=O)c2ccccc12
CN(Cc1cnc2nc(N)nc(N)c2n1)c1ccc(C(=O)NC(CCC(=O)O)C(O)CO)c2ccccc12
CC(=O)CCC(NC(=O)c1ccc(N(C)Cc2cnc3nc(N)nc(N)c3n2)c2ccccc12)C(=O)O
COc1ccc(C2=CC=CC=c3ccccc3=CC=CN2)nc1
O=C(NCC1CCCNC1)c1ccnc2[nH]c(-c3cccs3)nc12
O=C(NCC1CCCNC1)c1cccc2[nH]c(-c3cccs3)nc12
CC(=O)N[C@@H](CC(=O)NCC(COCc1ccccc1)COCc1ccccc1)C(=O)N[C@@H](CC(C)C)B(O)O
CC(=O)N[C@@H](CC(=O)NCC(COCc1ccccc1)COCc1ccccc1)C(=O)N[C@H](BO)CC(C)C
CC(C)C[C@@H](CO)NC(=O)[C@H](CC(=O)NCC(COCc1ccccc1)COCc1ccccc1)NCN(C)C
CC[C@H](C)[C@@H]1C(=O)O[C@H]1C(=O)NCCC(COCc1ccccc1)COCc1ccccc1
```

```
Cc1cc(Nc2c(C#N)cnc3c(Cl)cc(NCc4cn(CCN5CCCCC5)nn4)cc23)ccc1F
N#Cc1cnc2c(Cl)cc(NCc3cccc[n+]3[O-])cc2c1Nc1ccc(F)c(Cl)c1
NCCc1cnc2c(Cl)cc(NCC3=CN=CN=NN=N3)cc2c1Nc1ccc(F)c(Cl)c1
N#Cc1cnc2c(Br)cc(NCc3cn(CC(=O)O)nn3)cc2c1Nc1ccc(F)c(Cl)c1
OC1=c2c(cc3c4c2cn3-4)C=C1
Cn1cnc(-c2cc(-c3nnn[nH]3)ccn2)c1
Cn1cnc(-c2cc(C(=O)O)ccn2)c1-c1ccc(OCC2CC2)cc1
CCN(CCN(C)C)C(=O)CNCc1cc(C(=O)OC)ccn1
C=C1C=C2C=CC=CC=C2CC1C(=O)C(C(CC)OO)N(C)C
CCCC(OC)N(CC)CCN(CNC)C(=O)CNCC1=CC(C)=C1
CCCCOCC1=NC=CC(C(=O)OCC)=CC(O)CCCCCCCOC1
COc1ccc(-c2cncnc2-c2cc(C(=O)O)ccn2)cc1
Cc1cc2c(-c3cccnc3)ccc(N[C@@H]3CCNC[C@H]3CCC3CCS(=O)(=O)CC3)c2[nH]c1=O
Cc1cc2c(-c3cccnc3)ccc(N[C@@H]3CCCC[C@H]3OCC3CCS(=O)(=O)CC3)c2[nH]c1=O
Cc1cc2c(-c3cccnc3)cnc(N[C@@H]3CCNC[C@H]3OCC3CCS(=O)(=O)CC3)c2[nH]c1=O
Cc1cccc(-c2ccc(N[C@@H]3CCNC[C@H]3OCC3CCS(=O)(=O)CC3)c3[nH]c(=O)c(C)cc23)c1
Cc1cccc(-c2cnc(N[C@@H]3CCNC[C@H]3OCC3CCS(=O)(=O)CC3)c3[nH]c(=O)c(C)cc23)c1
Cc1cc2c(-c3cccnc3)ccc(N[C@@H]3CCNC[C@H]3OCC3CCS(=O)(=O)CC3)c2[nH]c1=O
Cc1cc2c(-c3ccccc3)ccc(N[C@@H]3CCNC[C@H]3OCC3CCS(=O)(=O)CC3)c2[nH]c1=O
Cc1cc2c(-c3ccccc3)cnc(N[C@@H]3CCNC[C@H]3OCC3CCS(=O)(=O)CC3)c2[nH]c1=O
Cc1cccc(-c2ccc(N[C@@H]3CCNC[C@H]3CCC3CCS(=O)(=O)CC3)c3[nH]c(=O)c(C)cc23)c1
Cc1cc2c(-c3cccnc3)ccc(N[C@@H]3CCCC[C@H]3CCC3CCS(=O)(=O)CC3)c2[nH]c1=O
Cc1cc2c(-c3cccnc3)cnc(N[C@@H]3CCNC[C@H]3CCC3CCS(=O)(=O)CC3)c2[nH]c1=O
Cc1cncc(-c2ccc(N[C@@H]3CCNC[C@H]3OCC3CCS(=O)(=O)CC3)c3[nH]c(=O)c(C)cc23)c1
Cc1cc2c(-c3cncnc3)cnc(N[C@@H]3CCNC[C@H]3OCC3CCS(=O)(=O)CC3)c2[nH]c1=O
Cc1cc2c(-c3ccccc3)ccc(N[C@@H]3CCCC[C@H]3CCC3CCS(=O)(=O)CC3)c2[nH]c1=O
Cc1cc2c(-c3ccccc3)cnc(N[C@@H]3CCNC[C@H]3CCC3CCS(=O)(=O)CC3)c2[nH]c1=O
Cc1cncc(-c2ccc(N[C@@H]3CCNC[C@H]3ONC3CCS(=O)(=O)CC3)c3[nH]c(=O)c(C)cc23)c1
Cc1cncc(-c2ccc(N[C@@H]3CCCC[C@H]3CCC3CCS(=O)(=O)CC3)c3[nH]c(=O)c(C)cc23)c1
Cc1cccc(-c2ccc(N[C@@H]3CCCC[C@H]3OCC3CCS(=O)(=O)CC3)c3[nH]c(=O)c(C)cc23)c1
COC1=CC=COC=C1COOC1C=C2C=CC=CC=CC(=C2)COOOCC1N
COC1=C2CC3=C(OO)C2=NC2C=C(C)C2C(=C3)C=C1
COC1=CC=COCC2C=CC=NC=CC=C2N=CC=CC=CC=C1
Nc1nc(-c2cc(C(=O)O)ccn2)cs1
Cc1ccnc(-c2csc(N)n2)c1
```

```
O=c1[nH]cnc2c(-n3cc(CCN4CCC(Cc5ccc(Cl)cc5)CC4)cn3)nccc12
O=c1[nH]cnc2c(-n3cc(CCN4CCC(Cc5ccc(Cl)c(Cl)c5)CC4)cn3)nccc12
CC1=C(n2cc(CCN3CCC(Cc4ccc(Cl)cc4)CC3)cn2)N=CC=C2C(=O)NC2N1
Cc1cc(CC2C=CN(CCc3cnn(-c4nccc5c4NCCNC5=O)c3)CC2)ccc1Cl
C=C1NC=Nc2c1ccnc2-n1cc(C=C2CCC(Cc3ccc(Cl)c(C)c3)CC2)cn1
OOC1NC=Nc2c1ccnc2-n1cc(CCN2CCC(Cc3ccc(Cl)cc3)CC2)cn1
COc1ccc(CC2CCC(CCc3cnn(-c4nccc5c(=O)[nH]cnc45)c3)CC2)cc1Cl
COc1ccc(CC2CCN(CCc3cnn(-c4nccc5c(=O)[nH]cnc45)c3)CC2)cc1
O=c1[nH]cnc2c(-n3cc(CCN4C=CC(Cc5ccc(Cl)cc5)CC4)cn3)nccc12
Cc1cc(CC2CCN(CCc3cnn(-c4nccc5c(=O)[nH]cnc45)c3)CC2)ccc1Cl
O=c1[nH]cnc2c(-n3cc(CCC4CCC(Cc5ccc(Cl)cc5)CC4)cn3)nccc12
CCc1cc(CC2CCN(CCc3cnn(-c4nccc5c(=O)[nH]cnc45)c3)CC2)ccc1Cl
Cc1cc(CC2CCN(CCc3cnn(-c4nccc5c4N=CNC5NO)c3)CC2)ccc1Cl
ONC1NC=Nc2c1ccnc2-n1cc(CCN2CCC(Cc3ccc(Cl)c(Cl)c3)CC2)cn1
O=c1[nH]nnc2c(-n3cc(CCN4CCC(Cc5ccc(Cl)c(Cl)c5)CC4)cn3)nccc12
O=C1NCCNc2c1ccnc2-n1cc(CCN2CCC(Cc3ccc(Cl)cc3)CC2)cn1
COc1cccc(CC2CCN(CCc3cnn(-c4nccc5c(=O)[nH]cnc45)c3)CC2)c1
ONC1NC=Nc2c1ccnc2-n1cc(CCN2CCC(Cc3ccc(Cl)cc3)CC2)cn1
Cc1cnc2c3cccccc(ccccn-3cc3cc-3c1)NCCNCC=N2
CCc1ccccncc2ccc3c(oon12)=C=CC=C3
CCC1=C2C=CC=CC2C(SCC(=O)N2CCOCC2)=CC1=O
```
